# High field asymmetric waveform ion mobility spectrometry improves *N*-homocysteinylation mapping in mouse liver and brain proteins

**DOI:** 10.1101/2025.07.21.664361

**Authors:** Joanna Perła-Kaján, Bianka Świderska, Agata Malinowska

## Abstract

*N*-Homocysteinylation has been shown to induce immunogenic, thrombogenic, and amyloidogenic properties of proteins. Although very important to gain insight into the mechanisms of homocysteine (Hcy) toxicity, proteome-wide studies of the effects of Hcy-thiolactone (HTL) protein modification remain challenging due to the low abundance of *N*-Hcy-proteins. High field asymmetric waveform ion mobility spectrometry (FAIMS) has been shown to improve the identification of other PTMs, we therefore expected it to facilitate the characterization of *N*-homocysteinylated proteins (*N*-Hcy-proteins) and help gain insight into their role in human disease. After extensive measurement optimization, we compared the yield of *N*-Hcy-protein/peptide identification across mouse liver and brain samples, either native or modified *in vitro* with HTL. Additionally, we examined the influence of different reduction and alkylation agents, namely DTT/IAA and TCEP/MMTS, on the number of identified *N*-Hcy-sites. FAIMS increased the number of *N*-Hcy-Lys-peptides and *N*-Hcy-proteins by 1.3-7-fold and 1.1-14-fold, respectively, regardless of alkylation method. We have identified 69 and 1,198 *in vivo* and *in vitro N*-Hcy-proteins, respectively. KEGG pathway term enrichment analysis showed that among *in vitro N*-Hcy-proteins, ten top KEGG pathways were Parkinson disease, prion disease, Huntington disease, oxidative phosphorylation, amyotrophic lateral sclerosis, pathways of neurodegeneration – multiple diseases, carbon metabolism, carcinogenesis – reactive oxygen species, Alzheimer disease, and diabetic cardiomyopathy. We conclude that FAIMS is a valuable addition to *N*-Hcy-proteome analysis workflow and facilitates the mapping of *N*-Hcy-sites. Data are available via ProteomeXchange with identifier PXD062860.

## INTRODUCTION

Hyperhomocysteinemia is associated with over a hundred human diseases [1]. One of the mechanisms of homocysteine (Hcy) toxicity involves its conversion to Hcy-thiolactone (HTL) catalyzed by Met-tRNA synthetase and spontaneous modification of protein Lys residues by HTL, a process called *N*-homocysteinylation [2]. As a result of protein *N*-homocysteinylation, an additional thiol group(s) is(are) added to the polypeptide chain, which can lead to protein oxidation and aggregation. Structural and functional consequences of modification with HTL have been studied for a number of individual proteins, e.g. fibrinogen [3, 4], collagen [5], albumin [6, 7], dynein [8], cytochrome c [9–11], α-synuclein [12], and recently DJ-1 (Parkinson’s disease protein 7) [13]. *N*-Homocysteinylated proteins (*N*-Hcy-proteins) have been shown to elicit autoimmune response [14], gain thrombogenic [3] and amyloidogenic properties [7], as well as reduce the effectiveness of DNA damage repair [15] and affect neuron function [8, 12, 13, 16]. However, due to the low abundance of modified proteins, their detection remains challenging. As a result, few proteome-wide studies of N-homocysteinylation sites have been performed. Previous subjects of research focused on *N*-Hcy-Lys site mapping include: yeast grown in presence of high Hcy or HTL [17], MARS-interacting proteins in cultured human colon cancer HCT116 cells [15], HeLa cells treated with Hcy-thiolactone [18], and mouse neuronal NE4C stem cells overexpressing MARS [16].

Post-translational modifications (PTMs) identification is usually achieved using a bottom-up proteomic approach, in which a mixture of proteins is digested into peptides, followed by separation via liquid chromatography, usually coupled with a mass spectrometer (LC-MS/MS). This strategy, while appropriate for identification of proteins present in the sample, has limited effectiveness in identifying low abundant entities like peptides carrying PTMs. Immunoenrichment often used to decrease the complexity of samples for identification of other PTMs is troublesome for *N*-Hcy-peptide discovery due to the lack of commercially available antibodies specific towards *N*-Hcy-proteins. With the development in instrumentation, the number of proteins identified in a single LC-MS run increases, which gives a better insight into the proteome, including isoforms and PTMs. Sample complexity can be reduced by ion mobility spectrometry (IMS), where ions are separated based on their mobility in an electric field. One of IMS types is high-field asymmetric waveform ion mobility spectrometry (FAIMS). In FAIMS, gas-phase ions transported by a flow of carrier gas, travel between inner and outer cylindrical electrodes to which an asymmetric waveform is applied, alternating between high and low electric fields of opposite polarity. During the travel of ions between the electrodes, they drift towards and eventually reach one of them. This leads to their discharging and neutralization. To prevent the ion drift, the FAIMS device applies a small direct current (DC) potential, namely the compensation voltage (CV), to the inner electrode. The CV required to compensate for the drift is ion dependent, thus CV is a selection parameter for transmitting a subset of ions to the mass spectrometer.

Using FAIMS has been shown to be advantageous in a number of research applications, e.g. separation of peptide isomers containing the same PTM but at different sites [19, 20], increasing the identification rate of multiphosphorylated peptides [21], improving protein and phosphosite identification [22] as well as identification of multi-PTM containing peptides, and detection of PTM crosstalk sites [23]. The addition of FAIMS to LC-MS/MS setup reduces chemical/endogenous background noise associated with highly-complex matrix, thereby improving the sensitivity via increasing the signal-to-noise ratio, making it highly advantageous in quantitative bioanalysis [24]. FAIMS reduces the chemical noise associated with singly charged ions and increases the overall population of detectable tryptic peptides. Venne et al. have shown a 6-12-fold improvement in signal-to-noise ratio for a wide range of multiply charged peptide ions and an increase of 20% in the number of detected peptides when compared to conventional LC-MS approach [25].

Klaeger et al. have reported that FAIMS increased the human leukocyte antigen class I (HLA-I)-I peptide identifications by up to 58% [26]. Yan et al. have used FAIMS for high-coverage profiling of the human cysteinome [27]. FAIMS increased phosphoproteomic coverage, and enabled the separation of phosphopeptide isomers that often coelute and can be misassigned in conventional LC−MS/MS experiments. It also provided more than 30% and 200% increase in the number of quantifiable peptides compared to LC−MS/MS performed with and without SCX fractionation, respectively. Furthermore, FAIMS reduced the occurrence of interfering isobaric ions and improved the accuracy of quantitative measurements [28].

We sought to determine whether FAIMS ion separation could improve the identification of *N*-Hcy-sites in proteins. In our study, a system consisting of the Evosep One chromatograph combined with the Orbitrap Exploris 480 spectrometer and the FAIMS Pro ion mobility module was used. Evosep is a chromatographic system that was redesigned for proteomics analysis, especially for measurements of long series of samples. Unlike traditional HPLC, the system uses disposable trap columns, which reduce sample cross-contamination and improve the repeatability of the measurements. In our set-up, ion mobility is installed between the ion source and the spectrometer to act as a peptide ion filter. By removing sets of ions during analysis, it gives the opportunity to detect low abundant peptides, during data-dependent acquisition, in which fragmentation is triggered for peptides generating highest MS signals. This instrumental setup has been shown to increase the number of identified proteins in complex samples [22, 29]. Novel FAIMS technology developed by Thermo Scientific allows to set various compensation voltage (CV) values during one LC-MS run, a feature already used for global proteomics and PTMs analysis studies [23, 30–32].

First, we ran a preliminary experiment to optimize the CV settings, to achieve the highest possible number of identified proteins and *N*-Hcy-sites per sample (Figure 1). In the next step we used the optimized CV settings to identify *N*-Hcy-sites in mouse brain and liver proteins. Here we demonstrate that FAIMS increases the number of identified *N*-Hcy-sites. Obtaining the information on the specific sites of *N*-homocysteinylation is essential for understanding their cellular roles, and how they affect protein properties, such as stability, translocation, interaction, catalytic activity, and as a result – function. Insight into *N*-homocysteinylome may help to identify biomarkers for disease diagnosis and prognosis, especially for neurodegenerative and cardiovascular disorders, which are known to be associated with HHcy. Gaining this information might also contribute to elucidating molecular mechanisms underlying these diseases and potentially lead to development of novel therapeutic targets.

**Figure 1.**
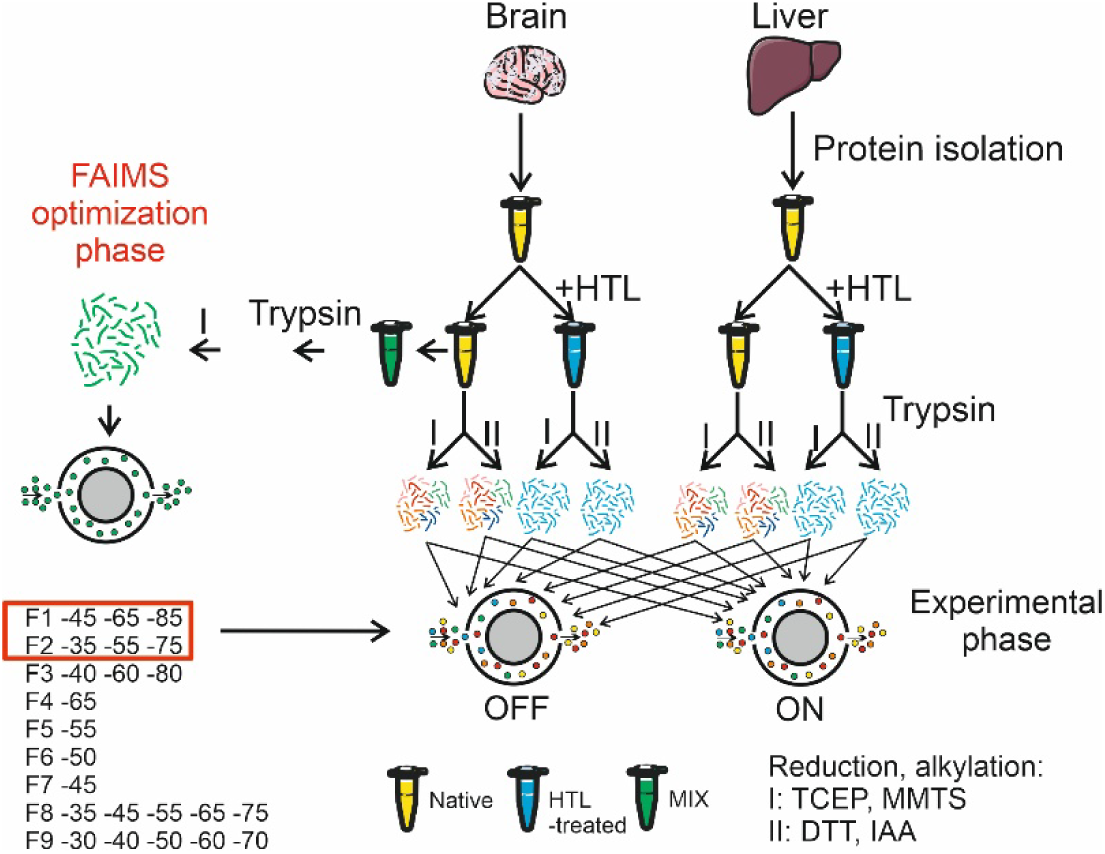
Schematic overview of experimental setup.

## METHODS

### Chemicals and Reagents

All materials were purchased from Merck unless otherwise stated.

### Mice

C57BL/6J mice were housed and bred at the Rutgers-New Jersey Medical School Animal Facility. The mice were fed with a standard rodent chow (LabDiet5010; Purina Mills International, St. Louis, MO, USA). Animal procedures were approved by the Institutional Animal Care and Use Committee at Rutgers-New Jersey Medical School. Liver and brain of 12-month-old male mice were used in the study.

#### Protein isolation and digestion

Protein was isolated from mouse liver and brain pulverized in liquid nitrogen. 40 mg of tissue was suspended in 10 fold volume of 50 mM ammonium bicarbonate with 0.1% SDS and 50% 2,2,2-trifluoroethanol (Merck, T63002) and sonicated on ice with 6 1-sec. strokes of 65-70% power (Bandelin Sonoplus). After centrifugation at 15,000xg for 15 min at 4°C protein concentration was estimated by Bradford method with Coomassie protein assay reagent (Merck, 27813). As a positive control for the detection of *N*-homocysteinylation, 100 μL of brain or liver protein was incubated with 100 mM HTL overnight at RT. 120 µg of protein from each sample or a sample mix (equal amounts of native and HTL-treated samples) was digested according to FASP protocol with some modifications [33]. Cysteine groups were reduced by 1 hour incubation at room temperature with 20 mM dithiothreitol (DTT) or at 60°C with 20 mM tris(2-carboxyethyl)phosphine (TCEP), for IAA and MMTS alkylation, respectively. Solutions were transferred onto Vivacon 30 kDa molecular weight cut-off filter (Sartorius Stedim). Filters were spun at 14 500 g for 15 min and washed with 200 µl urea solution (8 M urea in 100 mM ammonium bicarbonate). Cysteine residues were alkylated by 30 min incubation at room temperature with 50 mM iodoacetamide (IAA) or 50 mM methyl methanethiosulfonate (MMTS). Filters were washed three times with 8 M urea solution and with 100 mM ammonium bicarbonate, respectively. After each addition, samples were centrifuged until the cut-off filter was dry. Digestion was carried out overnight using 3 µg of trypsin (Promega) at 37°C (enzyme:protein ratio 1:40). Peptides were eluted from spin filters by two additions of 50 µl of 100 mM ammonium bicarbonate and one of 500 mM NaCl solution. The Speedvac dried samples were resuspended in 1.2 ml of 0.1% formic acid in water.

### Mass spectrometry

Samples were analyzed using an LC-MS system composed of Evosep One HPLC System (Evosep Biosystems) coupled to an Orbitrap Exploris 480 mass spectrometer (Thermo Scientific) working with FAIMS Pro interface (Thermo Scientific). 40 µl of peptide solutions (2 µg) were loaded onto Evotips C18 disposable trap columns as described previously [34]. Each sample was measured twice using single- or multi-CV FAIMS method with applied different CV values. Peptides were fractionated using 88 min (15 samples per day) predefined Evosep gradient at a flow rate of 220 nl/min on an analytical column (Dr Maisch C18 AQ, 1.9 µm beads, 150 µm ID, 15 cm long, Evosep Biosystems). The following FAIMS settings were used: FAIMS Standard Resolution, inner/outer electrode temperature 100°C, Total carrier gas flow = 4.6 L/min (nitrogen), Static. In the FAIMS optimization phase MIX of two samples was measured with 8 different single- or multi-FAIMS setting. For FAIMS optimization, CV settings designated F1-F9 were applied: F1: -45, -65, -85; F2: -35, -55, -75; F3: -40, -60, -80; F4: -65; F5: -55; F6: -50; F7: -45; F8: -35, -45, -55, -65, -75 or F9: -30, -40, -50, -60, -70 V. For native and HTL-treated brain and liver samples, three CV values were applied, each during MS acquisition, each during 1 s cycle: -35/-55/-75 or -45/-65/-85 V. Data-dependent acquisition parameters were as follows: cycle time 1s for each FAIMS CV value, collisional induced fragmentation NCE 30%, spray voltage 2.1 kV, funnel RF level 40, heated capillary temperature to 275°C. Full MS scans covered the mass range of 300–1700 m/z with a resolution of 60,000, maximum injection time was set to auto and a normalized AGC target to 200%. MS2 scans were acquired with a resolution of 15,000, an auto maximum injection time and a standard AGC target. Ion isolation window was set to 1.6 m/z, a dynamic exclusion to 20 s and a minimum intensity threshold at 5e3. For noFAIMS runs, the MS1 AGC target was set to 300%, and the top 40 precursors were selected for fragmentation in each cycle. All other parameters remained consistent with those used in FAIMS runs.

### Data analysis

Raw files were processed with Thermo Protein Discoverer (version 2.4, Thermo Scientific) using *Mus musculus* proteins deposited in Swissprot database (version 2021_04, 17062 sequences). Peptides and proteins were identified using Sequest HT module with following parameters: enzyme – Trypsin, precursor mass tolerance – 10ppm, fragment mass tolerance – 0.02 Da, max missed cleavages – 2, fixed modifications – Carbamidomethyl (C) for IAA alkylation, Methylthio (C) for MMTS alkylation, variable modifications – Oxidation (M) and Acetyl, Met-loss, Acetyl+Met-Loss at protein terminus. *N*-homocysteinylation was set as variable modification and identified based on mass increase of 174 Da or 163 Da for S-carbamidomethyl-Hcy-Lys (reduction with DTT, alkylation with IAA) or S-methylthio-Hcy-Lys (reduction with TCEP, alkylation with MMTS), respectively. FAIMS runs performed on the same sample with different CV values were treated as fractions and merged into one result. FDR was calculated using target/decoy strategy with Percolator module. Site probability threshold was set to 75. Peptide and protein-level FDR was calculated, with threshold value of 0.01. Only high probability proteins with at least 1 unique peptide and at least 2 peptides overall were considered, high confidence was also required for PTM sites and peptide groups.

Peptide properties, i.e. sequence length, hydrophobicity, GRAVY (Grand Average of Hydropathy, calculated as the sum of hydropathy values [35] of all the amino acids, divided by the number of residues in the sequence), molecular weight average, molecular weight monoisotopic and theoretical pI, were analyzed using Peptide Analyzing Tool available at www.thermofisher.com. Data on sequence length, hydrophobicity, GRAVY, molecular weight average, molecular weight monoisotopic and theoretical pI are presented as median, 25-75% and min-max values. The differences between groups of peptides acquired with different CVs were analyzed with One-way ANOVA and RIR Tukey’s post-hoc test. The alpha level that defines statistical significance was set at 0.05. Statistical analysis was performed with Statistica version 13 (TIBCO Software Inc.). Venn diagrams were generated with a DeepVenn web application (https://arxiv.org/abs/2210.04597; https://doi.org/10.48550/arXiv.2210.04597). Functional interaction network analysis of *in vivo N*-Hcy-proteins was performed using interaction data from the Search Tool for the Retrieval of Interacting Genes/Proteins (STRING) database, version 12.0 [35]. The following basic settings were used: full STRING network, active interaction sources: text mining, experiments, databases, co-expression, neighborhood, gene fusion, co-occurance; minimum required interaction score: medium confidence (0.4). KEGG pathway term enrichment analysis of *in vitro N*-Hcy-proteins was performed using DAVID Bioinformatics Resources [36, 37]. The positions of amino acids in protein chain are according to UniProtKB, without protein processing (initial M being the first AA).

The SAMPL Guidelines were followed for the statistical analyses.

### Data availability

The mass spectrometry proteomics data have been deposited to the ProteomeXchange Consortium via the PRIDE [38] partner repository with the dataset identifier PXD062860 and 10.6019/PXD062860.

## RESULTS

Previous studies have shown that ion separation strategies help to increase the number of identified proteins and peptides in complex samples. As FAIMS has, to our knowledge, not been evaluated for *N*-homocysteinylation mapping, the aim of this experiment was to examine whether FAIMS improves the identification of *N*-Hcy-sites in tissues. Samples were analyzed with Evosep One HPLC System (Evosep Biosystems) coupled to an Orbitrap Exploris 480 mass spectrometer (Thermo Scientific) working with FAIMS Pro interface (Thermo Scientific). FAIMS Pro interface is an ion mobility device installed between ESI ion source and MS, which allows for online ion fractionation and filtering to identify more proteins, low abundance peptides or PTMs.

The study workflow included FAIMS optimization phase and experimental phase (Figure 1). Fine tuning of FAIMS CV parameters was performed using a mixture (MIX) of equal amounts of mouse brain proteins in native and modified *in vitro* with HTL (referred to as HTL-treated) state. For the experimental phase, native and HTL-treated mouse brain and liver proteins were used. Protein samples were extracted from the brains (two animals) and livers (three animals) of one-year-old wild-type male mice as described in the Methods section. To evaluate the influence of alkylation agents on *N*-Hcy mapping effectiveness, either DTT and IAA or TCEP and MMTS were used for thiol groups reduction and alkylation. Identification of *N*-Hcy-Lys sites was based on the 174 Da or 163 Da increase in peptide mass due to S-carbamidomethyl- or S-methylthio-Hcy-Lys present in a peptide, respectively. In the optimization phase, MIX was analyzed using 9 different CV settings, either in single- or multi-CV FAIMS runs. In the experimental phase, each sample was measured either without FAIMS or with FAIMS. For the second option, two FAIMS-LC-MS runs per sample were performed, using the multi-CV FAIMS settings selected in the optimization phase (Figure 1).

### CV optimization phase

To develop a FAIMS-based method for improving the sensitivity of *N*-Hcy-site detection, we optimized CV settings as a first step. Mouse brain proteins were analyzed using multiple or single CVs and the number of identified proteins and *N*-Hcy-peptides was recorded. We tested five different multi-FAIMS settings, three with three CVs and two with five CV values (Figure 1). During multi-FAIMS, which can potentially increase the number of identified peptides, the spectrometer switches between selected CV parameter values and for each CV, full MS measurement and a series of MS2 fragmentations are performed.

#### Selecting optimal CV settings

The highest number of identified proteins and *N*-Hcy-peptides was recorded for multi-FAIMS with three CV values, while the lowest number of identified *N*-Hcy-peptides was obtained while using single CV settings (Figure 2). Then, we wanted to determine which two multi-CV methods yielded the highest number of identified proteins and *N*-Hcy peptides (Figure 2 A,B) in total.

**Figure 2.**
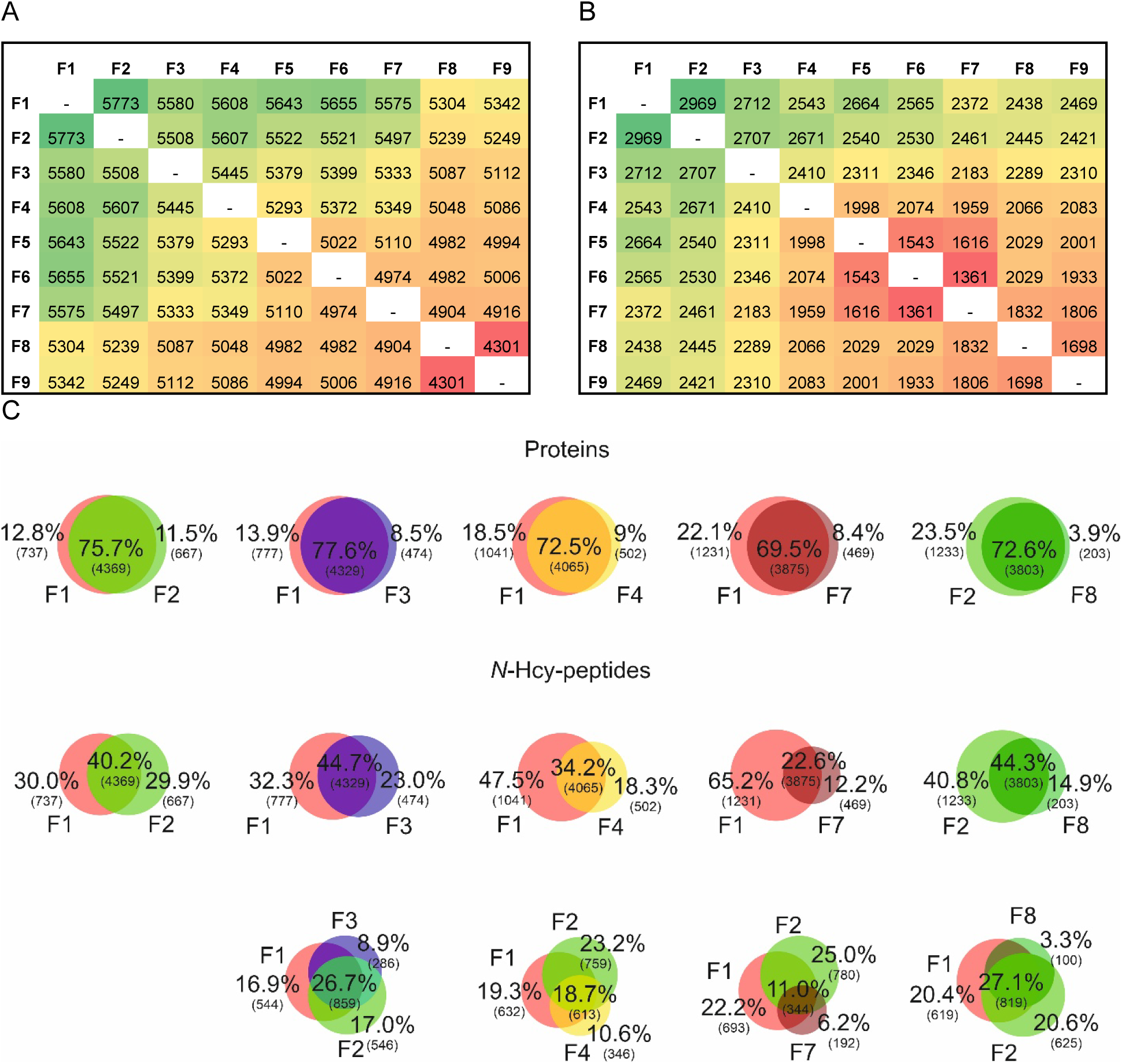
Optimization phase: the number of identified proteins (A) and N-Hcy-peptides (B); Venn diagrams of proteins and N-Hcy-peptides identified with various CV settings. Percentages and absolute numbers are shown (C). CV settings coding: F1: -45, -65, -85; F2: -35, -55, -75; F3: -40, -60, -80; F4: -65; F5: -55; F6: -50; F7: -45; F8: -35, -45, -55, -65, -75 or F9: -30, -40, -50, -60, -70 V.

As shown in Figure 2 A,B the highest number of identified proteins and *N*-Hcy-peptides was observed in data acquired with two triple-CV methods, i.e. F1 -45 -65 -85 and F2 -35 -55 -75 V. There was 75.7% overlap in proteins identified with F1 and F2 CV settings, with 12.8% and 11.5% of unique proteins identified with F1 and F2 CV settings, respectively. For *N*-Hcy-peptides, the overlap between F1 and F2 CV settings was 40.2%, with 30% and 29.8% of unique proteins acquired with F1 and F2 CV settings, respectively. Other CV settings yielded lower numbers of unique proteins and *N*-Hcy-peptides as compared to F1 (Figure 2 C).

Next, we sought to analyze whether sample measurement with a third CV setting would result in a significant increase in the number of identified *N*-Hcy-peptides. As shown in the Venn diagrams for three CV settings (Figure 2 C), in most cases the additional FAIMS measurement gives less than 10% increase in *N*-Hcy-peptides. Such limited improvement in detection did not, in our opinion, justify performing additional run, so in the experimental phase we decided to use two CV settings per sample, i.e. F1 and F2.

#### Properties of N-Hcy-peptides identified with different CVs

To check if various CV settings correlate with properties of identified *N*-Hcy-peptides, sequence length, hydrophobicity, and theoretical pI were analyzed. The median with ranges (Table 1 S) and histograms (Figure 3, Figure 1 S) of *N*-Hcy-peptide characteristics as a function of the CVs used are shown.

**Figure 3.**
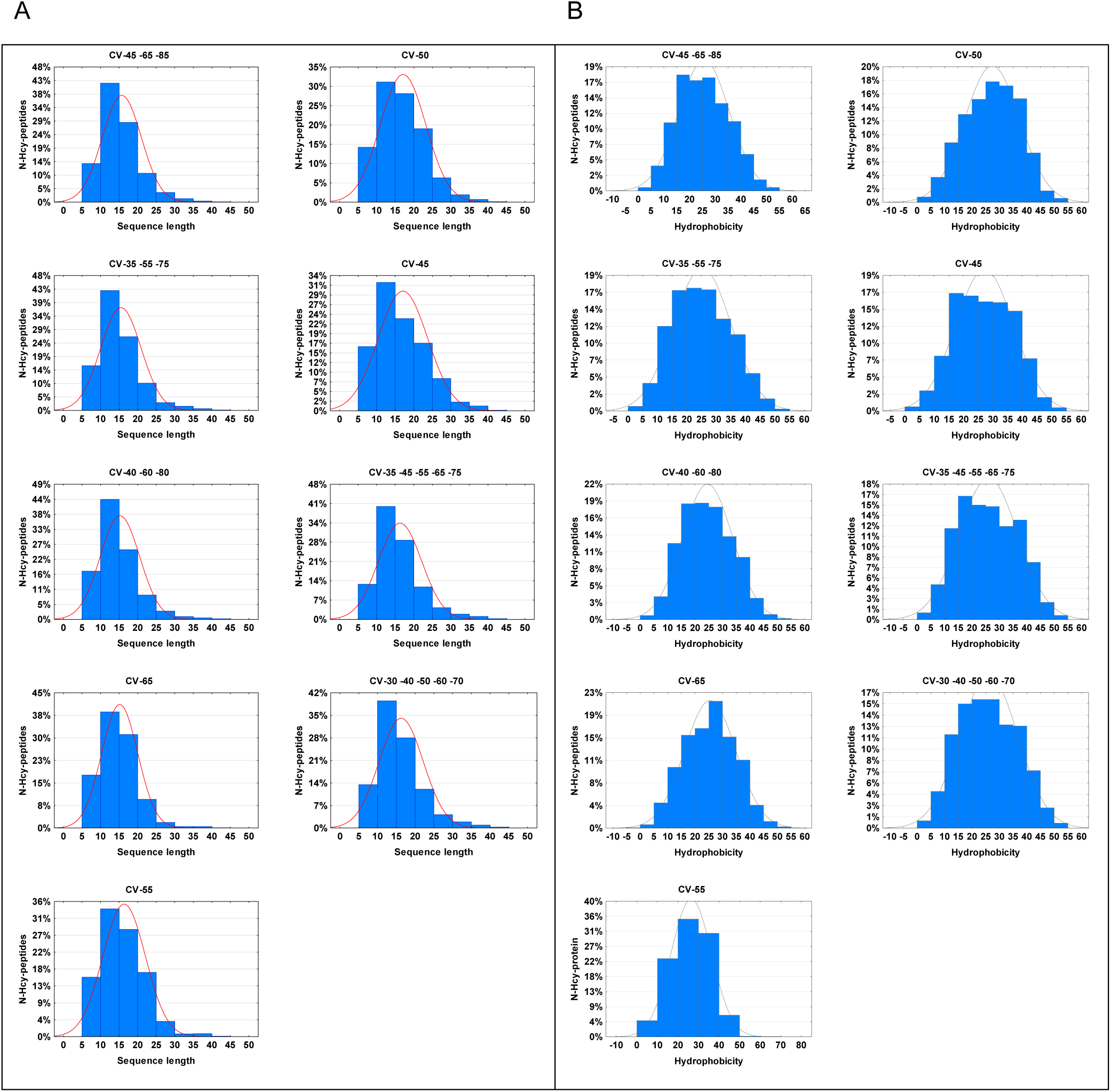
Histograms of N-Hcy-peptides’ sequence length (A) and hydrophobicity (B) for each CV setting tested during optimization phase.

Single and multi-CV settings resulted in the detection of *N*-Hcy-peptides with significantly different sequence lengths (Table 1 S, Figure 4). The average length of *N*-Hcy-peptides identified during the optimization phase was 16 and ranged from 6 to 45 amino acid residues. Within the population of *N*-Hcy-peptides, those with lengths between 10 and 15 residues are most abundant, reaching 40-44% in multi-CV settings, and 30-35% in single-CV conditions. CV settings designated as F5, F6 and F7 resulted in acquisition of *N*-Hcy-peptides with significantly longer median length. *N*-Hcy-peptides with the shortest median were identified using F2 and F3 CVs.

**Figure 4.**
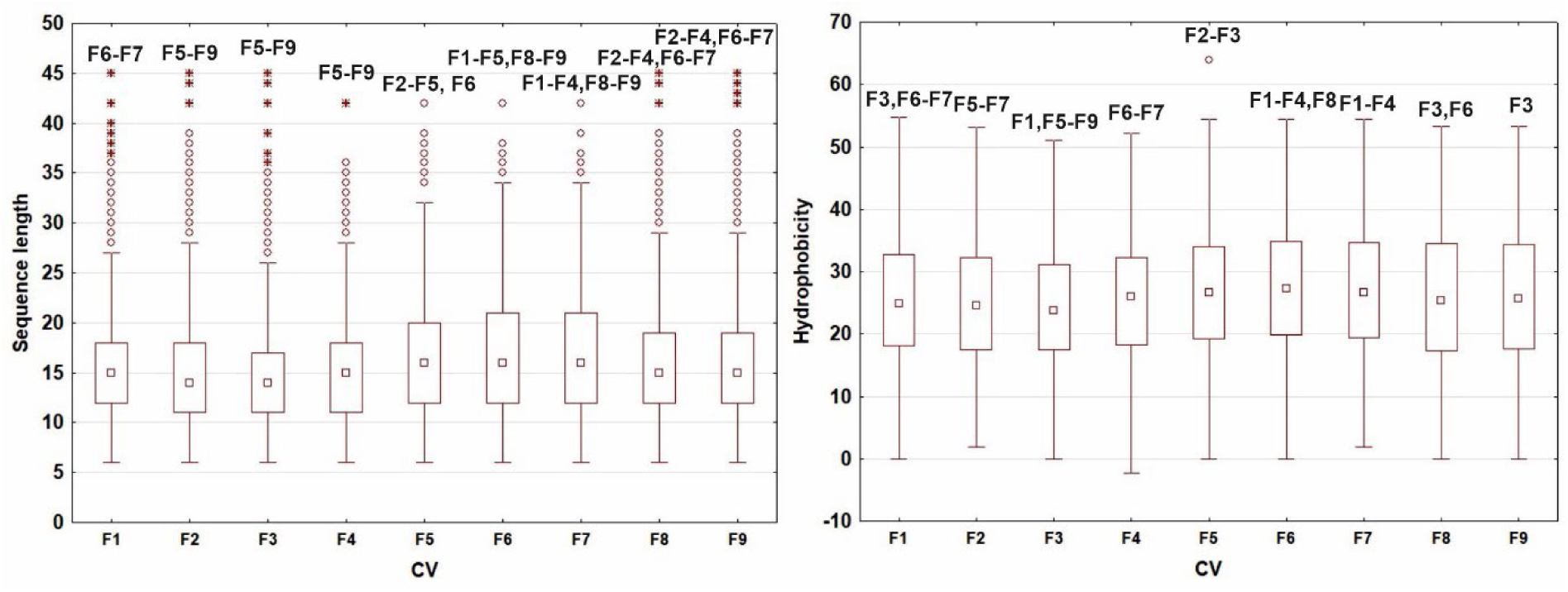
Properties of N-Hcy-peptides acquired across various CVs during optimization phase. Boxplot depicts upper and lower quartiles with median shown as a square. Whiskers – non-outliers, circles – outliers, stars – extreme data. Symbols F1-F9 above data indicate significant (p<0.05) difference with other CV’s peptides property. Detailed P-values are shown in Table 2 S.

Differences in CV settings also correlated with changes in hydrophobicity distribution of identified *N*-Hcy-peptides. The mean hydrophobicity of *N*-Hcy-peptides was 25.6 and ranged from -2.3 to 63.9 (Table 1 S, Figure 4). *N*-Hcy-peptides with the highest hydrophobicity were identified with F6, F5 and F7 CVs.

The average theoretical pI of *N*-Hcy-peptides was 7.33 and ranged from 3.40 to 12.50. No statistically significant differences in theoretical pI of *N*-Hcy-peptides were observed between peptides obtained with different CVs (Table 1 S, Figure 1 S).

### Experimental phase

In the experimental phase, native and HTL-treated samples of brain and liver protein were analyzed with and without FAIMS. Optimized FAIMS parameters were used, i.e. samples were measured with two CV settings, F1 -45 -65 -85 and F2 -35 -55 -75 V. Analyses were performed with two types of alkylation methods, namely DTT/IAA and TCEP/MMTS.

#### N-Hcy-peptide identification benefits from separation of ions in the gas phase

FAIMS increased the number of identified proteins by 30-40% (Figure 5 A, G) and peptides by 40-70% (Figure 5 D, J) in both organs and using both alkylation methods. In liver and brain samples, number of *N*-Hcy-Lys-peptides and *N*-Hcy-proteins increased 1.3-7-fold (Figure 5 E-F, K-L) and 1.1-14-fold (Figure 5 B-C, H-I), respectively. Both alkylation methods performed similarly, apparently not influencing FAIMS efficiency. Only in the brain IAA-alkylated sample, a 30% and 40% decrease in the number of acquired *N*-Hcy-peptides and *N*-Hcy-proteins, respectively, was observed upon using FAIMS.

**Figure 5.**
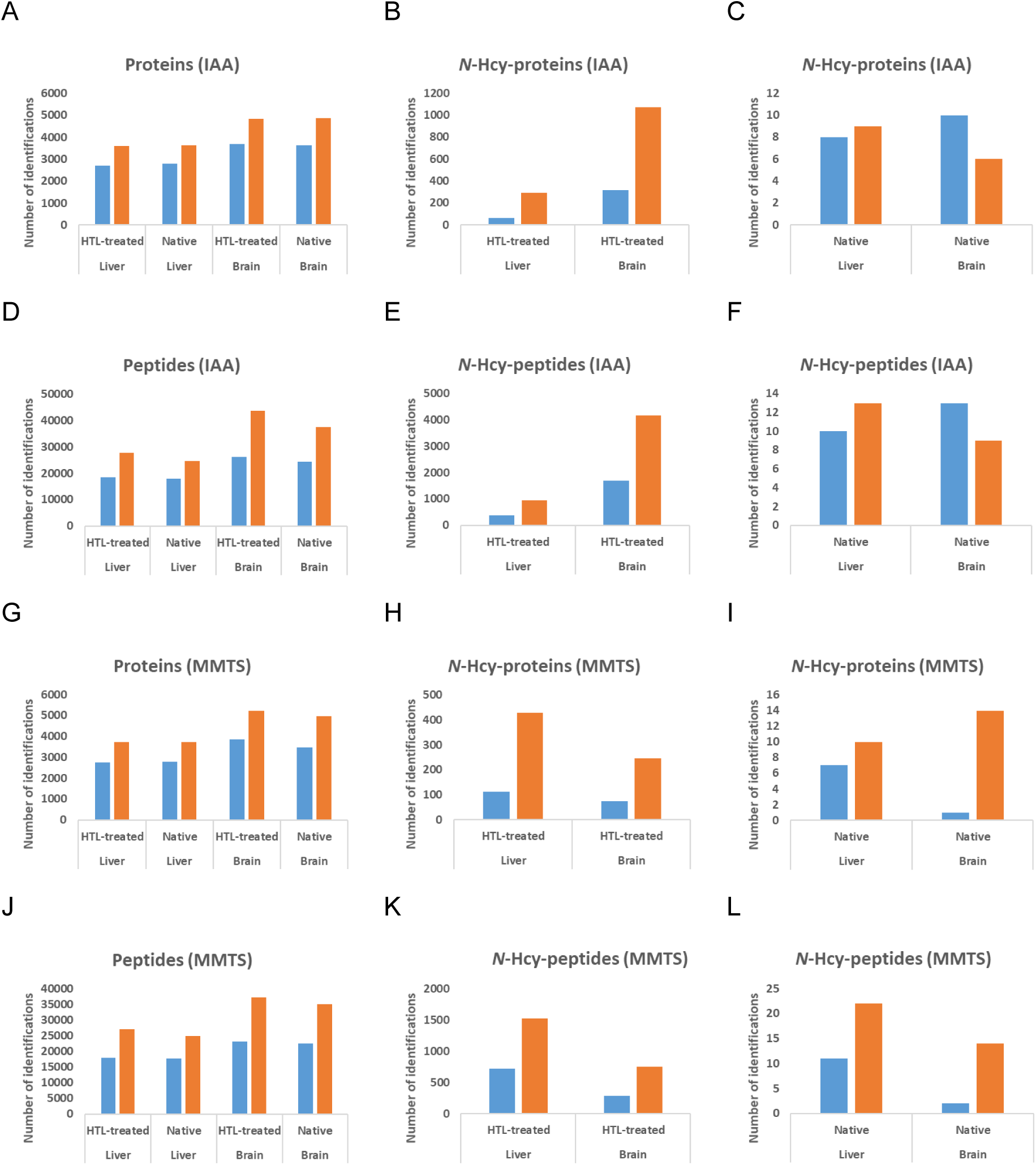
FAIMS (orange bars) improves identification yield of proteins, N-Hcy-proteins, N-Hcy-peptides as compared to no FAIMS acquisition (blue bars). A-F IAA-; G-L MMTS-alkylation.

#### FAIMS decreases isolation interference and signal to noise ratio

The increased number of identifications upon applying FAIMS ion separation could be related to improved data quality. One of the factors related to data quality is peptide spectrum match (PSM) isolation interference. The interference refers to the challenges encountered when multiple peptides coelute and are fragmented simultaneously during MS/MS analysis. This phenomenon can lead to inaccurate quantification and identification of peptides due to overlapping signals in the resulting spectra, and is particularly troublesome for highly complex samples like tissue. Since FAIMS acting as ion filter and separation technique could reduce interference, we therefore calculated PSM isolation interference for all analyzed samples (brain native and HTL-treated, as well as liver native and HTL-treated) separately for both alkylation methods.

Indeed, FAIMS increased the number of PTMs with isolation interference below or equal 5% from 20% to 25%, and doubled the percentage of PTMs with isolation interference in the range 5-10%. The percent of PTMs with isolation interference greater than 45% have decreased with the use of FAIMS as compared to conventional acquisition. The reduction in isolation interference induced by FAIMS was similar for both alkylation conditions (Figure 6).

**Figure 6.**
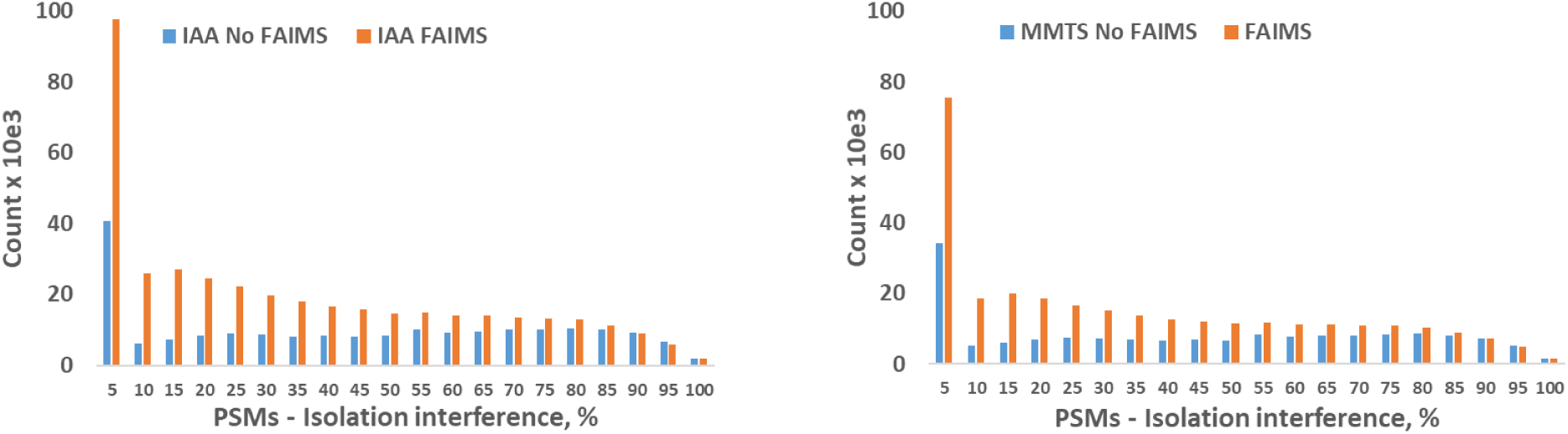
Isolation interference of PSMs for acquisition with FAIMS (orange bars) and conventional method (blue bars), using IAA or MMTS alkylation. Histograms are calculated for all samples examined in the measurement phase.

In previous studies, FAIMS has been shown to increase the population of peptide ions while reducing the contribution of chemical noise [25]. We have observed statistically significant increase of signal to noise ratio while using FAIMS, as compared to no-FAIMS analysis (no-FAIMS vs. CV -35, -55, -75 and no-FAIMS vs. CV -45, -65, -85 adjusted *P*-values 0.0000382 and 0.0000942, respectively) (Figure 7). Signal to noise ratio was not significantly different between CV -45, -65, -85 and CV -35, -55, -75 (adjusted *P*-value 0.2418).

**Figure 7.**
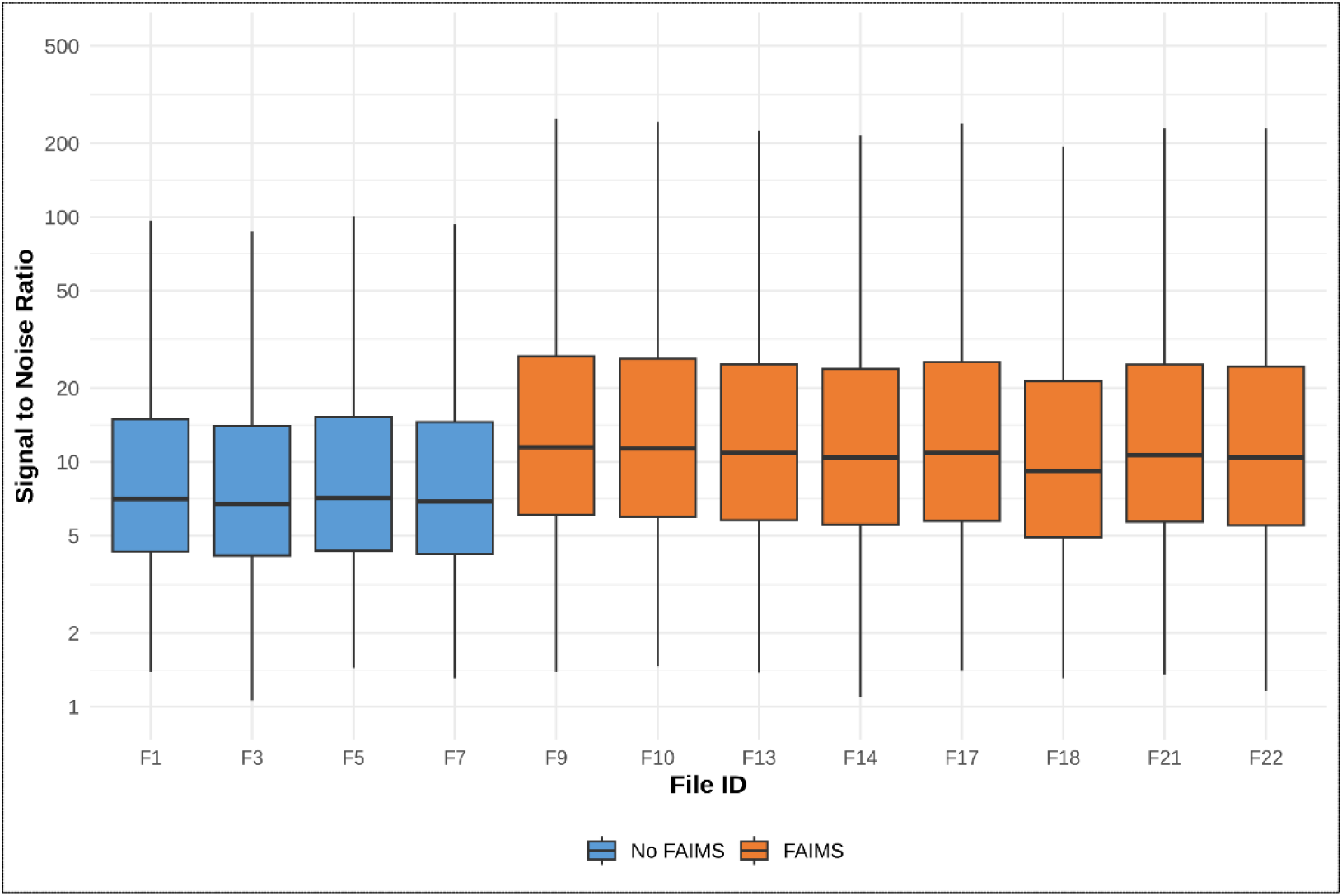
Signal to noise ratio. Median, interquartile range, min-max values are shown.

#### N-Hcy-peptides have higher precursor MH+ and charge state than unmodified peptides

We have noticed that *N*-Hcy-peptides have significantly higher median precursor MH+ compared to unmodified peptides (Table 1). This is probably due to miscleavages caused by Hcy residue bound to ε-amino group of Lys residue and thus decreasing the effectiveness of tryptic proteolysis. The increased miscleavage rate for *N*-Hcy-peptides was confirmed by the acquired data (Figure 3 S). We have also observed statistically significant difference in median precursor MH+ of *N*-Hcy-peptides depending on the alkylating agent, with IAA yielding higher precursor MH+ than MMTS, in case of both *N*-Hcy-peptides and peptides (Table 1).

**Table 1.** Median (min-max) precursor MH+ of N-Hcy-peptides and peptides identified in both alkylation conditions.

| Alkylating agent | Median precursor MH<sup>+</sup> (min-max) [Da] |  | *P*-value |
| --- | --- | --- | --- |
|  | *N*-Hcy-peptides | Peptides |  |
| IAA | 1,838.86 (902.55-4,984.13)<br>n=4,875 | 1,482.68 (600.41-4,996.59)<br>n=70,196 | 0.0000 |
| MMTS | 1,734.89 (907.47-4,282.94)<br>n=2,173 | 1,453.77 (606.36-4,996.59)<br>n=64,123 | 0.0000 |
| *P*-value | 0.0000 | 0.0000 |  |

Next we compared the charge state of *N*-Hcy-peptides with the charge state of peptides that were not *N*-homocysteinylated (Figure 2 S). The majority of *N*-Hcy-peptides carry the 3+ charge, whereas the majority of non-*N*-Hcy-peptides are in 2+ charged state. This observation is independent of the acquisition mode and holds true both for FAIMS and conventional method. The fact that *N*-Hcy-peptides carry, on average, more charges than non-*N*-Hcy-peptides is probably connected with their greater average length, which results in more potential sites of charge attachment.

#### Properties *of* in vitro N-Hcy-peptides, depending on organ, alkylation and acquisition method

We asked whether the introduction of FAIMS into analysis, and the type of cysteine alkylation (IAA or MMTS) correlates with the properties of identified *N*-Hcy-peptides. To answer this question, peptide length, hydrophobicity and theoretical pI of *in vitro* liver and brain *N*-Hcy-peptides were analyzed.

#### Sequence length

*N*-Hcy-peptides (liver and brain) alkylated with IAA and detected in FAIMS measurement are significantly longer than *N*-Hcy-peptides identified without FAIMS, while *N*-Hcy-peptides alkylated with MMTS do not differ significantly in length depending on acquiring method. *N*-Hcy-peptides acquired following IAA alkylation are significantly longer than those identified following MMTS alkylation, but only when analyzed together for both organs (Figure 8 A). When analyzed separately liver and brain *N*-Hcy-peptides, do not differ significantly in peptide length, depending on alkylation method (Figure 8 B).

**Figure 8.**
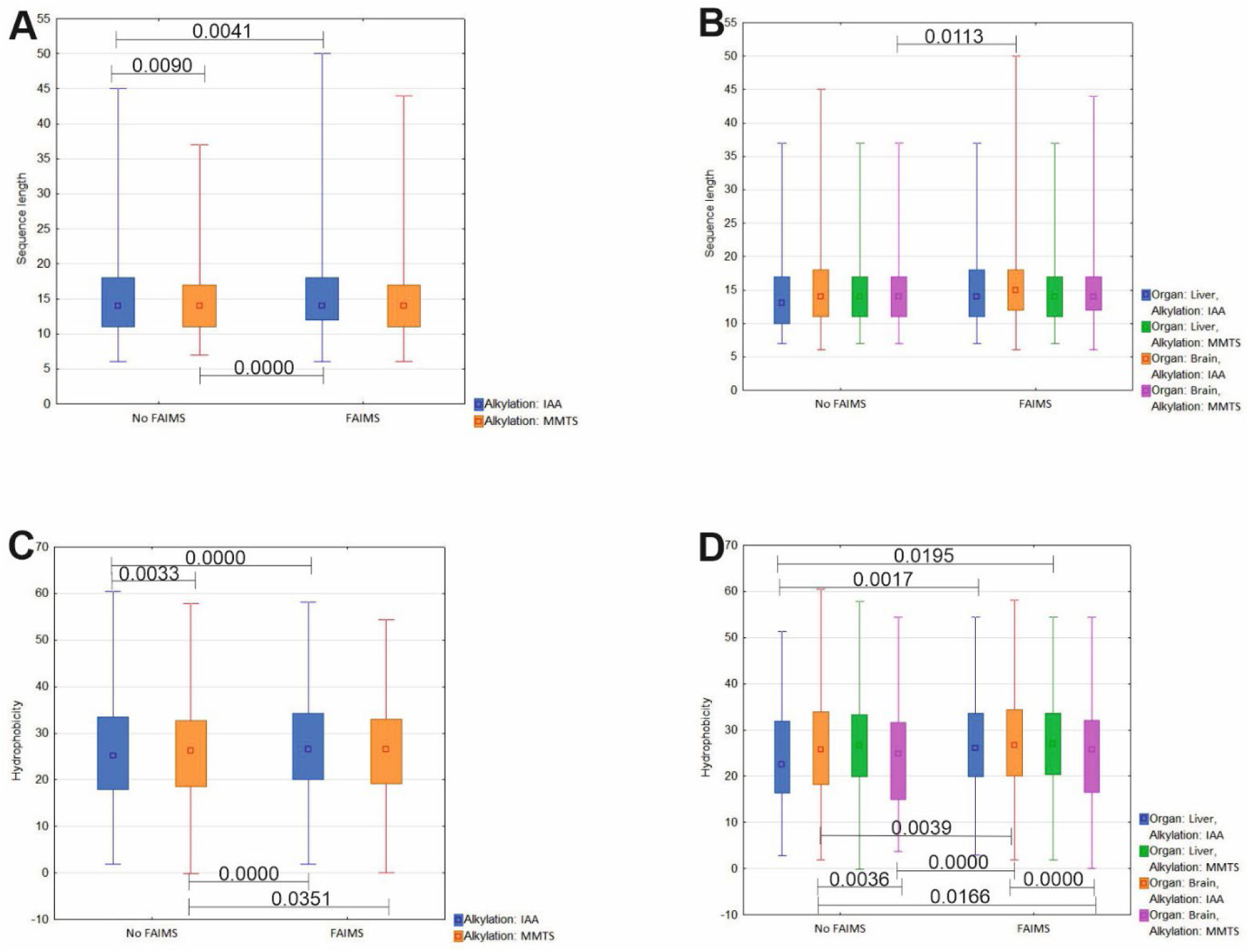
Box-plots of N-Hcy-peptides acquired from HTL-treated samples with FAIMS and no-FAIMS methods, either analyzed together for liver and brain (A, C) or separately for the organs (B, D). Median (square), upper and lower quartiles (box) and min-max values (whiskers) are shown. P-values were calculated with RIR Tuckey’s post-hoc test.

Sequence length of liver *in vitro N*-Hcy-peptides acquired with FAIMS is not significantly different from those acquired without FAIMS independently on alkylation method. Sequence length of brain *in vitro N*-Hcy-peptides acquired with FAIMS is also not significantly different from those acquired without FAIMS independently on alkylation method.

#### Hydrophobicity

*N*-Hcy-peptides (liver and brain) detected with FAIMS are significantly more hydrophobic than *N*-Hcy-peptides identified without FAIMS, regardless of alkylation method (Figure 8 C). In addition, *N*-Hcy-peptides identified following alkylation with IAA are more hydrophobic than those identified after MMTS treatment.

Hydrophobicity of liver *in vitro N*-Hcy-peptides acquired with FAIMS is significantly higher than those acquired without FAIMS in case of IAA but not-MMTS-alkylated peptides. Similar results were obtained for brain *in vitro N*-Hcy-peptides (Figure 8 D).

#### Theoretical pI

Theoretical pI of *N*-Hcy-peptides (both brain and liver) shows no significant difference with regard to acquisition and alkylation method (data not shown).

#### Unique and overlapping proteins, peptides, and N-Hcy-peptides detected with and without FAIMS

Next, we sought to investigate the degree of overlap of proteins, peptides and *N*-Hcy-peptides identified with and without FAIMS. The analyses were performed collectively for liver and brain, native and HTL-treated samples, alkylated with MMTS. As shown in Figure 9, there is 78, 60 and nearly 40% overlap in proteins, peptides and *N*-Hcy-peptides, respectively, identified with and without FAIMS. Twenty one, 34 and 56% of the identified proteins, peptides and *N*-Hcy-peptides, respectively, are unique for the FAIMS method, while the conventional approach gives a low fraction of unique identifications (1, 6 and 4% for proteins, peptides and *N*-Hcy-peptides, respectively). Overall, the most significant gain from introducing FAIMS is observed in the number of identified peptides carrying PTMs, which confirms its usefulness for detecting low-abundant entities, not easily reached during typical LC-MS DDA run.

**Figure 9.**
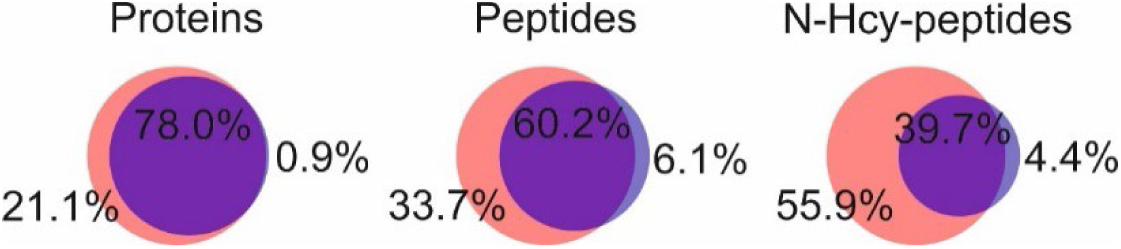
Venn diagrams showing the overlap of proteins, peptides and N-Hcy-peptides for FAIMS (orange) and no FAIMS (blue) acquisitions of brain and liver, native and HTL-treated samples, alkylated with MMTS.

#### Unique and overlapping N-Hcy-peptides detected with different reducing/alkylating agents

For native and HTL-treated samples (liver and brain), there is less than 3% and 12% overlap in *N*-Hcy-peptides identified with the use of two tested reducing/alkylating agents, respectively (Figure 10).

**Figure 10.**
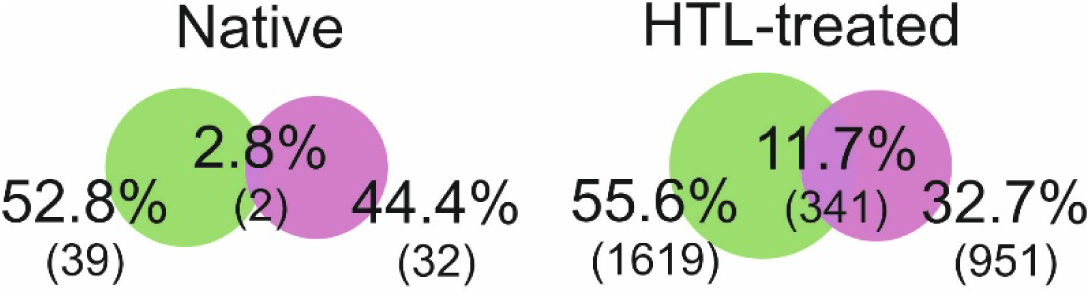
Venn diagrams showing the overlap of N-Hcy-peptides for MMTS (green) and IAA (fuchsia) alkylations of liver and brain protein samples (total of acquired with and without FAIMS). Separate Venn diagrams are shown for native and HTL-treated samples.

#### Biological relevance of identified in vivo N-Hcy-proteins

We have identified 94 and 69 *in vivo N*-Hcy-peptides and *N*-Hcy proteins, respectively. Forty two *N*-Hcy-proteins were found in the liver and 34 in the brain samples. Seven (10 %) of *N*-Hcy-proteins identified in our study, were detected in both organs, i.e. actin, septin-14, β-actin-like protein 2, plastin-3, pleckstrin homology domain-containing family O member 1, ubiquitin-conjugating enzyme E2 J2, and mitochondrial proton/calcium exchanger protein. The use of IAA alkylating agent yielded 45 and 34 of *N*-Hcy-peptides and *N*-Hcy-proteins, respectively, while MMTS resulted in the identification of 49 *N*-Hcy-peptides and 38 *N*-Hcy-proteins. The full list of identified *N*-Hcy-proteins and examples of MS/MS spectra of identified *N*-Hcy-peptides is shown in Supplementary Materials (Table 3 S, Figure 4 S, respectively).

Among *N*-Hcy-proteins are molecules involved in 1) **cell division**: G protein subunit alpha i3 (Gnai3), cell division cycle 42 (Cdc42), cyclin G2 (Ccng2), cyclin Y (Ccny), septin 14 (Septin14), septin 7 (Septin7); 2) **transport**: ADP ribosylation factor guanine nucleotide exchange factor 2 (Arfgef2), ATP synthase peripheral stalk subunit d (Atp5pd), Kv channel interacting protein 4 (Kcnip4), StAR related lipid transfer domain containing 10 (Stard10), USO1 vesicle docking factor (Uso1), adaptor-related protein complex 2, alpha 1 subunit (Ap2a1), fatty acid binding protein 1, live r (Fabp1), glycoprotein m6b (Gpm6b), hemoglobin alpha, adult chain 1 (Hba-a1), hemoglobin, beta adult major chain (Hbb-b1), leucine zipper-EF-hand containing transmembrane protein 1 (Letm1), major urinary protein 17 (Mup17), purinergic receptor P2X, ligand-gated ion channel, 1 (P2rx1), rabaptin, RAB GTPase binding effector protein 1 (Rabep1), vacuolar protein sorting 13B (Vps13b); and 3) **lipid metabolism**: arachidonate lipoxygenase 3 (Aloxe3), geranylgeranyl diphosphate synthase 1 (Ggps1), glutathione S-transferase, alpha 3 (Gsta3), glutathione S-transferase, pi 1 (Gstp1), glutathione S-transferase, pi 2 (Gstp2), peroxiredoxin 6 (Prdx6).

Next, the functional enrichment in the brain, liver and all *in vivo N*-Hcy-proteins was analyzed. STRING [35] analysis of brain *N*-Hcy-proteins has found functional enrichment in the network of cellular component and subcellular localization, where neuron projection was fund to be enriched (eleven and ten proteins, respectively; FDR q-value = 0.0356 and 0.0061, respectively). Among brain *N*-Hcy-proteins there are e.g. β-actin-like protein 2, actin, plastin-3, septin-14, and neurogranin. According to STRING [35] functional enrichment in the network of liver *N*-Hcy-proteins was found for biological processes (detoxification, response to toxic substance, and response to reactive oxygen species; FDR = 0.0034, 0.0034, 0.0061 and 0.0043, respectively) and molecular function (antioxidant activity, peroxidase activity, organic acid binding, haptoglobin binding and structural constituent of synapse; FDR = 0.0037, 0.0098, 0.0160, 0.0439 and 0.0404, respectively). When all *in vivo N*-Hcy-proteins identified in both organs where analyzed with STRING [35], functional enrichment in their network (Figure 12) was found in the following categories: biological processes, where two enriched terms were found, response to toxic substance (eight proteins; FDR q-value = 0.0171) and detoxification (six proteins; FDR = 0.0338); molecular function, where antioxidant activity was enriched (five proteins; FDR = 0.0443) and cellular component, with myelin sheath significantly enriched (seven proteins; FDR q-value = 0.0145).

**Figure 11.**
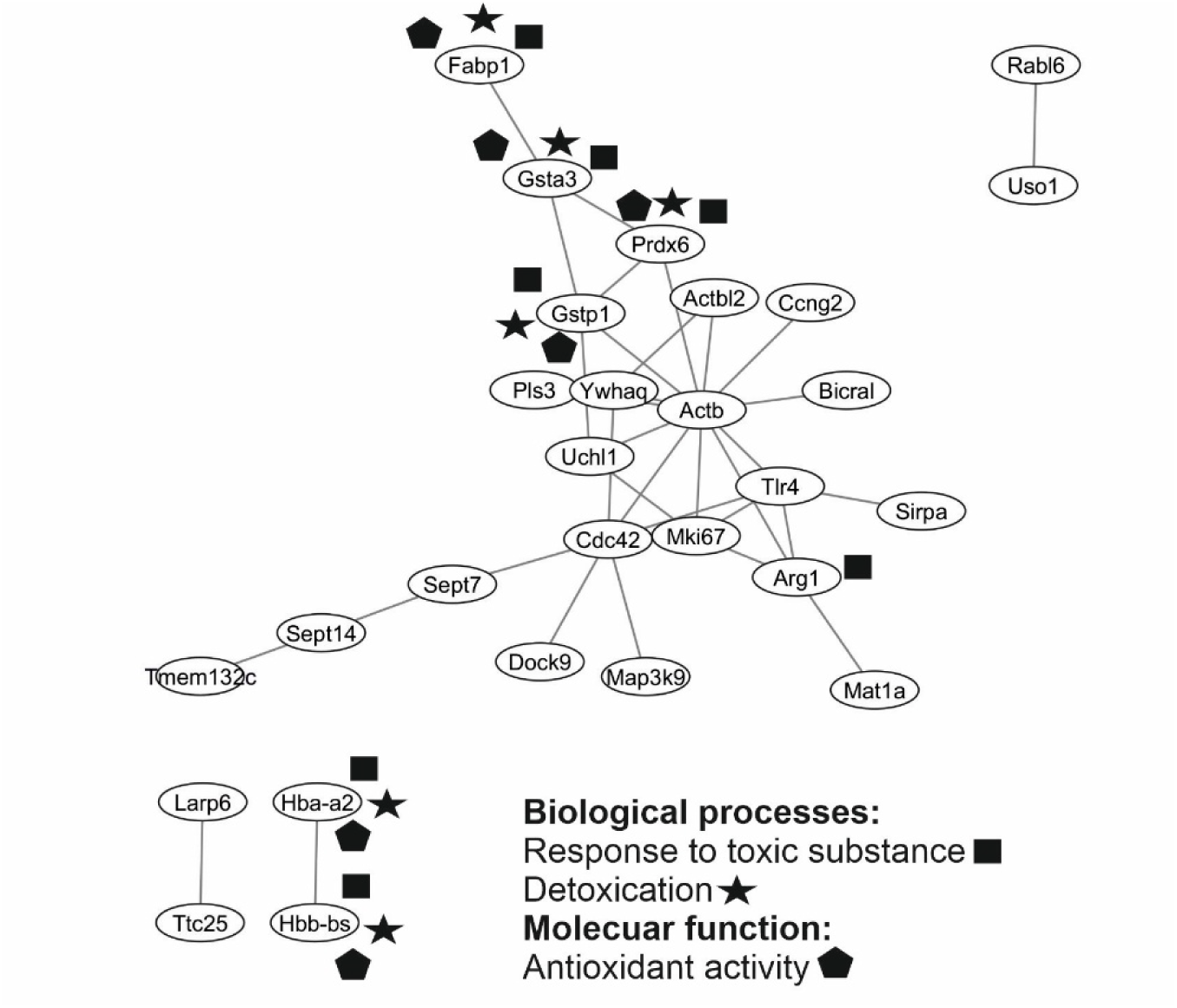
Functional interaction network of in vivo N-Hcy-proteins. Network generated with STRING.

**Figure 12.**
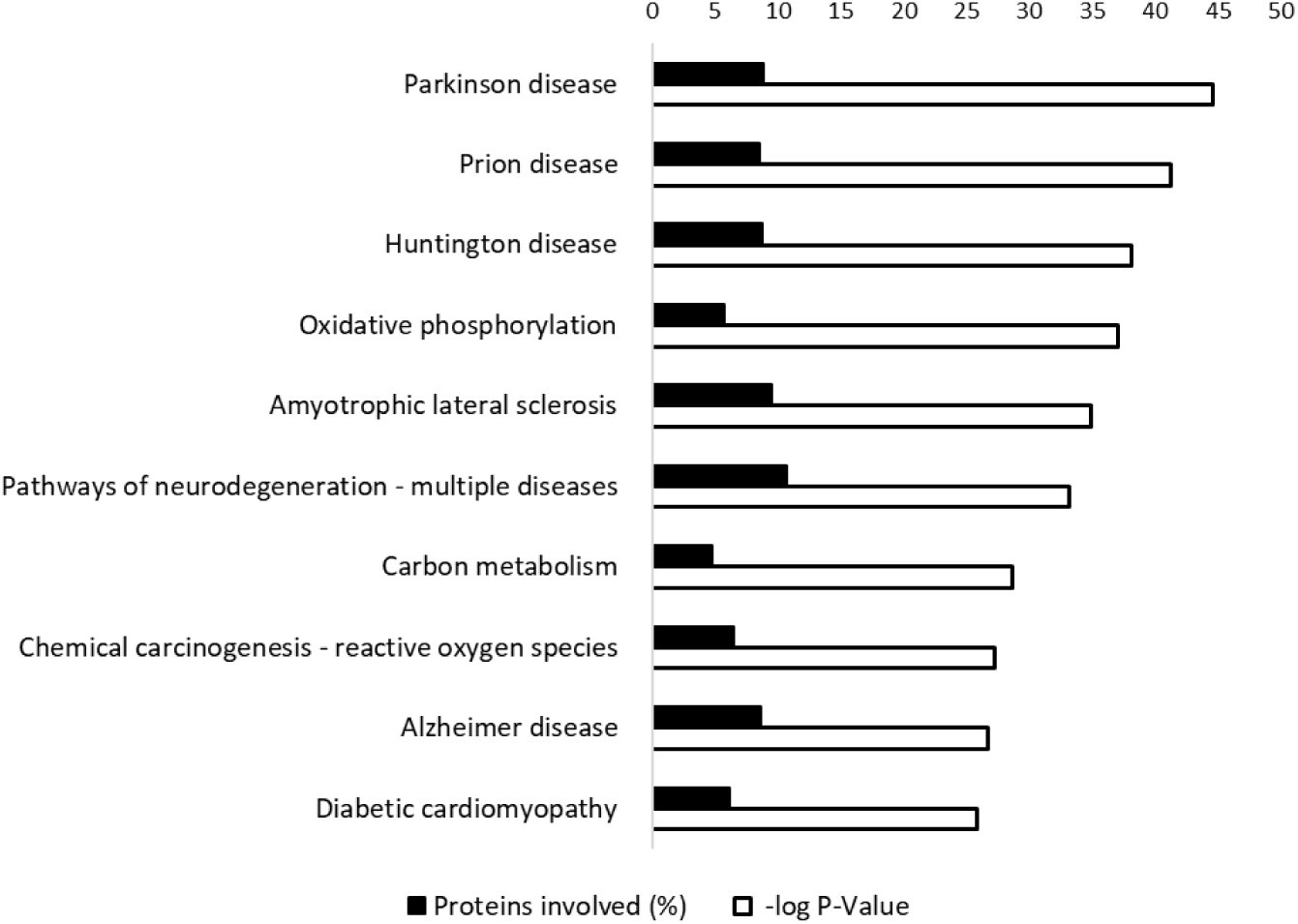
KEGG pathway term enrichment analysis. Functional annotation chart for in vitro N-Hcy-proteins.

#### N-Homocysteinylation in vitro

To better examine how FAIMS enhances the *N*-homocysteinylation detection, we introduced positive controls, in which liver and brain protein were incubated with HTL before MS analysis. In total 1,198 brain and liver *N*-Hcy-proteins were identified. The full list of *in vitro N*-Hcy-proteins is presented in Supplementary Materials (Table 4 S). KEGG pathway term enrichment analysis performed with DAVID [36, 37] showed that among *in vitro N*-Hcy-proteins, top ten KEGG pathways with highest – log *P*-value are Parkinson disease, prion disease, Huntington disease, oxidative phosphorylation, amyotrophic lateral sclerosis, pathways of neurodegeneration – multiple diseases, carbon metabolism, carcinogenesis – reactive oxygen species, Alzheimer disease, and diabetic cardiomyopathy (Figure 12).

## DISCUSSION

There is a limited number of proteome-wide studies of *N*-homocysteinylation, mainly due to the low rate of the modification in proteins, which complicates the detection when rich protein background is present. To our knowledge, this is the first proteome-wide study of protein *N*-homocysteinylation in a mouse *in vivo* model. Using liver and brain samples, we demonstrate that *N*-Hcy-sites mapping can be enhanced by FAIMS, particularly after diligent optimization of measurement parameters.

We began our study with FAIMS settings optimization. Yan et al. who employed FAIMS for human cysteinome profiling, have found that using three CV values in single run afforded more than twofold increase in coverage compared with one CV [27]. We also learnt that the highest yield of *N*-Hcy-proteins and *N*-Hcy-peptides was obtained using three-CV setting, i.e. F1 -45 -65 -85, F2 -35 -55 -75. Similarly to Hebert et al. [22], we demonstrated that combining multiple runs with single CVs (external CV stepping) is less efficient than longer runs of multi FAIMS with internal CV stepping. The CV settings and workflow optimized for the identification of *N*-Hcy-peptides could be a valuable for other experiments: as reported by Hebert et al. [22], peptides from different organisms behave similarly in FAIMS analysis, so our CV settings can presumably be successfully used to analyze samples of different origin. We expect that similar increase in the number of identifications could be observed also for other modifications, although each analyzed PTM type may require individual CV setting optimization.

In general, incorporating FAIMS into the workflow was advantageous for *N*-Hcy-sites identification, giving 1.3-7-fold increase in the yield of identified *N*-Hcy-peptides. Intriguingly, not all proteins, *N*-Hcy-proteins and *N*-Hcy-peptides acquired in the conventional approach, were identified with the use of FAIMS. We found that 1, 6 and 4% for proteins, peptides and *N*-Hcy-peptides, were uniquely found with the conventional approach, respectively. Similar observation was reported by Adoni et al. [23], who speculated that this effect may be due to differing frequency of MS/MS events when FAIMS is applied.

The *N*-Hcy-sites identified in this study are partially consistent with those identified in other studies. For example, we have identified three *in vivo N*-Hcy-sites in hemoglobin, one in the α-chain (K41) and two in β-chain (K67 and K83). Hemoglobin K67 and K83 were previously identified as *in vitro N*-homocysteinylation sites by Zang et al. [39]. Several *N*-Hcy-sites in both hemoglobin chains that were identified after *in vitro N*-homocysteinylation in our study also correspond to the sites identified by Zang et al. [39] in human plasma (α-chain: K8, K12, K17, K57, K140 and β-chain: K60, K83, K96, and K121).

Earlier, we discovered *N*-Hcy-sites *in vitro* in equine heart cytochrome c [9]. In this study, we identified the *N*-Hcy-site K100 as being modified *in vitro* in mice. We also detected three additional *in vitro N*-Hcy-sites of mouse cytochrome c: K28, K40 and K41.

Because histone *N*-homocysteinylation sites are particularly interesting due to possible cross-talk with other PTMs, which could influence pathomechanisms of HHcy-related diseases [40], we analyzed *N*-Hcy-sites identified in mouse histone proteins. We found that several histone *N*-Hcy-sites identified in our study overlap with those found previously. We confirmed the *in vitro N*-Hcy-site K57 of histone H3, which was reported by Xu at al. [40], in HTL-treated HEK293T cells (human kidney). We also identified additional *in vitro N*-Hcy-sites of histone H3: K80 and K123. K80 (K79) was also recently found to be *N*-homocysteinylated in the brain of human fetus with neural tube defects [41]. The histone H3 *N*-Hcy-site K80 was previously identified in *in vitro*-modified human histones [2], while the H3 *N*-Hcy-sites K57 and K80 were identified by Chen et al. [18] in HTL-treated HeLa cells and in this study. The other *N*-Hcy-sites identified in human histones by Chen et al. [18] were also identified as *N*-homocysteinylated *in vitro* in our study. In the histone H2A identified in the HTL-treated sample, we found the *N*-Hcy-K96, which was also identified by Zhang et al. [41]. In histone H4 we identified *in vitro N*-Hcy-sites, K60 and K80, which were recognized before [41], and in addition K32, K78, and K92 were found to be *N*-homocysteinylated in this study. In linker histone H1, we have identified K40, which was modified *in vitro* and, to our knowledge, had not been identified previously.

*N-*Homocysteinylation is increasingly linked to brain pathologies. For example, *N*-homocysteinylated tau is elevated in autopsy specimens of AD and vascular dementia [42]. α-Synuclein undergoes *N*-homocysteinylation and aggravates neurotoxicity in a TgA53T mouse model of Parkinson’s disease [12]. *N*-Homocysteinylation of amyloid β peptide increases its neurotoxicity *in vitro* [44]. We identified several *N*-Hcy-proteins that are associated with neurodegenerative diseases. For example, we showed that mouse brain microtubule-associated protein tau undergoes *N*-homocysteinylation *in vitro* at K466, K472, K555, K566 and K603, while amyloid β precursor protein is modified at K393. We also identified *N*-Hcy-sites in synuclein, i.e. K20 and K34, K12 and K57, K24 and K32 in α, β, γ chains, respectively. Zhou et al. [12] found K34, K43, K58, K60, K80 and K96 to be modified by Hcy-thiolactone. Targeting of *N*-homocysteinylated proteins associated with diseases could be a future therapeutic strategy against those diseases.

In conclusion, application of FAIMS in the analysis of protein *N*-homocysteinylation significantly increases the number of identified *N*-Hcy-Lys sites. Ninety-four *in vivo N*-Hcy-peptides were identified in mouse liver and brain proteins. Both reduction/alkylation methods resulted in a similar number of identified *N*-Hcy-peptides but due to low overlap in the identifications between the two approaches, we recommend to employ both alkylation methods. This optimized workflow helped us to establish a methodology suitable for deep proteome *N*-homocysteinylation profiling while running the sample only twice and thus reducing the overall cost and time needed for the analysis.

## ACKNOWLEDGEMENTS

We would like to thank Hieronim Jakubowski† for providing resources and funding acquisition for this study. The equipment used at the Institute of Biochemistry and Biophysics, PAS, Warsaw, was founded in part by the Centre for Preclinical Research and Technology (CePT), a project co-sponsored by European Regional Development Fund and Innovative Economy, The National Cohesion Strategy of Poland.

