## Supplementary Materials for "High field asymmetric waveform ion mobility spectrometry improves *N*-homocysteinylation mapping in mouse liver and brain proteins"

*Table 1 S. Median (min-max) of peptide characteristics depending on CVs used during optimization phase (F1: -45, -65, -85; F2: -35, -55, -75; F3: -40, -60, -80; F4: -65; F5: -55; F6: -50; F7: -45; F8: -35, -45, -55, -65, -75 or F9: -30, -40, -50, -60, -70 V).*

| | F1 | F2 | F3 | F4 | F5 | F6 | F7 | F8 | F9 |
|---|---|---|---|---|---|---|---|---|---|
| Sequence length | 15 (6-45) | 14 (6-45) | 14 (6-45) | 15 (6-42) | 16 (6-42) | 16 (6-42) | 16 (6-42) | 15 (6-45) | 15 (6-45) |
| Hydrophobicity | 24.92 (0.02-54.73) | 24.57 (1.84-53.11) | 23.75 (0.02-50.99) | 25.98 (-2.30-52.11) | 26.57 (0.02-63.91) | 27.29 (0.02-54.49) | 26.65 (1.84-54.49) | 25.41 (0.02-53.38) | 25.72 (0.02-53.38) |
| Theoretical pI | 7.1 (3.4-12.5) | 7.1 (3.4-12.5) | 7,1 (3.5-12.5) | 7.0 (3.6-12.5) | 7.1 (3.6-11.5) | 7.1 (3.4-12.5) | 7.1 (3.4-12.5) | 7.0 (3.5-12.5) | 7.0 (3.4-12.5) |

*Table 2 S. RIR Tukey's test P-values for comparisons of sequence length and hydrophobicity of N-Hcy-peptides acquired with different CV settings (F1: -45, -65, -85; F2: -35, -55, -75; F3: -40, -60, -80; F4: -65; F5: -55; F6: -50; F7: -45; F8: -35, -45, -55, -65, -75 or F9: -30, -40, -50, -60, -70 V).*

| | RIR Tukey's Test; variable: Sequence length, in red p <0.05000 | | | | | | | | |
|---|---|---|---|---|---|---|---|---|---|
| CV | {1} M=15.637 | {2} M=15.393 | {3} M=15.133 | {4} M=15.074 | {5} M=16.260 | {6} M=17.019 | {7} M=16.982 | {8} M=16.208 | {9} M=16.202 |
| F1 {1} | | 0.894582 | 0.109919 | 0.094756 | 0.064769 | 0.000010 | 0.000010 | 0.069039 | 0.078187 |
| F2 {2} | 0.894582 | | 0.877274 | 0.789332 | 0.000938 | 0.000010 | 0.000010 | 0.000687 | 0.000846 |
| F3 {3} | 0.109919 | 0.877274 | | 0.999998 | 0.000013 | 0.000010 | 0.000010 | 0.000012 | 0.000012 |
| F4 {4} | 0.094756 | 0.789332 | 0.999998 | | 0.000015 | 0.000010 | 0.000010 | 0.000013 | 0.000014 |
| F5 {5} | 0.064769 | 0.000938 | 0.000013 | 0.000015 | | 0.044626 | 0.108629 | 1.000000 | 0.999999 |
| F6 {6} | 0.000010 | 0.000010 | 0.000010 | 0.000010 | 0.044626 | | 1.000000 | 0.011683 | 0.010963 |
| F7 {7} | 0.000010 | 0.000010 | 0.000010 | 0.000010 | 0.108629 | 1.000000 | | 0.039519 | 0.037349 |
| F8 {8} | 0.069039 | 0.000687 | 0.000012 | 0.000013 | 1.000000 | 0.011683 | 0.039519 | | 1.000000 |
| F9 {9} | 0.078187 | 0.000846 | 0.000012 | 0.000014 | 0.999999 | 0.010963 | 0.037349 | 1.000000 | |

| | RIR Tukey's Test; variable: Hydrophobicity, in red p <0.05000 | | | | | | | | |
|---|---|---|---|---|---|---|---|---|---|
| CV | {1} M=25.455 | {2} M=25.094 | {3} M=24.357 | {4} M=25.335 | {5} M=26.270 | {6} M=27.188 | {7} M=26.943 | {8} M=25.838 | {9} M=25.955 |
| F1 {1} | | 0.961772 | 0.015507 | 0.999994 | 0.387633 | 0.000191 | 0.007738 | 0.969232 | 0.867863 |
| F2 {2} | 0.961772 | | 0.325817 | 0.998893 | 0.035788 | 0.000011 | 0.000217 | 0.406406 | 0.215190 |
| F3 {3} | 0.015507 | 0.325817 | | 0.131949 | 0.000021 | 0.000010 | 0.000010 | 0.000698 | 0.000188 |
| F4 {4} | 0.999994 | 0.998893 | 0.131949 | | 0.320601 | 0.000263 | 0.007387 | 0.918799 | 0.777774 |
| F5 {5} | 0.387633 | 0.035788 | 0.000021 | 0.320601 | | 0.441950 | 0.862395 | 0.974520 | 0.996932 |
| F6 {6} | 0.000191 | 0.000011 | 0.000010 | 0.000263 | 0.441950 | | 0.999848 | 0.023932 | 0.060032 |
| F7 {7} | 0.007738 | 0.000217 | 0.000010 | 0.007387 | 0.862395 | 0.999848 | | 0.201723 | 0.349763 |
| F8 {8} | 0.969232 | 0.406406 | 0.000698 | 0.918799 | 0.974520 | 0.023932 | 0.201723 | | 0.999997 |
| F9 {9} | 0.867863 | 0.215190 | 0.000188 | 0.777774 | 0.996932 | 0.060032 | 0.349763 | 0.999997 | |

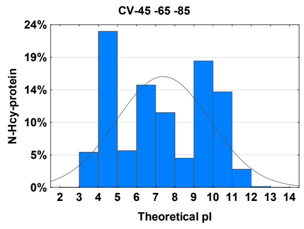


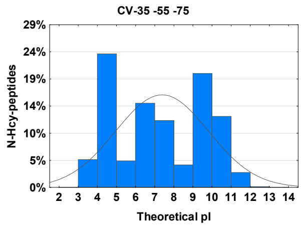


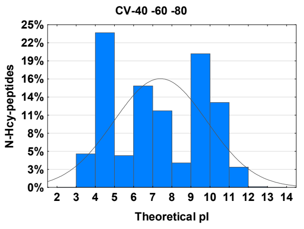

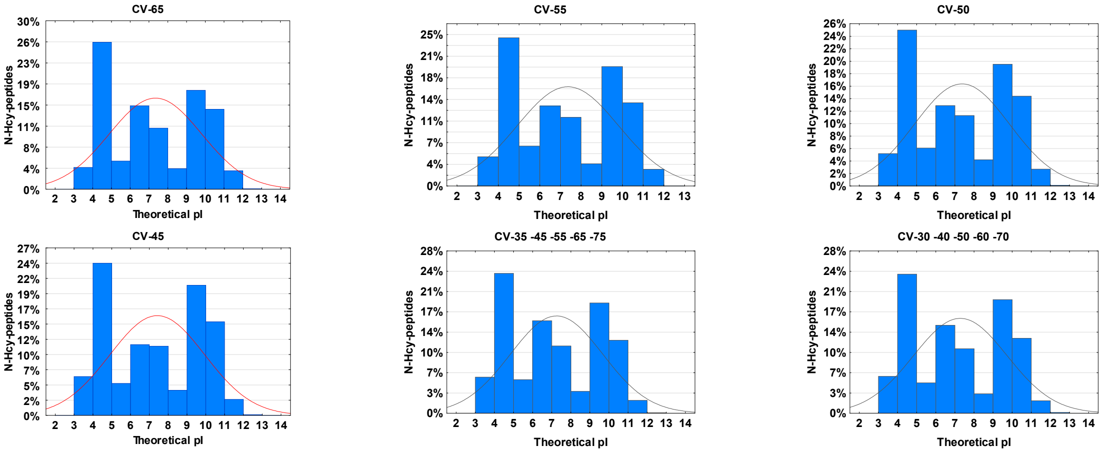


*Figure 1 S. Histograms of N-Hcy-peptides' theoretical pI distributions for each CV tested during optimization phase.*

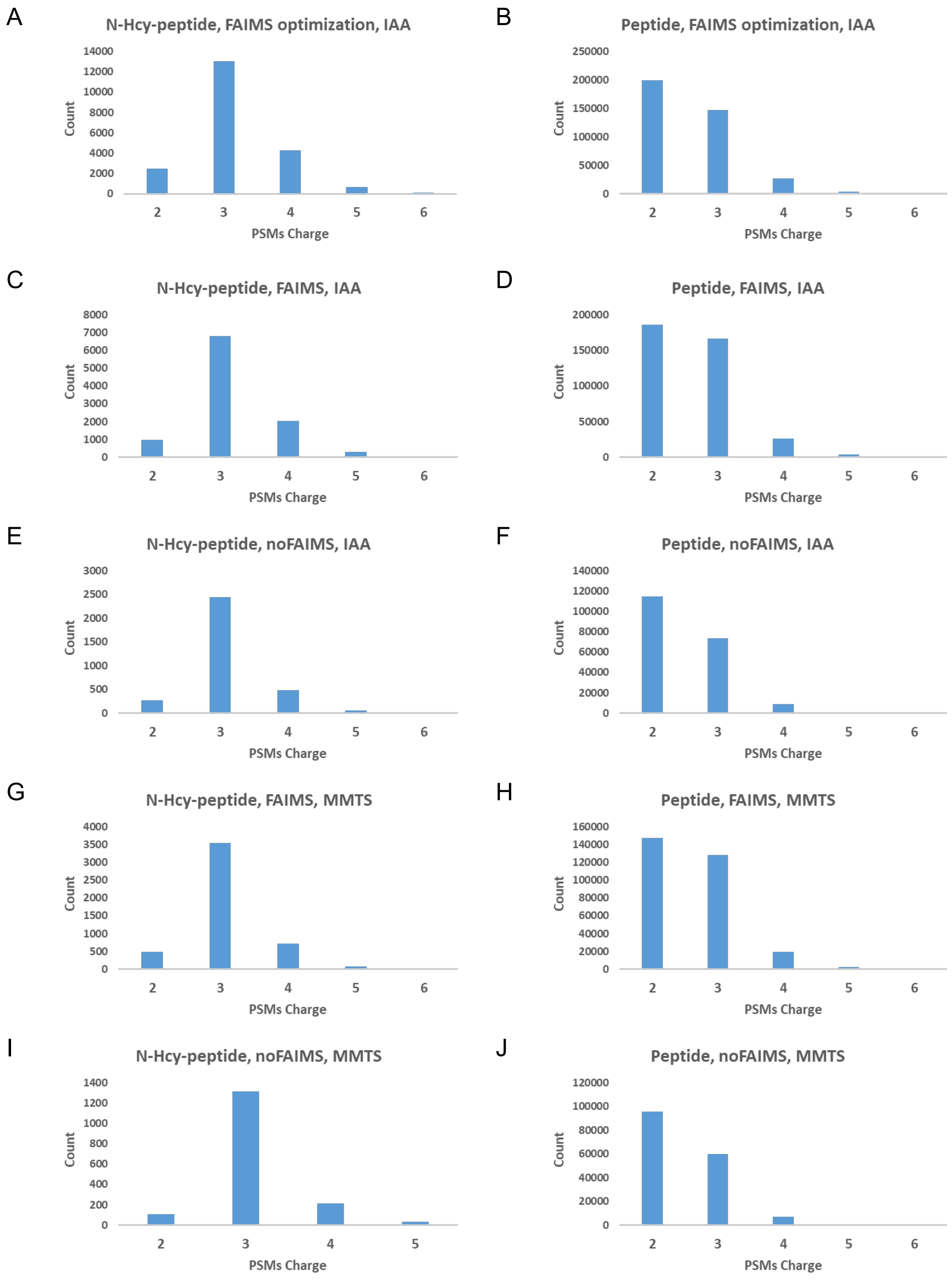


*Figure 2 S. Histograms showing charge states for fragmentation spectra (PTMs, Peptide-Spectrum Match) of N-Hcy-peptides (A, C, E, G, H) and non-N-Hcy-peptides (B, D, F, H, J) obtained during optimization phase (A, B). and during the experimental phase with IAA (C, D, E, F) and MMTS (G, H, I, J) alkylation method with (C, D, G, H) and without (E, F, I, J) FAIMS.*

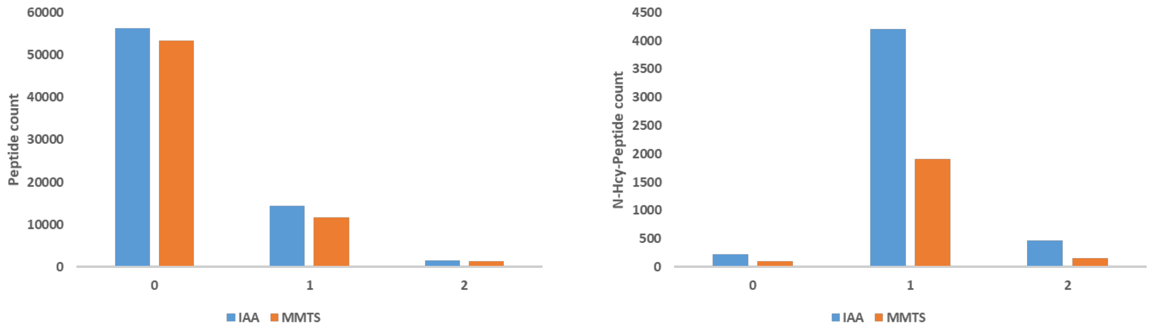


*Figure 3 S. The number of missed cleavages in all peptides (A) and N-Hcy-peptides (B).*

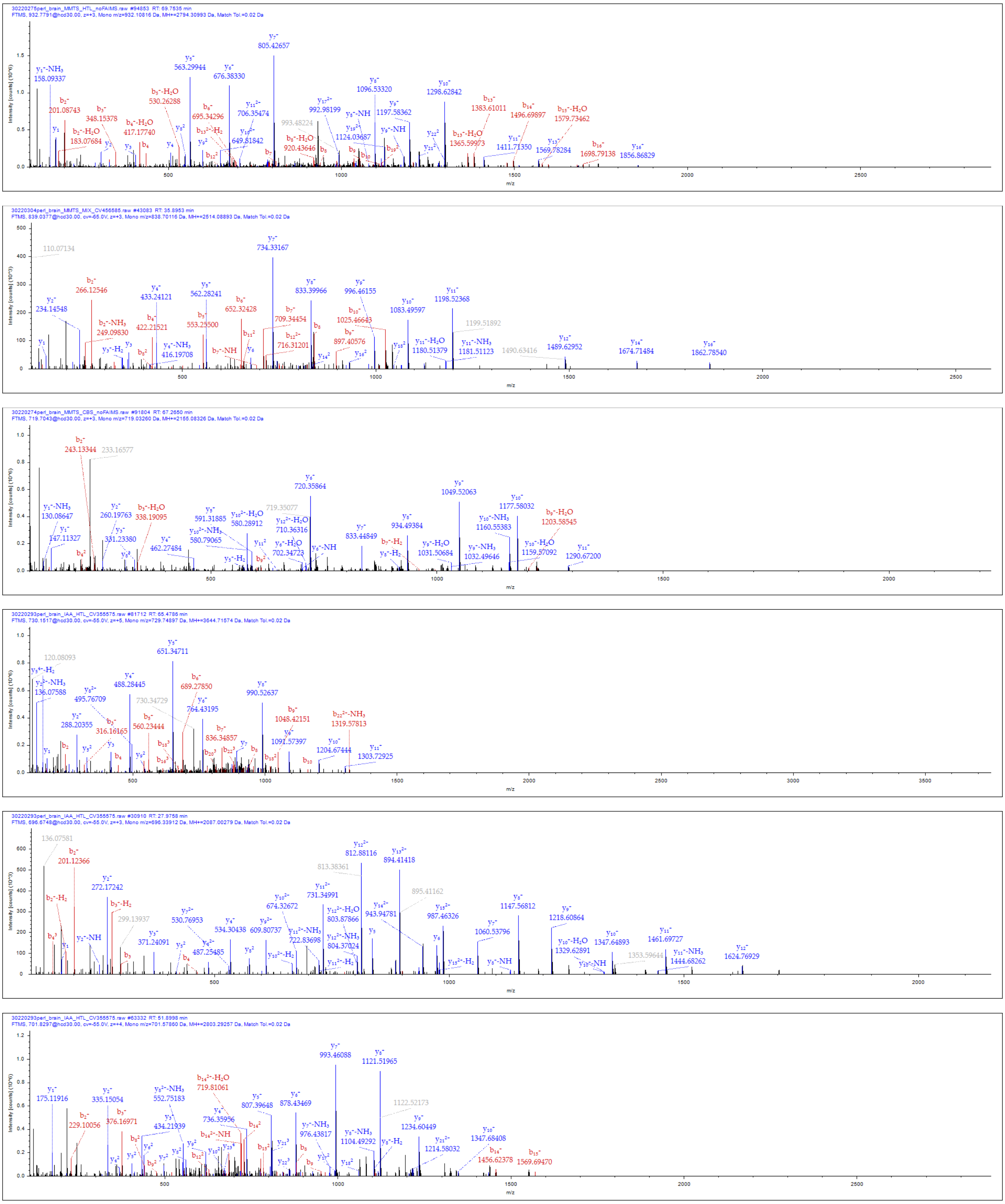

*Figure 4 S. Fragmentation spectra of N-Hcy-peptides identified in the study.*
