## Supplemental Table 3 S for "High field asymmetric waveform ion mobility spectrometry improves *N*-homocysteinylation mapping in mouse liver and brain proteins"

*Table 3 S. List of in vivo N-Hcy-proteins identified in the study.*

| | Master Protein Accessions | Protein name (Gene Name) | Annotated Sequence | Modifications | Positions in Master Proteins | Theo. MH+ [Da] | Organ/ Method/Alkylation |
|---|---|---|---|---|---|---|---|
| 1 | P60710; P62737; P68134; P63260 | actin, beta(Actb); actin alpha 2, smooth muscle, aorta(Acta2); actin alpha 1, skeletal muscle(Acta1); actin, gamma, cytoplasmic 1(Actg1) | [R].MQKEITALAPSTMK.[I] | 1xOxidation [M]; 1xHomocysteineIAA [K3] | [313-326]; [315-328]; [315-328]; [313-326] | 1738,85 | Liver/ FAIMS/IAA |
| 2 | P60710; P62737; P68134; P63260 | actin, beta(Actb); actin alpha 2, smooth muscle, aorta(Acta2); actin alpha 1, skeletal muscle(Acta1); actin, gamma, cytoplasmic 1(Actg1) | [R].MQKEITALAPSTMK.[I] | 1xOxidation [M]; 1xHomocysteineIAA [K3] | [313-326]; [315-328]; [315-328]; [313-326] | 1738,85 | Brain/ FAIMS/IAA |
| 3 | Q8BFZ3 | actin, beta-like 2(Actbl2) | [R].MQKEIVTLAPSTMK.[I] | 1xHomocysteineMMTS [K3] | [314-327] | 1739,86 | Liver/ FAIMS/MMTS |
| 4 | Q8BFZ3 | actin, beta-like 2(Actbl2) | [R].MQKEIVTLAPSTMK.[I] | 1xHomocysteineMMTS [K3] | [314-327] | 1739,86 | Brain/ FAIMS/MMTS |
| 5 | P17426 | adaptor-related protein complex 2, alpha 1 subunit(Ap2a1) | [R].SNAKQIVSEMLR.[Y] | 1xHomocysteineIAA [K4] | [400-411] | 1549,78 | Brain/ FAIMS/IAA |
| 6 | A2A5R2 | ADP ribosylation factor guanine nucleotide exchange factor 2(Arfgef2) | [K].LSMKPLGEGPPDPK.[S] | 1xOxidation [M3]; 1xHomocysteineIAA [K4] | [391-404] | 1655,81 | Brain/ No FAIMS/IAA |
| 7 | E9PVX6 | antigen identified by monoclonal antibody Ki 67(Mki67) | [R].NTLKEPVGDSINVEEVKK.[S] | 1xHomocysteineMMTS [K17] | [2691-2708] | 2162,08 | Liver/ No FAIMS/MMTS |
| 8 | Q9WV07 | arachidonate lipoxygenase 3(Aloxe3) | [R].DDGLKIWAAIER.[F] | 1xHomocysteineMMTS [K5] | [510-521] | 1549,75 | Liver/ FAIMS/MMTS |
| 9 | Q61176 | arginase, liver(Arg1) | [R].EGNHKPGTDYLKPPK.[-] | 1xHomocysteineMMTS [K] | [309-323] | 1843,88 | Liver/ No FAIMS/MMTS |
| 10 | Q61176 | arginase, liver(Arg1) | [R].DHGDLAFVDVPNDSSFQIVKNPR.[S] | 1xHomocysteineMMTS [K20] | [49-71] | 2733,28 | Liver/ FAIMS/MMTS |

| | | | | | | | |
|---|---|---|---|---|---|---|---|
| 11 | Q9DCX2 | ATP synthase peripheral stalk subunit d(Atp5pd) | [R].LASLSEKPPAIDWAYYR.[A] | 1xHomocysteineMMTS [K7] | [42-58] | 2143,04 | Liver/ No FAIMS/MMTS |
| 12 | Q91XV3 | brain abundant, membrane attached signal protein 1(Basp1) | [K].ESTEEKPKDAADGEAK.[A] | 1xHomocysteineIAA [K] | [53-68] | 1878,84 | Brain/ FAIMS/IAA |
| 13 | Q8CHH5 | BRD4 interacting chromatin remodeling complex associated protein like(Bicral) | [K].GSGEPQPDLQLTKSLEK.[T] | 2xHomocysteineMMTS [K13; K17] | [989-1005] | 2152,97 | Liver/ No FAIMS/MMTS |
| 14 | P16015 | carbonic anhydrase 3(Car3) | [K].YAAELHLVHWNPKYNTFGEALK.[Q] | 1xHomocysteineMMTS [K13] | [114-135] | 2764,34 | Liver/ No FAIMS/MMTS |
| 15 | P16015 | carbonic anhydrase 3(Car3) | [-].MAKEWGYASHNGPDHWHELYPIAK.[G] | 1xMet-loss+Acetyl [N-Term]; 1xHomocysteineMMTS [K3] | [1-24] | 2911,31 | Liver/ FAIMS/MMTS |
| 16 | P60766 | cell division cycle 42(Cdc42) | [-].MQTIKCVVVGDGAVGK.[T] | 1xMethylthio [C6]; 1xHomocysteineMMTS [K5] | [1-16] | 1813,85 | Brain/ FAIMS/MMTS |
| 17 | Q8BI22 | centrosomal protein 128(Cep128) | [R].FLSSPQFSHSLPVFTK.[R] | 1xHomocysteineIAA [K16] | [1049-1064] | 1996,00 | Liver/ No FAIMS/IAA |
| 18 | Q8BI22 | centrosomal protein 128(Cep128) | [R].FLSSPQFSHSLPVFTK.[R] | 1xHomocysteineIAA [K16] | [1049-1064] | 1996,00 | Liver/ FAIMS/IAA |
| 19 | O08918 | cyclin G2(Ccng2) | [R].VVFSKARPSVLALCLLNLEIETIK.[S] | 1xMethylthio [C14]; 2xHomocysteineMMTS [K5; K24] | [196-219] | 3028,56 | Brain/ FAIMS/MMTS |
| 20 | Q8BGU5 | cyclin Y(Ccny) | [R].SLAEANNLSFPLEPLSRERAHK.[L] | 1xHomocysteineMMTS [K22] | [280-301] | 2642,32 | Brain/ FAIMS/MMTS |
| 21 | Q9CYC6 | decapping mRNA 2(Dcp2) | [R].QPLQQKSHSNHGEVSDLLK.[A] | 1xHomocysteineIAA [K6] | [294-312] | 2319,15 | Liver/ FAIMS/IAA |
| 22 | Q8BIK4 | dedicator of cytokinesis 9(Dock9) | [K].DAVETQVGFSWLPLLKDGR.[V] | 1xHomocysteineIAA [K16] | [758-776] | 2305,16 | Liver/ No FAIMS/IAA |
| 23 | Q30KP6 | defensin beta 41(Defb41) | [K].KSYPEYGSLDLR.[K] | 1xHomocysteineIAA [K1] | [21-32] | 1601,76 | Liver/ FAIMS/IAA |
| 24 | P12710 | fatty acid binding protein 1, liver(Fabp1) | [K].GIKSVTELNGDTITNTMTLGDIVYK.[R] | 1xHomocysteineMMTS [K3] | [97-121] | 2846,40 | Liver/ No FAIMS/MMTS |
| 25 | P12710 | fatty acid binding protein 1, liver(Fabp1) | [K].GIKSVTELNGDTITNTMTLGDIVYK.[R] | 1xHomocysteineMMTS [K3] | [97-121] | 2846,40 | Liver/ FAIMS/MMTS |
| 26 | P12710 | fatty acid binding protein 1, liver(Fabp1) | [K].GIKSVTELNGDTITNTMTLGDIVYK.[R] | 1xOxidation [M17]; 1xHomocysteineMMTS [K3] | [97-121] | 2862,39 | Liver/ FAIMS/MMTS |

| | | | | | | | |
|---|---|---|---|---|---|---|---|
| 27 | P12710 | fatty acid binding protein 1, liver(Fabp1) | [-].MNFSGKYQLQSQENFEPFMK.[A] | 1xMet-loss [N-Term]; 1xHomocysteineIAA [K6] | [1-20] | 2496,13 | Liver/ No FAIMS/IAA |
| 28 | P12710 | fatty acid binding protein 1, liver(Fabp1) | [-].MNFSGKYQLQSQENFEPFMK.[A] | 1xMet-loss [N-Term]; 1xHomocysteineIAA [K6] | [1-20] | 2496,13 | Liver/ FAIMS/IAA |
| 29 | E9Q8I9 | FRY microtubule binding protein(Fry) | [R].ELVEELHPLMKEALER.[R] | 1xOxidation [M10]; 1xHomocysteineMMTS [K11] | [1000-1015] | 2115,03 | Liver/ FAIMS/MMTS |
| 30 | E9Q8I9 | FRY microtubule binding protein(Fry) | [R].ELVEELHPLMKEALER.[R] | 1xHomocysteineMMTS [K11] | [1000-1015] | 2099,03 | Liver/ FAIMS/MMTS |
| 31 | Q9DC51; P08752; B2RSH2 | G protein subunit alpha i3(Gnai3); G protein subunit alpha i2(Gnai2); G protein subunit alpha i1(Gnai1) | [KR].EVKLLLLGAGESGK.[S] | 1xHomocysteineIAA [K3] | [33-46] | 1587,88 | Brain/ No FAIMS/IAA |
| 32 | Q9WTN0 | geranylgeranyl diphosphate synthase 1(Ggps1) | [K].VLTLDHPDAVKLFTR.[Q] | 1xHomocysteineMMTS [K11] | [103-117] | 1887,98 | Brain/ FAIMS/MMTS |
| 33 | Q9JK38 | glucosamine-phosphate N-acetyltransferase 1(Gnpnat1) | [-].MKPDETPMFDPSLLK.[E] | 1xMet-loss [N-Term]; 1xHomocysteineIAA [K2] | [1-15] | 1791,87 | Liver/ FAIMS/IAA |
| 34 | P30115 | glutathione S-transferase, alpha 3(Gsta3) | [-].MAGKPVLHYFDGR.[G] | 1xHomocysteineMMTS [K4] | [1-13] | 1653,77 | Liver/ FAIMS/MMTS |
| 35 | P19157 | glutathione S-transferase, pi 2(Gstp2) | [K].YVTLIYTNYENGKNDYVK.[A] | 1xHomocysteineMMTS [K13] | [104-121] | 2360,09 | Liver/ FAIMS/MMTS |
| 36 | Q91XE0 | glycine-N-acyltransferase(Glyat) | [K].LSSLDVVHAALVNKFWLFGGNER.[S] | 1xHomocysteineMMTS [K14] | [170-192] | 2735,38 | Brain/ FAIMS/MMTS |
| 37 | P35803 | glycoprotein m6b(Gpm6b) | [-].MKPAMETAAEENTEQSQER.[K] | 1xMet-loss+Acetyl [N-Term]; 1xHomocysteineMMTS [K2] | [1-19] | 2253,94 | Brain/ FAIMS/MMTS |
| 38 | Q4U2R1 | HECT and RLD domain containing E3 ubiquitin protein ligase 2(Herc2) | [R].LGHGDTVPLEEPKVISAFSGK.[Q] | 2xHomocysteineIAA [K13; K21] | [528-548] | 2529,25 | Brain/ FAIMS/IAA |
| 39 | P01942 | hemoglobin alpha, adult chain 1(Hba-a1) | [R].MFASFPTTKTYFPHFDVSHGSAQVK.[G] | 1xOxidation [M1]; 1xHomocysteineMMTS [K9] | [33-57] | 3009,37 | Liver/ No FAIMS/MMTS |
| 40 | P02088 | hemoglobin, beta adult major chain(Hbb-b1) | [K].KVITAFNDGLNHLDSLK.[G] | 1xHomocysteineMMTS [K1] | [67-83] | 2048,03 | Liver/ FAIMS/MMTS |

| | | | | | | | |
|---|---|---|---|---|---|---|---|
| 41 | P02088 | hemoglobin, beta adult major chain(Hbb-b1) | [K].VITAFNDGLNHLDSLKGTFASLSELHCDK.[L] | 1xMethylthio [C27]; 1xHomocysteineMMTS [K16] | [68-96] | 3354,56 | Liver/ FAIMS/MMTS |
| 42 | P02088 | hemoglobin, beta adult major chain(Hbb-b1) | [K].KVITAFNDGLNHLDSLK.[G] | 1xHomocysteineIAA [K1] | [67-83] | 2059,06 | Liver/ FAIMS/IAA |
| 43 | Q7TQK1 | integrator complex subunit 7(Ints7) | [K].HLEKILNVDEFVK.[R] | 1xHomocysteineIAA [K4] | [93-105] | 1757,93 | Brain/ No FAIMS/IAA |
| 44 | Q91V64 | isochorismatase domain containing 1(Isoc1) | [K].GLGSTVQEIDLTGVK.[L] | 1xHomocysteineMMTS [K15] | [162-176] | 1679,83 | Liver/ FAIMS/MMTS |
| 45 | E0CZ16 | kelch-like 3(Klhl3) | [R].TVTVNAAHMGKAFK.[V] | 1xHomocysteineMMTS [K11] | [25-38] | 1637,80 | Liver/ No FAIMS/MMTS |
| 46 | Q6PHZ8 | Kv channel interacting protein 4(Kcnip4) | [R].HRPEALELLEAQSKFTK.[K] | 1xHomocysteineMMTS [K14] | [68-84] | 2160,09 | Brain/ FAIMS/MMTS |
| 47 | Q8BN59 | La ribonucleoprotein 6, translational regulator(Larp6) | [K].AVLIGMKPPK.[K] | 1xHomocysteineIAA [K7] | [288-297] | 1227,70 | Liver/ No FAIMS/IAA |
| 48 | Q9Z2I0 | leucine zipper-EF-hand containing transmembrane protein 1(Letm1) | [K].LELAKFLQDTIEEMALK.[N] | 1xHomocysteineMMTS [K5] | [257-273] | 2155,08 | Liver/ No FAIMS/MMTS |
| 49 | Q9Z2I0 | leucine zipper-EF-hand containing transmembrane protein 1(Letm1) | [K].LELAKFLQDTIEEMALK.[N] | 1xHomocysteineMMTS [K5] | [257-273] | 2155,08 | Brain/ No FAIMS/MMTS |
| 50 | B5X0G2; P11589; P11588 | major urinary protein 17(Mup17); 2(Mup2); 1(Mup1) | [R].NFNVEKINGEWHTIILASDK.[R] | 1xHomocysteineMMTS [K6] | [27-46] | 2491,21 | Liver/ FAIMS/MMTS |
| 51 | B5X0G2; P11589; P11588 | major urinary protein 17(Mup17); major urinary protein 2(Mup2); major urinary protein 1(Mup1) | [R].NFNVEKINGEWHTIILASDK.[R] | 1xHomocysteineMMTS [K6] | [27-46] | 2491,21 | Liver/ FAIMS/MMTS |
| 52 | Q91X83 | methionine adenosyltransferase 1A(Mat1a) | [K].VACETVCKTGMVLLCGEITSVAMVDYQR.[V] | 3xCarbamidomethyl [C3; C7; C15]; 1xHomocysteineIAA [K8] | [55-82] | 3364,55 | Liver/ No FAIMS/IAA |
| 53 | Q3U1V8 | mitogen-activated protein kinase kinase kinase 9(Map3k9) | [R].TPSDGALKPTAAPAVLGSR.[S] | 1xHomocysteineMMTS [K8] | [885-903] | 1972,00 | Liver/ FAIMS/MMTS |
| 54 | A2ABU4 | myomesin family, member 3(Myom3) | [K].KLSHEIR.[N] | 1xHomocysteineIAA [K1] | [998-1004] | 1056,56 | Liver/ No FAIMS/IAA |

| | | | | | | | |
|---|---|---|---|---|---|---|---|
| 55 | P60761 | neurogranin(Nrgn) | [-].MDCCTESACSKPDDDILDIPLDDPGANAAAAK.[I] | 3xCarbamidomethyl [C3; C4; C9]; 1xMet-loss [N-Term]; 1xHomocysteineIAA [K11] | [1-32] | 3479,47 | Brain/ No FAIMS/IAA |
| 56 | P60761 | neurogranin(Nrgn) | [-].MDCCTESACSKPDDDILDIPLDDPGANAAAAK.[I] | 3xCarbamidomethyl [C3; C4; C9]; 1xMet-loss [N-Term]; 1xHomocysteineIAA [K11] | [1-32] | 3479,47 | Brain/ FAIMS/IAA |
| 57 | Q9D4B2 | outer dynein arm complex subunit 4(Odad4) | [K].EYLEKLLLDEDLIK.[G] | 2xHomocysteineIAA [K5; K14] | [197-210] | 2082,05 | Brain/ FAIMS/IAA |
| 58 | O08709 | peroxiredoxin 6(Prdx6) | [K].DINAYNGETPTEKLPFPIIDDK.[G] | 1xHomocysteineMMTS [K13] | [85-106] | 2653,25 | Liver/ FAIMS/MMTS |
| 59 | Q99K51 | plastin 3 (T-isoform)(Pls3) | [-].MDEMATTQISKDELDELKEAFAK.[V] | 1xMet-loss [N-Term]; 1xHomocysteineIAA [K11] | [1-23] | 2686,26 | Liver/ No FAIMS/IAA |
| 60 | Q99K51 | plastin 3 (T-isoform)(Pls3) | [-].MDEMATTQISKDELDELKEAFAK.[V] | 1xMet-loss [N-Term]; 1xHomocysteineIAA [K11] | [1-23] | 2686,26 | Liver/ FAIMS/IAA |
| 61 | Q99K51 | plastin 3 (T-isoform)(Pls3) | [-].MDEMATTQISKDELDELKEAFAK.[V] | 1xMet-loss [N-Term]; 1xHomocysteineIAA [K11] | [1-23] | 2686,26 | Brain/ No FAIMS/IAA |
| 62 | Q9JIY0 | pleckstrin homology domain containing, family O member 1(Plekho1) | [R].KAKDPPQSPPDSESEQLLLETER.[L] | 2xHomocysteineIAA [K1; K3] | [334-356] | 2942,39 | Liver/ FAIMS/IAA |
| 63 | Q9JIY0 | pleckstrin homology domain containing, family O member 1(Plekho1) | [R].KAKDPPQSPPDSESEQLLLETER.[L] | 2xHomocysteineIAA [K1; K3] | [334-356] | 2942,39 | Brain/ No FAIMS/IAA |
| 64 | P51576 | purinergic receptor P2X, ligand-gated ion channel, 1(P2rx1) | [R].FDILVDGKAGK.[F] | 1xHomocysteineIAA [K8] | [315-325] | 1336,69 | Liver/ FAIMS/IAA |
| 65 | Q8R4E6 | purine-rich element binding protein G(Purg) | [K].AWTRFGENFIK.[Y] | 1xHomocysteineMMTS [K11] | [309-319] | 1531,72 | Brain/ FAIMS/MMTS |
| 66 | Q5U3K5 | RAB, member RAS oncogene family-like 6(Rabl6) | [K].NISLSSEEEAEGLAGHPRVAPQQCSEPETK.[W] | 1xCarbamidomethyl [C24]; 1xHomocysteineIAA [K30] | [478-507] | 3424,57 | Brain/ FAIMS/IAA |
| 67 | O35551 | rabaptin, RAB GTPase binding effector protein 1(Rabep1) | [K].EEIASISSLKAELER.[I] | 1xHomocysteineMMTS [K10] | [736-750] | 1837,90 | Liver/ FAIMS/MMTS |

| | | | | | | | |
|---|---|---|---|---|---|---|---|
| 68 | Q9EPQ2 | retinitis pigmentosa GTPase regulator interacting protein 1(Rpgrip1) | [K].TSKVEKPEQR.[S] | 1xHomocysteineMMTS [K3] | [206-215] | 1364,67 | Brain/ No FAIMS/MMTS |
| 69 | P97868 | retinoblastoma binding protein 6, ubiquitin ligase(Rbbp6) | [K].KPHDHKAPYETKRPCEETKPVDK.[N] | 1xCarbamidomethyl [C15]; 1xHomocysteineIAA [K12] | [1541-1563] | 2964,45 | Liver/ No FAIMS/IAA |
| 70 | Q8C310 | roundabout guidance receptor 4(Robo4) | [R].KHVPWTLEQLR.[A] | 1xHomocysteineMMTS [K1] | [453-463] | 1569,80 | Brain/ FAIMS/MMTS |
| 71 | A7XUZ6 | selection and upkeep of intraepithelial T cells 6(Skint6) | [K].VIQSSIQSFSVDLETK.[E] | 1xHomocysteineIAA [K16] | [580-595] | 1954,98 | Brain/ No FAIMS/IAA |
| 72 | Q9DA97 | septin 14(Septin14) | [-].MAEKPTNTSVPIPGSEDPQK.[E] | 1xMet-loss+Acetyl [N-Term]; 1xHomocysteineIAA [K4] | [1-20] | 2211,06 | Liver/ No FAIMS/IAA |
| 73 | Q9DA97 | septin 14(Septin14) | [-].MAEKPTNTSVPIPGSEDPQK.[E] | 1xMet-loss+Acetyl [N-Term]; 1xHomocysteineIAA [K4] | [1-20] | 2211,06 | Brain/ FAIMS/IAA |
| 74 | O55131 | septin 7(Septin7) | [R].THMQDLKDVTNNVHYENYR.[S] | 1xHomocysteineMMTS [K7] | [291-309] | 2540,11 | Liver/ No FAIMS/MMTS |
| 75 | O55131 | septin 7(Septin7) | [R].THMQDLKDVTNNVHYENYR.[S] | 1xHomocysteineMMTS [K7] | [291-309] | 2540,11 | Liver/ FAIMS/MMTS |
| 76 | Q8BZT2 | SH3 domain containing ring finger 2(Sh3rf2) | [R].TVHSPEGHQMVEISTPMLISSNSPSVLTQHGDK.[A] | 1xHomocysteineMMTS [K33] | [313-345] | 3706,73 | Brain/ FAIMS/MMTS |
| 77 | P97797 | signal-regulatory protein alpha(Sirpa) | [K].NLTKNTDGTYNYTSLFLVNSSAHR.[E] | 1xHomocysteineIAA [K4] | [302-325] | 2890,38 | Liver/ FAIMS/IAA |
| 78 | Q9JMD3 | StAR related lipid transfer domain containing 10(Stard10) | [-].MEKPAASTEPQGSRPALGR.[E] | 1xMet-loss [N-Term]; 1xHomocysteineIAA [K3] | [1-19] | 2026,01 | Liver/ FAIMS/IAA |
| 79 | Q3URK3 | tet methylcytosine dioxygenase 1(Tet1) | [K].SLNKPNHGFDINK.[I] | 1xHomocysteineMMTS [K4] | [1932-1944] | 1646,78 | Liver/ FAIMS/MMTS |
| 80 | Q9QUK6 | toll-like receptor 4(Tlr4) | [K].RVTEFSAFLSLEK.[L] | 1xHomocysteineIAA [K13] | [434-446] | 1700,87 | Liver/ FAIMS/IAA |
| 81 | Q8CEF9 | transmembrane protein 132C(Tmem132c) | [R].AETAFFLK.[E] | 1xHomocysteineMMTS [K8] | [57-64] | 1089,51 | Brain/ FAIMS/MMTS |

| | | | | | | | |
|---|---|---|---|---|---|---|---|
| 82 | P68254; Q9CQV8; P63101 | tyrosine 3-monooxygenase/tryptophan 5-monooxygenase activation protein theta(Ywhaq); tyrosine 3-monooxygenase/tryptophan 5-monooxygenase activation protein, beta polypeptide(Ywhab); tyrosine 3-monooxygenase/tryptophan 5-monooxygenase activation protein, zeta polypeptide(Ywhaz) | [K].KEMQPTHPIR.[L] | 1xHomocysteineIAA [K1] | [158-167]; [160-169]; [158-167] | 1410,70 | Brain/ No FAIMS/IAA |
| 83 | Q9R0P9 | ubiquitin carboxy-terminal hydrolase L1(Uchl1) | [-].MQLKPMEINPEMLNK.[V] | 1xMet-loss [N-Term]; 1xHomocysteineMMTS [K4] | [1-15] | 1847,89 | Brain/ FAIMS/MMTS |
| 84 | Q9R0P9 | ubiquitin carboxy-terminal hydrolase L1(Uchl1) | [-].MQLKPMEINPEMLNK.[V] | 1xOxidation [M6]; 1xMet-loss [N-Term]; 1xHomocysteineIAA [K4] | [1-15] | 1874,92 | Brain/ No FAIMS/IAA |
| 85 | Q9R0P9 | ubiquitin carboxy-terminal hydrolase L1(Uchl1) | [-].MQLKPMEINPEMLNK.[V] | 1xMet-loss [N-Term]; 1xHomocysteineIAA [K4] | [1-15] | 1858,92 | Brain/ No FAIMS/IAA |
| 86 | Q6P073 | ubiquitin-conjugating enzyme E2J 2(Ube2j2) | [K].QLAAQSLVFNLKDK.[V] | 1xHomocysteineIAA [K12] | [141-154] | 1748,94 | Liver/ No FAIMS/IAA |
| 87 | Q6P073 | ubiquitin-conjugating enzyme E2J 2(Ube2j2) | [K].QLAAQSLVFNLKDK.[V] | 1xHomocysteineIAA [K12] | [141-154] | 1748,94 | Brain/ No FAIMS/IAA |
| 88 | Q9Z1Z0 | USO1 vesicle docking factor(Uso1) | [K].LDLEVTDSKK.[E] | 1xHomocysteineMMTS [K9] | [895-904] | 1310,63 | Liver/ No FAIMS/MMTS |
| 89 | Q9Z1Z0 | USO1 vesicle docking factor(Uso1) | [K].LDLEVTDSKK.[E] | 1xHomocysteineMMTS [K9] | [895-904] | 1310,63 | Liver/ FAIMS/MMTS |
| 90 | Q80TY5 | vacuolar protein sorting 13B(Vps13b) | [K].SGIPPSLVTLHIKDFLNGPADIYLDVSKPLK.[A] | 2xHomocysteineMMTS [K13; K28] | [1996-2026] | 3673,89 | Liver/ FAIMS/MMTS |
| 91 | P62761 | visinin-like 1(Vsnl1) | [K].QNSKLAPEVMEDLVK.[S] | 1xOxidation [M10]; 1xHomocysteineIAA [K4] | [4-18] | 1890,93 | Brain/ No FAIMS/IAA |
| 92 | Q811F1 | zinc finger and BTB domain containing 41(Zbtb41) | [R].KLLSSLQYNKNLLK.[Y] | 1xHomocysteineMMTS [K10] | [63-76] | 1825,01 | Brain/ FAIMS/MMTS |

| | | | | | | | |
|---|---|---|---|---|---|---|---|
| 93 | Q8C8V1 | ZXD family zinc finger C(Zxdc) | [K].FTTVYNLKAHMK.[G] | 1xOxidation [M11]; 1xHomocysteineIAA [K8] | [250-261] | 1642,81 | Brain/ No FAIMS/IAA |
| 94 | Q8C8V1 | ZXD family zinc finger C(Zxdc) | [K].FTTVYNLKAHMK.[G] | 1xOxidation [M11]; 1xHomocysteineIAA [K8] | [250-261] | 1642,81 | Brain/ FAIMS/IAA |
