## Supplemental Table 4 S for "High field asymmetric waveform ion mobility spectrometry improves *N*-homocysteinylation mapping in mouse liver and brain proteins"

*Table 4 S. List of in vitro N-Hcy-proteins identified in the study.*

| Master Protein Accessions | Positions in Master Proteins | Annotated Sequence | Modifications | Qvality q-value | Theo. MH+ [Da] | Organ/ Method/Alkylation |
|---|---|---|---|---|---|---|
| A2A432 | A2A432 [597-611] | [K].FINAMKEAFETFINK.[R] | 1xHomocysteineIAA [K6] | 0,00960393 | 1976,96087 | Liver/ FAIMS/IAA |
| A2ABU4 | A2ABU4 [998-1004] | [K].KLSHEIR.[N] | 1xHomocysteineIAA [K1] | 0,00434736 | 1056,56187 | Liver/ FAIMS/IAA |
| A2AEY4 | A2AEY4 [578-588] | [R].KPEIMAEKLDK.[V] | 1xHomocysteineMMTS [K8] | 0,0069474 | 1464,7259 | Liver/ FAIMS/MMTS |
| A2AJI0 | A2AJI0 [818-834] | [R].TAEGLLPFAEAEAFLKK.[A] | 1xHomocysteineIAA [K16] | 0,000318237 | 2009,04122 | Brain/ FAIMS/IAA |
| A2AS89 | A2AS89 [214-224] | [R].SVDEGLLDSKR.[V] | 1xHomocysteineMMTS [K10] | 0,00805184 | 1381,64501 | Liver/ FAIMS/MMTS |
| A2ASS6 | A2ASS6 [11979-11995] | [R].TVPETVPEKTRPMAPPK.[K] | 1xOxidation [M13]; 1xHomocysteineIAA [K9] | 0,00976205 | 2068,05656 | Brain/ No FAIMS/IAA |
| A6X935 | A6X935 [908-920] | [R].TVKVQGVDYLATR.[E] | 1xHomocysteineMMTS [K3] | 0,00107839 | 1612,81856 | Liver/ FAIMS/MMTS |
| B2RSH2 | B2RSH2 [55-70] | [K].IIHEAGYSEEECKQYK.[A] | 1xCarbamidomethyl [C12]; 1xHomocysteineIAA [K13] | 4,32935E-05 | 2157,95797 | Brain/ FAIMS/IAA |
| B2RXS4 | B2RXS4 [109-123] | [R].LVECGSLFKGICALR.[A] | 2xCarbamidomethyl [C4; C12]; 1xHomocysteineIAA [K9] | 0,00626664 | 1896,94926 | Brain/ FAIMS/IAA |
| B5X0G2 | B5X0G2 [58-78] | [R].LFLEQIHVLENSLVLKVHTVR.[D] | 1xHomocysteineIAA [K16] | 6,30912E-05 | 2661,49089 | Liver/ FAIMS/IAA |
| B5X0G2 | B5X0G2 [48-57] | [R].EKIEEHGNFR.[L] | 1xHomocysteineMMTS [K2] | 0,00413569 | 1421,63003 | Liver/ FAIMS/MMTS |
| B5X0G2; P11589; P11588 | B5X0G2 [33-47]; P11589 [33-47]; P11588 [33-47] | [K].INGEWHTIILASDKR.[E] | 1xHomocysteineIAA [K14] | 4,32935E-05 | 1926,98544 | Liver/ No FAIMS/IAA |
| B5X0G2; P11589; P11588 | B5X0G2 [33-47]; P11589 [33-47]; P11588 [33-47] | [K].INGEWHTIILASDKR.[E] | 1xHomocysteineIAA [K14] | 4,32935E-05 | 1926,98544 | Liver/ FAIMS/IAA |
| B5X0G2; P11589; P11588 | B5X0G2 [27-47]; P11589 [27-47]; P11588 [27-47] | [R].NFNVEKINGEWHTIILASDKR.[E] | 1xHomocysteineMMTS [K6] | 4,49774E-05 | 2647,31194 | Liver/ No FAIMS/MMTS |
| B5X0G2; P11589; P11588 | B5X0G2 [27-46]; P11589 [27-46]; P11588 [27-46] | [R].NFNVEKINGEWHTIILASDK.[R] | 1xHomocysteineMMTS [K6] | 4,49774E-05 | 2491,21083 | Liver/ No FAIMS/MMTS |
| B5X0G2; P11589; P11588 | B5X0G2 [79-94]; P11589 [79-94]; P11588 [79-94] | [R].DEECSELSMVADKTEK.[A] | 1xMethylthio [C4]; 1xHomocysteineMMTS [K13] | 0,000302483 | 2022,7833 | Liver/ No FAIMS/MMTS |
| B5X0G2; P11589; P11588 | B5X0G2 [141-151]; P11589 [141-151]; P11588 [141-151] | [R].EPDLSSDIKER.[F] | 1xHomocysteineMMTS [K9] | 4,49774E-05 | 1451,65049 | Liver/ No FAIMS/MMTS |
| B5X0G2; P11589; P11588 | B5X0G2 [92-112]; P11589 [92-112]; P11588 [92-112] | [K].TEKAGEYSVTYDGFNTFTIPK.[T] | 1xHomocysteineMMTS [K3] | 4,49774E-05 | 2531,14689 | Liver/ No FAIMS/MMTS |
| B5X0G2; P11589; P11588 | B5X0G2 [33-47]; P11589 [33-47]; P11588 [33-47] | [K].INGEWHTIILASDKR.[E] | 1xHomocysteineMMTS [K14] | 4,49774E-05 | 1915,9517 | Liver/ No FAIMS/MMTS |
| B5X0G2; P11589; P11588 | B5X0G2 [141-151]; P11589 [141-151]; P11588 [141-151] | [R].EPDLSSDIKER.[F] | 1xHomocysteineMMTS [K9] | 4,49774E-05 | 1451,65049 | Liver/ FAIMS/MMTS |
| B5X0G2; P11589; P11588 | B5X0G2 [27-47]; P11589 [27-47]; P11588 [27-47] | [R].NFNVEKINGEWHTIILASDKR.[E] | 1xHomocysteineMMTS [K6] | 4,49774E-05 | 2647,31194 | Liver/ FAIMS/MMTS |
| B5X0G2; P11589; P11588 | B5X0G2 [27-46]; P11589 [27-46]; P11588 [27-46] | [R].NFNVEKINGEWHTIILASDK.[R] | 1xHomocysteineMMTS [K6] | 4,49774E-05 | 2491,21083 | Liver/ FAIMS/MMTS |
| B5X0G2; P11589; P11588 | B5X0G2 [92-112]; P11589 [92-112]; P11588 [92-112] | [K].TEKAGEYSVTYDGFNTFTIPK.[T] | 1xHomocysteineMMTS [K3] | 4,49774E-05 | 2531,14689 | Liver/ FAIMS/MMTS |
| B5X0G2; P11589; P11588 | B5X0G2 [33-47]; P11589 [33-47]; P11588 [33-47] | [K].INGEWHTIILASDKR.[E] | 1xHomocysteineMMTS [K14] | 4,49774E-05 | 1915,9517 | Liver/ FAIMS/MMTS |
| B5X0G2; P11589; P11588 | B5X0G2 [113-140]; P11589 [113-140]; P11588 [113-140] | [K].TDYDNFLMAHLINEKDGETFQLMGLYGR.[E] | 2xOxidation [M8; M23]; 1xHomocysteineMMTS [K15] | 4,49774E-05 | 3486,54729 | Liver/ FAIMS/MMTS |
| B5X0G2; P11589; P11588 | B5X0G2 [113-140]; P11589 [113-140]; P11588 [113-140] | [K].TDYDNFLMAHLINEKDGETFQLMGLYGR.[E] | 1xOxidation [M8]; 1xHomocysteineMMTS [K15] | 4,49774E-05 | 3470,55238 | Liver/ FAIMS/MMTS |
| B5X0G2; P11589; P11588 | B5X0G2 [79-94]; P11589 [79-94]; P11588 [79-94] | [R].DEECSELSMVADKTEK.[A] | 1xMethylthio [C4]; 1xHomocysteineMMTS [K13] | 0,000302483 | 2022,7833 | Liver/ FAIMS/MMTS |
| B9EKR1 | B9EKR1 [61-79] | [K].QSPINIDEDLTQVNVNLKK.[L] | 1xHomocysteineIAA [K18] | 4,32935E-05 | 2342,20204 | Brain/ No FAIMS/IAA |
| B9EKR1 | B9EKR1 [61-79] | [K].QSPINIDEDLTQVNVNLKK.[L] | 1xHomocysteineIAA [K18] | 4,32935E-05 | 2342,20204 | Brain/ FAIMS/IAA |
| B9EKR1 | B9EKR1 [82-100] | [K].FQGWEKASLENTFIHNTGK.[T] | 1xHomocysteineMMTS [K6] | 0,00239411 | 2370,10055 | Brain/ FAIMS/MMTS |
| D3YVF0 | D3YVF0 [345-360] | [K].EGKLSQAEETTVAQAK.[E] | 1xHomocysteineIAA [K3] | 4,32935E-05 | 1863,91167 | Brain/ FAIMS/IAA |
| D3Z6Q9; O08539; Q7TQF7 | D3Z6Q9 [145-153]; O08539 [146-154]; Q7TQF7 [141-149] | [R].KLVDYDSAR.[H] | 1xHomocysteineIAA [K1] | 9,61721E-05 | 1240,59905 | Brain/ No FAIMS/IAA |
| D3Z6Q9; O08539; Q7TQF7 | D3Z6Q9 [145-153]; O08539 [146-154]; Q7TQF7 [141-149] | [R].KLVDYDSAR.[H] | 1xHomocysteineIAA [K1] | 9,61721E-05 | 1240,59905 | Brain/ FAIMS/IAA |
| D3Z7P3 | D3Z7P3 [164-174] | [K].FITALKSTGLR.[T] | 1xHomocysteineIAA [K6] | 4,32935E-05 | 1380,76678 | Brain/ FAIMS/IAA |
| E9PUL5 | E9PUL5 [108-118] | [R].QEPASKPDVNR.[E] | 1xHomocysteineIAA [K6] | 4,32935E-05 | 1414,67434 | Brain/ No FAIMS/IAA |
| E9PUL5 | E9PUL5 [108-118] | [R].QEPASKPDVNR.[E] | 1xHomocysteineIAA [K6] | 4,32935E-05 | 1414,67434 | Brain/ FAIMS/IAA |
| E9PUL5 | E9PUL5 [13-26] | [K].GVEDSSKTQTEGPR.[H] | 1xHomocysteineMMTS [K7] | 0,000266738 | 1653,7207 | Brain/ FAIMS/MMTS |
| E9PV24 | E9PV24 [411-422] | [K].TITKTVTGPDGR.[R] | 1xHomocysteineMMTS [K4] | 0,000879882 | 1408,6923 | Liver/ FAIMS/MMTS |

| | | | | | | |
|---|---|---|---|---|---|---|
| E9PV24 | E9PV24 [240-256] | [K].SQLQEAPPEWKALTEMR.[Q] | 1xHomocysteineMMTS [K11] | 4,49774E-05 | 2177,01879 | Liver/ FAIMS/MMTS |
| E9PVX6 | E9PVX6 [2691-2708] | [R].NTLKEPVGDSINVEEVKK.[S] | 1xHomocysteineMMTS [K17] | 0,0060387 | 2162,08317 | Liver/ No FAIMS/MMTS |
| E9Q5G3 | E9Q5G3 [551-561] | [K].VISGQKLEIER.[L] | 1xHomocysteineIAA [K6] | 0,00738589 | 1445,77807 | Liver/ FAIMS/IAA |
| E9Q5G3 | E9Q5G3 [551-561] | [K].VISGQKLEIER.[L] | 1xHomocysteineIAA [K6] | 0,00738589 | 1445,77807 | Brain/ FAIMS/IAA |
| E9Q912 | E9Q912 [309-320] | [K].LFEGGKGSVFQR.[V] | 1xHomocysteineIAA [K6] | 4,32935E-05 | 1498,74711 | Brain/ FAIMS/IAA |
| F8VPU2 | F8VPU2 [183-195] | [K].NKYVPQQDALEDR.[I] | 1xHomocysteineIAA [K2] | 0,00012645 | 1749,82246 | Brain/ FAIMS/IAA |
| G3X982 | G3X982 [890-900] | [K].LENAYKIPNLR.[V] | 1xHomocysteineMMTS [K6] | 0,000192903 | 1493,76032 | Liver/ FAIMS/MMTS |
| G3X982 | G3X982 [807-823] | [K].ASKPGLLASVAAVAAQK.[T] | 1xHomocysteineMMTS [K3] | 4,49774E-05 | 1744,94482 | Liver/ FAIMS/MMTS |
| G3X982 | G3X982 [419-429] | [R].SSKWEFVSAFR.[Q] | 1xHomocysteineMMTS [K3] | 0,00107839 | 1506,68682 | Liver/ FAIMS/MMTS |
| G3X982 | G3X982 [987-996] | [K].KAVDEFNQQR.[F] | 1xHomocysteineMMTS [K1] | 0,00680313 | 1397,63003 | Liver/ FAIMS/MMTS |
| G5E829 | G5E829 [63-80] | [K].LKTSPNEGLSGNPADLER.[R] | 1xHomocysteineIAA [K2] | 4,32935E-05 | 2072,00769 | Brain/ FAIMS/IAA |
| G5E829; Q9R0K7 | G5E829 [743-754]; Q9R0K7 [721-732] | [R].IRNEKGEIEQER.[I] | 1xHomocysteineIAA [K5] | 0,00012645 | 1674,82279 | Brain/ FAIMS/IAA |
| G5E8K5 | G5E8K5 [1315-1330] | [K].FVVFAKTNDPVESSLR.[C] | 1xHomocysteineIAA [K6] | 0,000258769 | 1983,00042 | Brain/ FAIMS/IAA |
| O08532 | O08532 [100-114] | [R].LAMEAEKVQAAHQWR.[E] | 1xHomocysteineIAA [K7] | 4,32935E-05 | 1941,9422 | Brain/ FAIMS/IAA |
| O08539 | O08539 [273-292] | [K].QHGSNTFTVKAQPSDNAPEK.[G] | 1xHomocysteineIAA [K10] | 4,32935E-05 | 2330,08298 | Brain/ No FAIMS/IAA |
| O08539 | O08539 [273-292] | [K].QHGSNTFTVKAQPSDNAPEK.[G] | 1xHomocysteineIAA [K10] | 4,32935E-05 | 2330,08298 | Brain/ FAIMS/IAA |
| O08539 | O08539 [1-13] | [-].MAEMGSKGVTAGK.[I] | 1xMet-loss+Acetyl [N-Term]; 1xHomocysteineMMTS [K7] | 4,49774E-05 | 1340,6007 | Brain/ FAIMS/MMTS |
| O08539 | O08539 [255-272] | [K].EMSKLNQNLNDVLVSLEK.[Q] | 1xHomocysteineMMTS [K4] | 0,0097599 | 2237,09744 | Brain/ FAIMS/MMTS |
| O08539 | O08539 [235-258] | [R].VGFYVNTFQSIAGLEENFHKEMSK.[L] | 1xHomocysteineMMTS [K20] | 0,000437433 | 2938,35723 | Brain/ FAIMS/MMTS |
| O08539 | O08539 [259-282] | [K].LNQNLNDVLVSLEKQHGSNTFTVK.[A] | 1xHomocysteineMMTS [K14] | 4,49774E-05 | 2861,42842 | Brain/ FAIMS/MMTS |
| O08553 | O08553 [270-293] | [K].KGTVVYGEPITASLGTDGSHYWSK.[N] | 1xHomocysteineIAA [K1] | 4,32935E-05 | 2727,30829 | Brain/ No FAIMS/IAA |
| O08553 | O08553 [270-293] | [K].KGTVVYGEPITASLGTDGSHYWSK.[N] | 1xHomocysteineIAA [K1] | 4,32935E-05 | 2727,30829 | Brain/ FAIMS/IAA |
| O08553 | O08553 [512-525] | [K].TVTPASSAKTSPAK.[Q] | 1xHomocysteineMMTS [K9] | 4,49774E-05 | 1508,74473 | Brain/ No FAIMS/MMTS |
| O08553 | O08553 [521-532] | [K].TSPAKQQAPPVR.[N] | 1xHomocysteineMMTS [K5] | 4,49774E-05 | 1442,72426 | Brain/ No FAIMS/MMTS |
| O08553 | O08553 [255-268] | [K].VMSKSAAEVIAQAR.[K] | 1xHomocysteineMMTS [K4] | 4,49774E-05 | 1623,80153 | Brain/ No FAIMS/MMTS |
| O08553 | O08553 [472-480] | [R].KPFPDFVYK.[R] | 1xHomocysteineMMTS [K1] | 0,00180518 | 1303,62136 | Brain/ No FAIMS/MMTS |
| O08553 | O08553 [512-525] | [K].TVTPASSAKTSPAK.[Q] | 1xHomocysteineMMTS [K9] | 4,49774E-05 | 1508,74473 | Brain/ FAIMS/MMTS |
| O08553 | O08553 [362-374] | [R].MSVIWDKAVVTGK.[M] | 1xHomocysteineMMTS [K7] | 4,49774E-05 | 1596,79465 | Brain/ FAIMS/MMTS |
| O08553 | O08553 [521-532] | [K].TSPAKQQAPPVR.[N] | 1xHomocysteineMMTS [K5] | 4,49774E-05 | 1442,72426 | Brain/ FAIMS/MMTS |
| O08553 | O08553 [255-268] | [K].VMSKSAAEVIAQAR.[K] | 1xHomocysteineMMTS [K4] | 4,49774E-05 | 1623,80153 | Brain/ FAIMS/MMTS |
| O08553 | O08553 [401-423] | [R].ISVGSDADLVIWDPDSVKTISAK.[T] | 1xHomocysteineMMTS [K18] | 4,49774E-05 | 2579,27315 | Brain/ FAIMS/MMTS |
| O08553 | O08553 [441-467] | [R].GSPLVVISQGKIVLEDGTLHVTEGSGR.[Y] | 1xHomocysteineMMTS [K11] | 0,000530332 | 2911,50159 | Brain/ FAIMS/MMTS |
| O08553; O35098 | O08553 [17-23]; O35098 [17-23] | [R].LLIKGGK.[I] | 1xHomocysteineIAA [K4] | 0,00052325 | 902,54918 | Brain/ FAIMS/IAA |
| O08585 | O08585 [223-235] | [R].SVLISLKQAPLVH.[-] | 1xHomocysteineIAA [K7] | 4,32935E-05 | 1578,90361 | Brain/ No FAIMS/IAA |
| O08585 | O08585 [223-235] | [R].SVLISLKQAPLVH.[-] | 1xHomocysteineIAA [K7] | 4,32935E-05 | 1578,90361 | Brain/ FAIMS/IAA |
| O08585 | O08585 [223-235] | [R].SVLISLKQAPLVH.[-] | 1xHomocysteineMMTS [K7] | 4,49774E-05 | 1567,86987 | Brain/ No FAIMS/MMTS |
| O08585 | O08585 [223-235] | [R].SVLISLKQAPLVH.[-] | 1xHomocysteineMMTS [K7] | 4,49774E-05 | 1567,86987 | Brain/ FAIMS/MMTS |
| O08585 | O08585 [138-148] | [K].AIKELEEWYAR.[Q] | 1xHomocysteineMMTS [K3] | 6,70238E-05 | 1570,73925 | Brain/ FAIMS/MMTS |
| O08599 | O08599 [83-100] | [K].SVHSLISDFKDPPTAKYR.[A] | 1xHomocysteineIAA [K16] | 4,32935E-05 | 2235,12266 | Brain/ FAIMS/IAA |
| O08599 | O08599 [316-325] | [R].MNTGEKTTMR.[D] | 1xHomocysteineMMTS [K6] | 0,000152766 | 1331,55746 | Brain/ FAIMS/MMTS |
| O08599 | O08599 [8-20] | [K].AVVGEKIMHDVIK.[K] | 1xHomocysteineMMTS [K6] | 0,00166208 | 1601,8212 | Brain/ FAIMS/MMTS |
| O08599 | O08599 [209-225] | [K].LDAYKADDPTMGEGPDK.[A] | 1xHomocysteineMMTS [K5] | 0,00074635 | 1985,82892 | Brain/ FAIMS/MMTS |
| O08599 | O08599 [293-305] | [R].HKHIAEVSQEVTR.[S] | 1xHomocysteineMMTS [K2] | 4,49774E-05 | 1696,82577 | Brain/ FAIMS/MMTS |
| O08599 | O08599 [214-227] | [K].ADDPTMGEGPDKAR.[S] | 1xHomocysteineMMTS [K12] | 4,49774E-05 | 1622,66074 | Brain/ FAIMS/MMTS |
| O08599 | O08599 [484-497] | [K].DIMEDTIEDKLDTK.[H] | 1xHomocysteineMMTS [K10] | 4,49774E-05 | 1828,80131 | Brain/ FAIMS/MMTS |
| O08599; Q64324 | O08599 [1-13]; Q64324 [1-13] | [-].MAPLGLKAVVGEK.[I] | 1xMet-loss [N-Term]; 1xHomocysteineIAA [K7] | 4,32935E-05 | 1355,77153 | Brain/ No FAIMS/IAA |
| O08599; Q64324 | O08599 [1-13]; Q64324 [1-13] | [-].MAPLGLKAVVGEK.[I] | 1xMet-loss [N-Term]; 1xHomocysteineIAA [K7] | 4,32935E-05 | 1355,77153 | Brain/ FAIMS/IAA |
| O08677 | O08677 [389-399] | [R].SVTVQETKEGR.[T] | 1xHomocysteineMMTS [K8] | 0,000829616 | 1396,65591 | Liver/ FAIMS/MMTS |
| O08677 | O08677 [59-75] | [R].VIEGTKTDGSPTFYSFK.[Y] | 1xHomocysteineMMTS [K6] | 4,49774E-05 | 2039,94528 | Liver/ FAIMS/MMTS |
| O08709 | O08709 [65-84] | [R].NVKLIALSIDSVEDHLAWSK.[D] | 1xHomocysteineIAA [K3] | 4,32935E-05 | 2412,25916 | Liver/ FAIMS/IAA |
| O08709 | O08709 [183-199] | [K].KGESVMVVPTLSEEEAK.[Q] | 1xHomocysteineIAA [K1] | 4,32935E-05 | 2006,9773 | Brain/ FAIMS/IAA |
| O08709 | O08709 [85-106] | [K].DINAYNGETPTEKLPFPIIDDK.[G] | 1xHomocysteineMMTS [K13] | 4,49774E-05 | 2653,25242 | Liver/ No FAIMS/MMTS |
| O08709 | O08709 [183-199] | [K].KGESVMVVPTLSEEEAK.[Q] | 1xOxidation [M6]; 1xHomocysteineMMTS [K1] | 4,49774E-05 | 2011,93848 | Liver/ No FAIMS/MMTS |
| O08709 | O08709 [98-108] | [K].LPFPIIDDKGR.[D] | 1xHomocysteineMMTS [K9] | 4,49774E-05 | 1433,72795 | Liver/ No FAIMS/MMTS |
| O08709 | O08709 [133-142] | [R].VVFIFGPDKK.[L] | 1xHomocysteineMMTS [K9] | 0,000781536 | 1312,67921 | Liver/ No FAIMS/MMTS |
| O08709 | O08709 [205-215] | [K].GVFTKELPSGK.[K] | 1xHomocysteineMMTS [K5] | 4,49774E-05 | 1325,6592 | Liver/ No FAIMS/MMTS |
| O08709 | O08709 [65-84] | [R].NVKLIALSIDSVEDHLAWSK.[D] | 1xHomocysteineMMTS [K3] | 4,49774E-05 | 2401,22541 | Liver/ No FAIMS/MMTS |
| O08709 | O08709 [54-63] | [R].AAKLAPEFAK.[R] | 1xHomocysteineMMTS [K3] | 0,000342214 | 1208,61661 | Liver/ No FAIMS/MMTS |
| O08709 | O08709 [163-183] | [R].VVDSLQLTGTKPVATPVDWKK.[G] | 1xHomocysteineMMTS [K20] | 4,49774E-05 | 2445,28801 | Liver/ No FAIMS/MMTS |
| O08709 | O08709 [143-155] | [K].LKLSILYPATTGR.[N] | 1xHomocysteineMMTS [K2] | 0,000437433 | 1595,86478 | Liver/ No FAIMS/MMTS |
| O08709 | O08709 [163-182] | [R].VVDSLQLTGTKPVATPVDWK.[K] | 1xHomocysteineMMTS [K11] | 4,49774E-05 | 2317,19305 | Liver/ No FAIMS/MMTS |
| O08709 | O08709 [183-199] | [K].KGESVMVVPTLSEEEAK.[Q] | 1xHomocysteineMMTS [K1] | 4,49774E-05 | 1995,94356 | Liver/ No FAIMS/MMTS |
| O08709 | O08709 [183-199] | [K].KGESVMVVPTLSEEEAK.[Q] | 1xOxidation [M6]; 1xHomocysteineMMTS [K1] | 4,49774E-05 | 2011,93848 | Liver/ FAIMS/MMTS |
| O08709 | O08709 [98-108] | [K].LPFPIIDDKGR.[D] | 1xHomocysteineMMTS [K9] | 4,49774E-05 | 1433,72795 | Liver/ FAIMS/MMTS |
| O08709 | O08709 [133-142] | [R].VVFIFGPDKK.[L] | 1xHomocysteineMMTS [K9] | 0,000781536 | 1312,67921 | Liver/ FAIMS/MMTS |
| O08709 | O08709 [205-215] | [K].GVFTKELPSGK.[K] | 1xHomocysteineMMTS [K5] | 4,49774E-05 | 1325,6592 | Liver/ FAIMS/MMTS |
| O08709 | O08709 [143-155] | [K].LKLSILYPATTGR.[N] | 1xHomocysteineMMTS [K2] | 0,000437433 | 1595,86478 | Liver/ FAIMS/MMTS |
| O08709 | O08709 [163-182] | [R].VVDSLQLTGTKPVATPVDWK.[K] | 1xHomocysteineMMTS [K11] | 4,49774E-05 | 2317,19305 | Liver/ FAIMS/MMTS |
| O08709 | O08709 [183-199] | [K].KGESVMVVPTLSEEEAK.[Q] | 1xHomocysteineMMTS [K1] | 4,49774E-05 | 1995,94356 | Liver/ FAIMS/MMTS |
| O08709 | O08709 [54-63] | [R].AAKLAPEFAK.[R] | 1xHomocysteineMMTS [K3] | 0,000342214 | 1208,61661 | Liver/ FAIMS/MMTS |
| O08709 | O08709 [163-183] | [R].VVDSLQLTGTKPVATPVDWKK.[G] | 1xHomocysteineMMTS [K20] | 4,49774E-05 | 2445,28801 | Liver/ FAIMS/MMTS |
| O08709 | O08709 [85-106] | [K].DINAYNGETPTEKLPFPIIDDK.[G] | 1xHomocysteineMMTS [K13] | 4,49774E-05 | 2653,25242 | Liver/ FAIMS/MMTS |
| O08709 | O08709 [205-215] | [K].GVFTKELPSGK.[K] | 1xHomocysteineMMTS [K5] | 4,49774E-05 | 1325,6592 | Brain/ No FAIMS/MMTS |

| | | | | | | |
|---|---|---|---|---|---|---|
| O08709 | O08709 [65-84] | [R].NVKLIALSIDSVEDHLAWSK.[D] | 1xHomocysteineMMTS [K3] | 4,49774E-05 | 2401,22541 | Brain/ FAIMS/MMTS |
| O08709 | O08709 [163-183] | [R].VVDSLQLTGTKPVATPVDWKK.[G] | 1xHomocysteineMMTS [K20] | 4,49774E-05 | 2445,28801 | Brain/ FAIMS/MMTS |
| O08709 | O08709 [163-182] | [R].VVDSLQLTGTKPVATPVDWK.[K] | 1xHomocysteineMMTS [K11] | 4,49774E-05 | 2317,19305 | Brain/ FAIMS/MMTS |
| O08749 | O08749 [316-336] | [R].RPFTQNLGLEELGIELDPKGR.[I] | 1xHomocysteineIAA [K19] | 4,32935E-05 | 2556,32388 | Brain/ FAIMS/IAA |
| O08749 | O08749 [160-177] | [K].NQVTATKADGSTQVIDTK.[N] | 1xHomocysteineMMTS [K7] | 8,93346E-05 | 2039,97362 | Liver/ FAIMS/MMTS |
| O08749 | O08749 [160-177] | [K].NQVTATKADGSTQVIDTK.[N] | 1xHomocysteineMMTS [K7] | 8,93346E-05 | 2039,97362 | Brain/ FAIMS/MMTS |
| O08756 | O08756 [106-116] | [K].NKIHTLEDFQR.[V] | 1xHomocysteineIAA [K2] | 0,00444768 | 1574,77439 | Liver/ FAIMS/IAA |
| O08756 | O08756 [53-69] | [K].KLGESCIFAPANVTSEK.[E] | 1xCarbamidomethyl [C6]; 1xHomocysteineIAA [K1] | 9,61721E-05 | 2024,97797 | Brain/ FAIMS/IAA |
| O08756 | O08756 [106-116] | [K].NKIHTLEDFQR.[V] | 1xHomocysteineMMTS [K2] | 0,00576292 | 1563,74064 | Liver/ FAIMS/MMTS |
| O08795 | O08795 [159-167] | [K].LLELQAGKK.[S] | 1xHomocysteineIAA [K8] | 9,61721E-05 | 1173,666 | Brain/ FAIMS/IAA |
| O08795 | O08795 [178-188] | [R].AAKEEAERPEK.[E] | 1xHomocysteineMMTS [K3] | 0,00107839 | 1420,65591 | Liver/ No FAIMS/MMTS |
| O08917 | O08917 [153-168] | [K].DIHDDQDYLHSLGKAR.[T] | 1xHomocysteineIAA [K14] | 0,000162511 | 2056,95051 | Brain/ FAIMS/IAA |
| O09061 | O09061 [222-231] | [R].ICIVTKEGIR.[E] | 1xCarbamidomethyl [C2]; 1xHomocysteineIAA [K6] | 0,000889294 | 1362,7232 | Brain/ FAIMS/IAA |
| O09111 | O09111 [34-44] | [R].AVIAPSGVEKK.[R] | 1xHomocysteineIAA [K10] | 0,00012645 | 1272,69803 | Liver/ FAIMS/IAA |
| O09111 | O09111 [34-44] | [R].AVIAPSGVEKK.[R] | 1xHomocysteineIAA [K10] | 0,00012645 | 1272,69803 | Brain/ FAIMS/IAA |
| O09173 | O09173 [239-252] | [R].VPGGYTVINKFQGK.[L] | 1xHomocysteineMMTS [K10] | 0,00074635 | 1670,83929 | Liver/ FAIMS/MMTS |
| O35136 | O35136 [257-269] | [K].YILKGSNTELTVR.[N] | 1xHomocysteineIAA [K4] | 0,00012645 | 1667,87852 | Brain/ FAIMS/IAA |
| O35143 | O35143 [72-87] | [R].KHHEDEIDHHSKEIER.[L] | 1xHomocysteineIAA [K12] | 4,32935E-05 | 2213,01524 | Brain/ No FAIMS/IAA |
| O35143 | O35143 [72-87] | [R].KHHEDEIDHHSKEIER.[L] | 1xHomocysteineIAA [K12] | 4,32935E-05 | 2213,01524 | Brain/ FAIMS/IAA |
| O35143 | O35143 [63-71] | [K].TKEQLAALR.[K] | 1xHomocysteineMMTS [K2] | 0,00542903 | 1192,61767 | Brain/ FAIMS/MMTS |
| O35215 | O35215 [23-37] | [R].LCAATATILDKPEDR.[V] | 1xCarbamidomethyl [C2]; 1xHomocysteineIAA [K11] | 4,32935E-05 | 1847,89899 | Liver/ No FAIMS/IAA |
| O35215 | O35215 [23-37] | [R].LCAATATILDKPEDR.[V] | 1xCarbamidomethyl [C2]; 1xHomocysteineIAA [K11] | 4,32935E-05 | 1847,89899 | Liver/ FAIMS/IAA |
| O35215 | O35215 [76-95] | [R].THSASFFKFLTEELSLDQDR.[I] | 1xHomocysteineMMTS [K8] | 4,49774E-05 | 2534,16902 | Liver/ No FAIMS/MMTS |
| O35215 | O35215 [100-111] | [R].FFPLEAWQIGKK.[G] | 1xHomocysteineMMTS [K11] | 0,000152766 | 1626,8171 | Liver/ No FAIMS/MMTS |
| O35215 | O35215 [23-37] | [R].LCAATATILDKPEDR.[V] | 1xMethylthio [C2]; 1xHomocysteineMMTS [K11] | 4,49774E-05 | 1825,83151 | Liver/ FAIMS/MMTS |
| O35215 | O35215 [76-95] | [R].THSASFFKFLTEELSLDQDR.[I] | 1xHomocysteineMMTS [K8] | 4,49774E-05 | 2534,16902 | Liver/ FAIMS/MMTS |
| O35215 | O35215 [100-111] | [R].FFPLEAWQIGKK.[G] | 1xHomocysteineMMTS [K11] | 0,000152766 | 1626,8171 | Liver/ FAIMS/MMTS |
| O35226 | O35226 [26-42] | [R].LQAQQDAVNIVCHSKTR.[S] | 1xCarbamidomethyl [C12]; 1xHomocysteineIAA [K15] | 4,32935E-05 | 2142,05427 | Brain/ FAIMS/IAA |
| O35295 | O35295 [70-82] | [R].FLKIAEVGAGGSK.[S] | 1xHomocysteineIAA [K3] | 4,32935E-05 | 1450,77226 | Brain/ FAIMS/IAA |
| O35295; P42669 | O35295 [58-67]; P42669 [72-81] | [R].FYLDVKQNAK.[G] | 1xHomocysteineIAA [K6] | 4,32935E-05 | 1399,70385 | Brain/ FAIMS/IAA |
| O35381 | O35381 [15-28] | [R].TPSDVKELVLDNCK.[S] | 1xCarbamidomethyl [C13]; 1xHomocysteineIAA [K6] | 4,32935E-05 | 1791,86155 | Liver/ FAIMS/IAA |
| O35381 | O35381 [76-91] | [R].ISGDLEVLAEKCPNLK.[H] | 1xCarbamidomethyl [C12]; 1xHomocysteineIAA [K11] | 4,32935E-05 | 1959,98781 | Brain/ No FAIMS/IAA |
| O35381 | O35381 [76-91] | [R].ISGDLEVLAEKCPNLK.[H] | 1xCarbamidomethyl [C12]; 1xHomocysteineIAA [K11] | 4,32935E-05 | 1959,98781 | Brain/ FAIMS/IAA |
| O35405 | O35405 [303-317] | [R].TPDLKALLNVVDSAR.[S] | 1xHomocysteineIAA [K5] | 4,32935E-05 | 1785,95274 | Brain/ No FAIMS/IAA |
| O35405 | O35405 [303-317] | [R].TPDLKALLNVVDSAR.[S] | 1xHomocysteineIAA [K5] | 4,32935E-05 | 1785,95274 | Brain/ FAIMS/IAA |
| O35459 | O35459 [267-282] | [K].SPVAVQGSKINLIYSR.[D] | 1xHomocysteineMMTS [K9] | 4,49774E-05 | 1894,98775 | Liver/ FAIMS/MMTS |
| O35459 | O35459 [211-229] | [R].LPKVIGNQSLVNELTFSAR.[K] | 1xHomocysteineMMTS [K3] | 4,49774E-05 | 2249,17807 | Liver/ FAIMS/MMTS |
| O35490 | O35490 [36-57] | [K].RGYVKAGPWTPEAAVEHPEAVR.[Q] | 1xHomocysteineIAA [K5] | 4,32935E-05 | 2594,29325 | Liver/ FAIMS/IAA |
| O35490 | O35490 [378-395] | [K].GAAELMQQKEATTEQQLR.[E] | 1xOxidation [M6]; 1xHomocysteineMMTS [K9] | 4,49774E-05 | 2211,02025 | Liver/ No FAIMS/MMTS |
| O35490 | O35490 [66-86] | [R].AGSNVMQTFTFYASEDKLENR.[G] | 1xOxidation [M6]; 1xHomocysteineMMTS [K17] | 4,49774E-05 | 2587,12617 | Liver/ No FAIMS/MMTS |
| O35490 | O35490 [328-344] | [K].HGSWGSGLDMHTKPWIR.[A] | 1xOxidation [M10]; 1xHomocysteineMMTS [K13] | 4,49774E-05 | 2143,96228 | Liver/ No FAIMS/MMTS |
| O35490 | O35490 [94-108] | [K].ISGQKVNEAACDIAR.[Q] | 1xMethylthio [C11]; 1xHomocysteineMMTS [K5] | 4,49774E-05 | 1783,79579 | Liver/ No FAIMS/MMTS |
| O35490 | O35490 [378-395] | [K].GAAELMQQKEATTEQQLR.[E] | 1xHomocysteineMMTS [K9] | 4,49774E-05 | 2195,02533 | Liver/ No FAIMS/MMTS |
| O35490 | O35490 [143-151] | [R].QQLEVFMKK.[N] | 1xHomocysteineMMTS [K8] | 0,0074518 | 1313,64145 | Liver/ No FAIMS/MMTS |
| O35490 | O35490 [87-98] | [R].GNYVAEKISGQK.[V] | 1xHomocysteineMMTS [K7] | 4,49774E-05 | 1456,6923 | Liver/ No FAIMS/MMTS |
| O35490 | O35490 [279-286] | [R].WDIQKYAR.[E] | 1xHomocysteineMMTS [K5] | 0,000507453 | 1242,57581 | Liver/ No FAIMS/MMTS |
| O35490 | O35490 [37-57] | [R].GYVKAGPWTPEAAVEHPEAVR.[Q] | 1xHomocysteineMMTS [K4] | 4,49774E-05 | 2427,1584 | Liver/ No FAIMS/MMTS |
| O35490 | O35490 [230-239] | [K].LMKEGLEAAR.[L] | 1xHomocysteineMMTS [K3] | 4,49774E-05 | 1280,61596 | Liver/ No FAIMS/MMTS |
| O35490 | O35490 [17-36] | [R].LNAGEVVIGDGGFVFALEKR.[G] | 1xHomocysteineMMTS [K19] | 4,49774E-05 | 2254,13587 | Liver/ No FAIMS/MMTS |
| O35490 | O35490 [66-86] | [R].AGSNVMQTFTFYASEDKLENR.[G] | 1xHomocysteineMMTS [K17] | 4,49774E-05 | 2571,13125 | Liver/ No FAIMS/MMTS |
| O35490 | O35490 [328-344] | [K].HGSWGSGLDMHTKPWIR.[A] | 1xHomocysteineMMTS [K13] | 4,49774E-05 | 2127,96737 | Liver/ No FAIMS/MMTS |
| O35490 | O35490 [349-356] | [R].KEYWQNLR.[I] | 1xHomocysteineMMTS [K1] | 0,00206087 | 1299,59727 | Liver/ No FAIMS/MMTS |
| O35490 | O35490 [378-395] | [K].GAAELMQQKEATTEQQLR.[E] | 1xOxidation [M6]; 1xHomocysteineMMTS [K9] | 4,49774E-05 | 2211,02025 | Liver/ FAIMS/MMTS |
| O35490 | O35490 [66-86] | [R].AGSNVMQTFTFYASEDKLENR.[G] | 1xOxidation [M6]; 1xHomocysteineMMTS [K17] | 4,49774E-05 | 2587,12617 | Liver/ FAIMS/MMTS |
| O35490 | O35490 [230-239] | [K].LMKEGLEAAR.[L] | 1xOxidation [M2]; 1xHomocysteineMMTS [K3] | 0,000879882 | 1296,61087 | Liver/ FAIMS/MMTS |
| O35490 | O35490 [94-108] | [K].ISGQKVNEAACDIAR.[Q] | 1xMethylthio [C11]; 1xHomocysteineMMTS [K5] | 4,49774E-05 | 1783,79579 | Liver/ FAIMS/MMTS |
| O35490 | O35490 [378-395] | [K].GAAELMQQKEATTEQQLR.[E] | 1xHomocysteineMMTS [K9] | 4,49774E-05 | 2195,02533 | Liver/ FAIMS/MMTS |
| O35490 | O35490 [143-151] | [R].QQLEVFMKK.[N] | 1xHomocysteineMMTS [K8] | 0,0074518 | 1313,64145 | Liver/ FAIMS/MMTS |
| O35490 | O35490 [87-98] | [R].GNYVAEKISGQK.[V] | 1xHomocysteineMMTS [K7] | 4,49774E-05 | 1456,6923 | Liver/ FAIMS/MMTS |
| O35490 | O35490 [279-286] | [R].WDIQKYAR.[E] | 1xHomocysteineMMTS [K5] | 0,000507453 | 1242,57581 | Liver/ FAIMS/MMTS |
| O35490 | O35490 [37-57] | [R].GYVKAGPWTPEAAVEHPEAVR.[Q] | 1xHomocysteineMMTS [K4] | 4,49774E-05 | 2427,1584 | Liver/ FAIMS/MMTS |
| O35490 | O35490 [230-239] | [K].LMKEGLEAAR.[L] | 1xHomocysteineMMTS [K3] | 4,49774E-05 | 1280,61596 | Liver/ FAIMS/MMTS |
| O35490 | O35490 [17-36] | [R].LNAGEVVIGDGGFVFALEKR.[G] | 1xHomocysteineMMTS [K19] | 4,49774E-05 | 2254,13587 | Liver/ FAIMS/MMTS |
| O35490 | O35490 [66-86] | [R].AGSNVMQTFTFYASEDKLENR.[G] | 1xHomocysteineMMTS [K17] | 4,49774E-05 | 2571,13125 | Liver/ FAIMS/MMTS |

| | | | | | | |
|---|---|---|---|---|---|---|
| O35490 | O35490 [328-344] | [K].HGSWGSGLDMHTKPWIR.[A] | 1xHomocysteineMMTS [K13] | 4,49774E-05 | 2127,96737 | Liver/ FAIMS/MMTS |
| O35490 | O35490 [349-356] | [R].KEYWQNLR.[I] | 1xHomocysteineMMTS [K1] | 0,00206087 | 1299,59727 | Liver/ FAIMS/MMTS |
| O35490 | O35490 [319-344] | [R].GFLPPASEKHGSWGSGLDMHTKPWIR.[A] | 1xOxidation [M19]; 1xHomocysteineMMTS [K9] | 0,00356631 | 3070,44845 | Liver/ FAIMS/MMTS |
| O35490 | O35490 [328-344] | [K].HGSWGSGLDMHTKPWIR.[A] | 1xOxidation [M10]; 1xHomocysteineMMTS [K13] | 4,49774E-05 | 2143,96228 | Liver/ FAIMS/MMTS |
| O35490 | O35490 [319-344] | [R].GFLPPASEKHGSWGSGLDMHTKPWIR.[A] | 1xHomocysteineMMTS [K9] | 0,000211896 | 3054,45353 | Liver/ FAIMS/MMTS |
| O35490 | O35490 [36-57] | [K].RGYVKAGPWTPEAAVEHPEAVR.[Q] | 1xHomocysteineMMTS [K5] | 4,49774E-05 | 2583,25951 | Liver/ FAIMS/MMTS |
| O35490; Q91WS4 | O35490 [279-286]; Q91WS4 [270-277] | [R].WDIQKYAR.[E] | 1xHomocysteineIAA [K5] | 0,000194827 | 1253,60955 | Liver/ No FAIMS/IAA |
| O35490; Q91WS4 | O35490 [279-286]; Q91WS4 [270-277] | [R].WDIQKYAR.[E] | 1xHomocysteineIAA [K5] | 0,000194827 | 1253,60955 | Liver/ FAIMS/IAA |
| O35526 | O35526 [57-70] | [R].KHSAILASPNPDEK.[T] | 1xHomocysteineIAA [K1] | 4,32935E-05 | 1680,83738 | Brain/ No FAIMS/IAA |
| O35526 | O35526 [57-70] | [R].KHSAILASPNPDEK.[T] | 1xHomocysteineIAA [K1] | 4,32935E-05 | 1680,83738 | Brain/ FAIMS/IAA |
| O35526 | O35526 [47-56] | [K].IAENVEEVKR.[K] | 1xHomocysteineMMTS [K9] | 0,000599431 | 1349,65518 | Brain/ FAIMS/MMTS |
| O35526 | O35526 [93-108] | [K].LKSIEQSIEQEEGLNR.[S] | 1xHomocysteineMMTS [K2] | 4,49774E-05 | 2035,9787 | Brain/ FAIMS/MMTS |
| O35526 | O35526 [71-84] | [K].TKEELEELMSDIKK.[T] | 1xHomocysteineMMTS [K13] | 0,00221411 | 1855,88498 | Brain/ FAIMS/MMTS |
| O35526 | O35526 [57-70] | [R].KHSAILASPNPDEK.[T] | 1xHomocysteineMMTS [K1] | 0,000266738 | 1669,80364 | Brain/ FAIMS/MMTS |
| O35526 | O35526 [126-142] | [K].KFVEVMSEYNATQSDYR.[E] | 1xHomocysteineMMTS [K1] | 4,49774E-05 | 2229,96134 | Brain/ FAIMS/MMTS |
| O35633 | O35633 [9-23] | [K].LTNVATSVSNKSQAK.[V] | 1xHomocysteineIAA [K11] | 4,32935E-05 | 1721,88506 | Brain/ FAIMS/IAA |
| O35639 | O35639 [258-274] | [R].LHQALKGAGTDEFTLNR.[I] | 1xHomocysteineIAA [K6] | 0,00107087 | 2045,02328 | Brain/ FAIMS/IAA |
| O35658 | O35658 [78-91] | [K].AFVEFLTDEIKEEK.[K] | 1xHomocysteineIAA [K11] | 4,32935E-05 | 1871,90954 | Liver/ FAIMS/IAA |
| O35658 | O35658 [78-92] | [K].AFVEFLTDEIKEEKK.[I] | 1xHomocysteineIAA [K] | 4,32935E-05 | 2000,0045 | Brain/ FAIMS/IAA |
| O35685 | O35685 [210-228] | [R].VGLKGQPPVVDGELYNEVK.[V] | 1xHomocysteineIAA [K4] | 4,32935E-05 | 2215,14273 | Brain/ FAIMS/IAA |
| O35737 | O35737 [88-114] | [K].SNNVEMDWVLKHTGPNSPDTANDGFVR.[L] | 1xHomocysteineIAA [K11] | 0,000698725 | 3174,43676 | Brain/ FAIMS/IAA |
| O35943 | O35943 [133-149] | [K].LGGDLGTYVINKQTPNK.[Q] | 1xHomocysteineMMTS [K12] | 0,00175939 | 1980,98814 | Liver/ FAIMS/MMTS |
| O35945; P24549 | O35945 [396-410]; P24549 [396-410] | [R].IAKEEIFGPVQQIMK.[F] | 1xHomocysteineIAA [K3] | 0,00012645 | 1904,99725 | Liver/ FAIMS/IAA |
| O35945; P24549 | O35945 [88-98]; P24549 [88-98] | [R].LLNKLADLMER.[D] | 1xHomocysteineMMTS [K4] | 4,49774E-05 | 1478,75279 | Liver/ No FAIMS/MMTS |
| O35945; P24549 | O35945 [253-260]; P24549 [253-260] | [K].LIKEAAGK.[S] | 1xHomocysteineMMTS [K3] | 0,000706375 | 992,52673 | Liver/ No FAIMS/MMTS |
| O35945; P24549 | O35945 [88-98]; P24549 [88-98] | [R].LLNKLADLMER.[D] | 1xHomocysteineMMTS [K4] | 4,49774E-05 | 1478,75279 | Liver/ FAIMS/MMTS |
| O35945; P24549 | O35945 [396-410]; P24549 [396-410] | [R].IAKEEIFGPVQQIMK.[F] | 1xHomocysteineMMTS [K3] | 4,49774E-05 | 1893,96351 | Liver/ FAIMS/MMTS |
| O35945; P24549 | O35945 [88-98]; P24549 [88-98] | [R].LLNKLADLMER.[D] | 1xOxidation [M9]; 1xHomocysteineMMTS [K4] | 0,00107839 | 1494,7477 | Liver/ FAIMS/MMTS |
| O35945; P24549 | O35945 [253-260]; P24549 [253-260] | [K].LIKEAAGK.[S] | 1xHomocysteineMMTS [K3] | 0,000706375 | 992,52673 | Liver/ FAIMS/MMTS |
| O54901 | O54901 [51-66] | [R].CSLKTSQEPLIVTWQK.[K] | 1xCarbamidomethyl [C1]; 1xHomocysteineIAA [K4] | 6,30912E-05 | 2092,05656 | Brain/ FAIMS/IAA |
| O54983 | O54983 [171-185] | [R].ENAEKFASTVQGDVR.[V] | 1xHomocysteineIAA [K5] | 0,0014138 | 1824,85449 | Brain/ FAIMS/IAA |
| O54991 | O54991 [881-889] | [R].AEINVKQAR.[L] | 1xHomocysteineIAA [K6] | 0,000177335 | 1202,63102 | Brain/ No FAIMS/IAA |
| O54991 | O54991 [881-889] | [R].AEINVKQAR.[L] | 1xHomocysteineIAA [K6] | 0,000177335 | 1202,63102 | Brain/ FAIMS/IAA |
| O55013 | O55013 [150-164] | [K].FVQDTLKGDGVTEIR.[M] | 1xHomocysteineIAA [K7] | 9,61721E-05 | 1851,92692 | Brain/ FAIMS/IAA |
| O55022 | O55022 [105-119] | [R].KFYGPEGPYGVFAGR.[D] | 1xHomocysteineIAA [K1] | 4,32935E-05 | 1818,8632 | Liver/ No FAIMS/IAA |
| O55022 | O55022 [105-119] | [R].KFYGPEGPYGVFAGR.[D] | 1xHomocysteineIAA [K1] | 4,32935E-05 | 1818,8632 | Liver/ FAIMS/IAA |
| O55022 | O55022 [105-119] | [R].KFYGPEGPYGVFAGR.[D] | 1xHomocysteineIAA [K1] | 4,32935E-05 | 1818,8632 | Brain/ No FAIMS/IAA |
| O55022 | O55022 [105-119] | [R].KFYGPEGPYGVFAGR.[D] | 1xHomocysteineIAA [K1] | 4,32935E-05 | 1818,8632 | Brain/ FAIMS/IAA |
| O55022 | O55022 [97-104] | [K].VFDVTKGR.[K] | 1xHomocysteineMMTS [K6] | 0,00363897 | 1084,5278 | Liver/ No FAIMS/MMTS |
| O55022 | O55022 [105-119] | [R].KFYGPEGPYGVFAGR.[D] | 1xHomocysteineMMTS [K1] | 4,49774E-05 | 1807,82946 | Liver/ No FAIMS/MMTS |
| O55022 | O55022 [89-102] | [R].ILMAINGKVFDVTK.[G] | 1xOxidation [M3]; 1xHomocysteineMMTS [K8] | 4,49774E-05 | 1727,88928 | Liver/ FAIMS/MMTS |
| O55022 | O55022 [164-172] | [K].YHHVGKLLK.[E] | 1xHomocysteineMMTS [K6] | 0,000653201 | 1257,65948 | Liver/ FAIMS/MMTS |
| O55022 | O55022 [97-104] | [K].VFDVTKGR.[K] | 1xHomocysteineMMTS [K6] | 0,00363897 | 1084,5278 | Liver/ FAIMS/MMTS |
| O55022 | O55022 [105-119] | [R].KFYGPEGPYGVFAGR.[D] | 1xHomocysteineMMTS [K1] | 4,49774E-05 | 1807,82946 | Liver/ FAIMS/MMTS |
| O55022 | O55022 [89-102] | [R].ILMAINGKVFDVTK.[G] | 1xHomocysteineMMTS [K8] | 0,00347466 | 1711,89437 | Liver/ FAIMS/MMTS |
| O55022 | O55022 [105-119] | [R].KFYGPEGPYGVFAGR.[D] | 1xHomocysteineMMTS [K1] | 4,49774E-05 | 1807,82946 | Brain/ FAIMS/MMTS |
| O55042 | O55042 [1-10] | [-].MDVFMKGLSK.[A] | 1xAcetyl [N-Term]; 1xHomocysteineIAA [K6] | 4,32935E-05 | 1371,64693 | Brain/ No FAIMS/IAA |
| O55042 | O55042 [1-10] | [-].MDVFMKGLSK.[A] | 1xAcetyl [N-Term]; 1xHomocysteineIAA [K6] | 4,32935E-05 | 1371,64693 | Brain/ FAIMS/IAA |
| O55042 | O55042 [44-58] | [K].TKEGVVHGVTTVAEK.[T] | 1xHomocysteineMMTS [K2] | 4,49774E-05 | 1717,86115 | Brain/ FAIMS/MMTS |
| O55042 | O55042 [59-80] | [K].TKEQVTNVGGAVVTGVTAVAQK.[T] | 1xHomocysteineMMTS [K2] | 4,49774E-05 | 2320,19993 | Brain/ FAIMS/MMTS |
| O55042 | O55042 [11-21] | [K].AKEGVVAAAEK.[T] | 1xHomocysteineMMTS [K2] | 0,000250158 | 1235,61225 | Brain/ FAIMS/MMTS |
| O55091 | O55091 [286-302] | [R].NILVEKNFTNTPDESTK.[N] | 1xHomocysteineIAA [K6] | 0,000273428 | 2124,02776 | Brain/ FAIMS/IAA |
| O55100 | O55100 [1-22] | [-].MEGGAYGAGKAGGAFDPYTLVR.[Q] | 1xAcetyl [N-Term]; 1xHomocysteineIAA [K10] | 4,32935E-05 | 2404,10603 | Brain/ No FAIMS/IAA |
| O55100 | O55100 [1-22] | [-].MEGGAYGAGKAGGAFDPYTLVR.[Q] | 1xAcetyl [N-Term]; 1xHomocysteineIAA [K10] | 4,32935E-05 | 2404,10603 | Brain/ FAIMS/IAA |
| O55126 | O55126 [139-149] | [K].LKENQEFVNFR.[K] | 1xHomocysteineIAA [K2] | 0,00097531 | 1597,77914 | Brain/ FAIMS/IAA |
| O55131 | O55131 [351-363] | [K].KMEMEMEQVFEMK.[V] | 1xHomocysteineIAA [K1] | 4,32935E-05 | 1863,78178 | Brain/ FAIMS/IAA |
| O55131 | O55131 [291-309] | [R].THMQDLKDVTNNVHYENYR.[S] | 1xHomocysteineMMTS [K7] | 0,00230389 | 2540,11152 | Liver/ FAIMS/MMTS |
| O55142 | O55142 [5-15] | [R].LWCKAIFAGYK.[R] | 1xCarbamidomethyl [C3]; 1xHomocysteineIAA [K4] | 9,61721E-05 | 1530,75959 | Brain/ FAIMS/IAA |
| O70161 | O70161 [435-445] | [R].FFKFMSSTVFR.[K] | 1xHomocysteineIAA [K3] | 0,00290455 | 1570,7545 | Brain/ FAIMS/IAA |
| O70251 | O70251 [1-22] | [-].MGFGDLKTPAGLQVLNDYLADK.[S] | 1xMet-loss [N-Term]; 1xHomocysteineIAA [K7] | 4,32935E-05 | 2409,21187 | Liver/ FAIMS/IAA |

| | | | | | | |
|---|---|---|---|---|---|---|
| O70251 | O70251 [1-22] | [-].MGFGDLKTPAGLQVLNDYLADK.[S] | 1xMet-loss [N-Term]; 1xHomocysteineIAA [K7] | 4,32935E-05 | 2409,21187 | Brain/ FAIMS/IAA |
| O70251 | O70251 [1-22] | [-].MGFGDLKTPAGLQVLNDYLADK.[S] | 1xMet-loss [N-Term]; 1xHomocysteineMMTS [K7] | 4,49774E-05 | 2398,17813 | Liver/ FAIMS/MMTS |
| O70318; Q9Z2H5 | O70318 [438-444]; Q9Z2H5 [324-330] | [K].ILKISYK.[R] | 1xHomocysteineIAA [K3] | 0,00174544 | 1038,60161 | Brain/ No FAIMS/IAA |
| O70318; Q9Z2H5 | O70318 [438-444]; Q9Z2H5 [324-330] | [K].ILKISYK.[R] | 1xHomocysteineIAA [K3] | 0,00174544 | 1038,60161 | Brain/ FAIMS/IAA |
| O70400 | O70400 [240-254] | [K].APVTKVAASVGNAQK.[L] | 1xHomocysteineMMTS [K5] | 0,00824761 | 1603,82946 | Liver/ FAIMS/MMTS |
| O70400 | O70400 [18-32] | [R].LVGGKDFEQPLAISR.[V] | 1xHomocysteineMMTS [K5] | 0,000929608 | 1792,90844 | Liver/ FAIMS/MMTS |
| O70435 | O70435 [175-183] | [K].TEIEKLQMK.[E] | 1xHomocysteineIAA [K5] | 0,00746057 | 1293,65412 | Brain/ FAIMS/IAA |
| O70439 | O70439 [93-109] | [R].LVAEFTTSLTNFQKAQR.[Q] | 1xHomocysteineIAA [K14] | 4,32935E-05 | 2128,08555 | Brain/ FAIMS/IAA |
| O70443 | O70443 [68-83] | [K].EYKPLIIYNAIDSLTR.[I] | 1xHomocysteineIAA [K3] | 6,30912E-05 | 2083,08924 | Brain/ FAIMS/IAA |
| O70493 | O70493 [72-84] | [R].YSDFEWLKNELER.[D] | 1xHomocysteineIAA [K8] | 0,00416819 | 1902,86907 | Brain/ FAIMS/IAA |
| O70591 | O70591 [126-145] | [R].LMGEDEKPAAKENSEGAGAK.[A] | 1xHomocysteineIAA [K11] | 4,32935E-05 | 2206,01146 | Brain/ FAIMS/IAA |
| O70591 | O70591 [126-145] | [R].LMGEDEKPAAKENSEGAGAK.[A] | 1xHomocysteineMMTS [K11] | 0,0021186 | 2194,97771 | Brain/ FAIMS/MMTS |
| O88342 | O88342 [471-489] | [R].VYSILASTLKDEGKLLEAK.[G] | 1xHomocysteineIAA [K14] | 4,32935E-05 | 2252,22065 | Brain/ FAIMS/IAA |
| O88343 | O88343 [753-766] | [K].LIVPSEFKPTSPNR.[G] | 1xHomocysteineIAA [K8] | 4,32935E-05 | 1758,92072 | Brain/ FAIMS/IAA |
| O88448 | O88448 [357-368] | [R].LGPDDPNVAKTK.[N] | 1xHomocysteineIAA [K10] | 0,00347063 | 1428,71514 | Brain/ FAIMS/IAA |
| O88485 | O88485 [569-584] | [R].WAQGGKEVAVGDSEGR.[I] | 1xHomocysteineIAA [K6] | 0,000988467 | 1819,83917 | Brain/ FAIMS/IAA |
| O88531 | O88531 [180-196] | [R].LVQAQYWHDPIKESVYR.[N] | 1xHomocysteineIAA [K12] | 4,32935E-05 | 2306,13865 | Brain/ FAIMS/IAA |
| O88569 | O88569 [22-38] | [R].KLFIGGLSFETTEESLR.[N] | 1xHomocysteineIAA [K1] | 4,32935E-05 | 2101,06342 | Liver/ FAIMS/IAA |
| O88569 | O88569 [22-38] | [R].KLFIGGLSFETTEESLR.[N] | 1xHomocysteineIAA [K1] | 4,32935E-05 | 2101,06342 | Brain/ No FAIMS/IAA |
| O88569 | O88569 [154-173] | [R].GFGFVTFDDHDPVDKIVLQK.[Y] | 1xHomocysteineMMTS [K15] | 6,70238E-05 | 2440,16756 | Liver/ No FAIMS/MMTS |
| O88569 | O88569 [100-112] | [R].EESGKPGAHVTVK.[K] | 1xHomocysteineMMTS [K5] | 0,00669057 | 1501,71376 | Liver/ FAIMS/MMTS |
| O88569 | O88569 [154-173] | [R].GFGFVTFDDHDPVDKIVLQK.[Y] | 1xHomocysteineMMTS [K15] | 6,70238E-05 | 2440,16756 | Liver/ FAIMS/MMTS |
| O88569 | O88569 [154-173] | [R].GFGFVTFDDHDPVDKIVLQK.[Y] | 1xHomocysteineMMTS [K15] | 6,70238E-05 | 2440,16756 | Brain/ FAIMS/MMTS |
| O88587 | O88587 [205-227] | [R].KGTVLLADNVIVPGTPDFLAYVR.[G] | 1xHomocysteineIAA [K1] | 4,32935E-05 | 2632,41672 | Liver/ FAIMS/IAA |
| O88587 | O88587 [155-171] | [K].VSILIGASQDLIPQLKK.[K] | 1xHomocysteineMMTS [K16] | 0,00714378 | 1986,11262 | Liver/ FAIMS/MMTS |
| O88643 | O88643 [248-258] | [K].MSDEEILEKLR.[S] | 1xHomocysteineIAA [K9] | 4,32935E-05 | 1536,73964 | Brain/ No FAIMS/IAA |
| O88643 | O88643 [248-258] | [K].MSDEEILEKLR.[S] | 1xHomocysteineIAA [K9] | 4,32935E-05 | 1536,73964 | Brain/ FAIMS/IAA |
| O88643 | O88643 [248-258] | [K].MSDEEILEKLR.[S] | 1xHomocysteineMMTS [K9] | 0,00394153 | 1525,7059 | Brain/ FAIMS/MMTS |
| O88643; Q61036 | O88643 [309-320]; Q61036 [322-333] | [K].KELIINEILVMR.[E] | 1xHomocysteineIAA [K1] | 0,000508188 | 1644,91754 | Brain/ No FAIMS/IAA |
| O88685 | O88685 [148-174] | [K].LKPGDLVGVNKDSYLILETLPTEYDSR.[V] | 1xHomocysteineIAA [K11] | 9,61721E-05 | 3209,63986 | Brain/ FAIMS/IAA |
| O88696 | O88696 [208-222] | [K].HTKQSLQVIESAMER.[D] | 1xHomocysteineIAA [K3] | 4,32935E-05 | 1930,94734 | Brain/ FAIMS/IAA |
| O88737 | O88737 [88-103] | [R].TTSPTPKQASATAPGR.[E] | 1xHomocysteineIAA [K7] | 0,000243434 | 1744,86466 | Brain/ FAIMS/IAA |
| O88844 | O88844 [375-388] | [K].DLAACIKGLPNVQR.[S] | 1xCarbamidomethyl [C5]; 1xHomocysteineIAA [K7] | 0,000741565 | 1728,88837 | Liver/ FAIMS/IAA |
| O88844 | O88844 [375-388] | [K].DLAACIKGLPNVQR.[S] | 1xCarbamidomethyl [C5]; 1xHomocysteineIAA [K7] | 0,000741565 | 1728,88837 | Brain/ FAIMS/IAA |
| O88844 | O88844 [21-29] | [R].IIWELIKEK.[L] | 1xHomocysteineMMTS [K7] | 0,00413569 | 1334,72108 | Liver/ No FAIMS/MMTS |
| O88844 | O88844 [141-159] | [R].ATDFVVPGPGKVEITYTPK.[D] | 1xHomocysteineMMTS [K11] | 4,49774E-05 | 2182,09227 | Liver/ No FAIMS/MMTS |
| O88844 | O88844 [141-159] | [R].ATDFVVPGPGKVEITYTPK.[D] | 1xHomocysteineMMTS [K11] | 4,49774E-05 | 2182,09227 | Liver/ FAIMS/MMTS |
| O88844 | O88844 [152-164] | [K].VEITYTPKDGTQK.[V] | 1xHomocysteineMMTS [K8] | 0,0004709 | 1642,7815 | Liver/ FAIMS/MMTS |
| O88844 | O88844 [21-29] | [R].IIWELIKEK.[L] | 1xHomocysteineMMTS [K7] | 0,00413569 | 1334,72108 | Liver/ FAIMS/MMTS |
| O88844 | O88844 [4-20] | [R].KIQGGSVVEMQGDEMTR.[I] | 1xHomocysteineMMTS [K1] | 4,49774E-05 | 2027,90171 | Liver/ FAIMS/MMTS |
| O88844 | O88844 [21-29] | [R].IIWELIKEK.[L] | 1xHomocysteineMMTS [K7] | 0,00413569 | 1334,72108 | Brain/ FAIMS/MMTS |
| O88935 | O88935 [238-256] | [K].KLGTEEFPLIDQTFYPNHK.[E] | 1xHomocysteineIAA [K1] | 4,32935E-05 | 2451,20131 | Brain/ FAIMS/IAA |
| O88935 | O88935 [588-602] | [R].QGPPQKPPGPAGPTR.[Q] | 1xHomocysteineMMTS [K6] | 0,00109439 | 1647,80939 | Brain/ FAIMS/MMTS |
| O88935 | O88935 [115-131] | [R].VLLVIDEPHTDWAKYFK.[G] | 1xHomocysteineMMTS [K14] | 0,000738471 | 2237,11334 | Brain/ FAIMS/MMTS |
| O88952 | O88952 [17-26] | [R].AIELLEKLQR.[S] | 1xHomocysteineIAA [K7] | 4,32935E-05 | 1386,77735 | Brain/ FAIMS/IAA |
| O88998 | O88998 [125-135] | [K].MENQMKGLETK.[F] | 1xHomocysteineIAA [K6] | 0,000429391 | 1482,67493 | Brain/ FAIMS/IAA |
| O89023 | O89023 [245-262] | [R].LFGGSFTHQASVAKVVGK.[Q] | 1xHomocysteineIAA [K14] | 4,32935E-05 | 2007,04804 | Brain/ FAIMS/IAA |
| O89023 | O89023 [339-349] | [R].VNTEFMKAAAR.[G] | 1xOxidation [M6]; 1xHomocysteineMMTS [K7] | 0,00239411 | 1416,64324 | Liver/ FAIMS/MMTS |
| O89023 | O89023 [339-349] | [R].VNTEFMKAAAR.[G] | 1xHomocysteineMMTS [K7] | 0,000530332 | 1400,64832 | Liver/ FAIMS/MMTS |
| O89053 | O89053 [13-29] | [R].HVFGQPAKADQCYEDVR.[V] | 1xCarbamidomethyl [C12]; 1xHomocysteineIAA [K8] | 0,000289025 | 2193,98043 | Brain/ FAIMS/IAA |
| O89112 | O89112 [240-253] | [K].LHSLVKPSVDFVCR.[L] | 1xCarbamidomethyl [C13]; 1xHomocysteineIAA [K6] | 0,00596151 | 1830,93532 | Brain/ FAIMS/IAA |
| P00329 | P00329 [314-324] | [R].TWKGAIFGGFK.[S] | 1xHomocysteineIAA [K3] | 0,000210768 | 1385,70345 | Liver/ FAIMS/IAA |
| P00329 | P00329 [1-9] | [-].MSTAGKVIK.[C] | 1xMet-loss+Acetyl [N-Term]; 1xHomocysteineMMTS [K6] | 0,00288111 | 1008,52165 | Liver/ FAIMS/MMTS |
| P00329 | P00329 [314-324] | [R].TWKGAIFGGFK.[S] | 1xHomocysteineMMTS [K3] | 0,000530332 | 1374,66971 | Liver/ FAIMS/MMTS |
| P00405 | P00405 [152-178] | [R].MLISSEDVLHSWAVPSLGLKTDAIPGR.[L] | 1xHomocysteineIAA [K20] | 4,32935E-05 | 3066,57509 | Brain/ No FAIMS/IAA |
| P00493 | P00493 [157-166] | [K].MVKVASLLVK.[R] | 1xHomocysteineIAA [K3] | 8,35381E-05 | 1261,73706 | Brain/ FAIMS/IAA |
| P00920 | P00920 [28-57] | [R].QSPVDIDTATAQHDPALQPLLISYDKAASK.[S] | 1xHomocysteineIAA [K26] | 4,32935E-05 | 3367,68385 | Brain/ No FAIMS/IAA |
| P00920 | P00920 [28-57] | [R].QSPVDIDTATAQHDPALQPLLISYDKAASK.[S] | 1xHomocysteineIAA [K26] | 4,32935E-05 | 3367,68385 | Brain/ FAIMS/IAA |
| P00920 | P00920 [159-169] | [K].VLEALHSIKTK.[G] | 1xHomocysteineMMTS [K9] | 4,49774E-05 | 1401,75925 | Liver/ No FAIMS/MMTS |
| P00920 | P00920 [159-169] | [K].VLEALHSIKTK.[G] | 1xHomocysteineMMTS [K9] | 4,49774E-05 | 1401,75925 | Liver/ FAIMS/MMTS |
| P00920 | P00920 [159-169] | [K].VLEALHSIKTK.[G] | 1xHomocysteineMMTS [K9] | 4,49774E-05 | 1401,75925 | Brain/ No FAIMS/MMTS |
| P00920 | P00920 [159-169] | [K].VLEALHSIKTK.[G] | 1xHomocysteineMMTS [K9] | 4,49774E-05 | 1401,75925 | Brain/ FAIMS/MMTS |
| P00920 | P00920 [28-57] | [R].QSPVDIDTATAQHDPALQPLLISYDKAASK.[S] | 1xHomocysteineMMTS [K26] | 0,000173718 | 3356,6501 | Brain/ FAIMS/MMTS |
| P00920 | P00920 [227-253] | [R].TLNFNEEGDAEEAMVDNWRPAQPLKNR.[K] | 1xHomocysteineMMTS [K25] | 4,49774E-05 | 3307,49289 | Brain/ FAIMS/MMTS |
| P01831 | P01831 [61-76] | [R].KHVLSGTLGIPEHTYR.[S] | 1xHomocysteineIAA [K1] | 4,32935E-05 | 1982,02764 | Brain/ No FAIMS/IAA |

| | | | | | | |
|---|---|---|---|---|---|---|
| P01831 | P01831 [61-76] | [R].KHVLSGTLGIPEHTYR.[S] | 1xHomocysteineIAA [K1] | 4,32935E-05 | 1982,02764 | Brain/ FAIMS/IAA |
| P01831 | P01831 [61-76] | [R].KHVLSGTLGIPEHTYR.[S] | 1xHomocysteineMMTS [K1] | 0,000547722 | 1970,9939 | Brain/ No FAIMS/MMTS |
| P01831 | P01831 [61-76] | [R].KHVLSGTLGIPEHTYR.[S] | 1xHomocysteineMMTS [K1] | 0,000547722 | 1970,9939 | Brain/ FAIMS/MMTS |
| P01837 | P01837 [82-100] | [R].HNSYTCEATHKTSTSPIVK.[S] | 1xCarbamidomethyl [C6]; 1xHomocysteineIAA [K11] | 4,32935E-05 | 2335,08054 | Liver/ No FAIMS/IAA |
| P01837 | P01837 [82-100] | [R].HNSYTCEATHKTSTSPIVK.[S] | 1xCarbamidomethyl [C6]; 1xHomocysteineIAA [K11] | 4,32935E-05 | 2335,08054 | Liver/ FAIMS/IAA |
| P01837 | P01837 [93-104] | [K].TSTSPIVKSFNR.[N] | 1xHomocysteineMMTS [K8] | 4,49774E-05 | 1499,73449 | Liver/ No FAIMS/MMTS |
| P01837 | P01837 [41-48] | [K].WKIDGSER.[Q] | 1xHomocysteineMMTS [K2] | 0,0042423 | 1153,51288 | Liver/ No FAIMS/MMTS |
| P01837 | P01837 [93-104] | [K].TSTSPIVKSFNR.[N] | 1xHomocysteineMMTS [K8] | 4,49774E-05 | 1499,73449 | Liver/ FAIMS/MMTS |
| P01837 | P01837 [41-48] | [K].WKIDGSER.[Q] | 1xHomocysteineMMTS [K2] | 0,0042423 | 1153,51288 | Liver/ FAIMS/MMTS |
| P01872 | P01872 [152-170] | [K].DGKLVESGFTTDPVTIENK.[G] | 1xHomocysteineMMTS [K3] | 4,49774E-05 | 2213,04645 | Liver/ FAIMS/MMTS |
| P01878 | P01878 [245-254] | [R].AFNPKEVLVR.[W] | 1xHomocysteineMMTS [K5] | 0,000530332 | 1335,69117 | Liver/ No FAIMS/MMTS |
| P01878 | P01878 [245-254] | [R].AFNPKEVLVR.[W] | 1xHomocysteineMMTS [K5] | 0,000530332 | 1335,69117 | Liver/ FAIMS/MMTS |
| P01878 | P01878 [145-164] | [R].NPEGAVFTWEPSTGKDAVQK.[K] | 1xHomocysteineMMTS [K15] | 8,93346E-05 | 2324,06858 | Liver/ FAIMS/MMTS |
| P01887 | P01887 [66-78] | [K].IPKVEMSDMSFSK.[D] | 1xHomocysteineMMTS [K3] | 0,000530332 | 1661,74057 | Liver/ FAIMS/MMTS |
| P01898 | P01898 [195-205] | [R].YLELGKETLLR.[T] | 1xHomocysteineMMTS [K6] | 0,000530332 | 1497,78038 | Liver/ FAIMS/MMTS |
| P01942 | P01942 [9-17] | [K].SNIKAAWGK.[I] | 1xHomocysteineIAA [K4] | 4,32935E-05 | 1148,58809 | Liver/ No FAIMS/IAA |
| P01942 | P01942 [9-17] | [K].SNIKAAWGK.[I] | 1xHomocysteineIAA [K4] | 4,32935E-05 | 1148,58809 | Liver/ FAIMS/IAA |
| P01942 | P01942 [9-17] | [K].SNIKAAWGK.[I] | 1xHomocysteineIAA [K4] | 4,32935E-05 | 1148,58809 | Brain/ No FAIMS/IAA |
| P01942 | P01942 [9-17] | [K].SNIKAAWGK.[I] | 1xHomocysteineIAA [K4] | 4,32935E-05 | 1148,58809 | Brain/ FAIMS/IAA |
| P01942 | P01942 [33-57] | [R].MFASFPTTKTYFPHFDVSHGSAQVK.[G] | 1xOxidation [M1]; 1xHomocysteineMMTS [K9] | 4,49774E-05 | 3009,37322 | Liver/ No FAIMS/MMTS |
| P01942 | P01942 [1-17] | [-].MVLSGEDKSNIKAAWGK.[I] | 1xMet-loss [N-Term]; 2xHomocysteineMMTS [K8; K12] | 4,49774E-05 | 2028,93737 | Liver/ No FAIMS/MMTS |
| P01942 | P01942 [1-12] | [-].MVLSGEDKSNIK.[A] | 1xMet-loss [N-Term]; 1xHomocysteineMMTS [K8] | 4,49774E-05 | 1352,65485 | Liver/ No FAIMS/MMTS |
| P01942 | P01942 [1-17] | [-].MVLSGEDKSNIKAAWGK.[I] | 1xMet-loss [N-Term]; 1xHomocysteineMMTS [K12] | 4,49774E-05 | 1865,92481 | Liver/ No FAIMS/MMTS |
| P01942 | P01942 [33-57] | [R].MFASFPTTKTYFPHFDVSHGSAQVK.[G] | 1xHomocysteineMMTS [K9] | 4,49774E-05 | 2993,3783 | Liver/ No FAIMS/MMTS |
| P01942 | P01942 [13-32] | [K].AAWGKIGGHGAEYGAEALER.[M] | 1xHomocysteineMMTS [K5] | 4,49774E-05 | 2206,01682 | Liver/ No FAIMS/MMTS |
| P01942 | P01942 [9-17] | [K].SNIKAAWGK.[I] | 1xHomocysteineMMTS [K4] | 8,93346E-05 | 1137,55435 | Liver/ No FAIMS/MMTS |
| P01942 | P01942 [42-61] | [K].TYFPHFDVSHGSAQVKGHGK.[K] | 1xHomocysteineMMTS [K16] | 4,49774E-05 | 2362,08557 | Liver/ No FAIMS/MMTS |
| P01942 | P01942 [129-142] | [K].FLASVSTVLTSKYR.[-] | 1xHomocysteineMMTS [K12] | 4,49774E-05 | 1734,89172 | Liver/ No FAIMS/MMTS |
| P01942 | P01942 [33-57] | [R].MFASFPTTKTYFPHFDVSHGSAQVK.[G] | 1xOxidation [M1]; 1xHomocysteineMMTS [K9] | 4,49774E-05 | 3009,37322 | Liver/ FAIMS/MMTS |
| P01942 | P01942 [1-17] | [-].MVLSGEDKSNIKAAWGK.[I] | 1xMet-loss [N-Term]; 2xHomocysteineMMTS [K8; K12] | 4,49774E-05 | 2028,93737 | Liver/ FAIMS/MMTS |
| P01942 | P01942 [1-12] | [-].MVLSGEDKSNIK.[A] | 1xMet-loss [N-Term]; 1xHomocysteineMMTS [K8] | 4,49774E-05 | 1352,65485 | Liver/ FAIMS/MMTS |
| P01942 | P01942 [1-17] | [-].MVLSGEDKSNIKAAWGK.[I] | 1xMet-loss [N-Term]; 1xHomocysteineMMTS [K12] | 4,49774E-05 | 1865,92481 | Liver/ FAIMS/MMTS |
| P01942 | P01942 [33-57] | [R].MFASFPTTKTYFPHFDVSHGSAQVK.[G] | 1xHomocysteineMMTS [K9] | 4,49774E-05 | 2993,3783 | Liver/ FAIMS/MMTS |
| P01942 | P01942 [13-32] | [K].AAWGKIGGHGAEYGAEALER.[M] | 1xHomocysteineMMTS [K5] | 4,49774E-05 | 2206,01682 | Liver/ FAIMS/MMTS |
| P01942 | P01942 [9-17] | [K].SNIKAAWGK.[I] | 1xHomocysteineMMTS [K4] | 8,93346E-05 | 1137,55435 | Liver/ FAIMS/MMTS |
| P01942 | P01942 [42-61] | [K].TYFPHFDVSHGSAQVKGHGK.[K] | 1xHomocysteineMMTS [K16] | 4,49774E-05 | 2362,08557 | Liver/ FAIMS/MMTS |
| P01942 | P01942 [129-142] | [K].FLASVSTVLTSKYR.[-] | 1xHomocysteineMMTS [K12] | 4,49774E-05 | 1734,89172 | Liver/ FAIMS/MMTS |
| P01942 | P01942 [1-12] | [-].MVLSGEDKSNIK.[A] | 1xMet-loss+Acetyl [N-Term]; 1xHomocysteineMMTS [K8] | 4,49774E-05 | 1394,66541 | Liver/ FAIMS/MMTS |
| P01942 | P01942 [1-12] | [-].MVLSGEDKSNIK.[A] | 1xMet-loss [N-Term]; 1xHomocysteineMMTS [K8] | 4,49774E-05 | 1352,65485 | Brain/ No FAIMS/MMTS |
| P01942 | P01942 [1-17] | [-].MVLSGEDKSNIKAAWGK.[I] | 1xMet-loss [N-Term]; 1xHomocysteineMMTS [K12] | 4,49774E-05 | 1865,92481 | Brain/ No FAIMS/MMTS |
| P01942 | P01942 [33-57] | [R].MFASFPTTKTYFPHFDVSHGSAQVK.[G] | 1xHomocysteineMMTS [K9] | 4,49774E-05 | 2993,3783 | Brain/ No FAIMS/MMTS |
| P01942 | P01942 [13-32] | [K].AAWGKIGGHGAEYGAEALER.[M] | 1xHomocysteineMMTS [K5] | 4,49774E-05 | 2206,01682 | Brain/ No FAIMS/MMTS |
| P01942 | P01942 [9-17] | [K].SNIKAAWGK.[I] | 1xHomocysteineMMTS [K4] | 8,93346E-05 | 1137,55435 | Brain/ No FAIMS/MMTS |
| P01942 | P01942 [42-61] | [K].TYFPHFDVSHGSAQVKGHGK.[K] | 1xHomocysteineMMTS [K16] | 4,49774E-05 | 2362,08557 | Brain/ No FAIMS/MMTS |
| P01942 | P01942 [33-57] | [R].MFASFPTTKTYFPHFDVSHGSAQVK.[G] | 1xOxidation [M1]; 1xHomocysteineMMTS [K9] | 4,49774E-05 | 3009,37322 | Brain/ FAIMS/MMTS |
| P01942 | P01942 [1-12] | [-].MVLSGEDKSNIK.[A] | 1xMet-loss [N-Term]; 1xHomocysteineMMTS [K8] | 4,49774E-05 | 1352,65485 | Brain/ FAIMS/MMTS |
| P01942 | P01942 [1-17] | [-].MVLSGEDKSNIKAAWGK.[I] | 1xMet-loss [N-Term]; 1xHomocysteineMMTS [K12] | 4,49774E-05 | 1865,92481 | Brain/ FAIMS/MMTS |
| P01942 | P01942 [33-57] | [R].MFASFPTTKTYFPHFDVSHGSAQVK.[G] | 1xHomocysteineMMTS [K9] | 4,49774E-05 | 2993,3783 | Brain/ FAIMS/MMTS |
| P01942 | P01942 [13-32] | [K].AAWGKIGGHGAEYGAEALER.[M] | 1xHomocysteineMMTS [K5] | 4,49774E-05 | 2206,01682 | Brain/ FAIMS/MMTS |
| P01942 | P01942 [9-17] | [K].SNIKAAWGK.[I] | 1xHomocysteineMMTS [K4] | 8,93346E-05 | 1137,55435 | Brain/ FAIMS/MMTS |
| P01942 | P01942 [42-61] | [K].TYFPHFDVSHGSAQVKGHGK.[K] | 1xHomocysteineMMTS [K16] | 4,49774E-05 | 2362,08557 | Brain/ FAIMS/MMTS |
| P02088 | P02088 [67-96] | [K].KVITAFNDGLNHLDSLKGTFASLSELHCDK.[L] | 1xCarbamidomethyl [C28]; 1xHomocysteineIAA [K17] | 4,32935E-05 | 3504,725 | Liver/ FAIMS/IAA |
| P02088 | P02088 [67-96] | [K].KVITAFNDGLNHLDSLKGTFASLSELHCDK.[L] | 1xCarbamidomethyl [C28]; 1xHomocysteineIAA [K17] | 4,32935E-05 | 3504,725 | Brain/ No FAIMS/IAA |
| P02088 | P02088 [67-96] | [K].KVITAFNDGLNHLDSLKGTFASLSELHCDK.[L] | 1xCarbamidomethyl [C28]; 1xHomocysteineIAA [K17] | 4,32935E-05 | 3504,725 | Brain/ FAIMS/IAA |
| P02088 | P02088 [67-83] | [K].KVITAFNDGLNHLDSLK.[G] | 1xHomocysteineMMTS [K1] | 4,49774E-05 | 2048,03034 | Liver/ No FAIMS/MMTS |
| P02088 | P02088 [68-96] | [K].VITAFNDGLNHLDSLKGTFASLSELHCDK.[L] | 1xMethylthio [C27]; 1xHomocysteineMMTS [K16] | 0,00242157 | 3354,56255 | Liver/ No FAIMS/MMTS |

| | | | | | | |
|---|---|---|---|---|---|---|
| P02088 | P02088 [106-133] | [R].LLGNMIVIVLGHHLGKDFTPAAQAAFQK.[V] | 1xOxidation [M5]; 1xHomocysteineMMTS [K16] | 4,49774E-05 | 3168,6519 | Liver/ No FAIMS/MMTS |
| P02088 | P02088 [42-62] | [R].YFDSFGDLSSASAIMGNAKVK.[A] | 1xOxidation [M15]; 1xHomocysteineMMTS [K19] | 4,49774E-05 | 2387,07162 | Liver/ No FAIMS/MMTS |
| P02088 | P02088 [67-96] | [K].KVITAFNDGLNHLDSLKGTFASLSELHCDK.[L] | 1xMethylthio [C28]; 1xHomocysteineMMTS [K17] | 4,49774E-05 | 3482,65751 | Liver/ No FAIMS/MMTS |
| P02088 | P02088 [42-62] | [R].YFDSFGDLSSASAIMGNAKVK.[A] | 1xHomocysteineMMTS [K19] | 4,49774E-05 | 2371,0767 | Liver/ No FAIMS/MMTS |
| P02088 | P02088 [63-83] | [K].AHGKKVITAFNDGLNHLDSLK.[G] | 2xHomocysteineMMTS [K4; K5] | 4,49774E-05 | 2604,25535 | Liver/ FAIMS/MMTS |
| P02088 | P02088 [42-62] | [R].YFDSFGDLSSASAIMGNAKVK.[A] | 2xHomocysteineMMTS [K19; K21] | 4,49774E-05 | 2534,08926 | Liver/ FAIMS/MMTS |
| P02088 | P02088 [106-133] | [R].LLGNMIVIVLGHHLGKDFTPAAQAAFQK.[V] | 1xOxidation [M5]; 1xHomocysteineMMTS [K16] | 4,49774E-05 | 3168,6519 | Liver/ FAIMS/MMTS |
| P02088 | P02088 [42-62] | [R].YFDSFGDLSSASAIMGNAKVK.[A] | 1xOxidation [M15]; 2xHomocysteineMMTS [K19; K21] | 4,49774E-05 | 2550,08417 | Liver/ FAIMS/MMTS |
| P02088 | P02088 [42-62] | [R].YFDSFGDLSSASAIMGNAKVK.[A] | 1xOxidation [M15]; 1xHomocysteineMMTS [K19] | 4,49774E-05 | 2387,07162 | Liver/ FAIMS/MMTS |
| P02088 | P02088 [42-62] | [R].YFDSFGDLSSASAIMGNAKVK.[A] | 1xHomocysteineMMTS [K19] | 4,49774E-05 | 2371,0767 | Liver/ FAIMS/MMTS |
| P02088 | P02088 [67-83] | [K].KVITAFNDGLNHLDSLK.[G] | 1xHomocysteineMMTS [K1] | 4,49774E-05 | 2048,03034 | Liver/ FAIMS/MMTS |
| P02088 | P02088 [68-96] | [K].VITAFNDGLNHLDSLKGTFASLSELHCDK.[L] | 1xMethylthio [C27]; 1xHomocysteineMMTS [K16] | 0,00242157 | 3354,56255 | Liver/ FAIMS/MMTS |
| P02088 | P02088 [67-83] | [K].KVITAFNDGLNHLDSLK.[G] | 1xHomocysteineMMTS [K1] | 4,49774E-05 | 2048,03034 | Brain/ No FAIMS/MMTS |
| P02088 | P02088 [42-62] | [R].YFDSFGDLSSASAIMGNAKVK.[A] | 1xHomocysteineMMTS [K19] | 4,49774E-05 | 2371,0767 | Brain/ No FAIMS/MMTS |
| P02088 | P02088 [67-83] | [K].KVITAFNDGLNHLDSLK.[G] | 1xHomocysteineMMTS [K1] | 4,49774E-05 | 2048,03034 | Brain/ FAIMS/MMTS |
| P02088 | P02088 [42-62] | [R].YFDSFGDLSSASAIMGNAKVK.[A] | 1xOxidation [M15]; 1xHomocysteineMMTS [K19] | 4,49774E-05 | 2387,07162 | Brain/ FAIMS/MMTS |
| P02088 | P02088 [42-62] | [R].YFDSFGDLSSASAIMGNAKVK.[A] | 1xHomocysteineMMTS [K19] | 4,49774E-05 | 2371,0767 | Brain/ FAIMS/MMTS |
| P02088; P02089 | P02088 [84-105]; P02089 [84-105] | [K].GTFASLSELHCDKLHVDPENFR.[L] | 1xCarbamidomethyl [C11]; 1xHomocysteineIAA [K13] | 4,32935E-05 | 2746,27119 | Liver/ No FAIMS/IAA |
| P02088; P02089 | P02088 [84-105]; P02089 [84-105] | [K].GTFASLSELHCDKLHVDPENFR.[L] | 1xCarbamidomethyl [C11]; 1xHomocysteineIAA [K13] | 4,32935E-05 | 2746,27119 | Liver/ FAIMS/IAA |
| P02088; P02089 | P02088 [84-105]; P02089 [84-105] | [K].GTFASLSELHCDKLHVDPENFR.[L] | 1xCarbamidomethyl [C11]; 1xHomocysteineIAA [K13] | 4,32935E-05 | 2746,27119 | Brain/ No FAIMS/IAA |
| P02088; P02089 | P02088 [84-105]; P02089 [84-105] | [K].GTFASLSELHCDKLHVDPENFR.[L] | 1xCarbamidomethyl [C11]; 1xHomocysteineIAA [K13] | 4,32935E-05 | 2746,27119 | Brain/ FAIMS/IAA |
| P02088; P02089 | P02088 [84-105]; P02089 [84-105] | [K].GTFASLSELHCDKLHVDPENFR.[L] | 1xMethylthio [C11]; 1xHomocysteineMMTS [K13] | 4,49774E-05 | 2724,20371 | Liver/ No FAIMS/MMTS |
| P02088; P02089 | P02088 [84-105]; P02089 [84-105] | [K].GTFASLSELHCDKLHVDPENFR.[L] | 1xMethylthio [C11]; 1xHomocysteineMMTS [K13] | 4,49774E-05 | 2724,20371 | Liver/ FAIMS/MMTS |
| P02089 | P02089 [42-62] | [R].YFDSFGDLSSASAIMGNPKVK.[A] | 1xHomocysteineMMTS [K19] | 4,49774E-05 | 2397,09235 | Liver/ FAIMS/MMTS |
| P02468 | P02468 [1326-1339] | [K].GKAEQQTADQLLAR.[A] | 1xHomocysteineIAA [K2] | 4,32935E-05 | 1702,85409 | Brain/ FAIMS/IAA |
| P03930 | P03930 [49-57] | [K].TPWELKWTK.[I] | 1xHomocysteineIAA [K6] | 9,61721E-05 | 1362,68747 | Brain/ No FAIMS/IAA |
| P03930 | P03930 [49-57] | [K].TPWELKWTK.[I] | 1xHomocysteineIAA [K6] | 9,61721E-05 | 1362,68747 | Brain/ FAIMS/IAA |
| P03930 | P03930 [41-48] | [K].SLTTMKVK.[T] | 1xHomocysteineMMTS [K6] | 0,00680313 | 1070,54067 | Liver/ No FAIMS/MMTS |
| P03930 | P03930 [49-57] | [K].TPWELKWTK.[I] | 1xHomocysteineMMTS [K6] | 0,00599528 | 1351,65373 | Liver/ No FAIMS/MMTS |
| P03930 | P03930 [49-57] | [K].TPWELKWTK.[I] | 1xHomocysteineMMTS [K6] | 0,00599528 | 1351,65373 | Liver/ FAIMS/MMTS |
| P03930 | P03930 [41-48] | [K].SLTTMKVK.[T] | 1xHomocysteineMMTS [K6] | 0,00680313 | 1070,54067 | Liver/ FAIMS/MMTS |
| P03930 | P03930 [49-57] | [K].TPWELKWTK.[I] | 1xHomocysteineMMTS [K6] | 0,00599528 | 1351,65373 | Brain/ No FAIMS/MMTS |
| P03930 | P03930 [49-57] | [K].TPWELKWTK.[I] | 1xHomocysteineMMTS [K6] | 0,00599528 | 1351,65373 | Brain/ FAIMS/MMTS |
| P03995 | P03995 [186-198] | [R].KVESLEEEIQFLR.[K] | 1xHomocysteineIAA [K1] | 4,32935E-05 | 1793,91021 | Brain/ FAIMS/IAA |
| P04247 | P04247 [80-97] | [K].KGQHAAEIQPLAQSHATK.[H] | 1xHomocysteineIAA [K1] | 4,32935E-05 | 2089,06073 | Liver/ No FAIMS/IAA |
| P04247 | P04247 [80-97] | [K].KGQHAAEIQPLAQSHATK.[H] | 1xHomocysteineIAA [K1] | 4,32935E-05 | 2089,06073 | Liver/ FAIMS/IAA |
| P04247 | P04247 [52-64] | [K].SEEDMKGSEDLKK.[H] | 1xOxidation [M5]; 1xHomocysteineMMTS [K6] | 0,000530332 | 1674,70193 | Liver/ No FAIMS/MMTS |
| P04247 | P04247 [80-99] | [K].KGQHAAEIQPLAQSHATKHK.[I] | 1xHomocysteineMMTS [K18] | 0,000530332 | 2343,18086 | Liver/ No FAIMS/MMTS |
| P04247 | P04247 [18-46] | [K].VEADLAGHGQEVLIGLFKTHPETLDKFDK.[F] | 1xHomocysteineMMTS [K18] | 4,49774E-05 | 3370,68101 | Liver/ No FAIMS/MMTS |
| P04247 | P04247 [80-97] | [K].KGQHAAEIQPLAQSHATK.[H] | 1xHomocysteineMMTS [K1] | 4,49774E-05 | 2078,02699 | Liver/ No FAIMS/MMTS |
| P04247 | P04247 [36-48] | [K].THPETLDKFDKFK.[N] | 1xHomocysteineMMTS [K] | 4,49774E-05 | 1768,83969 | Liver/ No FAIMS/MMTS |
| P04247 | P04247 [52-64] | [K].SEEDMKGSEDLKK.[H] | 1xOxidation [M5]; 1xHomocysteineMMTS [K6] | 0,000530332 | 1674,70193 | Liver/ FAIMS/MMTS |
| P04247 | P04247 [36-46] | [K].THPETLDKFDK.[F] | 1xHomocysteineMMTS [K8] | 0,000437433 | 1493,67631 | Liver/ FAIMS/MMTS |
| P04247 | P04247 [80-99] | [K].KGQHAAEIQPLAQSHATKHK.[I] | 1xHomocysteineMMTS [K18] | 0,000530332 | 2343,18086 | Liver/ FAIMS/MMTS |
| P04247 | P04247 [18-46] | [K].VEADLAGHGQEVLIGLFKTHPETLDKFDK.[F] | 1xHomocysteineMMTS [K18] | 4,49774E-05 | 3370,68101 | Liver/ FAIMS/MMTS |
| P04247 | P04247 [81-99] | [K].GQHAAEIQPLAQSHATKHK.[I] | 1xHomocysteineMMTS [K17] | 4,49774E-05 | 2215,0859 | Liver/ FAIMS/MMTS |
| P04247 | P04247 [120-140] | [R].HSGDFGADAQGAMSKALELFR.[N] | 1xOxidation [M13]; 1xHomocysteineMMTS [K15] | 4,49774E-05 | 2387,0577 | Liver/ FAIMS/MMTS |
| P04247 | P04247 [79-97] | [K].KKGQHAAEIQPLAQSHATK.[H] | 1xHomocysteineMMTS [K2] | 0,00261874 | 2206,12195 | Liver/ FAIMS/MMTS |
| P04247 | P04247 [18-43] | [K].VEADLAGHGQEVLIGLFKTHPETLDK.[F] | 1xHomocysteineMMTS [K18] | 4,49774E-05 | 2980,49069 | Liver/ FAIMS/MMTS |
| P04247 | P04247 [120-140] | [R].HSGDFGADAQGAMSKALELFR.[N] | 1xHomocysteineMMTS [K15] | 4,49774E-05 | 2371,06278 | Liver/ FAIMS/MMTS |
| P04247 | P04247 [80-97] | [K].KGQHAAEIQPLAQSHATK.[H] | 1xHomocysteineMMTS [K1] | 4,49774E-05 | 2078,02699 | Liver/ FAIMS/MMTS |
| P04247 | P04247 [36-48] | [K].THPETLDKFDKFK.[N] | 1xHomocysteineMMTS [K] | 4,49774E-05 | 1768,83969 | Liver/ FAIMS/MMTS |
| P04370 | P04370 [144-157] | [R].SKYLATASTMDHAR.[H] | 1xOxidation [M10]; 1xHomocysteineIAA [K2] | 4,32935E-05 | 1741,79961 | Brain/ No FAIMS/IAA |
| P04370 | P04370 [144-157] | [R].SKYLATASTMDHAR.[H] | 1xOxidation [M10]; 1xHomocysteineIAA [K2] | 4,32935E-05 | 1741,79961 | Brain/ FAIMS/IAA |
| P04370 | P04370 [144-157] | [R].SKYLATASTMDHAR.[H] | 1xOxidation [M10]; 1xHomocysteineMMTS [K2] | 4,49774E-05 | 1730,76587 | Brain/ No FAIMS/MMTS |
| P04370 | P04370 [144-157] | [R].SKYLATASTMDHAR.[H] | 1xHomocysteineMMTS [K2] | 4,49774E-05 | 1714,77096 | Brain/ No FAIMS/MMTS |
| P04370 | P04370 [211-228] | [R].TQDENPVVHFFKNIVTPR.[T] | 1xHomocysteineMMTS [K12] | 4,49774E-05 | 2304,12637 | Brain/ No FAIMS/MMTS |

| | | | | | | |
|---|---|---|---|---|---|---|
| P04370 | P04370 [196-210] | [R].TTHYGSLPQKSQHGR.[T] | 1xHomocysteineMMTS [K10] | 4,49774E-05 | 1859,86395 | Brain/ No FAIMS/MMTS |
| P04370 | P04370 [144-157] | [R].SKYLATASTMDHAR.[H] | 1xOxidation [M10]; 1xHomocysteineMMTS [K2] | 4,49774E-05 | 1730,76587 | Brain/ FAIMS/MMTS |
| P04370 | P04370 [144-157] | [R].SKYLATASTMDHAR.[H] | 1xHomocysteineMMTS [K2] | 4,49774E-05 | 1714,77096 | Brain/ FAIMS/MMTS |
| P04370 | P04370 [211-228] | [R].TQDENPVVHFFKNIVTPR.[T] | 1xHomocysteineMMTS [K12] | 4,49774E-05 | 2304,12637 | Brain/ FAIMS/MMTS |
| P04370 | P04370 [196-210] | [R].TTHYGSLPQKSQHGR.[T] | 1xHomocysteineMMTS [K10] | 4,49774E-05 | 1859,86395 | Brain/ FAIMS/MMTS |
| P04925 | P04925 [208-228] | [R].VVEQMCVTQYQKESQAYYDGR.[R] | 1xCarbamidomethyl [C6]; 1xHomocysteineIAA [K12] | 4,32935E-05 | 2756,2113 | Brain/ FAIMS/IAA |
| P04939 | P04939 [132-144] | [K].DGKTFQLMELYGR.[EK] | 1xHomocysteineIAA [K3] | 9,61721E-05 | 1731,81929 | Liver/ FAIMS/IAA |
| P04939; P11590 | P04939 [132-144]; P11590 [126-138] | [K].DGKTFQLMELYGR.[EK] | 1xHomocysteineMMTS [K3] | 0,00107839 | 1720,78554 | Liver/ FAIMS/MMTS |
| P04939; P11590; Q5FW60 | P04939 [52-61]; P11590 [46-55]; Q5FW60 [49-58] | [R].EKIEEHGSMR.[AV] | 1xHomocysteineMMTS [K2] | 4,49774E-05 | 1378,5912 | Liver/ No FAIMS/MMTS |
| P04939; P11590; Q5FW60 | P04939 [52-61]; P11590 [46-55]; Q5FW60 [49-58] | [R].EKIEEHGSMR.[AV] | 1xHomocysteineMMTS [K2] | 4,49774E-05 | 1378,5912 | Liver/ FAIMS/MMTS |
| P04939; Q5FW60 | P04939 [117-131]; Q5FW60 [114-128] | [K].TDYDNYIMIHLINKK.[D] | 1xHomocysteineIAA [K14] | 0,000243434 | 2055,00379 | Liver/ FAIMS/IAA |
| P04939; Q5FW60 | P04939 [117-131]; Q5FW60 [114-128] | [K].TDYDNYIMIHLINKK.[D] | 1xHomocysteineMMTS [K14] | 6,70238E-05 | 2043,97005 | Liver/ No FAIMS/MMTS |
| P04939; Q5FW60 | P04939 [117-131]; Q5FW60 [114-128] | [K].TDYDNYIMIHLINKK.[D] | 1xHomocysteineMMTS [K14] | 6,70238E-05 | 2043,97005 | Liver/ FAIMS/MMTS |
| P05063 | P05063 [72-87] | [K].KCIGGVIFFHETLYQK.[D] | 1xCarbamidomethyl [C2]; 1xHomocysteineIAA [K1] | 4,32935E-05 | 2114,05616 | Brain/ No FAIMS/IAA |
| P05063 | P05063 [72-87] | [K].KCIGGVIFFHETLYQK.[D] | 1xCarbamidomethyl [C2]; 1xHomocysteineIAA [K1] | 4,32935E-05 | 2114,05616 | Brain/ FAIMS/IAA |
| P05063 | P05063 [102-111] | [K].GILVGIKVDK.[G] | 1xHomocysteineMMTS [K7] | 0,00911075 | 1204,67921 | Brain/ No FAIMS/MMTS |
| P05063 | P05063 [23-42] | [R].IVTPGKGILAADESVGSMAK.[R] | 1xHomocysteineMMTS [K6] | 4,49774E-05 | 2107,05959 | Brain/ No FAIMS/MMTS |
| P05063 | P05063 [109-134] | [K].VDKGVVPLAGTDGETTTQGLDGLLER.[C] | 1xHomocysteineMMTS [K3] | 4,49774E-05 | 2804,38047 | Brain/ No FAIMS/MMTS |
| P05063 | P05063 [29-43] | [K].GILAADESVGSMAKR.[L] | 1xHomocysteineMMTS [K14] | 4,49774E-05 | 1667,79136 | Brain/ No FAIMS/MMTS |
| P05063 | P05063 [14-22] | [K].KELSDIALR.[I] | 1xHomocysteineMMTS [K1] | 0,000829616 | 1207,61734 | Brain/ No FAIMS/MMTS |
| P05063 | P05063 [97-108] | [R].TIQDKGILVGIK.[V] | 1xHomocysteineMMTS [K5] | 0,000266738 | 1447,80112 | Brain/ FAIMS/MMTS |
| P05063 | P05063 [109-134] | [K].VDKGVVPLAGTDGETTTQGLDGLLER.[C] | 1xHomocysteineMMTS [K3] | 4,49774E-05 | 2804,38047 | Brain/ FAIMS/MMTS |
| P05063 | P05063 [29-43] | [K].GILAADESVGSMAKR.[L] | 1xHomocysteineMMTS [K14] | 4,49774E-05 | 1667,79136 | Brain/ FAIMS/MMTS |
| P05063 | P05063 [319-331] | [R].DNAGAATEEFIKR.[A] | 1xHomocysteineMMTS [K12] | 4,49774E-05 | 1584,71449 | Brain/ FAIMS/MMTS |
| P05063 | P05063 [14-22] | [K].KELSDIALR.[I] | 1xHomocysteineMMTS [K1] | 0,000829616 | 1207,61734 | Brain/ FAIMS/MMTS |
| P05063; P05064 | P05063 [202-215]; P05064 [202-215] | [R].CQYVTEKVLAAVYK.[A] | 1xCarbamidomethyl [C1]; 1xHomocysteineIAA [K7] | 4,32935E-05 | 1845,92375 | Brain/ No FAIMS/IAA |
| P05063; P05064 | P05063 [202-215]; P05064 [202-215] | [R].CQYVTEKVLAAVYK.[A] | 1xCarbamidomethyl [C1]; 1xHomocysteineIAA [K7] | 4,32935E-05 | 1845,92375 | Brain/ FAIMS/IAA |
| P05063; P05064 | P05063 [141-149]; P05064 [141-149] | [K].DGADFAKWR.[C] | 1xHomocysteineMMTS [K7] | 0,00074635 | 1228,52377 | Brain/ No FAIMS/MMTS |
| P05063; P05064 | P05063 [141-149]; P05064 [141-149] | [K].DGADFAKWR.[C] | 1xHomocysteineMMTS [K7] | 0,00074635 | 1228,52377 | Brain/ FAIMS/MMTS |
| P05063; P05064 | P05063 [140-147]; P05064 [140-147] | [K].KDGADFAK.[W] | 1xHomocysteineMMTS [K1] | 0,00677626 | 1014,43831 | Brain/ FAIMS/MMTS |
| P05064 | P05064 [14-22] | [K].KELSDIAHR.[I] | 1xHomocysteineIAA [K1] | 4,32935E-05 | 1242,62593 | Brain/ No FAIMS/IAA |
| P05064 | P05064 [14-22] | [K].KELSDIAHR.[I] | 1xHomocysteineIAA [K1] | 4,32935E-05 | 1242,62593 | Brain/ FAIMS/IAA |
| P05064 | P05064 [323-331] | [K].AAQEEYIKR.[A] | 1xHomocysteineMMTS [K8] | 0,000250158 | 1270,59185 | Liver/ No FAIMS/MMTS |
| P05064 | P05064 [23-42] | [R].IVAPGKGILAADESTGSIAK.[R] | 1xHomocysteineMMTS [K6] | 4,49774E-05 | 2061,07187 | Liver/ No FAIMS/MMTS |
| P05064 | P05064 [323-331] | [K].AAQEEYIKR.[A] | 1xHomocysteineMMTS [K8] | 0,000250158 | 1270,59185 | Liver/ FAIMS/MMTS |
| P05064 | P05064 [102-111] | [K].GGVVGIKVDK.[G] | 1xHomocysteineMMTS [K7] | 4,49774E-05 | 1134,60096 | Liver/ FAIMS/MMTS |
| P05064 | P05064 [23-42] | [R].IVAPGKGILAADESTGSIAK.[R] | 1xHomocysteineMMTS [K6] | 4,49774E-05 | 2061,07187 | Liver/ FAIMS/MMTS |
| P05064 | P05064 [29-43] | [K].GILAADESTGSIAKR.[L] | 1xHomocysteineMMTS [K14] | 4,49774E-05 | 1651,8142 | Liver/ FAIMS/MMTS |
| P05064 | P05064 [14-22] | [K].KELSDIAHR.[I] | 1xHomocysteineMMTS [K1] | 4,49774E-05 | 1231,59219 | Liver/ FAIMS/MMTS |
| P05064 | P05064 [323-331] | [K].AAQEEYIKR.[A] | 1xHomocysteineMMTS [K8] | 0,000250158 | 1270,59185 | Brain/ FAIMS/MMTS |
| P05064 | P05064 [1-14] | [-].MPHPYPALTPEQKK.[E] | 1xMet-loss [N-Term]; 1xHomocysteineMMTS [K13] | 4,49774E-05 | 1668,82364 | Brain/ FAIMS/MMTS |
| P05064 | P05064 [323-331] | [K].AAQEEYIKR.[A] | 1xHomocysteineMMTS [K8] | 0,000250158 | 1270,59185 | Brain/ FAIMS/MMTS |
| P05064 | P05064 [102-111] | [K].GGVVGIKVDK.[G] | 1xHomocysteineMMTS [K7] | 4,49774E-05 | 1134,60096 | Brain/ FAIMS/MMTS |
| P05064 | P05064 [23-42] | [R].IVAPGKGILAADESTGSIAK.[R] | 1xHomocysteineMMTS [K6] | 4,49774E-05 | 2061,07187 | Brain/ FAIMS/MMTS |
| P05064 | P05064 [319-330] | [K].ENLKAAQEEYIK.[R] | 1xHomocysteineMMTS [K4] | 0,00452171 | 1598,75529 | Brain/ FAIMS/MMTS |
| P05064 | P05064 [29-43] | [K].GILAADESTGSIAKR.[L] | 1xHomocysteineMMTS [K14] | 4,49774E-05 | 1651,8142 | Brain/ FAIMS/MMTS |
| P05064 | P05064 [88-101] | [K].ADDGRPFPQVIKSK.[G] | 1xHomocysteineMMTS [K12] | 0,00199616 | 1720,85092 | Brain/ FAIMS/MMTS |
| P05064 | P05064 [14-22] | [K].KELSDIAHR.[I] | 1xHomocysteineMMTS [K1] | 4,49774E-05 | 1231,59219 | Brain/ FAIMS/MMTS |
| P05132; P68181 | P05132 [258-271]; P68181 [258-271] | [R].FPSHFSSDLKDLLR.[N] | 1xHomocysteineIAA [K10] | 4,32935E-05 | 1835,91088 | Brain/ FAIMS/IAA |
| P05201 | P05201 [306-321] | [R].IVAATLSDPELFKEWK.[G] | 1xHomocysteineIAA [K13] | 4,32935E-05 | 2021,04122 | Brain/ No FAIMS/IAA |
| P05201 | P05201 [306-321] | [R].IVAATLSDPELFKEWK.[G] | 1xHomocysteineIAA [K13] | 4,32935E-05 | 2021,04122 | Brain/ FAIMS/IAA |
| P05201 | P05201 [268-283] | [R].VGNLTVVGKESDSVLR.[V] | 1xHomocysteineMMTS [K9] | 0,00137282 | 1835,93538 | Liver/ FAIMS/MMTS |
| P05201 | P05201 [306-321] | [R].IVAATLSDPELFKEWK.[G] | 1xHomocysteineMMTS [K13] | 0,000152766 | 2010,00748 | Liver/ FAIMS/MMTS |
| P05201 | P05201 [284-293] | [R].VLSQMEKIVR.[I] | 1xHomocysteineMMTS [K7] | 0,0004709 | 1365,70511 | Brain/ FAIMS/MMTS |
| P05201 | P05201 [306-321] | [R].IVAATLSDPELFKEWK.[G] | 1xHomocysteineMMTS [K13] | 0,000152766 | 2010,00748 | Brain/ FAIMS/MMTS |
| P05202 | P05202 [408-430] | [R].ISVAGVTSGNVGYLAHAIHQVTK.[-] | 1xHomocysteineIAA [K23] | 4,32935E-05 | 2496,30275 | Brain/ FAIMS/IAA |
| P05202 | P05202 [394-404] | [R].LTKEFSVYMTK.[D] | 1xHomocysteineMMTS [K3] | 0,00074635 | 1509,71501 | Brain/ FAIMS/MMTS |
| P05202 | P05202 [408-430] | [R].ISVAGVTSGNVGYLAHAIHQVTK.[-] | 1xHomocysteineMMTS [K23] | 0,00413569 | 2485,26901 | Brain/ FAIMS/MMTS |
| P05214; P68369; P68368 | P05214 [265-304]; P68369 [265-304]; P68368 [265-304] | [R].IHFPLATYAPVISAEKAYHEQLSVAEITNACFEPANQMVK.[C] | 1xCarbamidomethyl [C31]; 1xHomocysteineIAA [K16] | 4,32935E-05 | 4662,28295 | Brain/ No FAIMS/IAA |

| | | | | | | |
|---|---|---|---|---|---|---|
| P05214; P68369; P68368 | P05214 [265-304]; P68369 [265-304]; P68368 [265-304] | [R].IHFPLATYAPVISAEKAYHEQLSVAEITNACFEPANQMVK.[C] | 1xCarbamidomethyl [C31]; 1xHomocysteineIAA [K16] | 4,32935E-05 | 4662,28295 | Brain/ FAIMS/IAA |
| P05214; P68369; P68373 | P05214 [327-338]; P68369 [327-338]; P68373 [327-338] | [K].DVNAAIATIKTK.[R] | 1xHomocysteineIAA [K] | 4,32935E-05 | 1418,76718 | Brain/ No FAIMS/IAA |
| P05214; P68369; P68373 | P05214 [327-338]; P68369 [327-338]; P68373 [327-338] | [K].DVNAAIATIKTK.[R] | 1xHomocysteineIAA [K] | 4,32935E-05 | 1418,76718 | Brain/ FAIMS/IAA |
| P05214; P68369; P68373 | P05214 [327-338]; P68369 [327-338]; P68373 [327-338] | [K].DVNAAIATIKTK.[R] | 1xHomocysteineMMTS [K10] | 4,49774E-05 | 1407,73343 | Liver/ FAIMS/MMTS |
| P05214; P68369; P68373 | P05214 [321-336]; P68369 [321-336]; P68373 [321-336] | [R].GDVVPKDVNAAIATIK.[T] | 1xHomocysteineMMTS [K6] | 4,49774E-05 | 1773,92375 | Brain/ No FAIMS/MMTS |
| P05214; P68369; P68373 | P05214 [41-64]; P68369 [41-64]; P68373 [41-64] | [K].TIGGGDDSFNTFFSETGAGKHVPR.[A] | 1xHomocysteineMMTS [K20] | 4,49774E-05 | 2660,1868 | Brain/ No FAIMS/MMTS |
| P05214; P68369; P68373 | P05214 [327-338]; P68369 [327-338]; P68373 [327-338] | [K].DVNAAIATIKTK.[R] | 1xHomocysteineMMTS [K10] | 4,49774E-05 | 1407,73343 | Brain/ No FAIMS/MMTS |
| P05214; P68369; P68373 | P05214 [321-336]; P68369 [321-336]; P68373 [321-336] | [R].GDVVPKDVNAAIATIK.[T] | 1xHomocysteineMMTS [K6] | 4,49774E-05 | 1773,92375 | Brain/ FAIMS/MMTS |
| P05214; P68369; P68373 | P05214 [41-64]; P68369 [41-64]; P68373 [41-64] | [K].TIGGGDDSFNTFFSETGAGKHVPR.[A] | 1xHomocysteineMMTS [K20] | 4,49774E-05 | 2660,1868 | Brain/ FAIMS/MMTS |
| P05214; P68369; P68373 | P05214 [327-338]; P68369 [327-338]; P68373 [327-338] | [K].DVNAAIATIKTK.[R] | 1xHomocysteineMMTS [K10] | 4,49774E-05 | 1407,73343 | Brain/ FAIMS/MMTS |
| P05214; P68369; P68373; P68368 | P05214 [157-164]; P68369 [157-164]; P68373 [157-164]; P68368 [157-164] | [R].LSVDYGKK.[S] | 1xHomocysteineIAA [K7] | 0,000162511 | 1083,55031 | Brain/ No FAIMS/IAA |
| P05214; P68369; P68373; P68368 | P05214 [157-164]; P68369 [157-164]; P68373 [157-164]; P68368 [157-164] | [R].LSVDYGKK.[S] | 1xHomocysteineIAA [K7] | 0,000162511 | 1083,55031 | Brain/ FAIMS/IAA |
| P05214; Q9JJZ2; P68369; P68373 | P05214 [340-370]; Q9JJZ2 [340-370]; P68369 [340-370]; P68373 [340-370] | [R].TIQFVDWCPTGFKVGINYQPPTVVPGGDLAK.[V] | 1xCarbamidomethyl [C8]; 1xHomocysteineIAA [K13] | 4,32935E-05 | 3578,78106 | Brain/ No FAIMS/IAA |
| P05214; Q9JJZ2; P68369; P68373 | P05214 [340-370]; Q9JJZ2 [340-370]; P68369 [340-370]; P68373 [340-370] | [R].TIQFVDWCPTGFKVGINYQPPTVVPGGDLAK.[V] | 1xCarbamidomethyl [C8]; 1xHomocysteineIAA [K13] | 4,32935E-05 | 3578,78106 | Brain/ FAIMS/IAA |
| P05214; Q9JJZ2; P68369; P68373; P68368 | P05214 [85-105]; Q9JJZ2 [85-105]; P68369 [85-105]; P68373 [85-105]; P68368 [85-105] | [R].QLFHPEQLITGKEDAANNYAR.[G] | 1xHomocysteineIAA [K12] | 4,32935E-05 | 2589,25144 | Brain/ No FAIMS/IAA |
| P05214; Q9JJZ2; P68369; P68373; P68368 | P05214 [85-105]; Q9JJZ2 [85-105]; P68369 [85-105]; P68373 [85-105]; P68368 [85-105] | [R].QLFHPEQLITGKEDAANNYAR.[G] | 1xHomocysteineIAA [K12] | 4,32935E-05 | 2589,25144 | Brain/ FAIMS/IAA |
| P05214; Q9JJZ2; P68369; P68373; P68368 | P05214 [395-402]; Q9JJZ2 [395-402]; P68369 [395-402]; P68373 [395-402]; P68368 [395-402] | [K].FDLMYAKR.[A] | 1xHomocysteineMMTS [K7] | 0,000599431 | 1206,54682 | Liver/ FAIMS/MMTS |
| P05214; Q9JJZ2; P68369; P68373; P68368 | P05214 [395-402]; Q9JJZ2 [395-402]; P68369 [395-402]; P68373 [395-402]; P68368 [395-402] | [K].FDLMYAKR.[A] | 1xHomocysteineMMTS [K7] | 0,000599431 | 1206,54682 | Brain/ No FAIMS/MMTS |
| P05214; Q9JJZ2; P68369; P68373; P68368 | P05214 [353-373]; Q9JJZ2 [353-373]; P68369 [353-373]; P68373 [353-373]; P68368 [353-373] | [K].VGINYQPPTVVPGGDLAKVQR.[A] | 1xHomocysteineMMTS [K18] | 4,49774E-05 | 2371,22608 | Brain/ No FAIMS/MMTS |
| P05214; Q9JJZ2; P68369; P68373; P68368 | P05214 [395-402]; Q9JJZ2 [395-402]; P68369 [395-402]; P68373 [395-402]; P68368 [395-402] | [K].FDLMYAKR.[A] | 1xHomocysteineMMTS [K7] | 0,000599431 | 1206,54682 | Brain/ FAIMS/MMTS |
| P05214; Q9JJZ2; P68369; P68373; P68368 | P05214 [353-373]; Q9JJZ2 [353-373]; P68369 [353-373]; P68373 [353-373]; P68368 [353-373] | [K].VGINYQPPTVVPGGDLAKVQR.[A] | 1xHomocysteineMMTS [K18] | 4,49774E-05 | 2371,22608 | Brain/ FAIMS/MMTS |

| | | | | | | |
|---|---|---|---|---|---|---|
| P05214; | P05214 [85-105]; | | | | | |
| Q9JJZ2; | Q9JJZ2 [85-105]; | | | | | |
| P68369; | P68369 [85-105]; | | | | | |
| P68373; | P68373 [85-105]; | | | | | |
| P68368 | P68368 [85-105] | [R].QLFHPEQLITGKEDAANNYAR.[G] | 1xHomocysteineMMTS [K12] | 4,49774E-05 | 2578,2177 | Brain/ FAIMS/MMTS |
| P05214; | P05214 [391-401]; | | | | | |
| Q9JJZ2; | Q9JJZ2 [391-401]; | | | | | |
| Q3UX10; | Q3UX10 [398-408]; | | | | | |
| P68369; | P68369 [391-401]; | | | | | |
| P68373; | P68373 [391-401]; | | | | | |
| P68368 | P68368 [391-401] | [R].LDHKFDLMYAK.[RK] | 1xHomocysteineIAA [K4] | 4,32935E-05 | 1554,74433 | Brain/ No FAIMS/IAA |
| P05214; | P05214 [391-401]; | | | | | |
| Q9JJZ2; | Q9JJZ2 [391-401]; | | | | | |
| Q3UX10; | Q3UX10 [398-408]; | | | | | |
| P68369; | P68369 [391-401]; | | | | | |
| P68373; | P68373 [391-401]; | | | | | |
| P68368 | P68368 [391-401] | [R].LDHKFDLMYAK.[RK] | 1xHomocysteineIAA [K4] | 4,32935E-05 | 1554,74433 | Brain/ FAIMS/IAA |
| P05214; | P05214 [391-401]; | | | | | |
| Q9JJZ2; | Q9JJZ2 [391-401]; | | | | | |
| Q3UX10; | Q3UX10 [398-408]; | | | | | |
| P68369; | P68369 [391-401]; | | | | | |
| P68373; | P68373 [391-401]; | | | | | |
| P68368 | P68368 [391-401] | [R].LDHKFDLMYAK.[RK] | 1xHomocysteineMMTS [K4] | 0,00107839 | 1543,71059 | Brain/ FAIMS/MMTS |
| P05784 | P05784 [90-98] | [K].VKSLETENR.[R] | 1xHomocysteineMMTS [K2] | 0,00459586 | 1238,58677 | Liver/ FAIMS/MMTS |
| P05784 | P05784 [180-189] | [R].KVVDDTNITR.[L] | 1xHomocysteineMMTS [K1] | 0,00924953 | 1323,63953 | Liver/ FAIMS/MMTS |
| P05977 | P05977 [5-28] | [K].KDVKKPAAAPAPAPAPAPAPAKPK.[E] | 1xHomocysteineIAA [K] | 4,32935E-05 | 2463,39045 | Liver/ No FAIMS/IAA |
| P05977 | P05977 [5-28] | [K].KDVKKPAAAPAPAPAPAPAPAKPK.[E] | 1xHomocysteineIAA [K] | 4,32935E-05 | 2463,39045 | Liver/ FAIMS/IAA |
| | | | 1xOxidation [M10]; | | | |
| P05977 | P05977 [88-101] | [K].VLGNPSNEEMNAKK.[I] | 1xHomocysteineMMTS [K13] | 4,49774E-05 | 1709,76554 | Liver/ No FAIMS/MMTS |
| P05977 | P05977 [32-43] | [K].IDLSAIKIEFSK.[E] | 1xHomocysteineMMTS [K7] | 4,49774E-05 | 1526,7957 | Liver/ No FAIMS/MMTS |
| P05977 | P05977 [44-58] | [K].EQQEDFKEAFLLFDR.[T] | 1xHomocysteineMMTS [K7] | 4,49774E-05 | 2077,93577 | Liver/ No FAIMS/MMTS |
| P05977 | P05977 [39-50] | [K].IEFSKEQQEDFK.[E] | 1xHomocysteineMMTS [K5] | 4,49774E-05 | 1690,74512 | Liver/ No FAIMS/MMTS |
| P05977 | P05977 [29-38] | [K].EEKIDLSAIK.[I] | 1xHomocysteineMMTS [K3] | 0,000211896 | 1308,65379 | Liver/ No FAIMS/MMTS |
| P05977 | P05977 [75-87] | [R].ALGTNPTNAEVKK.[V] | 1xHomocysteineMMTS [K12] | 4,49774E-05 | 1505,74506 | Liver/ No FAIMS/MMTS |
| P05977 | P05977 [87-100] | [K].KVLGNPSNEEMNAK.[K] | 1xHomocysteineMMTS [K1] | 4,49774E-05 | 1693,77062 | Liver/ No FAIMS/MMTS |
| P05977 | P05977 [9-28] | [K].KPAAAPAPAPAPAPAPAKPK.[E] | 1xHomocysteineMMTS [K] | 4,49774E-05 | 1982,07142 | Liver/ No FAIMS/MMTS |
| P05977 | P05977 [9-31] | [K].KPAAAPAPAPAPAPAPAKPKEEK.[I] | 1xHomocysteineMMTS [K] | 4,49774E-05 | 2368,25157 | Liver/ No FAIMS/MMTS |
| | | | 1xMethylthio [C4]; | | | |
| P05977 | P05977 [59-74] | [R].TGECKITLSQVGDVLR.[A] | 1xHomocysteineMMTS [K5] | 0,000266738 | 1927,91082 | Liver/ FAIMS/MMTS |
| P05977 | P05977 [32-43] | [K].IDLSAIKIEFSK.[E] | 1xHomocysteineMMTS [K7] | 4,49774E-05 | 1526,7957 | Liver/ FAIMS/MMTS |
| P05977 | P05977 [44-58] | [K].EQQEDFKEAFLLFDR.[T] | 1xHomocysteineMMTS [K7] | 4,49774E-05 | 2077,93577 | Liver/ FAIMS/MMTS |
| P05977 | P05977 [39-58] | [K].IEFSKEQQEDFKEAFLLFDR.[T] | 1xHomocysteineMMTS [K5] | 4,49774E-05 | 2682,25784 | Liver/ FAIMS/MMTS |
| P05977 | P05977 [29-38] | [K].EEKIDLSAIK.[I] | 1xHomocysteineMMTS [K3] | 0,000211896 | 1308,65379 | Liver/ FAIMS/MMTS |
| P05977 | P05977 [88-101] | [K].VLGNPSNEEMNAKK.[I] | 1xHomocysteineMMTS [K13] | 4,49774E-05 | 1693,77062 | Liver/ FAIMS/MMTS |
| P05977 | P05977 [75-87] | [R].ALGTNPTNAEVKK.[V] | 1xHomocysteineMMTS [K12] | 4,49774E-05 | 1505,74506 | Liver/ FAIMS/MMTS |
| P05977 | P05977 [87-100] | [K].KVLGNPSNEEMNAK.[K] | 1xHomocysteineMMTS [K1] | 4,49774E-05 | 1693,77062 | Liver/ FAIMS/MMTS |
| P05977 | P05977 [9-28] | [K].KPAAAPAPAPAPAPAPAKPK.[E] | 1xHomocysteineMMTS [K] | 4,49774E-05 | 1982,07142 | Liver/ FAIMS/MMTS |
| P05977 | P05977 [9-31] | [K].KPAAAPAPAPAPAPAPAKPKEEK.[I] | 1xHomocysteineMMTS [K] | 4,49774E-05 | 2368,25157 | Liver/ FAIMS/MMTS |
| | | | 1xOxidation [M11]; | | | |
| P05977 | P05977 [87-100] | [K].KVLGNPSNEEMNAK.[K] | 1xHomocysteineMMTS [K1] | 6,70238E-05 | 1709,76554 | Liver/ FAIMS/MMTS |
| P05977 | P05977 [39-50] | [K].IEFSKEQQEDFK.[E] | 1xHomocysteineMMTS [K5] | 4,49774E-05 | 1690,74512 | Liver/ FAIMS/MMTS |
| P05977 | P05977 [5-28] | [K].KDVKKPAAAPAPAPAPAPAPAKPK.[E] | 1xHomocysteineMMTS [K5] | 4,49774E-05 | 2452,3567 | Liver/ FAIMS/MMTS |
| P05977 | P05977 [6-28] | [K].DVKKPAAAPAPAPAPAPAPAKPK.[E] | 1xHomocysteineMMTS [K4] | 4,49774E-05 | 2324,26174 | Liver/ FAIMS/MMTS |
| P06151 | P06151 [213-228] | [K].SLNPELGTDADKEQWK.[E] | 1xHomocysteineIAA [K12] | 4,32935E-05 | 2004,93313 | Brain/ FAIMS/IAA |
| P06728 | P06728 [256-265] | [K].VSAKIDQLQK.[N] | 1xHomocysteineMMTS [K4] | 0,0021186 | 1292,6701 | Liver/ FAIMS/MMTS |
| P06745 | P06745 [235-252] | [K].DPSAVAKHFVALSTNTAK.[V] | 1xHomocysteineIAA [K7] | 4,32935E-05 | 2031,03279 | Brain/ FAIMS/IAA |
| P06745 | P06745 [82-96] | [R].DNMFSGSKINYTENR.[A] | 1xHomocysteineMMTS [K8] | 0,000437433 | 1938,81428 | Brain/ FAIMS/MMTS |
| P06745 | P06745 [19-27] | [R].ANSANLKLR.[E] | 1xHomocysteineMMTS [K7] | 0,0046895 | 1149,58671 | Brain/ FAIMS/MMTS |
| P06745 | P06745 [235-252] | [K].DPSAVAKHFVALSTNTAK.[V] | 1xHomocysteineMMTS [K7] | 6,70238E-05 | 2019,99904 | Brain/ FAIMS/MMTS |
| P06837 | P06837 [81-98] | [K].KEGDGSATTDAAPATSPK.[A] | 1xHomocysteineIAA [K1] | 4,32935E-05 | 1877,85455 | Brain/ No FAIMS/IAA |
| P06837 | P06837 [81-98] | [K].KEGDGSATTDAAPATSPK.[A] | 1xHomocysteineIAA [K1] | 4,32935E-05 | 1877,85455 | Brain/ FAIMS/IAA |
| P06837 | P06837 [147-157] | [K].AEDGPAKEEPK.[Q] | 1xHomocysteineMMTS [K7] | 0,001242 | 1333,57626 | Brain/ No FAIMS/MMTS |
| P06837 | P06837 [99-113] | [K].AEEPSKAGDAPSEEK.[K] | 1xHomocysteineMMTS [K6] | 4,49774E-05 | 1707,72003 | Brain/ No FAIMS/MMTS |
| P06837 | P06837 [213-227] | [R].QDEGKEDPEADQEHA.[-] | 1xHomocysteineMMTS [K5] | 0,00074635 | 1860,70108 | Brain/ No FAIMS/MMTS |
| | | [K].AAQPPTETAESSQAEEEKDAVDEAKPKES | | | | |
| P06837 | P06837 [182-212] | AR.[Q] | 1xHomocysteineMMTS [K27] | 4,49774E-05 | 3462,56353 | Brain/ No FAIMS/MMTS |
| P06837 | P06837 [114-136] | [K].KGEGDAAPSEEKAGSAETESAAK.[A] | 1xHomocysteineMMTS [K12] | 4,49774E-05 | 2383,03879 | Brain/ No FAIMS/MMTS |
| P06837 | P06837 [126-146] | [K].AGSAETESAAKATTDNSPSSK.[A] | 1xHomocysteineMMTS [K11] | 4,49774E-05 | 2172,93835 | Brain/ No FAIMS/MMTS |
| P06837 | P06837 [56-67] | [K].KGDAPAAEAEAK.[E] | 1xHomocysteineMMTS [K1] | 4,49774E-05 | 1320,59225 | Brain/ No FAIMS/MMTS |
| | | [K].AAQPPTETAESSQAEEEKDAVDEAKPK.[ | | | | |
| P06837 | P06837 [182-208] | E] | 1xHomocysteineMMTS [K] | 4,49774E-05 | 3019,35069 | Brain/ No FAIMS/MMTS |
| P06837 | P06837 [147-157] | [K].AEDGPAKEEPK.[Q] | 1xHomocysteineMMTS [K7] | 0,001242 | 1333,57626 | Brain/ FAIMS/MMTS |
| P06837 | P06837 [99-113] | [K].AEEPSKAGDAPSEEK.[K] | 1xHomocysteineMMTS [K6] | 4,49774E-05 | 1707,72003 | Brain/ FAIMS/MMTS |
| P06837 | P06837 [213-227] | [R].QDEGKEDPEADQEHA.[-] | 1xHomocysteineMMTS [K5] | 0,00074635 | 1860,70108 | Brain/ FAIMS/MMTS |
| P06837 | P06837 [82-104] | [K].EGDGSATTDAAPATSPKAEEPSK.[A] | 1xHomocysteineMMTS [K17] | 4,49774E-05 | 2380,02789 | Brain/ FAIMS/MMTS |
| P06837 | P06837 [114-136] | [K].KGEGDAAPSEEKAGSAETESAAK.[A] | 1xHomocysteineMMTS [K12] | 4,49774E-05 | 2383,03879 | Brain/ FAIMS/MMTS |
| P06837 | P06837 [126-146] | [K].AGSAETESAAKATTDNSPSSK.[A] | 1xHomocysteineMMTS [K11] | 4,49774E-05 | 2172,93835 | Brain/ FAIMS/MMTS |
| P06837 | P06837 [137-157] | [K].ATTDNSPSSKAEDGPAKEEPK.[Q] | 1xHomocysteineMMTS [K10] | 4,49774E-05 | 2322,02242 | Brain/ FAIMS/MMTS |
| P06837 | P06837 [105-125] | [K].AGDAPSEEKKGEGDAAPSEEK.[A] | 1xHomocysteineMMTS [K10] | 0,000706375 | 2264,96457 | Brain/ FAIMS/MMTS |
| P06837 | P06837 [81-98] | [K].KEGDGSATTDAAPATSPK.[A] | 1xHomocysteineMMTS [K1] | 4,49774E-05 | 1866,8208 | Brain/ FAIMS/MMTS |
| P06837 | P06837 [56-67] | [K].KGDAPAAEAEAK.[E] | 1xHomocysteineMMTS [K1] | 4,49774E-05 | 1320,59225 | Brain/ FAIMS/MMTS |
| P06837 | P06837 [114-125] | [K].KGEGDAAPSEEK.[A] | 1xHomocysteineMMTS [K1] | 0,000879882 | 1380,57699 | Brain/ FAIMS/MMTS |

| | | | | | | |
|---|---|---|---|---|---|---|
| P06837 | P06837 [182-208] | [K].AAQPPTETAESSQAEEEKDAVDEAKPK.[E] | 1xHomocysteineMMTS [K] | 4,49774E-05 | 3019,35069 | Brain/ FAIMS/MMTS |
| P07309 | P07309 [69-90] | [K].TAESGELHGLTTDEKFVEGVYR.[V] | 1xHomocysteineIAA [K15] | 4,32935E-05 | 2612,22971 | Liver/ FAIMS/IAA |
| P07309 | P07309 [42-54] | [R].GSPAVDVAVKVFK.[K] | 1xHomocysteineMMTS [K10] | 0,000192903 | 1479,76982 | Liver/ No FAIMS/MMTS |
| P07309 | P07309 [69-90] | [K].TAESGELHGLTTDEKFVEGVYR.[V] | 1xHomocysteineMMTS [K15] | 0,000211896 | 2601,19596 | Liver/ FAIMS/MMTS |
| P07309 | P07309 [56-83] | [K].TSEGSWEPFASGKTAESGELHGLTTDEK.[F] | 1xHomocysteineMMTS [K13] | 4,49774E-05 | 3114,36667 | Liver/ FAIMS/MMTS |
| P07309 | P07309 [42-54] | [R].GSPAVDVAVKVFK.[K] | 1xHomocysteineMMTS [K10] | 0,000192903 | 1479,76982 | Liver/ FAIMS/MMTS |
| P07309 | P07309 [55-68] | [K].KTSEGSWEPFASGK.[T] | 1xHomocysteineMMTS [K1] | 0,00650587 | 1673,7298 | Liver/ FAIMS/MMTS |
| P07310 | P07310 [157-172] | [K].LSVEALNSLTGEFKGK.[Y] | 1xHomocysteineIAA [K14] | 4,32935E-05 | 1866,96297 | Liver/ FAIMS/IAA |
| P07310 | P07310 [33-43] | [K].VLTPDLYNKLR.[D] | 1xHomocysteineMMTS [K9] | 4,49774E-05 | 1494,78072 | Liver/ No FAIMS/MMTS |
| P07310 | P07310 [293-304] | [R].GGVHVKLANLSK.[H] | 1xHomocysteineMMTS [K6] | 0,00145127 | 1385,73919 | Liver/ No FAIMS/MMTS |
| P07310 | P07310 [153-170] | [R].AVEKLSVEALNSLTGEFK.[G] | 1xHomocysteineMMTS [K4] | 4,49774E-05 | 2098,05589 | Liver/ No FAIMS/MMTS |
| P07310 | P07310 [305-314] | [K].HPKFEEILTR.[L] | 1xHomocysteineMMTS [K3] | 0,00325546 | 1432,70755 | Liver/ No FAIMS/MMTS |
| P07310 | P07310 [10-25] | [K].FKLNYKPQEEYPDLSK.[H] | 1xHomocysteineMMTS [K2] | 4,49774E-05 | 2162,02967 | Liver/ No FAIMS/MMTS |
| P07310 | P07310 [106-116] | [K].HKTDLNHENLK.[G] | 1xHomocysteineMMTS [K2] | 4,49774E-05 | 1511,70934 | Liver/ No FAIMS/MMTS |
| P07310 | P07310 [157-172] | [K].LSVEALNSLTGEFKGK.[Y] | 1xHomocysteineMMTS [K14] | 4,49774E-05 | 1855,92923 | Liver/ No FAIMS/MMTS |
| P07310 | P07310 [366-381] | [K].KLEKGQSIDDMIPAQK.[-] | 1xHomocysteineMMTS [K4] | 0,000437433 | 1963,96496 | Liver/ FAIMS/MMTS |
| P07310 | P07310 [10-25] | [K].FKLNYKPQEEYPDLSK.[H] | 1xHomocysteineMMTS [K2] | 4,49774E-05 | 2162,02967 | Liver/ FAIMS/MMTS |
| P07310 | P07310 [157-172] | [K].LSVEALNSLTGEFKGK.[Y] | 1xHomocysteineMMTS [K14] | 4,49774E-05 | 1855,92923 | Liver/ FAIMS/MMTS |
| P07310 | P07310 [370-381] | [K].GQSIDDMIPAQK.[-] | 1xOxidation [M7]; 1xHomocysteineMMTS [K12] | 6,70238E-05 | 1481,6433 | Liver/ FAIMS/MMTS |
| P07310 | P07310 [366-381] | [K].KLEKGQSIDDMIPAQK.[-] | 1xOxidation [M11]; 1xHomocysteineMMTS [K4] | 0,000929608 | 1979,95988 | Liver/ FAIMS/MMTS |
| P07310 | P07310 [108-130] | [K].TDLNHENLKGGDDLDPNYVLSSR.[V] | 1xHomocysteineMMTS [K9] | 4,49774E-05 | 2735,23995 | Liver/ FAIMS/MMTS |
| P07310 | P07310 [33-43] | [K].VLTPDLYNKLR.[D] | 1xHomocysteineMMTS [K9] | 4,49774E-05 | 1494,78072 | Liver/ FAIMS/MMTS |
| P07310 | P07310 [293-304] | [R].GGVHVKLANLSK.[H] | 1xHomocysteineMMTS [K6] | 0,00145127 | 1385,73919 | Liver/ FAIMS/MMTS |
| P07310 | P07310 [153-170] | [R].AVEKLSVEALNSLTGEFK.[G] | 1xHomocysteineMMTS [K4] | 4,49774E-05 | 2098,05589 | Liver/ FAIMS/MMTS |
| P07310 | P07310 [12-25] | [K].LNYKPQEEYPDLSK.[H] | 1xHomocysteineMMTS [K4] | 4,49774E-05 | 1886,8663 | Liver/ FAIMS/MMTS |
| P07310 | P07310 [305-314] | [K].HPKFEEILTR.[L] | 1xHomocysteineMMTS [K3] | 0,00325546 | 1432,70755 | Liver/ FAIMS/MMTS |
| P07310 | P07310 [106-116] | [K].HKTDLNHENLK.[G] | 1xHomocysteineMMTS [K2] | 4,49774E-05 | 1511,70934 | Liver/ FAIMS/MMTS |
| P07310; P30275; Q6P8J7 | P07310 [237-247]; P30275 [271-281]; Q6P8J7 [271-281] | [R].VISMEKGGNMK.[ER] | 1xHomocysteineIAA [K6] | 0,000162511 | 1367,64799 | Brain/ No FAIMS/IAA |
| P07310; P30275; Q6P8J7 | P07310 [237-247]; P30275 [271-281]; Q6P8J7 [271-281] | [R].VISMEKGGNMK.[ER] | 1xHomocysteineIAA [K6] | 0,000162511 | 1367,64799 | Brain/ FAIMS/IAA |
| P07310; P30275; Q6P8J7 | P07310 [237-247]; P30275 [271-281]; Q6P8J7 [271-281] | [R].VISMEKGGNMK.[ER] | 1xHomocysteineMMTS [K6] | 4,49774E-05 | 1356,61425 | Liver/ FAIMS/MMTS |
| P07724 | P07724 [509-527] | [R].RPCFSALTVDETYVPKEFK.[A] | 1xCarbamidomethyl [C3]; 1xHomocysteineIAA [K16] | 4,32935E-05 | 2461,18903 | Liver/ No FAIMS/IAA |
| P07724 | P07724 [509-527] | [R].RPCFSALTVDETYVPKEFK.[A] | 1xCarbamidomethyl [C3]; 1xHomocysteineIAA [K16] | 4,32935E-05 | 2461,18903 | Liver/ FAIMS/IAA |
| P07724 | P07724 [509-527] | [R].RPCFSALTVDETYVPKEFK.[A] | 1xCarbamidomethyl [C3]; 1xHomocysteineIAA [K16] | 4,32935E-05 | 2461,18903 | Brain/ No FAIMS/IAA |
| P07724 | P07724 [509-527] | [R].RPCFSALTVDETYVPKEFK.[A] | 1xCarbamidomethyl [C3]; 1xHomocysteineIAA [K16] | 4,32935E-05 | 2461,18903 | Brain/ FAIMS/IAA |
| P07724 | P07724 [585-602] | [K].AADKDTCFSTEGPNLVTR.[C] | 1xMethylthio [C7]; 1xHomocysteineMMTS [K4] | 4,49774E-05 | 2133,90719 | Liver/ No FAIMS/MMTS |
| P07724 | P07724 [58-75] | [K].CSYDEHAKLVQEVTDFAK.[T] | 1xMethylthio [C1]; 1xHomocysteineMMTS [K8] | 4,49774E-05 | 2291,98036 | Liver/ No FAIMS/MMTS |
| P07724 | P07724 [550-562] | [K].QTALAELVKHKPK.[A] | 1xHomocysteineMMTS [K9] | 4,49774E-05 | 1625,88658 | Liver/ No FAIMS/MMTS |
| P07724 | P07724 [258-267] | [K].LATDLTKVNK.[E] | 1xHomocysteineMMTS [K7] | 4,49774E-05 | 1265,6592 | Liver/ No FAIMS/MMTS |
| P07724 | P07724 [435-452] | [R].YTQKAPQVSTPTLVEAAR.[N] | 1xHomocysteineMMTS [K4] | 4,49774E-05 | 2123,06237 | Liver/ No FAIMS/MMTS |
| P07724 | P07724 [559-569] | [K].HKPKATAEQLK.[T] | 1xHomocysteineMMTS [K4] | 0,000133296 | 1413,7341 | Liver/ No FAIMS/MMTS |
| P07724 | P07724 [25-34] | [R].EAHKSEIAHR.[Y] | 1xHomocysteineMMTS [K4] | 0,00074635 | 1340,6198 | Liver/ No FAIMS/MMTS |
| P07724 | P07724 [234-242] | [R].AFKAWAVAR.[L] | 1xHomocysteineMMTS [K3] | 0,00138774 | 1182,59107 | Liver/ No FAIMS/MMTS |
| P07724 | P07724 [211-219] | [K].EKALVSSVR.[Q] | 1xHomocysteineMMTS [K2] | 0,00115861 | 1151,59113 | Liver/ No FAIMS/MMTS |
| P07724 | P07724 [243-264] | [R].LSQTFPNADFAEITKLATDLTK.[V] | 1xHomocysteineMMTS [K15] | 4,49774E-05 | 2587,27824 | Liver/ No FAIMS/MMTS |
| P07724 | P07724 [549-562] | [K].KQTALAELVKHKPK.[A] | 1xHomocysteineMMTS [K10] | 6,70238E-05 | 1753,98154 | Liver/ No FAIMS/MMTS |
| P07724 | P07724 [376-383] | [K].KYEATLEK.[C] | 1xHomocysteineMMTS [K1] | 0,000211896 | 1144,53769 | Liver/ No FAIMS/MMTS |
| P07724 | P07724 [549-558] | [K].KQTALAELVK.[H] | 1xHomocysteineMMTS [K1] | 4,49774E-05 | 1263,67994 | Liver/ No FAIMS/MMTS |
| P07724 | P07724 [76-97] | [K].TCVADESAANCDKSLHTLFGDK.[L] | 2xMethylthio [C2; C11]; 1xHomocysteineMMTS [K13] | 4,49774E-05 | 2580,03657 | Liver/ FAIMS/MMTS |
| P07724 | P07724 [410-421] | [K].NLVKTNCDLYEK.[L] | 1xMethylthio [C7]; 1xHomocysteineMMTS [K4] | 0,0004709 | 1648,72017 | Liver/ FAIMS/MMTS |
| P07724 | P07724 [585-602] | [K].AADKDTCFSTEGPNLVTR.[C] | 1xMethylthio [C7]; 1xHomocysteineMMTS [K4] | 4,49774E-05 | 2133,90719 | Liver/ FAIMS/MMTS |
| P07724 | P07724 [89-105] | [K].SLHTLFGDKLCAIPNLR.[E] | 1xMethylthio [C11]; 1xHomocysteineMMTS [K9] | 0,000250158 | 2107,03194 | Liver/ FAIMS/MMTS |
| P07724 | P07724 [528-545] | [K].AETFTFHSDICTLPEKEK.[Q] | 1xMethylthio [C11]; 1xHomocysteineMMTS [K16] | 0,000361246 | 2305,00076 | Liver/ FAIMS/MMTS |
| P07724 | P07724 [58-75] | [K].CSYDEHAKLVQEVTDFAK.[T] | 1xMethylthio [C1]; 1xHomocysteineMMTS [K8] | 4,49774E-05 | 2291,98036 | Liver/ FAIMS/MMTS |
| P07724 | P07724 [550-562] | [K].QTALAELVKHKPK.[A] | 1xHomocysteineMMTS [K9] | 4,49774E-05 | 1625,88658 | Liver/ FAIMS/MMTS |
| P07724 | P07724 [258-267] | [K].LATDLTKVNK.[E] | 1xHomocysteineMMTS [K7] | 4,49774E-05 | 1265,6592 | Liver/ FAIMS/MMTS |
| P07724 | P07724 [435-452] | [R].YTQKAPQVSTPTLVEAAR.[N] | 1xHomocysteineMMTS [K4] | 4,49774E-05 | 2123,06237 | Liver/ FAIMS/MMTS |
| P07724 | P07724 [234-242] | [R].AFKAWAVAR.[L] | 1xHomocysteineMMTS [K3] | 0,00138774 | 1182,59107 | Liver/ FAIMS/MMTS |
| P07724 | P07724 [211-219] | [K].EKALVSSVR.[Q] | 1xHomocysteineMMTS [K2] | 0,00115861 | 1151,59113 | Liver/ FAIMS/MMTS |
| P07724 | P07724 [549-558] | [K].KQTALAELVK.[H] | 1xHomocysteineMMTS [K1] | 4,49774E-05 | 1263,67994 | Liver/ FAIMS/MMTS |
| P07724 | P07724 [376-383] | [K].KYEATLEK.[C] | 1xHomocysteineMMTS [K1] | 0,000211896 | 1144,53769 | Liver/ FAIMS/MMTS |

| | | | | | | |
|---|---|---|---|---|---|---|
| P07724 | P07724 [549-558] | [K].KQTALAELVK.[H] | 1xHomocysteineMMTS [K1] | 4,49774E-05 | 1263,67994 | Brain/ No FAIMS/MMTS |
| P07724 | P07724 [435-452] | [R].YTQKAPQVSTPTLVEAAR.[N] | 1xHomocysteineMMTS [K4] | 4,49774E-05 | 2123,06237 | Brain/ FAIMS/MMTS |
| P07724 | P07724 [211-219] | [K].EKALVSSVR.[Q] | 1xHomocysteineMMTS [K2] | 0,00115861 | 1151,59113 | Brain/ FAIMS/MMTS |
| P07724 | P07724 [549-558] | [K].KQTALAELVK.[H] | 1xHomocysteineMMTS [K1] | 4,49774E-05 | 1263,67994 | Brain/ FAIMS/MMTS |
| P07758 | P07758 [214-235] | [K].KPFDPENTEEAEFHVDESTTVK.[V] | 1xHomocysteineIAA [K1] | 4,32935E-05 | 2723,21412 | Liver/ FAIMS/IAA |
| P07758 | P07758 [278-292] | [K].MQHLEQTLSKELISK.[F] | 1xHomocysteineMMTS [K10]<br>1xOxidation [M13]; | 4,49774E-05 | 1947,97005 | Liver/ No FAIMS/MMTS |
| P07758 | P07758 [309-328] | [R].LSISGEYNLKTLMSPLGITR.[I] | 1xHomocysteineMMTS [K10] | 8,93346E-05 | 2372,20224 | Liver/ FAIMS/MMTS |
| P07758 | P07758 [309-328] | [R].LSISGEYNLKTLMSPLGITR.[I] | 1xHomocysteineMMTS [K10]<br>1xOxidation [M1]; | 4,49774E-05 | 2356,20732 | Liver/ FAIMS/MMTS |
| P07758 | P07758 [278-292] | [K].MQHLEQTLSKELISK.[F] | 1xHomocysteineMMTS [K10] | 0,0046895 | 1963,96497 | Liver/ FAIMS/MMTS |
| P07758 | P07758 [278-292] | [K].MQHLEQTLSKELISK.[F] | 1xHomocysteineMMTS [K10] | 4,49774E-05 | 1947,97005 | Liver/ FAIMS/MMTS |
| P07758 | P07758 [214-235] | [K].KPFDPENTEEAEFHVDESTTVK.[V] | 1xHomocysteineMMTS [K1] | 4,49774E-05 | 2712,18037 | Liver/ FAIMS/MMTS |
| P07758;<br>P22599;<br>Q00898;<br>Q00897 | P07758 [144-153];<br>P22599 [144-153];<br>Q00898 [144-153];<br>Q00897 [144-153] | [K].LVEKFLEEAK.[N] | 1xHomocysteineIAA [K4] | 4,32935E-05 | 1379,72391 | Liver/ FAIMS/IAA |
| P07758;<br>P22599;<br>Q00898;<br>Q00897 | P07758 [144-153];<br>P22599 [144-153];<br>Q00898 [144-153];<br>Q00897 [144-153] | [K].LVEKFLEEAK.[N] | 1xHomocysteineIAA [K4] | 4,32935E-05 | 1379,72391 | Brain/ FAIMS/IAA |
| P07758;<br>P22599;<br>Q00898;<br>Q00897 | P07758 [174-186];<br>P22599 [174-186];<br>Q00898 [174-186];<br>Q00897 [174-186] | [K].VINDFVEKGTQGK.[I] | 1xHomocysteineMMTS [K8] | 4,49774E-05 | 1597,77127 | Liver/ No FAIMS/MMTS |
| P07758;<br>P22599;<br>Q00898;<br>Q00897 | P07758 [144-153];<br>P22599 [144-153];<br>Q00898 [144-153];<br>Q00897 [144-153] | [K].LVEKFLEEAK.[N] | 1xHomocysteineMMTS [K4] | 4,49774E-05 | 1368,69017 | Liver/ No FAIMS/MMTS |
| P07758;<br>P22599;<br>Q00898;<br>Q00897 | P07758 [173-181];<br>P22599 [173-181];<br>Q00898 [173-181];<br>Q00897 [173-181] | [K].KVINDFVEK.[G] | 1xHomocysteineMMTS [K1] | 4,49774E-05 | 1254,62209 | Liver/ No FAIMS/MMTS |
| P07758;<br>P22599;<br>Q00898;<br>Q00897 | P07758 [174-186];<br>P22599 [174-186];<br>Q00898 [174-186];<br>Q00897 [174-186] | [K].VINDFVEKGTQGK.[I] | 1xHomocysteineMMTS [K8] | 4,49774E-05 | 1597,77127 | Liver/ FAIMS/MMTS |
| P07758;<br>P22599;<br>Q00898;<br>Q00897 | P07758 [144-153];<br>P22599 [144-153];<br>Q00898 [144-153];<br>Q00897 [144-153] | [K].LVEKFLEEAK.[N] | 1xHomocysteineMMTS [K4] | 4,49774E-05 | 1368,69017 | Liver/ FAIMS/MMTS |
| P07758;<br>P22599;<br>Q00898;<br>Q00897 | P07758 [154-173];<br>P22599 [154-173];<br>Q00898 [154-173];<br>Q00897 [154-173] | [K].NHYQAEVFSVNFAESEEAKK.[V] | 1xHomocysteineMMTS [K19] | 4,49774E-05 | 2490,10642 | Liver/ FAIMS/MMTS |
| P07758;<br>P22599;<br>Q00898;<br>Q00897 | P07758 [407-413];<br>P22599 [407-413];<br>Q00898 [407-413];<br>Q00897 [407-413] | [K].VVDPTHK.[-] | 1xHomocysteineMMTS [K7] | 0,00338636 | 958,44848 | Liver/ FAIMS/MMTS |
| P07758;<br>P22599;<br>Q00898;<br>Q00897 | P07758 [173-181];<br>P22599 [173-181];<br>Q00898 [173-181];<br>Q00897 [173-181] | [K].KVINDFVEK.[G] | 1xHomocysteineMMTS [K1] | 4,49774E-05 | 1254,62209 | Liver/ FAIMS/MMTS |
| P07758;<br>Q00898 | P07758 [288-297];<br>Q00898 [288-297] | [K].ELISKFLLNR.[R] | 1xHomocysteineIAA [K5] | 0,000876609 | 1406,78243 | Liver/ No FAIMS/IAA |
| P07758;<br>Q00898 | P07758 [288-297];<br>Q00898 [288-297] | [K].ELISKFLLNR.[R] | 1xHomocysteineIAA [K5] | 0,000876609 | 1406,78243 | Liver/ FAIMS/IAA |
| P07758;<br>Q00898 | P07758 [288-297];<br>Q00898 [288-297] | [K].ELISKFLLNR.[R] | 1xHomocysteineMMTS [K5] | 0,000192903 | 1395,74869 | Liver/ No FAIMS/MMTS |
| P07758;<br>Q00898 | P07758 [288-297];<br>Q00898 [288-297] | [K].ELISKFLLNR.[R] | 1xHomocysteineMMTS [K5] | 0,000192903 | 1395,74869 | Liver/ FAIMS/MMTS |
| P07759 | P07759 [92-103] | [K].GKTMEEILEGLK.[F] | 1xHomocysteineIAA [K2] | 4,32935E-05 | 1521,76513 | Liver/ FAIMS/IAA |
| P07759 | P07759 [92-103] | [K].GKTMEEILEGLK.[F] | 1xHomocysteineIAA [K2] | 4,32935E-05 | 1521,76513 | Brain/ FAIMS/IAA |
| P07759 | P07759 [92-103] | [K].GKTMEEILEGLK.[F] | 1xHomocysteineMMTS [K2] | 4,49774E-05 | 1510,73138 | Liver/ No FAIMS/MMTS |
| P07759 | P07759 [351-360] | [K].KLSVSQVVHK.[A] | 1xHomocysteineMMTS [K1] | 4,49774E-05 | 1287,69117 | Liver/ No FAIMS/MMTS |
| P07759 | P07759 [92-103] | [K].GKTMEEILEGLK.[F] | 1xHomocysteineMMTS [K2] | 4,49774E-05 | 1510,73138 | Liver/ FAIMS/MMTS |
| P07759 | P07759 [351-360] | [K].KLSVSQVVHK.[A] | 1xHomocysteineMMTS [K1] | 4,49774E-05 | 1287,69117 | Liver/ FAIMS/MMTS |
| P07759;<br>Q03734 | P07759 [145-157];<br>Q03734 [144-156] | [K].DLQILAEFHEKTR.[A] | 1xHomocysteineIAA [K11] | 4,32935E-05 | 1773,89523 | Liver/ FAIMS/IAA |
| P07759;<br>Q03734 | P07759 [145-157];<br>Q03734 [144-156] | [K].DLQILAEFHEKTR.[A] | 1xHomocysteineMMTS [K11] | 4,49774E-05 | 1762,86149 | Liver/ No FAIMS/MMTS |
| P07759;<br>Q03734 | P07759 [218-236];<br>Q03734 [217-235] | [K].ISFDPQDTFESEFYLDEKR.[S] | 1xHomocysteineMMTS [K18] | 4,49774E-05 | 2529,09485 | Liver/ FAIMS/MMTS |
| P07759;<br>Q03734 | P07759 [145-157];<br>Q03734 [144-156] | [K].DLQILAEFHEKTR.[A] | 1xHomocysteineMMTS [K11] | 4,49774E-05 | 1762,86149 | Liver/ FAIMS/MMTS |
| P07901 | P07901 [633-648] | [K].KHLEINPDHSIIETLR.[Q] | 1xHomocysteineIAA [K1] | 4,32935E-05 | 2089,08588 | Brain/ No FAIMS/IAA |
| P07901 | P07901 [633-648] | [K].KHLEINPDHSIIETLR.[Q] | 1xHomocysteineIAA [K1] | 4,32935E-05 | 2089,08588 | Brain/ FAIMS/IAA |
| P07901 | P07901 [536-547] | [K].EFEGKTLVSVTK.[E] | 1xHomocysteineMMTS [K5] | 0,00459586 | 1500,74366 | Liver/ FAIMS/MMTS |
| P07901 | P07901 [445-457] | [K].NIKLGIHEDSQNR.[K] | 1xHomocysteineMMTS [K3] | 4,49774E-05 | 1686,80503 | Liver/ FAIMS/MMTS |
| P07901 | P07901 [47-60] | [R].ELISNSSDALDKIR.[Y] | 1xHomocysteineMMTS [K12] | 0,00309423 | 1723,83533 | Liver/ FAIMS/MMTS |
| P07901 | P07901 [633-648] | [K].KHLEINPDHSIIETLR.[Q] | 1xHomocysteineMMTS [K1] | 4,49774E-05 | 2078,05214 | Liver/ FAIMS/MMTS |
| P07901 | P07901 [536-547] | [K].EFEGKTLVSVTK.[E] | 1xHomocysteineMMTS [K5] | 0,00459586 | 1500,74366 | Brain/ FAIMS/MMTS |

| | | | | | | |
|---|---|---|---|---|---|---|
| P07901 | P07901 [445-457] | [K].NIKLGIHEDSQNR.[K] | 1xHomocysteineMMTS [K3] | 4,49774E-05 | 1686,80503 | Brain/ FAIMS/MMTS |
| P07901 | P07901 [633-648] | [K].KHLEINPDHSIIETLR.[Q] | 1xHomocysteineMMTS [K1] | 4,49774E-05 | 2078,05214 | Brain/ FAIMS/MMTS |
| P07901; P11499 | P07901 [183-191]; P11499 [178-186] | [R].GTKVILHLK.[E] | 1xHomocysteineIAA [K3] | 4,32935E-05 | 1182,70272 | Liver/ No FAIMS/IAA |
| P07901; P11499 | P07901 [281-293]; P11499 [272-284] | [K].IKEKYIDQEELNK.[T] | 1xHomocysteineIAA [K] | 4,32935E-05 | 1823,92078 | Liver/ FAIMS/IAA |
| P07901; P11499 | P07901 [183-191]; P11499 [178-186] | [R].GTKVILHLK.[E] | 1xHomocysteineIAA [K3] | 4,32935E-05 | 1182,70272 | Brain/ No FAIMS/IAA |
| P07901; P11499 | P07901 [183-191]; P11499 [178-186] | [R].GTKVILHLK.[E] | 1xHomocysteineIAA [K3] | 4,32935E-05 | 1182,70272 | Brain/ FAIMS/IAA |
| P07901; P11499 | P07901 [61-74]; P11499 [56-69] | [R].YESLTDPSKLDSGK.[E] | 1xHomocysteineMMTS [K9] | 4,49774E-05 | 1702,76625 | Liver/ No FAIMS/MMTS |
| P07901; P11499 | P07901 [186-201]; P11499 [181-196] | [K].VILHLKEDQTEYLEER.[R] | 1xHomocysteineMMTS [K6] | 4,49774E-05 | 2178,05695 | Liver/ No FAIMS/MMTS |
| P07901; P11499 | P07901 [294-300]; P11499 [285-291] | [K].TKPIWTR.[N] | 1xHomocysteineMMTS [K2] | 0,00166208 | 1064,53797 | Liver/ No FAIMS/MMTS |
| P07901; P11499 | P07901 [101-116]; P11499 [96-111] | [K].ADLINNLGTIAKSGTK.[A] | 1xHomocysteineMMTS [K12] | 0,000530332 | 1778,91392 | Liver/ No FAIMS/MMTS |
| P07901; P11499 | P07901 [402-411]; P11499 [393-402] | [R].EMLQQSKILK.[V] | 1xHomocysteineMMTS [K7] 1xOxidation [M2]; | 0,00367611 | 1380,70477 | Liver/ FAIMS/MMTS |
| P07901; P11499 | P07901 [614-621]; P11499 [605-612] | [R].IMKAQALR.[D] | 1xHomocysteineMMTS [K3] | 0,00830327 | 1109,5628 | Liver/ FAIMS/MMTS |
| P07901; P11499 | P07901 [285-300]; P11499 [276-291] | [K].YIDQEELNKTKPIWTR.[N] | 1xHomocysteineMMTS [K9] | 4,49774E-05 | 2197,07802 | Liver/ FAIMS/MMTS |
| P07901; P11499 | P07901 [61-74]; P11499 [56-69] | [R].YESLTDPSKLDSGK.[E] | 1xHomocysteineMMTS [K9] | 4,49774E-05 | 1702,76625 | Liver/ FAIMS/MMTS |
| P07901; P11499 | P07901 [186-201]; P11499 [181-196] | [K].VILHLKEDQTEYLEER.[R] | 1xHomocysteineMMTS [K6] | 4,49774E-05 | 2178,05695 | Liver/ FAIMS/MMTS |
| P07901; P11499 | P07901 [183-191]; P11499 [178-186] | [R].GTKVILHLK.[E] | 1xHomocysteineMMTS [K3] | 0,00107839 | 1171,66898 | Liver/ FAIMS/MMTS |
| P07901; P11499 | P07901 [614-621]; P11499 [605-612] | [R].IMKAQALR.[D] | 1xHomocysteineMMTS [K3] | 0,00322925 | 1093,56789 | Liver/ FAIMS/MMTS |
| P07901; P11499 | P07901 [281-293]; P11499 [272-284] | [K].IKEKYIDQEELNK.[T] | 1xHomocysteineMMTS [K2] | 0,00312202 | 1812,88703 | Liver/ FAIMS/MMTS |
| P07901; P11499 | P07901 [548-560]; P11499 [539-551] | [K].EGLELPEDEEEKK.[K] | 1xHomocysteineMMTS [K12] | 0,000211896 | 1707,74518 | Liver/ FAIMS/MMTS |
| P07901; P11499 | P07901 [61-74]; P11499 [56-69] | [R].YESLTDPSKLDSGK.[E] | 1xHomocysteineMMTS [K9] | 4,49774E-05 | 1702,76625 | Brain/ FAIMS/MMTS |
| P07901; P11499 | P07901 [101-116]; P11499 [96-111] | [K].ADLINNLGTIAKSGTK.[A] | 1xHomocysteineMMTS [K12] | 0,000530332 | 1778,91392 | Brain/ FAIMS/MMTS |
| P08003 | P08003 [149-158] | [R].TQEEIVAKVR.[E] | 1xHomocysteineIAA [K8] | 9,61721E-05 | 1346,70966 | Liver/ FAIMS/IAA |
| P08003 | P08003 [623-631] | [K].FIDEHATKR.[S] | 1xHomocysteineMMTS [K8] | 0,0033721 | 1279,59219 | Liver/ No FAIMS/MMTS |
| P08003 | P08003 [149-158] | [R].TQEEIVAKVR.[E] | 1xHomocysteineMMTS [K8] | 0,00501021 | 1335,67592 | Liver/ FAIMS/MMTS |
| P08003 | P08003 [616-630] | [R].DLEHLSKFIDEHATK.[R] | 1xHomocysteineMMTS [K7] | 0,00467787 | 1945,91464 | Liver/ FAIMS/MMTS |
| P08003 | P08003 [228-239] | [K].VDATEQTDLAKR.[F] | 1xHomocysteineMMTS [K11] | 0,00175939 | 1509,70359 | Liver/ FAIMS/MMTS |
| P08113 | P08113 [725-738] | [R].SGYLLPDTKAYGDR.[I] | 1xHomocysteineIAA [K9] | 4,32935E-05 | 1729,8214 | Liver/ FAIMS/IAA |
| P08113 | P08113 [725-738] | [R].SGYLLPDTKAYGDR.[I] | 1xHomocysteineIAA [K9] | 4,32935E-05 | 1729,8214 | Brain/ FAIMS/IAA |
| P08113 | P08113 [157-168] | [R].EELVKNLGTIAK.[S] | 1xHomocysteineMMTS [K5] | 0,0004709 | 1477,7753 | Liver/ No FAIMS/MMTS |
| P08113 | P08113 [609-623] | [R].EATEKEFEPLLNWMK.[D] | 1xHomocysteineMMTS [K5] | 0,000724331 | 2027,92752 | Liver/ No FAIMS/MMTS |
| P08113 | P08113 [661-671] | [R].IMKAQAYQTGK.[D] | 1xHomocysteineMMTS [K3] | 0,000173718 | 1401,66872 | Liver/ No FAIMS/MMTS |
| P08113 | P08113 [672-683] | [K].DISTNYYASQKK.[T] | 1xHomocysteineMMTS [K11] | 4,49774E-05 | 1580,70834 | Liver/ No FAIMS/MMTS |
| P08113 | P08113 [547-557] | [R].KEAESSPFVER.[L] | 1xHomocysteineMMTS [K1] | 0,00386874 | 1441,64501 | Liver/ No FAIMS/MMTS |
| P08113 | P08113 [157-168] | [R].EELVKNLGTIAK.[S] | 1xHomocysteineMMTS [K5] | 0,0004709 | 1477,7753 | Liver/ FAIMS/MMTS |
| P08113 | P08113 [609-623] | [R].EATEKEFEPLLNWMK.[D] | 1xHomocysteineMMTS [K5] | 0,000724331 | 2027,92752 | Liver/ FAIMS/MMTS |
| P08113 | P08113 [535-546] | [K].QDKIYFMAGSSR.[K] | 1xHomocysteineMMTS [K3] 1xOxidation [M2]; | 4,49774E-05 | 1565,69092 | Liver/ FAIMS/MMTS |
| P08113 | P08113 [661-671] | [R].IMKAQAYQTGK.[D] | 1xHomocysteineMMTS [K3] | 0,00115861 | 1417,66364 | Liver/ FAIMS/MMTS |
| P08113 | P08113 [725-738] | [R].SGYLLPDTKAYGDR.[I] | 1xHomocysteineMMTS [K9] | 4,49774E-05 | 1718,78765 | Liver/ FAIMS/MMTS |
| P08113 | P08113 [664-682] | [K].AQAYQTGKDISTNYYASQK.[K] | 1xHomocysteineMMTS [K8] | 4,49774E-05 | 2300,03219 | Liver/ FAIMS/MMTS |
| P08113 | P08113 [162-177] | [K].NLGTIAKSGTSEFLNK.[M] | 1xHomocysteineMMTS [K7] | 4,49774E-05 | 1842,90883 | Liver/ FAIMS/MMTS |
| P08113 | P08113 [480-493] | [K].YNDTFWKEFGTNIK.[L] | 1xHomocysteineMMTS [K7] | 0,000530332 | 1925,85607 | Liver/ FAIMS/MMTS |
| P08113 | P08113 [487-503] | [K].EFGTNIKLGVIEDHSNR.[T] | 1xHomocysteineMMTS [K7] | 4,49774E-05 | 2091,99502 | Liver/ FAIMS/MMTS |
| P08113 | P08113 [429-448] | [K].YLNFVKGVVDSDDLPLNVSR.[E] | 1xHomocysteineMMTS [K6] | 4,49774E-05 | 2413,18903 | Liver/ FAIMS/MMTS |
| P08113 | P08113 [588-597] | [R].FQNVAKEGVK.[F] | 1xHomocysteineMMTS [K6] | 0,00207528 | 1282,62824 | Liver/ FAIMS/MMTS |
| P08113 | P08113 [73-84] | [K].SEKFAFQAEVNR.[M] | 1xHomocysteineMMTS [K3] | 0,00074635 | 1588,72466 | Liver/ FAIMS/MMTS |
| P08113 | P08113 [661-671] | [R].IMKAQAYQTGK.[D] | 1xHomocysteineMMTS [K3] | 0,000173718 | 1401,66872 | Liver/ FAIMS/MMTS |
| P08113 | P08113 [416-434] | [R].VFITDDFHDMMPKYLNFVK.[G] | 1xHomocysteineMMTS [K13] | 0,000764279 | 2523,15793 | Liver/ FAIMS/MMTS |
| P08113 | P08113 [672-683] | [K].DISTNYYASQKK.[T] | 1xHomocysteineMMTS [K11] | 4,49774E-05 | 1580,70834 | Liver/ FAIMS/MMTS |
| P08113 | P08113 [396-405] | [R].GLFDEYGSKK.[S] | 1xHomocysteineMMTS [K] | 0,00288111 | 1306,58062 | Liver/ FAIMS/MMTS |
| P08113 | P08113 [609-623] | [R].EATEKEFEPLLNWMK.[D] | 1xHomocysteineMMTS [K5] | 0,000724331 | 2027,92752 | Brain/ FAIMS/MMTS |
| P08226 | P08226 [86-100] | [K].KELEEQLGPVAEETR.[A] | 1xHomocysteineIAA [K1] | 4,32935E-05 | 1901,92732 | Liver/ FAIMS/IAA |
| P08226 | P08226 [86-100] | [K].KELEEQLGPVAEETR.[A] | 1xHomocysteineIAA [K1] | 4,32935E-05 | 1901,92732 | Brain/ FAIMS/IAA |
| P08226 | P08226 [103-113] | [R].LGKEVQAAQAR.[L] | 1xHomocysteineMMTS [K3] 1xOxidation [M3]; | 0,00074635 | 1333,6715 | Liver/ FAIMS/MMTS |
| P08226 | P08226 [251-261] | [R].SKMEEQTQQIR.[L] | 1xHomocysteineMMTS [K2] | 0,00618488 | 1556,68656 | Liver/ FAIMS/MMTS |
| P08226 | P08226 [251-261] | [R].SKMEEQTQQIR.[L] | 1xHomocysteineMMTS [K2] | 0,0031519 | 1540,69164 | Liver/ FAIMS/MMTS |
| P08228 | P08228 [124-144] | [K].QDDLGKGGNEESTKTGNAGSR.[L] | 1xHomocysteineIAA [K14] | 4,32935E-05 | 2295,02659 | Liver/ FAIMS/IAA |
| P08228 | P08228 [130-144] | [K].GGNEESTKTGNAGSR.[L] | 1xHomocysteineMMTS [K8] | 4,49774E-05 | 1627,67989 | Liver/ No FAIMS/MMTS |
| P08228 | P08228 [117-129] | [R].TMVVHEKQDDLGK.[G] | 1xHomocysteineMMTS [K7] | 4,49774E-05 | 1662,76481 | Liver/ No FAIMS/MMTS |
| P08228 | P08228 [117-137] | [R].TMVVHEKQDDLGKGGNEESTK.[T] | 1xHomocysteineMMTS [K7] | 4,49774E-05 | 2465,11052 | Liver/ No FAIMS/MMTS |
| P08228 | P08228 [124-137] | [K].QDDLGKGGNEESTK.[T] | 1xHomocysteineMMTS [K6] | 4,49774E-05 | 1640,68906 | Liver/ No FAIMS/MMTS |
| P08228 | P08228 [81-103] | [R].HVGDLGNVTAGKDGVANVSIEDR.[V] | 1xHomocysteineMMTS [K12] | 4,49774E-05 | 2486,17623 | Liver/ No FAIMS/MMTS |

| | | | | | | |
|---|---|---|---|---|---|---|
| P08228 | P08228 [117-129] | [R].TMVVHEKQDDLGK.[G] | 1xHomocysteineMMTS [K7] | 4,49774E-05 | 1662,76481 | Liver/ FAIMS/MMTS |
| P08228 | P08228 [117-137] | [R].TMVVHEKQDDLGKGGNEESTK.[T] | 1xHomocysteineMMTS [K7] | 4,49774E-05 | 2465,11052 | Liver/ FAIMS/MMTS |
| P08228 | P08228 [124-137] | [K].QDDLGKGGNEESTK.[T] | 1xHomocysteineMMTS [K6] | 4,49774E-05 | 1640,68906 | Liver/ FAIMS/MMTS |
| P08228 | P08228 [81-103] | [R].HVGDLGNVTAGKDGVANVSIEDR.[V] | 1xHomocysteineMMTS [K12] | 4,49774E-05 | 2486,17623 | Liver/ FAIMS/MMTS |
| P08228 | P08228 [117-129] | [R].TMVVHEKQDDLGK.[G] | 1xOxidation [M2];<br>1xHomocysteineMMTS [K7] | 0,00145127 | 1678,75972 | Liver/ FAIMS/MMTS |
| P08228 | P08228 [117-129] | [R].TMVVHEKQDDLGK.[G] | 1xHomocysteineMMTS [K7] | 4,49774E-05 | 1662,76481 | Brain/ FAIMS/MMTS |
| P08228 | P08228 [117-137] | [R].TMVVHEKQDDLGKGGNEESTK.[T] | 1xHomocysteineMMTS [K7] | 4,49774E-05 | 2465,11052 | Brain/ FAIMS/MMTS |
| P08228 | P08228 [124-137] | [K].QDDLGKGGNEESTK.[T] | 1xHomocysteineMMTS [K6] | 4,49774E-05 | 1640,68906 | Brain/ FAIMS/MMTS |
| P08249 | P08249 [158-176] | [K].HGVYNPNKIFGVTTLDIVR.[A] | 1xHomocysteineIAA [K8] | 4,32935E-05 | 2317,21215 | Liver/ FAIMS/IAA |
| P08249 | P08249 [329-338] | [K].KGEDFVKNMK.[-] | 1xHomocysteineIAA [K7] | 4,32935E-05 | 1369,66027 | Brain/ No FAIMS/IAA |
| P08249 | P08249 [329-338] | [K].KGEDFVKNMK.[-] | 1xHomocysteineIAA [K7] | 4,32935E-05 | 1369,66027 | Brain/ FAIMS/IAA |
| P08249 | P08249 [308-324] | [K].ITPFEEKMIAEAIPELK.[A] | 1xOxidation [M8];<br>1xHomocysteineMMTS [K7] | 0,000599431 | 2138,0582 | Liver/ No FAIMS/MMTS |
| P08249 | P08249 [177-191] | [R].ANTFVAELKGLDPAR.[V] | 1xHomocysteineMMTS [K9] | 4,49774E-05 | 1764,87714 | Liver/ No FAIMS/MMTS |
| P08249 | P08249 [158-176] | [K].HGVYNPNKIFGVTTLDIVR.[A] | 1xHomocysteineMMTS [K8] | 4,49774E-05 | 2306,1784 | Liver/ No FAIMS/MMTS |
| P08249 | P08249 [298-307] | [K].GLEKNLGIGK.[I] | 1xHomocysteineMMTS [K4] | 6,70238E-05 | 1191,62243 | Liver/ No FAIMS/MMTS |
| P08249 | P08249 [315-328] | [K].MIAEAIPELKASIK.[K] | 1xHomocysteineMMTS [K10] | 4,49774E-05 | 1676,87838 | Liver/ No FAIMS/MMTS |
| P08249 | P08249 [177-191] | [R].ANTFVAELKGLDPAR.[V] | 1xHomocysteineMMTS [K9] | 4,49774E-05 | 1764,87714 | Liver/ FAIMS/MMTS |
| P08249 | P08249 [302-314] | [K].NLGIGKITPFEEK.[M] | 1xHomocysteineMMTS [K6] | 4,49774E-05 | 1608,81241 | Liver/ FAIMS/MMTS |
| P08249 | P08249 [230-241] | [R].IQEAGTEVVKAK.[A] | 1xHomocysteineMMTS [K10] | 4,49774E-05 | 1435,72835 | Liver/ FAIMS/MMTS |
| P08249 | P08249 [315-328] | [K].MIAEAIPELKASIK.[K] | 1xHomocysteineMMTS [K10] | 4,49774E-05 | 1676,87838 | Liver/ FAIMS/MMTS |
| P08249 | P08249 [308-324] | [K].ITPFEEKMIAEAIPELK.[A] | 1xOxidation [M8];<br>1xHomocysteineMMTS [K7] | 0,000599431 | 2138,0582 | Liver/ FAIMS/MMTS |
| P08249 | P08249 [240-257] | [K].AKAGAGSATLSMAYAGAR.[F] | 1xOxidation [M12];<br>1xHomocysteineMMTS [K2] | 4,49774E-05 | 1832,84518 | Liver/ FAIMS/MMTS |
| P08249 | P08249 [315-328] | [K].MIAEAIPELKASIK.[K] | 1xOxidation [M1];<br>1xHomocysteineMMTS [K10] | 0,000152766 | 1692,8733 | Liver/ FAIMS/MMTS |
| P08249 | P08249 [158-176] | [K].HGVYNPNKIFGVTTLDIVR.[A] | 1xHomocysteineMMTS [K8] | 4,49774E-05 | 2306,1784 | Liver/ FAIMS/MMTS |
| P08249 | P08249 [297-307] | [K].KGLEKNLGIGK.[I] | 1xHomocysteineMMTS [K5] | 0,00869467 | 1319,71739 | Liver/ FAIMS/MMTS |
| P08249 | P08249 [177-191] | [R].ANTFVAELKGLDPAR.[V] | 1xHomocysteineMMTS [K9] | 4,49774E-05 | 1764,87714 | Brain/ No FAIMS/MMTS |
| P08249 | P08249 [158-176] | [K].HGVYNPNKIFGVTTLDIVR.[A] | 1xHomocysteineMMTS [K8] | 4,49774E-05 | 2306,1784 | Brain/ No FAIMS/MMTS |
| P08249 | P08249 [308-324] | [K].ITPFEEKMIAEAIPELK.[A] | 1xHomocysteineMMTS [K7] | 4,49774E-05 | 2122,06328 | Brain/ No FAIMS/MMTS |
| P08249 | P08249 [302-314] | [K].NLGIGKITPFEEK.[M] | 1xHomocysteineMMTS [K6] | 4,49774E-05 | 1608,81241 | Brain/ No FAIMS/MMTS |
| P08249 | P08249 [298-307] | [K].GLEKNLGIGK.[I] | 1xHomocysteineMMTS [K4] | 6,70238E-05 | 1191,62243 | Brain/ No FAIMS/MMTS |
| P08249 | P08249 [230-241] | [R].IQEAGTEVVKAK.[A] | 1xHomocysteineMMTS [K10] | 4,49774E-05 | 1435,72835 | Brain/ No FAIMS/MMTS |
| P08249 | P08249 [315-328] | [K].MIAEAIPELKASIK.[K] | 1xHomocysteineMMTS [K10] | 4,49774E-05 | 1676,87838 | Brain/ No FAIMS/MMTS |
| P08249 | P08249 [177-191] | [R].ANTFVAELKGLDPAR.[V] | 1xHomocysteineMMTS [K9] | 4,49774E-05 | 1764,87714 | Brain/ FAIMS/MMTS |
| P08249 | P08249 [158-176] | [K].HGVYNPNKIFGVTTLDIVR.[A] | 1xHomocysteineMMTS [K8] | 4,49774E-05 | 2306,1784 | Brain/ FAIMS/MMTS |
| P08249 | P08249 [302-314] | [K].NLGIGKITPFEEK.[M] | 1xHomocysteineMMTS [K6] | 4,49774E-05 | 1608,81241 | Brain/ FAIMS/MMTS |
| P08249 | P08249 [297-307] | [K].KGLEKNLGIGK.[I] | 1xHomocysteineMMTS [K5] | 0,00869467 | 1319,71739 | Brain/ FAIMS/MMTS |
| P08249 | P08249 [298-307] | [K].GLEKNLGIGK.[I] | 1xHomocysteineMMTS [K4] | 6,70238E-05 | 1191,62243 | Brain/ FAIMS/MMTS |
| P08249 | P08249 [230-241] | [R].IQEAGTEVVKAK.[A] | 1xHomocysteineMMTS [K10] | 4,49774E-05 | 1435,72835 | Brain/ FAIMS/MMTS |
| P08249 | P08249 [315-328] | [K].MIAEAIPELKASIK.[K] | 1xHomocysteineMMTS [K10] | 4,49774E-05 | 1676,87838 | Brain/ FAIMS/MMTS |
| P08551 | P08551 [341-360] | [K].QNADISAMQDTINKLENELR.[S] | 1xHomocysteineIAA [K14] | 4,32935E-05 | 2477,1759 | Brain/ No FAIMS/IAA |
| P08551 | P08551 [341-360] | [K].QNADISAMQDTINKLENELR.[S] | 1xHomocysteineIAA [K14] | 4,32935E-05 | 2477,1759 | Brain/ FAIMS/IAA |
| P08551 | P08551 [88-100] | [R].TQEKAQLQDLNDR.[F] | 1xHomocysteineMMTS [K4] | 0,000152766 | 1721,79453 | Brain/ No FAIMS/MMTS |
| P08551 | P08551 [301-310] | [R].AAKDEVSESR.[R] | 1xHomocysteineMMTS [K3] | 0,00230389 | 1254,5453 | Brain/ No FAIMS/MMTS |
| P08551 | P08551 [285-300] | [R].FTVLTESAAKNTDAVR.[A] | 1xHomocysteineMMTS [K10] | 4,49774E-05 | 1885,91464 | Brain/ No FAIMS/MMTS |
| P08551 | P08551 [148-164] | [R].LAAEDATNEKQALQGER.[E] | 1xHomocysteineMMTS [K10] | 4,49774E-05 | 2006,927 | Brain/ No FAIMS/MMTS |
| P08551 | P08551 [197-206] | [R].KGADEAALAR.[A] | 1xHomocysteineMMTS [K1] | 0,0004709 | 1164,54999 | Brain/ No FAIMS/MMTS |
| P08551 | P08551 [269-282] | [K].LAAKNMQNAEEWFK.[S] | 1xHomocysteineMMTS [K4] | 4,49774E-05 | 1842,83356 | Brain/ FAIMS/MMTS |
| P08551 | P08551 [88-100] | [R].TQEKAQLQDLNDR.[F] | 1xHomocysteineMMTS [K4] | 0,000152766 | 1721,79453 | Brain/ FAIMS/MMTS |
| P08551 | P08551 [369-380] | [R].YLKEYQDLLNVK.[M] | 1xHomocysteineMMTS [K3] | 0,000530332 | 1688,83863 | Brain/ FAIMS/MMTS |
| P08551 | P08551 [301-310] | [R].AAKDEVSESR.[R] | 1xHomocysteineMMTS [K3] | 0,00230389 | 1254,5453 | Brain/ FAIMS/MMTS |
| P08551 | P08551 [285-300] | [R].FTVLTESAAKNTDAVR.[A] | 1xHomocysteineMMTS [K10] | 4,49774E-05 | 1885,91464 | Brain/ FAIMS/MMTS |
| P08551 | P08551 [148-164] | [R].LAAEDATNEKQALQGER.[E] | 1xHomocysteineMMTS [K10] | 4,49774E-05 | 2006,927 | Brain/ FAIMS/MMTS |
| P08551 | P08551 [197-206] | [R].KGADEAALAR.[A] | 1xHomocysteineMMTS [K1] | 0,0004709 | 1164,54999 | Brain/ FAIMS/MMTS |
| P08551;<br>P08553;<br>P46660 | P08551 [392-400];<br>P08553 [401-409];<br>P46660 [398-406] | [R].KLLEGEETR.[LF] | 1xHomocysteineIAA [K1] | 0,000570625 | 1248,62526 | Brain/ No FAIMS/IAA |
| P08551;<br>P08553;<br>P46660 | P08551 [392-400];<br>P08553 [401-409];<br>P46660 [398-406] | [R].KLLEGEETR.[LF] | 1xHomocysteineIAA [K1] | 0,000570625 | 1248,62526 | Brain/ FAIMS/IAA |
| P08551;<br>P08553;<br>P46660 | P08551 [392-400];<br>P08553 [401-409];<br>P46660 [398-406] | [R].KLLEGEETR.[LF] | 1xHomocysteineMMTS [K1] | 0,000266738 | 1237,59152 | Brain/ FAIMS/MMTS |
| P08553 | P08553 [426-437] | [R].QPSVTISSKIQK.[T] | 1xHomocysteineIAA [K9] | 4,32935E-05 | 1489,80429 | Brain/ FAIMS/IAA |
| P08553 | P08553 [297-309] | [K].LTEAAEQNKEAIR.[S] | 1xHomocysteineMMTS [K9] | 4,49774E-05 | 1635,7829 | Liver/ FAIMS/MMTS |
| P08553 | P08553 [297-309] | [K].LTEAAEQNKEAIR.[S] | 1xHomocysteineMMTS [K9] | 4,49774E-05 | 1635,7829 | Brain/ No FAIMS/MMTS |
| P08553 | P08553 [262-272] | [K].TDISTALKEIR.[S] | 1xHomocysteineMMTS [K8] | 4,49774E-05 | 1409,7127 | Brain/ No FAIMS/MMTS |
| P08553 | P08553 [775-788] | [K].KGEDSSDDKVVVTK.[K] | 1xHomocysteineMMTS [K9] | 0,00460899 | 1669,77715 | Brain/ FAIMS/MMTS |
| P08553 | P08553 [297-309] | [K].LTEAAEQNKEAIR.[S] | 1xHomocysteineMMTS [K9] | 4,49774E-05 | 1635,7829 | Brain/ FAIMS/MMTS |
| P08553 | P08553 [117-135] | [K].VHYLEQQNKEIEAEIQALR.[Q] | 1xHomocysteineMMTS [K9] | 0,00107839 | 2474,21664 | Brain/ FAIMS/MMTS |
| P08553 | P08553 [262-272] | [K].TDISTALKEIR.[S] | 1xHomocysteineMMTS [K8] | 4,49774E-05 | 1409,7127 | Brain/ FAIMS/MMTS |
| P08553 | P08553 [110-125] | [R].FAGYIEKVHYLEQQNK.[E] | 1xHomocysteineMMTS [K7] | 4,49774E-05 | 2130,01469 | Brain/ FAIMS/MMTS |
| P08553 | P08553 [321-333] | [R].QLQSKSIELESVR.[G] | 1xHomocysteineMMTS [K5] | 0,0017273 | 1679,8455 | Brain/ FAIMS/MMTS |
| P08553 | P08553 [831-842] | [K].KVEKVTSHAIVK.[E] | 1xHomocysteineMMTS [K4] | 0,000133296 | 1501,82292 | Brain/ FAIMS/MMTS |
| P08553 | P08553 [789-803] | [K].KVEKITSEGGDGATK.[Y] | 1xHomocysteineMMTS [K4] | 4,49774E-05 | 1682,80878 | Brain/ FAIMS/MMTS |
| P08553 | P08553 [804-814] | [K].YITKSVTVTQK.[V] | 1xHomocysteineMMTS [K4] | 0,00452171 | 1430,73818 | Brain/ FAIMS/MMTS |
| P08553 | P08553 [294-305] | [R].YAKLTEAAEQNK.[E] | 1xHomocysteineMMTS [K3] | 0,000530332 | 1528,71343 | Brain/ FAIMS/MMTS |
| P08553 | P08553 [310-319] | [R].SAKEEIAEYR.[R] | 1xHomocysteineMMTS [K3] | 0,00466355 | 1358,6079 | Brain/ FAIMS/MMTS |

| | | | | | | |
|---|---|---|---|---|---|---|
| P08553 | P08553 [136-153] | [R].QKQASHAQLGDAYDQEIR.[E] | 1xHomocysteineMMTS [K2] | 4,49774E-05 | 2221,01246 | Brain/ FAIMS/MMTS |
| P08553 | P08553 [221-233] | [K].KVQSLQDEVAFLR.[S] | 1xHomocysteineMMTS [K1] | 0,000738471 | 1695,85567 | Brain/ FAIMS/MMTS |
| P08752 | P08752 [308-314] | [K].FEDLNKR.[K] | 1xHomocysteineIAA [K6] | 0,00863543 | 1095,52515 | Brain/ No FAIMS/IAA |
| P08752; B2RSH2 | P08752 [316-331]; B2RSH2 [315-330] | [K].DTKEIYTHFTCATDTK.[N] | 1xCarbamidomethyl [C11]; 1xHomocysteineIAA [K3] | 4,32935E-05 | 2104,93142 | Brain/ FAIMS/IAA |
| P08752; B2RSH2 | P08752 [194-206]; B2RSH2 [193-205] | [K].DLHFKMFDVGGQR.[S] | 1xHomocysteineMMTS [K5] | 0,00952012 | 1712,77056 | Brain/ FAIMS/MMTS |
| P09041 | P09041 [7-17] | [K].LTLDKVDLKGK.[R] | 1xHomocysteineIAA [K9] | 0,000595249 | 1403,79266 | Brain/ FAIMS/IAA |
| P09041; P09411 | P09041 [23-39]; P09411 [23-39] | [R].VDFNVPMKNNQITNNQR.[I] | 1xHomocysteineIAA [K8] | 4,32935E-05 | 2206,04918 | Brain/ FAIMS/IAA |
| P09103 | P09103 [286-302] | [K].GKILFIFIDSDHTDNQR.[I] | 1xHomocysteineIAA [K2] | 4,32935E-05 | 2193,07571 | Liver/ FAIMS/IAA |
| P09103 | P09103 [257-273] | [K].THILLFLPKSVSDYDGK.[L] | 1xHomocysteineMMTS [K9] | 4,49774E-05 | 2096,05549 | Liver/ No FAIMS/MMTS |
| P09103 | P09103 [216-226] | [R].NNFEGEITKEK.[L] | 1xHomocysteineMMTS [K9] | 4,49774E-05 | 1471,65558 | Liver/ No FAIMS/MMTS |
| P09103 | P09103 [266-278] | [K].SVSDYDGKLSSFK.[R] | 1xHomocysteineMMTS [K8] | 4,49774E-05 | 1595,70801 | Liver/ No FAIMS/MMTS |
| P09103 | P09103 [439-454] | [K].VHSFPTLKFFPASADR.[T] | 1xHomocysteineMMTS [K8] | 4,49774E-05 | 1982,96153 | Liver/ No FAIMS/MMTS |
| P09103 | P09103 [303-311] | [R].ILEFFGLKK.[E] | 1xHomocysteineMMTS [K8] | 0,00127368 | 1257,6734 | Liver/ No FAIMS/MMTS |
| P09103 | P09103 [60-71] | [K].ALAPEYAKAAAK.[L] | 1xHomocysteineMMTS [K8] | 4,49774E-05 | 1366,68575 | Liver/ No FAIMS/MMTS |
| P09103 | P09103 [412-426] | [K].LGETYKDHENIIIAK.[M] | 1xHomocysteineMMTS [K6] | 6,70238E-05 | 1906,94013 | Liver/ No FAIMS/MMTS |
| P09103 | P09103 [100-108] | [R].GYPTIKFFK.[N] | 1xHomocysteineMMTS [K6] | 0,00312202 | 1263,62645 | Liver/ No FAIMS/MMTS |
| P09103 | P09103 [198-209] | [K].YQLDKDGVVLFK.[K] | 1xHomocysteineMMTS [K5] | 4,49774E-05 | 1587,79095 | Liver/ No FAIMS/MMTS |
| P09103 | P09103 [81-99] | [R].LAKVDATEESDLAQQYGVR.[G] | 1xHomocysteineMMTS [K3] | 4,49774E-05 | 2256,06349 | Liver/ No FAIMS/MMTS |
| P09103 | P09103 [225-232] | [K].EKLLDFIK.[H] | 1xHomocysteineMMTS [K2] | 0,00246851 | 1168,61046 | Liver/ No FAIMS/MMTS |
| P09103 | P09103 [329-340] | [K].YKPESDELTAEK.[I] | 1xHomocysteineMMTS [K2] | 4,49774E-05 | 1572,69202 | Liver/ No FAIMS/MMTS |
| P09103 | P09103 [72-80] | [K].LKAEGSEIR.[L] | 1xHomocysteineMMTS [K2] | 0,00180518 | 1165,57039 | Liver/ No FAIMS/MMTS |
| P09103 | P09103 [427-446] | [K].MDSTANEVEAVKVHSFPTLK.[F] | 1xHomocysteineMMTS [K12] | 4,49774E-05 | 2366,1189 | Liver/ No FAIMS/MMTS |
| P09103 | P09103 [123-133] | [R].EADDIVNWLKK.[R] | 1xHomocysteineMMTS [K10] | 0,00135666 | 1493,7127 | Liver/ No FAIMS/MMTS |
| P09103 | P09103 [427-446] | [K].MDSTANEVEAVKVHSFPTLK.[F] | 1xOxidation [M1]; 1xHomocysteineMMTS [K12] | 4,49774E-05 | 2382,11381 | Liver/ FAIMS/MMTS |
| P09103 | P09103 [216-226] | [R].NNFEGEITKEK.[L] | 1xHomocysteineMMTS [K9] | 4,49774E-05 | 1471,65558 | Liver/ FAIMS/MMTS |
| P09103 | P09103 [266-278] | [K].SVSDYDGKLSSFK.[R] | 1xHomocysteineMMTS [K8] | 4,49774E-05 | 1595,70801 | Liver/ FAIMS/MMTS |
| P09103 | P09103 [439-454] | [K].VHSFPTLKFFPASADR.[T] | 1xHomocysteineMMTS [K8] | 4,49774E-05 | 1982,96153 | Liver/ FAIMS/MMTS |
| P09103 | P09103 [303-311] | [R].ILEFFGLKK.[E] | 1xHomocysteineMMTS [K8] | 0,00127368 | 1257,6734 | Liver/ FAIMS/MMTS |
| P09103 | P09103 [60-71] | [K].ALAPEYAKAAAK.[L] | 1xHomocysteineMMTS [K8] | 4,49774E-05 | 1366,68575 | Liver/ FAIMS/MMTS |
| P09103 | P09103 [100-108] | [R].GYPTIKFFK.[N] | 1xHomocysteineMMTS [K6] | 0,00312202 | 1263,62645 | Liver/ FAIMS/MMTS |
| P09103 | P09103 [198-209] | [K].YQLDKDGVVLFK.[K] | 1xHomocysteineMMTS [K5] | 4,49774E-05 | 1587,79095 | Liver/ FAIMS/MMTS |
| P09103 | P09103 [81-99] | [R].LAKVDATEESDLAQQYGVR.[G] | 1xHomocysteineMMTS [K3] | 4,49774E-05 | 2256,06349 | Liver/ FAIMS/MMTS |
| P09103 | P09103 [225-232] | [K].EKLLDFIK.[H] | 1xHomocysteineMMTS [K2] | 0,00246851 | 1168,61046 | Liver/ FAIMS/MMTS |
| P09103 | P09103 [72-80] | [K].LKAEGSEIR.[L] | 1xHomocysteineMMTS [K2] | 0,00180518 | 1165,57039 | Liver/ FAIMS/MMTS |
| P09103 | P09103 [427-446] | [K].MDSTANEVEAVKVHSFPTLK.[F] | 1xHomocysteineMMTS [K12] | 4,49774E-05 | 2366,1189 | Liver/ FAIMS/MMTS |
| P09103 | P09103 [329-347] | [K].YKPESDELTAEKITEFCHR.[F] | 1xMethylthio [C17]; 1xHomocysteineMMTS [K12] | 4,49774E-05 | 2505,0917 | Liver/ FAIMS/MMTS |
| P09103 | P09103 [257-273] | [K].THILLFLPKSVSDYDGK.[L] | 1xHomocysteineMMTS [K9] | 4,49774E-05 | 2096,05549 | Liver/ FAIMS/MMTS |
| P09103 | P09103 [109-122] | [K].NGDTASPKEYTAGR.[E] | 1xHomocysteineMMTS [K8] | 0,000547722 | 1629,69957 | Liver/ FAIMS/MMTS |
| P09103 | P09103 [412-426] | [K].LGETYKDHENIIIAK.[M] | 1xHomocysteineMMTS [K6] | 6,70238E-05 | 1906,94013 | Liver/ FAIMS/MMTS |
| P09103 | P09103 [106-116] | [K].FFKNGDTASPK.[E] | 1xHomocysteineMMTS [K3] | 0,0025135 | 1374,61807 | Liver/ FAIMS/MMTS |
| P09103 | P09103 [329-340] | [K].YKPESDELTAEK.[I] | 1xHomocysteineMMTS [K2] | 4,49774E-05 | 1572,69202 | Liver/ FAIMS/MMTS |
| P09103 | P09103 [353-368] | [K].IKPHLMSQEVPEDWDK.[Q] | 1xHomocysteineMMTS [K2] | 4,49774E-05 | 2114,97078 | Liver/ FAIMS/MMTS |
| P09103 | P09103 [412-438] | [K].LGETYKDHENIIIAKMDSTANEVEAVK.[V] | 1xHomocysteineMMTS [K15] | 4,49774E-05 | 3181,5214 | Liver/ FAIMS/MMTS |
| P09103 | P09103 [373-388] | [K].VLVGANFEEVAFDEKK.[N] | 1xHomocysteineMMTS [K15] | 0,000706375 | 1957,9398 | Liver/ FAIMS/MMTS |
| P09103 | P09103 [319-340] | [R].LITLEEEMTKYKPESDELTAEK.[I] | 1xHomocysteineMMTS [K10] | 4,49774E-05 | 2760,3028 | Liver/ FAIMS/MMTS |
| P09103 | P09103 [123-133] | [R].EADDIVNWLKK.[R] | 1xHomocysteineMMTS [K10] | 0,00135666 | 1493,7127 | Liver/ FAIMS/MMTS |
| P09103 | P09103 [33-44] | [K].KSNFEEALAAHK.[Y] | 1xHomocysteineMMTS [K1] | 4,49774E-05 | 1507,70319 | Liver/ FAIMS/MMTS |
| P09411 | P09411 [354-365] | [K].SLMDEVVKATSR.[G] | 1xHomocysteineIAA [K8] | 4,32935E-05 | 1509,73997 | Liver/ FAIMS/IAA |
| P09411 | P09411 [354-365] | [K].SLMDEVVKATSR.[G] | 1xHomocysteineIAA [K8] | 4,32935E-05 | 1509,73997 | Brain/ No FAIMS/IAA |
| P09411 | P09411 [354-365] | [K].SLMDEVVKATSR.[G] | 1xHomocysteineIAA [K8] | 4,32935E-05 | 1509,73997 | Brain/ FAIMS/IAA |
| P09411 | P09411 [1-15] | [-].MSLSNKLTLDKLDVK.[G] | 1xMet-loss+Acetyl [N-Term]; 1xHomocysteineMMTS [K6] | 4,49774E-05 | 1778,93907 | Liver/ No FAIMS/MMTS |
| P09411 | P09411 [87-97] | [K].SLLGKDVLFLK.[D] | 1xHomocysteineMMTS [K5] | 0,000133296 | 1395,77384 | Liver/ No FAIMS/MMTS |
| P09411 | P09411 [354-365] | [K].SLMDEVVKATSR.[G] | 1xHomocysteineMMTS [K8] | 4,49774E-05 | 1498,70623 | Liver/ FAIMS/MMTS |
| P09411 | P09411 [1-11] | [-].MSLSNKLTLDK.[L] | 1xMet-loss+Acetyl [N-Term]; 1xHomocysteineMMTS [K6] | 4,49774E-05 | 1323,66468 | Liver/ FAIMS/MMTS |
| P09411 | P09411 [1-15] | [-].MSLSNKLTLDKLDVK.[G] | 1xMet-loss+Acetyl [N-Term]; 1xHomocysteineMMTS [K6] | 4,49774E-05 | 1778,93907 | Liver/ FAIMS/MMTS |
| P09411 | P09411 [7-17] | [K].LTLDKLDVKGK.[R] | 1xHomocysteineMMTS [K9] | 4,49774E-05 | 1392,75892 | Liver/ FAIMS/MMTS |
| P09411 | P09411 [124-139] | [R].FHVEEEGKGKDASGNK.[V] | 1xHomocysteineMMTS [K8] | 0,00135666 | 1894,84221 | Liver/ FAIMS/MMTS |
| P09411 | P09411 [152-171] | [R].ASLSKLGDVYVNDAFGTAHR.[A] | 1xHomocysteineMMTS [K5] | 4,49774E-05 | 2284,0849 | Liver/ FAIMS/MMTS |
| P09411 | P09411 [7-15] | [K].LTLDKLDVK.[G] | 1xHomocysteineMMTS [K5] | 0,00756679 | 1207,64249 | Liver/ FAIMS/MMTS |
| P09411 | P09411 [87-97] | [K].SLLGKDVLFLK.[D] | 1xHomocysteineMMTS [K5] | 0,000133296 | 1395,77384 | Liver/ FAIMS/MMTS |
| P09411 | P09411 [217-230] | [K].VADKIQLINNMLDK.[V] | 1xHomocysteineMMTS [K4] | 0,00107839 | 1777,90091 | Liver/ FAIMS/MMTS |
| P09411 | P09411 [351-361] | [R].GTKSLMDEVVK.[A] | 1xHomocysteineMMTS [K3] | 0,00830327 | 1369,6524 | Liver/ FAIMS/MMTS |
| P09411 | P09411 [280-297] | [K].ITLPVDFVTADKFDENAK.[T] | 1xHomocysteineMMTS [K12] | 4,49774E-05 | 2186,0508 | Liver/ FAIMS/MMTS |
| P09411 | P09411 [1-11] | [-].MSLSNKLTLDK.[L] | 1xMet-loss+Acetyl [N-Term]; 1xHomocysteineMMTS [K6] | 4,49774E-05 | 1323,66468 | Brain/ FAIMS/MMTS |
| P09411 | P09411 [354-365] | [K].SLMDEVVKATSR.[G] | 1xHomocysteineMMTS [K8] | 4,49774E-05 | 1498,70623 | Brain/ FAIMS/MMTS |
| P09411 | P09411 [152-171] | [R].ASLSKLGDVYVNDAFGTAHR.[A] | 1xHomocysteineMMTS [K5] | 4,49774E-05 | 2284,0849 | Brain/ FAIMS/MMTS |
| P09411 | P09411 [7-15] | [K].LTLDKLDVK.[G] | 1xHomocysteineMMTS [K5] | 0,00756679 | 1207,64249 | Brain/ FAIMS/MMTS |
| P09411 | P09411 [87-97] | [K].SLLGKDVLFLK.[D] | 1xHomocysteineMMTS [K5] | 0,000133296 | 1395,77384 | Brain/ FAIMS/MMTS |
| P09411 | P09411 [280-297] | [K].ITLPVDFVTADKFDENAK.[T] | 1xHomocysteineMMTS [K12] | 4,49774E-05 | 2186,0508 | Brain/ FAIMS/MMTS |
| P09528 | P09528 [158-173] | [R].KMGAPEAGMAEYLFDK.[H] | 1xHomocysteineIAA [K1] | 4,32935E-05 | 1931,87 | Brain/ No FAIMS/IAA |
| P09528 | P09528 [70-77] | [K].LMKLQNQR.[G] | 1xHomocysteineMMTS [K3] | 0,00461842 | 1193,59516 | Liver/ No FAIMS/MMTS |
| P09528 | P09528 [70-77] | [K].LMKLQNQR.[G] | 1xHomocysteineMMTS [K3] | 0,00461842 | 1193,59516 | Liver/ FAIMS/MMTS |

| | | | | | | |
|---|---|---|---|---|---|---|
| P09528 | P09528 [145-157] | [K].SIKELGDHVTNLR.[K] | 1xHomocysteineMMTS [K3] | 0,000250158 | 1644,81962 | Liver/ FAIMS/MMTS |
| P09528 | P09528 [159-182] | [K].MGAPEAGMAEYLFDKHTLGHGDES.[-] | 1xHomocysteineMMTS [K15] | 4,49774E-05 | 2726,13535 | Liver/ FAIMS/MMTS |
| P09528 | P09528 [145-157] | [K].SIKELGDHVTNLR.[K] | 1xHomocysteineMMTS [K3] | 0,000250158 | 1644,81962 | Brain/ FAIMS/MMTS |
| P09528 | P09528 [159-182] | [K].MGAPEAGMAEYLFDKHTLGHGDES.[-] | 1xHomocysteineMMTS [K15] | 4,49774E-05 | 2726,13535 | Brain/ FAIMS/MMTS |
| P09542 | P09542 [15-42] | [K].AAAPKAAPAPAAAPAAAPAAAPEPERPK.[E] | 1xHomocysteineIAA [K5] | 4,32935E-05 | 2705,41918 | Liver/ No FAIMS/IAA |
| P09542 | P09542 [15-42] | [K].AAAPKAAPAPAAAPAAAPAAAPEPERPK.[E] | 1xHomocysteineIAA [K5] | 4,32935E-05 | 2705,41918 | Liver/ FAIMS/IAA |
| P09542 | P09542 [15-42] | [K].AAAPKAAPAPAAAPAAAPAAAPEPERPK.[E] | 1xHomocysteineMMTS [K5] | 4,49774E-05 | 2694,38544 | Liver/ FAIMS/MMTS |
| P09542 | P09542 [133-147] | [K].NKDTGTYEDFVEGLR.[V] | 1xHomocysteineMMTS [K2] | 0,00169472 | 1906,83097 | Liver/ FAIMS/MMTS |
| P09542 | P09542 [51-64] | [K].IKIEFTPEQIEEFK.[E] | 1xHomocysteineMMTS [K2] | 6,70238E-05 | 1913,93873 | Liver/ FAIMS/MMTS |
| P09671 | P09671 [124-132] | [R].DFGSFEKFK.[E] | 1xHomocysteineIAA [K7] | 0,000258769 | 1278,58233 | Liver/ FAIMS/IAA |
| P09671 | P09671 [195-216] | [K].NVRPDYLKAIWNVINWENVTER.[Y] | 1xHomocysteineIAA [K8] | 4,32935E-05 | 2903,46211 | Brain/ FAIMS/IAA |
| P09671 | P09671 [115-123] | [K].GELLEAIKR.[D] | 1xHomocysteineMMTS [K8] | 0,000915247 | 1191,62243 | Liver/ No FAIMS/MMTS |
| P09671 | P09671 [124-132] | [R].DFGSFEKFK.[E] | 1xHomocysteineMMTS [K7] | 0,00246851 | 1267,54859 | Liver/ No FAIMS/MMTS |
| P09671 | P09671 [115-123] | [K].GELLEAIKR.[D] | 1xHomocysteineMMTS [K8] | 0,000915247 | 1191,62243 | Liver/ FAIMS/MMTS |
| P09671 | P09671 [69-89] | [K].YHEALAKGDVTTQVALQPALK.[F] | 1xHomocysteineMMTS [K7] | 4,49774E-05 | 2416,23631 | Liver/ FAIMS/MMTS |
| P09671 | P09671 [124-132] | [R].DFGSFEKFK.[E] | 1xHomocysteineMMTS [K7] | 0,00246851 | 1267,54859 | Liver/ FAIMS/MMTS |
| P09671 | P09671 [109-122] | [K].GGGEPKGELLEAIK.[R] | 1xHomocysteineMMTS [K6] | 4,49774E-05 | 1560,77603 | Liver/ FAIMS/MMTS |
| P09671 | P09671 [124-132] | [R].DFGSFEKFK.[E] | 1xHomocysteineMMTS [K7] | 0,00246851 | 1267,54859 | Brain/ No FAIMS/MMTS |
| P09671 | P09671 [115-123] | [K].GELLEAIKR.[D] | 1xHomocysteineMMTS [K8] | 0,000915247 | 1191,62243 | Brain/ FAIMS/MMTS |
| P09813 | P09813 [54-67] | [K].TSEIQSQAKAYFEK.[T] | 1xHomocysteineMMTS [K9] | 0,000133296 | 1792,82443 | Liver/ No FAIMS/MMTS |
| P09813 | P09813 [52-62] | [K].AKTSEIQSQAK.[A] | 1xHomocysteineMMTS [K2] | 0,000192903 | 1353,6501 | Liver/ No FAIMS/MMTS |
| P09813 | P09813 [54-67] | [K].TSEIQSQAKAYFEK.[T] | 1xHomocysteineMMTS [K9] | 0,000133296 | 1792,82443 | Liver/ FAIMS/MMTS |
| P09813 | P09813 [63-77] | [K].AYFEKTHEQLTPLVR.[S] | 1xHomocysteineMMTS [K5] | 0,00576292 | 1994,98266 | Liver/ FAIMS/MMTS |
| P09813 | P09813 [52-62] | [K].AKTSEIQSQAK.[A] | 1xHomocysteineMMTS [K2] | 0,000192903 | 1353,6501 | Liver/ FAIMS/MMTS |
| P0DN34 | P0DN34 [47-57] | [R].ELRPNEEVTWK.[-] | 1xHomocysteineIAA [K11] | 0,000876609 | 1574,76315 | Brain/ FAIMS/IAA |
| P0DP27 | P0DP27 [23-38] | [K].DGDGTITTKELGTVMR.[S] | 1xHomocysteineIAA [K9] | 4,32935E-05 | 1867,88882 | Liver/ FAIMS/IAA |
| P0DP27 | P0DP27 [1-31] | [-].MADQLTEEQIAEFKEAFSLFDKDGDGTITTK.[E] | 1xMet-loss+Acetyl [N-Term]; 1xHomocysteineIAA [K14] | 4,32935E-05 | 3563,6734 | Brain/ No FAIMS/IAA |
| P0DP27 | P0DP27 [1-31] | [-].MADQLTEEQIAEFKEAFSLFDKDGDGTITTK.[E] | 1xMet-loss+Acetyl [N-Term]; 1xHomocysteineIAA [K14] | 4,32935E-05 | 3563,6734 | Brain/ FAIMS/IAA |
| P0DP27 | P0DP27 [15-31] | [K].EAFSLFDKDGDGTITTK.[E] | 1xHomocysteineMMTS [K8] | 4,49774E-05 | 2007,9038 | Liver/ No FAIMS/MMTS |
| P0DP27 | P0DP27 [92-107] | [R].VFDKDGNGYISAAELR.[H] | 1xHomocysteineMMTS [K4] | 4,49774E-05 | 1917,88334 | Liver/ No FAIMS/MMTS |
| P0DP27 | P0DP27 [15-38] | [K].EAFSLFDKDGDGTITTKELGTVMR.[S] | 1xHomocysteineMMTS [K17] | 4,49774E-05 | 2794,30961 | Liver/ No FAIMS/MMTS |
| P0DP27 | P0DP27 [23-38] | [K].DGDGTITTKELGTVMR.[S] | 1xHomocysteineMMTS [K9] | 4,49774E-05 | 1856,85508 | Liver/ FAIMS/MMTS |
| P0DP27 | P0DP27 [15-31] | [K].EAFSLFDKDGDGTITTK.[E] | 1xHomocysteineMMTS [K8] | 4,49774E-05 | 2007,9038 | Liver/ FAIMS/MMTS |
| P0DP27 | P0DP27 [92-107] | [R].VFDKDGNGYISAAELR.[H] | 1xHomocysteineMMTS [K4] | 4,49774E-05 | 1917,88334 | Liver/ FAIMS/MMTS |
| P0DP27 | P0DP27 [15-38] | [K].EAFSLFDKDGDGTITTKELGTVMR.[S] | 1xHomocysteineMMTS [K17] | 4,49774E-05 | 2794,30961 | Liver/ FAIMS/MMTS |
| P0DP27 | P0DP27 [23-38] | [K].DGDGTITTKELGTVMR.[S] | 1xHomocysteineMMTS [K9] | 4,49774E-05 | 1856,85508 | Brain/ No FAIMS/MMTS |
| P0DP27 | P0DP27 [15-31] | [K].EAFSLFDKDGDGTITTK.[E] | 1xHomocysteineMMTS [K8] | 4,49774E-05 | 2007,9038 | Brain/ No FAIMS/MMTS |
| P0DP27 | P0DP27 [92-107] | [R].VFDKDGNGYISAAELR.[H] | 1xHomocysteineMMTS [K4] | 4,49774E-05 | 1917,88334 | Brain/ No FAIMS/MMTS |
| P0DP27 | P0DP27 [15-38] | [K].EAFSLFDKDGDGTITTKELGTVMR.[S] | 1xHomocysteineMMTS [K17] | 4,49774E-05 | 2794,30961 | Brain/ No FAIMS/MMTS |
| P0DP27 | P0DP27 [15-38] | [K].EAFSLFDKDGDGTITTKELGTVMR.[S] | 1xOxidation [M23]; 1xHomocysteineMMTS [K17] | 0,000282457 | 2810,30453 | Brain/ FAIMS/MMTS |
| P0DP27 | P0DP27 [23-38] | [K].DGDGTITTKELGTVMR.[S] | 1xHomocysteineMMTS [K9] | 4,49774E-05 | 1856,85508 | Brain/ FAIMS/MMTS |
| P0DP27 | P0DP27 [15-31] | [K].EAFSLFDKDGDGTITTK.[E] | 1xHomocysteineMMTS [K8] | 4,49774E-05 | 2007,9038 | Brain/ FAIMS/MMTS |
| P0DP27 | P0DP27 [92-107] | [R].VFDKDGNGYISAAELR.[H] | 1xHomocysteineMMTS [K4] | 4,49774E-05 | 1917,88334 | Brain/ FAIMS/MMTS |
| P0DP27 | P0DP27 [15-38] | [K].EAFSLFDKDGDGTITTKELGTVMR.[S] | 1xHomocysteineMMTS [K17] | 4,49774E-05 | 2794,30961 | Brain/ FAIMS/MMTS |
| P0DP27; Q9D6P8 | P0DP27 [76-87]; Q9D6P8 [76-87] | [R].KMKDTDSEEEIR.[E] | 1xHomocysteineIAA [K] | 4,32935E-05 | 1654,7411 | Liver/ No FAIMS/IAA |
| P0DP27; Q9D6P8 | P0DP27 [77-91]; Q9D6P8 [77-91] | [K].MKDTDSEEEIREAFR.[V] | 1xHomocysteineIAA [K2] | 4,32935E-05 | 2029,89536 | Liver/ FAIMS/IAA |
| P0DP27; Q9D6P8 | P0DP27 [76-87]; Q9D6P8 [76-87] | [R].KMKDTDSEEEIR.[E] | 1xHomocysteineIAA [K] | 4,32935E-05 | 1654,7411 | Brain/ No FAIMS/IAA |
| P0DP27; Q9D6P8 | P0DP27 [76-87]; Q9D6P8 [76-87] | [R].KMKDTDSEEEIR.[E] | 1xHomocysteineIAA [K] | 4,32935E-05 | 1654,7411 | Brain/ FAIMS/IAA |
| P0DP27; Q9D6P8 | P0DP27 [76-87]; Q9D6P8 [76-87] | [R].KMKDTDSEEEIR.[E] | 1xOxidation [M2]; 1xHomocysteineMMTS [K3] | 0,000530332 | 1659,70227 | Liver/ No FAIMS/MMTS |
| P0DP27; Q9D6P8 | P0DP27 [77-87]; Q9D6P8 [77-87] | [K].MKDTDSEEEIR.[E] | 1xHomocysteineMMTS [K2] | 4,49774E-05 | 1515,61239 | Liver/ FAIMS/MMTS |
| P0DP27; Q9D6P8 | P0DP27 [76-87]; Q9D6P8 [76-87] | [R].KMKDTDSEEEIR.[E] | 1xOxidation [M2]; 1xHomocysteineMMTS [K3] | 0,000530332 | 1659,70227 | Liver/ FAIMS/MMTS |
| P0DP27; Q9D6P8 | P0DP27 [76-87]; Q9D6P8 [76-87] | [R].KMKDTDSEEEIR.[E] | 1xHomocysteineMMTS [K] | 4,49774E-05 | 1643,70735 | Liver/ FAIMS/MMTS |
| P0DP27; Q9D6P8 | P0DP27 [76-87]; Q9D6P8 [76-87] | [R].KMKDTDSEEEIR.[E] | 1xOxidation [M2]; 1xHomocysteineMMTS [K3] | 0,000530332 | 1659,70227 | Brain/ No FAIMS/MMTS |
| P0DP27; Q9D6P8 | P0DP27 [77-87]; Q9D6P8 [77-87] | [K].MKDTDSEEEIR.[E] | 1xHomocysteineMMTS [K2] | 4,49774E-05 | 1515,61239 | Brain/ No FAIMS/MMTS |
| P0DP27; Q9D6P8 | P0DP27 [77-91]; Q9D6P8 [77-91] | [K].MKDTDSEEEIREAFR.[V] | 1xHomocysteineMMTS [K2] | 4,49774E-05 | 2018,86162 | Brain/ No FAIMS/MMTS |
| P0DP27; Q9D6P8 | P0DP27 [76-87]; Q9D6P8 [76-87] | [R].KMKDTDSEEEIR.[E] | 1xHomocysteineMMTS [K] | 4,49774E-05 | 1643,70735 | Brain/ No FAIMS/MMTS |
| P0DP27; Q9D6P8 | P0DP27 [76-87]; Q9D6P8 [76-87] | [R].KMKDTDSEEEIR.[E] | 1xOxidation [M2]; 1xHomocysteineMMTS [K3] | 0,000530332 | 1659,70227 | Brain/ FAIMS/MMTS |
| P0DP27; Q9D6P8 | P0DP27 [77-87]; Q9D6P8 [77-87] | [K].MKDTDSEEEIR.[E] | 1xHomocysteineMMTS [K2] | 4,49774E-05 | 1515,61239 | Brain/ FAIMS/MMTS |

| | | | | | | |
|---|---|---|---|---|---|---|
| P0DP27; Q9D6P8 | P0DP27 [77-91]; Q9D6P8 [77-91] | [K].MKDTDSEEEIREAFR.[V] | 1xHomocysteineMMTS [K2] | 4,49774E-05 | 2018,86162 | Brain/ FAIMS/MMTS |
| P0DP27; Q9D6P8 | P0DP27 [76-87]; Q9D6P8 [76-87] | [R].KMKDTDSEEEIR.[E] | 1xHomocysteineMMTS [K] | 4,49774E-05 | 1643,70735 | Brain/ FAIMS/MMTS |
| P10126 | P10126 [220-247] | [K].DGSASGTTLLEALDCILPPTRPTDKPLR.[L] | 1xCarbamidomethyl [C15]; 1xHomocysteineIAA [K25] | 4,32935E-05 | 3168,60276 | Brain/ FAIMS/IAA |
| P10126 | P10126 [167-179] | [R].YEEIVKEVSTYIK.[K] | 1xHomocysteineMMTS [K6] | 0,000547722 | 1763,85942 | Liver/ FAIMS/MMTS |
| P10126; P62631 | P10126 [248-266]; P62631 [248-266] | [R].LPLQDVYKIGGIGTVPVGR.[V] | 1xHomocysteineIAA [K8] | 4,32935E-05 | 2156,18962 | Liver/ FAIMS/IAA |
| P10126; P62631 | P10126 [248-266]; P62631 [248-266] | [R].LPLQDVYKIGGIGTVPVGR.[V] | 1xHomocysteineIAA [K8] | 4,32935E-05 | 2156,18962 | Brain/ No FAIMS/IAA |
| P10126; P62631 | P10126 [248-266]; P62631 [248-266] | [R].LPLQDVYKIGGIGTVPVGR.[V] | 1xHomocysteineIAA [K8] | 4,32935E-05 | 2156,18962 | Brain/ FAIMS/IAA |
| P10126; P62631 | P10126 [248-266]; P62631 [248-266] | [R].LPLQDVYKIGGIGTVPVGR.[V] | 1xHomocysteineMMTS [K8] | 4,49774E-05 | 2145,15588 | Liver/ FAIMS/MMTS |
| P10126; P62631 | P10126 [248-266]; P62631 [248-266] | [R].LPLQDVYKIGGIGTVPVGR.[V] | 1xHomocysteineMMTS [K8] | 4,49774E-05 | 2145,15588 | Brain/ FAIMS/MMTS |
| P10605 | P10605 [210-223] | [K].SCEAGYSPSYKEDK.[H] | 1xCarbamidomethyl [C2]; 1xHomocysteineIAA [K11] | 0,00677865 | 1794,73093 | Liver/ FAIMS/IAA |
| P10605 | P10605 [210-223] | [K].SCEAGYSPSYKEDK.[H] | 1xCarbamidomethyl [C2]; 1xHomocysteineIAA [K11] | 0,00677865 | 1794,73093 | Brain/ No FAIMS/IAA |
| P10637 | P10637 [591-609] | [K].HVPGGGSVQIVYKPVDLSK.[V] | 1xHomocysteineIAA [K13] | 4,32935E-05 | 2154,13758 | Brain/ No FAIMS/IAA |
| P10637 | P10637 [591-609] | [K].HVPGGGSVQIVYKPVDLSK.[V] | 1xHomocysteineIAA [K13] | 4,32935E-05 | 2154,13758 | Brain/ FAIMS/IAA |
| P10637 | P10637 [448-462] | [R].GAASPAQKGTSNATR.[I] | 1xHomocysteineMMTS [K8] | 0,000211896 | 1579,73153 | Brain/ No FAIMS/MMTS |
| P10637 | P10637 [463-472] | [R].IPAKTTPSPK.[T] | 1xHomocysteineMMTS [K4] | 0,00110986 | 1202,62718 | Brain/ No FAIMS/MMTS |
| P10637 | P10637 [448-462] | [R].GAASPAQKGTSNATR.[I] | 1xHomocysteineMMTS [K8] | 0,000211896 | 1579,73153 | Brain/ FAIMS/MMTS |
| P10637 | P10637 [560-572] | [K].HQPGGGKVQIINK.[K] | 1xHomocysteineMMTS [K7] | 0,00531384 | 1538,79301 | Brain/ FAIMS/MMTS |
| P10637 | P10637 [467-482] | [K].TTPSPKTPPGSGEPPK.[S] | 1xHomocysteineMMTS [K6] | 0,00980048 | 1740,82952 | Brain/ FAIMS/MMTS |
| P10639 | P10639 [4-21] | [K].LIESKEAFQEALAAAGDK.[L] | 1xHomocysteineIAA [K5] | 4,32935E-05 | 2065,02703 | Liver/ No FAIMS/IAA |
| P10639 | P10639 [4-21] | [K].LIESKEAFQEALAAAGDK.[L] | 1xHomocysteineIAA [K5] | 4,32935E-05 | 2065,02703 | Liver/ FAIMS/IAA |
| P10639 | P10639 [4-21] | [K].LIESKEAFQEALAAAGDK.[L] | 1xHomocysteineIAA [K5] | 4,32935E-05 | 2065,02703 | Brain/ No FAIMS/IAA |
| P10639 | P10639 [4-21] | [K].LIESKEAFQEALAAAGDK.[L] | 1xHomocysteineIAA [K5] | 4,32935E-05 | 2065,02703 | Brain/ FAIMS/IAA |
| P10639 | P10639 [86-96] | [K].VGEFSGANKEK.[L] | 1xHomocysteineMMTS [K9] | 4,49774E-05 | 1328,59733 | Liver/ No FAIMS/MMTS |
| P10639 | P10639 [4-21] | [K].LIESKEAFQEALAAAGDK.[L] | 1xHomocysteineMMTS [K5] | 4,49774E-05 | 2053,99329 | Liver/ No FAIMS/MMTS |
| P10639 | P10639 [83-94] | [K].GQKVGEFSGANK.[E] | 1xHomocysteineMMTS [K3] | 4,49774E-05 | 1384,63478 | Liver/ No FAIMS/MMTS |
| P10639 | P10639 [86-96] | [K].VGEFSGANKEK.[L] | 1xHomocysteineMMTS [K9] | 4,49774E-05 | 1328,59733 | Liver/ FAIMS/MMTS |
| P10639 | P10639 [83-94] | [K].GQKVGEFSGANK.[E] | 1xHomocysteineMMTS [K3] | 4,49774E-05 | 1384,63478 | Liver/ FAIMS/MMTS |
| P10639 | P10639 [4-21] | [K].LIESKEAFQEALAAAGDK.[L] | 1xHomocysteineMMTS [K5] | 4,49774E-05 | 2053,99329 | Liver/ FAIMS/MMTS |
| P10639 | P10639 [82-94] | [K].KGQKVGEFSGANK.[E] | 1xHomocysteineMMTS [K4] | 0,000282457 | 1512,72974 | Liver/ FAIMS/MMTS |
| P10639 | P10639 [86-96] | [K].VGEFSGANKEK.[L] | 1xHomocysteineMMTS [K9] | 4,49774E-05 | 1328,59733 | Brain/ FAIMS/MMTS |
| P10639 | P10639 [83-94] | [K].GQKVGEFSGANK.[E] | 1xHomocysteineMMTS [K3] | 4,49774E-05 | 1384,63478 | Brain/ FAIMS/MMTS |
| P10648; P30115 | P10648 [139-152]; P30115 [139-152] | [K].VLKSHGQDYLVGNR.[L] | 1xHomocysteineIAA [K3] | 4,32935E-05 | 1759,89081 | Liver/ No FAIMS/IAA |
| P10648; P30115 | P10648 [139-152]; P30115 [139-152] | [K].VLKSHGQDYLVGNR.[L] | 1xHomocysteineIAA [K3] | 4,32935E-05 | 1759,89081 | Liver/ FAIMS/IAA |
| P10648; P30115 | P10648 [139-152]; P30115 [139-152] | [K].VLKSHGQDYLVGNR.[L] | 1xHomocysteineMMTS [K3] | 4,49774E-05 | 1748,85707 | Liver/ No FAIMS/MMTS |
| P10648; P30115 | P10648 [139-152]; P30115 [139-152] | [K].VLKSHGQDYLVGNR.[L] | 1xHomocysteineMMTS [K3] | 4,49774E-05 | 1748,85707 | Liver/ FAIMS/MMTS |
| P10649 | P10649 [19-32] | [R].MLLEYTDSSYDEKR.[Y] | 1xHomocysteineIAA [K13] | 4,32935E-05 | 1923,84629 | Liver/ FAIMS/IAA |
| P10649 | P10649 [19-32] | [R].MLLEYTDSSYDEKR.[Y] | 1xHomocysteineIAA [K13] | 4,32935E-05 | 1923,84629 | Brain/ FAIMS/IAA |
| P10649 | P10649 [19-32] | [R].MLLEYTDSSYDEKR.[Y] | 1xOxidation [M1]; 1xHomocysteineMMTS [K13] | 4,49774E-05 | 1928,80746 | Liver/ No FAIMS/MMTS |
| P10649 | P10649 [123-134] | [K].QKPEFLKTIPEK.[M] | 1xHomocysteineMMTS [K7] | 0,00924953 | 1620,8488 | Liver/ No FAIMS/MMTS |
| P10649 | P10649 [44-52] | [R].SQWLNEKFK.[L] | 1xHomocysteineMMTS [K7] | 0,00367611 | 1342,62824 | Liver/ No FAIMS/MMTS |
| P10649 | P10649 [19-32] | [R].MLLEYTDSSYDEKR.[Y] | 1xHomocysteineMMTS [K13] | 4,49774E-05 | 1912,81255 | Liver/ No FAIMS/MMTS |
| P10649 | P10649 [83-94] | [R].KHHLDGETEEER.[I] | 1xHomocysteineMMTS [K1] | 4,49774E-05 | 1642,69481 | Liver/ No FAIMS/MMTS |
| P10649 | P10649 [44-52] | [R].SQWLNEKFK.[L] | 1xHomocysteineMMTS [K7] | 0,00367611 | 1342,62824 | Liver/ FAIMS/MMTS |
| P10649 | P10649 [19-32] | [R].MLLEYTDSSYDEKR.[Y] | 1xHomocysteineMMTS [K13] | 4,49774E-05 | 1912,81255 | Liver/ FAIMS/MMTS |
| P10649 | P10649 [19-32] | [R].MLLEYTDSSYDEKR.[Y] | 1xOxidation [M1]; 1xHomocysteineMMTS [K13] | 4,49774E-05 | 1928,80746 | Liver/ FAIMS/MMTS |
| P10649 | P10649 [123-134] | [K].QKPEFLKTIPEK.[M] | 1xHomocysteineMMTS [K7] | 0,00924953 | 1620,8488 | Liver/ FAIMS/MMTS |
| P10649 | P10649 [83-94] | [R].KHHLDGETEEER.[I] | 1xHomocysteineMMTS [K1] | 4,49774E-05 | 1642,69481 | Liver/ FAIMS/MMTS |
| P10649; P15626; P19639 | P10649 [135-144]; P15626 [135-144]; P19639 [135-144] | [K].MKLYSEFLGK.[RQ] | 1xHomocysteineIAA [K2] | 0,00012645 | 1389,69051 | Liver/ No FAIMS/IAA |
| P10649; P15626; P19639 | P10649 [135-144]; P15626 [135-144]; P19639 [135-144] | [K].MKLYSEFLGK.[RQ] | 1xHomocysteineIAA [K2] | 0,00012645 | 1389,69051 | Liver/ FAIMS/IAA |
| P10649; P15626; P19639 | P10649 [135-144]; P15626 [135-144]; P19639 [135-144] | [K].MKLYSEFLGK.[RQ] | 1xHomocysteineIAA [K2] | 0,00012645 | 1389,69051 | Brain/ No FAIMS/IAA |
| P10649; P15626; P19639 | P10649 [135-144]; P15626 [135-144]; P19639 [135-144] | [K].MKLYSEFLGK.[RQ] | 1xHomocysteineIAA [K2] | 0,00012645 | 1389,69051 | Brain/ FAIMS/IAA |
| P10649; P15626; P19639 | P10649 [135-144]; P15626 [135-144]; P19639 [135-144] | [K].MKLYSEFLGK.[RQ] | 1xOxidation [M1]; 1xHomocysteineMMTS [K2] | 0,000282457 | 1394,65168 | Liver/ No FAIMS/MMTS |
| P10649; P15626; P19639 | P10649 [135-144]; P15626 [135-144]; P19639 [135-144] | [K].MKLYSEFLGK.[RQ] | 1xHomocysteineMMTS [K2] | 4,49774E-05 | 1378,65676 | Liver/ No FAIMS/MMTS |

| | | | | | | |
|---|---|---|---|---|---|---|
| P10649; P15626; P19639 | P10649 [135-144]; P15626 [135-144]; P19639 [135-144] | [K].MKLYSEFLGK.[RQ] | 1xHomocysteineMMTS [K2] | 4,49774E-05 | 1378,65676 | Liver/ FAIMS/MMTS |
| P10649; P15626; P19639 | P10649 [135-144]; P15626 [135-144]; P19639 [135-144] | [K].MKLYSEFLGK.[RQ] | 1xOxidation [M1]; 1xHomocysteineMMTS [K2] | 0,000282457 | 1394,65168 | Liver/ FAIMS/MMTS |
| P10649; P19639 | P10649 [169-182]; P19639 [169-182] | [R].MFEPKCLDAFPNLR.[D] | 1xCarbamidomethyl [C6]; 1xOxidation [M1]; 1xHomocysteineIAA [K5] | 4,32935E-05 | 1927,88632 | Liver/ FAIMS/IAA |
| P10649; P19639 | P10649 [169-182]; P19639 [169-182] | [R].MFEPKCLDAFPNLR.[D] | 1xCarbamidomethyl [C6]; 1xOxidation [M1]; 1xHomocysteineIAA [K5] | 4,32935E-05 | 1927,88632 | Brain/ FAIMS/IAA |
| P10649; P19639 | P10649 [123-129]; P19639 [123-129] | [K].QKPEFLK.[TA] | 1xHomocysteineMMTS [K2] | 0,00107839 | 1052,52673 | Liver/ No FAIMS/MMTS |
| P10649; P19639 | P10649 [194-202]; P19639 [194-202] | [K].ISAYMKSSR.[YF] | 1xOxidation [M5]; 1xHomocysteineMMTS [K6] | 0,000530332 | 1221,54246 | Liver/ FAIMS/MMTS |
| P10649; P19639 | P10649 [123-129]; P19639 [123-129] | [K].QKPEFLK.[TA] | 1xHomocysteineMMTS [K2] | 0,00107839 | 1052,52673 | Liver/ FAIMS/MMTS |
| P10649; P19639; Q80W21 | P10649 [137-152]; P19639 [137-152]; Q80W21 [137-152] | [KR].LYSEFLGKRPWFAGDK.[VI] | 1xHomocysteineIAA [K8] | 4,32935E-05 | 2088,03714 | Liver/ FAIMS/IAA |
| P10649; P19639; Q80W21 | P10649 [137-152]; P19639 [137-152]; Q80W21 [137-152] | [KR].LYSEFLGKRPWFAGDK.[VI] | 1xHomocysteineIAA [K8] | 4,32935E-05 | 2088,03714 | Brain/ FAIMS/IAA |
| P10852 | P10852 [172-191] | [K].QINPTLGSQEDFKDLLQSAK.[K] | 1xHomocysteineIAA [K13] | 4,32935E-05 | 2406,19695 | Brain/ FAIMS/IAA |
| P10852 | P10852 [54-64] | [K].FTGLSKEELLK.[V] | 1xHomocysteineMMTS [K6] | 0,00367611 | 1427,72728 | Brain/ FAIMS/MMTS |
| P10852 | P10852 [160-171] | [K].NQKDEINETDLK.[Q] | 1xHomocysteineMMTS [K3] | 0,00074635 | 1609,71963 | Brain/ FAIMS/MMTS |
| P10852 | P10852 [224-242] | [K].MKEALSSWLQDGVDGFQFR.[D] | 1xHomocysteineMMTS [K2] | 0,00334305 | 2377,07737 | Brain/ FAIMS/MMTS |
| P10922 | P10922 [28-42] | [K].YSDMIVAAIQAEKNR.[A] | 1xHomocysteineIAA [K13] | 4,32935E-05 | 1882,91498 | Brain/ FAIMS/IAA |
| P11352 | P11352 [156-175] | [R].NDIAWNFEKFLVGPDGVPVR.[R] | 1xHomocysteineIAA [K9] | 0,000114608 | 2447,21763 | Liver/ FAIMS/IAA |
| P11352 | P11352 [53-63] | [R].DYTEMNDLQKR.[L] | 1xHomocysteineMMTS [K10] | 0,000152766 | 1575,66001 | Liver/ FAIMS/MMTS |
| P11404 | P11404 [32-45] | [R].QVASMTKPTTIIEK.[N] | 1xHomocysteineIAA [K7] | 4,32935E-05 | 1720,8972 | Brain/ FAIMS/IAA |
| P11404 | P11404 [81-91] | [K].VKSLVTLDGGK.[L] | 1xHomocysteineMMTS [K2] | 0,00325546 | 1279,67485 | Liver/ No FAIMS/MMTS |
| P11404 | P11404 [16-31] | [K].NFDDYMKSLGVGFATR.[Q] | 1xHomocysteineMMTS [K7] | 0,00122601 | 1983,87615 | Liver/ FAIMS/MMTS |
| P11404 | P11404 [32-45] | [R].QVASMTKPTTIIEK.[N] | 1xHomocysteineMMTS [K7] | 0,00876208 | 1709,86346 | Liver/ FAIMS/MMTS |
| P11404 | P11404 [108-127] | [R].ELVDGKLILTLTHGSVVSTR.[T] | 1xHomocysteineMMTS [K6] | 0,0014351 | 2301,2305 | Liver/ FAIMS/MMTS |
| P11404 | P11404 [11-22] | [K].LVDSKNFDDYMK.[S] | 1xHomocysteineMMTS [K5] | 0,00618488 | 1637,70081 | Liver/ FAIMS/MMTS |
| P11404 | P11404 [81-91] | [K].VKSLVTLDGGK.[L] | 1xHomocysteineMMTS [K2] | 0,00325546 | 1279,67485 | Liver/ FAIMS/MMTS |
| P11404 | P11404 [16-31] | [K].NFDDYMKSLGVGFATR.[Q] | 1xHomocysteineMMTS [K7] | 0,00122601 | 1983,87615 | Brain/ FAIMS/MMTS |
| P11404 | P11404 [32-45] | [R].QVASMTKPTTIIEK.[N] | 1xHomocysteineMMTS [K7] | 0,00876208 | 1709,86346 | Brain/ FAIMS/MMTS |
| P11438 | P11438 [141-148] | [K].ADINKAYR.[C] | 1xHomocysteineMMTS [K5] | 0,00885916 | 1113,51796 | Liver/ FAIMS/MMTS |
| P11499 | P11499 [436-448] | [K].NLKLGIHEDSTNR.[R] | 1xHomocysteineIAA [K3] | 4,32935E-05 | 1670,82788 | Liver/ FAIMS/IAA |
| P11499 | P11499 [624-639] | [K].KHLEINPDHPIVETLR.[Q] | 1xHomocysteineIAA [K1] | 4,32935E-05 | 2085,09097 | Brain/ No FAIMS/IAA |
| P11499 | P11499 [624-639] | [K].KHLEINPDHPIVETLR.[Q] | 1xHomocysteineIAA [K1] | 4,32935E-05 | 2085,09097 | Brain/ FAIMS/IAA |
| P11499 | P11499 [624-639] | [K].KHLEINPDHPIVETLR.[Q] | 1xHomocysteineMMTS [K1] | 4,49774E-05 | 2074,05723 | Liver/ No FAIMS/MMTS |
| P11499 | P11499 [527-538] | [K].EFDGKSLVSVTK.[E] | 1xHomocysteineMMTS [K5] | 0,0021186 | 1472,71236 | Liver/ FAIMS/MMTS |
| P11499 | P11499 [624-639] | [K].KHLEINPDHPIVETLR.[Q] | 1xHomocysteineMMTS [K1] | 4,49774E-05 | 2074,05723 | Liver/ FAIMS/MMTS |
| P11499 | P11499 [429-438] | [K].FYEAFSKNLK.[L] | 1xHomocysteineMMTS [K7] | 0,0004709 | 1409,6592 | Liver/ FAIMS/MMTS |
| P11499 | P11499 [478-491] | [K].ETQKSIYYITGESK.[E] | 1xHomocysteineMMTS [K4] | 0,00255629 | 1809,83975 | Liver/ FAIMS/MMTS |
| P11499 | P11499 [436-448] | [K].NLKLGIHEDSTNR.[R] | 1xHomocysteineMMTS [K3] | 0,000266738 | 1659,79413 | Liver/ FAIMS/MMTS |
| P11499 | P11499 [56-72] | [R].YESLTDPSKLDSGKELK.[I] | 1xHomocysteineMMTS [K14] | 0,00579131 | 2072,98787 | Liver/ FAIMS/MMTS |
| P11499 | P11499 [527-538] | [K].EFDGKSLVSVTK.[E] | 1xHomocysteineMMTS [K5] | 0,0021186 | 1472,71236 | Brain/ FAIMS/MMTS |
| P11499 | P11499 [478-491] | [K].ETQKSIYYITGESK.[E] | 1xHomocysteineMMTS [K4] | 0,00255629 | 1809,83975 | Brain/ FAIMS/MMTS |
| P11499; P08113 | P11499 [42-55]; P08113 [103-116] | [R].ELISNASDALDKIR.[YL] | 1xHomocysteineIAA [K12] | 4,32935E-05 | 1718,87416 | Liver/ No FAIMS/IAA |
| P11499; P08113 | P11499 [42-55]; P08113 [103-116] | [R].ELISNASDALDKIR.[YL] | 1xHomocysteineIAA [K12] | 4,32935E-05 | 1718,87416 | Liver/ FAIMS/IAA |
| P11499; P08113 | P11499 [42-55]; P08113 [103-116] | [R].ELISNASDALDKIR.[YL] | 1xHomocysteineMMTS [K12] | 4,49774E-05 | 1707,84042 | Liver/ No FAIMS/MMTS |
| P11499; P08113 | P11499 [42-55]; P08113 [103-116] | [R].ELISNASDALDKIR.[YL] | 1xHomocysteineMMTS [K12] | 4,49774E-05 | 1707,84042 | Liver/ FAIMS/MMTS |
| P11499; P08113 | P11499 [42-55]; P08113 [103-116] | [R].ELISNASDALDKIR.[YL] | 1xHomocysteineMMTS [K12] | 4,49774E-05 | 1707,84042 | Brain/ No FAIMS/MMTS |
| P11499; P08113 | P11499 [42-55]; P08113 [103-116] | [R].ELISNASDALDKIR.[YL] | 1xHomocysteineMMTS [K12] | 4,49774E-05 | 1707,84042 | Brain/ FAIMS/MMTS |
| P11588 | P11588 [152-163] | [R].FAQLCEKHGILR.[E] | 1xCarbamidomethyl [C5]; 1xHomocysteineIAA [K7] | 4,32935E-05 | 1645,83013 | Liver/ No FAIMS/IAA |
| P11588 | P11588 [152-163] | [R].FAQLCEKHGILR.[E] | 1xCarbamidomethyl [C5]; 1xHomocysteineIAA [K7] | 4,32935E-05 | 1645,83013 | Liver/ FAIMS/IAA |
| P11589 | P11589 [58-73] | [R].LFLEQIHVLEKSLVLK.[F] | 1xHomocysteineIAA [K11] | 4,32935E-05 | 2083,19839 | Liver/ No FAIMS/IAA |
| P11589 | P11589 [58-73] | [R].LFLEQIHVLEKSLVLK.[F] | 1xHomocysteineIAA [K11] | 4,32935E-05 | 2083,19839 | Liver/ FAIMS/IAA |
| P11589 | P11589 [69-78] | [K].SLVLKFHTVR.[D] | 1xHomocysteineMMTS [K5] | 0,000250158 | 1362,73846 | Liver/ No FAIMS/MMTS |
| P11589 | P11589 [58-73] | [R].LFLEQIHVLEKSLVLK.[F] | 1xHomocysteineMMTS [K11] | 4,49774E-05 | 2072,16465 | Liver/ No FAIMS/MMTS |
| P11589 | P11589 [58-73] | [R].LFLEQIHVLEKSLVLK.[F] | 1xHomocysteineMMTS [K11] | 4,49774E-05 | 2072,16465 | Liver/ FAIMS/MMTS |
| P11589 | P11589 [69-78] | [K].SLVLKFHTVR.[D] | 1xHomocysteineMMTS [K5] | 0,000250158 | 1362,73846 | Liver/ FAIMS/MMTS |
| P11589; P11588 | P11589 [74-94]; P11588 [74-94] | [K].FHTVRDEECSELSMVADKTEK.[A] | 1xCarbamidomethyl [C9]; 1xOxidation [M14]; 1xHomocysteineIAA [K18] | 4,32935E-05 | 2701,19023 | Liver/ FAIMS/IAA |
| P11589; P11588 | P11589 [74-94]; P11588 [74-94] | [K].FHTVRDEECSELSMVADKTEK.[A] | 1xMethylthio [C9]; 1xHomocysteineMMTS [K18] | 4,49774E-05 | 2663,12783 | Liver/ No FAIMS/MMTS |
| P11589; P11588 | P11589 [48-57]; P11588 [48-57] | [R].EKIEDNGNFR.[L] | 1xHomocysteineMMTS [K2] | 0,00107839 | 1384,5984 | Liver/ No FAIMS/MMTS |

| | | | | | | |
|---|---|---|---|---|---|---|
| P11589; P11588 | P11589 [74-94]; P11588 [74-94] | [K].FHTVRDEECSELSMVADKTEK.[A] | 1xMethylthio [C9]; 1xHomocysteineMMTS [K18] | 4,49774E-05 | 2663,12783 | Liver/ FAIMS/MMTS |
| P11589; P11588 | P11589 [48-57]; P11588 [48-57] | [R].EKIEDNGNFR.[L] | 1xHomocysteineMMTS [K2] | 0,00107839 | 1384,5984 | Liver/ FAIMS/MMTS |
| P11589; P11588 | P11589 [74-94]; P11588 [74-94] | [K].FHTVRDEECSELSMVADKTEK.[A] | 1xOxidation [M14]; 1xMethylthio [C9]; 1xHomocysteineMMTS [K18] | 6,70238E-05 | 2679,12274 | Liver/ FAIMS/MMTS |
| P11627 | P11627 [623-646] | [R].HLLKQSQVHLSWSPAEDHNSPIEK.[Y] | 1xHomocysteineIAA [K4] | 4,32935E-05 | 2954,45775 | Brain/ FAIMS/IAA |
| P11679 | P11679 [323-334] | [R].LQAEIEALKGQR.[A] | 1xHomocysteineIAA [K9] | 6,30912E-05 | 1529,81044 | Liver/ No FAIMS/IAA |
| P11679 | P11679 [323-334] | [R].LQAEIEALKGQR.[A] | 1xHomocysteineIAA [K9] | 6,30912E-05 | 1529,81044 | Liver/ FAIMS/IAA |
| P11679 | P11679 [323-334] | [R].LQAEIEALKGQR.[A] | 1xHomocysteineMMTS [K9] | 4,49774E-05 | 1518,77669 | Liver/ No FAIMS/MMTS |
| P11679 | P11679 [309-318] | [R].TKTEISEMNR.[N] | 1xHomocysteineMMTS [K2] | 0,00687493 | 1371,60652 | Liver/ No FAIMS/MMTS |
| P11679 | P11679 [103-114] | [K].SLNNKFASFIDK.[V] | 1xHomocysteineMMTS [K5] | 0,00115861 | 1546,73925 | Liver/ FAIMS/MMTS |
| P11679 | P11679 [323-334] | [R].LQAEIEALKGQR.[A] | 1xHomocysteineMMTS [K9] | 4,49774E-05 | 1518,77669 | Liver/ FAIMS/MMTS |
| P11679 | P11679 [309-318] | [R].TKTEISEMNR.[N] | 1xHomocysteineMMTS [K2] | 0,00687493 | 1371,60652 | Liver/ FAIMS/MMTS |
| P11679; Q9DCV7 | P11679 [117-128]; Q9DCV7 [105-116] | [R].FLEQQNKMLETK.[W] | 1xHomocysteineMMTS [K7] | 0,0024085 | 1671,7903 | Liver/ FAIMS/MMTS |
| P11725 | P11725 [142-153] | [R].VYKQSDLDTLAK.[E] | 1xHomocysteineIAA [K3] | 4,32935E-05 | 1554,78322 | Liver/ No FAIMS/IAA |
| P11725 | P11725 [142-153] | [R].VYKQSDLDTLAK.[E] | 1xHomocysteineIAA [K3] | 4,32935E-05 | 1554,78322 | Liver/ FAIMS/IAA |
| P11725 | P11725 [81-89] | [K].SLGMIFEKR.[S] | 1xOxidation [M4]; 1xHomocysteineMMTS [K8] | 0,00501021 | 1259,5945 | Liver/ No FAIMS/MMTS |
| P11725 | P11725 [81-89] | [K].SLGMIFEKR.[S] | 1xHomocysteineMMTS [K8] | 0,000489972 | 1243,59958 | Liver/ No FAIMS/MMTS |
| P11725 | P11725 [142-153] | [R].VYKQSDLDTLAK.[E] | 1xHomocysteineMMTS [K3] | 4,49774E-05 | 1543,74948 | Liver/ No FAIMS/MMTS |
| P11725 | P11725 [69-80] | [K].QKGEYLPLLQGK.[S] | 1xHomocysteineMMTS [K2] | 0,00843159 | 1536,79128 | Liver/ No FAIMS/MMTS |
| P11725 | P11725 [256-275] | [R].GGNVLITDTWISMGQEDEKK.[K] | 1xHomocysteineMMTS [K19] | 4,49774E-05 | 2384,09308 | Liver/ No FAIMS/MMTS |
| P11725 | P11725 [222-238] | [K].GYEPDPNIVKLAEQYAK.[E] | 1xHomocysteineMMTS [K10] | 4,49774E-05 | 2097,99837 | Liver/ No FAIMS/MMTS |
| P11725 | P11725 [307-320] | [R].KPEEVDDEVFYSPR.[S] | 1xHomocysteineMMTS [K1] | 4,49774E-05 | 1872,81426 | Liver/ No FAIMS/MMTS |
| P11725 | P11725 [239-255] | [K].ENGTKLSMTNDPLEAAR.[G] | 1xHomocysteineMMTS [K5] | 0,00905461 | 2009,90891 | Liver/ FAIMS/MMTS |
| P11725 | P11725 [142-153] | [R].VYKQSDLDTLAK.[E] | 1xHomocysteineMMTS [K3] | 4,49774E-05 | 1543,74948 | Liver/ FAIMS/MMTS |
| P11725 | P11725 [222-238] | [K].GYEPDPNIVKLAEQYAK.[E] | 1xHomocysteineMMTS [K10] | 4,49774E-05 | 2097,99837 | Liver/ FAIMS/MMTS |
| P11725 | P11725 [81-89] | [K].SLGMIFEKR.[S] | 1xOxidation [M4]; 1xHomocysteineMMTS [K8] | 0,00501021 | 1259,5945 | Liver/ FAIMS/MMTS |
| P11725 | P11725 [81-89] | [K].SLGMIFEKR.[S] | 1xHomocysteineMMTS [K8] | 0,000489972 | 1243,59958 | Liver/ FAIMS/MMTS |
| P11725 | P11725 [232-243] | [K].LAEQYAKENGTK.[L] | 1xHomocysteineMMTS [K7] | 0,00738083 | 1514,69778 | Liver/ FAIMS/MMTS |
| P11725 | P11725 [256-275] | [R].GGNVLITDTWISMGQEDEKK.[K] | 1xHomocysteineMMTS [K19] | 4,49774E-05 | 2384,09308 | Liver/ FAIMS/MMTS |
| P11725 | P11725 [278-292] | [R].LQAFQGYQVTMKTAK.[V] | 1xHomocysteineMMTS [K12] | 4,49774E-05 | 1876,91181 | Liver/ FAIMS/MMTS |
| P11725 | P11725 [307-320] | [R].KPEEVDDEVFYSPR.[S] | 1xHomocysteineMMTS [K1] | 4,49774E-05 | 1872,81426 | Liver/ FAIMS/MMTS |
| P11798 | P11798 [260-274] | [R].ITAAEALKHPWISHR.[S] | 1xHomocysteineIAA [K8] | 4,32935E-05 | 1903,99595 | Brain/ No FAIMS/IAA |
| P11798 | P11798 [260-274] | [R].ITAAEALKHPWISHR.[S] | 1xHomocysteineIAA [K8] | 4,32935E-05 | 1903,99595 | Brain/ FAIMS/IAA |
| P11798 | P11798 [246-258] | [K].DLINKMLTINPSK.[R] | 1xHomocysteineMMTS [K5] | 0,00312202 | 1649,84233 | Brain/ FAIMS/MMTS |
| P11798; P28652; Q6PHZ2 | P11798 [299-311]; P28652 [300-312]; Q6PHZ2 [300-312] | [K].LKGAILTTMLATR.[N] | 1xHomocysteineIAA [K2] | 4,32935E-05 | 1562,87568 | Brain/ No FAIMS/IAA |
| P11798; P28652; Q6PHZ2 | P11798 [299-311]; P28652 [300-312]; Q6PHZ2 [300-312] | [K].LKGAILTTMLATR.[N] | 1xHomocysteineIAA [K2] | 4,32935E-05 | 1562,87568 | Brain/ FAIMS/IAA |
| P11798; P28652; Q923T9; Q6PHZ2 | P11798 [221-245]; P28652 [222-246]; Q923T9 [222-246]; Q6PHZ2 [222-246] | [RK].LYQQIKAGAYDFPSPEWDTVTPEAK.[DN] | 1xHomocysteineIAA [K6] | 4,32935E-05 | 3028,4397 | Brain/ FAIMS/IAA |
| P11798; P28652; Q923T9; Q6PHZ2 | P11798 [135-146]; P28652 [136-147]; Q923T9 [136-147]; Q6PHZ2 [136-147] | [R].DLKPENLLLASK.[LCS] | 1xHomocysteineMMTS [K3] | 0,000211896 | 1503,79095 | Brain/ FAIMS/MMTS |
| P11798; Q6PHZ2 | P11798 [227-250]; Q6PHZ2 [228-251] | [K].AGAYDFPSPEWDTVTPEAKDLINK.[M] | 1xHomocysteineIAA [K19] | 4,32935E-05 | 2838,32909 | Brain/ FAIMS/IAA |
| P11798; Q923T9; Q6PHZ2 | P11798 [66-74]; Q923T9 [67-75]; Q6PHZ2 [67-75] | [R].LLKHPNIVR.[L] | 1xHomocysteineIAA [K3] | 9,61721E-05 | 1263,73542 | Brain/ No FAIMS/IAA |
| P11798; Q923T9; Q6PHZ2 | P11798 [66-74]; Q923T9 [67-75]; Q6PHZ2 [67-75] | [R].LLKHPNIVR.[L] | 1xHomocysteineIAA [K3] | 9,61721E-05 | 1263,73542 | Brain/ FAIMS/IAA |
| P11983 | P11983 [391-401] | [R].SLHDALCVVKR.[V] | 1xCarbamidomethyl [C7]; 1xHomocysteineIAA [K10] | 4,32935E-05 | 1471,75081 | Brain/ FAIMS/IAA |
| P12023 | P12023 [378-395] | [K].YLETPGDENEHAHFQKAK.[E] | 1xHomocysteineIAA [K16] | 0,00012645 | 2288,04006 | Brain/ FAIMS/IAA |
| P12658 | P12658 [109-124] | [R].KYDTDHSGFIETEELK.[N] | 1xHomocysteineIAA [K1] | 4,32935E-05 | 2085,94336 | Brain/ No FAIMS/IAA |
| P12658 | P12658 [109-124] | [R].KYDTDHSGFIETEELK.[N] | 1xHomocysteineIAA [K1] | 4,32935E-05 | 2085,94336 | Brain/ FAIMS/IAA |
| P12658 | P12658 [153-169] | [K].LFDSNNDGKLELTEMAR.[L] | 1xHomocysteineMMTS [K9] | 0,000173718 | 2115,95077 | Brain/ FAIMS/MMTS |
| P12658 | P12658 [137-152] | [K].TVDDTKLAEYTDLMLK.[L] | 1xHomocysteineMMTS [K6] | 4,49774E-05 | 2018,94831 | Brain/ FAIMS/MMTS |
| P12658 | P12658 [48-59] | [R].KKAGLELSPEMK.[S] | 1xHomocysteineMMTS [K2] | 0,00729558 | 1493,75245 | Brain/ FAIMS/MMTS |
| P12658 | P12658 [50-68] | [K].AGLELSPEMKSFVDQYGQR.[D] | 1xHomocysteineMMTS [K10] | 4,49774E-05 | 2318,06138 | Brain/ FAIMS/MMTS |
| P12658 | P12658 [109-124] | [R].KYDTDHSGFIETEELK.[N] | 1xHomocysteineMMTS [K1] | 0,00576292 | 2074,90962 | Brain/ FAIMS/MMTS |
| P12710 | P12710 [47-57] | [K].KIKLTITYGPK.[V] | 1xHomocysteineIAA [K3] | 4,32935E-05 | 1435,83413 | Liver/ FAIMS/IAA |
| P12710 | P12710 [97-121] | [K].GIKSVTELNGDTITNTMTLGDIVYK.[R] | 1xHomocysteineMMTS [K3] | 4,49774E-05 | 2846,39843 | Liver/ No FAIMS/MMTS |
| P12710 | P12710 [97-121] | [K].GIKSVTELNGDTITNTMTLGDIVYK.[R] | 1xOxidation [M17]; 1xHomocysteineMMTS [K3] | 4,49774E-05 | 2862,39334 | Liver/ No FAIMS/MMTS |
| P12710 | P12710 [34-47] | [K].DIKGVSEIVHEGKK.[I] | 2xHomocysteineMMTS [K3; K13] | 4,49774E-05 | 1864,87879 | Liver/ No FAIMS/MMTS |
| P12710 | P12710 [21-33] | [K].AIGLPEDLIQKGK.[D] | 2xHomocysteineMMTS [K11; K13] | 0,00272554 | 1707,83005 | Liver/ No FAIMS/MMTS |
| P12710 | P12710 [37-49] | [K].GVSEIVHEGKKIK.[L] | 2xHomocysteineMMTS [K10; K11] | 0,00328452 | 1749,85185 | Liver/ No FAIMS/MMTS |

| | | | | | | |
|---|---|---|---|---|---|---|
| P12710 | P12710 [37-47] | [K].GVSEIVHEGKK.[I] | 2xHomocysteineMMTS [K10; K11] | 0,000829616 | 1508,67282 | Liver/ No FAIMS/MMTS |
| P12710 | P12710 [85-96] | [K].LEGDNKMVTTFK.[G] | 1xOxidation [M7]; 1xHomocysteineMMTS [K6] | 4,49774E-05 | 1561,7059 | Liver/ No FAIMS/MMTS |
| P12710 | P12710 [100-122] | [K].SVTELNGDTITNTMTLGDIVYKR.[V] | 1xOxidation [M14]; 1xHomocysteineMMTS [K22] | 4,49774E-05 | 2720,29396 | Liver/ No FAIMS/MMTS |
| P12710 | P12710 [7-31] | [K].YQLQSQENFEPFMKAIGLPEDLIQK.[G] | 1xOxidation [M13]; 1xHomocysteineMMTS [K14] | 4,49774E-05 | 3145,50429 | Liver/ No FAIMS/MMTS |
| P12710 | P12710 [61-80] | [R].NEFTLGEECELETMTGEKVK.[A] | 1xMethylthio [C9]; 1xHomocysteineMMTS [K18] | 4,49774E-05 | 2496,04712 | Liver/ No FAIMS/MMTS |
| P12710 | P12710 [50-60] | [K].LTITYGPKVVR.[N] | 1xHomocysteineMMTS [K8] | 4,49774E-05 | 1409,76434 | Liver/ No FAIMS/MMTS |
| P12710 | P12710 [85-96] | [K].LEGDNKMVTTFK.[G] | 1xHomocysteineMMTS [K6] | 4,49774E-05 | 1545,71098 | Liver/ No FAIMS/MMTS |
| P12710 | P12710 [32-46] | [K].GKDIKGVSEIVHEGK.[K] | 1xHomocysteineMMTS [K5] | 4,49774E-05 | 1758,8877 | Liver/ No FAIMS/MMTS |
| P12710 | P12710 [81-90] | [K].AVVKLEGDNK.[M] | 1xHomocysteineMMTS [K4] | 4,49774E-05 | 1235,61225 | Liver/ No FAIMS/MMTS |
| P12710 | P12710 [34-46] | [K].DIKGVSEIVHEGK.[K] | 1xHomocysteineMMTS [K3] | 4,49774E-05 | 1573,77127 | Liver/ No FAIMS/MMTS |
| P12710 | P12710 [34-47] | [K].DIKGVSEIVHEGKK.[I] | 1xHomocysteineMMTS [K3] | 4,49774E-05 | 1701,86624 | Liver/ No FAIMS/MMTS |
| P12710 | P12710 [97-122] | [K].GIKSVTELNGDTITNTMTLGDIVYKR.[V] | 1xHomocysteineMMTS [K3] | 4,49774E-05 | 3002,49954 | Liver/ No FAIMS/MMTS |
| P12710 | P12710 [100-122] | [K].SVTELNGDTITNTMTLGDIVYKR.[V] | 1xHomocysteineMMTS [K22] | 4,49774E-05 | 2704,29905 | Liver/ No FAIMS/MMTS |
| P12710 | P12710 [48-57] | [K].IKLTITYGPK.[V] | 1xHomocysteineMMTS [K2] | 6,70238E-05 | 1296,70543 | Liver/ No FAIMS/MMTS |
| P12710 | P12710 [7-31] | [K].YQLQSQENFEPFMKAIGLPEDLIQK.[G] | 1xHomocysteineMMTS [K14] | 4,49774E-05 | 3129,50938 | Liver/ No FAIMS/MMTS |
| P12710 | P12710 [21-33] | [K].AIGLPEDLIQKGK.[D] | 1xHomocysteineMMTS [K11] | 4,49774E-05 | 1544,8175 | Liver/ No FAIMS/MMTS |
| P12710 | P12710 [48-60] | [K].IKLTITYGPKVVR.[N] | 1xHomocysteineMMTS [K10] | 0,000400718 | 1650,94337 | Liver/ No FAIMS/MMTS |
| P12710 | P12710 [37-47] | [K].GVSEIVHEGKK.[I] | 1xHomocysteineMMTS [K] | 4,49774E-05 | 1345,66027 | Liver/ No FAIMS/MMTS |
| P12710 | P12710 [91-99] | [K].MVTTFKGIK.[S] | 1xHomocysteineMMTS [K] | 0,000250158 | 1187,59852 | Liver/ No FAIMS/MMTS |
| P12710 | P12710 [1-20] | [-].MNFSGKYQLQSQENFEPFMK.[A] | 1xAcetyl [N-Term]; 1xOxidation [M19]; 1xHomocysteineMMTS [K6] | 4,49774E-05 | 2674,14446 | Liver/ No FAIMS/MMTS |
| P12710 | P12710 [1-20] | [-].MNFSGKYQLQSQENFEPFMK.[A] | 1xAcetyl [N-Term]; 1xHomocysteineMMTS [K6] | 4,49774E-05 | 2658,14955 | Liver/ No FAIMS/MMTS |
| P12710 | P12710 [21-33] | [K].AIGLPEDLIQKGK.[D] | 2xHomocysteineMMTS [K11; K13] | 0,00272554 | 1707,83005 | Liver/ FAIMS/MMTS |
| P12710 | P12710 [85-96] | [K].LEGDNKMVTTFK.[G] | 1xOxidation [M7]; 1xHomocysteineMMTS [K6] | 4,49774E-05 | 1561,7059 | Liver/ FAIMS/MMTS |
| P12710 | P12710 [97-121] | [K].GIKSVTELNGDTITNTMTLGDIVYK.[R] | 1xOxidation [M17]; 1xHomocysteineMMTS [K3] | 4,49774E-05 | 2862,39334 | Liver/ FAIMS/MMTS |
| P12710 | P12710 [61-80] | [R].NEFTLGEECELETMTGEKVK.[A] | 1xOxidation [M14]; 1xMethylthio [C9]; 1xHomocysteineMMTS [K18] | 4,49774E-05 | 2512,04203 | Liver/ FAIMS/MMTS |
| P12710 | P12710 [100-122] | [K].SVTELNGDTITNTMTLGDIVYKR.[V] | 1xOxidation [M14]; 1xHomocysteineMMTS [K22] | 4,49774E-05 | 2720,29396 | Liver/ FAIMS/MMTS |
| P12710 | P12710 [7-31] | [K].YQLQSQENFEPFMKAIGLPEDLIQK.[G] | 1xOxidation [M13]; 1xHomocysteineMMTS [K14] | 4,49774E-05 | 3145,50429 | Liver/ FAIMS/MMTS |
| P12710 | P12710 [61-80] | [R].NEFTLGEECELETMTGEKVK.[A] | 1xMethylthio [C9]; 1xHomocysteineMMTS [K18] | 4,49774E-05 | 2496,04712 | Liver/ FAIMS/MMTS |
| P12710 | P12710 [50-60] | [K].LTITYGPKVVR.[N] | 1xHomocysteineMMTS [K8] | 4,49774E-05 | 1409,76434 | Liver/ FAIMS/MMTS |
| P12710 | P12710 [85-96] | [K].LEGDNKMVTTFK.[G] | 1xHomocysteineMMTS [K6] | 4,49774E-05 | 1545,71098 | Liver/ FAIMS/MMTS |
| P12710 | P12710 [32-46] | [K].GKDIKGVSEIVHEGK.[K] | 1xHomocysteineMMTS [K5] | 4,49774E-05 | 1758,8877 | Liver/ FAIMS/MMTS |
| P12710 | P12710 [81-90] | [K].AVVKLEGDNK.[M] | 1xHomocysteineMMTS [K4] | 4,49774E-05 | 1235,61225 | Liver/ FAIMS/MMTS |
| P12710 | P12710 [34-46] | [K].DIKGVSEIVHEGK.[K] | 1xHomocysteineMMTS [K3] | 4,49774E-05 | 1573,77127 | Liver/ FAIMS/MMTS |
| P12710 | P12710 [34-47] | [K].DIKGVSEIVHEGKK.[I] | 1xHomocysteineMMTS [K3] | 4,49774E-05 | 1701,86624 | Liver/ FAIMS/MMTS |
| P12710 | P12710 [97-121] | [K].GIKSVTELNGDTITNTMTLGDIVYK.[R] | 1xHomocysteineMMTS [K3] | 4,49774E-05 | 2846,39843 | Liver/ FAIMS/MMTS |
| P12710 | P12710 [100-122] | [K].SVTELNGDTITNTMTLGDIVYKR.[V] | 1xHomocysteineMMTS [K22] | 4,49774E-05 | 2704,29905 | Liver/ FAIMS/MMTS |
| P12710 | P12710 [48-57] | [K].IKLTITYGPK.[V] | 1xHomocysteineMMTS [K2] | 6,70238E-05 | 1296,70543 | Liver/ FAIMS/MMTS |
| P12710 | P12710 [7-31] | [K].YQLQSQENFEPFMKAIGLPEDLIQK.[G] | 1xHomocysteineMMTS [K14] | 4,49774E-05 | 3129,50938 | Liver/ FAIMS/MMTS |
| P12710 | P12710 [21-33] | [K].AIGLPEDLIQKGK.[D] | 1xHomocysteineMMTS [K11] | 4,49774E-05 | 1544,8175 | Liver/ FAIMS/MMTS |
| P12710 | P12710 [81-96] | [K].AVVKLEGDNKMVTTFK.[G] | 1xHomocysteineMMTS [K10] | 4,49774E-05 | 1942,97989 | Liver/ FAIMS/MMTS |
| P12710 | P12710 [37-47] | [K].GVSEIVHEGKK.[I] | 1xHomocysteineMMTS [K] | 4,49774E-05 | 1345,66027 | Liver/ FAIMS/MMTS |
| P12710 | P12710 [91-99] | [K].MVTTFKGIK.[S] | 1xHomocysteineMMTS [K] | 0,000250158 | 1187,59852 | Liver/ FAIMS/MMTS |
| P12710 | P12710 [1-20] | [-].MNFSGKYQLQSQENFEPFMK.[A] | 1xAcetyl [N-Term]; 1xHomocysteineMMTS [K6] | 4,49774E-05 | 2658,14955 | Liver/ FAIMS/MMTS |
| P12710 | P12710 [34-47] | [K].DIKGVSEIVHEGKK.[I] | 2xHomocysteineMMTS [K3; K13] | 4,49774E-05 | 1864,87879 | Liver/ FAIMS/MMTS |
| P12710 | P12710 [32-46] | [K].GKDIKGVSEIVHEGK.[K] | 2xHomocysteineMMTS [K2; K5] | 8,93346E-05 | 1921,90026 | Liver/ FAIMS/MMTS |
| P12710 | P12710 [37-49] | [K].GVSEIVHEGKKIK.[L] | 2xHomocysteineMMTS [K10; K11] | 0,00328452 | 1749,85185 | Liver/ FAIMS/MMTS |
| P12710 | P12710 [37-47] | [K].GVSEIVHEGKK.[I] | 2xHomocysteineMMTS [K10; K11] | 0,000829616 | 1508,67282 | Liver/ FAIMS/MMTS |
| P12710 | P12710 [91-99] | [K].MVTTFKGIK.[S] | 1xOxidation [M1]; 1xHomocysteineMMTS [K6] | 0,00624351 | 1203,59343 | Liver/ FAIMS/MMTS |
| P12710 | P12710 [1-20] | [-].MNFSGKYQLQSQENFEPFMK.[A] | 1xHomocysteineMMTS [K6] | 4,49774E-05 | 2616,13898 | Liver/ FAIMS/MMTS |
| P12710 | P12710 [47-57] | [K].KIKLTITYGPK.[V] | 1xHomocysteineMMTS [K3] | 0,00367611 | 1424,80039 | Liver/ FAIMS/MMTS |
| P12710 | P12710 [97-122] | [K].GIKSVTELNGDTITNTMTLGDIVYKR.[V] | 1xHomocysteineMMTS [K3] | 4,49774E-05 | 3002,49954 | Liver/ FAIMS/MMTS |
| P12710 | P12710 [48-60] | [K].IKLTITYGPKVVR.[N] | 1xHomocysteineMMTS [K10] | 0,000400718 | 1650,94337 | Liver/ FAIMS/MMTS |
| P12710 | P12710 [1-20] | [-].MNFSGKYQLQSQENFEPFMK.[A] | 1xAcetyl [N-Term]; 1xOxidation [M19]; 1xHomocysteineMMTS [K6] | 4,49774E-05 | 2674,14446 | Liver/ FAIMS/MMTS |
| P12787 | P12787 [68-83] | [R].KGMNTLVGYDLVPEPK.[I] | 1xHomocysteineIAA [K1] | 4,32935E-05 | 1934,97143 | Liver/ FAIMS/IAA |
| P12787 | P12787 [68-83] | [R].KGMNTLVGYDLVPEPK.[I] | 1xHomocysteineIAA [K1] | 4,32935E-05 | 1934,97143 | Brain/ No FAIMS/IAA |

| | | | | | | |
|---|---|---|---|---|---|---|
| P12787 | P12787 [68-83] | [R].KGMNTLVGYDLVPEPK.[I] | 1xHomocysteineIAA [K1] | 4,32935E-05 | 1934,97143 | Brain/ FAIMS/IAA |
| | | | 1xOxidation [M2]; | | | |
| P12787 | P12787 [69-90] | [K].GMNTLVGYDLVPEPKIIDAALR.[A] | 1xHomocysteineMMTS [K15] | 4,49774E-05 | 2564,29211 | Liver/ No FAIMS/MMTS |
| P12787 | P12787 [52-67] | [R].WVTYFNKPDIDAWELR.[K] | 1xHomocysteineMMTS [K7] | 4,49774E-05 | 2216,03034 | Liver/ No FAIMS/MMTS |
| P12787 | P12787 [68-83] | [R].KGMNTLVGYDLVPEPK.[I] | 1xHomocysteineMMTS [K1] | 4,49774E-05 | 1923,93769 | Liver/ No FAIMS/MMTS |
| | | | 1xOxidation [M2]; | | | |
| P12787 | P12787 [69-90] | [K].GMNTLVGYDLVPEPKIIDAALR.[A] | 1xHomocysteineMMTS [K15] | 4,49774E-05 | 2564,29211 | Liver/ FAIMS/MMTS |
| P12787 | P12787 [52-67] | [R].WVTYFNKPDIDAWELR.[K] | 1xHomocysteineMMTS [K7] | 4,49774E-05 | 2216,03034 | Liver/ FAIMS/MMTS |
| P12787 | P12787 [104-111] | [R].ILEVVKDK.[A] | 1xHomocysteineMMTS [K6] | 0,000829616 | 1106,59481 | Liver/ FAIMS/MMTS |
| P12787 | P12787 [69-90] | [K].GMNTLVGYDLVPEPKIIDAALR.[A] | 1xHomocysteineMMTS [K15] | 4,49774E-05 | 2548,2972 | Liver/ FAIMS/MMTS |
| P12787 | P12787 [68-83] | [R].KGMNTLVGYDLVPEPK.[I] | 1xHomocysteineMMTS [K1] | 4,49774E-05 | 1923,93769 | Liver/ FAIMS/MMTS |
| | | | 1xOxidation [M3]; | | | |
| P12787 | P12787 [68-83] | [R].KGMNTLVGYDLVPEPK.[I] | 1xHomocysteineMMTS [K1] | 4,49774E-05 | 1939,9326 | Liver/ FAIMS/MMTS |
| P12787 | P12787 [52-67] | [R].WVTYFNKPDIDAWELR.[K] | 1xHomocysteineMMTS [K7] | 4,49774E-05 | 2216,03034 | Brain/ No FAIMS/MMTS |
| P12787 | P12787 [104-111] | [R].ILEVVKDK.[A] | 1xHomocysteineMMTS [K6] | 0,000829616 | 1106,59481 | Brain/ No FAIMS/MMTS |
| P12787 | P12787 [68-83] | [R].KGMNTLVGYDLVPEPK.[I] | 1xHomocysteineMMTS [K1] | 4,49774E-05 | 1923,93769 | Brain/ No FAIMS/MMTS |
| P12787 | P12787 [52-67] | [R].WVTYFNKPDIDAWELR.[K] | 1xHomocysteineMMTS [K7] | 4,49774E-05 | 2216,03034 | Brain/ FAIMS/MMTS |
| P12787 | P12787 [104-111] | [R].ILEVVKDK.[A] | 1xHomocysteineMMTS [K6] | 0,000829616 | 1106,59481 | Brain/ FAIMS/MMTS |
| P12787 | P12787 [69-90] | [K].GMNTLVGYDLVPEPKIIDAALR.[A] | 1xHomocysteineMMTS [K15] | 4,49774E-05 | 2548,2972 | Brain/ FAIMS/MMTS |
| P12787 | P12787 [68-83] | [R].KGMNTLVGYDLVPEPK.[I] | 1xHomocysteineMMTS [K1] | 4,49774E-05 | 1923,93769 | Brain/ FAIMS/MMTS |
| P12960 | P12960 [277-296] | [R].KVLEPMPSTAEISTSGAVLK.[I] | 1xHomocysteineIAA [K1] | 4,32935E-05 | 2232,16142 | Brain/ No FAIMS/IAA |
| P12960 | P12960 [701-723] | [R].IKTDGAAPNVAPSDVGGGGGTNR.[E] | 1xHomocysteineMMTS [K2] | 4,49774E-05 | 2274,06014 | Brain/ FAIMS/MMTS |
| P12970 | P12970 [11-20] | [K].KVAPAPAVVK.[K] | 1xHomocysteineIAA [K1] | 0,000741565 | 1153,67618 | Brain/ FAIMS/IAA |
| P12970 | P12970 [122-131] | [K].AAGKGDVPTK.[R] | 1xHomocysteineMMTS [K4] | 0,00248361 | 1106,53328 | Liver/ No FAIMS/MMTS |
| P12970 | P12970 [12-21] | [K].VAPAPAVVKK.[Q] | 1xHomocysteineMMTS [K9] | 0,000282457 | 1142,64243 | Liver/ FAIMS/MMTS |
| P12970 | P12970 [122-131] | [K].AAGKGDVPTK.[R] | 1xHomocysteineMMTS [K4] | 0,00248361 | 1106,53328 | Liver/ FAIMS/MMTS |
| P12970 | P12970 [38-49] | [K].NFGIGQDIQPKR.[D] | 1xHomocysteineMMTS [K11] | 0,00427174 | 1535,74573 | Liver/ FAIMS/MMTS |
| P12970 | P12970 [236-249] | [R].HWGGNVLGPKSVAR.[I] | 1xHomocysteineMMTS [K10] | 0,00107839 | 1640,81481 | Liver/ FAIMS/MMTS |
| P12970 | P12970 [11-20] | [K].KVAPAPAVVK.[K] | 1xHomocysteineMMTS [K1] | 4,49774E-05 | 1142,64243 | Liver/ FAIMS/MMTS |
| P13020 | P13020 [614-624] | [K].TGAQELLKVLR.[S] | 1xHomocysteineIAA [K8] | 0,00252218 | 1401,78825 | Brain/ FAIMS/IAA |
| | | | 1xCarbamidomethyl [C12]; | | | |
| P13595 | P13595 [277-293] | [R].NVDKNDEAEYVCIAENK.[A] | 1xHomocysteineIAA [K4] | 4,32935E-05 | 2184,95361 | Brain/ No FAIMS/IAA |
| | | | 1xCarbamidomethyl [C12]; | | | |
| P13595 | P13595 [277-293] | [R].NVDKNDEAEYVCIAENK.[A] | 1xHomocysteineIAA [K4] | 4,32935E-05 | 2184,95361 | Brain/ FAIMS/IAA |
| P13595 | P13595 [620-633] | [K].VNLIKQDDGGSPIR.[H] | 1xHomocysteineMMTS [K5] | 4,49774E-05 | 1674,83019 | Brain/ No FAIMS/MMTS |
| P13595 | P13595 [555-572] | [K].EANMEGIVTIMGLKPETR.[Y] | 1xHomocysteineMMTS [K14] | 4,49774E-05 | 2152,02691 | Brain/ No FAIMS/MMTS |
| P13595 | P13595 [1084-1099] | [K].SEPPESEAKPAPTEVK.[T] | 1xHomocysteineMMTS [K9] | 0,00164853 | 1858,85613 | Brain/ FAIMS/MMTS |
| P13595 | P13595 [641-651] | [R].ALASEWKPEIR.[L] | 1xHomocysteineMMTS [K7] | 0,000530332 | 1462,71812 | Brain/ FAIMS/MMTS |
| P13595 | P13595 [620-633] | [K].VNLIKQDDGGSPIR.[H] | 1xHomocysteineMMTS [K5] | 4,49774E-05 | 1674,83019 | Brain/ FAIMS/MMTS |
| P13595 | P13595 [118-130] | [K].LMFKNAPTPQEFK.[E] | 1xHomocysteineMMTS [K4] | 0,0004709 | 1713,81612 | Brain/ FAIMS/MMTS |
| P13595 | P13595 [250-276] | [K].DGEPIENEEEDDEKHIFSDDSSELTIR.[N] | 1xHomocysteineMMTS [K14] | 4,49774E-05 | 3311,38384 | Brain/ FAIMS/MMTS |
| P13595 | P13595 [607-624] | [K].LEGQMGEDGNSIKVNLIK.[Q] | 1xHomocysteineMMTS [K13] | 4,49774E-05 | 2108,01846 | Brain/ FAIMS/MMTS |
| P13595 | P13595 [584-600] | [K].GLGEISAATEFKTQPVR.[E] | 1xHomocysteineMMTS [K12] | 4,49774E-05 | 1966,97249 | Brain/ FAIMS/MMTS |
| P13707 | P13707 [49-62] | [R].KLTEIINTQHENVK.[Y] | 1xHomocysteineIAA [K1] | 0,000988467 | 1840,95856 | Brain/ FAIMS/IAA |
| | | | 1xCarbamidomethyl [C8]; | | | |
| P14094 | P14094 [119-136] | [K].DDMIFEDCGNVPSEPKER.[G] | 1xHomocysteineIAA [K16] | 4,32935E-05 | 2311,96279 | Brain/ No FAIMS/IAA |
| | | | 1xCarbamidomethyl [C8]; | | | |
| P14094 | P14094 [119-136] | [K].DDMIFEDCGNVPSEPKER.[G] | 1xHomocysteineIAA [K16] | 4,32935E-05 | 2311,96279 | Brain/ FAIMS/IAA |
| P14094 | P14094 [72-96] | [R].VAPPGLTQIPQIQKTEISFRPNDPK.[S] | 1xHomocysteineMMTS [K14] | 4,49774E-05 | 2937,53249 | Brain/ No FAIMS/MMTS |
| P14094 | P14094 [279-291] | [K].AYGENIGYSEKDR.[F] | 1xHomocysteineMMTS [K11] | 4,49774E-05 | 1664,70432 | Brain/ No FAIMS/MMTS |
| P14094 | P14094 [72-96] | [R].VAPPGLTQIPQIQKTEISFRPNDPK.[S] | 1xHomocysteineMMTS [K14] | 4,49774E-05 | 2937,53249 | Brain/ FAIMS/MMTS |
| P14094 | P14094 [279-291] | [K].AYGENIGYSEKDR.[F] | 1xHomocysteineMMTS [K11] | 4,49774E-05 | 1664,70432 | Brain/ FAIMS/MMTS |
| P14094 | P14094 [279-295] | [K].AYGENIGYSEKDRFQGR.[F] | 1xHomocysteineMMTS [K11] | 0,000133296 | 2152,95388 | Brain/ FAIMS/MMTS |
| P14094 | P14094 [22-27] | [K].KEFLGR.[T] | 1xHomocysteineMMTS [K1] | 0,00534072 | 912,443 | Brain/ FAIMS/MMTS |
| P14115 | P14115 [93-105] | [K].NKTGVAPIIDVVR.[S] | 1xHomocysteineMMTS [K2] | 0,00202818 | 1544,82873 | Liver/ No FAIMS/MMTS |
| P14115 | P14115 [43-55] | [R].INFDKYHPGYFGK.[V] | 1xHomocysteineMMTS [K5] | 4,49774E-05 | 1748,79234 | Liver/ FAIMS/MMTS |
| P14115 | P14115 [106-114] | [R].SGYYKVLGK.[G] | 1xHomocysteineMMTS [K5] | 0,00448015 | 1177,57441 | Liver/ FAIMS/MMTS |
| P14115 | P14115 [93-105] | [K].NKTGVAPIIDVVR.[S] | 1xHomocysteineMMTS [K2] | 0,00202818 | 1544,82873 | Liver/ FAIMS/MMTS |
| P14131 | P14131 [51-62] | [K].LLEPVLLLGKER.[F] | 1xHomocysteineIAA [K10] | 6,30912E-05 | 1553,90836 | Brain/ FAIMS/IAA |
| P14131 | P14131 [51-62] | [K].LLEPVLLLGKER.[F] | 1xHomocysteineMMTS [K10] | 0,000231655 | 1542,87462 | Liver/ No FAIMS/MMTS |
| P14131 | P14131 [51-62] | [K].LLEPVLLLGKER.[F] | 1xHomocysteineMMTS [K10] | 0,000231655 | 1542,87462 | Liver/ FAIMS/MMTS |
| P14148 | P14148 [223-234] | [R].FKEANNFLWPFK.[L] | 1xHomocysteineIAA [K2] | 4,32935E-05 | 1714,84101 | Liver/ FAIMS/IAA |
| P14148 | P14148 [223-234] | [R].FKEANNFLWPFK.[L] | 1xHomocysteineIAA [K2] | 4,32935E-05 | 1714,84101 | Brain/ FAIMS/IAA |
| P14148 | P14148 [171-179] | [K].SVNELIYKR.[G] | 1xHomocysteineMMTS [K8] | 0,0069474 | 1284,64389 | Liver/ FAIMS/MMTS |
| P14148 | P14148 [100-116] | [K].AGNFYVPAEPKLAFVIR.[I] | 1xHomocysteineMMTS [K11] | 4,49774E-05 | 2055,05543 | Liver/ FAIMS/MMTS |
| P14148 | P14148 [223-234] | [R].FKEANNFLWPFK.[L] | 1xHomocysteineMMTS [K2] | 0,0076359 | 1703,80727 | Liver/ FAIMS/MMTS |
| P14148 | P14148 [245-264] | [K].KTTHFVEGGDAGNREDQINR.[L] | 1xHomocysteineMMTS [K1] | 4,49774E-05 | 2407,08775 | Liver/ FAIMS/MMTS |
| | | | 1xAcetyl [N-Term]; | | | |
| P14148 | P14148 [1-8] | [-].MEAVPEKK.[K] | 1xHomocysteineMMTS [K7] | 0,000863044 | 1136,51485 | Liver/ FAIMS/MMTS |
| P14152 | P14152 [165-179] | [K].SQIALKLGVTADDVK.[N] | 1xHomocysteineIAA [K6] | 4,32935E-05 | 1731,93095 | Liver/ FAIMS/IAA |
| P14152 | P14152 [239-255] | [R].KLSSAMSAAKAIADHIR.[D] | 1xHomocysteineIAA [K10] | 4,32935E-05 | 1944,01536 | Brain/ FAIMS/IAA |
| P14152 | P14152 [296-310] | [K].TWKFVEGLPINDFSR.[E] | 1xHomocysteineMMTS [K3] | 4,49774E-05 | 1971,94555 | Liver/ No FAIMS/MMTS |
| P14152 | P14152 [311-318] | [R].EKMDLTAK.[E] | 1xHomocysteineMMTS [K2] | 0,00138774 | 1098,4992 | Liver/ No FAIMS/MMTS |
| P14152 | P14152 [165-179] | [K].SQIALKLGVTADDVK.[N] | 1xHomocysteineMMTS [K6] | 4,49774E-05 | 1720,8972 | Liver/ FAIMS/MMTS |
| P14152 | P14152 [296-310] | [K].TWKFVEGLPINDFSR.[E] | 1xHomocysteineMMTS [K3] | 4,49774E-05 | 1971,94555 | Liver/ FAIMS/MMTS |
| P14152 | P14152 [311-318] | [R].EKMDLTAK.[E] | 1xHomocysteineMMTS [K2] | 0,00138774 | 1098,4992 | Liver/ FAIMS/MMTS |
| P14152 | P14152 [239-248] | [R].KLSSAMSAAK.[A] | 1xHomocysteineMMTS [K1] | 0,00833937 | 1156,5523 | Liver/ FAIMS/MMTS |
| P14152 | P14152 [296-310] | [K].TWKFVEGLPINDFSR.[E] | 1xHomocysteineMMTS [K3] | 4,49774E-05 | 1971,94555 | Brain/ No FAIMS/MMTS |
| P14152 | P14152 [165-179] | [K].SQIALKLGVTADDVK.[N] | 1xHomocysteineMMTS [K6] | 4,49774E-05 | 1720,8972 | Brain/ FAIMS/MMTS |
| P14152 | P14152 [296-310] | [K].TWKFVEGLPINDFSR.[E] | 1xHomocysteineMMTS [K3] | 4,49774E-05 | 1971,94555 | Brain/ FAIMS/MMTS |
| P14152 | P14152 [108-118] | [K].IFKSQGTALEK.[Y] | 1xHomocysteineMMTS [K3] | 0,000437433 | 1384,69632 | Brain/ FAIMS/MMTS |

| | | | | | | |
|---|---|---|---|---|---|---|
| P14152 | P14152 [311-318] | [R].EKMDLTAK.[E] | 1xHomocysteineMMTS [K2] | 0,00138774 | 1098,4992 | Brain/ FAIMS/MMTS |
| P14206 | P14206 [86-102] | [R].AVLKFAAATGATPIAGR.[F] | 1xHomocysteineIAA [K4] | 4,32935E-05 | 1788,9789 | Liver/ FAIMS/IAA |
| P14206 | P14206 [86-102] | [R].AVLKFAAATGATPIAGR.[F] | 1xHomocysteineIAA [K4] | 4,32935E-05 | 1788,9789 | Brain/ FAIMS/IAA |
| P14206 | P14206 [43-53] | [K].SDGIYIINLKR.[T] | 1xHomocysteineMMTS [K10] | 0,000635569 | 1454,74942 | Liver/ No FAIMS/MMTS |
| P14206 | P14206 [42-52] | [R].KSDGIYIINLK.[R] | 1xHomocysteineMMTS [K1] | 0,00505108 | 1426,74327 | Liver/ No FAIMS/MMTS |
| P14206 | P14206 [86-102] | [R].AVLKFAAATGATPIAGR.[F] | 1xHomocysteineMMTS [K4] | 4,49774E-05 | 1777,94516 | Liver/ FAIMS/MMTS |
| P14206 | P14206 [43-53] | [K].SDGIYIINLKR.[T] | 1xHomocysteineMMTS [K10] | 0,000635569 | 1454,74942 | Liver/ FAIMS/MMTS |
| P14206 | P14206 [42-52] | [R].KSDGIYIINLK.[R] | 1xHomocysteineMMTS [K1] | 0,00505108 | 1426,74327 | Liver/ FAIMS/MMTS |
| P14211 | P14211 [207-222] | [K].KIKDPDAAKPEDWDER.[A] | 1xHomocysteineIAA [K] | 4,32935E-05 | 2086,98623 | Liver/ No FAIMS/IAA |
| P14211 | P14211 [207-222] | [K].KIKDPDAAKPEDWDER.[A] | 1xHomocysteineIAA [K] | 4,32935E-05 | 2086,98623 | Liver/ FAIMS/IAA |
| P14211 | P14211 [207-222] | [K].KIKDPDAAKPEDWDER.[A] | 1xHomocysteineIAA [K] | 4,32935E-05 | 2086,98623 | Brain/ FAIMS/IAA |
| P14211 | P14211 [74-87] | [R].FYALSAKFEPFSNK.[G] | 1xHomocysteineMMTS [K7] | 4,49774E-05 | 1811,84952 | Liver/ No FAIMS/MMTS |
| P14211 | P14211 [49-62] | [K].FVLSSGKFYGDLEK.[D] | 1xHomocysteineMMTS [K7] | 4,49774E-05 | 1752,83354 | Liver/ No FAIMS/MMTS |
| P14211 | P14211 [207-222] | [K].KIKDPDAAKPEDWDER.[A] | 1xHomocysteineMMTS [K3] | 0,000173718 | 2075,95249 | Liver/ No FAIMS/MMTS |
| P14211 | P14211 [63-73] | [K].DKGLQTSQDAR.[F] | 1xHomocysteineMMTS [K2] | 0,000724331 | 1381,61986 | Liver/ No FAIMS/MMTS |
| P14211 | P14211 [225-248] | [K].IDDPTDSKPEDWDKPEHIPDPDAK.[K] | 1xHomocysteineMMTS [K14] | 4,49774E-05 | 2923,27606 | Liver/ No FAIMS/MMTS |
| P14211 | P14211 [143-151] | [K].KVHVIFNYK.[G] | 1xHomocysteineMMTS [K1] | 0,000133296 | 1310,6748 | Liver/ No FAIMS/MMTS |
| P14211 | P14211 [208-222] | [K].IKDPDAAKPEDWDER.[A] | 1xHomocysteineMMTS [K] | 4,49774E-05 | 1947,85752 | Liver/ No FAIMS/MMTS |
| P14211 | P14211 [74-87] | [R].FYALSAKFEPFSNK.[G] | 1xHomocysteineMMTS [K7] | 4,49774E-05 | 1811,84952 | Liver/ FAIMS/MMTS |
| P14211 | P14211 [49-62] | [K].FVLSSGKFYGDLEK.[D] | 1xHomocysteineMMTS [K7] | 4,49774E-05 | 1752,83354 | Liver/ FAIMS/MMTS |
| P14211 | P14211 [207-222] | [K].KIKDPDAAKPEDWDER.[A] | 1xHomocysteineMMTS [K3] | 0,000173718 | 2075,95249 | Liver/ FAIMS/MMTS |
| P14211 | P14211 [63-73] | [K].DKGLQTSQDAR.[F] | 1xHomocysteineMMTS [K2] | 0,000724331 | 1381,61986 | Liver/ FAIMS/MMTS |
| P14211 | P14211 [143-151] | [K].KVHVIFNYK.[G] | 1xHomocysteineMMTS [K1] | 0,000133296 | 1310,6748 | Liver/ FAIMS/MMTS |
| P14211 | P14211 [56-73] | [K].FYGDLEKDKGLQTSQDAR.[F] | 1xHomocysteineMMTS [K9] | 0,00842005 | 2234,02163 | Liver/ FAIMS/MMTS |
| P14211 | P14211 [49-64] | [K].FVLSSGKFYGDLEKDK.[G] | 1xHomocysteineMMTS [K7] | 0,00905461 | 1995,95545 | Liver/ FAIMS/MMTS |
| P14211 | P14211 [81-98] | [K].FEPFSNKGQTLVVQFTVK.[H] | 1xHomocysteineMMTS [K7] | 0,000192903 | 2232,11916 | Liver/ FAIMS/MMTS |
| P14211 | P14211 [56-64] | [K].FYGDLEKDK.[G] | 1xHomocysteineMMTS [K7] | 0,00182026 | 1277,55407 | Liver/ FAIMS/MMTS |
| P14211 | P14211 [154-162] | [K].NVLINKDIR.[C] | 1xHomocysteineMMTS [K6] | 0,00199616 | 1247,65987 | Liver/ FAIMS/MMTS |
| P14211 | P14211 [44-55] | [K].SDFGKFVLSSGK.[F] | 1xHomocysteineMMTS [K5] | 0,000980684 | 1434,67558 | Liver/ FAIMS/MMTS |
| P14211 | P14211 [186-207] | [K].IDNSQVESGSLEDDWDFLPPKK.[I] | 1xHomocysteineMMTS [K21] | 4,49774E-05 | 2682,20619 | Liver/ FAIMS/MMTS |
| P14211 | P14211 [223-248] | [R].AKIDDPTDSKPEDWDKPEHIPDPDAK.[K] | 1xHomocysteineMMTS [K2] | 0,000192903 | 3122,40814 | Liver/ FAIMS/MMTS |
| P14211 | P14211 [225-248] | [K].IDDPTDSKPEDWDKPEHIPDPDAK.[K] | 1xHomocysteineMMTS [K14] | 4,49774E-05 | 2923,27606 | Liver/ FAIMS/MMTS |
| P14211 | P14211 [208-222] | [K].IKDPDAAKPEDWDER.[A] | 1xHomocysteineMMTS [K] | 4,49774E-05 | 1947,85752 | Liver/ FAIMS/MMTS |
| P14211 | P14211 [207-222] | [K].KIKDPDAAKPEDWDER.[A] | 1xHomocysteineMMTS [K3] | 0,000173718 | 2075,95249 | Brain/ FAIMS/MMTS |
| P14211 | P14211 [63-73] | [K].DKGLQTSQDAR.[F] | 1xHomocysteineMMTS [K2] | 0,000724331 | 1381,61986 | Brain/ FAIMS/MMTS |
| P14231 | P14231 [9-25] | [K].SCGQVVEEWKEFVWNPR.[T] | 1xCarbamidomethyl [C2]; 1xHomocysteineIAA [K10] | 4,32935E-05 | 2324,05868 | Brain/ FAIMS/IAA |
| P14231 | P14231 [263-280] | [R].INAANIATDDERDKFAGR.[V] | 1xHomocysteineMMTS [K14] | 4,49774E-05 | 2139,991 | Brain/ FAIMS/MMTS |
| P14602 | P14602 [193-209] | [R].AQIGGPEAGKSEQSGAK.[-] | 1xHomocysteineMMTS [K10] | 0,00980048 | 1777,82074 | Liver/ FAIMS/MMTS |
| P14824 | P14824 [241-250] | [K].LMLAVVKCIR.[S] | 1xCarbamidomethyl [C8]; 1xHomocysteineIAA [K7] | 9,61721E-05 | 1376,75748 | Brain/ FAIMS/IAA |
| P14824 | P14824 [447-465] | [K].AMEGAGTDEKTLIEILATR.[T] | 1xHomocysteineMMTS [K10] | 0,00775032 | 2182,05524 | Liver/ FAIMS/MMTS |
| P14869 | P14869 [84-99] | [R].GNVGFVFTKEDLTEIR.[D] | 1xHomocysteineIAA [K9] | 4,32935E-05 | 1998,99534 | Liver/ FAIMS/IAA |
| P14869 | P14869 [84-99] | [R].GNVGFVFTKEDLTEIR.[D] | 1xHomocysteineIAA [K9] | 4,32935E-05 | 1998,99534 | Brain/ FAIMS/IAA |
| P14869 | P14869 [84-99] | [R].GNVGFVFTKEDLTEIR.[D] | 1xHomocysteineMMTS [K9] | 4,49774E-05 | 1987,96159 | Liver/ FAIMS/MMTS |
| P14869 | P14869 [248-266] | [R].VLALSVETEYTFPLTEKVK.[A] | 1xHomocysteineMMTS [K17] | 0,00439215 | 2330,20222 | Liver/ FAIMS/MMTS |
| P14873 | P14873 [1491-1510] | [R].KLGGDVSPTQIDVSQFGSFK.[E] | 1xHomocysteineIAA [K1] | 4,32935E-05 | 2284,12781 | Brain/ FAIMS/IAA |
| P15105 | P15105 [1-14] | [-].MATSASSHLNKGIK.[Q] | 1xMet-loss+Acetyl [N-Term]; 1xHomocysteineIAA [K11] | 4,32935E-05 | 1529,77405 | Liver/ FAIMS/IAA |
| P15105 | P15105 [91-103] | [R].KDPNKLVLCEVFK.[Y] | 1xCarbamidomethyl [C9]; 1xHomocysteineIAA [K5] | 4,32935E-05 | 1763,91827 | Brain/ No FAIMS/IAA |
| P15105 | P15105 [91-103] | [R].KDPNKLVLCEVFK.[Y] | 1xCarbamidomethyl [C9]; 1xHomocysteineIAA [K5] | 4,32935E-05 | 1763,91827 | Brain/ FAIMS/IAA |
| P15105 | P15105 [1-14] | [-].MATSASSHLNKGIK.[Q] | 1xMet-loss+Acetyl [N-Term]; 1xHomocysteineMMTS [K11] | 4,49774E-05 | 1518,74031 | Liver/ No FAIMS/MMTS |
| P15105 | P15105 [287-298] | [R].AYDPKGGLDNAR.[R] | 1xHomocysteineMMTS [K5] | 0,000266738 | 1439,64059 | Liver/ No FAIMS/MMTS |
| P15105 | P15105 [107-114] | [R].KPAETNLR.[H] | 1xHomocysteineMMTS [K1] | 0,0004709 | 1091,53361 | Liver/ No FAIMS/MMTS |
| P15105 | P15105 [1-14] | [-].MATSASSHLNKGIK.[Q] | 1xMet-loss+Acetyl [N-Term]; 1xHomocysteineMMTS [K11] | 4,49774E-05 | 1518,74031 | Liver/ FAIMS/MMTS |
| P15105 | P15105 [287-298] | [R].AYDPKGGLDNAR.[R] | 1xHomocysteineMMTS [K5] | 0,000266738 | 1439,64059 | Liver/ FAIMS/MMTS |
| P15105 | P15105 [12-25] | [K].GIKQMYMSLPQGEK.[V] | 1xHomocysteineMMTS [K3] | 4,49774E-05 | 1772,82022 | Liver/ FAIMS/MMTS |
| P15105 | P15105 [107-114] | [R].KPAETNLR.[H] | 1xHomocysteineMMTS [K1] | 0,0004709 | 1091,53361 | Liver/ FAIMS/MMTS |
| P15105 | P15105 [1-14] | [-].MATSASSHLNKGIK.[Q] | 1xMet-loss+Acetyl [N-Term]; 1xHomocysteineMMTS [K11] | 4,49774E-05 | 1518,74031 | Brain/ No FAIMS/MMTS |
| P15105 | P15105 [287-298] | [R].AYDPKGGLDNAR.[R] | 1xHomocysteineMMTS [K5] | 0,000266738 | 1439,64059 | Brain/ No FAIMS/MMTS |
| P15105 | P15105 [1-14] | [-].MATSASSHLNKGIK.[Q] | 1xMet-loss+Acetyl [N-Term]; 1xHomocysteineMMTS [K11] | 4,49774E-05 | 1518,74031 | Brain/ FAIMS/MMTS |
| P15105 | P15105 [287-298] | [R].AYDPKGGLDNAR.[R] | 1xHomocysteineMMTS [K5] | 0,000266738 | 1439,64059 | Brain/ FAIMS/MMTS |
| P15105 | P15105 [12-25] | [K].GIKQMYMSLPQGEK.[V] | 1xHomocysteineMMTS [K3] | 4,49774E-05 | 1772,82022 | Brain/ FAIMS/MMTS |
| P15116 | P15116 [534-545] | [R].YTKLSDPANWLK.[I] | 1xHomocysteineIAA [K3] | 9,61721E-05 | 1609,80429 | Brain/ FAIMS/IAA |
| P15209 | P15209 [105-113] | [K].FVAYKAFLK.[N] | 1xHomocysteineIAA [K5] | 0,000177335 | 1260,68093 | Brain/ No FAIMS/IAA |
| P15209 | P15209 [105-113] | [K].FVAYKAFLK.[N] | 1xHomocysteineIAA [K5] | 0,000177335 | 1260,68093 | Brain/ FAIMS/IAA |
| P15532 | P15532 [125-143] | [K].SAEKEISLWFQPEELVEYK.[S] | 1xHomocysteineIAA [K4] | 4,32935E-05 | 2499,2112 | Brain/ No FAIMS/IAA |
| P15532 | P15532 [125-143] | [K].SAEKEISLWFQPEELVEYK.[S] | 1xHomocysteineIAA [K4] | 4,32935E-05 | 2499,2112 | Brain/ FAIMS/IAA |
| P16015 | P16015 [227-253] | [R].SLFSSAENEPPVPLVGNWRPPQPVKGR.[V] | 1xHomocysteineIAA [K25] | 4,32935E-05 | 3132,60475 | Liver/ No FAIMS/IAA |
| P16015 | P16015 [227-253] | [R].SLFSSAENEPPVPLVGNWRPPQPVKGR.[V] | 1xHomocysteineIAA [K25] | 4,32935E-05 | 3132,60475 | Liver/ FAIMS/IAA |
| P16015 | P16015 [1-24] | [-].MAKEWGYASHNGPDHWHELYPIAK.[G] | 1xMet-loss+Acetyl [N-Term]; 1xHomocysteineMMTS [K3] | 4,49774E-05 | 2911,30791 | Liver/ No FAIMS/MMTS |
| P16015 | P16015 [114-135] | [K].YAAELHLVHWNPKYNTFGEALK.[Q] | 1xHomocysteineMMTS [K13] | 4,49774E-05 | 2764,33742 | Liver/ No FAIMS/MMTS |

| | | | | | | |
|---|---|---|---|---|---|---|
| P16015 | P16015 [37-57] | [K].DIKHDPSLQPWSASYDPGSAK.[T] | 1xHomocysteineMMTS [K3] | 4,49774E-05 | 2462,11151 | Liver/ No FAIMS/MMTS |
| P16015 | P16015 [227-253] | [R].SLFSSAENEPPVPLVGNWRPPQPVKGR.[V] | 1xHomocysteineMMTS [K25] | 4,49774E-05 | 3121,571 | Liver/ No FAIMS/MMTS |
| P16015 | P16015 [152-165] | [R].EKGEFQILLDALDK.[I] | 1xHomocysteineMMTS [K2] | 6,70238E-05 | 1781,88122 | Liver/ No FAIMS/MMTS |
| P16015 | P16015 [25-39] | [K].GDNQSPIELHTKDIK.[H] | 1xHomocysteineMMTS [K12] | 4,49774E-05 | 1857,88334 | Liver/ No FAIMS/MMTS |
| P16015 | P16015 [213-226] | [K].EPMTVSSDQMAKLR.[S] | 2xOxidation [M3; M10]; 1xHomocysteineMMTS [K12] | 4,49774E-05 | 1787,77947 | Liver/ FAIMS/MMTS |
| P16015 | P16015 [213-226] | [K].EPMTVSSDQMAKLR.[S] | 1xOxidation [M]; 1xHomocysteineMMTS [K12] | 4,49774E-05 | 1771,78456 | Liver/ FAIMS/MMTS |
| P16015 | P16015 [1-24] | [-].MAKEWGYASHNGPDHWHELYPIAK.[G] | 1xMet-loss+Acetyl [N-Term]; 1xHomocysteineMMTS [K3] | 4,49774E-05 | 2911,30791 | Liver/ FAIMS/MMTS |
| P16015 | P16015 [37-57] | [K].DIKHDPSLQPWSASYDPGSAK.[T] | 1xHomocysteineMMTS [K3] | 4,49774E-05 | 2462,11151 | Liver/ FAIMS/MMTS |
| P16015 | P16015 [227-253] | [R].SLFSSAENEPPVPLVGNWRPPQPVKGR.[V] | 1xHomocysteineMMTS [K25] | 4,49774E-05 | 3121,571 | Liver/ FAIMS/MMTS |
| P16015 | P16015 [152-165] | [R].EKGEFQILLDALDK.[I] | 1xHomocysteineMMTS [K2] | 6,70238E-05 | 1781,88122 | Liver/ FAIMS/MMTS |
| P16015 | P16015 [114-135] | [K].YAAELHLVHWNPKYNTFGEALK.[Q] | 1xHomocysteineMMTS [K13] | 4,49774E-05 | 2764,33742 | Liver/ FAIMS/MMTS |
| P16015 | P16015 [25-39] | [K].GDNQSPIELHTKDIK.[H] | 1xHomocysteineMMTS [K12] | 4,49774E-05 | 1857,88334 | Liver/ FAIMS/MMTS |
| P16015 | P16015 [4-36] | [K].EWGYASHNGPDHWHELYPIAKGDNQSPIELHTK.[D] | 1xHomocysteineMMTS [K21] | 0,00820834 | 3989,81225 | Liver/ FAIMS/MMTS |
| P16125 | P16125 [319-329] | [R].KSADTLWDIQK.[D] | 1xHomocysteineIAA [K1] | 4,32935E-05 | 1478,73079 | Brain/ FAIMS/IAA |
| P16125 | P16125 [309-318] | [K].LKDDEVAQLR.[K] | 1xHomocysteineMMTS [K2] | 0,00869467 | 1349,65518 | Brain/ No FAIMS/MMTS |
| P16125 | P16125 [78-91] | [K].IVADKDYSVTANSK.[I] | 1xHomocysteineMMTS [K5] | 4,49774E-05 | 1673,78732 | Brain/ FAIMS/MMTS |
| P16125 | P16125 [309-318] | [K].LKDDEVAQLR.[K] | 1xHomocysteineMMTS [K2] | 0,00869467 | 1349,65518 | Brain/ FAIMS/MMTS |
| P16125 | P16125 [234-246] | [K].MVVDSAYEVIKLK.[G] | 1xHomocysteineMMTS [K11] | 0,000211896 | 1657,83618 | Brain/ FAIMS/MMTS |
| P16330 | P16330 [244-260] | [K].RPPGVLHCTTKFCDYGK.[A] | 2xCarbamidomethyl [C8; C13]; 1xHomocysteineIAA [K11] | 4,32935E-05 | 2210,03036 | Brain/ FAIMS/IAA |
| P16330 | P16330 [224-234] | [R].HFISGDEPKEK.[L] | 1xHomocysteineMMTS [K9] | 0,00794311 | 1449,6501 | Brain/ No FAIMS/MMTS |
| P16330 | P16330 [80-93] | [K].MVSADAYKIIPGSR.[A] | 1xHomocysteineMMTS [K8] | 4,49774E-05 | 1670,80628 | Brain/ No FAIMS/MMTS |
| P16330 | P16330 [57-68] | [R].GLPGSGKSTLAR.[L] | 1xHomocysteineMMTS [K7] | 0,0014351 | 1306,6606 | Brain/ No FAIMS/MMTS |
| P16330 | P16330 [69-79] | [R].LILEKYHDGTK.[M] | 1xHomocysteineMMTS [K5] | 0,00074635 | 1479,73343 | Brain/ No FAIMS/MMTS |
| P16330 | P16330 [233-243] | [K].EKLELVSYFGK.[R] | 1xHomocysteineMMTS [K2] | 0,000192903 | 1475,72728 | Brain/ No FAIMS/MMTS |
| P16330 | P16330 [261-275] | [K].AAGAEEYAQQEVVKR.[S] | 1xHomocysteineMMTS [K14] | 4,49774E-05 | 1811,84148 | Brain/ No FAIMS/MMTS |
| P16330 | P16330 [204-219] | [K].AGQVFLEELGNHKAFK.[K] | 1xHomocysteineMMTS [K13] | 4,49774E-05 | 1950,95645 | Brain/ No FAIMS/MMTS |
| P16330 | P16330 [224-234] | [R].HFISGDEPKEK.[L] | 1xHomocysteineMMTS [K9] | 0,00794311 | 1449,6501 | Brain/ FAIMS/MMTS |
| P16330 | P16330 [80-93] | [K].MVSADAYKIIPGSR.[A] | 1xHomocysteineMMTS [K8] | 4,49774E-05 | 1670,80628 | Brain/ FAIMS/MMTS |
| P16330 | P16330 [94-102] | [R].ADFSEAYKR.[L] | 1xHomocysteineMMTS [K8] | 4,49774E-05 | 1249,534 | Brain/ FAIMS/MMTS |
| P16330 | P16330 [74-87] | [K].YHDGTKMVSADAYK.[I] | 1xHomocysteineMMTS [K6] | 6,70238E-05 | 1748,74407 | Brain/ FAIMS/MMTS |
| P16330 | P16330 [69-79] | [R].LILEKYHDGTK.[M] | 1xHomocysteineMMTS [K5] | 0,00074635 | 1479,73343 | Brain/ FAIMS/MMTS |
| P16330 | P16330 [233-243] | [K].EKLELVSYFGK.[R] | 1xHomocysteineMMTS [K2] | 0,000192903 | 1475,72728 | Brain/ FAIMS/MMTS |
| P16330 | P16330 [162-174] | [K].EKNQWQLSADDLK.[K] | 1xHomocysteineMMTS [K2] | 0,000530332 | 1737,79347 | Brain/ FAIMS/MMTS |
| P16330 | P16330 [203-219] | [R].KAGQVFLEELGNHKAFK.[K] | 1xHomocysteineMMTS [K14] | 0,000282457 | 2079,05141 | Brain/ FAIMS/MMTS |
| P16330 | P16330 [261-275] | [K].AAGAEEYAQQEVVKR.[S] | 1xHomocysteineMMTS [K14] | 4,49774E-05 | 1811,84148 | Brain/ FAIMS/MMTS |
| P16330 | P16330 [204-219] | [K].AGQVFLEELGNHKAFK.[K] | 1xHomocysteineMMTS [K13] | 4,49774E-05 | 1950,95645 | Brain/ FAIMS/MMTS |
| P16330 | P16330 [224-243] | [R].HFISGDEPKEKLELVSYFGK.[R] | 1xHomocysteineMMTS [K11] | 0,000152766 | 2486,20943 | Brain/ FAIMS/MMTS |
| P16331 | P16331 [131-155] | [R].FANQILSYGAELDADHPGFKDPVYR.[A] | 1xHomocysteineIAA [K20] | 4,32935E-05 | 2997,41997 | Liver/ FAIMS/IAA |
| P16331 | P16331 [131-155] | [R].FANQILSYGAELDADHPGFKDPVYR.[A] | 1xHomocysteineMMTS [K20] | 4,49774E-05 | 2986,38622 | Liver/ FAIMS/MMTS |
| P16331 | P16331 [43-53] | [K].EEVGALAKVLR.[L] | 1xHomocysteineMMTS [K8] | 8,93346E-05 | 1347,7123 | Liver/ FAIMS/MMTS |
| P16388; P16390; B2RQA1; P63141 | P16388 [308-320]; P16390 [333-345]; B2RQA1 [357-369]; P63141 [310-322] | [R].HSKGLQILGQTLK.[A] | 1xHomocysteineIAA [K3] | 9,61721E-05 | 1596,88902 | Brain/ FAIMS/IAA |
| P16460 | P16460 [210-228] | [K].TQDPAKAPNSPDVLEIEFK.[K] | 1xHomocysteineIAA [K6] | 4,32935E-05 | 2273,11182 | Liver/ FAIMS/IAA |
| P16460 | P16460 [113-127] | [K].YVSHGATGKGNDQVR.[F] | 1xHomocysteineMMTS [K9] | 0,00109439 | 1751,7952 | Liver/ No FAIMS/MMTS |
| P16460 | P16460 [109-121] | [R].EGAKYVSHGATGK.[G] | 1xHomocysteineMMTS [K4] | 4,49774E-05 | 1467,67189 | Liver/ No FAIMS/MMTS |
| P16460 | P16460 [210-228] | [K].TQDPAKAPNSPDVLEIEFK.[K] | 1xHomocysteineMMTS [K6] | 4,49774E-05 | 2262,07808 | Liver/ FAIMS/MMTS |
| P16460 | P16460 [113-127] | [K].YVSHGATGKGNDQVR.[F] | 1xHomocysteineMMTS [K9] | 0,00109439 | 1751,7952 | Liver/ FAIMS/MMTS |
| P16460 | P16460 [158-176] | [R].NDLMEYAKQHGIPIPVTPK.[S] | 1xHomocysteineMMTS [K8] | 0,00460899 | 2314,13924 | Liver/ FAIMS/MMTS |
| P16460 | P16460 [216-229] | [K].APNSPDVLEIEFKK.[G] | 1xHomocysteineMMTS [K13] | 0,000530332 | 1749,855 | Liver/ FAIMS/MMTS |
| P16460 | P16460 [200-215] | [K].NQAPPGLYTKTQDPAK.[A] | 1xHomocysteineMMTS [K10] | 4,49774E-05 | 1891,90408 | Liver/ FAIMS/MMTS |
| P16546 | P16546 [1911-1923] | [K].KHEAFETDFTVHK.[D] | 1xHomocysteineIAA [K1] | 4,32935E-05 | 1762,82173 | Brain/ FAIMS/IAA |
| P16546 | P16546 [2048-2057] | [K].HIQSKAIEAR.[H] | 1xHomocysteineMMTS [K5] | 0,0046895 | 1315,66094 | Liver/ FAIMS/MMTS |
| P16546 | P16546 [2048-2057] | [K].HIQSKAIEAR.[H] | 1xHomocysteineMMTS [K5] | 0,0046895 | 1315,66094 | Brain/ FAIMS/MMTS |
| P16546 | P16546 [273-285] | [K].EKEQLMASDDFGR.[D] | 1xHomocysteineMMTS [K2] | 0,00413569 | 1688,70769 | Brain/ FAIMS/MMTS |
| P16627 | P16627 [328-344] | [K].MDKAKIHDIVLVGGSTR.[I] | 1xHomocysteineIAA [K5] | 4,32935E-05 | 2014,05723 | Brain/ FAIMS/IAA |
| P16858 | P16858 [259-269] | [K].VVKQASEGPLK.[G] | 1xHomocysteineIAA [K3] | 4,32935E-05 | 1329,7195 | Liver/ FAIMS/IAA |
| P16858 | P16858 [65-78] | [K].LVINGKPITIFQER.[D] | 1xHomocysteineIAA [K6] | 4,32935E-05 | 1801,9993 | Brain/ No FAIMS/IAA |
| P16858 | P16858 [65-78] | [K].LVINGKPITIFQER.[D] | 1xHomocysteineIAA [K6] | 4,32935E-05 | 1801,9993 | Brain/ FAIMS/IAA |
| P16858 | P16858 [259-269] | [K].VVKQASEGPLK.[G] | 1xHomocysteineMMTS [K3] | 4,49774E-05 | 1318,68575 | Liver/ No FAIMS/MMTS |
| P16858 | P16858 [259-269] | [K].VVKQASEGPLK.[G] | 1xHomocysteineMMTS [K3] | 4,49774E-05 | 1318,68575 | Liver/ FAIMS/MMTS |
| P16858 | P16858 [218-232] | [K].VIPELNGKLTGMAFR.[V] | 1xHomocysteineMMTS [K8] | 4,49774E-05 | 1808,92198 | Liver/ FAIMS/MMTS |
| P16858 | P16858 [185-195] | [K].TVDGPSGKLWR.[D] | 1xHomocysteineMMTS [K8] | 4,49774E-05 | 1378,6606 | Liver/ FAIMS/MMTS |
| P16858 | P16858 [247-258] | [R].LEKPAKYDDIKK.[V] | 1xHomocysteineMMTS [K6] | 0,000530332 | 1610,82806 | Liver/ FAIMS/MMTS |
| P16858 | P16858 [85-111] | [K].WGEAGAEYVVESTGVFTTMEKAGAHLK.[G] | 1xHomocysteineMMTS [K21] | 4,49774E-05 | 3031,39982 | Liver/ FAIMS/MMTS |
| P16858 | P16858 [322-333] | [R].VVDLMAYMASKE.[-] | 1xHomocysteineMMTS [K11] | 4,49774E-05 | 1519,66634 | Liver/ FAIMS/MMTS |
| P16858 | P16858 [185-195] | [K].TVDGPSGKLWR.[D] | 1xHomocysteineMMTS [K8] | 4,49774E-05 | 1378,6606 | Brain/ No FAIMS/MMTS |
| P16858 | P16858 [259-269] | [K].VVKQASEGPLK.[G] | 1xHomocysteineMMTS [K3] | 4,49774E-05 | 1318,68575 | Brain/ No FAIMS/MMTS |
| P16858 | P16858 [85-111] | [K].WGEAGAEYVVESTGVFTTMEKAGAHLK.[G] | 1xHomocysteineMMTS [K21] | 4,49774E-05 | 3031,39982 | Brain/ No FAIMS/MMTS |
| P16858 | P16858 [199-217] | [R].GAAQNIIPASTGAAKAVGK.[V] | 1xHomocysteineMMTS [K15] | 4,49774E-05 | 1887,97791 | Brain/ No FAIMS/MMTS |

| | | | | | | |
|---|---|---|---|---|---|---|
| P16858 | P16858 [322-333] | [R].VVDLMAYMASKE.[-] | 1xHomocysteineMMTS [K11] | 4,49774E-05 | 1519,66634 | Brain/ No FAIMS/MMTS |
| P16858 | P16858 [218-232] | [K].VIPELNGKLTGMAFR.[V] | 1xHomocysteineMMTS [K8] | 4,49774E-05 | 1808,92198 | Brain/ FAIMS/MMTS |
| P16858 | P16858 [185-195] | [K].TVDGPSGKLWR.[D] | 1xHomocysteineMMTS [K8] | 4,49774E-05 | 1378,6606 | Brain/ FAIMS/MMTS |
| P16858 | P16858 [247-258] | [R].LEKPAKYDDIKK.[V] | 1xHomocysteineMMTS [K6] | 0,000530332 | 1610,82806 | Brain/ FAIMS/MMTS |
| P16858 | P16858 [65-78] | [K].LVINGKPITIFQER.[D] | 1xHomocysteineMMTS [K6] | 0,000282457 | 1790,96556 | Brain/ FAIMS/MMTS |
| P16858 | P16858 [258-269] | [K].KVVKQASEGPLK.[G] | 1xHomocysteineMMTS [K4] | 0,0020903 | 1446,78072 | Brain/ FAIMS/MMTS |
| P16858 | P16858 [259-269] | [K].VVKQASEGPLK.[G] | 1xHomocysteineMMTS [K3] | 4,49774E-05 | 1318,68575 | Brain/ FAIMS/MMTS |
| P16858 | P16858 [117-143] | [R].VIISAPSADAPMFVMGVNHEKYDNSLK.[I] | 1xHomocysteineMMTS [K21] | 4,49774E-05 | 3096,46613 | Brain/ FAIMS/MMTS |
| P16858 | P16858 [85-111] | [K].WGEAGAEYVVESTGVFTTMEKAGAHLK.[G] | 1xHomocysteineMMTS [K21] | 4,49774E-05 | 3031,39982 | Brain/ FAIMS/MMTS |
| P16858 | P16858 [199-217] | [R].GAAQNIIPASTGAAKAVGK.[V] | 1xHomocysteineMMTS [K15] | 4,49774E-05 | 1887,97791 | Brain/ FAIMS/MMTS |
| P16858 | P16858 [322-333] | [R].VVDLMAYMASKE.[-] | 1xHomocysteineMMTS [K11] | 4,49774E-05 | 1519,66634 | Brain/ FAIMS/MMTS |
| P17047 | P17047 [135-148] | [K].NPENFKVPLDVIFK.[C] | 1xHomocysteineIAA [K6] | 6,30912E-05 | 1833,95677 | Liver/ FAIMS/IAA |
| P17047 | P17047 [135-148] | [K].NPENFKVPLDVIFK.[C] | 1xHomocysteineIAA [K6] | 6,30912E-05 | 1833,95677 | Brain/ FAIMS/IAA |
| P17047 | P17047 [135-148] | [K].NPENFKVPLDVIFK.[C] | 1xHomocysteineMMTS [K6] | 4,49774E-05 | 1822,92302 | Liver/ FAIMS/MMTS |
| P17156 | P17156 [573-586] | [K].ILDKCQEVINWLDR.[N] | 1xCarbamidomethyl [C5]; 1xHomocysteineIAA [K4] | 0,00012645 | 1975,97283 | Brain/ FAIMS/IAA |
| P17182 | P17182 [61-80] | [K].GVSQAVEHINKTIAPALVSK.[K] | 1xHomocysteineIAA [K11] | 4,32935E-05 | 2236,21181 | Liver/ FAIMS/IAA |
| P17182 | P17182 [81-92] | [K].KVNVVEQEKIDK.[L] | 1xHomocysteineIAA [K] | 4,32935E-05 | 1602,85197 | Brain/ No FAIMS/IAA |
| P17182 | P17182 [81-92] | [K].KVNVVEQEKIDK.[L] | 1xHomocysteineIAA [K] | 4,32935E-05 | 1602,85197 | Brain/ FAIMS/IAA |
| P17182 | P17182 [72-81] | [K].TIAPALVSKK.[V] | 1xHomocysteineMMTS [K9] | 0,000530332 | 1190,66356 | Liver/ No FAIMS/MMTS |
| P17182 | P17182 [413-422] | [R].IEEELGSKAK.[F] | 1xHomocysteineMMTS [K8] | 0,000266738 | 1266,60683 | Liver/ No FAIMS/MMTS |
| P17182 | P17182 [222-233] | [K].EALELLKTAIAK.[A] | 1xHomocysteineMMTS [K7] | 4,49774E-05 | 1462,80078 | Liver/ No FAIMS/MMTS |
| P17182 | P17182 [254-262] | [R].SGKYDLDFK.[S] | 1xHomocysteineMMTS [K3] | 0,00832764 | 1235,54351 | Liver/ No FAIMS/MMTS |
| P17182 | P17182 [61-80] | [K].GVSQAVEHINKTIAPALVSK.[K] | 1xHomocysteineMMTS [K11] | 4,49774E-05 | 2225,17807 | Liver/ No FAIMS/MMTS |
| P17182 | P17182 [81-89] | [K].KVNVVEQEK.[I] | 1xHomocysteineMMTS [K1] | 0,000282457 | 1235,61225 | Liver/ No FAIMS/MMTS |
| P17182 | P17182 [413-422] | [R].IEEELGSKAK.[F] | 1xHomocysteineMMTS [K8] | 0,000266738 | 1266,60683 | Liver/ FAIMS/MMTS |
| P17182 | P17182 [222-233] | [K].EALELLKTAIAK.[A] | 1xHomocysteineMMTS [K7] | 4,49774E-05 | 1462,80078 | Liver/ FAIMS/MMTS |
| P17182 | P17182 [90-103] | [K].IDKLMIEMDGTENK.[S] | 1xHomocysteineMMTS [K3] | 0,00213323 | 1799,80462 | Liver/ FAIMS/MMTS |
| P17182 | P17182 [203-228] | [K].DATNVGDEGGFAPNILENKEALELLK.[T] | 1xHomocysteineMMTS [K19] | 4,49774E-05 | 2920,40668 | Liver/ FAIMS/MMTS |
| P17182 | P17182 [61-80] | [K].GVSQAVEHINKTIAPALVSK.[K] | 1xHomocysteineMMTS [K11] | 4,49774E-05 | 2225,17807 | Liver/ FAIMS/MMTS |
| P17182 | P17182 [184-197] | [R].IGAEVYHNLKNVIK.[E] | 1xHomocysteineMMTS [K10] | 4,49774E-05 | 1760,91861 | Liver/ FAIMS/MMTS |
| P17182 | P17182 [90-103] | [K].IDKLMIEMDGTENK.[S] | 1xOxidation [M]; 1xHomocysteineMMTS [K3] | 0,00386874 | 1815,79954 | Liver/ FAIMS/MMTS |
| P17182 | P17182 [72-81] | [K].TIAPALVSKK.[V] | 1xHomocysteineMMTS [K9] | 0,000530332 | 1190,66356 | Liver/ FAIMS/MMTS |
| P17182 | P17182 [121-132] | [K].AGAVEKGVPLYR.[H] | 1xHomocysteineMMTS [K6] | 4,49774E-05 | 1422,7232 | Liver/ FAIMS/MMTS |
| P17182 | P17182 [57-71] | [R].FMGKGVSQAVEHINK.[T] | 1xHomocysteineMMTS [K4] | 0,000863044 | 1807,86519 | Liver/ FAIMS/MMTS |
| P17182 | P17182 [404-412] | [R].LAKYNQILR.[I] | 1xHomocysteineMMTS [K3] | 0,00278756 | 1281,68061 | Liver/ FAIMS/MMTS |
| P17182 | P17182 [200-221] | [K].YGKDATNVGDEGGFAPNILENK.[E] | 1xHomocysteineMMTS [K3] | 4,49774E-05 | 2472,11698 | Liver/ FAIMS/MMTS |
| P17182 | P17182 [307-327] | [K].FTASAGIQVVGDDLTVTNPKR.[I] | 1xHomocysteineMMTS [K20] | 4,49774E-05 | 2352,16863 | Liver/ FAIMS/MMTS |
| P17182 | P17182 [93-105] | [K].LMIEMDGTENKSK.[F] | 1xHomocysteineMMTS [K11] | 0,00298406 | 1658,72565 | Liver/ FAIMS/MMTS |
| P17182 | P17182 [81-89] | [K].KVNVVEQEK.[I] | 1xHomocysteineMMTS [K1] | 0,000282457 | 1235,61225 | Liver/ FAIMS/MMTS |
| P17182 | P17182 [72-81] | [K].TIAPALVSKK.[V] | 1xHomocysteineMMTS [K9] | 0,000530332 | 1190,66356 | Brain/ FAIMS/MMTS |
| P17182 | P17182 [413-422] | [R].IEEELGSKAK.[F] | 1xHomocysteineMMTS [K8] | 0,000266738 | 1266,60683 | Brain/ FAIMS/MMTS |
| P17182 | P17182 [82-92] | [K].VNVVEQEKIDK.[L] | 1xHomocysteineMMTS [K8] | 0,000152766 | 1463,72326 | Brain/ FAIMS/MMTS |
| P17182 | P17182 [222-233] | [K].EALELLKTAIAK.[A] | 1xHomocysteineMMTS [K7] | 4,49774E-05 | 1462,80078 | Brain/ FAIMS/MMTS |
| P17182 | P17182 [404-412] | [R].LAKYNQILR.[I] | 1xHomocysteineMMTS [K3] | 0,00278756 | 1281,68061 | Brain/ FAIMS/MMTS |
| P17182 | P17182 [203-228] | [K].DATNVGDEGGFAPNILENKEALELLK.[T] | 1xHomocysteineMMTS [K19] | 4,49774E-05 | 2920,40668 | Brain/ FAIMS/MMTS |
| P17182 | P17182 [61-80] | [K].GVSQAVEHINKTIAPALVSK.[K] | 1xHomocysteineMMTS [K11] | 4,49774E-05 | 2225,17807 | Brain/ FAIMS/MMTS |
| P17182 | P17182 [184-197] | [R].IGAEVYHNLKNVIK.[E] | 1xHomocysteineMMTS [K10] | 4,49774E-05 | 1760,91861 | Brain/ FAIMS/MMTS |
| P17182; P17183 | P17182 [16-32]; P17183 [16-32] | [R].GNPTVEVDLYTAKGLFR.[A] | 1xHomocysteineIAA [K13] | 4,32935E-05 | 2054,03754 | Brain/ No FAIMS/IAA |
| P17182; P17183 | P17182 [16-32]; P17183 [16-32] | [R].GNPTVEVDLYTAKGLFR.[A] | 1xHomocysteineIAA [K13] | 4,32935E-05 | 2054,03754 | Brain/ FAIMS/IAA |
| P17182; P17183 | P17182 [16-32]; P17183 [16-32] | [R].GNPTVEVDLYTAKGLFR.[A] | 1xHomocysteineMMTS [K13] | 4,49774E-05 | 2043,00379 | Liver/ No FAIMS/MMTS |
| P17182; P17183 | P17182 [16-32]; P17183 [16-32] | [R].GNPTVEVDLYTAKGLFR.[A] | 1xHomocysteineMMTS [K13] | 4,49774E-05 | 2043,00379 | Liver/ FAIMS/MMTS |
| P17182; P17183 | P17182 [16-32]; P17183 [16-32] | [R].GNPTVEVDLYTAKGLFR.[A] | 1xHomocysteineMMTS [K13] | 4,49774E-05 | 2043,00379 | Brain/ No FAIMS/MMTS |
| P17182; P17183 | P17182 [16-32]; P17183 [16-32] | [R].GNPTVEVDLYTAKGLFR.[A] | 1xHomocysteineMMTS [K13] | 4,49774E-05 | 2043,00379 | Brain/ FAIMS/MMTS |
| P17183 | P17183 [61-72] | [K].GVLKAVDHINSR.[I] | 1xHomocysteineIAA [K4] | 4,32935E-05 | 1482,78456 | Brain/ No FAIMS/IAA |
| P17183 | P17183 [61-72] | [K].GVLKAVDHINSR.[I] | 1xHomocysteineIAA [K4] | 4,32935E-05 | 1482,78456 | Brain/ FAIMS/IAA |
| P17183 | P17183 [1-9] | [-].MSIEKIWAR.[E] | 1xMet-loss+Acetyl [N-Term]; 1xHomocysteineMMTS [K5] | 0,000282457 | 1207,59621 | Brain/ No FAIMS/MMTS |
| P17183 | P17183 [61-72] | [K].GVLKAVDHINSR.[I] | 1xHomocysteineMMTS [K4] | 4,49774E-05 | 1471,75081 | Brain/ No FAIMS/MMTS |
| P17183 | P17183 [184-197] | [R].LGAEVYHTLKGVIK.[D] | 1xHomocysteineMMTS [K10] | 4,49774E-05 | 1690,90189 | Brain/ No FAIMS/MMTS |
| P17183 | P17183 [1-9] | [-].MSIEKIWAR.[E] | 1xMet-loss+Acetyl [N-Term]; 1xHomocysteineMMTS [K5] | 0,000282457 | 1207,59621 | Brain/ FAIMS/MMTS |
| P17183 | P17183 [229-239] | [K].EAIDKAGYTEK.[M] | 1xHomocysteineMMTS [K5] | 0,00188612 | 1387,62321 | Brain/ FAIMS/MMTS |
| P17183 | P17183 [61-72] | [K].GVLKAVDHINSR.[I] | 1xHomocysteineMMTS [K4] | 4,49774E-05 | 1471,75081 | Brain/ FAIMS/MMTS |
| P17183 | P17183 [307-327] | [K].FTANVGIQIVGDDLTVTNPKR.[I] | 1xHomocysteineMMTS [K20] | 4,49774E-05 | 2421,22648 | Brain/ FAIMS/MMTS |
| P17183 | P17183 [184-197] | [R].LGAEVYHTLKGVIK.[D] | 1xHomocysteineMMTS [K10] | 4,49774E-05 | 1690,90189 | Brain/ FAIMS/MMTS |
| P17426 | P17426 [548-564] | [K].FINLFPETKATIQGVLR.[A] | 1xHomocysteineIAA [K9] | 4,32935E-05 | 2121,15251 | Brain/ FAIMS/IAA |
| P17427; P17426 | P17427 [111-123]; P17426 [111-123] | [R].LINNAIKNDLASR.[N] | 1xHomocysteineIAA [K7] | 4,32935E-05 | 1615,85845 | Brain/ No FAIMS/IAA |
| P17427; P17426 | P17427 [111-123]; P17426 [111-123] | [R].LINNAIKNDLASR.[N] | 1xHomocysteineIAA [K7] | 4,32935E-05 | 1615,85845 | Brain/ FAIMS/IAA |

| | | | | | | |
|---|---|---|---|---|---|---|
| P17563 | P17563 [399-419] | [R].LYATTSLYSAWDKQFYPDLIR.[E] | 1xHomocysteineIAA [K13] | 9,61721E-05 | 2725,33305 | Liver/ FAIMS/IAA |
| P17563 | P17563 [277-289] | [R].FYKNAEGTWSVEK.[V] | 1xHomocysteineMMTS [K3] | 0,000617167 | 1721,76619 | Liver/ FAIMS/MMTS |
| P17563 | P17563 [280-296] | [K].NAEGTWSVEKVIQVPSK.[K] | 1xHomocysteineMMTS [K10] | 4,49774E-05 | 2034,99871 | Liver/ FAIMS/MMTS |
| P17665 | P17665 [26-36] | [K].NLPFSVENKWR.[L] | 1xHomocysteineIAA [K9] | 4,32935E-05 | 1563,77366 | Liver/ FAIMS/IAA |
| P17665 | P17665 [17-34] | [R].SHYEEGPGKNLPFSVENK.[W] | 1xHomocysteineIAA [K9] | 4,32935E-05 | 2206,02334 | Brain/ No FAIMS/IAA |
| P17665 | P17665 [17-34] | [R].SHYEEGPGKNLPFSVENK.[W] | 1xHomocysteineIAA [K9] | 4,32935E-05 | 2206,02334 | Brain/ FAIMS/IAA |
| P17665 | P17665 [17-34] | [R].SHYEEGPGKNLPFSVENK.[W] | 1xHomocysteineMMTS [K9] | 4,49774E-05 | 2194,9896 | Liver/ No FAIMS/MMTS |
| P17665 | P17665 [26-36] | [K].NLPFSVENKWR.[L] | 1xHomocysteineMMTS [K9] | 0,000152766 | 1552,73991 | Liver/ No FAIMS/MMTS |
| P17665 | P17665 [17-34] | [R].SHYEEGPGKNLPFSVENK.[W] | 1xHomocysteineMMTS [K9] | 4,49774E-05 | 2194,9896 | Liver/ FAIMS/MMTS |
| P17665 | P17665 [26-36] | [K].NLPFSVENKWR.[L] | 1xHomocysteineMMTS [K9] | 0,000152766 | 1552,73991 | Liver/ FAIMS/MMTS |
| P17665 | P17665 [17-34] | [R].SHYEEGPGKNLPFSVENK.[W] | 1xHomocysteineMMTS [K9] | 4,49774E-05 | 2194,9896 | Brain/ FAIMS/MMTS |
| P17665 | P17665 [26-36] | [K].NLPFSVENKWR.[L] | 1xHomocysteineMMTS [K9] | 0,000152766 | 1552,73991 | Brain/ FAIMS/MMTS |
| P17710 | P17710 [836-850] | [R].GIFETKFLSQIESDR.[L] | 1xHomocysteineIAA [K6] | 4,32935E-05 | 1943,95314 | Brain/ FAIMS/IAA |
| P17717 | P17717 [175-187] | [R].FSPGYQIEKSSGR.[F] | 1xHomocysteineIAA [K9] | 0,00012645 | 1629,76897 | Liver/ FAIMS/IAA |
| P17742 | P17742 [77-91] | [R].SIYGEKFEDENFILK.[H] | 1xHomocysteineIAA [K6] | 4,32935E-05 | 2005,95755 | Liver/ FAIMS/IAA |
| P17742 | P17742 [77-91] | [R].SIYGEKFEDENFILK.[H] | 1xHomocysteineIAA [K6] | 4,32935E-05 | 2005,95755 | Brain/ No FAIMS/IAA |
| P17742 | P17742 [77-91] | [R].SIYGEKFEDENFILK.[H] | 1xHomocysteineIAA [K6] | 4,32935E-05 | 2005,95755 | Brain/ FAIMS/IAA |
| P17742 | P17742 [20-31] | [R].VSFELFADKVPK.[T] | 1xHomocysteineMMTS [K9] | 4,49774E-05 | 1542,76948 | Liver/ No FAIMS/MMTS |
| P17742 | P17742 [38-49] | [R].ALSTGEKGFGYK.[G] | 1xHomocysteineMMTS [K7] | 4,49774E-05 | 1420,65993 | Liver/ No FAIMS/MMTS |
| P17742 | P17742 [119-131] | [K].TEWLDGKHVVFGK.[V] | 1xHomocysteineMMTS [K7] | 0,000530332 | 1678,80799 | Liver/ No FAIMS/MMTS |
| P17742 | P17742 [77-91] | [R].SIYGEKFEDENFILK.[H] | 1xHomocysteineMMTS [K6] | 4,49774E-05 | 1994,92381 | Liver/ No FAIMS/MMTS |
| P17742 | P17742 [29-37] | [K].VPKTAENFR.[A] | 1xHomocysteineMMTS [K3] | 0,000526479 | 1224,58637 | Liver/ No FAIMS/MMTS |
| P17742 | P17742 [132-144] | [K].VKEGMNIVEAMER.[F] | 1xHomocysteineMMTS [K2] | 0,000173718 | 1668,75761 | Liver/ No FAIMS/MMTS |
| P17742 | P17742 [20-37] | [R].VSFELFADKVPKTAENFR.[A] | 1xHomocysteineMMTS [K12] | 4,49774E-05 | 2261,10932 | Liver/ No FAIMS/MMTS |
| P17742 | P17742 [132-144] | [K].VKEGMNIVEAMER.[F] | 1xOxidation [M5]; 1xHomocysteineMMTS [K2] | 4,49774E-05 | 1684,75253 | Liver/ FAIMS/MMTS |
| P17742 | P17742 [20-31] | [R].VSFELFADKVPK.[T] | 1xHomocysteineMMTS [K9] | 4,49774E-05 | 1542,76948 | Liver/ FAIMS/MMTS |
| P17742 | P17742 [38-49] | [R].ALSTGEKGFGYK.[G] | 1xHomocysteineMMTS [K7] | 4,49774E-05 | 1420,65993 | Liver/ FAIMS/MMTS |
| P17742 | P17742 [77-91] | [R].SIYGEKFEDENFILK.[H] | 1xHomocysteineMMTS [K6] | 4,49774E-05 | 1994,92381 | Liver/ FAIMS/MMTS |
| P17742 | P17742 [29-37] | [K].VPKTAENFR.[A] | 1xHomocysteineMMTS [K3] | 0,000526479 | 1224,58637 | Liver/ FAIMS/MMTS |
| P17742 | P17742 [132-144] | [K].VKEGMNIVEAMER.[F] | 1xHomocysteineMMTS [K2] | 0,000173718 | 1668,75761 | Liver/ FAIMS/MMTS |
| P17742 | P17742 [20-37] | [R].VSFELFADKVPKTAENFR.[A] | 1xHomocysteineMMTS [K12] | 4,49774E-05 | 2261,10932 | Liver/ FAIMS/MMTS |
| P17742 | P17742 [119-131] | [K].TEWLDGKHVVFGK.[V] | 1xHomocysteineMMTS [K7] | 0,000530332 | 1678,80799 | Liver/ FAIMS/MMTS |
| P17742 | P17742 [45-55] | [K].GFGYKGSSFHR.[I] | 1xHomocysteineMMTS [K5] | 0,000152766 | 1405,61399 | Liver/ FAIMS/MMTS |
| P17742 | P17742 [20-31] | [R].VSFELFADKVPK.[T] | 1xHomocysteineMMTS [K9] | 4,49774E-05 | 1542,76948 | Brain/ No FAIMS/MMTS |
| P17742 | P17742 [38-49] | [R].ALSTGEKGFGYK.[G] | 1xHomocysteineMMTS [K7] | 4,49774E-05 | 1420,65993 | Brain/ No FAIMS/MMTS |
| P17742 | P17742 [77-91] | [R].SIYGEKFEDENFILK.[H] | 1xHomocysteineMMTS [K6] | 4,49774E-05 | 1994,92381 | Brain/ No FAIMS/MMTS |
| P17742 | P17742 [132-144] | [K].VKEGMNIVEAMER.[F] | 1xHomocysteineMMTS [K2] | 0,000173718 | 1668,75761 | Brain/ No FAIMS/MMTS |
| P17742 | P17742 [20-37] | [R].VSFELFADKVPKTAENFR.[A] | 1xHomocysteineMMTS [K12] | 4,49774E-05 | 2261,10932 | Brain/ No FAIMS/MMTS |
| P17742 | P17742 [20-31] | [R].VSFELFADKVPK.[T] | 1xHomocysteineMMTS [K9] | 4,49774E-05 | 1542,76948 | Brain/ FAIMS/MMTS |
| P17742 | P17742 [38-49] | [R].ALSTGEKGFGYK.[G] | 1xHomocysteineMMTS [K7] | 4,49774E-05 | 1420,65993 | Brain/ FAIMS/MMTS |
| P17742 | P17742 [119-131] | [K].TEWLDGKHVVFGK.[V] | 1xHomocysteineMMTS [K7] | 0,000530332 | 1678,80799 | Brain/ FAIMS/MMTS |
| P17742 | P17742 [77-91] | [R].SIYGEKFEDENFILK.[H] | 1xHomocysteineMMTS [K6] | 4,49774E-05 | 1994,92381 | Brain/ FAIMS/MMTS |
| P17742 | P17742 [45-55] | [K].GFGYKGSSFHR.[I] | 1xHomocysteineMMTS [K5] | 0,000152766 | 1405,61399 | Brain/ FAIMS/MMTS |
| P17742 | P17742 [29-37] | [K].VPKTAENFR.[A] | 1xHomocysteineMMTS [K3] | 0,000526479 | 1224,58637 | Brain/ FAIMS/MMTS |
| P17742 | P17742 [132-144] | [K].VKEGMNIVEAMER.[F] | 1xHomocysteineMMTS [K2] | 0,000173718 | 1668,75761 | Brain/ FAIMS/MMTS |
| P17742 | P17742 [20-37] | [R].VSFELFADKVPKTAENFR.[A] | 1xHomocysteineMMTS [K12] | 4,49774E-05 | 2261,10932 | Brain/ FAIMS/MMTS |
| P17751 | P17751 [191-206] | [R].GWLKSNVNDGVAQSTR.[I] | 1xHomocysteineIAA [K4] | 4,32935E-05 | 1905,92357 | Liver/ FAIMS/IAA |
| P17751 | P17751 [54-69] | [R].QKLDPKIAVAAQNCYK.[V] | 1xCarbamidomethyl [C14]; 1xHomocysteineIAA [K6] | 4,32935E-05 | 2021,03068 | Brain/ FAIMS/IAA |
| P17751 | P17751 [143-156] | [K].VVFEQTKVIADNVK.[D] | 1xHomocysteineMMTS [K7] | 4,49774E-05 | 1752,90229 | Liver/ No FAIMS/MMTS |
| P17751 | P17751 [191-206] | [R].GWLKSNVNDGVAQSTR.[I] | 1xHomocysteineMMTS [K4] | 4,49774E-05 | 1894,88983 | Liver/ No FAIMS/MMTS |
| P17751 | P17751 [161-188] | [K].VVLAYEPVWAIGTGKTATPQQAQEVHEK.[L] | 1xHomocysteineMMTS [K15] | 4,49774E-05 | 3213,60712 | Liver/ No FAIMS/MMTS |
| P17751 | P17751 [176-190] | [K].TATPQQAQEVHEKLR.[G] | 1xHomocysteineMMTS [K13] | 4,49774E-05 | 1898,92113 | Liver/ No FAIMS/MMTS |
| P17751 | P17751 [143-156] | [K].VVFEQTKVIADNVK.[D] | 1xHomocysteineMMTS [K7] | 4,49774E-05 | 1752,90229 | Liver/ FAIMS/MMTS |
| P17751 | P17751 [191-206] | [R].GWLKSNVNDGVAQSTR.[I] | 1xHomocysteineMMTS [K4] | 4,49774E-05 | 1894,88983 | Liver/ FAIMS/MMTS |
| P17751 | P17751 [176-190] | [K].TATPQQAQEVHEKLR.[G] | 1xHomocysteineMMTS [K13] | 4,49774E-05 | 1898,92113 | Liver/ FAIMS/MMTS |
| P17751 | P17751 [136-149] | [R].EAGITEKVVFEQTK.[V] | 1xHomocysteineMMTS [K7] | 4,49774E-05 | 1741,84992 | Liver/ FAIMS/MMTS |
| P17751 | P17751 [161-188] | [K].VVLAYEPVWAIGTGKTATPQQAQEVHEK.[L] | 1xHomocysteineMMTS [K15] | 4,49774E-05 | 3213,60712 | Liver/ FAIMS/MMTS |
| P17751 | P17751 [143-156] | [K].VVFEQTKVIADNVK.[D] | 1xHomocysteineMMTS [K7] | 4,49774E-05 | 1752,90229 | Brain/ No FAIMS/MMTS |
| P17751 | P17751 [150-160] | [K].VIADNVKDWSK.[V] | 1xHomocysteineMMTS [K7] | 4,49774E-05 | 1437,68648 | Brain/ No FAIMS/MMTS |
| P17751 | P17751 [176-190] | [K].TATPQQAQEVHEKLR.[G] | 1xHomocysteineMMTS [K13] | 4,49774E-05 | 1898,92113 | Brain/ No FAIMS/MMTS |
| P17751 | P17751 [143-156] | [K].VVFEQTKVIADNVK.[D] | 1xHomocysteineMMTS [K7] | 4,49774E-05 | 1752,90229 | Brain/ FAIMS/MMTS |
| P17751 | P17751 [150-160] | [K].VIADNVKDWSK.[V] | 1xHomocysteineMMTS [K7] | 4,49774E-05 | 1437,68648 | Brain/ FAIMS/MMTS |
| P17751 | P17751 [136-149] | [R].EAGITEKVVFEQTK.[V] | 1xHomocysteineMMTS [K7] | 4,49774E-05 | 1741,84992 | Brain/ FAIMS/MMTS |
| P17751 | P17751 [191-206] | [R].GWLKSNVNDGVAQSTR.[I] | 1xHomocysteineMMTS [K4] | 4,49774E-05 | 1894,88983 | Brain/ FAIMS/MMTS |
| P17751 | P17751 [161-188] | [K].VVLAYEPVWAIGTGKTATPQQAQEVHEK.[L] | 1xHomocysteineMMTS [K15] | 4,49774E-05 | 3213,60712 | Brain/ FAIMS/MMTS |
| P17751 | P17751 [176-190] | [K].TATPQQAQEVHEKLR.[G] | 1xHomocysteineMMTS [K13] | 4,49774E-05 | 1898,92113 | Brain/ FAIMS/MMTS |
| P18052 | P18052 [781-795] | [K].AEGILDVFQTVKSLR.[L] | 1xHomocysteineIAA [K12] | 0,00988041 | 1849,98404 | Brain/ FAIMS/IAA |
| P18242 | P18242 [175-192] | [K].QIFGEATKQPGIVFVAAK.[F] | 1xHomocysteineIAA [K8] | 4,32935E-05 | 2078,11031 | Brain/ FAIMS/IAA |
| P18242 | P18242 [221-233] | [K].LVDKNIFSFYLNR.[D] | 1xHomocysteineMMTS [K4] | 4,49774E-05 | 1791,89206 | Liver/ No FAIMS/MMTS |
| P18242 | P18242 [221-233] | [K].LVDKNIFSFYLNR.[D] | 1xHomocysteineMMTS [K4] | 4,49774E-05 | 1791,89206 | Liver/ FAIMS/MMTS |
| P18242 | P18242 [352-361] | [K].YILKVSQGGK.[T] | 1xHomocysteineMMTS [K4] | 0,00952012 | 1255,65373 | Liver/ FAIMS/MMTS |
| P18242 | P18242 [172-182] | [K].VEKQIFGEATK.[Q] | 1xHomocysteineMMTS [K3] | 0,00760806 | 1412,69123 | Liver/ FAIMS/MMTS |
| P18572 | P18572 [177-197] | [K].VLQEDTLPDLHTKYIVDADDR.[S] | 1xHomocysteineIAA [K13] | 4,32935E-05 | 2630,27666 | Liver/ FAIMS/IAA |
| P18572 | P18572 [354-370] | [R].KPDQTLDEDDPGAAPLK.[G] | 1xHomocysteineIAA [K1] | 4,32935E-05 | 1983,9328 | Brain/ FAIMS/IAA |
| P18760 | P18760 [133-146] | [K].HELQANCYEEVKDR.[C] | 1xCarbamidomethyl [C7]; 1xHomocysteineIAA [K12] | 4,32935E-05 | 1964,85892 | Liver/ FAIMS/IAA |
| P18760 | P18760 [96-112] | [K].KEDLVFIFWAPENAPLK.[S] | 1xHomocysteineIAA [K1] | 4,32935E-05 | 2191,12562 | Brain/ No FAIMS/IAA |

| | | | | | | |
|---|---|---|---|---|---|---|
| P18760 | P18760 [96-112] | [K].KEDLVFIFWAPENAPLK.[S] | 1xHomocysteineIAA [K1] | 4,32935E-05 | 2191,12562 | Brain/ FAIMS/IAA |
| P18760 | P18760 [153-166] | [K].LGGSAVISLEGKPL.[-] | 1xHomocysteineMMTS [K12] | 4,49774E-05 | 1503,79095 | Brain/ FAIMS/MMTS |
| P18872 | P18872 [52-67] | [K].QMKIIHEDGFSGEDVK.[Q] | 1xHomocysteineIAA [K3] | 4,32935E-05 | 2006,93102 | Brain/ No FAIMS/IAA |
| P18872 | P18872 [52-67] | [K].QMKIIHEDGFSGEDVK.[Q] | 1xHomocysteineIAA [K3] | 4,32935E-05 | 2006,93102 | Brain/ FAIMS/IAA |
| P18872 | P18872 [33-46] | [K].DVKLLLLGAGESGK.[S] | 1xHomocysteineMMTS [K3] | 4,49774E-05 | 1562,82806 | Brain/ FAIMS/MMTS |
| P18872 | P18872 [52-67] | [K].QMKIIHEDGFSGEDVK.[Q] | 1xHomocysteineMMTS [K3] | 0,00394153 | 1995,89728 | Brain/ FAIMS/MMTS |
| P19096 | P19096 [1479-1500] | [K].LDPGSPELQQVLKHDLVMNVYR.[D] | 1xHomocysteineIAA [K13] | 4,32935E-05 | 2725,38002 | Brain/ FAIMS/IAA |
| P19157 | P19157 [104-121] | [K].YVTLIYTNYENGKNDYVK.[A] | 1xHomocysteineMMTS [K13] | 4,49774E-05 | 2360,09373 | Liver/ FAIMS/MMTS |
| P19157 | P19157 [102-116] | [R].GKYVTLIYTNYENGK.[N] | 1xHomocysteineMMTS [K2] | 0,000437433 | 1925,91358 | Liver/ FAIMS/MMTS |
| P19246 | P19246 [448-463] | [K].SEKETVIVEGQTEEIR.[V] | 1xHomocysteineIAA [K3] | 4,32935E-05 | 2020,98556 | Brain/ No FAIMS/IAA |
| P19246 | P19246 [448-463] | [K].SEKETVIVEGQTEEIR.[V] | 1xHomocysteineIAA [K3] | 4,32935E-05 | 2020,98556 | Brain/ FAIMS/IAA |
| P19253 | P19253 [149-160] | [K].YQAVTATLEEKR.[K] | 1xHomocysteineIAA [K11] | 8,35381E-05 | 1582,78937 | Liver/ FAIMS/IAA |
| P19253 | P19253 [149-160] | [K].YQAVTATLEEKR.[K] | 1xHomocysteineIAA [K11] | 8,35381E-05 | 1582,78937 | Brain/ FAIMS/IAA |
| P19253 | P19253 [118-128] | [R].MVVPAALKVVR.[L] | 1xHomocysteineMMTS [K8] | 4,49774E-05 | 1345,75167 | Liver/ No FAIMS/MMTS |
| P19253 | P19253 [118-128] | [R].MVVPAALKVVR.[L] | 1xHomocysteineMMTS [K8] | 4,49774E-05 | 1345,75167 | Liver/ FAIMS/MMTS |
| P19253 | P19253 [141-159] | [R].LAHEVGWKYQAVTATLEEK.[R] | 1xHomocysteineMMTS [K8] | 4,49774E-05 | 2336,14135 | Liver/ FAIMS/MMTS |
| P19253 | P19253 [149-160] | [K].YQAVTATLEEKR.[K] | 1xHomocysteineMMTS [K11] | 4,49774E-05 | 1571,75562 | Liver/ FAIMS/MMTS |
| P19536 | P19536 [68-85] | [K].AASGTKEDPNLVPSISNK.[R] | 1xHomocysteineIAA [K6] | 4,32935E-05 | 2001,99098 | Liver/ FAIMS/IAA |
| P19536 | P19536 [87-111] | [R].IVGCICEEDNCTVIWFWLHKGESQR.[C] | 3xCarbamidomethyl [C4; C6; C11]; 1xHomocysteineIAA [K20] | 4,32935E-05 | 3310,49005 | Brain/ FAIMS/IAA |
| P19536 | P19536 [68-85] | [K].AASGTKEDPNLVPSISNK.[R] | 1xHomocysteineMMTS [K6] | 4,49774E-05 | 1990,95724 | Liver/ No FAIMS/MMTS |
| P19536 | P19536 [74-86] | [K].EDPNLVPSISNKR.[I] | 1xHomocysteineMMTS [K12] | 0,00115861 | 1631,78799 | Liver/ No FAIMS/MMTS |
| P19536 | P19536 [68-85] | [K].AASGTKEDPNLVPSISNK.[R] | 1xHomocysteineMMTS [K6] | 4,49774E-05 | 1990,95724 | Liver/ FAIMS/MMTS |
| P19536 | P19536 [49-67] | [R].EIMIAAQKGLDPYNMLPPK.[A] | 1xOxidation [M15]; 1xHomocysteineMMTS [K8] | 0,0021186 | 2308,12081 | Liver/ FAIMS/MMTS |
| P19536 | P19536 [49-67] | [R].EIMIAAQKGLDPYNMLPPK.[A] | 1xHomocysteineMMTS [K8] | 0,000879882 | 2292,1259 | Liver/ FAIMS/MMTS |
| P19536 | P19536 [74-86] | [K].EDPNLVPSISNKR.[I] | 1xHomocysteineMMTS [K12] | 0,00115861 | 1631,78799 | Liver/ FAIMS/MMTS |
| P19536 | P19536 [49-67] | [R].EIMIAAQKGLDPYNMLPPK.[A] | 1xHomocysteineMMTS [K8] | 0,000879882 | 2292,1259 | Brain/ FAIMS/MMTS |
| P19536 | P19536 [68-85] | [K].AASGTKEDPNLVPSISNK.[R] | 1xHomocysteineMMTS [K6] | 4,49774E-05 | 1990,95724 | Brain/ FAIMS/MMTS |
| P19536 | P19536 [74-86] | [K].EDPNLVPSISNKR.[I] | 1xHomocysteineMMTS [K12] | 0,00115861 | 1631,78799 | Brain/ FAIMS/MMTS |
| P19783 | P19783 [43-63] | [R].DYPLPDVAHVTMLSASQKALK.[E] | 1xHomocysteineIAA [K18] | 4,32935E-05 | 2458,24688 | Liver/ FAIMS/IAA |
| P19783 | P19783 [145-159] | [R].MLDMKANPIQGFSAK.[W] | 1xHomocysteineIAA [K5] | 4,32935E-05 | 1824,88051 | Brain/ No FAIMS/IAA |
| P19783 | P19783 [145-159] | [R].MLDMKANPIQGFSAK.[W] | 1xHomocysteineIAA [K5] | 4,32935E-05 | 1824,88051 | Brain/ FAIMS/IAA |
| P19783 | P19783 [23-41] | [R].AHGSVVKSEDYAFPTYADR.[R] | 1xHomocysteineMMTS [K7] | 4,49774E-05 | 2276,01106 | Liver/ No FAIMS/MMTS |
| P19783 | P19783 [43-63] | [R].DYPLPDVAHVTMLSASQKALK.[E] | 1xHomocysteineMMTS [K18] | 4,49774E-05 | 2447,21313 | Liver/ No FAIMS/MMTS |
| P19783 | P19783 [23-41] | [R].AHGSVVKSEDYAFPTYADR.[R] | 1xHomocysteineMMTS [K7] | 4,49774E-05 | 2276,01106 | Liver/ FAIMS/MMTS |
| P19783 | P19783 [43-63] | [R].DYPLPDVAHVTMLSASQKALK.[E] | 1xOxidation [M12]; 1xHomocysteineMMTS [K18] | 4,49774E-05 | 2463,20805 | Liver/ FAIMS/MMTS |
| P19783 | P19783 [145-159] | [R].MLDMKANPIQGFSAK.[W] | 1xOxidation [M1]; 1xHomocysteineMMTS [K5] | 0,00760806 | 1829,84168 | Liver/ FAIMS/MMTS |
| P19783 | P19783 [145-159] | [R].MLDMKANPIQGFSAK.[W] | 1xHomocysteineMMTS [K5] | 4,49774E-05 | 1813,84676 | Liver/ FAIMS/MMTS |
| P19783 | P19783 [43-63] | [R].DYPLPDVAHVTMLSASQKALK.[E] | 1xHomocysteineMMTS [K18] | 4,49774E-05 | 2447,21313 | Liver/ FAIMS/MMTS |
| P19783 | P19783 [150-168] | [K].ANPIQGFSAKWDYDKNEWK.[K] | 1xHomocysteineMMTS [K10] | 0,000250158 | 2460,11111 | Liver/ FAIMS/MMTS |
| P19783 | P19783 [43-63] | [R].DYPLPDVAHVTMLSASQKALK.[E] | 1xHomocysteineMMTS [K18] | 4,49774E-05 | 2447,21313 | Brain/ No FAIMS/MMTS |
| P19783 | P19783 [23-41] | [R].AHGSVVKSEDYAFPTYADR.[R] | 1xHomocysteineMMTS [K7] | 4,49774E-05 | 2276,01106 | Brain/ FAIMS/MMTS |
| P19783 | P19783 [145-159] | [R].MLDMKANPIQGFSAK.[W] | 1xHomocysteineMMTS [K5] | 4,49774E-05 | 1813,84676 | Brain/ FAIMS/MMTS |
| P19783 | P19783 [43-63] | [R].DYPLPDVAHVTMLSASQKALK.[E] | 1xHomocysteineMMTS [K18] | 4,49774E-05 | 2447,21313 | Brain/ FAIMS/MMTS |
| P19783 | P19783 [150-168] | [K].ANPIQGFSAKWDYDKNEWK.[K] | 1xHomocysteineMMTS [K10] | 0,000250158 | 2460,11111 | Brain/ FAIMS/MMTS |
| P20029 | P20029 [621-634] | [K].KKELEEIVQPIISK.[L] | 1xHomocysteineIAA [K2] | 4,32935E-05 | 1828,02485 | Liver/ No FAIMS/IAA |
| P20029 | P20029 [621-634] | [K].KKELEEIVQPIISK.[L] | 1xHomocysteineIAA [K2] | 4,32935E-05 | 1828,02485 | Liver/ FAIMS/IAA |
| P20029 | P20029 [621-634] | [K].KKELEEIVQPIISK.[L] | 1xHomocysteineIAA [K2] | 4,32935E-05 | 1828,02485 | Brain/ FAIMS/IAA |
| P20029 | P20029 [76-97] | [R].LIGDAAKNQLTSNPENTVFDAK.[R] | 1xHomocysteineMMTS [K7] | 4,49774E-05 | 2509,20613 | Liver/ No FAIMS/MMTS |
| P20029 | P20029 [512-522] | [R].VTAEDKGTGNK.[N] | 1xHomocysteineMMTS [K6] | 0,000322194 | 1282,5766 | Liver/ No FAIMS/MMTS |
| P20029 | P20029 [598-618] | [K].AVEEKIEWLESHQDADIEDFK.[A] | 1xHomocysteineMMTS [K5] | 4,49774E-05 | 2694,20619 | Liver/ No FAIMS/MMTS |
| P20029 | P20029 [115-123] | [K].FLPFKVVEK.[K] | 1xHomocysteineMMTS [K5] | 0,00107839 | 1269,6734 | Liver/ No FAIMS/MMTS |
| P20029 | P20029 [154-164] | [K].MKETAEAYLGK.[K] | 1xHomocysteineMMTS [K2] | 0,000564742 | 1403,63675 | Liver/ No FAIMS/MMTS |
| P20029 | P20029 [326-337] | [R].AKFEELNMDLFR.[S] | 1xHomocysteineMMTS [K2] | 0,000192903 | 1675,76408 | Liver/ No FAIMS/MMTS |
| P20029 | P20029 [603-620] | [K].IEWLESHQDADIEDFKAK.[K] | 1xHomocysteineMMTS [K16] | 4,49774E-05 | 2337,05259 | Liver/ No FAIMS/MMTS |
| P20029 | P20029 [199-215] | [R].IINEPTAAAIAYGLDKR.[E] | 1xHomocysteineMMTS [K16] | 4,49774E-05 | 1979,00888 | Liver/ No FAIMS/MMTS |
| P20029 | P20029 [83-98] | [K].NQLTSNPENTVFDAKR.[L] | 1xHomocysteineMMTS [K15] | 4,49774E-05 | 1996,92152 | Liver/ No FAIMS/MMTS |
| P20029 | P20029 [125-153] | [K].TKPYIQVDIGGGQTKTFAPEEISAMVLTK.[M] | 1xHomocysteineMMTS [K15] | 0,00188612 | 3285,65677 | Liver/ No FAIMS/MMTS |
| P20029 | P20029 [103-119] | [R].TWNDPSVQQDIKFLPFK.[V] | 1xHomocysteineMMTS [K12] | 4,49774E-05 | 2226,07221 | Liver/ No FAIMS/MMTS |
| P20029 | P20029 [564-582] | [R].NELESYAYSLKNQIGDKEK.[L] | 1xHomocysteineMMTS [K11] | 4,49774E-05 | 2392,11592 | Liver/ No FAIMS/MMTS |
| P20029 | P20029 [354-368] | [K].KSDIDEIVLVGGSTR.[I] | 1xHomocysteineMMTS [K1] | 0,00129036 | 1751,86663 | Liver/ No FAIMS/MMTS |
| P20029 | P20029 [448-465] | [K].KSQIFSTASDNQPTVTIK.[V] | 1xHomocysteineMMTS [K1] | 4,49774E-05 | 2128,0413 | Liver/ No FAIMS/MMTS |
| P20029 | P20029 [263-272] | [R].VMEHFIKLYK.[K] | 1xHomocysteineMMTS [K7] | 4,49774E-05 | 1470,7306 | Liver/ FAIMS/MMTS |
| P20029 | P20029 [76-97] | [R].LIGDAAKNQLTSNPENTVFDAK.[R] | 1xHomocysteineMMTS [K7] | 4,49774E-05 | 2509,20613 | Liver/ FAIMS/MMTS |
| P20029 | P20029 [115-123] | [K].FLPFKVVEK.[K] | 1xHomocysteineMMTS [K5] | 0,00107839 | 1269,6734 | Liver/ FAIMS/MMTS |
| P20029 | P20029 [621-634] | [K].KKELEEIVQPIISK.[L] | 1xHomocysteineMMTS [K2] | 4,49774E-05 | 1816,9911 | Liver/ FAIMS/MMTS |
| P20029 | P20029 [199-215] | [R].IINEPTAAAIAYGLDKR.[E] | 1xHomocysteineMMTS [K16] | 4,49774E-05 | 1979,00888 | Liver/ FAIMS/MMTS |
| P20029 | P20029 [83-98] | [K].NQLTSNPENTVFDAKR.[L] | 1xHomocysteineMMTS [K15] | 4,49774E-05 | 1996,92152 | Liver/ FAIMS/MMTS |
| P20029 | P20029 [103-119] | [R].TWNDPSVQQDIKFLPFK.[V] | 1xHomocysteineMMTS [K12] | 4,49774E-05 | 2226,07221 | Liver/ FAIMS/MMTS |
| P20029 | P20029 [564-582] | [R].NELESYAYSLKNQIGDKEK.[L] | 1xHomocysteineMMTS [K11] | 4,49774E-05 | 2392,11592 | Liver/ FAIMS/MMTS |
| P20029 | P20029 [448-465] | [K].KSQIFSTASDNQPTVTIK.[V] | 1xHomocysteineMMTS [K1] | 4,49774E-05 | 2128,0413 | Liver/ FAIMS/MMTS |
| P20029 | P20029 [263-272] | [R].VMEHFIKLYK.[K] | 1xOxidation [M2]; 1xHomocysteineMMTS [K7] | 0,0021186 | 1486,72551 | Liver/ FAIMS/MMTS |
| P20029 | P20029 [542-555] | [R].MVNDAEKFAEEDKK.[L] | 1xOxidation [M1]; 1xHomocysteineMMTS [K7] | 0,00207528 | 1832,78633 | Liver/ FAIMS/MMTS |
| P20029 | P20029 [372-383] | [K].IQQLVKEFFNGK.[E] | 1xHomocysteineMMTS [K6] | 0,000266738 | 1613,81783 | Liver/ FAIMS/MMTS |
| P20029 | P20029 [378-387] | [K].EFFNGKEPSR.[G] | 1xHomocysteineMMTS [K6] | 0,0025135 | 1373,59767 | Liver/ FAIMS/MMTS |

| | | | | | | |
|---|---|---|---|---|---|---|
| P20029 | P20029 [598-618] | [K].AVEEKIEWLESHQDADIEDFK.[A] | 1xHomocysteineMMTS [K5] | 4,49774E-05 | 2694,20619 | Liver/ FAIMS/MMTS |
| P20029 | P20029 [124-139] | [K].KTKPYIQVDIGGGQTK.[T] | 1xHomocysteineMMTS [K3] | 4,49774E-05 | 1895,97176 | Liver/ FAIMS/MMTS |
| P20029 | P20029 [523-533] | [K].NKITITNDQNR.[L] | 1xHomocysteineMMTS [K2] | 0,000282457 | 1479,70426 | Liver/ FAIMS/MMTS |
| P20029 | P20029 [154-164] | [K].MKETAEAYLGK.[K] | 1xHomocysteineMMTS [K2] | 0,000564742 | 1403,63675 | Liver/ FAIMS/MMTS |
| P20029 | P20029 [326-337] | [R].AKFEELNMDLFR.[S] | 1xHomocysteineMMTS [K2] | 0,000192903 | 1675,76408 | Liver/ FAIMS/MMTS |
| P20029 | P20029 [603-620] | [K].IEWLESHQDADIEDFKAK.[K] | 1xHomocysteineMMTS [K16] | 4,49774E-05 | 2337,05259 | Liver/ FAIMS/MMTS |
| P20029 | P20029 [354-368] | [K].KSDIDEIVLVGGSTR.[I] | 1xHomocysteineMMTS [K1] | 0,00129036 | 1751,86663 | Liver/ FAIMS/MMTS |
| P20029 | P20029 [83-98] | [K].NQLTSNPENTVFDAKR.[L] | 1xHomocysteineMMTS [K15] | 4,49774E-05 | 1996,92152 | Brain/ No FAIMS/MMTS |
| P20029 | P20029 [523-533] | [K].NKITITNDQNR.[L] | 1xHomocysteineMMTS [K2] | 0,000282457 | 1479,70426 | Brain/ FAIMS/MMTS |
| P20029 | P20029 [199-215] | [R].IINEPTAAAIAYGLDKR.[E] | 1xHomocysteineMMTS [K16] | 4,49774E-05 | 1979,00888 | Brain/ FAIMS/MMTS |
| P20065 | P20065 [33-45] | [K].NPLPSKETIEQEK.[Q] | 1xHomocysteineIAA [K6] | 4,32935E-05 | 1686,83671 | Brain/ FAIMS/IAA |
| P20108 | P20108 [250-257] | [K].EYFEKVHQ.[-] | 1xHomocysteineIAA [K5] | 0,00106329 | 1253,56193 | Liver/ FAIMS/IAA |
| P20108 | P20108 [186-208] | [R].GLFIIDPNGVVKHLSVNDLPVGR.[S] | 1xHomocysteineIAA [K12] | 4,32935E-05 | 2633,4232 | Brain/ FAIMS/IAA |
| P20108 | P20108 [186-208] | [R].GLFIIDPNGVVKHLSVNDLPVGR.[S] | 1xHomocysteineMMTS [K12] | 0,0021186 | 2622,38946 | Brain/ FAIMS/MMTS |
| P20152 | P20152 [114-122] | [R].FANYIDKVR.[F] | 1xHomocysteineIAA [K7] | 0,0015926 | 1299,65142 | Brain/ FAIMS/IAA |
| P20357 | P20357 [273-291] | [K].KDEWGLAAPISPGPLTPMR.[E] | 1xHomocysteineIAA [K1] | 4,32935E-05 | 2210,10966 | Brain/ FAIMS/IAA |
| P20357 | P20357 [514-528] | [K].TLGKVTSEPEAVSER.[R] | 1xHomocysteineMMTS [K4] | 0,00363897 | 1765,8459 | Brain/ FAIMS/MMTS |
| P20357 | P20357 [273-291] | [K].KDEWGLAAPISPGPLTPMR.[E] | 1xHomocysteineMMTS [K1] | 4,49774E-05 | 2199,07591 | Brain/ FAIMS/MMTS |
| P20801 | P20801 [22-45] | [K].AAFDMFDADGGGDISVKELGTVMR.[M] | 1xHomocysteineIAA [K17] | 4,32935E-05 | 2676,21023 | Liver/ No FAIMS/IAA |
| P20801 | P20801 [22-45] | [K].AAFDMFDADGGGDISVKELGTVMR.[M] | 1xHomocysteineIAA [K17] | 4,32935E-05 | 2676,21023 | Liver/ FAIMS/IAA |
| P20801 | P20801 [22-45] | [K].AAFDMFDADGGGDISVKELGTVMR.[M] | 1xHomocysteineMMTS [K17] | 4,49774E-05 | 2665,17649 | Liver/ No FAIMS/MMTS |
| P20801 | P20801 [22-45] | [K].AAFDMFDADGGGDISVKELGTVMR.[M] | 1xHomocysteineMMTS [K17] | 4,49774E-05 | 2665,17649 | Liver/ FAIMS/MMTS |
| P20917 | P20917 [426-449] | [R].DTVQCLCVVKSNPEPSVAFELPSR.[N] | 2xCarbamidomethyl [C5; C7]; 1xHomocysteineIAA [K10] | 4,32935E-05 | 2906,38451 | Brain/ No FAIMS/IAA |
| P20917 | P20917 [426-449] | [R].DTVQCLCVVKSNPEPSVAFELPSR.[N] | 2xCarbamidomethyl [C5; C7]; 1xHomocysteineIAA [K10] | 4,32935E-05 | 2906,38451 | Brain/ FAIMS/IAA |
| P21107 | P21107 [154-168] | [K].HIAEEADRKYEEVAR.[K] | 1xHomocysteineIAA [K9] | 4,32935E-05 | 1989,9447 | Liver/ FAIMS/IAA |
| P21107 | P21107 [154-168] | [K].HIAEEADRKYEEVAR.[K] | 1xHomocysteineIAA [K9] | 4,32935E-05 | 1989,9447 | Brain/ FAIMS/IAA |
| P21107 | P21107 [151-161] | [K].EAKHIAEEADR.[K] | 1xHomocysteineMMTS [K3] | 0,000361246 | 1431,63551 | Liver/ No FAIMS/MMTS |
| P21107 | P21107 [169-179] | [R].KLVIIEGDLER.[T] | 1xHomocysteineMMTS [K1] | 4,49774E-05 | 1447,76473 | Liver/ No FAIMS/MMTS |
| P21107 | P21107 [151-161] | [K].EAKHIAEEADR.[K] | 1xHomocysteineMMTS [K3] | 0,000361246 | 1431,63551 | Liver/ FAIMS/MMTS |
| P21107 | P21107 [23-36] | [R].AEQAEAEQKQAEER.[S] | 1xHomocysteineMMTS [K9] | 0,000738471 | 1779,76362 | Liver/ FAIMS/MMTS |
| P21107 | P21107 [253-267] | [K].TIDDLEDELYAQKLK.[Y] | 1xHomocysteineMMTS [K13] | 4,49774E-05 | 1956,92929 | Liver/ FAIMS/MMTS |
| P21107 | P21107 [169-179] | [R].KLVIIEGDLER.[T] | 1xHomocysteineMMTS [K1] | 4,49774E-05 | 1447,76473 | Liver/ FAIMS/MMTS |
| P21107 | P21107 [151-161] | [K].EAKHIAEEADR.[K] | 1xHomocysteineMMTS [K3] | 0,000361246 | 1431,63551 | Brain/ No FAIMS/MMTS |
| P21107 | P21107 [151-161] | [K].EAKHIAEEADR.[K] | 1xHomocysteineMMTS [K3] | 0,000361246 | 1431,63551 | Brain/ FAIMS/MMTS |
| P21107; P58771 | P21107 [53-71]; P58771 [52-70] | [K].GTEDELDKYSEALKDAQEK.[L] | 1xHomocysteineIAA [K14] | 4,32935E-05 | 2343,06566 | Liver/ FAIMS/IAA |
| P21107; P58771 | P21107 [142-153]; P58771 [141-152] | [K].MELQEIQLKEAK.[H] | 1xHomocysteineMMTS [K9] | 4,49774E-05 | 1622,79505 | Liver/ No FAIMS/MMTS |
| P21107; P58771 | P21107 [207-218]; P58771 [206-217] | [K].SLEAQAEKYSQK.[E] | 1xHomocysteineMMTS [K8] | 0,000250158 | 1544,70834 | Liver/ No FAIMS/MMTS |
| P21107; P58771 | P21107 [215-227]; P58771 [214-226] | [K].YSQKEDKYEEEIK.[IV] | 1xHomocysteineMMTS [K] | 4,49774E-05 | 1851,81393 | Liver/ No FAIMS/MMTS |
| P21107; P58771 | P21107 [142-153]; P58771 [141-152] | [K].MELQEIQLKEAK.[H] | 1xHomocysteineMMTS [K9] | 4,49774E-05 | 1622,79505 | Liver/ FAIMS/MMTS |
| P21107; P58771 | P21107 [207-218]; P58771 [206-217] | [K].SLEAQAEKYSQK.[E] | 1xHomocysteineMMTS [K8] | 0,000250158 | 1544,70834 | Liver/ FAIMS/MMTS |
| P21107; P58771 | P21107 [215-227]; P58771 [214-226] | [K].YSQKEDKYEEEIK.[IV] | 1xHomocysteineMMTS [K] | 4,49774E-05 | 1851,81393 | Liver/ FAIMS/MMTS |
| P21107; P58771 | P21107 [53-66]; P58771 [52-65] | [K].GTEDELDKYSEALK.[D] | 1xHomocysteineMMTS [K8] | 0,00074635 | 1760,77173 | Liver/ FAIMS/MMTS |
| P21107; P58771 | P21107 [53-71]; P58771 [52-70] | [K].GTEDELDKYSEALKDAQEK.[L] | 1xHomocysteineMMTS [K14] | 0,000250158 | 2332,03192 | Liver/ FAIMS/MMTS |
| P21107; P58771 | P21107 [142-153]; P58771 [141-152] | [K].MELQEIQLKEAK.[H] | 1xHomocysteineMMTS [K9] | 4,49774E-05 | 1622,79505 | Brain/ FAIMS/MMTS |
| P21107; P58771 | P21107 [215-227]; P58771 [214-226] | [K].YSQKEDKYEEEIK.[IV] | 1xHomocysteineMMTS [K] | 4,49774E-05 | 1851,81393 | Brain/ FAIMS/MMTS |
| P21278; P21279 | P21278 [248-256]; P21279 [248-256] | [R].MEESKALFR.[T] | 1xHomocysteineIAA [K5] | 0,00328792 | 1284,6075 | Brain/ FAIMS/IAA |
| P21300; P45377 | P21300 [170-177]; P45377 [170-177] | [R].LLNKPGLK.[H] | 1xHomocysteineIAA [K4] | 0,000305143 | 1056,62341 | Brain/ FAIMS/IAA |
| P21550 | P21550 [413-426] | [R].IEEALGDKAVFAGR.[K] | 1xHomocysteineMMTS [K8] | 0,000231655 | 1638,79782 | Liver/ FAIMS/MMTS |
| P21550; P17183 | P21550 [404-412]; P17183 [404-412] | [R].LAKYNQLMR.[I] | 1xHomocysteineIAA [K3] | 0,000194827 | 1310,67077 | Brain/ No FAIMS/IAA |
| P21550; P17183 | P21550 [404-412]; P17183 [404-412] | [R].LAKYNQLMR.[I] | 1xHomocysteineIAA [K3] | 0,000194827 | 1310,67077 | Brain/ FAIMS/IAA |
| P21550; P17183 | P21550 [404-412]; P17183 [404-412] | [R].LAKYNQLMR.[I] | 1xHomocysteineMMTS [K3] | 0,00236635 | 1299,63703 | Brain/ FAIMS/MMTS |
| P21614 | P21614 [428-439] | [R].TKTPNTSPAELK.[D] | 1xHomocysteineIAA [K2] | 0,00414251 | 1460,74135 | Liver/ FAIMS/IAA |
| P21614 | P21614 [364-373] | [K].VLEPTLKTLR.[E] | 1xHomocysteineMMTS [K7] | 0,00107839 | 1332,73779 | Liver/ No FAIMS/MMTS |
| P21614 | P21614 [364-373] | [K].VLEPTLKTLR.[E] | 1xHomocysteineMMTS [K7] | 0,00107839 | 1332,73779 | Liver/ FAIMS/MMTS |
| P21995 | P21995 [132-142] | [R].FIVFNSKQLGK.[Y] | 1xHomocysteineIAA [K7] | 4,32935E-05 | 1454,78243 | Brain/ FAIMS/IAA |
| P22599 | P22599 [214-235] | [K].KPFDPENTEEAEFHVDKSTTVK.[V] | 1xHomocysteineIAA [K17] | 6,30912E-05 | 2722,26649 | Liver/ FAIMS/IAA |
| P22599 | P22599 [288-297] | [K].ELISKILLNR.[R] | 1xHomocysteineMMTS [K5] | 0,000738471 | 1361,76434 | Liver/ No FAIMS/MMTS |
| P22599 | P22599 [214-235] | [K].KPFDPENTEEAEFHVDKSTTVK.[V] | 1xHomocysteineMMTS [K17] | 0,00138774 | 2711,23274 | Liver/ No FAIMS/MMTS |
| P22599 | P22599 [309-328] | [R].LSISGDYNLKTLMSPLGITR.[I] | 1xOxidation [M13]; 1xHomocysteineMMTS [K10] | 4,49774E-05 | 2358,18659 | Liver/ FAIMS/MMTS |

| | | | | | | |
|---|---|---|---|---|---|---|
| P22599 | P22599 [309-328] | [R].LSISGDYNLKTLMSPLGITR.[I] | 1xHomocysteineMMTS [K10] | 4,49774E-05 | 2342,19167 | Liver/ FAIMS/MMTS |
| P22599 | P22599 [214-230] | [K].KPFDPENTEEAEFHVDK.[S] | 1xHomocysteineMMTS [K1] | 0,000798864 | 2194,94198 | Liver/ FAIMS/MMTS |
| P22599; Q00898 | P22599 [278-292]; Q00898 [278-292] | [K].MQHLEQTLNKELISK.[IF] | 1xOxidation [M1]; 1xHomocysteineMMTS [K10] | 0,00103228 | 1990,97586 | Liver/ FAIMS/MMTS |
| P22599; Q00898 | P22599 [278-292]; Q00898 [278-292] | [K].MQHLEQTLNKELISK.[IF] | 1xHomocysteineMMTS [K10] | 0,00222764 | 1974,98095 | Liver/ FAIMS/MMTS |
| P22599; Q00898; Q00897 | P22599 [182-192]; Q00898 [182-192]; Q00897 [182-192] | [K].GTQGKIVEAVK.[EK] | 1xHomocysteineIAA [K5] | 0,0015926 | 1303,70385 | Liver/ FAIMS/IAA |
| P23242 | P23242 [244-264] | [R].SDPYHATTGPLSPSKDCGSPK.[Y] | 1xCarbamidomethyl [C17]; 1xHomocysteineIAA [K15] | 0,000595249 | 2376,05947 | Brain/ FAIMS/IAA |
| P23506 | P23506 [1-18] | [-].MAWKSGGASHSELIHNLR.[K] | 1xMet-loss+Acetyl [N-Term]; 1xHomocysteineIAA [K4] | 4,32935E-05 | 2079,01887 | Brain/ FAIMS/IAA |
| P23819 | P23819 [347-361] | [R].ALKQVQVEGLSGNIK.[F] | 1xHomocysteineIAA [K3] | 4,32935E-05 | 1757,95783 | Brain/ FAIMS/IAA |
| P23927 | P23927 [93-107] | [K].VLGDVIEVHGKHEER.[Q] | 1xHomocysteineIAA [K11] | 4,32935E-05 | 1890,94906 | Brain/ FAIMS/IAA |
| P23953 | P23953 [287-299] | [R].QKTESELLEISGK.[L] | 1xHomocysteineMMTS [K2] | 0,00129036 | 1624,79207 | Liver/ FAIMS/MMTS |
| P24270 | P24270 [1-16] | [-].MSDSRDPASDQMKQWK.[E] | 1xMet-loss+Acetyl [N-Term]; 1xHomocysteineIAA [K13] | 4,32935E-05 | 1994,86948 | Liver/ FAIMS/IAA |
| P24270 | P24270 [307-320] | [K].DYPLIPVGKLVLNK.[N] | 1xHomocysteineMMTS [K9] | 4,49774E-05 | 1731,9536 | Liver/ No FAIMS/MMTS |
| P24270 | P24270 [469-477] | [K].DAQLFIQKK.[A] | 1xHomocysteineMMTS [K8] | 0,000530332 | 1253,63808 | Liver/ No FAIMS/MMTS |
| P24270 | P24270 [238-252] | [K].TDQGIKNLPVGEAGR.[L] | 1xHomocysteineMMTS [K6] | 4,49774E-05 | 1717,836 | Liver/ No FAIMS/MMTS |
| P24270 | P24270 [73-93] | [R].VVHAKGAGAFGYFEVTHDITR.[Y] | 1xHomocysteineMMTS [K5] | 4,49774E-05 | 2438,17438 | Liver/ No FAIMS/MMTS |
| P24270 | P24270 [445-456] | [R].TFYTKVLNEEER.[K] | 1xHomocysteineMMTS [K5] | 4,49774E-05 | 1691,77675 | Liver/ No FAIMS/MMTS |
| P24270 | P24270 [97-105] | [K].AKVFEHIGK.[R] | 1xHomocysteineMMTS [K2] | 4,49774E-05 | 1191,6013 | Liver/ No FAIMS/MMTS |
| P24270 | P24270 [507-524] | [K].NAIHTYTQAGSHMAAKGK.[A] | 1xHomocysteineMMTS [K16] | 0,000437433 | 2048,9463 | Liver/ No FAIMS/MMTS |
| P24270 | P24270 [157-170] | [R].DAILFPSFIHSQKR.[N] | 1xHomocysteineMMTS [K13] | 4,49774E-05 | 1821,91386 | Liver/ No FAIMS/MMTS |
| P24270 | P24270 [307-320] | [K].DYPLIPVGKLVLNK.[N] | 1xHomocysteineMMTS [K9] | 4,49774E-05 | 1731,9536 | Liver/ FAIMS/MMTS |
| P24270 | P24270 [6-16] | [R].DPASDQMKQWK.[E] | 1xHomocysteineMMTS [K8] | 4,49774E-05 | 1496,63307 | Liver/ FAIMS/MMTS |
| P24270 | P24270 [469-477] | [K].DAQLFIQKK.[A] | 1xHomocysteineMMTS [K8] | 0,000530332 | 1253,63808 | Liver/ FAIMS/MMTS |
| P24270 | P24270 [238-252] | [K].TDQGIKNLPVGEAGR.[L] | 1xHomocysteineMMTS [K6] | 4,49774E-05 | 1717,836 | Liver/ FAIMS/MMTS |
| P24270 | P24270 [445-456] | [R].TFYTKVLNEEER.[K] | 1xHomocysteineMMTS [K5] | 4,49774E-05 | 1691,77675 | Liver/ FAIMS/MMTS |
| P24270 | P24270 [97-105] | [K].AKVFEHIGK.[R] | 1xHomocysteineMMTS [K2] | 4,49774E-05 | 1191,6013 | Liver/ FAIMS/MMTS |
| P24270 | P24270 [157-170] | [R].DAILFPSFIHSQKR.[N] | 1xHomocysteineMMTS [K13] | 4,49774E-05 | 1821,91386 | Liver/ FAIMS/MMTS |
| P24270 | P24270 [20-47] | [R].ASQRPDVLTTGGGNPIGDKLNIMTAGSR.[G] | 1xOxidation [M23]; 1xHomocysteineMMTS [K19] | 4,49774E-05 | 3005,46013 | Liver/ FAIMS/MMTS |
| P24270 | P24270 [1-16] | [-].MSDSRDPASDQMKQWK.[E] | 1xMet-loss+Acetyl [N-Term]; 1xHomocysteineMMTS [K13] | 4,49774E-05 | 1983,83574 | Liver/ FAIMS/MMTS |
| P24270 | P24270 [493-506] | [R].IQALLDKYNAEKPK.[N] | 1xHomocysteineMMTS [K7] | 4,49774E-05 | 1793,92884 | Liver/ FAIMS/MMTS |
| P24270 | P24270 [73-93] | [R].VVHAKGAGAFGYFEVTHDITR.[Y] | 1xHomocysteineMMTS [K5] | 4,49774E-05 | 2438,17438 | Liver/ FAIMS/MMTS |
| P24270 | P24270 [302-315] | [K].VWPHKDYPLIPVGK.[L] | 1xHomocysteineMMTS [K5] | 0,00196333 | 1811,93353 | Liver/ FAIMS/MMTS |
| P24270 | P24270 [234-243] | [K].FHYKTDQGIK.[N] | 1xHomocysteineMMTS [K4] | 0,00872051 | 1399,6497 | Liver/ FAIMS/MMTS |
| P24270 | P24270 [478-492] | [K].AVKNFTDVHPDYGAR.[I] | 1xHomocysteineMMTS [K3] | 4,49774E-05 | 1852,8469 | Liver/ FAIMS/MMTS |
| P24270 | P24270 [20-47] | [R].ASQRPDVLTTGGGNPIGDKLNIMTAGSR.[G] | 1xHomocysteineMMTS [K19] | 4,49774E-05 | 2989,46522 | Liver/ FAIMS/MMTS |
| P24270 | P24270 [507-524] | [K].NAIHTYTQAGSHMAAKGK.[A] | 1xHomocysteineMMTS [K16] | 0,000437433 | 2048,9463 | Liver/ FAIMS/MMTS |
| P24369 | P24369 [191-204] | [R].DKPLKDVIIVDSGK.[I] | 1xHomocysteineIAA [K5] | 9,61721E-05 | 1700,92513 | Liver/ FAIMS/IAA |
| P24369 | P24369 [191-204] | [R].DKPLKDVIIVDSGK.[I] | 1xHomocysteineIAA [K5] | 9,61721E-05 | 1700,92513 | Brain/ FAIMS/IAA |
| P24369 | P24369 [110-122] | [R].GDGTGGKSIYGER.[F] | 1xHomocysteineMMTS [K7] | 0,000530332 | 1459,63042 | Liver/ FAIMS/MMTS |
| P24369 | P24369 [191-204] | [R].DKPLKDVIIVDSGK.[I] | 1xHomocysteineMMTS [K] | 4,49774E-05 | 1689,89139 | Liver/ FAIMS/MMTS |
| P24369 | P24369 [159-171] | [K].TSWLDGKHVVFGK.[V] | 1xHomocysteineMMTS [K7] | 0,0061262 | 1636,79743 | Liver/ FAIMS/MMTS |
| P24369 | P24369 [166-180] | [K].HVVFGKVLEGMDVVR.[K] | 1xHomocysteineMMTS [K6] | 0,000250158 | 1847,93288 | Liver/ FAIMS/MMTS |
| P24369 | P24369 [205-215] | [K].IEVEKPFAIAK.[E] | 1xHomocysteineMMTS [K5] | 4,49774E-05 | 1407,73746 | Liver/ FAIMS/MMTS |
| P24369 | P24369 [96-109] | [R].VIKDFMIQGGDFTR.[G] | 1xHomocysteineMMTS [K3] | 0,00110986 | 1789,84339 | Liver/ FAIMS/MMTS |
| P24369 | P24369 [72-89] | [K].TVDNFVALATGEKGFGYK.[N] | 1xHomocysteineMMTS [K13] | 4,49774E-05 | 2079,98781 | Liver/ FAIMS/MMTS |
| P24369 | P24369 [205-216] | [K].IEVEKPFAIAKE.[-] | 1xHomocysteineMMTS [K11] | 0,00384072 | 1536,78005 | Liver/ FAIMS/MMTS |
| P24457; P11714 | P24457 [273-286]; P11714 [273-286] | [R].NLTDAFLAEIEKAK.[G] | 1xHomocysteineMMTS [K12] | 0,000322194 | 1725,855 | Liver/ FAIMS/MMTS |
| P24472 | P24472 [116-127] | [K].EKEESYDLILSR.[A] | 1xHomocysteineIAA [K2] | 0,00148217 | 1655,79451 | Brain/ FAIMS/IAA |
| P24472; P30115 | P24472 [196-204]; P30115 [196-204] | [K].KFLQPGSQR.[K] | 1xHomocysteineMMTS [K1] | 0,000915247 | 1223,60236 | Liver/ No FAIMS/MMTS |
| P24472; P30115 | P24472 [196-204]; P30115 [196-204] | [K].KFLQPGSQR.[K] | 1xHomocysteineMMTS [K1] | 0,000915247 | 1223,60236 | Liver/ FAIMS/MMTS |
| P24549 | P24549 [242-255] | [K].VAFTGSTQVGKLIK.[E] | 1xHomocysteineIAA [K11] | 4,32935E-05 | 1622,89344 | Liver/ FAIMS/IAA |
| P24549 | P24549 [1-22] | [-].MSSPAQPAVPAPLADLKIQHTK.[I] | 1xMet-loss+Acetyl [N-Term]; 1xHomocysteineMMTS [K17] | 0,000250158 | 2374,22575 | Liver/ No FAIMS/MMTS |
| P24549 | P24549 [411-419] | [K].FKSVDDVIK.[R] | 1xHomocysteineMMTS [K2] | 0,00394153 | 1213,59554 | Liver/ No FAIMS/MMTS |
| P24549 | P24549 [354-363] | [K].ILDLIESGKK.[E] | 1xHomocysteineMMTS [K9] | 0,00285008 | 1278,67961 | Liver/ FAIMS/MMTS |
| P24549 | P24549 [349-362] | [K].EQHDKILDLIESGK.[K] | 1xHomocysteineMMTS [K5] | 0,000437433 | 1787,86663 | Liver/ FAIMS/MMTS |
| P24549 | P24549 [242-255] | [K].VAFTGSTQVGKLIK.[E] | 1xHomocysteineMMTS [K11] | 4,49774E-05 | 1611,8597 | Liver/ FAIMS/MMTS |
| P25444 | P25444 [247-263] | [K].TYSYLTPDLWKETVFTK.[S] | 1xHomocysteineIAA [K11] | 0,0015926 | 2266,11003 | Brain/ FAIMS/IAA |
| P25444 | P25444 [264-279] | [K].SPYQEFTDHLVKTHTR.[V] | 1xHomocysteineMMTS [K12] | 6,70238E-05 | 2121,98445 | Liver/ FAIMS/MMTS |
| P25444 | P25444 [247-263] | [K].TYSYLTPDLWKETVFTK.[S] | 1xHomocysteineMMTS [K11] | 4,49774E-05 | 2255,07629 | Liver/ FAIMS/MMTS |
| P25444 | P25444 [55-68] | [K].AEDKEWIPVTKLGR.[L] | 1xHomocysteineMMTS [K11] | 0,0031519 | 1804,90844 | Liver/ FAIMS/MMTS |
| P25688 | P25688 [272-290] | [K].MGLINKEEVLLPLDNPYGK.[I] | 1xHomocysteineIAA [K6] | 4,32935E-05 | 2317,19305 | Liver/ No FAIMS/IAA |
| P25688 | P25688 [272-290] | [K].MGLINKEEVLLPLDNPYGK.[I] | 1xHomocysteineIAA [K6] | 4,32935E-05 | 2317,19305 | Liver/ FAIMS/IAA |
| P25688 | P25688 [272-290] | [K].MGLINKEEVLLPLDNPYGK.[I] | 1xOxidation [M1]; 1xHomocysteineMMTS [K6] | 4,49774E-05 | 2322,15422 | Liver/ No FAIMS/MMTS |
| P25688 | P25688 [73-82] | [K].NTVHVLAKLR.[G] | 1xHomocysteineMMTS [K8] | 0,000599431 | 1313,71806 | Liver/ No FAIMS/MMTS |
| P25688 | P25688 [221-236] | [K].FAGPYDKGEYSPSVQK.[T] | 1xHomocysteineMMTS [K7] | 4,49774E-05 | 1935,86155 | Liver/ No FAIMS/MMTS |
| P25688 | P25688 [272-290] | [K].MGLINKEEVLLPLDNPYGK.[I] | 1xHomocysteineMMTS [K6] | 4,49774E-05 | 2306,15931 | Liver/ No FAIMS/MMTS |
| P25688 | P25688 [215-227] | [R].DIVLQKFAGPYDK.[G] | 1xHomocysteineMMTS [K6] | 4,49774E-05 | 1656,81241 | Liver/ No FAIMS/MMTS |
| P25688 | P25688 [162-175] | [K].VLKTTQSGFEGFLK.[D] | 1xHomocysteineMMTS [K3] | 4,49774E-05 | 1717,86517 | Liver/ No FAIMS/MMTS |

| | | | | | | |
|---|---|---|---|---|---|---|
| P25688 | P25688 [34-41] | [R].DGKYHSIK.[E] | 1xHomocysteineMMTS [K3] | 0,00182026 | 1110,50706 | Liver/ No FAIMS/MMTS |
| P25688 | P25688 [272-296] | [K].MGLINKEEVLLPLDNPYGKITGTVK.[R] | 1xHomocysteineMMTS [K19] | 4,49774E-05 | 2905,52357 | Liver/ No FAIMS/MMTS |
| P25688 | P25688 [56-80] | [K].DYLHGDNSDIIPTDTIKNTVHVLAK.[L] | 1xHomocysteineMMTS [K17] | 4,49774E-05 | 2942,43865 | Liver/ No FAIMS/MMTS |
| P25688 | P25688 [278-296] | [K].EEVLLPLDNPYGKITGTVK.[R] | 1xHomocysteineMMTS [K13] | 4,49774E-05 | 2249,1556 | Liver/ No FAIMS/MMTS |
| P25688 | P25688 [165-185] | [K].TTQSGFEGFLKDQFTTLPEVK.[D] | 1xHomocysteineMMTS [K11] | 4,49774E-05 | 2536,20982 | Liver/ No FAIMS/MMTS |
| P25688 | P25688 [108-119] | [R].AHVYVEEVPWKR.[F] | 1xHomocysteineMMTS [K11] | 0,000192903 | 1675,80833 | Liver/ No FAIMS/MMTS |
| P25688 | P25688 [176-187] | [K].DQFTTLPEVKDR.[C] | 1xHomocysteineMMTS [K10] | 0,000530332 | 1611,75054 | Liver/ No FAIMS/MMTS |
| P25688 | P25688 [221-236] | [K].FAGPYDKGEYSPSVQK.[T] | 1xHomocysteineMMTS [K7] | 4,49774E-05 | 1935,86155 | Liver/ FAIMS/MMTS |
| P25688 | P25688 [272-290] | [K].MGLINKEEVLLPLDNPYGK.[I] | 1xHomocysteineMMTS [K6] | 4,49774E-05 | 2306,15931 | Liver/ FAIMS/MMTS |
| P25688 | P25688 [215-227] | [R].DIVLQKFAGPYDK.[G] | 1xHomocysteineMMTS [K6] | 4,49774E-05 | 1656,81241 | Liver/ FAIMS/MMTS |
| P25688 | P25688 [162-175] | [K].VLKTTQSGFEGFLK.[D] | 1xHomocysteineMMTS [K3] | 4,49774E-05 | 1717,86517 | Liver/ FAIMS/MMTS |
| P25688 | P25688 [278-296] | [K].EEVLLPLDNPYGKITGTVK.[R] | 1xHomocysteineMMTS [K13] | 4,49774E-05 | 2249,1556 | Liver/ FAIMS/MMTS |
| P25688 | P25688 [165-185] | [K].TTQSGFEGFLKDQFTTLPEVK.[D] | 1xHomocysteineMMTS [K11] | 4,49774E-05 | 2536,20982 | Liver/ FAIMS/MMTS |
| P25688 | P25688 [272-290] | [K].MGLINKEEVLLPLDNPYGK.[I] | 1xOxidation [M1]; 1xHomocysteineMMTS [K6] | 4,49774E-05 | 2322,15422 | Liver/ FAIMS/MMTS |
| P25688 | P25688 [1-18] | [-].MAHYHDNYGKNDEVEFVR.[T] | 1xMet-loss+Acetyl [N-Term]; 1xHomocysteineMMTS [K10] | 4,49774E-05 | 2297,97026 | Liver/ FAIMS/MMTS |
| P25688 | P25688 [73-82] | [K].NTVHVLAKLR.[G] | 1xHomocysteineMMTS [K8] | 0,000599431 | 1313,71806 | Liver/ FAIMS/MMTS |
| P25688 | P25688 [37-52] | [K].YHSIKEVATSVQLTLR.[S] | 1xHomocysteineMMTS [K5] | 4,49774E-05 | 2008,03543 | Liver/ FAIMS/MMTS |
| P25688 | P25688 [24-33] | [K].DMVKVLHIQR.[D] | 1xHomocysteineMMTS [K4] | 4,49774E-05 | 1401,71634 | Liver/ FAIMS/MMTS |
| P25688 | P25688 [272-296] | [K].MGLINKEEVLLPLDNPYGKITGTVK.[R] | 1xHomocysteineMMTS [K19] | 4,49774E-05 | 2905,52357 | Liver/ FAIMS/MMTS |
| P25688 | P25688 [56-80] | [K].DYLHGDNSDIIPTDTIKNTVHVLAK.[L] | 1xHomocysteineMMTS [K17] | 4,49774E-05 | 2942,43865 | Liver/ FAIMS/MMTS |
| P25688 | P25688 [108-119] | [R].AHVYVEEVPWKR.[F] | 1xHomocysteineMMTS [K11] | 0,000192903 | 1675,80833 | Liver/ FAIMS/MMTS |
| P25688 | P25688 [148-161] | [R].NGPPVIHSGIKDLK.[V] | 1xHomocysteineMMTS [K11] | 4,49774E-05 | 1637,85019 | Liver/ FAIMS/MMTS |
| P25688 | P25688 [176-187] | [K].DQFTTLPEVKDR.[C] | 1xHomocysteineMMTS [K10] | 0,000530332 | 1611,75054 | Liver/ FAIMS/MMTS |
| P25688 | P25688 [55-72] | [K].KDYLHGDNSDIIPTDTIK.[N] | 1xHomocysteineMMTS [K1] | 4,49774E-05 | 2208,03113 | Liver/ FAIMS/MMTS |
| P26041; P26040 | P26041 [28-40]; P26040 [28-40] | [K].QLFDQVVKTIGLR.[E] | 1xHomocysteineIAA [K8] | 0,006291 | 1690,93089 | Brain/ FAIMS/IAA |
| P26043 | P26043 [361-370] | [K].AQKELEEQTR.[K] | 1xHomocysteineMMTS [K3] | 0,000829616 | 1394,64026 | Liver/ FAIMS/MMTS |
| P26043 | P26043 [163-171] | [K].LTKEQWEER.[I] | 1xHomocysteineMMTS [K3] | 0,0062711 | 1381,62388 | Liver/ FAIMS/MMTS |
| P26043; P26041; P26040 | P26043 [234-246]; P26041 [234-246]; P26040 [234-246] | [KR].LTPKIGFPWSEIR.[N] | 1xHomocysteineIAA [K4] | 4,32935E-05 | 1717,90942 | Brain/ FAIMS/IAA |
| P26350 | P26350 [1-18] | [-].MSDAAVDTSSEITTKDLK.[E] | 1xMet-loss+Acetyl [N-Term]; 1xHomocysteineIAA [K15] | 4,32935E-05 | 1996,93794 | Brain/ No FAIMS/IAA |
| P26350 | P26350 [91-104] | [R].VAEDDEDDDVDTKK.[Q] | 1xHomocysteineMMTS [K13] | 4,49774E-05 | 1756,68879 | Brain/ FAIMS/MMTS |
| P26443 | P26443 [85-92] | [K].LVEDLKTR.[E] | 1xHomocysteineIAA [K6] | 4,32935E-05 | 1147,61397 | Liver/ FAIMS/IAA |
| P26443 | P26443 [212-231] | [K].KGFIGPGIDVPAPDMSTGER.[E] | 1xHomocysteineIAA [K1] | 4,32935E-05 | 2218,0631 | Brain/ FAIMS/IAA |
| P26443 | P26443 [192-203] | [K].NYTDNELEKITR.[R] | 1xHomocysteineMMTS [K9] | 0,00542903 | 1658,75127 | Liver/ FAIMS/MMTS |
| P26443 | P26443 [497-516] | [R].ISGASEKDIVHSGLAYTMER.[S] | 1xHomocysteineMMTS [K7] | 4,49774E-05 | 2327,08285 | Liver/ FAIMS/MMTS |
| P26443 | P26443 [85-92] | [K].LVEDLKTR.[E] | 1xHomocysteineMMTS [K6] | 0,0018995 | 1136,58023 | Liver/ FAIMS/MMTS |
| P26443 | P26443 [188-200] | [K].INPKNYTDNELEK.[I] | 1xHomocysteineMMTS [K4] | 0,000689517 | 1740,79313 | Liver/ FAIMS/MMTS |
| P26443 | P26443 [212-231] | [K].KGFIGPGIDVPAPDMSTGER.[E] | 1xHomocysteineMMTS [K1] | 4,49774E-05 | 2207,02936 | Liver/ FAIMS/MMTS |
| P26443 | P26443 [497-516] | [R].ISGASEKDIVHSGLAYTMER.[S] | 1xHomocysteineMMTS [K7] | 4,49774E-05 | 2327,08285 | Brain/ FAIMS/MMTS |
| P26443 | P26443 [524-535] | [R].TAMKYNLGLDLR.[T] | 1xHomocysteineMMTS [K4] | 0,0004709 | 1557,7586 | Brain/ FAIMS/MMTS |
| P26645 | P26645 [8-30] | [K].TAAKGEATAERPGEAAVASSPSK.[A] | 1xHomocysteineIAA [K4] | 4,32935E-05 | 2360,15106 | Brain/ No FAIMS/IAA |
| P26645 | P26645 [8-30] | [K].TAAKGEATAERPGEAAVASSPSK.[A] | 1xHomocysteineIAA [K4] | 4,32935E-05 | 2360,15106 | Brain/ FAIMS/IAA |
| P26645 | P26645 [8-30] | [K].TAAKGEATAERPGEAAVASSPSK.[A] | 1xHomocysteineMMTS [K4] | 0,00271031 | 2349,11732 | Brain/ FAIMS/MMTS |
| P26645 | P26645 [131-146] | [K].AEDGAAPSPSSETPKK.[K] | 1xHomocysteineMMTS [K15] | 4,49774E-05 | 1734,76731 | Brain/ FAIMS/MMTS |
| P26883 | P26883 [19-36] | [K].RGQTCVVHYTGMLEDGKK.[F] | 1xCarbamidomethyl [C5]; 1xHomocysteineIAA [K17] | 4,32935E-05 | 2253,0573 | Brain/ FAIMS/IAA |
| P26883 | P26883 [49-58] | [K].FTLGKQEVIR.[G] | 1xHomocysteineMMTS [K5] | 0,00074635 | 1353,70174 | Brain/ No FAIMS/MMTS |
| P26883 | P26883 [49-58] | [K].FTLGKQEVIR.[G] | 1xHomocysteineMMTS [K5] | 0,00074635 | 1353,70174 | Brain/ FAIMS/MMTS |
| P27546 | P27546 [822-833] | [R].MTAKSASADLSR.[S] | 1xHomocysteineIAA [K4] | 9,61721E-05 | 1411,66681 | Brain/ FAIMS/IAA |
| P27573 | P27573 [131-149] | [K].NPPDIVGKTSQVTLYVFEK.[V] | 1xHomocysteineIAA [K8] | 0,00722104 | 2309,18459 | Brain/ FAIMS/IAA |
| P27600; P27601 | P27600 [69-76]; P27601 [62-69] | [K].STFLKQMR.[I] | 1xHomocysteineIAA [K5] | 0,000348152 | 1184,59146 | Brain/ FAIMS/IAA |
| P27659 | P27659 [298-312] | [K].LIKNNASTDYDLSDK.[S] | 1xHomocysteineIAA [K3] | 4,32935E-05 | 1870,88512 | Liver/ FAIMS/IAA |
| P27659 | P27659 [298-312] | [K].LIKNNASTDYDLSDK.[S] | 1xHomocysteineIAA [K3] | 4,32935E-05 | 1870,88512 | Brain/ FAIMS/IAA |
| P27659 | P27659 [363-373] | [K].IDLKFIDTTSK.[F] | 1xHomocysteineMMTS [K4] | 0,000133296 | 1443,7222 | Liver/ No FAIMS/MMTS |
| P27659 | P27659 [287-297] | [K].IGQGYLIKDGK.[L] | 1xHomocysteineMMTS [K8] | 0,000946421 | 1354,68575 | Liver/ FAIMS/MMTS |
| P27659 | P27659 [363-373] | [K].IDLKFIDTTSK.[F] | 1xHomocysteineMMTS [K4] | 0,000133296 | 1443,7222 | Liver/ FAIMS/MMTS |
| P27659 | P27659 [284-294] | [K].IYKIGQGYLIK.[D] | 1xHomocysteineMMTS [K3] | 0,00074635 | 1458,78474 | Liver/ FAIMS/MMTS |
| P27659 | P27659 [298-312] | [K].LIKNNASTDYDLSDK.[S] | 1xHomocysteineMMTS [K3] | 4,49774E-05 | 1859,85137 | Liver/ FAIMS/MMTS |
| P27773 | P27773 [162-179] | [R].DLFSDGHSEFLKAASNLR.[D] | 1xHomocysteineIAA [K12] | 4,32935E-05 | 2181,03933 | Liver/ FAIMS/IAA |
| P27773 | P27773 [130-140] | [K].KQAGPASVPLR.[T] | 1xHomocysteineIAA [K1] | 4,32935E-05 | 1297,70452 | Brain/ No FAIMS/IAA |
| P27773 | P27773 [130-140] | [K].KQAGPASVPLR.[T] | 1xHomocysteineIAA [K1] | 4,32935E-05 | 1297,70452 | Brain/ FAIMS/IAA |
| P27773 | P27773 [364-379] | [R].YLKSEPIPESNEGPVK.[V] | 1xHomocysteineMMTS [K3] | 4,49774E-05 | 1949,93471 | Liver/ No FAIMS/MMTS |
| P27773 | P27773 [423-433] | [K].LSKDPNIVIAK.[M] | 1xHomocysteineMMTS [K3] | 0,00239411 | 1360,7327 | Liver/ No FAIMS/MMTS |
| P27773 | P27773 [290-304] | [K].FLDAGHKLNFAVASR.[K] | 1xHomocysteineMMTS [K7] | 4,49774E-05 | 1808,89346 | Liver/ FAIMS/MMTS |
| P27773 | P27773 [148-161] | [K].FISDKDASVVGFFR.[D] | 1xHomocysteineMMTS [K5] | 4,49774E-05 | 1750,82912 | Liver/ FAIMS/MMTS |
| P27773 | P27773 [364-379] | [R].YLKSEPIPESNEGPVK.[V] | 1xHomocysteineMMTS [K3] | 4,49774E-05 | 1949,93471 | Liver/ FAIMS/MMTS |
| P27773 | P27773 [333-344] | [K].GEKFVMQEEFSR.[D] | 1xOxidation [M6]; 1xHomocysteineMMTS [K3] | 0,000133296 | 1665,70696 | Liver/ FAIMS/MMTS |
| P27773 | P27773 [423-433] | [K].LSKDPNIVIAK.[M] | 1xHomocysteineMMTS [K3] | 0,00239411 | 1360,7327 | Liver/ FAIMS/MMTS |
| P27773 | P27773 [483-496] | [R].EATNPPIIQEEKPK.[K] | 1xHomocysteineMMTS [K12] | 0,00177533 | 1756,86082 | Liver/ FAIMS/MMTS |
| P27773 | P27773 [305-329] | [R].KTFSHELSDFGLESTTGEVPVVAIR.[T] | 1xHomocysteineMMTS [K1] | 4,49774E-05 | 2882,40629 | Liver/ FAIMS/MMTS |
| P28352 | P28352 [58-72] | [K].SATLKICSWNVDGLR.[A] | 1xCarbamidomethyl [C7]; 1xHomocysteineIAA [K5] | 0,000305143 | 1893,93096 | Brain/ FAIMS/IAA |
| P28474 | P28474 [108-115] | [K].TNLCQKIR.[V] | 1xCarbamidomethyl [C4]; 1xHomocysteineIAA [K6] | 0,00166226 | 1206,60817 | Brain/ FAIMS/IAA |

| | | | | | | |
|---|---|---|---|---|---|---|
| P28652 | P28652 [34-48] | [K].LCTGHEYAAKIINTK.[K] | 1xCarbamidomethyl [C2]; 1xHomocysteineIAA [K10] | 0,000364707 | 1892,93571 | Brain/ FAIMS/IAA |
| P28652; Q923T9 | P28652 [247-260]; Q923T9 [247-260] | [K].NLINQMLTINPAKR.[I] | 1xHomocysteineIAA [K13] | 9,61721E-05 | 1799,96187 | Brain/ FAIMS/IAA |
| P28663 | P28663 [246-264] | [K].KLLEAHEEQNSEAYTEAVK.[E] | 1xHomocysteineIAA [K1] | 4,32935E-05 | 2363,11837 | Brain/ FAIMS/IAA |
| P28663 | P28663 [223-239] | [K].LALEKYEEMFPAFTDSR.[E] | 1xHomocysteineMMTS [K5] | 0,00130669 | 2209,99666 | Brain/ FAIMS/MMTS |
| P28663 | P28663 [1-8] | [-].MDNAGKER.[E] | 1xAcetyl [N-Term]; 1xHomocysteineMMTS [K6] | 0,000211896 | 1125,44856 | Brain/ FAIMS/MMTS |
| P28665 | P28665 [645-659] | [K].HTNLVPHGTEKDVYR.[Y] | 1xHomocysteineIAA [K11] | 0,000584831 | 1939,9443 | Liver/ FAIMS/IAA |
| P28665 | P28665 [645-659] | [K].HTNLVPHGTEKDVYR.[Y] | 1xHomocysteineIAA [K11] | 0,000584831 | 1939,9443 | Brain/ FAIMS/IAA |
| P28665 | P28665 [645-659] | [K].HTNLVPHGTEKDVYR.[Y] | 1xHomocysteineMMTS [K11] | 0,000400718 | 1928,91056 | Liver/ No FAIMS/MMTS |
| P28665 | P28665 [645-659] | [K].HTNLVPHGTEKDVYR.[Y] | 1xHomocysteineMMTS [K11] | 0,000400718 | 1928,91056 | Liver/ FAIMS/MMTS |
| P28665 | P28665 [721-734] | [R].KDPSSNDPLTETIR.[K] | 1xHomocysteineMMTS [K1] | 0,0014351 | 1735,79895 | Liver/ FAIMS/MMTS |
| P28667 | P28667 [145-166] | [K].AAATPESQEPQAKGAEASAASK.[E] | 1xHomocysteineIAA [K13] | 4,32935E-05 | 2274,06666 | Brain/ FAIMS/IAA |
| P28740 | P28740 [330-343] | [K].NQDCSKGIYALAAR.[D] | 1xCarbamidomethyl [C4]; 1xHomocysteineIAA [K6] | 0,00684106 | 1740,8156 | Brain/ FAIMS/IAA |
| P28843 | P28843 [40-52] | [R].TYSLADYLKSTFR.[V] | 1xHomocysteineIAA [K9] | 0,00199357 | 1738,84688 | Liver/ FAIMS/IAA |
| P28843 | P28843 [606-616] | [R].QFVKMGFVDSK.[R] | 1xHomocysteineMMTS [K4] | 0,00427174 | 1448,67347 | Liver/ FAIMS/MMTS |
| P29391 | P29391 [77-92] | [R].ALFQDVQKPSQDEWGK.[T] | 1xHomocysteineIAA [K8] | 4,32935E-05 | 2049,96985 | Liver/ No FAIMS/IAA |
| P29391 | P29391 [77-92] | [R].ALFQDVQKPSQDEWGK.[T] | 1xHomocysteineIAA [K8] | 4,32935E-05 | 2049,96985 | Liver/ FAIMS/IAA |
| P29391 | P29391 [77-92] | [R].ALFQDVQKPSQDEWGK.[T] | 1xHomocysteineIAA [K8] | 4,32935E-05 | 2049,96985 | Brain/ FAIMS/IAA |
| P29391 | P29391 [144-153] | [K].KMGNHLTNLR.[R] | 1xOxidation [M2]; 1xHomocysteineMMTS [K1] | 0,000112422 | 1362,64391 | Liver/ No FAIMS/MMTS |
| P29391 | P29391 [144-153] | [K].KMGNHLTNLR.[R] | 1xHomocysteineMMTS [K1] | 8,93346E-05 | 1346,64899 | Liver/ No FAIMS/MMTS |
| P29391 | P29391 [77-92] | [R].ALFQDVQKPSQDEWGK.[T] | 1xHomocysteineMMTS [K8] | 4,49774E-05 | 2038,93611 | Liver/ FAIMS/MMTS |
| P29391 | P29391 [144-153] | [K].KMGNHLTNLR.[R] | 1xOxidation [M2]; 1xHomocysteineMMTS [K1] | 0,000112422 | 1362,64391 | Liver/ FAIMS/MMTS |
| P29391 | P29391 [77-105] | [R].ALFQDVQKPSQDEWGKTQEAMEAALAMEK.[N] | 1xHomocysteineMMTS [K16] | 0,000437433 | 3442,57859 | Liver/ FAIMS/MMTS |
| P29391 | P29391 [144-153] | [K].KMGNHLTNLR.[R] | 1xHomocysteineMMTS [K1] | 8,93346E-05 | 1346,64899 | Liver/ FAIMS/MMTS |
| P29595 | P29595 [43-54] | [R].LIYSGKQMNDEK.[T] | 1xHomocysteineIAA [K6] | 0,00012645 | 1599,75054 | Brain/ FAIMS/IAA |
| P29758 | P29758 [32-46] | [K].KTEQGPPSSEYIFER.[E] | 1xHomocysteineIAA [K1] | 4,32935E-05 | 1941,9011 | Liver/ FAIMS/IAA |
| P29758 | P29758 [414-426] | [R].LAPPLVIKEDEIR.[E] | 1xHomocysteineMMTS [K8] | 4,49774E-05 | 1655,88591 | Liver/ No FAIMS/MMTS |
| P29758 | P29758 [32-46] | [K].KTEQGPPSSEYIFER.[E] | 1xHomocysteineMMTS [K1] | 0,000173718 | 1930,86736 | Liver/ FAIMS/MMTS |
| P29758 | P29758 [359-372] | [K].ELMKLPSDVVTSVR.[G] | 1xOxidation [M3]; 1xHomocysteineMMTS [K4] | 0,0033721 | 1752,86927 | Liver/ FAIMS/MMTS |
| P29758 | P29758 [414-426] | [R].LAPPLVIKEDEIR.[E] | 1xHomocysteineMMTS [K8] | 4,49774E-05 | 1655,88591 | Liver/ FAIMS/MMTS |
| P29758 | P29758 [359-372] | [K].ELMKLPSDVVTSVR.[G] | 1xHomocysteineMMTS [K4] | 4,49774E-05 | 1736,87436 | Liver/ FAIMS/MMTS |
| P30115 | P30115 [1-13] | [-].MAGKPVLHYFDGR.[G] | 1xMet-loss+Acetyl [N-Term]; 1xHomocysteineIAA [K4] | 4,32935E-05 | 1575,77366 | Liver/ No FAIMS/IAA |
| P30115 | P30115 [1-13] | [-].MAGKPVLHYFDGR.[G] | 1xMet-loss+Acetyl [N-Term]; 1xHomocysteineIAA [K4] | 4,32935E-05 | 1575,77366 | Liver/ FAIMS/IAA |
| P30115 | P30115 [1-13] | [-].MAGKPVLHYFDGR.[G] | 1xMet-loss+Acetyl [N-Term]; 1xHomocysteineMMTS [K4] | 4,49774E-05 | 1564,73991 | Liver/ No FAIMS/MMTS |
| P30115 | P30115 [188-196] | [R].VSNLPTVKK.[F] | 1xHomocysteineMMTS [K8] | 0,000724331 | 1148,61661 | Liver/ No FAIMS/MMTS |
| P30115 | P30115 [132-141] | [R].YFPAFEKVLK.[S] | 1xHomocysteineMMTS [K7] | 0,000250158 | 1404,70543 | Liver/ No FAIMS/MMTS |
| P30115 | P30115 [118-125] | [K].EASLAKIK.[E] | 1xHomocysteineMMTS [K6] | 0,00804194 | 1022,5373 | Liver/ No FAIMS/MMTS |
| P30115 | P30115 [65-78] | [K].LVQTKAILNYIASK.[Y] | 1xHomocysteineMMTS [K5] | 4,49774E-05 | 1724,94376 | Liver/ No FAIMS/MMTS |
| P30115 | P30115 [205-211] | [R].KPFDDAK.[C] | 1xHomocysteineMMTS [K1] | 0,00466355 | 983,4325 | Liver/ No FAIMS/MMTS |
| P30115 | P30115 [1-13] | [-].MAGKPVLHYFDGR.[G] | 1xMet-loss+Acetyl [N-Term]; 1xHomocysteineMMTS [K4] | 4,49774E-05 | 1564,73991 | Liver/ FAIMS/MMTS |
| P30115 | P30115 [132-141] | [R].YFPAFEKVLK.[S] | 1xHomocysteineMMTS [K7] | 0,000250158 | 1404,70543 | Liver/ FAIMS/MMTS |
| P30115 | P30115 [65-78] | [K].LVQTKAILNYIASK.[Y] | 1xHomocysteineMMTS [K5] | 4,49774E-05 | 1724,94376 | Liver/ FAIMS/MMTS |
| P30115 | P30115 [46-69] | [R].SDGSLMFQQVPMVEIDGMKLVQTK.[A] | 2xOxidation [M6; M12]; 1xHomocysteineMMTS [K19] | 0,0018995 | 2876,33709 | Liver/ FAIMS/MMTS |
| P30115 | P30115 [46-69] | [R].SDGSLMFQQVPMVEIDGMKLVQTK.[A] | 1xOxidation [M12]; 1xHomocysteineMMTS [K19] | 4,49774E-05 | 2860,34218 | Liver/ FAIMS/MMTS |
| P30115 | P30115 [205-217] | [R].KPFDDAKCVESAK.[K] | 1xMethylthio [C8]; 1xHomocysteineMMTS [K7] | 0,00482082 | 1646,70452 | Liver/ FAIMS/MMTS |
| P30115 | P30115 [188-196] | [R].VSNLPTVKK.[F] | 1xHomocysteineMMTS [K8] | 0,000724331 | 1148,61661 | Liver/ FAIMS/MMTS |
| P30275 | P30275 [250-270] | [R].GIWHNNEKSFLIWVNEEDHTR.[V] | 1xHomocysteineIAA [K8] | 4,32935E-05 | 2798,31036 | Brain/ No FAIMS/IAA |
| P30275 | P30275 [250-270] | [R].GIWHNNEKSFLIWVNEEDHTR.[V] | 1xHomocysteineIAA [K8] | 4,32935E-05 | 2798,31036 | Brain/ FAIMS/IAA |
| P30275 | P30275 [138-149] | [R].TMKHTTDLDASK.[I] | 1xHomocysteineMMTS [K3] | 0,000211896 | 1510,66985 | Brain/ FAIMS/MMTS |
| P30275 | P30275 [191-206] | [R].VVVDALSGLKGDLAGR.[Y] | 1xHomocysteineMMTS [K10] | 4,49774E-05 | 1732,90844 | Brain/ FAIMS/MMTS |
| P30416 | P30416 [427-444] | [K].AEVAAGDHPTDAEMKGER.[N] | 1xHomocysteineIAA [K15] | 0,00033298 | 2057,90151 | Liver/ FAIMS/IAA |
| P31001; P20152 | P31001 [105-117]; P20152 [101-113] | [R].TNEKVELQELNDR.[F] | 1xHomocysteineMMTS [K4] | 0,000282457 | 1750,80984 | Liver/ FAIMS/MMTS |
| P31648 | P31648 [1-28] | [-].MATDNSKVADGQISTEVSEAPVASDKPK.[T] | 1xMet-loss+Acetyl [N-Term]; 1xHomocysteineIAA [K7] | 4,32935E-05 | 2960,41533 | Brain/ No FAIMS/IAA |
| P31648 | P31648 [1-28] | [-].MATDNSKVADGQISTEVSEAPVASDKPK.[T] | 1xMet-loss+Acetyl [N-Term]; 1xHomocysteineIAA [K7] | 4,32935E-05 | 2960,41533 | Brain/ FAIMS/IAA |
| P31650 | P31650 [601-612] | [R].TVTVNDCEAKVK.[G] | 1xCarbamidomethyl [C7]; 1xHomocysteineIAA [K10] | 4,32935E-05 | 1537,73489 | Brain/ No FAIMS/IAA |
| P31650 | P31650 [601-612] | [R].TVTVNDCEAKVK.[G] | 1xCarbamidomethyl [C7]; 1xHomocysteineIAA [K10] | 4,32935E-05 | 1537,73489 | Brain/ FAIMS/IAA |
| P31650 | P31650 [593-600] | [R].GKLGASPR.[T] | 1xHomocysteineMMTS [K2] | 0,0089835 | 948,47537 | Brain/ No FAIMS/MMTS |
| P31786 | P31786 [34-53] | [K].QATVGDVNTDRPGLLDLKGK.[A] | 1xHomocysteineIAA [K18] | 4,32935E-05 | 2271,17615 | Liver/ No FAIMS/IAA |
| P31786 | P31786 [34-53] | [K].QATVGDVNTDRPGLLDLKGK.[A] | 1xHomocysteineIAA [K18] | 4,32935E-05 | 2271,17615 | Liver/ FAIMS/IAA |
| P31786 | P31786 [34-53] | [K].QATVGDVNTDRPGLLDLKGK.[A] | 1xHomocysteineIAA [K18] | 4,32935E-05 | 2271,17615 | Brain/ FAIMS/IAA |

| | | | | | | |
|---|---|---|---|---|---|---|
| P31786 | P31786 [16-33] | [R].LKTQPTDEEMLFIYSHFK.[Q] | 1xOxidation [M10]; 1xHomocysteineMMTS [K2] | 0,000530332 | 2406,11784 | Liver/ No FAIMS/MMTS |
| P31786 | P31786 [1-14] | [-].MSQAEFDKAAEEVK.[R] | 1xMet-loss+Acetyl [N-Term]; 1xHomocysteineMMTS [K8] | 4,49774E-05 | 1656,72438 | Liver/ No FAIMS/MMTS |
| P31786 | P31786 [1-15] | [-].MSQAEFDKAAEEVKR.[L] | 1xMet-loss+Acetyl [N-Term]; 1xHomocysteineMMTS [K] | 4,49774E-05 | 1812,82549 | Liver/ No FAIMS/MMTS |
| P31786 | P31786 [56-63] | [K].WDSWNKLK.[G] | 1xHomocysteineMMTS [K6] | 0,00210568 | 1239,56491 | Liver/ No FAIMS/MMTS |
| P31786 | P31786 [73-82] | [K].TYVEKVDELK.[K] | 1xHomocysteineMMTS [K5] | 4,49774E-05 | 1386,66435 | Liver/ No FAIMS/MMTS |
| P31786 | P31786 [73-83] | [K].TYVEKVDELKK.[K] | 1xHomocysteineMMTS [K5] | 4,49774E-05 | 1514,75931 | Liver/ No FAIMS/MMTS |
| P31786 | P31786 [68-77] | [K].ESAMKTYVEK.[V] | 1xHomocysteineMMTS [K5] | 4,49774E-05 | 1348,59456 | Liver/ No FAIMS/MMTS |
| P31786 | P31786 [54-61] | [K].AKWDSWNK.[L] | 1xHomocysteineMMTS [K2] | 0,00207528 | 1197,51796 | Liver/ No FAIMS/MMTS |
| P31786 | P31786 [16-33] | [R].LKTQPTDEEMLFIYSHFK.[Q] | 1xHomocysteineMMTS [K2] | 4,49774E-05 | 2390,12292 | Liver/ No FAIMS/MMTS |
| P31786 | P31786 [34-53] | [K].QATVGDVNTDRPGLLDLKGK.[A] | 1xHomocysteineMMTS [K18] | 4,49774E-05 | 2260,14241 | Liver/ No FAIMS/MMTS |
| P31786 | P31786 [18-51] | [K].TQPTDEEMLFIYSHFKQATVGDVNTDRPGLLDLK.[G] | 1xHomocysteineMMTS [K16] | 4,49774E-05 | 4041,93948 | Liver/ No FAIMS/MMTS |
| P31786 | P31786 [18-51] | [K].TQPTDEEMLFIYSHFKQATVGDVNTDRPGLLDLK.[G] | 1xOxidation [M8]; 1xHomocysteineMMTS [K16] | 4,49774E-05 | 4057,9344 | Liver/ FAIMS/MMTS |
| P31786 | P31786 [16-33] | [R].LKTQPTDEEMLFIYSHFK.[Q] | 1xOxidation [M10]; 1xHomocysteineMMTS [K2] | 0,000530332 | 2406,11784 | Liver/ FAIMS/MMTS |
| P31786 | P31786 [1-14] | [-].MSQAEFDKAAEEVK.[R] | 1xMet-loss+Acetyl [N-Term]; 1xHomocysteineMMTS [K8] | 4,49774E-05 | 1656,72438 | Liver/ FAIMS/MMTS |
| P31786 | P31786 [1-15] | [-].MSQAEFDKAAEEVKR.[L] | 1xMet-loss+Acetyl [N-Term]; 1xHomocysteineMMTS [K] | 4,49774E-05 | 1812,82549 | Liver/ FAIMS/MMTS |
| P31786 | P31786 [56-63] | [K].WDSWNKLK.[G] | 1xHomocysteineMMTS [K6] | 0,00210568 | 1239,56491 | Liver/ FAIMS/MMTS |
| P31786 | P31786 [73-82] | [K].TYVEKVDELK.[K] | 1xHomocysteineMMTS [K5] | 4,49774E-05 | 1386,66435 | Liver/ FAIMS/MMTS |
| P31786 | P31786 [73-83] | [K].TYVEKVDELKK.[K] | 1xHomocysteineMMTS [K5] | 4,49774E-05 | 1514,75931 | Liver/ FAIMS/MMTS |
| P31786 | P31786 [54-61] | [K].AKWDSWNK.[L] | 1xHomocysteineMMTS [K2] | 0,00207528 | 1197,51796 | Liver/ FAIMS/MMTS |
| P31786 | P31786 [16-33] | [R].LKTQPTDEEMLFIYSHFK.[Q] | 1xHomocysteineMMTS [K2] | 4,49774E-05 | 2390,12292 | Liver/ FAIMS/MMTS |
| P31786 | P31786 [34-53] | [K].QATVGDVNTDRPGLLDLKGK.[A] | 1xHomocysteineMMTS [K18] | 4,49774E-05 | 2260,14241 | Liver/ FAIMS/MMTS |
| P31786 | P31786 [18-51] | [K].TQPTDEEMLFIYSHFKQATVGDVNTDRPGLLDLK.[G] | 1xHomocysteineMMTS [K16] | 4,49774E-05 | 4041,93948 | Liver/ FAIMS/MMTS |
| P31786 | P31786 [68-77] | [K].ESAMKTYVEK.[V] | 1xHomocysteineMMTS [K5] | 4,49774E-05 | 1348,59456 | Liver/ FAIMS/MMTS |
| P31786 | P31786 [1-15] | [-].MSQAEFDKAAEEVKR.[L] | 1xMet-loss+Acetyl [N-Term]; 1xHomocysteineMMTS [K] | 4,49774E-05 | 1812,82549 | Brain/ No FAIMS/MMTS |
| P31786 | P31786 [16-33] | [R].LKTQPTDEEMLFIYSHFK.[Q] | 1xHomocysteineMMTS [K2] | 4,49774E-05 | 2390,12292 | Brain/ No FAIMS/MMTS |
| P31786 | P31786 [1-15] | [-].MSQAEFDKAAEEVKR.[L] | 1xMet-loss+Acetyl [N-Term]; 1xHomocysteineMMTS [K] | 4,49774E-05 | 1812,82549 | Brain/ FAIMS/MMTS |
| P31786 | P31786 [73-82] | [K].TYVEKVDELK.[K] | 1xHomocysteineMMTS [K5] | 4,49774E-05 | 1386,66435 | Brain/ FAIMS/MMTS |
| P31786 | P31786 [68-77] | [K].ESAMKTYVEK.[V] | 1xHomocysteineMMTS [K5] | 4,49774E-05 | 1348,59456 | Brain/ FAIMS/MMTS |
| P31786 | P31786 [54-61] | [K].AKWDSWNK.[L] | 1xHomocysteineMMTS [K2] | 0,00207528 | 1197,51796 | Brain/ FAIMS/MMTS |
| P31786 | P31786 [16-33] | [R].LKTQPTDEEMLFIYSHFK.[Q] | 1xHomocysteineMMTS [K2] | 4,49774E-05 | 2390,12292 | Brain/ FAIMS/MMTS |
| P31786 | P31786 [34-53] | [K].QATVGDVNTDRPGLLDLKGK.[A] | 1xHomocysteineMMTS [K18] | 4,49774E-05 | 2260,14241 | Brain/ FAIMS/MMTS |
| P31786 | P31786 [18-51] | [K].TQPTDEEMLFIYSHFKQATVGDVNTDRPGLLDLK.[G] | 1xHomocysteineMMTS [K16] | 4,49774E-05 | 4041,93948 | Brain/ FAIMS/MMTS |
| P31938 | P31938 [169-181] | [K].VSIAVIKGLTYLR.[E] | 1xHomocysteineIAA [K7] | 9,61721E-05 | 1606,93491 | Brain/ FAIMS/IAA |
| P32020 | P32020 [443-454] | [K].KLEEEGEQFVKK.[I] | 1xHomocysteineIAA [K11] | 4,32935E-05 | 1637,82033 | Liver/ No FAIMS/IAA |
| P32020 | P32020 [443-454] | [K].KLEEEGEQFVKK.[I] | 1xHomocysteineIAA [K11] | 4,32935E-05 | 1637,82033 | Liver/ FAIMS/IAA |
| P32020 | P32020 [512-524] | [K].MNPQSAFFQGKLK.[I] | 1xOxidation [M1]; 1xHomocysteineMMTS [K11] | 4,49774E-05 | 1674,78007 | Liver/ No FAIMS/MMTS |
| P32020 | P32020 [455-464] | [K].IGGIFAFKVK.[D] | 1xHomocysteineMMTS [K8] | 0,0004709 | 1242,67373 | Liver/ No FAIMS/MMTS |
| P32020 | P32020 [463-479] | [K].VKDGPGGKEATWVVDVK.[N] | 1xHomocysteineMMTS [K8] | 4,49774E-05 | 1947,96668 | Liver/ No FAIMS/MMTS |
| P32020 | P32020 [137-150] | [R].TTPTDKHIEVLIDK.[Y] | 1xHomocysteineMMTS [K6] | 4,49774E-05 | 1772,89212 | Liver/ No FAIMS/MMTS |
| P32020 | P32020 [178-187] | [K].VEHFAKIGWK.[N] | 1xHomocysteineMMTS [K6] | 0,000530332 | 1377,68061 | Liver/ No FAIMS/MMTS |
| P32020 | P32020 [465-479] | [K].DGPGGKEATWVVDVK.[N] | 1xHomocysteineMMTS [K6] | 4,49774E-05 | 1720,8033 | Liver/ No FAIMS/MMTS |
| P32020 | P32020 [433-442] | [K].ANLVFKEIEK.[K] | 1xHomocysteineMMTS [K6] | 0,000231655 | 1353,6905 | Liver/ No FAIMS/MMTS |
| P32020 | P32020 [174-183] | [K].YGTKVEHFAK.[I] | 1xHomocysteineMMTS [K4] | 0,000781536 | 1342,62824 | Liver/ No FAIMS/MMTS |
| P32020 | P32020 [271-282] | [K].SIIKVVGYDMSK.[E] | 1xHomocysteineMMTS [K4] | 4,49774E-05 | 1502,74155 | Liver/ No FAIMS/MMTS |
| P32020 | P32020 [512-524] | [K].MNPQSAFFQGKLK.[I] | 1xHomocysteineMMTS [K11] | 4,49774E-05 | 1658,78515 | Liver/ No FAIMS/MMTS |
| P32020 | P32020 [443-454] | [K].KLEEEGEQFVKK.[I] | 1xHomocysteineMMTS [K11] | 4,49774E-05 | 1626,78659 | Liver/ No FAIMS/MMTS |
| P32020 | P32020 [535-546] | [K].LQNLQLQPGKAK.[L] | 1xHomocysteineMMTS [K10] | 0,00074635 | 1500,80251 | Liver/ No FAIMS/MMTS |
| P32020 | P32020 [525-544] | [K].IAGNMGLAMKLQNLQLQPGK.[A] | 1xHomocysteineMMTS [K10] | 4,49774E-05 | 2288,17458 | Liver/ No FAIMS/MMTS |
| P32020 | P32020 [444-454] | [K].LEEEGEQFVKK.[I] | 1xHomocysteineMMTS [K10] | 4,49774E-05 | 1498,69163 | Liver/ No FAIMS/MMTS |
| P32020 | P32020 [443-453] | [K].KLEEEGEQFVK.[K] | 1xHomocysteineMMTS [K1] | 0,000282457 | 1498,69163 | Liver/ No FAIMS/MMTS |
| P32020 | P32020 [454-462] | [K].KIGGIFAFK.[V] | 1xHomocysteineMMTS [K1] | 4,49774E-05 | 1143,60532 | Liver/ No FAIMS/MMTS |
| P32020 | P32020 [483-492] | [K].GSVLPNSDKK.[A] | 1xHomocysteineMMTS [K] | 0,000282457 | 1207,58095 | Liver/ No FAIMS/MMTS |
| P32020 | P32020 [512-524] | [K].MNPQSAFFQGKLK.[I] | 1xOxidation [M1]; 1xHomocysteineMMTS [K11] | 4,49774E-05 | 1674,78007 | Liver/ FAIMS/MMTS |
| P32020 | P32020 [525-544] | [K].IAGNMGLAMKLQNLQLQPGK.[A] | 1xOxidation [M]; 1xHomocysteineMMTS [K10] | 4,49774E-05 | 2304,1695 | Liver/ FAIMS/MMTS |
| P32020 | P32020 [334-353] | [R].GDNTYGGKWVINPSGGLISK.[G] | 1xHomocysteineMMTS [K8] | 4,49774E-05 | 2226,06818 | Liver/ FAIMS/MMTS |
| P32020 | P32020 [455-464] | [K].IGGIFAFKVK.[D] | 1xHomocysteineMMTS [K8] | 0,0004709 | 1242,67373 | Liver/ FAIMS/MMTS |
| P32020 | P32020 [465-479] | [K].DGPGGKEATWVVDVK.[N] | 1xHomocysteineMMTS [K6] | 4,49774E-05 | 1720,8033 | Liver/ FAIMS/MMTS |
| P32020 | P32020 [433-442] | [K].ANLVFKEIEK.[K] | 1xHomocysteineMMTS [K6] | 0,000231655 | 1353,6905 | Liver/ FAIMS/MMTS |
| P32020 | P32020 [512-524] | [K].MNPQSAFFQGKLK.[I] | 1xHomocysteineMMTS [K11] | 4,49774E-05 | 1658,78515 | Liver/ FAIMS/MMTS |
| P32020 | P32020 [443-454] | [K].KLEEEGEQFVKK.[I] | 1xHomocysteineMMTS [K11] | 4,49774E-05 | 1626,78659 | Liver/ FAIMS/MMTS |
| P32020 | P32020 [535-546] | [K].LQNLQLQPGKAK.[L] | 1xHomocysteineMMTS [K10] | 0,00074635 | 1500,80251 | Liver/ FAIMS/MMTS |
| P32020 | P32020 [525-544] | [K].IAGNMGLAMKLQNLQLQPGK.[A] | 1xHomocysteineMMTS [K10] | 4,49774E-05 | 2288,17458 | Liver/ FAIMS/MMTS |
| P32020 | P32020 [443-453] | [K].KLEEEGEQFVK.[K] | 1xHomocysteineMMTS [K1] | 0,000282457 | 1498,69163 | Liver/ FAIMS/MMTS |
| P32020 | P32020 [454-462] | [K].KIGGIFAFK.[V] | 1xHomocysteineMMTS [K1] | 4,49774E-05 | 1143,60532 | Liver/ FAIMS/MMTS |
| P32020 | P32020 [483-492] | [K].GSVLPNSDKK.[A] | 1xHomocysteineMMTS [K] | 0,000282457 | 1207,58095 | Liver/ FAIMS/MMTS |
| P32020 | P32020 [463-479] | [K].VKDGPGGKEATWVVDVK.[N] | 1xHomocysteineMMTS [K8] | 4,49774E-05 | 1947,96668 | Liver/ FAIMS/MMTS |
| P32020 | P32020 [34-45] | [R].DYPDMAKEAGQK.[A] | 1xHomocysteineMMTS [K7] | 4,49774E-05 | 1515,62765 | Liver/ FAIMS/MMTS |

| | | | | | | |
|---|---|---|---|---|---|---|
| P32020 | P32020 [137-150] | [R].TTPTDKHIEVLIDK.[Y] | 1xHomocysteineMMTS [K6] | 4,49774E-05 | 1772,89212 | Liver/ FAIMS/MMTS |
| P32020 | P32020 [178-187] | [K].VEHFAKIGWK.[N] | 1xHomocysteineMMTS [K6] | 0,000530332 | 1377,68061 | Liver/ FAIMS/MMTS |
| P32020 | P32020 [127-136] | [R].GSIGTKFSDR.[T] | 1xHomocysteineMMTS [K6] | 0,0033721 | 1230,56055 | Liver/ FAIMS/MMTS |
| P32020 | P32020 [174-183] | [K].YGTKVEHFAK.[I] | 1xHomocysteineMMTS [K4] | 0,000781536 | 1342,62824 | Liver/ FAIMS/MMTS |
| P32020 | P32020 [271-282] | [K].SIIKVVGYDMSK.[E] | 1xHomocysteineMMTS [K4] | 4,49774E-05 | 1502,74155 | Liver/ FAIMS/MMTS |
| P32020 | P32020 [444-454] | [K].LEEEGEQFVKK.[I] | 1xHomocysteineMMTS [K10] | 4,49774E-05 | 1498,69163 | Liver/ FAIMS/MMTS |
| P32037 | P32037 [249-262] | [R].MSQEKQVTVLELFR.[S] | 1xHomocysteineIAA [K5] | 4,32935E-05 | 1881,95611 | Brain/ FAIMS/IAA |
| P32848 | P32848 [46-55] | [K].KVFHILDKDK.[S] | 1xHomocysteineIAA [K1] | 4,32935E-05 | 1416,76678 | Liver/ No FAIMS/IAA |
| P32848 | P32848 [46-55] | [K].KVFHILDKDK.[S] | 1xHomocysteineIAA [K1] | 4,32935E-05 | 1416,76678 | Liver/ FAIMS/IAA |
| P32848 | P32848 [1-14] | [-].MSMTDVLSAEDIKK.[A] | 1xMet-loss+Acetyl [N-Term]; 1xHomocysteineMMTS [K13] | 4,49774E-05 | 1641,75324 | Liver/ No FAIMS/MMTS |
| P32848 | P32848 [77-84] | [R].DLSAKETK.[T] | 1xHomocysteineMMTS [K5] | 0,00338636 | 1054,49074 | Liver/ No FAIMS/MMTS |
| P32848 | P32848 [82-97] | [K].ETKTLLAAGDKDGDGK.[I] | 1xHomocysteineMMTS [K3] | 4,49774E-05 | 1781,84081 | Liver/ No FAIMS/MMTS |
| P32848 | P32848 [15-29] | [K].AIGAFAAADSFDHKK.[F] | 1xHomocysteineMMTS [K14] | 4,49774E-05 | 1711,79307 | Liver/ No FAIMS/MMTS |
| P32848 | P32848 [46-55] | [K].KVFHILDKDK.[S] | 1xHomocysteineMMTS [K1] | 6,70238E-05 | 1405,73304 | Liver/ No FAIMS/MMTS |
| P32848 | P32848 [29-37] | [K].KFFQMVGLK.[K] | 1xHomocysteineMMTS [K1] | 0,00624351 | 1260,63015 | Liver/ No FAIMS/MMTS |
| P32848 | P32848 [1-14] | [-].MSMTDVLSAEDIKK.[A] | 1xMet-loss+Acetyl [N-Term]; 1xHomocysteineMMTS [K13] | 4,49774E-05 | 1641,75324 | Liver/ FAIMS/MMTS |
| P32848 | P32848 [85-97] | [K].TLLAAGDKDGDGK.[I] | 1xHomocysteineMMTS [K8] | 4,49774E-05 | 1423,65558 | Liver/ FAIMS/MMTS |
| P32848 | P32848 [54-69] | [K].DKSGFIEEDELGSILK.[G] | 1xHomocysteineMMTS [K2] | 4,49774E-05 | 1942,91364 | Liver/ FAIMS/MMTS |
| P32848 | P32848 [15-29] | [K].AIGAFAAADSFDHKK.[F] | 1xHomocysteineMMTS [K14] | 4,49774E-05 | 1711,79307 | Liver/ FAIMS/MMTS |
| P32848 | P32848 [30-38] | [K].FFQMVGLKK.[K] | 1xOxidation [M4]; 1xHomocysteineMMTS [K8] | 0,00791642 | 1276,62507 | Liver/ FAIMS/MMTS |
| P32848 | P32848 [77-84] | [R].DLSAKETK.[T] | 1xHomocysteineMMTS [K5] | 0,00338636 | 1054,49074 | Liver/ FAIMS/MMTS |
| P32848 | P32848 [82-97] | [K].ETKTLLAAGDKDGDGK.[I] | 1xHomocysteineMMTS [K3] | 4,49774E-05 | 1781,84081 | Liver/ FAIMS/MMTS |
| P32848 | P32848 [46-55] | [K].KVFHILDKDK.[S] | 1xHomocysteineMMTS [K1] | 6,70238E-05 | 1405,73304 | Liver/ FAIMS/MMTS |
| P32848 | P32848 [29-37] | [K].KFFQMVGLK.[K] | 1xHomocysteineMMTS [K1] | 0,00624351 | 1260,63015 | Liver/ FAIMS/MMTS |
| P32848 | P32848 [1-14] | [-].MSMTDVLSAEDIKK.[A] | 1xMet-loss+Acetyl [N-Term]; 1xHomocysteineMMTS [K13] | 4,49774E-05 | 1641,75324 | Brain/ No FAIMS/MMTS |
| P32848 | P32848 [85-97] | [K].TLLAAGDKDGDGK.[I] | 1xHomocysteineMMTS [K8] | 4,49774E-05 | 1423,65558 | Brain/ FAIMS/MMTS |
| P34022 | P34022 [58-70] | [R].FASENDLPEWKER.[G] | 1xHomocysteineIAA [K11] | 9,61721E-05 | 1794,81156 | Brain/ FAIMS/IAA |
| P34022 | P34022 [179-189] | [K].VAEKLEALSVR.[E] | 1xHomocysteineMMTS [K4] | 0,000929608 | 1377,72287 | Liver/ FAIMS/MMTS |
| P34022 | P34022 [179-189] | [K].VAEKLEALSVR.[E] | 1xHomocysteineMMTS [K4] | 0,000929608 | 1377,72287 | Brain/ No FAIMS/MMTS |
| P34884 | P34884 [75-87] | [R].NYSKLLCGLLSDR.[L] | 1xCarbamidomethyl [C7]; 1xHomocysteineIAA [K4] | 4,32935E-05 | 1712,84584 | Liver/ FAIMS/IAA |
| P34884 | P34884 [75-87] | [R].NYSKLLCGLLSDR.[L] | 1xCarbamidomethyl [C7]; 1xHomocysteineIAA [K4] | 4,32935E-05 | 1712,84584 | Brain/ No FAIMS/IAA |
| P34884 | P34884 [75-87] | [R].NYSKLLCGLLSDR.[L] | 1xCarbamidomethyl [C7]; 1xHomocysteineIAA [K4] | 4,32935E-05 | 1712,84584 | Brain/ FAIMS/IAA |
| P34927 | P34927 [114-134] | [K].DLTEDHSSLLLHVKQLVSDVR.[S] | 1xHomocysteineIAA [K14] | 4,32935E-05 | 2578,32936 | Liver/ FAIMS/IAA |
| P34927 | P34927 [114-134] | [K].DLTEDHSSLLLHVKQLVSDVR.[S] | 1xHomocysteineMMTS [K14] | 4,49774E-05 | 2567,29562 | Liver/ FAIMS/MMTS |
| P35278; Q9CQD1; P61021 | P35278 [24-40]; Q9CQD1 [23-39]; P61021 [23-39] | [K].LVLLGESAVGKSSLVLR.[F] | 1xHomocysteineIAA [K11] | 4,32935E-05 | 1915,10449 | Brain/ FAIMS/IAA |
| P35282 | P35282 [166-177] | [K].GIEELFLDLCKR.[M] | 1xCarbamidomethyl [C10]; 1xHomocysteineIAA [K11] | 0,00459966 | 1666,82912 | Brain/ FAIMS/IAA |
| P35486 | P35486 [275-288] | [R].SGKGPILMELQTYR.[Y] | 1xHomocysteineIAA [K3] | 4,32935E-05 | 1766,89279 | Brain/ FAIMS/IAA |
| P35486; P35487 | P35486 [74-83]; P35487 [75-84] | [R].MELKADQLYK.[Q] | 1xHomocysteineIAA [K4] | 0,000318237 | 1412,69123 | Brain/ FAIMS/IAA |
| P35505 | P35505 [369-381] | [K].GTKAIDVEQGQTR.[T] | 1xHomocysteineMMTS [K3] | 0,000738471 | 1565,74104 | Liver/ FAIMS/MMTS |
| P35564 | P35564 [49-62] | [K].SDASTPPSPKVTYK.[A] | 1xHomocysteineIAA [K10] | 4,32935E-05 | 1651,7996 | Liver/ FAIMS/IAA |
| P35564 | P35564 [402-416] | [R].KIPNPDFFEDLEPFK.[M] | 1xHomocysteineIAA [K1] | 4,32935E-05 | 2009,96773 | Brain/ No FAIMS/IAA |
| P35564 | P35564 [402-416] | [R].KIPNPDFFEDLEPFK.[M] | 1xHomocysteineIAA [K1] | 4,32935E-05 | 2009,96773 | Brain/ FAIMS/IAA |
| P35564 | P35564 [115-126] | [K].LPGDKGLVLMSR.[A] | 1xHomocysteineMMTS [K5] | 4,49774E-05 | 1448,74222 | Liver/ No FAIMS/MMTS |
| P35564 | P35564 [47-58] | [K].SKSDASTPPSPK.[V] | 1xHomocysteineMMTS [K2] | 0,008514 | 1364,61846 | Liver/ No FAIMS/MMTS |
| P35564 | P35564 [115-126] | [K].LPGDKGLVLMSR.[A] | 1xHomocysteineMMTS [K5] | 4,49774E-05 | 1448,74222 | Liver/ FAIMS/MMTS |
| P35564 | P35564 [59-78] | [K].VTYKAPVPTGEVYFADSFDR.[G] | 1xHomocysteineMMTS [K4] | 8,93346E-05 | 2425,12028 | Liver/ FAIMS/MMTS |
| P35564 | P35564 [49-62] | [K].SDASTPPSPKVTYK.[A] | 1xHomocysteineMMTS [K10] | 0,000266738 | 1640,76585 | Liver/ FAIMS/MMTS |
| P35564 | P35564 [115-126] | [K].LPGDKGLVLMSR.[A] | 1xOxidation [M10]; 1xHomocysteineMMTS [K5] | 0,00531384 | 1464,73714 | Liver/ FAIMS/MMTS |
| P35564 | P35564 [294-312] | [K].IADPDAVKPDDWDEDAPSK.[I] | 1xHomocysteineMMTS [K8] | 4,49774E-05 | 2246,95802 | Liver/ FAIMS/MMTS |
| P35564 | P35564 [168-183] | [K].LLSKTAELSLDQFHDK.[T] | 1xHomocysteineMMTS [K4] | 0,00563517 | 2007,98781 | Liver/ FAIMS/MMTS |
| P35564 | P35564 [172-194] | [K].TAELSLDQFHDKTPYTIMFGPDK.[C] | 1xHomocysteineMMTS [K12] | 0,00325546 | 2817,29323 | Liver/ FAIMS/MMTS |
| P35564 | P35564 [402-416] | [R].KIPNPDFFEDLEPFK.[M] | 1xHomocysteineMMTS [K1] | 0,00631419 | 1998,93398 | Liver/ FAIMS/MMTS |
| P35564 | P35564 [284-293] | [R].KPEDWDERPK.[I] | 1xHomocysteineMMTS [K1] | 4,49774E-05 | 1462,64535 | Liver/ FAIMS/MMTS |
| P35564 | P35564 [518-535] | [K].TDAPQPDVKDEEGKEEEK.[N] | 1xHomocysteineMMTS [K] | 4,49774E-05 | 2206,94785 | Liver/ FAIMS/MMTS |
| P35564 | P35564 [115-126] | [K].LPGDKGLVLMSR.[A] | 1xHomocysteineMMTS [K5] | 4,49774E-05 | 1448,74222 | Brain/ FAIMS/MMTS |
| P35564 | P35564 [172-194] | [K].TAELSLDQFHDKTPYTIMFGPDK.[C] | 1xHomocysteineMMTS [K12] | 0,00325546 | 2817,29323 | Brain/ FAIMS/MMTS |
| P35564 | P35564 [49-62] | [K].SDASTPPSPKVTYK.[A] | 1xHomocysteineMMTS [K10] | 0,000266738 | 1640,76585 | Brain/ FAIMS/MMTS |
| P35564 | P35564 [518-535] | [K].TDAPQPDVKDEEGKEEEK.[N] | 1xHomocysteineMMTS [K] | 4,49774E-05 | 2206,94785 | Brain/ FAIMS/MMTS |
| P35700 | P35700 [94-110] | [K].QGGLGPMNIPLISDPKR.[T] | 1xHomocysteineIAA [K16] | 4,32935E-05 | 1967,02011 | Liver/ FAIMS/IAA |
| P35700 | P35700 [94-110] | [K].QGGLGPMNIPLISDPKR.[T] | 1xHomocysteineIAA [K16] | 4,32935E-05 | 1967,02011 | Brain/ FAIMS/IAA |
| P35700 | P35700 [94-110] | [K].QGGLGPMNIPLISDPKR.[T] | 1xOxidation [M7]; 1xHomocysteineMMTS [K16] | 4,49774E-05 | 1971,98128 | Liver/ No FAIMS/MMTS |
| P35700 | P35700 [1-16] | [-].MSSGNAKIGYPAPNFK.[A] | 1xMet-loss+Acetyl [N-Term]; 1xHomocysteineMMTS [K7] | 4,49774E-05 | 1755,81929 | Liver/ No FAIMS/MMTS |
| P35700 | P35700 [129-140] | [R].GLFIIDDKGILR.[Q] | 1xHomocysteineMMTS [K8] | 4,49774E-05 | 1522,81202 | Liver/ No FAIMS/MMTS |
| P35700 | P35700 [94-110] | [K].QGGLGPMNIPLISDPKR.[T] | 1xHomocysteineMMTS [K16] | 4,49774E-05 | 1955,98637 | Liver/ No FAIMS/MMTS |
| P35700 | P35700 [17-35] | [K].ATAVMPDGQFKDISLSEYK.[G] | 1xHomocysteineMMTS [K11] | 4,49774E-05 | 2263,04434 | Liver/ No FAIMS/MMTS |
| P35700 | P35700 [1-16] | [-].MSSGNAKIGYPAPNFK.[A] | 1xMet-loss+Acetyl [N-Term]; 1xHomocysteineMMTS [K7] | 4,49774E-05 | 1755,81929 | Liver/ FAIMS/MMTS |
| P35700 | P35700 [129-140] | [R].GLFIIDDKGILR.[Q] | 1xHomocysteineMMTS [K8] | 4,49774E-05 | 1522,81202 | Liver/ FAIMS/MMTS |

| | | | | | | |
|---|---|---|---|---|---|---|
| P35700 | P35700 [94-110] | [K].QGGLGPMNIPLISDPKR.[T] | 1xHomocysteineMMTS [K16] | 4,49774E-05 | 1955,98637 | Liver/ FAIMS/MMTS |
| P35700 | P35700 [17-35] | [K].ATAVMPDGQFKDISLSEYK.[G] | 1xHomocysteineMMTS [K11] | 4,49774E-05 | 2263,04434 | Liver/ FAIMS/MMTS |
| P35700 | P35700 [93-109] | [K].KQGGLGPMNIPLISDPK.[R] | 1xHomocysteineMMTS [K1] | 4,49774E-05 | 1927,98022 | Liver/ FAIMS/MMTS |
| P35700 | P35700 [93-109] | [K].KQGGLGPMNIPLISDPK.[R] | 1xOxidation [M8]; 1xHomocysteineMMTS [K1] | 4,49774E-05 | 1943,97514 | Liver/ FAIMS/MMTS |
| P35700 | P35700 [94-110] | [K].QGGLGPMNIPLISDPKR.[T] | 1xOxidation [M7]; 1xHomocysteineMMTS [K16] | 4,49774E-05 | 1971,98128 | Liver/ FAIMS/MMTS |
| P35700 | P35700 [17-35] | [K].ATAVMPDGQFKDISLSEYK.[G] | 1xOxidation [M5]; 1xHomocysteineMMTS [K11] | 4,49774E-05 | 2279,03925 | Liver/ FAIMS/MMTS |
| P35700 | P35700 [8-27] | [K].IGYPAPNFKATAVMPDGQFK.[D] | 1xOxidation [M14]; 1xHomocysteineMMTS [K9] | 0,00106681 | 2331,09704 | Liver/ FAIMS/MMTS |
| P35700 | P35700 [8-27] | [K].IGYPAPNFKATAVMPDGQFK.[D] | 1xHomocysteineMMTS [K9] | 0,000211896 | 2315,10213 | Liver/ FAIMS/MMTS |
| P35700 | P35700 [28-37] | [K].DISLSEYKGK.[Y] | 1xHomocysteineMMTS [K8] | 0,000342214 | 1302,60683 | Liver/ FAIMS/MMTS |
| P35700 | P35700 [111-128] | [R].TIAQDYGVLKADEGISFR.[G] | 1xHomocysteineMMTS [K10] | 4,49774E-05 | 2146,03074 | Liver/ FAIMS/MMTS |
| P35700 | P35700 [129-140] | [R].GLFIIDDKGILR.[Q] | 1xHomocysteineMMTS [K8] | 4,49774E-05 | 1522,81202 | Brain/ No FAIMS/MMTS |
| P35700 | P35700 [1-16] | [-].MSSGNAKIGYPAPNFK.[A] | 1xMet-loss+Acetyl [N-Term]; 1xHomocysteineMMTS [K7] | 4,49774E-05 | 1755,81929 | Brain/ FAIMS/MMTS |
| P35700 | P35700 [129-140] | [R].GLFIIDDKGILR.[Q] | 1xHomocysteineMMTS [K8] | 4,49774E-05 | 1522,81202 | Brain/ FAIMS/MMTS |
| P35700 | P35700 [94-110] | [K].QGGLGPMNIPLISDPKR.[T] | 1xHomocysteineMMTS [K16] | 4,49774E-05 | 1955,98637 | Brain/ FAIMS/MMTS |
| P35700 | P35700 [111-128] | [R].TIAQDYGVLKADEGISFR.[G] | 1xHomocysteineMMTS [K10] | 4,49774E-05 | 2146,03074 | Brain/ FAIMS/MMTS |
| P35700 | P35700 [93-109] | [K].KQGGLGPMNIPLISDPK.[R] | 1xHomocysteineMMTS [K1] | 4,49774E-05 | 1927,98022 | Brain/ FAIMS/MMTS |
| P35762 | P35762 [145-171] | [K].AVVKTFHETLNCCGSNALTTLTTTILR.[N] | 2xCarbamidomethyl [C12; C13]; 1xHomocysteineIAA [K4] | 6,30912E-05 | 3195,5959 | Brain/ FAIMS/IAA |
| P35802 | P35802 [190-200] | [K].KICTASENFLR.[M] | 1xCarbamidomethyl [C3]; 1xHomocysteineIAA [K1] | 4,32935E-05 | 1512,72974 | Brain/ No FAIMS/IAA |
| P35802 | P35802 [190-200] | [K].KICTASENFLR.[M] | 1xCarbamidomethyl [C3]; 1xHomocysteineIAA [K1] | 4,32935E-05 | 1512,72974 | Brain/ FAIMS/IAA |
| P35802 | P35802 [256-269] | [K].SKEEQELHDIHSTR.[S] | 1xHomocysteineMMTS [K2] | 4,49774E-05 | 1871,83746 | Brain/ No FAIMS/MMTS |
| P35802 | P35802 [256-269] | [K].SKEEQELHDIHSTR.[S] | 1xHomocysteineMMTS [K2] | 4,49774E-05 | 1871,83746 | Brain/ FAIMS/MMTS |
| P35803 | P35803 [1-19] | [-].MKPAMETAAEENTEQSQER.[K] | 1xOxidation [M5]; 1xMet-loss [N-Term]; 1xHomocysteineIAA [K2] | 4,32935E-05 | 2238,96015 | Brain/ No FAIMS/IAA |
| P35803 | P35803 [1-19] | [-].MKPAMETAAEENTEQSQER.[K] | 1xOxidation [M5]; 1xMet-loss [N-Term]; 1xHomocysteineIAA [K2] | 4,32935E-05 | 2238,96015 | Brain/ FAIMS/IAA |
| P35803 | P35803 [1-19] | [-].MKPAMETAAEENTEQSQER.[K] | 1xMet-loss+Acetyl [N-Term]; 1xHomocysteineMMTS [K2] | 0,000282457 | 2253,94206 | Brain/ FAIMS/MMTS |
| P35803 | P35803 [307-319] | [K].AKEEQELQDIQSR.[S] | 1xHomocysteineMMTS [K2] | 4,49774E-05 | 1736,79419 | Brain/ FAIMS/MMTS |
| P35980 | P35980 [116-132] | [K].AGGKILTFDQLALESPK.[G] | 1xHomocysteineIAA [K4] | 4,32935E-05 | 1962,03647 | Brain/ FAIMS/IAA |
| P35980 | P35980 [116-132] | [K].AGGKILTFDQLALESPK.[G] | 1xHomocysteineMMTS [K4] | 4,49774E-05 | 1951,00273 | Liver/ FAIMS/MMTS |
| P35980 | P35980 [120-134] | [K].ILTFDQLALESPKGR.[G] | 1xHomocysteineMMTS [K13] | 0,0046895 | 1850,9503 | Liver/ FAIMS/MMTS |
| P36536; Q9CQC9 | P36536 [25-38]; Q9CQC9 [25-38] | [K].SGKLVFLGLDNAGK.[T] | 1xHomocysteineIAA [K3] | 9,61721E-05 | 1592,84649 | Brain/ FAIMS/IAA |
| P38060 | P38060 [180-190] | [K].VSPAKVAEVAK.[K] | 1xHomocysteineIAA [K5] | 0,00467963 | 1272,69803 | Liver/ FAIMS/IAA |
| P38060 | P38060 [49-58] | [K].SIVPTPVKIR.[L] | 1xHomocysteineMMTS [K8] | 0,000282457 | 1272,71666 | Liver/ FAIMS/MMTS |
| P38647 | P38647 [203-218] | [R].QATKDAGQISGLNVLR.[V] | 1xHomocysteineIAA [K4] | 4,32935E-05 | 1844,96471 | Liver/ FAIMS/IAA |
| P38647 | P38647 [468-485] | [K].KSQVFSTAADGQTQVEIK.[V] | 1xHomocysteineIAA [K1] | 4,32935E-05 | 2111,04374 | Brain/ FAIMS/IAA |
| P38647 | P38647 [127-138] | [R].RYDDPEVQKDTK.[N] | 1xHomocysteineMMTS [K9] | 0,00340109 | 1656,73562 | Liver/ No FAIMS/MMTS |
| P38647 | P38647 [620-634] | [R].ALLAGKDSETGENIR.[Q] | 1xHomocysteineMMTS [K6] | 4,49774E-05 | 1736,83058 | Liver/ No FAIMS/MMTS |
| P38647 | P38647 [203-218] | [R].QATKDAGQISGLNVLR.[V] | 1xHomocysteineMMTS [K4] | 0,000599431 | 1833,93096 | Liver/ No FAIMS/MMTS |
| P38647 | P38647 [564-574] | [K].NAEKYAEEDRR.[K] | 1xHomocysteineMMTS [K4] | 0,000599431 | 1543,66279 | Liver/ No FAIMS/MMTS |
| P38647 | P38647 [564-573] | [K].NAEKYAEEDR.[R] | 1xHomocysteineMMTS [K4] | 0,00347466 | 1387,56168 | Liver/ No FAIMS/MMTS |
| P38647 | P38647 [349-361] | [R].AQFEGIVTDLIKR.[T] | 1xHomocysteineMMTS [K12] | 0,000599431 | 1652,84986 | Liver/ No FAIMS/MMTS |
| P38647 | P38647 [660-678] | [R].EGSGSSGTGEQKEDQKEEK.[Q] | 1xHomocysteineMMTS [K12] | 0,000689517 | 2172,90197 | Liver/ No FAIMS/MMTS |
| P38647 | P38647 [392-405] | [R].MPKVQQTVQDLFGR.[A] | 1xOxidation [M1]; 1xHomocysteineMMTS [K3] | 4,49774E-05 | 1825,87576 | Liver/ FAIMS/MMTS |
| P38647 | P38647 [596-610] | [K].MEEFKDQLPADECNK.[L] | 1xMethylthio [C13]; 1xHomocysteineMMTS [K5] | 0,000282457 | 2005,78324 | Liver/ FAIMS/MMTS |
| P38647 | P38647 [203-218] | [R].QATKDAGQISGLNVLR.[V] | 1xHomocysteineMMTS [K4] | 0,000599431 | 1833,93096 | Liver/ FAIMS/MMTS |
| P38647 | P38647 [564-574] | [K].NAEKYAEEDRR.[K] | 1xHomocysteineMMTS [K4] | 0,000599431 | 1543,66279 | Liver/ FAIMS/MMTS |
| P38647 | P38647 [392-405] | [R].MPKVQQTVQDLFGR.[A] | 1xHomocysteineMMTS [K3] | 4,49774E-05 | 1809,88084 | Liver/ FAIMS/MMTS |
| P38647 | P38647 [127-138] | [R].RYDDPEVQKDTK.[N] | 1xHomocysteineMMTS [K9] | 0,00340109 | 1656,73562 | Liver/ FAIMS/MMTS |
| P38647 | P38647 [556-567] | [K].DDIENMVKNAEK.[Y] | 1xHomocysteineMMTS [K8] | 0,0004709 | 1568,67533 | Liver/ FAIMS/MMTS |
| P38647 | P38647 [620-634] | [R].ALLAGKDSETGENIR.[Q] | 1xHomocysteineMMTS [K6] | 4,49774E-05 | 1736,83058 | Liver/ FAIMS/MMTS |
| P38647 | P38647 [564-573] | [K].NAEKYAEEDR.[R] | 1xHomocysteineMMTS [K4] | 0,00347466 | 1387,56168 | Liver/ FAIMS/MMTS |
| P38647 | P38647 [174-187] | [K].MKETAENYLGHTAK.[N] | 1xHomocysteineMMTS [K2] | 4,49774E-05 | 1755,78627 | Liver/ FAIMS/MMTS |
| P38647 | P38647 [219-238] | [R].VINEPTAAALAYGLDKSEDK.[V] | 1xHomocysteineMMTS [K16] | 4,49774E-05 | 2268,08864 | Liver/ FAIMS/MMTS |
| P38647 | P38647 [108-122] | [R].QAVTNPNNTFYATKR.[L] | 1xHomocysteineMMTS [K14] | 4,49774E-05 | 1887,88401 | Liver/ FAIMS/MMTS |
| P38647 | P38647 [635-653] | [R].QAASSLQQASLKLFEMAYK.[K] | 1xHomocysteineMMTS [K12] | 0,00312202 | 2277,10761 | Liver/ FAIMS/MMTS |
| P38647 | P38647 [468-485] | [K].KSQVFSTAADGQTQVEIK.[V] | 1xHomocysteineMMTS [K1] | 4,49774E-05 | 2100,01 | Liver/ FAIMS/MMTS |
| P38647 | P38647 [293-307] | [R].ETGVDLTKDNMALQR.[V] | 1xHomocysteineMMTS [K8] | 4,49774E-05 | 1853,85541 | Brain/ FAIMS/MMTS |
| P39053 | P39053 [257-266] | [R].KFFLSHPSYR.[H] | 1xHomocysteineIAA [K1] | 4,32935E-05 | 1455,72017 | Brain/ FAIMS/IAA |
| P39053 | P39053 [563-573] | [K].YMLSVDNLKLR.[D] | 1xHomocysteineMMTS [K9] | 0,000507453 | 1514,75279 | Brain/ FAIMS/MMTS |
| P39053 | P39053 [189-199] | [K].IAKEVDPQGQR.[T] | 1xHomocysteineMMTS [K3] | 0,000599431 | 1403,67698 | Brain/ FAIMS/MMTS |
| P39053 | P39053 [298-309] | [R].NKLQSQLLSIEK.[E] | 1xHomocysteineMMTS [K2] | 0,000211896 | 1563,82331 | Brain/ FAIMS/MMTS |
| P39053; P39054 | P39053 [679-694]; P39054 [673-688] | [R].DLMPKTIMHLMINNTK.[EA] | 1xHomocysteineIAA [K5] | 4,32935E-05 | 2074,0316 | Brain/ FAIMS/IAA |
| P39053; P39054; Q8BZ98 | P39053 [200-217]; P39054 [200-217]; Q8BZ98 [200-217] | [R].TIGVITKLDLMDEGTDAR.[D] | 1xHomocysteineIAA [K7] | 4,32935E-05 | 2122,05187 | Brain/ FAIMS/IAA |

| | | | | | | |
|---|---|---|---|---|---|---|
| P39053; P39054; Q8BZ98 | P39053 [200-217]; P39054 [200-217]; Q8BZ98 [200-217] | [R].TIGVITKLDLMDEGTDAR.[D] | 1xHomocysteineMMTS [K7] | 4,49774E-05 | 2111,01812 | Brain/ FAIMS/MMTS |
| P39053; Q8BZ98 | P39053 [387-399]; Q8BZ98 [387-399] | [R].EISYAIKNIHGIR.[T] | 1xHomocysteineIAA [K7] | 6,30912E-05 | 1687,89484 | Brain/ FAIMS/IAA |
| P39054; Q8BZ98 | P39054 [257-266]; Q8BZ98 [257-266] | [R].KFFLSHPAYR.[H] | 1xHomocysteineIAA [K1] | 0,000595249 | 1439,72525 | Brain/ FAIMS/IAA |
| P40124 | P40124 [71-83] | [K].HAEMVHTGLKLER.[A] | 1xHomocysteineIAA [K10] | 4,32935E-05 | 1694,84651 | Brain/ FAIMS/IAA |
| P40142 | P40142 [439-465] | [R].SVPMSTVFYPSDGVATEKAVELAANTK.[G] | 1xHomocysteineIAA [K18] | 4,32935E-05 | 2986,45363 | Brain/ FAIMS/IAA |
| P40142 | P40142 [531-542] | [R].VLDPFTIKPLDR.[K] | 1xHomocysteineMMTS [K8] | 0,000250158 | 1576,82258 | Liver/ No FAIMS/MMTS |
| P40142 | P40142 [531-542] | [R].VLDPFTIKPLDR.[K] | 1xHomocysteineMMTS [K8] | 0,000250158 | 1576,82258 | Liver/ FAIMS/MMTS |
| P40142 | P40142 [531-542] | [R].VLDPFTIKPLDR.[K] | 1xHomocysteineMMTS [K8] | 0,000250158 | 1576,82258 | Brain/ FAIMS/MMTS |
| P40142 | P40142 [604-617] | [K].MFGIDKDAIVQAVK.[G] | 1xHomocysteineMMTS [K6] | 4,49774E-05 | 1697,84233 | Brain/ FAIMS/MMTS |
| P40237 | P40237 [179-189] | [K].IKEEDNQLIVK.[K] | 1xHomocysteineIAA [K2] | 0,00251002 | 1502,7883 | Brain/ FAIMS/IAA |
| P40240 | P40240 [118-129] | [K].ELQEFYKDTYQK.[L] | 1xHomocysteineIAA [K7] | 4,32935E-05 | 1765,81016 | Brain/ FAIMS/IAA |
| P40336 | P40336 [286-296] | [R].YFKQQEIILWR.[K] | 1xHomocysteineIAA [K3] | 0,000228034 | 1697,88321 | Brain/ FAIMS/IAA |
| P40936 | P40936 [5-19] | [K].VYIGGEDYEKEFTPK.[D] | 1xHomocysteineIAA [K10] | 4,32935E-05 | 1948,89971 | Liver/ FAIMS/IAA |
| P40936 | P40936 [5-19] | [K].VYIGGEDYEKEFTPK.[D] | 1xHomocysteineMMTS [K10] | 4,49774E-05 | 1937,86596 | Liver/ FAIMS/MMTS |
| P40936 | P40936 [212-224] | [K].FSGVYLEKEVVEK.[A] | 1xHomocysteineMMTS [K8] | 0,000266738 | 1689,82264 | Liver/ FAIMS/MMTS |
| P40936 | P40936 [81-100] | [R].EIIVTDYTPQNLQELQKWLK.[K] | 1xHomocysteineMMTS [K17] | 0,00074635 | 2622,33061 | Liver/ FAIMS/MMTS |
| P40936 | P40936 [1-14] | [-].MEGKVYIGGEDYEK.[E] | 1xAcetyl [N-Term]; 1xHomocysteineMMTS [K4] | 4,49774E-05 | 1822,76962 | Liver/ FAIMS/MMTS |
| P41105 | P41105 [23-35] | [K].QTYSTEPNNLKAR.[N] | 1xHomocysteineIAA [K11] | 0,00554312 | 1695,81189 | Brain/ FAIMS/IAA |
| P42125 | P42125 [207-224] | [K].VGVVDEVVPEDQVHSKAR.[S] | 1xHomocysteineMMTS [K16] | 4,49774E-05 | 2126,03688 | Liver/ FAIMS/MMTS |
| P42669 | P42669 [230-244] | [R].FFFDVGSNKYGVFMR.[V] | 1xHomocysteineIAA [K9] | 4,32935E-05 | 1987,91934 | Brain/ FAIMS/IAA |
| P42669 | P42669 [84-96] | [R].FLKIAEVGAGGNK.[S] | 1xHomocysteineMMTS [K3] | 0,00074635 | 1466,74942 | Brain/ FAIMS/MMTS |
| P42932 | P42932 [32-44] | [R].NIQACKELAQTTR.[T] | 1xCarbamidomethyl [C5]; 1xHomocysteineIAA [K6] | 4,32935E-05 | 1706,83125 | Brain/ FAIMS/IAA |
| P43006 | P43006 [158-173] | [K].KNDEVSSLDAFLDLIR.[N] | 1xHomocysteineIAA [K1] | 4,32935E-05 | 2009,00082 | Brain/ FAIMS/IAA |
| P43006 | P43006 [478-508] | [R].TSVNVVGDSFGAGIVYHLSKSELDTIDSQHR.[M] | 1xHomocysteineMMTS [K20] | 4,49774E-05 | 3494,66788 | Brain/ No FAIMS/MMTS |
| P43006 | P43006 [18-31] | [R].MHDSHLSSDEPKHR.[N] | 1xHomocysteineMMTS [K12] | 4,49774E-05 | 1838,77308 | Brain/ No FAIMS/MMTS |
| P43006 | P43006 [1-17] | [-].MASTEGANNMPKQVEVR.[M] | 1xMet-loss+Acetyl [N-Term]; 1xHomocysteineMMTS [K12] | 4,49774E-05 | 1935,87213 | Brain/ FAIMS/MMTS |
| P43006 | P43006 [478-508] | [R].TSVNVVGDSFGAGIVYHLSKSELDTIDSQHR.[M] | 1xHomocysteineMMTS [K20] | 4,49774E-05 | 3494,66788 | Brain/ FAIMS/MMTS |
| P43006 | P43006 [18-31] | [R].MHDSHLSSDEPKHR.[N] | 1xHomocysteineMMTS [K12] | 4,49774E-05 | 1838,77308 | Brain/ FAIMS/MMTS |
| P43024 | P43024 [83-111] | [R].TKPFPWGDGNHTLFHNPHVNPLPTGYEDE.[-] | 1xHomocysteineIAA [K2] | 4,32935E-05 | 3490,59095 | Brain/ No FAIMS/IAA |
| P43024 | P43024 [83-111] | [R].TKPFPWGDGNHTLFHNPHVNPLPTGYEDE.[-] | 1xHomocysteineIAA [K2] | 4,32935E-05 | 3490,59095 | Brain/ FAIMS/IAA |
| P43274 | P43274 [35-52] | [K].TSGPPVSELITKAVAASK.[E] | 1xHomocysteineIAA [K12] | 4,32935E-05 | 1930,03139 | Brain/ No FAIMS/IAA |
| P43274 | P43274 [35-52] | [K].TSGPPVSELITKAVAASK.[E] | 1xHomocysteineIAA [K12] | 4,32935E-05 | 1930,03139 | Brain/ FAIMS/IAA |
| P43274 | P43274 [35-52] | [K].TSGPPVSELITKAVAASK.[E] | 1xHomocysteineMMTS [K12] | 0,00394153 | 1918,99765 | Liver/ FAIMS/MMTS |
| P43274; P15864; P43277 | P43274 [64-75]; P15864 [64-75]; P43277 [65-76] | [K].KALAAAGYDVEK.[N] | 1xHomocysteineIAA [K1] | 4,32935E-05 | 1409,70933 | Brain/ No FAIMS/IAA |
| P43274; P15864; P43277 | P43274 [64-75]; P15864 [64-75]; P43277 [65-76] | [K].KALAAAGYDVEK.[N] | 1xHomocysteineIAA [K1] | 4,32935E-05 | 1409,70933 | Brain/ FAIMS/IAA |
| P43274; P43275; P15864; P43277 | P43274 [55-64]; P43275 [57-66]; P15864 [55-64]; P43277 [56-65] | [R].SGVSLAALKK.[AS] | 1xHomocysteineIAA [K] | 6,30912E-05 | 1147,65035 | Brain/ FAIMS/IAA |
| P43274; P43276; P15864; P43277 | P43274 [82-90]; P43276 [82-90]; P15864 [82-90]; P43277 [83-91] | [K].LGLKSLVSK.[G] | 1xHomocysteineIAA [K4] | 0,000318237 | 1118,66019 | Brain/ FAIMS/IAA |
| P43274; P43276; P43277 | P43274 [91-106]; P43276 [91-106]; P43277 [92-107] | [K].GTLVQTKGTGASGSFK.[L] | 1xHomocysteineIAA [K7] | 4,32935E-05 | 1712,86359 | Brain/ FAIMS/IAA |
| P45376 | P45376 [257-269] | [R].NLVVIPKSVTPVR.[I] | 1xHomocysteineIAA [K7] | 0,00012645 | 1595,93016 | Brain/ FAIMS/IAA |
| P45591 | P45591 [1-19] | [-].MASGVTVNDEVIKVFNDMK.[V] | 1xMet-loss [N-Term]; 1xHomocysteineMMTS [K13] | 0,00918178 | 2129,00756 | Brain/ FAIMS/MMTS |
| P45591; P18760 | P45591 [82-95]; P18760 [82-95] | [R].YALYDATYETKESK.[K] | 1xHomocysteineIAA [K] | 4,32935E-05 | 1855,84186 | Liver/ FAIMS/IAA |
| P45591; P18760 | P45591 [82-95]; P18760 [82-95] | [R].YALYDATYETKESK.[K] | 1xHomocysteineIAA [K] | 4,32935E-05 | 1855,84186 | Brain/ No FAIMS/IAA |
| P45591; P18760 | P45591 [82-95]; P18760 [82-95] | [R].YALYDATYETKESK.[K] | 1xHomocysteineIAA [K] | 4,32935E-05 | 1855,84186 | Brain/ FAIMS/IAA |
| P45591; P18760 | P45591 [14-21]; P18760 [14-21] | [K].VFNDMKVR.[K] | 1xHomocysteineMMTS [K6] | 0,00489249 | 1171,54207 | Liver/ No FAIMS/MMTS |
| P45591; P18760 | P45591 [82-95]; P18760 [82-95] | [R].YALYDATYETKESK.[K] | 1xHomocysteineMMTS [K11] | 4,49774E-05 | 1844,80811 | Liver/ FAIMS/MMTS |
| P45591; P18760 | P45591 [82-95]; P18760 [82-95] | [R].YALYDATYETKESK.[K] | 1xHomocysteineMMTS [K11] | 4,49774E-05 | 1844,80811 | Brain/ No FAIMS/MMTS |
| P45591; P18760 | P45591 [14-21]; P18760 [14-21] | [K].VFNDMKVR.[K] | 1xHomocysteineMMTS [K6] | 0,00489249 | 1171,54207 | Brain/ FAIMS/MMTS |
| P45591; P18760 | P45591 [82-95]; P18760 [82-95] | [R].YALYDATYETKESK.[K] | 1xHomocysteineMMTS [K11] | 4,49774E-05 | 1844,80811 | Brain/ FAIMS/MMTS |
| P45878 | P45878 [101-113] | [R].KLVIPSELGYGER.[G] | 1xHomocysteineIAA [K1] | 9,61721E-05 | 1634,85705 | Brain/ FAIMS/IAA |

| | | | | | | |
|---|---|---|---|---|---|---|
| P45952 | P45952 [257-271] | [R].VPKENVLIGEGAGFK.[I] | 1xHomocysteineMMTS [K3] | 4,49774E-05 | 1720,87607 | Liver/ FAIMS/MMTS |
| P45952 | P45952 [58-79] | [R].EEIIPVAPEYDKSGEYPFPLIK.[R] | 1xHomocysteineMMTS [K12] | 0,00324404 | 2697,31904 | Liver/ FAIMS/MMTS |
| P45952 | P45952 [70-80] | [K].SGEYPFPLIKR.[A] | 1xHomocysteineMMTS [K10] | 0,000706375 | 1469,72795 | Liver/ FAIMS/MMTS |
| P46096 | P46096 [399-421] | [R].RPIAQWHTLQVEEEVDAMLAVKK.[-] | 1xHomocysteineIAA [K22] | 4,32935E-05 | 2865,47498 | Brain/ No FAIMS/IAA |
| P46096 | P46096 [399-421] | [R].RPIAQWHTLQVEEEVDAMLAVKK.[-] | 1xHomocysteineIAA [K22] | 4,32935E-05 | 2865,47498 | Brain/ FAIMS/IAA |
| P46096 | P46096 [289-300] | [K].LTVVILEAKNLK.[K] | 1xHomocysteineMMTS [K9] | 4,49774E-05 | 1503,86372 | Brain/ No FAIMS/MMTS |
| P46096 | P46096 [200-213] | [R].KTLNPVFNEQFTFK.[V] | 1xHomocysteineMMTS [K1] | 6,70238E-05 | 1875,91319 | Brain/ No FAIMS/MMTS |
| P46096 | P46096 [289-300] | [K].LTVVILEAKNLK.[K] | 1xHomocysteineMMTS [K9] | 4,49774E-05 | 1503,86372 | Brain/ FAIMS/MMTS |
| P46096 | P46096 [261-272] | [R].DLQSAEKEEQEK.[L] | 1xHomocysteineMMTS [K7] | 4,49774E-05 | 1596,688 | Brain/ FAIMS/MMTS |
| P46096 | P46096 [370-388] | [K].NDAIGKVFVGYNSTGAELR.[H] | 1xHomocysteineMMTS [K6] | 4,49774E-05 | 2174,03688 | Brain/ FAIMS/MMTS |
| P46096 | P46096 [234-244] | [R].FSKHDIIGEFK.[V] | 1xHomocysteineMMTS [K3] | 0,00074635 | 1483,70722 | Brain/ FAIMS/MMTS |
| P46096 | P46096 [367-375] | [K].IGKNDAIGK.[V] | 1xHomocysteineMMTS [K3] | 0,00207528 | 1078,53836 | Brain/ FAIMS/MMTS |
| P46096 | P46096 [200-213] | [R].KTLNPVFNEQFTFK.[V] | 1xHomocysteineMMTS [K1] | 6,70238E-05 | 1875,91319 | Brain/ FAIMS/MMTS |
| P46096;<br>Q9R0N5 | P46096 [268-281];<br>Q9R0N5 [235-248] | [K].EEQEKLGDICFSLR.[Y] | 1xCarbamidomethyl [C10];<br>1xHomocysteineIAA [K5] | 0,000258769 | 1897,87826 | Brain/ FAIMS/IAA |
| P46097 | P46097 [235-245] | [R].FSKHDIIGEVK.[V] | 1xHomocysteineIAA [K3] | 4,32935E-05 | 1446,74096 | Brain/ FAIMS/IAA |
| P46097;<br>P46096 | P46097 [184-191];<br>P46096 [183-190] | [K].VFLLPDKK.[K] | 1xHomocysteineIAA [K] | 0,00130032 | 1133,63873 | Brain/ No FAIMS/IAA |
| P46097;<br>P46096 | P46097 [184-191];<br>P46096 [183-190] | [K].VFLLPDKK.[K] | 1xHomocysteineIAA [K] | 0,00130032 | 1133,63873 | Brain/ FAIMS/IAA |
| P46097;<br>P46096;<br>Q9R0N5 | P46097 [302-314];<br>P46096 [301-313];<br>Q9R0N5 [268-280] | [K].KMDVGGLSDPYVK.[IV] | 1xHomocysteineIAA [K1] | 4,32935E-05 | 1582,76038 | Brain/ No FAIMS/IAA |
| P46097;<br>P46096;<br>Q9R0N5 | P46097 [302-314];<br>P46096 [301-313];<br>Q9R0N5 [268-280] | [K].KMDVGGLSDPYVK.[IV] | 1xHomocysteineIAA [K1] | 4,32935E-05 | 1582,76038 | Brain/ FAIMS/IAA |
| P46425;<br>P19157 | P46425 [76-85]; P19157<br>[76-85] | [R].SLGLYGKNQR.[E] | 1xHomocysteineIAA [K7] | 4,32935E-05 | 1309,66813 | Liver/ No FAIMS/IAA |
| P46425;<br>P19157 | P46425 [76-85]; P19157<br>[76-85] | [R].SLGLYGKNQR.[E] | 1xHomocysteineIAA [K7] | 4,32935E-05 | 1309,66813 | Liver/ FAIMS/IAA |
| P46425;<br>P19157 | P46425 [76-85]; P19157<br>[76-85] | [R].SLGLYGKNQR.[E] | 1xHomocysteineIAA [K7] | 4,32935E-05 | 1309,66813 | Brain/ No FAIMS/IAA |
| P46425;<br>P19157 | P46425 [76-85]; P19157<br>[76-85] | [R].SLGLYGKNQR.[E] | 1xHomocysteineIAA [K7] | 4,32935E-05 | 1309,66813 | Brain/ FAIMS/IAA |
| P46425;<br>P19157 | P46425 [122-141];<br>P19157 [122-141] | [K].ALPGHLKPFETLLSQNQGGK.[A] | 1xHomocysteineMMTS [K7] | 4,49774E-05 | 2298,17332 | Liver/ No FAIMS/MMTS |
| P46425;<br>P19157 | P46425 [76-85]; P19157<br>[76-85] | [R].SLGLYGKNQR.[E] | 1xHomocysteineMMTS [K7] | 0,000152766 | 1298,63439 | Liver/ No FAIMS/MMTS |
| P46425;<br>P19157 | P46425 [117-141];<br>P19157 [117-141] | [K].NDYVKALPGHLKPFETLLSQNQGGK.[A] | 1xHomocysteineMMTS [K] | 4,49774E-05 | 2917,46989 | Liver/ No FAIMS/MMTS |
| P46425;<br>P19157 | P46425 [122-141];<br>P19157 [122-141] | [K].ALPGHLKPFETLLSQNQGGK.[A] | 1xHomocysteineMMTS [K7] | 4,49774E-05 | 2298,17332 | Liver/ FAIMS/MMTS |
| P46425;<br>P19157 | P46425 [76-85]; P19157<br>[76-85] | [R].SLGLYGKNQR.[E] | 1xHomocysteineMMTS [K7] | 0,000152766 | 1298,63439 | Liver/ FAIMS/MMTS |
| P46425;<br>P19157 | P46425 [117-141];<br>P19157 [117-141] | [K].NDYVKALPGHLKPFETLLSQNQGGK.[A] | 1xHomocysteineMMTS [K] | 4,49774E-05 | 2917,46989 | Liver/ FAIMS/MMTS |
| P46425;<br>P19157 | P46425 [76-85]; P19157<br>[76-85] | [R].SLGLYGKNQR.[E] | 1xHomocysteineMMTS [K7] | 0,000152766 | 1298,63439 | Brain/ FAIMS/MMTS |
| P46460 | P46460 [206-225] | [R].QSIINPDWNFEKMGIGGLDK.[E] | 1xHomocysteineIAA [K12] | 4,32935E-05 | 2436,16863 | Brain/ FAIMS/IAA |
| P46460 | P46460 [218-232] | [K].MGIGGLDKEFSDIFR.[R] | 1xHomocysteineMMTS [K8] | 0,000173718 | 1847,84887 | Brain/ No FAIMS/MMTS |
| P46460 | P46460 [218-232] | [K].MGIGGLDKEFSDIFR.[R] | 1xHomocysteineMMTS [K8] | 0,000173718 | 1847,84887 | Brain/ FAIMS/MMTS |
| P46460 | P46460 [517-533] | [R].VLDDGELLVQQTKNSDR.[T] | 1xHomocysteineMMTS [K13] | 0,000361246 | 2093,00016 | Brain/ FAIMS/MMTS |
| P46471 | P46471 [403-415] | [K].IATEKDFLEAVNK.[V] | 1xHomocysteineIAA [K5] | 0,00250017 | 1651,83598 | Brain/ FAIMS/IAA |
| P46638 | P46638 [14-30] | [K].VVLIGDSGVGKSNLLSR.[F] | 1xHomocysteineIAA [K11] | 4,32935E-05 | 1888,03206 | Brain/ FAIMS/IAA |
| P46660 | P46660 [448-463] | [K].KEEEEEEEASKEVSK.[K] | 1xHomocysteineIAA [K12] | 4,32935E-05 | 2082,90195 | Brain/ No FAIMS/IAA |
| P46660 | P46660 [448-463] | [K].KEEEEEEEASKEVSK.[K] | 1xHomocysteineIAA [K12] | 4,32935E-05 | 2082,90195 | Brain/ FAIMS/IAA |
| P46660 | P46660 [216-228] | [K].KVESLLDELAFVR.[Q] | 1xHomocysteineMMTS [K1] | 4,49774E-05 | 1681,86517 | Brain/ No FAIMS/MMTS |
| P46660 | P46660 [105-120] | [R].FAVFIEKVHQLETQNR.[A] | 1xHomocysteineMMTS [K7] | 0,000282457 | 2122,05723 | Brain/ FAIMS/MMTS |
| P46660 | P46660 [92-104] | [R].TNEKEQLQGLNDR.[F] | 1xHomocysteineMMTS [K4] | 0,000530332 | 1707,77888 | Brain/ FAIMS/MMTS |
| P46660 | P46660 [216-228] | [K].KVESLLDELAFVR.[Q] | 1xHomocysteineMMTS [K1] | 4,49774E-05 | 1681,86517 | Brain/ FAIMS/MMTS |
| P47708 | P47708 [171-192] | [K].KTKPQQPAGEPATQEQPTPESR.[H] | 1xHomocysteineIAA [K3] | 4,32935E-05 | 2579,25184 | Brain/ FAIMS/IAA |
| P47738 | P47738 [158-174] | [K].YHGKTIPIDGDFFSYTR.[H] | 1xHomocysteineIAA [K4] | 4,32935E-05 | 2191,0277 | Liver/ FAIMS/IAA |
| P47738 | P47738 [158-174] | [K].YHGKTIPIDGDFFSYTR.[H] | 1xHomocysteineIAA [K4] | 4,32935E-05 | 2191,0277 | Brain/ FAIMS/IAA |
| P47738 | P47738 [358-371] | [R].TEQGPQVDETQFKK.[I] | 1xHomocysteineMMTS [K13] | 4,49774E-05 | 1797,8146 | Liver/ No FAIMS/MMTS |
| P47738 | P47738 [397-416] | [R].GYFIQPTVFGDVKDGMTIAK.[E] | 1xHomocysteineMMTS [K13] | 4,49774E-05 | 2350,12801 | Liver/ No FAIMS/MMTS |
| P47738 | P47738 [444-457] | [K].YGLAAAVFTKDLDK.[A] | 1xHomocysteineMMTS [K10] | 0,000706375 | 1674,82298 | Liver/ No FAIMS/MMTS |
| P47738 | P47738 [372-385] | [K].ILGYIKSGQQEGAK.[L] | 1xHomocysteineMMTS [K6] | 4,49774E-05 | 1654,82912 | Liver/ FAIMS/MMTS |
| P47738 | P47738 [158-174] | [K].YHGKTIPIDGDFFSYTR.[H] | 1xHomocysteineMMTS [K4] | 4,49774E-05 | 2179,99396 | Liver/ FAIMS/MMTS |
| P47738 | P47738 [358-371] | [R].TEQGPQVDETQFKK.[I] | 1xHomocysteineMMTS [K13] | 4,49774E-05 | 1797,8146 | Liver/ FAIMS/MMTS |
| P47738 | P47738 [397-416] | [R].GYFIQPTVFGDVKDGMTIAK.[E] | 1xOxidation [M16];<br>1xHomocysteineMMTS [K13] | 0,000152766 | 2366,12292 | Liver/ FAIMS/MMTS |
| P47738 | P47738 [439-453] | [R].ANDSKYGLAAAVFTK.[D] | 1xHomocysteineMMTS [K5] | 0,000530332 | 1718,82404 | Liver/ FAIMS/MMTS |
| P47738 | P47738 [260-283] | [K].VAFTGSTEVGHLIQVAAGSSNLKR.[V] | 1xHomocysteineMMTS [K23] | 4,49774E-05 | 2605,3225 | Liver/ FAIMS/MMTS |
| P47738 | P47738 [429-438] | [K].FKTIEEVVGR.[A] | 1xHomocysteineMMTS [K2] | 0,000266738 | 1340,6701 | Liver/ FAIMS/MMTS |
| P47738 | P47738 [495-513] | [R].ELGEYGLQAYTEVKTVTVK.[V] | 1xHomocysteineMMTS [K14] | 4,49774E-05 | 2291,12978 | Liver/ FAIMS/MMTS |
| P47738 | P47738 [397-416] | [R].GYFIQPTVFGDVKDGMTIAK.[E] | 1xHomocysteineMMTS [K13] | 4,49774E-05 | 2350,12801 | Liver/ FAIMS/MMTS |
| P47753;<br>P47754 | P47753 [274-281];<br>P47754 [274-281] | [K].ILSYKIGK.[E] | 1xHomocysteineIAA [K5] | 0,00012645 | 1095,62308 | Brain/ No FAIMS/IAA |
| P47753;<br>P47754 | P47753 [274-281];<br>P47754 [274-281] | [K].ILSYKIGK.[E] | 1xHomocysteineIAA [K5] | 0,00012645 | 1095,62308 | Brain/ FAIMS/IAA |
| P47757 | P47757 [95-108] | [R].KLEVEANNAFDQYR.[D] | 1xHomocysteineIAA [K1] | 4,32935E-05 | 1870,87522 | Brain/ No FAIMS/IAA |
| P47757 | P47757 [95-108] | [R].KLEVEANNAFDQYR.[D] | 1xHomocysteineIAA [K1] | 4,32935E-05 | 1870,87522 | Brain/ FAIMS/IAA |

| | | | | | | |
|---|---|---|---|---|---|---|
| P47857 | P47857 [656-673] | [R].KNVLGHMQQGGSPTPFDR.[N] | 1xHomocysteineIAA [K1] 1xOxidation [M9]; | 4,32935E-05 | 2143,01715 | Brain/ FAIMS/IAA |
| P47867 | P47867 [308-318] | [K].TLIDFVKMMVK.[Y] | 1xHomocysteineIAA [K7] | 0,00471647 | 1514,77794 | Brain/ FAIMS/IAA |
| P47911 | P47911 [235-254] | [R].HQEGEIFDTEKEKYEITEQR.[K] | 1xHomocysteineIAA [K] | 4,32935E-05 | 2683,23043 | Liver/ FAIMS/IAA |
| P47911 | P47911 [235-254] | [R].HQEGEIFDTEKEKYEITEQR.[K] | 1xHomocysteineIAA [K] | 4,32935E-05 | 2683,23043 | Brain/ FAIMS/IAA |
| P47911 | P47911 [260-270] | [K].AVDLQILPKIK.[A] | 1xHomocysteineMMTS [K9] | 0,00074635 | 1400,80039 | Liver/ FAIMS/MMTS |
| P47911 | P47911 [255-268] | [R].KADQKAVDLQILPK.[I] | 1xHomocysteineMMTS [K5] | 4,49774E-05 | 1729,93392 | Liver/ FAIMS/MMTS |
| P47911 | P47911 [197-208] | [R].THQKFVIATSTK.[V] | 1xHomocysteineMMTS [K4] | 0,000152766 | 1523,77088 | Liver/ FAIMS/MMTS |
| P47911 | P47911 [269-280] | [K].IKAVPQLQGYLR.[S] | 1xHomocysteineMMTS [K2] | 4,49774E-05 | 1548,8389 | Liver/ FAIMS/MMTS |
| P47911 | P47911 [235-254] | [R].HQEGEIFDTEKEKYEITEQR.[K] | 1xHomocysteineMMTS [K] | 4,49774E-05 | 2672,19669 | Liver/ FAIMS/MMTS |
| P47915 | P47915 [135-151] | [K].AQASAPAQAPKGAQAPK.[G] | 1xHomocysteineIAA [K11] | 4,32935E-05 | 1765,90138 | Liver/ No FAIMS/IAA |
| P47915 | P47915 [135-151] | [K].AQASAPAQAPKGAQAPK.[G] | 1xHomocysteineIAA [K11] | 4,32935E-05 | 1765,90138 | Liver/ FAIMS/IAA |
| P47915 | P47915 [146-158] | [K].GAQAPKGAQAPVK.[A] | 1xHomocysteineMMTS [K6] | 0,000946421 | 1385,7028 | Liver/ No FAIMS/MMTS |
| P47915 | P47915 [131-145] | [K].APAKAQASAPAQAPK.[G] | 1xHomocysteineMMTS [K4] | 4,49774E-05 | 1569,78759 | Liver/ No FAIMS/MMTS |
| P47915 | P47915 [135-151] | [K].AQASAPAQAPKGAQAPK.[G] | 1xHomocysteineMMTS [K11] | 0,000192903 | 1754,86763 | Liver/ No FAIMS/MMTS |
| P47962 | P47962 [153-164] | [R].TTTGNKVFGALK.[G] | 1xHomocysteineIAA [K6] | 0,00107087 | 1410,74096 | Brain/ FAIMS/IAA |
| P47962 | P47962 [197-209] | [R].KHIMGQNVADYMR.[Y] | 1xHomocysteineMMTS [K1] | 0,00213323 | 1725,76918 | Liver/ FAIMS/MMTS |
| P47962 | P47962 [180-196] | [R].FPGYDSESKEFNAEVHR.[K] | 1xHomocysteineMMTS [K9] | 0,000173718 | 2174,927 | Liver/ FAIMS/MMTS |
| P47963 | P47963 [168-177] | [R].VITEEEKNFK.[A] | 1xHomocysteineIAA [K7] | 9,61721E-05 | 1410,69334 | Brain/ FAIMS/IAA |
| P47963 | P47963 [168-177] | [R].VITEEEKNFK.[A] | 1xHomocysteineMMTS [K7] | 0,000361246 | 1399,6596 | Liver/ No FAIMS/MMTS |
| P47963 | P47963 [199-209] | [R].AKEAAEQDVEK.[K] | 1xHomocysteineMMTS [K2] | 0,0004709 | 1380,61338 | Liver/ No FAIMS/MMTS |
| P47963 | P47963 [104-116] | [R].NKSTESLQANVQR.[L] | 1xHomocysteineMMTS [K2] | 0,000530332 | 1637,7734 | Liver/ FAIMS/MMTS |
| P47963 | P47963 [199-209] | [R].AKEAAEQDVEK.[K] | 1xHomocysteineMMTS [K2] | 0,0004709 | 1380,61338 | Liver/ FAIMS/MMTS |
| P47963 | P47963 [168-177] | [R].VITEEEKNFK.[A] | 1xHomocysteineMMTS [K7] | 0,000361246 | 1399,6596 | Liver/ FAIMS/MMTS |
| P48024 | P48024 [42-56] | [R].KTLTTVQGIADDYDK.[K] | 1xHomocysteineMMTS [K1] | 0,000266738 | 1830,86121 | Liver/ FAIMS/MMTS |
| P48036 | P48036 [62-74] | [R].DLVDDLKSELTGK.[F] | 1xHomocysteineIAA [K7] | 4,32935E-05 | 1606,79926 | Liver/ FAIMS/IAA |
| P48036 | P48036 [49-61] | [R].QEIAQEFKTLFGR.[D] | 1xHomocysteineIAA [K8] | 4,32935E-05 | 1740,87377 | Brain/ No FAIMS/IAA |
| P48036 | P48036 [49-61] | [R].QEIAQEFKTLFGR.[D] | 1xHomocysteineIAA [K8] | 4,32935E-05 | 1740,87377 | Brain/ FAIMS/IAA |
| P48036 | P48036 [49-61] | [R].QEIAQEFKTLFGR.[D] | 1xHomocysteineMMTS [K8] | 0,000173718 | 1729,84002 | Liver/ No FAIMS/MMTS |
| P48036 | P48036 [100-115] | [K].GAGTDEKVLTEIIASR.[T] | 1xHomocysteineMMTS [K7] | 4,49774E-05 | 1822,90374 | Liver/ FAIMS/MMTS |
| P48036 | P48036 [88-99] | [R].LYDAYELKHALK.[G] | 1xHomocysteineMMTS [K8] | 6,70238E-05 | 1626,80185 | Liver/ FAIMS/MMTS |
| P48036 | P48036 [49-61] | [R].QEIAQEFKTLFGR.[D] | 1xHomocysteineMMTS [K8] | 0,000173718 | 1729,84002 | Liver/ FAIMS/MMTS |
| P48036 | P48036 [185-199] | [K].WGTDEEKFITIFGTR.[S] | 1xHomocysteineMMTS [K7] | 0,000266738 | 1962,90883 | Liver/ FAIMS/MMTS |
| P48036 | P48036 [25-43] | [K].AMKGLGTDEDSILNLLTSR.[S] | 1xHomocysteineMMTS [K3] | 0,00230389 | 2197,06614 | Liver/ FAIMS/MMTS |
| P48036 | P48036 [49-61] | [R].QEIAQEFKTLFGR.[D] | 1xHomocysteineMMTS [K8] | 0,000173718 | 1729,84002 | Brain/ No FAIMS/MMTS |
| P48036 | P48036 [185-199] | [K].WGTDEEKFITIFGTR.[S] | 1xHomocysteineMMTS [K7] | 0,000266738 | 1962,90883 | Brain/ FAIMS/MMTS |
| P48036 | P48036 [100-115] | [K].GAGTDEKVLTEIIASR.[T] | 1xHomocysteineMMTS [K7] | 4,49774E-05 | 1822,90374 | Brain/ FAIMS/MMTS |
| P48320 | P48320 [526-534] | [R].LSKVAPVIK.[A] | 1xHomocysteineIAA [K3] | 0,00927232 | 1128,68093 | Brain/ FAIMS/IAA |
| P48455; P63328; P48453 | P48455 [315-328]; P63328 [319-332]; P48453 [328-341] | [K].AAVLKYENNVMNIR.[Q] | 1xHomocysteineIAA [K5] | 4,32935E-05 | 1808,91458 | Brain/ FAIMS/IAA |
| P48678 | P48678 [134-144] | [R].LKDLEALLNSK.[E] | 1xHomocysteineIAA [K2] | 0,00554312 | 1417,77193 | Brain/ FAIMS/IAA |
| P48722 | P48722 [347-356] | [R].IPAVKEQVTR.[F] | 1xHomocysteineIAA [K5] | 4,32935E-05 | 1314,71983 | Brain/ FAIMS/IAA |
| P48771 | P48771 [34-56] | [K].LFQEDNGMPVHLKGGASDALLYR.[A] | 1xHomocysteineIAA [K13] | 4,32935E-05 | 2705,31742 | Brain/ No FAIMS/IAA |
| P48771 | P48771 [34-56] | [K].LFQEDNGMPVHLKGGASDALLYR.[A] | 1xHomocysteineIAA [K13] 1xOxidation [M8]; | 4,32935E-05 | 2705,31742 | Brain/ FAIMS/IAA |
| P48771 | P48771 [34-56] | [K].LFQEDNGMPVHLKGGASDALLYR.[A] | 1xHomocysteineMMTS [K13] | 4,49774E-05 | 2710,27859 | Liver/ FAIMS/MMTS |
| P48771 | P48771 [34-56] | [K].LFQEDNGMPVHLKGGASDALLYR.[A] | 1xHomocysteineMMTS [K13] 1xOxidation [M8]; | 4,49774E-05 | 2694,28367 | Liver/ FAIMS/MMTS |
| P48771 | P48771 [34-56] | [K].LFQEDNGMPVHLKGGASDALLYR.[A] | 1xHomocysteineMMTS [K13] | 4,49774E-05 | 2710,27859 | Brain/ FAIMS/MMTS |
| P48774 | P48774 [71-81] | [K].NKITQSNAILR.[Y] | 1xHomocysteineIAA [K2] | 9,61721E-05 | 1431,77366 | Brain/ FAIMS/IAA |
| P48962 | P48962 [34-49] | [K].LLLQVQHASKQISAEK.[Q] | 1xHomocysteineIAA [K10] | 4,32935E-05 | 1967,07426 | Brain/ No FAIMS/IAA |
| P48962 | P48962 [34-49] | [K].LLLQVQHASKQISAEK.[Q] | 1xHomocysteineIAA [K10] | 4,32935E-05 | 1967,07426 | Brain/ FAIMS/IAA |
| P48962 | P48962 [141-152] | [R].LAADVGKGSSQR.[E] | 1xHomocysteineMMTS [K7] | 4,49774E-05 | 1351,64568 | Brain/ No FAIMS/MMTS |
| P48962 | P48962 [264-280] | [K].DEGANAFFKGAWSNVLR.[G] | 1xHomocysteineMMTS [K9] | 0,000192903 | 2044,93678 | Brain/ FAIMS/MMTS |
| P48962 | P48962 [141-152] | [R].LAADVGKGSSQR.[E] | 1xHomocysteineMMTS [K7] | 4,49774E-05 | 1351,64568 | Brain/ FAIMS/MMTS |
| P48962 | P48962 [261-272] | [K].IAKDEGANAFFK.[G] | 1xHomocysteineMMTS [K3] | 0,00393061 | 1473,68648 | Brain/ FAIMS/MMTS |
| P48962 | P48962 [61-72] | [R].IPKEQGFLSFWR.[G] | 1xHomocysteineMMTS [K3] | 4,49774E-05 | 1670,81817 | Brain/ FAIMS/MMTS |
| P48962 | P48962 [34-49] | [K].LLLQVQHASKQISAEK.[Q] | 1xHomocysteineMMTS [K10] | 0,00107839 | 1956,04051 | Brain/ FAIMS/MMTS |
| P48962; P51881 | P48962 [32-43]; P51881 [32-43] | [R].VKLLLQVQHASK.[Q] | 1xHomocysteineIAA [K2] | 4,32935E-05 | 1537,88829 | Brain/ No FAIMS/IAA |
| P48962; P51881 | P48962 [32-43]; P51881 [32-43] | [R].VKLLLQVQHASK.[Q] | 1xHomocysteineIAA [K2] | 4,32935E-05 | 1537,88829 | Brain/ FAIMS/IAA |
| P48962; P51881 | P48962 [81-94]; P51881 [81-94] | [R].YFPTQALNFAFKDK.[Y] | 1xHomocysteineMMTS [K12] | 0,00466355 | 1852,87607 | Brain/ FAIMS/MMTS |
| P49312 | P49312 [106-122] | [K].KIFVGGIKEDTEEHHLR.[D] | 1xHomocysteineIAA [K1] | 4,32935E-05 | 2182,10735 | Brain/ FAIMS/IAA |
| P49429 | P49429 [133-155] | [K].FAVLQTYGDTTHTLVEKINYTGR.[F] | 1xHomocysteineMMTS [K17] | 4,49774E-05 | 2790,35895 | Liver/ FAIMS/MMTS |
| P49429 | P49429 [285-300] | [R].GTEFLAAPSSYYKLLR.[E] | 1xHomocysteineMMTS [K13] | 0,000152766 | 1978,97652 | Liver/ FAIMS/MMTS |
| P49429 | P49429 [156-172] | [R].FLPGFEAPTYKDTLLPK.[L] | 1xHomocysteineMMTS [K11] | 0,000829616 | 2100,05443 | Liver/ FAIMS/MMTS |
| P50114 | P50114 [35-56] | [K].ELINNELSHFLEEIKEQEVVDK.[V] | 1xHomocysteineIAA [K15] | 4,32935E-05 | 2829,3975 | Brain/ No FAIMS/IAA |
| P50114 | P50114 [35-56] | [K].ELINNELSHFLEEIKEQEVVDK.[V] | 1xHomocysteineIAA [K15] | 4,32935E-05 | 2829,3975 | Brain/ FAIMS/IAA |
| P50172 | P50172 [37-48] | [K].VIVTGASKGIGR.[E] | 1xHomocysteineIAA [K8] | 4,32935E-05 | 1331,74638 | Liver/ FAIMS/IAA |
| P50172 | P50172 [37-48] | [K].VIVTGASKGIGR.[E] | 1xHomocysteineMMTS [K8] | 4,49774E-05 | 1320,71264 | Liver/ FAIMS/MMTS |
| P50172 | P50172 [67-77] | [R].SEEGLQKVVSR.[C] | 1xHomocysteineMMTS [K7] | 4,49774E-05 | 1394,67665 | Liver/ FAIMS/MMTS |
| P50172 | P50172 [254-273] | [K].SEVYYDKSPLTPILLGNPGR.[K] | 1xHomocysteineMMTS [K7] | 4,49774E-05 | 2382,18321 | Liver/ FAIMS/MMTS |
| P50172 | P50172 [36-44] | [K].KVIVTGASK.[G] | 1xHomocysteineMMTS [K1] | 0,000282457 | 1065,5795 | Liver/ FAIMS/MMTS |
| P50172 | P50172 [274-282] | [R].KIMEFFSLR.[Y] | 1xHomocysteineMMTS [K1] | 0,00107839 | 1333,64653 | Liver/ FAIMS/MMTS |
| P50247 | P50247 [20-34] | [R].KALDIAENEMPGLMR.[M] | 1xHomocysteineMMTS [K1] 1xCarbamidomethyl [C7]; | 0,000282457 | 1850,86314 | Liver/ FAIMS/MMTS |
| P50396 | P50396 [329-348] | [R].KSDIYVCMISYAHNVAAQGK.[Y] | 1xHomocysteineIAA [K1] | 4,32935E-05 | 2429,14103 | Brain/ FAIMS/IAA |
| P50396 | P50396 [36-55] | [R].NPYYGGESSSITPLEELYKR.[F] | 1xHomocysteineMMTS [K19] | 4,49774E-05 | 2466,13157 | Brain/ FAIMS/MMTS |
| P50396 | P50396 [424-436] | [R].MAGSAFDFENMKR.[K] | 1xHomocysteineMMTS [K12] | 0,00427174 | 1666,68445 | Brain/ FAIMS/MMTS |

| | | | | | | |
|---|---|---|---|---|---|---|
| P50396; Q61598 | P50396 [219-240]; Q61598 [219-240] | [R].YGKSPYLYPLYGLGELPQGFAR.[L] | 1xHomocysteineIAA [K3] | 4,32935E-05 | 2663,33266 | Brain/ No FAIMS/IAA |
| P50396; Q61598 | P50396 [219-240]; Q61598 [219-240] | [R].YGKSPYLYPLYGLGELPQGFAR.[L] | 1xHomocysteineIAA [K3] | 4,32935E-05 | 2663,33266 | Brain/ FAIMS/IAA |
| P50396; Q61598 | P50396 [209-218]; Q61598 [209-218] | [R].IKLYSESLAR.[Y] | 1xHomocysteineMMTS [K2] | 0,00359453 | 1342,68575 | Brain/ FAIMS/MMTS |
| P50431 | P50431 [446-460] | [K].LAGDEKIQSAVATLR.[E] | 1xHomocysteineMMTS [K6] | 0,000361246 | 1734,8877 | Liver/ FAIMS/MMTS |
| P50516 | P50516 [443-456] | [R].KHFPSVNWLISYSK.[Y] | 1xHomocysteineIAA [K1] | 4,32935E-05 | 1879,95235 | Brain/ FAIMS/IAA |
| P50518 | P50518 [14-33] | [K].HMMAFIEQEANEKAEEIDAK.[A] | 1xHomocysteineIAA [K13] | 4,32935E-05 | 2508,12036 | Brain/ FAIMS/IAA |
| P50571; P63080; P63137 | P50571 [129-139]; P63080 [129-139]; P63137 [128-138] | [K].SFVHGVTVKNR.[M] | 1xHomocysteineIAA [K9] | 0,00596151 | 1417,73688 | Brain/ FAIMS/IAA |
| P50580 | P50580 [156-172] | [R].LVKPGNQNTQVTEAWNK.[V] | 1xHomocysteineMMTS [K3] | 0,000781536 | 2090,01575 | Liver/ FAIMS/MMTS |
| P51150 | P51150 [114-132] | [R].DPENFPFVVLGNKIDLENR.[Q] | 1xHomocysteineIAA [K13] | 4,32935E-05 | 2390,18091 | Brain/ No FAIMS/IAA |
| P51150 | P51150 [114-132] | [R].DPENFPFVVLGNKIDLENR.[Q] | 1xHomocysteineIAA [K13] | 4,32935E-05 | 2390,18091 | Brain/ FAIMS/IAA |
| P51410 | P51410 [72-89] | [R].TICSHVQNMIKGVTLGFR.[Y] | 1xCarbamidomethyl [C3]; 1xHomocysteineIAA [K11] | 0,00151111 | 2235,11951 | Brain/ FAIMS/IAA |
| P51859 | P51859 [80-96] | [R].KGFSEGLWEIENNPTVK.[A] | 1xHomocysteineMMTS [K1] | 0,000211896 | 2110,99362 | Liver/ FAIMS/MMTS |
| P51863 | P51863 [266-288] | [K].NVADYYPEYKLLFEGAGSNPGDK.[T] | 1xHomocysteineIAA [K10] | 4,32935E-05 | 2721,25011 | Brain/ No FAIMS/IAA |
| P51881 | P51881 [97-106] | [K].QIFLGGVDKR.[T] | 1xHomocysteineIAA [K9] | 4,32935E-05 | 1306,69362 | Brain/ FAIMS/IAA |
| P51881 | P51881 [141-152] | [R].LAADVGKAGAER.[E] | 1xHomocysteineMMTS [K7] | 4,49774E-05 | 1320,63987 | Brain/ FAIMS/MMTS |
| P51910 | P51910 [46-58] | [R].WYEIEKIPASFEK.[G] | 1xHomocysteineIAA [K6] | 0,000210768 | 1813,88293 | Brain/ FAIMS/IAA |
| P52196 | P52196 [164-176] | [K].TYEQVLENLQSKR.[F] | 1xHomocysteineIAA [K12] | 0,00918796 | 1781,88506 | Brain/ FAIMS/IAA |
| P52196 | P52196 [164-176] | [K].TYEQVLENLQSKR.[F] | 1xHomocysteineMMTS [K12] | 0,000724331 | 1770,85132 | Liver/ FAIMS/MMTS |
| P52480 | P52480 [207-224] | [K].KGVNLPGAAVDLPAVSEK.[D] | 1xHomocysteineIAA [K1] | 4,32935E-05 | 1939,03172 | Brain/ No FAIMS/IAA |
| P52480 | P52480 [207-224] | [K].KGVNLPGAAVDLPAVSEK.[D] | 1xHomocysteineIAA [K1] | 4,32935E-05 | 1939,03172 | Brain/ FAIMS/IAA |
| P52480 | P52480 [57-66] | [R].SVEMLKEMIK.[S] | 1xHomocysteineMMTS [K6] | 0,000653201 | 1370,65505 | Brain/ No FAIMS/MMTS |
| P52480 | P52480 [267-278] | [K].IISKIENHEGVR.[R] | 1xHomocysteineMMTS [K4] | 6,70238E-05 | 1557,78759 | Brain/ No FAIMS/MMTS |
| P52480 | P52480 [247-255] | [R].KAADVHEVR.[K] | 1xHomocysteineMMTS [K1] | 4,49774E-05 | 1187,56597 | Brain/ No FAIMS/MMTS |
| P52480 | P52480 [57-66] | [R].SVEMLKEMIK.[S] | 1xHomocysteineMMTS [K6] | 0,000653201 | 1370,65505 | Brain/ FAIMS/MMTS |
| P52480 | P52480 [121-135] | [R].TGLIKGSGTAEVELK.[K] | 1xHomocysteineMMTS [K5] | 4,49774E-05 | 1665,855 | Brain/ FAIMS/MMTS |
| P52480 | P52480 [267-278] | [K].IISKIENHEGVR.[R] | 1xHomocysteineMMTS [K4] | 6,70238E-05 | 1557,78759 | Brain/ FAIMS/MMTS |
| P52480 | P52480 [93-120] | [R].EATESFASDPILYRPVAVALDTKGPEIR.[T] | 1xHomocysteineMMTS [K23] | 0,00074635 | 3208,6017 | Brain/ FAIMS/MMTS |
| P52480 | P52480 [207-224] | [K].KGVNLPGAAVDLPAVSEK.[D] | 1xHomocysteineMMTS [K1] | 0,000282457 | 1927,99798 | Brain/ FAIMS/MMTS |
| P52480; P53657 | P52480 [295-311]; P53657 [338-354] | [R].GDLGIEIPAEKVFLAQK.[M] | 1xHomocysteineIAA [K11] | 4,32935E-05 | 2002,06777 | Brain/ No FAIMS/IAA |
| P52480; P53657 | P52480 [295-311]; P53657 [338-354] | [R].GDLGIEIPAEKVFLAQK.[M] | 1xHomocysteineIAA [K11] | 4,32935E-05 | 2002,06777 | Brain/ FAIMS/IAA |
| P52480; P53657 | P52480 [295-311]; P53657 [338-354] | [R].GDLGIEIPAEKVFLAQK.[M] | 1xHomocysteineMMTS [K11] | 4,49774E-05 | 1991,03403 | Brain/ No FAIMS/MMTS |
| P52480; P53657 | P52480 [295-311]; P53657 [338-354] | [R].GDLGIEIPAEKVFLAQK.[M] | 1xHomocysteineMMTS [K11] | 4,49774E-05 | 1991,03403 | Brain/ FAIMS/MMTS |
| P52503 | P52503 [31-44] | [K].ITHTGQVYDEKDYR.[R] | 1xHomocysteineIAA [K11] | 9,61721E-05 | 1898,87014 | Brain/ FAIMS/IAA |
| P52760 | P52760 [61-78] | [K].NLGEILKAAGCDFNNVVK.[T] | 1xCarbamidomethyl [C11]; 1xHomocysteineIAA [K7] | 0,00124346 | 2136,05762 | Liver/ FAIMS/IAA |
| P52760 | P52760 [57-67] | [K].QALKNLGEILK.[A] | 1xHomocysteineMMTS [K4] | 0,00367611 | 1389,75925 | Liver/ No FAIMS/MMTS |
| P52760 | P52760 [30-60] | [R].TIYISGQVGLDPSSGQLVPGGVVEEAKQALK.[N] | 1xHomocysteineMMTS [K27] | 4,49774E-05 | 3303,69632 | Liver/ No FAIMS/MMTS |
| P52760 | P52760 [121-135] | [R].VEIEAIAVQGPFIKA.[-] | 1xHomocysteineMMTS [K14] | 4,49774E-05 | 1747,91212 | Liver/ No FAIMS/MMTS |
| P52760 | P52760 [8-29] | [K].VISTTKAPAAIGPYSQAVQVDR.[T] | 1xHomocysteineMMTS [K6] | 4,49774E-05 | 2435,24213 | Liver/ FAIMS/MMTS |
| P52760 | P52760 [57-67] | [K].QALKNLGEILK.[A] | 1xHomocysteineMMTS [K4] | 0,00367611 | 1389,75925 | Liver/ FAIMS/MMTS |
| P52760 | P52760 [121-135] | [R].VEIEAIAVQGPFIKA.[-] | 1xHomocysteineMMTS [K14] | 4,49774E-05 | 1747,91212 | Liver/ FAIMS/MMTS |
| P53395 | P53395 [188-199] | [K].LSEVVGSGKDGR.[I] | 1xHomocysteineIAA [K9] | 0,00544356 | 1377,67909 | Liver/ FAIMS/IAA |
| P53564 | P53564 [743-764] | [K].HLSASPMSTVSTYPPLAISLKK.[T] | 1xOxidation [M7]; 1xHomocysteineMMTS [K21] | 0,00927584 | 2507,27065 | Brain/ FAIMS/MMTS |
| P53810 | P53810 [172-186] | [R].GPLGPNWKQELVNQK.[D] | 1xHomocysteineIAA [K8] | 0,000114608 | 1881,96398 | Brain/ FAIMS/IAA |
| P53994 | P53994 [47-56] | [R].MITIDGKQIK.[L] | 1xHomocysteineIAA [K7] | 0,00115802 | 1320,7014 | Brain/ FAIMS/IAA |
| P53994; P59279 | P53994 [54-69]; P59279 [54-69] | [K].QIKLQIWDTAGQESFR.[S] | 1xHomocysteineIAA [K3] | 4,32935E-05 | 2094,04368 | Brain/ FAIMS/IAA |
| P54071 | P54071 [262-272] | [R].FKDIFQEIFDK.[H] | 1xHomocysteineIAA [K2] | 0,00049316 | 1603,78249 | Liver/ FAIMS/IAA |
| P54071 | P54071 [262-272] | [R].FKDIFQEIFDK.[H] | 1xHomocysteineIAA [K2] | 0,00049316 | 1603,78249 | Brain/ FAIMS/IAA |
| P54071 | P54071 [194-205] | [K].LVFTPKDGSSAK.[E] | 1xHomocysteineMMTS [K6] | 0,0018995 | 1412,69123 | Liver/ FAIMS/MMTS |
| P54071 | P54071 [252-261] | [K].NTILKAYDGR.[F] | 1xHomocysteineMMTS [K5] | 0,00140305 | 1313,63405 | Liver/ FAIMS/MMTS |
| P54071 | P54071 [70-89] | [K].LILPHVDVQLKYFDLGLPNR.[D] | 1xHomocysteineMMTS [K11] | 0,001242 | 2513,34072 | Liver/ FAIMS/MMTS |
| P54099 | P54099 [427-444] | [R].YLTEAQNTYEELQREMKK.[S] | 1xOxidation [M16]; 1xHomocysteineIAA [K17] | 0,00606366 | 2464,14829 | Liver/ No FAIMS/IAA |
| P54227 | P54227 [63-75] | [K].SHEAEVLKQLAEK.[R] | 1xHomocysteineIAA [K8] | 4,32935E-05 | 1655,84213 | Brain/ FAIMS/IAA |
| P54227 | P54227 [1-13] | [-].MASSDIQVKELEK.[R] | 1xMet-loss+Acetyl [N-Term]; 1xHomocysteineMMTS [K9] | 0,000152766 | 1551,73931 | Brain/ FAIMS/MMTS |
| P54227 | P54227 [63-75] | [K].SHEAEVLKQLAEK.[R] | 1xHomocysteineMMTS [K8] | 0,00869467 | 1644,80839 | Brain/ FAIMS/MMTS |
| P54227 | P54227 [113-122] | [R].EAQMAAKLER.[L] | 1xHomocysteineMMTS [K7] | 8,93346E-05 | 1309,60612 | Brain/ FAIMS/MMTS |
| P54227 | P54227 [96-104] | [K].MAEEKLTHK.[M] | 1xHomocysteineMMTS [K5] | 0,00820834 | 1249,57376 | Brain/ FAIMS/MMTS |
| P54227; P55821 | P54227 [42-52]; P55821 [76-86] | [K].KKDLSLEEIQK.[K] | 1xHomocysteineIAA [K2] | 0,000289025 | 1504,80395 | Brain/ FAIMS/IAA |
| P54728 | P54728 [46-60] | [K].LIYAGKILSDDTALK.[E] | 1xHomocysteineIAA [K6] | 4,32935E-05 | 1794,967 | Brain/ FAIMS/IAA |
| P54869 | P54869 [340-350] | [R].GLKLEETYTNK.[D] | 1xHomocysteineMMTS [K3] | 0,00222764 | 1458,69671 | Liver/ FAIMS/MMTS |
| P55050 | P55050 [48-57] | [K].FTVKESSNFR.[N] | 1xHomocysteineIAA [K4] | 0,00113008 | 1388,66271 | Liver/ FAIMS/IAA |
| P55088 | P55088 [285-310] | [R].SQVETEDLILKPGVVHVIDIDRGEEK.[K] | 1xHomocysteineIAA [K11] | 4,32935E-05 | 3092,59324 | Brain/ FAIMS/IAA |
| P55264 | P55264 [249-258] | [R].EQGFETKDIK.[E] | 1xHomocysteineIAA [K7] | 0,00012645 | 1368,64639 | Liver/ No FAIMS/IAA |
| P55264 | P55264 [249-258] | [R].EQGFETKDIK.[E] | 1xHomocysteineIAA [K7] | 0,00012645 | 1368,64639 | Liver/ FAIMS/IAA |

| | | | | | | |
|---|---|---|---|---|---|---|
| P55264 | P55264 [264-273] | [K].AQALPKVNSK.[R] | 1xHomocysteineMMTS [K6] | 0,000564742 | 1218,63332 | Liver/ No FAIMS/MMTS |
| P55264 | P55264 [63-71] | [K].ELFDELVKK.[F] | 1xHomocysteineMMTS [K8] | 0,00263388 | 1283,63741 | Liver/ FAIMS/MMTS |
| P55264 | P55264 [264-273] | [K].AQALPKVNSK.[R] | 1xHomocysteineMMTS [K6] | 0,000564742 | 1218,63332 | Liver/ FAIMS/MMTS |
| P55264 | P55264 [47-60] | [K].YSLKPNDQILAEDK.[H] | 1xHomocysteineMMTS [K4] | 0,000980684 | 1796,85573 | Liver/ FAIMS/MMTS |
| P55264 | P55264 [163-170] | [K].EKHLDLER.[N] | 1xHomocysteineMMTS [K2] | 0,00489249 | 1202,56564 | Liver/ FAIMS/MMTS |
| P55264 | P55264 [47-62] | [K].YSLKPNDQILAEDKHK.[E] | 1xHomocysteineMMTS [K14] | 0,0025135 | 2062,00961 | Liver/ FAIMS/MMTS |
| P56135 | P56135 [12-27] | [K].LMEVKLGELPSWIMMR.[D] | 1xHomocysteineIAA [K5] | 4,32935E-05 | 2107,05709 | Brain/ No FAIMS/IAA |
| P56135 | P56135 [12-27] | [K].LMEVKLGELPSWIMMR.[D] | 1xHomocysteineIAA [K5] | 4,32935E-05 | 2107,05709 | Brain/ FAIMS/IAA |
| P56375 | P56375 [49-65] | [K].NTSKGTVTGQVQGPEEK.[V] | 1xHomocysteineIAA [K4] | 0,00033298 | 1933,92838 | Brain/ FAIMS/IAA |
| P56375 | P56375 [93-105] | [K].TISKLEYSDFSIR.[Y] | 1xHomocysteineMMTS [K4] | 0,00369001 | 1721,8237 | Brain/ FAIMS/MMTS |
| P56376 | P56376 [46-60] | [R].GTVQGQLQGPVSKVR.[F] | 1xHomocysteineIAA [K13] | 6,30912E-05 | 1727,92211 | Brain/ FAIMS/IAA |
| P56382 | P56382 [15-24] | [R].FSQICAKAVR.[D] | 1xCarbamidomethyl [C5]; 1xHomocysteineIAA [K7] | 6,30912E-05 | 1353,67659 | Liver/ FAIMS/IAA |
| P56382 | P56382 [15-24] | [R].FSQICAKAVR.[D] | 1xCarbamidomethyl [C5]; 1xHomocysteineIAA [K7] | 6,30912E-05 | 1353,67659 | Brain/ No FAIMS/IAA |
| P56382 | P56382 [15-24] | [R].FSQICAKAVR.[D] | 1xCarbamidomethyl [C5]; 1xHomocysteineIAA [K7] | 6,30912E-05 | 1353,67659 | Brain/ FAIMS/IAA |
| P56391 | P56391 [60-76] | [R].VYKSLCPVSWVSAWDDR.[I] | 1xCarbamidomethyl [C6]; 1xHomocysteineIAA [K3] | 4,32935E-05 | 2242,04197 | Liver/ FAIMS/IAA |
| P56391 | P56391 [60-76] | [R].VYKSLCPVSWVSAWDDR.[I] | 1xCarbamidomethyl [C6]; 1xHomocysteineIAA [K3] | 4,32935E-05 | 2242,04197 | Brain/ No FAIMS/IAA |
| P56391 | P56391 [60-76] | [R].VYKSLCPVSWVSAWDDR.[I] | 1xCarbamidomethyl [C6]; 1xHomocysteineIAA [K3] | 4,32935E-05 | 2242,04197 | Brain/ FAIMS/IAA |
| P56391 | P56391 [1-8] | [-].MAEDIKTK.[I] | 1xMet-loss+Acetyl [N-Term]; 1xHomocysteineMMTS [K6] | 0,00074635 | 1009,46928 | Liver/ No FAIMS/MMTS |
| P56391 | P56391 [11-20] | [K].NYKTAPFDSR.[F] | 1xHomocysteineMMTS [K3] | 0,000530332 | 1361,59767 | Liver/ No FAIMS/MMTS |
| P56391 | P56391 [1-8] | [-].MAEDIKTK.[I] | 1xMet-loss+Acetyl [N-Term]; 1xHomocysteineMMTS [K6] | 0,00074635 | 1009,46928 | Liver/ FAIMS/MMTS |
| P56391 | P56391 [11-20] | [K].NYKTAPFDSR.[F] | 1xHomocysteineMMTS [K3] | 0,000530332 | 1361,59767 | Liver/ FAIMS/MMTS |
| P56391 | P56391 [1-8] | [-].MAEDIKTK.[I] | 1xMet-loss+Acetyl [N-Term]; 1xHomocysteineMMTS [K6] | 0,00074635 | 1009,46928 | Brain/ No FAIMS/MMTS |
| P56391 | P56391 [11-20] | [K].NYKTAPFDSR.[F] | 1xHomocysteineMMTS [K3] | 0,000530332 | 1361,59767 | Brain/ No FAIMS/MMTS |
| P56391 | P56391 [1-8] | [-].MAEDIKTK.[I] | 1xMet-loss+Acetyl [N-Term]; 1xHomocysteineMMTS [K6] | 0,00074635 | 1009,46928 | Brain/ FAIMS/MMTS |
| P56391 | P56391 [77-86] | [R].IAEGTFPGKI.[-] | 1xHomocysteineMMTS [K9] | 0,0004709 | 1195,58498 | Brain/ FAIMS/MMTS |
| P56393 | P56393 [72-78] | [R].VTPKEWR.[D] | 1xHomocysteineIAA [K4] | 0,000756219 | 1089,55098 | Liver/ No FAIMS/IAA |
| P56393 | P56393 [72-78] | [R].VTPKEWR.[D] | 1xHomocysteineIAA [K4] | 0,000756219 | 1089,55098 | Liver/ FAIMS/IAA |
| P56393 | P56393 [72-78] | [R].VTPKEWR.[D] | 1xHomocysteineIAA [K4] | 0,000756219 | 1089,55098 | Brain/ No FAIMS/IAA |
| P56393 | P56393 [72-78] | [R].VTPKEWR.[D] | 1xHomocysteineIAA [K4] | 0,000756219 | 1089,55098 | Brain/ FAIMS/IAA |
| P56395 | P56395 [25-39] | [K].STWVILHHKVYDLTK.[F] | 1xHomocysteineIAA [K9] | 4,32935E-05 | 2014,05788 | Liver/ No FAIMS/IAA |
| P56395 | P56395 [25-39] | [K].STWVILHHKVYDLTK.[F] | 1xHomocysteineIAA [K9] | 4,32935E-05 | 2014,05788 | Liver/ FAIMS/IAA |
| P56395 | P56395 [1-10] | [-].MAGQSDKDVK.[Y] | 1xMet-loss+Acetyl [N-Term]; 1xHomocysteineMMTS [K7] | 0,000192903 | 1152,50237 | Liver/ No FAIMS/MMTS |
| P56395 | P56395 [1-19] | [-].MAGQSDKDVKYYTLEEIQK.[H] | 1xMet-loss+Acetyl [N-Term]; 1xHomocysteineMMTS [K10] | 4,49774E-05 | 2320,08356 | Liver/ No FAIMS/MMTS |
| P56395 | P56395 [25-39] | [K].STWVILHHKVYDLTK.[F] | 1xHomocysteineMMTS [K9] | 4,49774E-05 | 2003,02413 | Liver/ No FAIMS/MMTS |
| P56395 | P56395 [11-21] | [K].YYTLEEIQKHK.[D] | 1xHomocysteineMMTS [K9] | 4,49774E-05 | 1614,76546 | Liver/ No FAIMS/MMTS |
| P56395 | P56395 [20-33] | [K].HKDSKSTWVILHHK.[V] | 1xHomocysteineMMTS [K5] | 0,00182026 | 1878,94655 | Liver/ No FAIMS/MMTS |
| P56395 | P56395 [74-89] | [R].ELSKTYIIGELHPDDR.[S] | 1xHomocysteineMMTS [K4] | 4,49774E-05 | 2048,97797 | Liver/ No FAIMS/MMTS |
| P56395 | P56395 [22-33] | [K].DSKSTWVILHHK.[V] | 1xHomocysteineMMTS [K3] | 4,49774E-05 | 1613,79268 | Liver/ No FAIMS/MMTS |
| P56395 | P56395 [1-10] | [-].MAGQSDKDVK.[Y] | 1xMet-loss+Acetyl [N-Term]; 1xHomocysteineMMTS [K7] | 0,000192903 | 1152,50237 | Liver/ FAIMS/MMTS |
| P56395 | P56395 [1-19] | [-].MAGQSDKDVKYYTLEEIQK.[H] | 1xMet-loss+Acetyl [N-Term]; 1xHomocysteineMMTS [K10] | 4,49774E-05 | 2320,08356 | Liver/ FAIMS/MMTS |
| P56395 | P56395 [25-39] | [K].STWVILHHKVYDLTK.[F] | 1xHomocysteineMMTS [K9] | 4,49774E-05 | 2003,02413 | Liver/ FAIMS/MMTS |
| P56395 | P56395 [11-21] | [K].YYTLEEIQKHK.[D] | 1xHomocysteineMMTS [K9] | 4,49774E-05 | 1614,76546 | Liver/ FAIMS/MMTS |
| P56395 | P56395 [34-52] | [K].VYDLTKFLEEHPGGEEVLR.[E] | 1xHomocysteineMMTS [K6] | 4,49774E-05 | 2394,14683 | Liver/ FAIMS/MMTS |
| P56395 | P56395 [74-89] | [R].ELSKTYIIGELHPDDR.[S] | 1xHomocysteineMMTS [K4] | 4,49774E-05 | 2048,97797 | Liver/ FAIMS/MMTS |
| P56395 | P56395 [20-33] | [K].HKDSKSTWVILHHK.[V] | 1xHomocysteineMMTS [K5] | 0,00182026 | 1878,94655 | Liver/ FAIMS/MMTS |
| P56395 | P56395 [22-33] | [K].DSKSTWVILHHK.[V] | 1xHomocysteineMMTS [K3] | 4,49774E-05 | 1613,79268 | Liver/ FAIMS/MMTS |
| P56395 | P56395 [8-19] | [K].DVKYYTLEEIQK.[H] | 1xHomocysteineMMTS [K3] | 0,000266738 | 1691,80191 | Liver/ FAIMS/MMTS |
| P56399 | P56399 [416-423] | [R].MFKALIGK.[G] | 1xHomocysteineIAA [K3] | 0,00137306 | 1081,58967 | Brain/ FAIMS/IAA |
| P56480 | P56480 [122-133] | [R].GQKVLDSGAPIK.[I] | 1xHomocysteineIAA [K3] | 4,32935E-05 | 1386,74096 | Liver/ FAIMS/IAA |
| P56480 | P56480 [351-387] | [K].KGSITSVQAIYVPADDLTDPAPATTFAHL DATTVLSR.[A] | 1xHomocysteineIAA [K1] | 4,32935E-05 | 4017,02737 | Brain/ No FAIMS/IAA |
| P56480 | P56480 [351-387] | [K].KGSITSVQAIYVPADDLTDPAPATTFAHL DATTVLSR.[A] | 1xHomocysteineIAA [K1] | 4,32935E-05 | 4017,02737 | Brain/ FAIMS/IAA |
| P56480 | P56480 [463-485] | [R].FLSQPFQVAEVFTGHMGKLVPLK.[E] | 1xOxidation [M16]; 1xHomocysteineMMTS [K18] | 4,49774E-05 | 2752,40233 | Liver/ No FAIMS/MMTS |
| P56480 | P56480 [125-143] | [K].VLDSGAPIKIPVGPETLGR.[I] | 1xHomocysteineMMTS [K9] | 4,49774E-05 | 2082,10859 | Liver/ No FAIMS/MMTS |
| P56480 | P56480 [260-279] | [K].DATSKVALVYGQMNEPPGAR.[A] | 1xHomocysteineMMTS [K5] | 4,49774E-05 | 2267,06172 | Liver/ No FAIMS/MMTS |
| P56480 | P56480 [481-489] | [K].LVPLKETIK.[G] | 1xHomocysteineMMTS [K5] | 0,00375052 | 1203,68396 | Liver/ No FAIMS/MMTS |
| P56480 | P56480 [423-432] | [R].GVQKILQDYK.[S] | 1xHomocysteineMMTS [K4] | 4,49774E-05 | 1354,68575 | Liver/ No FAIMS/MMTS |
| P56480 | P56480 [199-212] | [K].GGKIGLFGGAGVGK.[T] | 1xHomocysteineMMTS [K3] | 4,49774E-05 | 1380,71264 | Liver/ No FAIMS/MMTS |
| P56480 | P56480 [122-133] | [R].GQKVLDSGAPIK.[I] | 1xHomocysteineMMTS [K3] | 4,49774E-05 | 1375,70722 | Liver/ No FAIMS/MMTS |
| P56480 | P56480 [520-529] | [K].ADKLAEEHGS.[-] | 1xHomocysteineMMTS [K3] | 0,00848892 | 1219,50818 | Liver/ No FAIMS/MMTS |
| P56480 | P56480 [189-201] | [K].VVDLLAPYAKGGK.[I] | 1xHomocysteineMMTS [K10] | 4,49774E-05 | 1493,78547 | Liver/ No FAIMS/MMTS |
| P56480 | P56480 [351-387] | [K].KGSITSVQAIYVPADDLTDPAPATTFAHL DATTVLSR.[A] | 1xHomocysteineMMTS [K1] | 4,49774E-05 | 4005,99363 | Liver/ No FAIMS/MMTS |
| P56480 | P56480 [463-485] | [R].FLSQPFQVAEVFTGHMGKLVPLK.[E] | 1xOxidation [M16]; 1xHomocysteineMMTS [K18] | 4,49774E-05 | 2752,40233 | Liver/ FAIMS/MMTS |
| P56480 | P56480 [125-143] | [K].VLDSGAPIKIPVGPETLGR.[I] | 1xHomocysteineMMTS [K9] | 4,49774E-05 | 2082,10859 | Liver/ FAIMS/MMTS |

| | | | | | | |
|---|---|---|---|---|---|---|
| P56480 | P56480 [481-489] | [K].LVPLKETIK.[G] | 1xHomocysteineMMTS [K5] | 0,00375052 | 1203,68396 | Liver/ FAIMS/MMTS |
| P56480 | P56480 [423-432] | [R].GVQKILQDYK.[S] | 1xHomocysteineMMTS [K4] | 4,49774E-05 | 1354,68575 | Liver/ FAIMS/MMTS |
| P56480 | P56480 [199-212] | [K].GGKIGLFGGAGVGK.[T] | 1xHomocysteineMMTS [K3] | 4,49774E-05 | 1380,71264 | Liver/ FAIMS/MMTS |
| P56480 | P56480 [122-133] | [R].GQKVLDSGAPIK.[I] | 1xHomocysteineMMTS [K3] | 4,49774E-05 | 1375,70722 | Liver/ FAIMS/MMTS |
| P56480 | P56480 [189-201] | [K].VVDLLAPYAKGGK.[I] | 1xHomocysteineMMTS [K10] | 4,49774E-05 | 1493,78547 | Liver/ FAIMS/MMTS |
| P56480 | P56480 [242-264] | [R].EGNDLYHEMIESGVINLKDATSK.[V] | 1xOxidation [M9]; 1xHomocysteineMMTS [K18] | 0,000211896 | 2742,24193 | Liver/ FAIMS/MMTS |
| P56480 | P56480 [520-529] | [K].ADKLAEEHGS.[-] | 1xHomocysteineMMTS [K3] | 0,00848892 | 1219,50818 | Liver/ FAIMS/MMTS |
| P56480 | P56480 [242-264] | [R].EGNDLYHEMIESGVINLKDATSK.[V] | 1xHomocysteineMMTS [K18] | 4,49774E-05 | 2726,24701 | Liver/ FAIMS/MMTS |
| P56480 | P56480 [351-387] | [K].KGSITSVQAIYVPADDLTDPAPATTFAHL DATTVLSR.[A] | 1xHomocysteineMMTS [K1] | 4,49774E-05 | 4005,99363 | Liver/ FAIMS/MMTS |
| P56480 | P56480 [125-143] | [K].VLDSGAPIKIPVGPETLGR.[I] | 1xHomocysteineMMTS [K9] | 4,49774E-05 | 2082,10859 | Brain/ No FAIMS/MMTS |
| P56480 | P56480 [260-279] | [K].DATSKVALVYGQMNEPPGAR.[A] | 1xHomocysteineMMTS [K5] | 4,49774E-05 | 2267,06172 | Brain/ No FAIMS/MMTS |
| P56480 | P56480 [481-489] | [K].LVPLKETIK.[G] | 1xHomocysteineMMTS [K5] | 0,00375052 | 1203,68396 | Brain/ No FAIMS/MMTS |
| P56480 | P56480 [199-212] | [K].GGKIGLFGGAGVGK.[T] | 1xHomocysteineMMTS [K3] | 4,49774E-05 | 1380,71264 | Brain/ No FAIMS/MMTS |
| P56480 | P56480 [122-143] | [R].GQKVLDSGAPIKIPVGPETLGR.[I] | 1xHomocysteineMMTS [K3] | 0,00382737 | 2395,2836 | Brain/ No FAIMS/MMTS |
| P56480 | P56480 [520-529] | [K].ADKLAEEHGS.[-] | 1xHomocysteineMMTS [K3] | 0,00848892 | 1219,50818 | Brain/ No FAIMS/MMTS |
| P56480 | P56480 [189-201] | [K].VVDLLAPYAKGGK.[I] | 1xHomocysteineMMTS [K10] | 4,49774E-05 | 1493,78547 | Brain/ No FAIMS/MMTS |
| P56480 | P56480 [125-143] | [K].VLDSGAPIKIPVGPETLGR.[I] | 1xHomocysteineMMTS [K9] | 4,49774E-05 | 2082,10859 | Brain/ FAIMS/MMTS |
| P56480 | P56480 [260-279] | [K].DATSKVALVYGQMNEPPGAR.[A] | 1xHomocysteineMMTS [K5] | 4,49774E-05 | 2267,06172 | Brain/ FAIMS/MMTS |
| P56480 | P56480 [481-489] | [K].LVPLKETIK.[G] | 1xHomocysteineMMTS [K5] | 0,00375052 | 1203,68396 | Brain/ FAIMS/MMTS |
| P56480 | P56480 [423-432] | [R].GVQKILQDYK.[S] | 1xHomocysteineMMTS [K4] | 4,49774E-05 | 1354,68575 | Brain/ FAIMS/MMTS |
| P56480 | P56480 [199-212] | [K].GGKIGLFGGAGVGK.[T] | 1xHomocysteineMMTS [K3] | 4,49774E-05 | 1380,71264 | Brain/ FAIMS/MMTS |
| P56480 | P56480 [122-133] | [R].GQKVLDSGAPIK.[I] | 1xHomocysteineMMTS [K3] | 4,49774E-05 | 1375,70722 | Brain/ FAIMS/MMTS |
| P56480 | P56480 [520-529] | [K].ADKLAEEHGS.[-] | 1xHomocysteineMMTS [K3] | 0,00848892 | 1219,50818 | Brain/ FAIMS/MMTS |
| P56480 | P56480 [242-264] | [R].EGNDLYHEMIESGVINLKDATSK.[V] | 1xHomocysteineMMTS [K18] | 4,49774E-05 | 2726,24701 | Brain/ FAIMS/MMTS |
| P56480 | P56480 [189-201] | [K].VVDLLAPYAKGGK.[I] | 1xHomocysteineMMTS [K10] | 4,49774E-05 | 1493,78547 | Brain/ FAIMS/MMTS |
| P56546; O88712 | P56546 [280-289]; O88712 [274-283] | [K].ALAQALKEGR.[I] | 1xHomocysteineIAA [K7] | 9,61721E-05 | 1230,66232 | Brain/ FAIMS/IAA |
| P56564 | P56564 [521-542] | [K].KPYQLIAQDNEPEKPVADSETK.[M] | 1xHomocysteineIAA [K] | 4,32935E-05 | 2674,30287 | Brain/ No FAIMS/IAA |
| P56564 | P56564 [521-542] | [K].KPYQLIAQDNEPEKPVADSETK.[M] | 1xHomocysteineIAA [K] | 4,32935E-05 | 2674,30287 | Brain/ FAIMS/IAA |
| P56564 | P56564 [521-543] | [K].KPYQLIAQDNEPEKPVADSETKM.[-] | 1xHomocysteineMMTS [K22] | 4,49774E-05 | 2794,30961 | Brain/ FAIMS/MMTS |
| P56564 | P56564 [521-542] | [K].KPYQLIAQDNEPEKPVADSETK.[M] | 1xHomocysteineMMTS [K1] | 4,49774E-05 | 2663,26913 | Brain/ FAIMS/MMTS |
| P56564; O35544 | P56564 [176-191]; O35544 [181-196] | [R].NMFPPNLVEACFKQFK.[T] | 1xCarbamidomethyl [C11]; 1xHomocysteineIAA [K13] | 4,32935E-05 | 2144,01258 | Brain/ No FAIMS/IAA |
| P56564; O35544 | P56564 [176-191]; O35544 [181-196] | [R].NMFPPNLVEACFKQFK.[T] | 1xCarbamidomethyl [C11]; 1xHomocysteineIAA [K13] | 4,32935E-05 | 2144,01258 | Brain/ FAIMS/IAA |
| P56565 | P56565 [36-57] | [K].DLLQTELSGFLDVQKDADAVDK.[V] | 1xHomocysteineIAA [K15] | 4,32935E-05 | 2594,26542 | Brain/ FAIMS/IAA |
| P56959 | P56959 [442-465] | [K].APKPDGPGGGPGGSHMGGNYGDDR.[R ] | 1xHomocysteineIAA [K3] | 4,32935E-05 | 2427,02006 | Brain/ FAIMS/IAA |
| P57746 | P57746 [14-22] | [R].MAQTIMKAR.[L] | 1xHomocysteineIAA [K7] | 0,00117224 | 1223,60573 | Brain/ No FAIMS/IAA |
| P57746 | P57746 [14-22] | [R].MAQTIMKAR.[L] | 1xHomocysteineIAA [K7] | 0,00117224 | 1223,60573 | Brain/ FAIMS/IAA |
| P57759 | P57759 [180-194] | [R].QAILKQGQDGLLSVK.[E] | 1xHomocysteineIAA [K5] | 0,00012645 | 1771,97348 | Brain/ FAIMS/IAA |
| P57776 | P57776 [1-15] | [-].MATNFLAHEKIWFDK.[F] | 1xMet-loss+Acetyl [N-Term]; 1xHomocysteineMMTS [K10] | 4,49774E-05 | 1924,90844 | Liver/ FAIMS/MMTS |
| P57776 | P57776 [112-123] | [R].LSSLEKSSPTPR.[A] | 1xHomocysteineMMTS [K6] | 4,49774E-05 | 1464,71851 | Liver/ FAIMS/MMTS |
| P57780; Q7TPR4 | P57780 [412-420]; Q7TPR4 [392-400] | [R].LDHLAEKFR.[Q] | 1xHomocysteineIAA [K7] | 4,32935E-05 | 1302,66232 | Brain/ FAIMS/IAA |
| P57780; Q9JI91; Q7TPR4 | P57780 [456-470]; Q9JI91 [443-457]; Q7TPR4 [436-450] | [RK].KHEAFESDLAAHQDR.[V] | 1xHomocysteineIAA [K1] | 4,32935E-05 | 1927,87153 | Brain/ No FAIMS/IAA |
| P57780; Q9JI91; Q7TPR4 | P57780 [456-470]; Q9JI91 [443-457]; Q7TPR4 [436-450] | [RK].KHEAFESDLAAHQDR.[V] | 1xHomocysteineIAA [K1] | 4,32935E-05 | 1927,87153 | Brain/ FAIMS/IAA |
| P57780; Q9JI91; Q7TPR4 | P57780 [456-470]; Q9JI91 [443-457]; Q7TPR4 [436-450] | [RK].KHEAFESDLAAHQDR.[V] | 1xHomocysteineMMTS [K1] | 0,000564742 | 1916,83779 | Liver/ FAIMS/MMTS |
| P58252 | P58252 [236-249] | [K].FAAKGEGQLSAAER.[A] | 1xHomocysteineIAA [K4] | 9,61721E-05 | 1608,77987 | Brain/ FAIMS/IAA |
| P58281 | P58281 [215-231] | [K].IDQLQEELLHTQLKYQR.[I] | 1xHomocysteineIAA [K14] | 9,61721E-05 | 2329,19689 | Brain/ FAIMS/IAA |
| P58771 | P58771 [252-266] | [K].SIDDLEDELYAQKLK.[Y] | 1xHomocysteineIAA [K13] | 4,32935E-05 | 1953,94738 | Liver/ No FAIMS/IAA |
| P58771 | P58771 [252-266] | [K].SIDDLEDELYAQKLK.[Y] | 1xHomocysteineIAA [K13] | 4,32935E-05 | 1953,94738 | Liver/ FAIMS/IAA |
| P58771 | P58771 [183-198] | [R].AELSEGKCAELEEELK.[T] | 1xMethylthio [C8]; 1xHomocysteineMMTS [K7] | 4,49774E-05 | 1986,8527 | Liver/ No FAIMS/MMTS |
| P58771 | P58771 [199-213] | [K].TVTNNLKSLEAQAEK.[Y] | 1xHomocysteineMMTS [K7] | 4,49774E-05 | 1808,88809 | Liver/ No FAIMS/MMTS |
| P58771 | P58771 [150-160] | [K].EAKHIAEDADR.[K] | 1xHomocysteineMMTS [K3] | 0,000266738 | 1417,61986 | Liver/ No FAIMS/MMTS |
| P58771 | P58771 [36-48] | [R].SKQLEDELVSLQK.[K] | 1xHomocysteineMMTS [K2] | 4,49774E-05 | 1679,83427 | Liver/ No FAIMS/MMTS |
| P58771 | P58771 [252-266] | [K].SIDDLEDELYAQKLK.[Y] | 1xHomocysteineMMTS [K13] | 4,49774E-05 | 1942,91364 | Liver/ No FAIMS/MMTS |
| P58771 | P58771 [38-49] | [K].QLEDELVSLQKK.[L] | 1xHomocysteineMMTS [K11] | 0,00130669 | 1592,80224 | Liver/ No FAIMS/MMTS |
| P58771 | P58771 [168-178] | [R].KLVIIESDLER.[A] | 1xHomocysteineMMTS [K1] | 4,49774E-05 | 1477,7753 | Liver/ No FAIMS/MMTS |
| P58771 | P58771 [134-149] | [R].AQKDEEKMEIQEIQLK.[E] | 1xHomocysteineMMTS [K] | 4,49774E-05 | 2123,01812 | Liver/ No FAIMS/MMTS |
| P58771 | P58771 [183-198] | [R].AELSEGKCAELEEELK.[T] | 1xMethylthio [C8]; 1xHomocysteineMMTS [K7] | 4,49774E-05 | 1986,8527 | Liver/ FAIMS/MMTS |
| P58771 | P58771 [199-213] | [K].TVTNNLKSLEAQAEK.[Y] | 1xHomocysteineMMTS [K7] | 4,49774E-05 | 1808,88809 | Liver/ FAIMS/MMTS |
| P58771 | P58771 [36-48] | [R].SKQLEDELVSLQK.[K] | 1xHomocysteineMMTS [K2] | 4,49774E-05 | 1679,83427 | Liver/ FAIMS/MMTS |
| P58771 | P58771 [267-284] | [K].YKAISEELDHALNDMTSI.[-] | 1xHomocysteineMMTS [K2] | 0,00995323 | 2212,9923 | Liver/ FAIMS/MMTS |
| P58771 | P58771 [38-49] | [K].QLEDELVSLQKK.[L] | 1xHomocysteineMMTS [K11] | 0,00130669 | 1592,80224 | Liver/ FAIMS/MMTS |
| P58771 | P58771 [168-178] | [R].KLVIIESDLER.[A] | 1xHomocysteineMMTS [K1] | 4,49774E-05 | 1477,7753 | Liver/ FAIMS/MMTS |
| P58771 | P58771 [150-160] | [K].EAKHIAEDADR.[K] | 1xHomocysteineMMTS [K3] | 0,000266738 | 1417,61986 | Liver/ FAIMS/MMTS |
| P58771 | P58771 [249-264] | [K].LEKSIDDLEDELYAQK.[L] | 1xHomocysteineMMTS [K3] | 4,49774E-05 | 2071,95623 | Liver/ FAIMS/MMTS |
| P58771 | P58771 [252-266] | [K].SIDDLEDELYAQKLK.[Y] | 1xHomocysteineMMTS [K13] | 4,49774E-05 | 1942,91364 | Liver/ FAIMS/MMTS |
| P58771 | P58771 [22-35] | [R].AEQAEADKKAAEDR.[S] | 1xHomocysteineMMTS [K] | 4,49774E-05 | 1694,74724 | Liver/ FAIMS/MMTS |
| P58771 | P58771 [134-149] | [R].AQKDEEKMEIQEIQLK.[E] | 1xHomocysteineMMTS [K] | 4,49774E-05 | 2123,01812 | Liver/ FAIMS/MMTS |

| | | | | | | |
|---|---|---|---|---|---|---|
| P58771 | P58771 [168-178] | [R].KLVIIESDLER.[A] | 1xHomocysteineMMTS [K1] | 4,49774E-05 | 1477,7753 | Brain/ No FAIMS/MMTS |
| P58774 | P58774 [38-49] | [K].QLEEEQQALQKK.[L] | 1xHomocysteineIAA [K11] | 4,32935E-05 | 1645,8214 | Liver/ No FAIMS/IAA |
| P58774 | P58774 [38-49] | [K].QLEEEQQALQKK.[L] | 1xHomocysteineIAA [K11] | 4,32935E-05 | 1645,8214 | Liver/ FAIMS/IAA |
| P58774 | P58774 [183-198] | [R].AEVAESKCGDLEEELK.[I] | 1xMethylthio [C8]; 1xHomocysteineMMTS [K7] | 4,49774E-05 | 1958,8214 | Liver/ No FAIMS/MMTS |
| P58774 | P58774 [206-217] | [K].SLEAQADKYSTK.[E] | 1xHomocysteineMMTS [K8] | 4,49774E-05 | 1503,68179 | Liver/ No FAIMS/MMTS |
| P58774 | P58774 [199-213] | [K].IVTNNLKSLEAQADK.[Y] | 1xHomocysteineMMTS [K7] | 4,49774E-05 | 1806,90883 | Liver/ No FAIMS/MMTS |
| P58774 | P58774 [60-70] | [K].YSESVKDAQEK.[L] | 1xHomocysteineMMTS [K6] | 4,49774E-05 | 1446,62394 | Liver/ No FAIMS/MMTS |
| P58774 | P58774 [66-76] | [K].DAQEKLEQAEK.[K] | 1xHomocysteineMMTS [K5] | 0,0018995 | 1451,65049 | Liver/ No FAIMS/MMTS |
| P58774 | P58774 [150-160] | [K].EAKHIAEDSDR.[K] | 1xHomocysteineMMTS [K3] | 0,000211896 | 1433,61477 | Liver/ No FAIMS/MMTS |
| P58774 | P58774 [249-264] | [K].LEKTIDDLEDEVYAQK.[M] | 1xHomocysteineMMTS [K3] | 4,49774E-05 | 2071,95623 | Liver/ No FAIMS/MMTS |
| P58774 | P58774 [50-59] | [K].LKGTEDEVEK.[Y] | 1xHomocysteineMMTS [K2] | 0,00334305 | 1310,59666 | Liver/ No FAIMS/MMTS |
| P58774 | P58774 [252-266] | [K].TIDDLEDEVYAQKMK.[Y] | 1xHomocysteineMMTS [K13] | 4,49774E-05 | 1960,87006 | Liver/ No FAIMS/MMTS |
| P58774 | P58774 [168-178] | [R].KLVILEGELER.[S] | 1xHomocysteineMMTS [K1] | 4,49774E-05 | 1461,78038 | Liver/ No FAIMS/MMTS |
| P58774 | P58774 [183-198] | [R].AEVAESKCGDLEEELK.[I] | 1xMethylthio [C8]; 1xHomocysteineMMTS [K7] | 4,49774E-05 | 1958,8214 | Liver/ FAIMS/MMTS |
| P58774 | P58774 [206-217] | [K].SLEAQADKYSTK.[E] | 1xHomocysteineMMTS [K8] | 4,49774E-05 | 1503,68179 | Liver/ FAIMS/MMTS |
| P58774 | P58774 [199-213] | [K].IVTNNLKSLEAQADK.[Y] | 1xHomocysteineMMTS [K7] | 4,49774E-05 | 1806,90883 | Liver/ FAIMS/MMTS |
| P58774 | P58774 [60-70] | [K].YSESVKDAQEK.[L] | 1xHomocysteineMMTS [K6] | 4,49774E-05 | 1446,62394 | Liver/ FAIMS/MMTS |
| P58774 | P58774 [150-160] | [K].EAKHIAEDSDR.[K] | 1xHomocysteineMMTS [K3] | 0,000211896 | 1433,61477 | Liver/ FAIMS/MMTS |
| P58774 | P58774 [50-59] | [K].LKGTEDEVEK.[Y] | 1xHomocysteineMMTS [K2] | 0,00334305 | 1310,59666 | Liver/ FAIMS/MMTS |
| P58774 | P58774 [168-178] | [R].KLVILEGELER.[S] | 1xHomocysteineMMTS [K1] | 4,49774E-05 | 1461,78038 | Liver/ FAIMS/MMTS |
| P58774 | P58774 [252-266] | [K].TIDDLEDEVYAQKMK.[Y] | 1xOxidation [M14]; 1xHomocysteineMMTS [K13] | 0,000152766 | 1976,86498 | Liver/ FAIMS/MMTS |
| P58774 | P58774 [36-48] | [R].CKQLEEEQQALQK.[K] | 1xMethylthio [C1]; 1xHomocysteineMMTS [K2] | 0,000211896 | 1783,78456 | Liver/ FAIMS/MMTS |
| P58774 | P58774 [141-152] | [K].MELQEMQLKEAK.[H] | 1xHomocysteineMMTS [K9] | 0,000342214 | 1640,75147 | Liver/ FAIMS/MMTS |
| P58774 | P58774 [22-35] | [R].AEQAEADKKQAEDR.[C] | 1xHomocysteineMMTS [K9] | 0,00101551 | 1751,76871 | Liver/ FAIMS/MMTS |
| P58774 | P58774 [66-76] | [K].DAQEKLEQAEK.[K] | 1xHomocysteineMMTS [K5] | 0,0018995 | 1451,65049 | Liver/ FAIMS/MMTS |
| P58774 | P58774 [214-226] | [K].YSTKEDKYEEEIK.[L] | 1xHomocysteineMMTS [K4] | 4,49774E-05 | 1824,80303 | Liver/ FAIMS/MMTS |
| P58774 | P58774 [249-264] | [K].LEKTIDDLEDEVYAQK.[M] | 1xHomocysteineMMTS [K3] | 4,49774E-05 | 2071,95623 | Liver/ FAIMS/MMTS |
| P58774 | P58774 [252-266] | [K].TIDDLEDEVYAQKMK.[Y] | 1xHomocysteineMMTS [K13] | 4,49774E-05 | 1960,87006 | Liver/ FAIMS/MMTS |
| P58774 | P58774 [50-65] | [K].LKGTEDEVEKYSESVK.[D] | 1xHomocysteineMMTS [K10] | 0,00257148 | 2003,93002 | Liver/ FAIMS/MMTS |
| P58774; P21107; P58771 | P58774 [113-125]; P21107 [114-126]; P58771 [113-125] | [K].LEEAEKAADESER.[G] | 1xHomocysteineIAA [K6] | 4,32935E-05 | 1650,72755 | Liver/ No FAIMS/IAA |
| P58774; P21107; P58771 | P58774 [113-125]; P21107 [114-126]; P58771 [113-125] | [K].LEEAEKAADESER.[G] | 1xHomocysteineIAA [K6] | 4,32935E-05 | 1650,72755 | Liver/ FAIMS/IAA |
| P58774; P21107; P58771 | P58774 [113-125]; P21107 [114-126]; P58771 [113-125] | [K].LEEAEKAADESER.[G] | 1xHomocysteineIAA [K6] | 4,32935E-05 | 1650,72755 | Brain/ FAIMS/IAA |
| P58774; P21107; P58771 | P58774 [106-118]; P21107 [107-119]; P58771 [106-118] | [R].LATALQKLEEAEK.[A] | 1xHomocysteineMMTS [K7] | 4,49774E-05 | 1606,81789 | Liver/ No FAIMS/MMTS |
| P58774; P21107; P58771 | P58774 [113-125]; P21107 [114-126]; P58771 [113-125] | [K].LEEAEKAADESER.[G] | 1xHomocysteineMMTS [K6] | 4,49774E-05 | 1639,69381 | Liver/ No FAIMS/MMTS |
| P58774; P21107; P58771 | P58774 [106-118]; P21107 [107-119]; P58771 [106-118] | [R].LATALQKLEEAEK.[A] | 1xHomocysteineMMTS [K7] | 4,49774E-05 | 1606,81789 | Liver/ FAIMS/MMTS |
| P58774; P21107; P58771 | P58774 [113-125]; P21107 [114-126]; P58771 [113-125] | [K].LEEAEKAADESER.[G] | 1xHomocysteineMMTS [K6] | 4,49774E-05 | 1639,69381 | Liver/ FAIMS/MMTS |
| P58774; P21107; P58771 | P58774 [106-118]; P21107 [107-119]; P58771 [106-118] | [R].LATALQKLEEAEK.[A] | 1xHomocysteineMMTS [K7] | 4,49774E-05 | 1606,81789 | Brain/ No FAIMS/MMTS |
| P58774; P21107; P58771 | P58774 [106-118]; P21107 [107-119]; P58771 [106-118] | [R].LATALQKLEEAEK.[A] | 1xHomocysteineMMTS [K7] | 4,49774E-05 | 1606,81789 | Brain/ FAIMS/MMTS |
| P58774; P21107; P58771 | P58774 [113-125]; P21107 [114-126]; P58771 [113-125] | [K].LEEAEKAADESER.[G] | 1xHomocysteineMMTS [K6] | 4,49774E-05 | 1639,69381 | Brain/ FAIMS/MMTS |
| P58774; P21107; Q6IRU2 | P58774 [126-133]; P21107 [127-134]; Q6IRU2 [90-97] | [R].GMKVIENR.[A] | 1xHomocysteineIAA [K3] | 0,00590083 | 1120,56016 | Liver/ FAIMS/IAA |
| P58774; P21107; Q6IRU2 | P58774 [126-133]; P21107 [127-134]; Q6IRU2 [90-97] | [R].GMKVIENR.[A] | 1xHomocysteineIAA [K3] | 0,00590083 | 1120,56016 | Brain/ No FAIMS/IAA |
| P58774; P21107; Q6IRU2 | P58774 [126-133]; P21107 [127-134]; Q6IRU2 [90-97] | [R].GMKVIENR.[A] | 1xHomocysteineIAA [K3] | 0,00590083 | 1120,56016 | Brain/ FAIMS/IAA |
| P58774; P21107; Q6IRU2; P58771 | P58774 [161-167]; P21107 [162-168]; Q6IRU2 [125-131]; P58771 [161-167] | [R].KYEEVAR.[K] | 1xHomocysteineMMTS [K1] | 0,00703166 | 1057,48051 | Liver/ No FAIMS/MMTS |
| P58774; P21107; Q6IRU2; P58771 | P58774 [161-167]; P21107 [162-168]; Q6IRU2 [125-131]; P58771 [161-167] | [R].KYEEVAR.[K] | 1xHomocysteineMMTS [K1] | 0,00703166 | 1057,48051 | Liver/ FAIMS/MMTS |
| P58774; P58771 | P58774 [77-90]; P58771 [77-90] | [K].KATDAEADVASLNR.[R] | 1xHomocysteineIAA [K1] | 4,32935E-05 | 1634,78026 | Liver/ No FAIMS/IAA |

| | | | | | | |
|---|---|---|---|---|---|---|
| P58774; P58771 | P58774 [77-90]; P58771 [77-90] | [K].KATDAEADVASLNR.[R] | 1xHomocysteineIAA [K1] | 4,32935E-05 | 1634,78026 | Liver/ FAIMS/IAA |
| P58774; P58771 | P58774 [77-90]; P58771 [77-90] | [K].KATDAEADVASLNR.[R] | 1xHomocysteineIAA [K1] | 4,32935E-05 | 1634,78026 | Brain/ FAIMS/IAA |
| P58774; P58771 | P58774 [8-21]; P58771 [8-21] | [K].MQMLKLDKENALDR.[A] | 1xHomocysteineMMTS [K5] | 4,49774E-05 | 1867,88969 | Liver/ No FAIMS/MMTS |
| P58774; P58771 | P58774 [13-21]; P58771 [13-21] | [K].LDKENALDR.[A] | 1xHomocysteineMMTS [K3] | 4,49774E-05 | 1236,57112 | Liver/ No FAIMS/MMTS |
| P58774; P58771 | P58774 [77-90]; P58771 [77-90] | [K].KATDAEADVASLNR.[R] | 1xHomocysteineMMTS [K1] | 4,49774E-05 | 1623,74652 | Liver/ No FAIMS/MMTS |
| P58774; P58771 | P58774 [8-21]; P58771 [8-21] | [K].MQMLKLDKENALDR.[A] | 1xHomocysteineMMTS [K5] | 4,49774E-05 | 1867,88969 | Liver/ FAIMS/MMTS |
| P58774; P58771 | P58774 [13-21]; P58771 [13-21] | [K].LDKENALDR.[A] | 1xHomocysteineMMTS [K3] | 4,49774E-05 | 1236,57112 | Liver/ FAIMS/MMTS |
| P58774; P58771 | P58774 [77-90]; P58771 [77-90] | [K].KATDAEADVASLNR.[R] | 1xHomocysteineMMTS [K1] | 4,49774E-05 | 1623,74652 | Liver/ FAIMS/MMTS |
| P59648 | P59648 [65-80] | [K].SCKSELPSSAPGGGGV.[-] | 1xCarbamidomethyl [C2]; 1xHomocysteineIAA [K3] | 4,32935E-05 | 1663,74143 | Brain/ No FAIMS/IAA |
| P59648 | P59648 [65-80] | [K].SCKSELPSSAPGGGGV.[-] | 1xCarbamidomethyl [C2]; 1xHomocysteineIAA [K3] | 4,32935E-05 | 1663,74143 | Brain/ FAIMS/IAA |
| P59999 | P59999 [72-84] | [R].VSIAVKQADEIEK.[I] | 1xHomocysteineIAA [K6] | 9,61721E-05 | 1603,83598 | Brain/ FAIMS/IAA |
| P60202 | P60202 [99-111] | [R].QIFGDYKTTICGK.[G] | 1xCarbamidomethyl [C11]; 1xHomocysteineIAA [K7] | 4,32935E-05 | 1704,80839 | Brain/ No FAIMS/IAA |
| P60202 | P60202 [99-111] | [R].QIFGDYKTTICGK.[G] | 1xCarbamidomethyl [C11]; 1xHomocysteineIAA [K7] | 4,32935E-05 | 1704,80839 | Brain/ FAIMS/IAA |
| P60202 | P60202 [112-124] | [K].GLSATVTGGQKGR.[G] | 1xHomocysteineMMTS [K11] | 4,49774E-05 | 1394,68788 | Brain/ FAIMS/MMTS |
| P60469 | P60469 [619-630] | [K].LIQEEKETTEQR.[A] | 1xHomocysteineIAA [K6] | 0,00544356 | 1677,81122 | Brain/ FAIMS/IAA |
| P60521 | P60521 [36-47] | [K].VSGSQIVDIDKR.[K] | 1xHomocysteineIAA [K11] | 9,61721E-05 | 1490,76315 | Brain/ FAIMS/IAA |
| P60710 | P60710 [1-28] | [-].MDDDIAALVVDNGSGMCKAGFAGDDAPR.[A] | 1xCarbamidomethyl [C17]; 1xMet-loss+Acetyl [N-Term]; 1xHomocysteineIAA [K18] | 4,32935E-05 | 2953,27608 | Brain/ No FAIMS/IAA |
| P60710 | P60710 [1-28] | [-].MDDDIAALVVDNGSGMCKAGFAGDDAPR.[A] | 1xCarbamidomethyl [C17]; 1xMet-loss+Acetyl [N-Term]; 1xHomocysteineIAA [K18] | 4,32935E-05 | 2953,27608 | Brain/ FAIMS/IAA |
| P60710; P62737; P68134; P63260 | P60710 [40-61]; P62737 [42-63]; P68134 [42-63]; P63260 [40-61] | [R].HQGVMVGMGQKDSYVGDEAQSK.[R] | 1xHomocysteineIAA [K11] | 4,32935E-05 | 2525,12175 | Liver/ FAIMS/IAA |
| P60710; P62737; P68134; P63260 | P60710 [40-62]; P62737 [42-64]; P68134 [42-64]; P63260 [40-62] | [R].HQGVMVGMGQKDSYVGDEAQSKR.[G] | 2xHomocysteineIAA [K11; K22] | 4,32935E-05 | 2855,26916 | Brain/ FAIMS/IAA |
| P60710; P62737; P68134; P63260 | P60710 [313-326]; P62737 [315-328]; P68134 [315-328]; P63260 [313-326] | [R].MQKEITALAPSTMK.[I] | 2xOxidation [M1; M13]; 1xHomocysteineMMTS [K3] | 4,49774E-05 | 1743,8148 | Liver/ No FAIMS/MMTS |
| P60710; P62737; P68134; P63260 | P60710 [316-328]; P62737 [318-330]; P68134 [318-330]; P63260 [316-328] | [K].EITALAPSTMKIK.[I] | 1xOxidation [M10]; 1xHomocysteineMMTS [K11] | 4,49774E-05 | 1581,80488 | Liver/ No FAIMS/MMTS |
| P60710; P62737; P68134; P63260 | P60710 [313-326]; P62737 [315-328]; P68134 [315-328]; P63260 [313-326] | [R].MQKEITALAPSTMK.[I] | 1xOxidation [M]; 1xHomocysteineMMTS [K3] | 4,49774E-05 | 1727,81988 | Liver/ No FAIMS/MMTS |
| P60710; P62737; P68134; P63260 | P60710 [40-61]; P62737 [42-63]; P68134 [42-63]; P63260 [40-61] | [R].HQGVMVGMGQKDSYVGDEAQSK.[R] | 1xOxidation [M]; 1xHomocysteineMMTS [K11] | 4,49774E-05 | 2530,08292 | Liver/ No FAIMS/MMTS |
| P60710; P62737; P68134; P63260 | P60710 [313-326]; P62737 [315-328]; P68134 [315-328]; P63260 [313-326] | [R].MQKEITALAPSTMK.[I] | 1xHomocysteineMMTS [K3] | 4,49774E-05 | 1711,82497 | Liver/ No FAIMS/MMTS |
| P60710; P62737; P68134; P63260 | P60710 [316-328]; P62737 [318-330]; P68134 [318-330]; P63260 [316-328] | [K].EITALAPSTMKIK.[I] | 1xHomocysteineMMTS [K11] | 4,49774E-05 | 1565,80997 | Liver/ No FAIMS/MMTS |
| P60710; P62737; P68134; P63260 | P60710 [40-61]; P62737 [42-63]; P68134 [42-63]; P63260 [40-61] | [R].HQGVMVGMGQKDSYVGDEAQSK.[R] | 1xHomocysteineMMTS [K11] | 4,49774E-05 | 2514,08801 | Liver/ No FAIMS/MMTS |
| P60710; P62737; P68134; P63260 | P60710 [51-62]; P62737 [53-64]; P68134 [53-64]; P63260 [51-62] | [K].DSYVGDEAQSKR.[G] | 1xHomocysteineMMTS [K11] | 4,49774E-05 | 1517,6359 | Liver/ No FAIMS/MMTS |
| P60710; P62737; P68134; P63260 | P60710 [316-328]; P62737 [318-330]; P68134 [318-330]; P63260 [316-328] | [K].EITALAPSTMKIK.[I] | 1xOxidation [M10]; 1xHomocysteineMMTS [K11] | 4,49774E-05 | 1581,80488 | Liver/ FAIMS/MMTS |
| P60710; P62737; P68134; P63260 | P60710 [313-326]; P62737 [315-328]; P68134 [315-328]; P63260 [313-326] | [R].MQKEITALAPSTMK.[I] | 1xOxidation [M]; 1xHomocysteineMMTS [K3] | 4,49774E-05 | 1727,81988 | Liver/ FAIMS/MMTS |

| | | | | | | |
|---|---|---|---|---|---|---|
| P60710; P62737; P68134; P63260 | P60710 [40-61]; P62737 [42-63]; P68134 [42-63]; P63260 [40-61] | [R].HQGVMVGMGQKDSYVGDEAQSK.[R] | 1xOxidation [M]; 1xHomocysteineMMTS [K11] | 4,49774E-05 | 2530,08292 | Liver/ FAIMS/MMTS |
| P60710; P62737; P68134; P63260 | P60710 [313-326]; P62737 [315-328]; P68134 [315-328]; P63260 [313-326] | [R].MQKEITALAPSTMK.[I] | 1xHomocysteineMMTS [K3] | 4,49774E-05 | 1711,82497 | Liver/ FAIMS/MMTS |
| P60710; P62737; P68134; P63260 | P60710 [316-328]; P62737 [318-330]; P68134 [318-330]; P63260 [316-328] | [K].EITALAPSTMKIK.[I] | 1xHomocysteineMMTS [K11] | 4,49774E-05 | 1565,80997 | Liver/ FAIMS/MMTS |
| P60710; P62737; P68134; P63260 | P60710 [40-61]; P62737 [42-63]; P68134 [42-63]; P63260 [40-61] | [R].HQGVMVGMGQKDSYVGDEAQSK.[R] | 1xHomocysteineMMTS [K11] | 4,49774E-05 | 2514,08801 | Liver/ FAIMS/MMTS |
| P60710; P62737; P68134; P63260 | P60710 [51-62]; P62737 [53-64]; P68134 [53-64]; P63260 [51-62] | [K].DSYVGDEAQSKR.[G] | 1xHomocysteineMMTS [K11] | 4,49774E-05 | 1517,6359 | Liver/ FAIMS/MMTS |
| P60710; P62737; P68134; P63260 | P60710 [40-61]; P62737 [42-63]; P68134 [42-63]; P63260 [40-61] | [R].HQGVMVGMGQKDSYVGDEAQSK.[R] | 2xOxidation [M5; M8]; 1xHomocysteineMMTS [K11] | 4,49774E-05 | 2546,07784 | Liver/ FAIMS/MMTS |
| P60710; P62737; P68134; P63260 | P60710 [313-326]; P62737 [315-328]; P68134 [315-328]; P63260 [313-326] | [R].MQKEITALAPSTMK.[I] | 2xOxidation [M1; M13]; 1xHomocysteineMMTS [K3] | 4,49774E-05 | 1743,8148 | Liver/ FAIMS/MMTS |
| P60710; P62737; P68134; P63260 | P60710 [313-326]; P62737 [315-328]; P68134 [315-328]; P63260 [313-326] | [R].MQKEITALAPSTMK.[I] | 1xHomocysteineMMTS [K3] | 4,49774E-05 | 1711,82497 | Brain/ No FAIMS/MMTS |
| P60710; P62737; P68134; P63260 | P60710 [316-328]; P62737 [318-330]; P68134 [318-330]; P63260 [316-328] | [K].EITALAPSTMKIK.[I] | 1xHomocysteineMMTS [K11] | 4,49774E-05 | 1565,80997 | Brain/ No FAIMS/MMTS |
| P60710; P62737; P68134; P63260 | P60710 [40-61]; P62737 [42-63]; P68134 [42-63]; P63260 [40-61] | [R].HQGVMVGMGQKDSYVGDEAQSK.[R] | 1xHomocysteineMMTS [K11] | 4,49774E-05 | 2514,08801 | Brain/ No FAIMS/MMTS |
| P60710; P62737; P68134; P63260 | P60710 [51-62]; P62737 [53-64]; P68134 [53-64]; P63260 [51-62] | [K].DSYVGDEAQSKR.[G] | 1xHomocysteineMMTS [K11] | 4,49774E-05 | 1517,6359 | Brain/ No FAIMS/MMTS |
| P60710; P62737; P68134; P63260 | P60710 [316-328]; P62737 [318-330]; P68134 [318-330]; P63260 [316-328] | [K].EITALAPSTMKIK.[I] | 1xOxidation [M10]; 1xHomocysteineMMTS [K11] | 4,49774E-05 | 1581,80488 | Brain/ FAIMS/MMTS |
| P60710; P62737; P68134; P63260 | P60710 [40-61]; P62737 [42-63]; P68134 [42-63]; P63260 [40-61] | [R].HQGVMVGMGQKDSYVGDEAQSK.[R] | 1xOxidation [M]; 1xHomocysteineMMTS [K11] | 4,49774E-05 | 2530,08292 | Brain/ FAIMS/MMTS |
| P60710; P62737; P68134; P63260 | P60710 [313-326]; P62737 [315-328]; P68134 [315-328]; P63260 [313-326] | [R].MQKEITALAPSTMK.[I] | 1xHomocysteineMMTS [K3] | 4,49774E-05 | 1711,82497 | Brain/ FAIMS/MMTS |
| P60710; P62737; P68134; P63260 | P60710 [316-328]; P62737 [318-330]; P68134 [318-330]; P63260 [316-328] | [K].EITALAPSTMKIK.[I] | 1xHomocysteineMMTS [K11] | 4,49774E-05 | 1565,80997 | Brain/ FAIMS/MMTS |
| P60710; P62737; P68134; P63260 | P60710 [40-61]; P62737 [42-63]; P68134 [42-63]; P63260 [40-61] | [R].HQGVMVGMGQKDSYVGDEAQSK.[R] | 1xHomocysteineMMTS [K11] | 4,49774E-05 | 2514,08801 | Brain/ FAIMS/MMTS |
| P60710; P62737; P68134; P63260 | P60710 [51-62]; P62737 [53-64]; P68134 [53-64]; P63260 [51-62] | [K].DSYVGDEAQSKR.[G] | 1xHomocysteineMMTS [K11] | 4,49774E-05 | 1517,6359 | Brain/ FAIMS/MMTS |
| P60710; P63260 | P60710 [291-312]; P63260 [291-312] | [R].KDLYANTVLSGGTTMYPGIADR.[M] | 1xHomocysteineIAA [K1] | 4,32935E-05 | 2517,21122 | Liver/ FAIMS/IAA |
| P60710; P63260 | P60710 [291-312]; P63260 [291-312] | [R].KDLYANTVLSGGTTMYPGIADR.[M] | 1xHomocysteineIAA [K1] | 4,32935E-05 | 2517,21122 | Brain/ No FAIMS/IAA |
| P60710; P63260 | P60710 [291-312]; P63260 [291-312] | [R].KDLYANTVLSGGTTMYPGIADR.[M] | 1xHomocysteineIAA [K1] | 4,32935E-05 | 2517,21122 | Brain/ FAIMS/IAA |
| P60710; P63260 | P60710 [291-312]; P63260 [291-312] | [R].KDLYANTVLSGGTTMYPGIADR.[M] | 1xOxidation [M15]; 1xHomocysteineMMTS [K1] | 4,49774E-05 | 2522,17239 | Liver/ No FAIMS/MMTS |
| P60710; P63260 | P60710 [96-116]; P63260 [96-116] | [R].VAPEEHPVLLTEAPLNPKANR.[E] | 1xHomocysteineMMTS [K18] | 4,49774E-05 | 2458,25811 | Liver/ No FAIMS/MMTS |
| P60710; P63260 | P60710 [291-312]; P63260 [291-312] | [R].KDLYANTVLSGGTTMYPGIADR.[M] | 1xHomocysteineMMTS [K1] | 4,49774E-05 | 2506,17748 | Liver/ No FAIMS/MMTS |
| P60710; P63260 | P60710 [96-116]; P63260 [96-116] | [R].VAPEEHPVLLTEAPLNPKANR.[E] | 1xHomocysteineMMTS [K18] | 4,49774E-05 | 2458,25811 | Liver/ FAIMS/MMTS |

| | | | | | | |
|---|---|---|---|---|---|---|
| P60710; P63260 | P60710 [291-312]; P63260 [291-312] | [R].KDLYANTVLSGGTTMYPGIADR.[M] | 1xHomocysteineMMTS [K1] | 4,49774E-05 | 2506,17748 | Liver/ FAIMS/MMTS |
| P60710; P63260 | P60710 [291-312]; P63260 [291-312] | [R].KDLYANTVLSGGTTMYPGIADR.[M] | 1xOxidation [M15]; 1xHomocysteineMMTS [K1] | 4,49774E-05 | 2522,17239 | Liver/ FAIMS/MMTS |
| P60710; P63260 | P60710 [69-95]; P63260 [69-95] | [K].YPIEHGIVTNWDDMEKIWHHTFYNELR.[V] | 1xOxidation [M14]; 1xHomocysteineMMTS [K16] | 0,00110986 | 3622,63408 | Liver/ FAIMS/MMTS |
| P60710; P63260 | P60710 [69-95]; P63260 [69-95] | [K].YPIEHGIVTNWDDMEKIWHHTFYNELR.[V] | 1xOxidation [M14]; 1xHomocysteineMMTS [K16] | 0,00110986 | 3622,63408 | Brain/ No FAIMS/MMTS |
| P60710; P63260 | P60710 [96-116]; P63260 [96-116] | [R].VAPEEHPVLLTEAPLNPKANR.[E] | 1xHomocysteineMMTS [K18] | 4,49774E-05 | 2458,25811 | Brain/ No FAIMS/MMTS |
| P60710; P63260 | P60710 [291-312]; P63260 [291-312] | [R].KDLYANTVLSGGTTMYPGIADR.[M] | 1xHomocysteineMMTS [K1] | 4,49774E-05 | 2506,17748 | Brain/ No FAIMS/MMTS |
| P60710; P63260 | P60710 [291-312]; P63260 [291-312] | [R].KDLYANTVLSGGTTMYPGIADR.[M] | 1xOxidation [M15]; 1xHomocysteineMMTS [K1] | 4,49774E-05 | 2522,17239 | Brain/ FAIMS/MMTS |
| P60710; P63260 | P60710 [96-116]; P63260 [96-116] | [R].VAPEEHPVLLTEAPLNPKANR.[E] | 1xHomocysteineMMTS [K18] | 4,49774E-05 | 2458,25811 | Brain/ FAIMS/MMTS |
| P60710; P63260 | P60710 [291-312]; P63260 [291-312] | [R].KDLYANTVLSGGTTMYPGIADR.[M] | 1xHomocysteineMMTS [K1] | 4,49774E-05 | 2506,17748 | Brain/ FAIMS/MMTS |
| P60755 | P60755 [717-731] | [R].LTPLTKFGEGDSTIR.[V] | 1xHomocysteineIAA [K6] | 0,00540576 | 1808,92111 | Liver/ No FAIMS/IAA |
| P60761 | P60761 [1-32] | [-].MDCCTESACSKPDDDILDIPLDDPGANAAAAK.[I] | 3xCarbamidomethyl [C3; C4; C9]; 1xMet-loss [N-Term]; 1xHomocysteineIAA [K11] | 4,32935E-05 | 3479,47056 | Brain/ No FAIMS/IAA |
| P60761 | P60761 [1-32] | [-].MDCCTESACSKPDDDILDIPLDDPGANAAAAK.[I] | 3xCarbamidomethyl [C3; C4; C9]; 1xMet-loss [N-Term]; 1xHomocysteineIAA [K11] | 4,32935E-05 | 3479,47056 | Brain/ FAIMS/IAA |
| P60761 | P60761 [54-68] | [R].KGPGPGGPGGAGGAR.[G] | 1xHomocysteineMMTS [K1] | 4,49774E-05 | 1355,6307 | Brain/ No FAIMS/MMTS |
| P60761 | P60761 [54-68] | [R].KGPGPGGPGGAGGAR.[G] | 1xHomocysteineMMTS [K1] | 4,49774E-05 | 1355,6307 | Brain/ FAIMS/MMTS |
| P60764; P63001 | P60764 [1-16]; P63001 [1-16] | [-].MQAIKCVVVGDGAVGK.[T] | 1xAcetyl [N-Term]; 1xCarbamidomethyl [C6]; 1xHomocysteineIAA [K5] | 4,32935E-05 | 1847,91762 | Brain/ FAIMS/IAA |
| P60766 | P60766 [134-147] | [K].QKPITPETAEKLAR.[D] | 1xHomocysteineIAA [K11] | 0,000243434 | 1755,94218 | Brain/ FAIMS/IAA |
| P60867 | P60867 [5-19] | [K].DTGKTPVEPEVAIHR.[I] | 1xHomocysteineIAA [K4] | 4,32935E-05 | 1822,91161 | Liver/ FAIMS/IAA |
| P60867 | P60867 [5-19] | [K].DTGKTPVEPEVAIHR.[I] | 1xHomocysteineIAA [K4] | 4,32935E-05 | 1822,91161 | Brain/ FAIMS/IAA |
| P60867 | P60867 [5-19] | [K].DTGKTPVEPEVAIHR.[I] | 1xHomocysteineMMTS [K4] | 0,0046895 | 1811,87786 | Liver/ FAIMS/MMTS |
| P60879 | P60879 [103-119] | [K].KAWGNNQDGVVASQPAR.[V] | 1xHomocysteineIAA [K1] | 4,32935E-05 | 1971,94537 | Brain/ No FAIMS/IAA |
| P60879 | P60879 [103-119] | [K].KAWGNNQDGVVASQPAR.[V] | 1xHomocysteineIAA [K1] | 4,32935E-05 | 1971,94537 | Brain/ FAIMS/IAA |
| P60879 | P60879 [32-45] | [R].MLQLVEESKDAGIR.[T] | 1xHomocysteineMMTS [K9] | 4,49774E-05 | 1751,84887 | Brain/ No FAIMS/MMTS |
| P60879 | P60879 [32-45] | [R].MLQLVEESKDAGIR.[T] | 1xHomocysteineMMTS [K9] | 4,49774E-05 | 1751,84887 | Brain/ FAIMS/MMTS |
| P60879 | P60879 [60-72] | [R].IEEGMDQINKDMK.[E] | 1xHomocysteineMMTS [K10] | 0,000250158 | 1713,73146 | Brain/ FAIMS/MMTS |
| P60904 | P60904 [38-56] | [K].LALKYHPDKNPDNPEAADK.[F] | 1xHomocysteineIAA [K4] | 4,32935E-05 | 2310,11831 | Brain/ No FAIMS/IAA |
| P60904 | P60904 [38-56] | [K].LALKYHPDKNPDNPEAADK.[F] | 1xHomocysteineIAA [K4] | 4,32935E-05 | 2310,11831 | Brain/ FAIMS/IAA |
| P60904 | P60904 [42-58] | [K].YHPDKNPDNPEAADKFK.[E] | 1xHomocysteineMMTS [K] | 0,000489972 | 2148,94773 | Brain/ FAIMS/MMTS |
| P61021 | P61021 [166-180] | [K].TAMNVNDLFLAIAKK.[L] | 1xHomocysteineIAA [K14] | 0,000273428 | 1822,95539 | Brain/ FAIMS/IAA |
| P61027 | P61027 [96-105] | [K].SFENISKWLR.[N] | 1xHomocysteineIAA [K7] | 9,61721E-05 | 1453,72564 | Brain/ FAIMS/IAA |
| P61027; P63011; Q91V41; O35963; Q8BHC1; P61028; P55258; P62823; P35276 | P61027 [58-70]; P63011 [71-83]; Q91V41 [60-72]; O35963 [82-94]; Q8BHC1 [58-70]; P61028 [57-69]; P55258 [57-69]; P62823 [79-91]; P35276 [71-83] | [KR].IKLQIWDTAGQER.[FY] | 1xHomocysteineIAA [K2] | 4,32935E-05 | 1731,88466 | Brain/ No FAIMS/IAA |
| P61027; P63011; Q91V41; O35963; Q8BHC1; P61028; P55258; P62823; P35276 | P61027 [58-70]; P63011 [71-83]; Q91V41 [60-72]; O35963 [82-94]; Q8BHC1 [58-70]; P61028 [57-69]; P55258 [57-69]; P62823 [79-91]; P35276 [71-83] | [KR].IKLQIWDTAGQER.[FY] | 1xHomocysteineIAA [K2] | 4,32935E-05 | 1731,88466 | Brain/ FAIMS/IAA |
| P61027; P63011; Q91V41; O35963; Q8BHC1; P61028; P55258; P62823; P35276 | P61027 [58-70]; P63011 [71-83]; Q91V41 [60-72]; O35963 [82-94]; Q8BHC1 [58-70]; P61028 [57-69]; P55258 [57-69]; P62823 [79-91]; P35276 [71-83] | [KR].IKLQLWDTAGQER.[FY] | 1xHomocysteineMMTS [K2] | 0,000250158 | 1720,85092 | Brain/ FAIMS/MMTS |
| P61082 | P61082 [34-45] | [R].IQKDINELNLPK.[T] | 1xHomocysteineIAA [K3] | 0,00432311 | 1598,85705 | Brain/ FAIMS/IAA |
| P61087 | P61087 [165-176] | [K].KIENLCAMGFDR.[N] | 1xCarbamidomethyl [C6]; 1xHomocysteineIAA [K1] | 0,000624482 | 1627,73893 | Brain/ FAIMS/IAA |
| P61089 | P61089 [75-85] | [K].IYHPNVDKLGR.[I] | 1xHomocysteineIAA [K8] | 4,32935E-05 | 1485,76309 | Brain/ FAIMS/IAA |
| P61089 | P61089 [54-70] | [K].LELFLPEEYPMAAPKVR.[F] | 1xHomocysteineMMTS [K15] | 4,49774E-05 | 2166,0796 | Brain/ FAIMS/MMTS |
| P61164; Q8R5C5 | P61164 [314-320]; Q8R5C5 [314-320] | [R].LLSEVKK.[L] | 1xHomocysteineIAA [K6] | 0,000639442 | 990,56523 | Brain/ No FAIMS/IAA |
| P61164; Q8R5C5 | P61164 [314-320]; Q8R5C5 [314-320] | [R].LLSEVKK.[L] | 1xHomocysteineIAA [K6] | 0,000639442 | 990,56523 | Brain/ FAIMS/IAA |
| P61205 | P61205 [118-142] | [R].DAVLLVFANKQDLPNAMNAAEITDK.[L] | 1xHomocysteineIAA [K10] | 4,32935E-05 | 2875,43284 | Brain/ FAIMS/IAA |

| | | | | | | |
|---|---|---|---|---|---|---|
| P61205; Q8BSL7 | P61205 [128-149]; Q8BSL7 [128-149] | [K].QDLPNAMNAAEITDKLGLHSLR.[HQ] | 1xHomocysteineIAA [K15] | 4,32935E-05 | 2581,28612 | Brain/ No FAIMS/IAA |
| P61205; Q8BSL7 | P61205 [128-149]; Q8BSL7 [128-149] | [K].QDLPNAMNAAEITDKLGLHSLR.[HQ] | 1xHomocysteineIAA [K15] | 4,32935E-05 | 2581,28612 | Brain/ FAIMS/IAA |
| P61255 | P61255 [135-142] | [K].YKEETIEK.[M] | 1xHomocysteineMMTS [K2] | 0,00479272 | 1202,54317 | Liver/ FAIMS/MMTS |
| P61264 | P61264 [56-82] | [K].QHSAILAAPNPDEKTKQELEDLTADIK.[K] | 1xHomocysteineIAA [K] | 4,32935E-05 | 3149,57832 | Brain/ FAIMS/IAA |
| P61264 | P61264 [173-188] | [K].LAIFTDDIKMDSQMTK.[Q] | 1xHomocysteineMMTS [K9] | 4,49774E-05 | 2019,9258 | Brain/ No FAIMS/MMTS |
| P61264 | P61264 [182-197] | [K].MDSQMTKQALNEIETR.[H] | 1xHomocysteineMMTS [K7] | 4,49774E-05 | 2057,91228 | Brain/ No FAIMS/MMTS |
| P61264 | P61264 [55-69] | [K].KQHSAILAAPNPDEK.[T] | 1xHomocysteineMMTS [K1] | 4,49774E-05 | 1781,8673 | Brain/ No FAIMS/MMTS |
| P61264 | P61264 [173-188] | [K].LAIFTDDIKMDSQMTK.[Q] | 1xHomocysteineMMTS [K9] | 4,49774E-05 | 2019,9258 | Brain/ FAIMS/MMTS |
| P61264 | P61264 [182-197] | [K].MDSQMTKQALNEIETR.[H] | 1xHomocysteineMMTS [K7] | 4,49774E-05 | 2057,91228 | Brain/ FAIMS/MMTS |
| P61264 | P61264 [198-209] | [R].HNEIIKLETSIR.[E] | 1xHomocysteineMMTS [K6] | 0,000173718 | 1615,82946 | Brain/ FAIMS/MMTS |
| P61264 | P61264 [70-82] | [K].TKQELEDLTADIK.[K] | 1xHomocysteineMMTS [K2] | 4,49774E-05 | 1666,80263 | Brain/ FAIMS/MMTS |
| P61264 | P61264 [92-107] | [K].LKAIEQSIEQEEGLNR.[S] | 1xHomocysteineMMTS [K2] | 4,49774E-05 | 2019,98379 | Brain/ FAIMS/MMTS |
| P61264 | P61264 [125-139] | [R].KFVEVMTEYNATQSK.[Y] | 1xHomocysteineMMTS [K1] | 4,49774E-05 | 1937,88057 | Brain/ FAIMS/MMTS |
| P61264 | P61264 [55-69] | [K].KQHSAILAAPNPDEK.[T] | 1xHomocysteineMMTS [K1] | 4,49774E-05 | 1781,8673 | Brain/ FAIMS/MMTS |
| P61294 | P61294 [159-168] | [K].TGYNVKQLFR.[R] | 1xHomocysteineIAA [K6] | 4,32935E-05 | 1399,71508 | Brain/ FAIMS/IAA |
| P61294; P35279 | P61294 [16-32]; P35279 [16-32] | [K].LVFLGEQSVGKTSLITR.[F] | 1xHomocysteineIAA [K11] | 4,32935E-05 | 2022,10522 | Brain/ FAIMS/IAA |
| P61329 | P61329 [172-185] | [R].AWFLGLNKEGQIMK.[G] | 1xHomocysteineIAA [K8] | 4,32935E-05 | 1808,91861 | Brain/ FAIMS/IAA |
| P61358 | P61358 [85-98] | [R].YSVDIPLDKTVVNK.[D] | 1xHomocysteineMMTS [K9] | 4,49774E-05 | 1753,8863 | Liver/ No FAIMS/MMTS |
| P61358 | P61358 [85-98] | [R].YSVDIPLDKTVVNK.[D] | 1xHomocysteineMMTS [K9] | 4,49774E-05 | 1753,8863 | Liver/ FAIMS/MMTS |
| P61358 | P61358 [94-102] | [K].TVVNKDVFR.[D] | 1xHomocysteineMMTS [K5] | 0,00164853 | 1240,61767 | Liver/ FAIMS/MMTS |
| P61458 | P61458 [37-45] | [K].QFHFKDFNR.[A] | 1xHomocysteineIAA [K5] | 0,000318237 | 1412,65281 | Liver/ No FAIMS/IAA |
| P61458 | P61458 [32-41] | [R].DAIFKQFHFK.[D] | 1xHomocysteineMMTS [K5] | 0,000361246 | 1443,69117 | Liver/ No FAIMS/MMTS |
| P61458 | P61458 [37-45] | [K].QFHFKDFNR.[A] | 1xHomocysteineMMTS [K5] | 0,00687493 | 1401,61907 | Liver/ No FAIMS/MMTS |
| P61458 | P61458 [53-72] | [R].VALQAEKLDHHPEWFNVYNK.[V] | 1xHomocysteineMMTS [K7] | 0,00074635 | 2601,23771 | Liver/ FAIMS/MMTS |
| P61458 | P61458 [32-41] | [R].DAIFKQFHFK.[D] | 1xHomocysteineMMTS [K5] | 0,000361246 | 1443,69117 | Liver/ FAIMS/MMTS |
| P61458 | P61458 [37-45] | [K].QFHFKDFNR.[A] | 1xHomocysteineMMTS [K5] | 0,00687493 | 1401,61907 | Liver/ FAIMS/MMTS |
| P61514 | P61514 [8-17] | [K].VGIVGKYGTR.[Y] | 1xHomocysteineIAA [K6] | 0,000114608 | 1223,6565 | Brain/ FAIMS/IAA |
| P61750; P61205; Q8BSL7; P84084 | P61750 [20-36]; P61205 [20-36]; Q8BSL7 [20-36]; P84084 [20-36] | [R].ILMVGLDAAGKTTILYK.[L] | 1xHomocysteineIAA [K11] | 4,32935E-05 | 1981,08607 | Brain/ FAIMS/IAA |
| P61922 | P61922 [232-250] | [K].AIHKIDIPSFDWPIAPFPR.[L] | 1xHomocysteineIAA [K4] | 4,32935E-05 | 2394,24272 | Brain/ FAIMS/IAA |
| P61924 | P61924 [26-39] | [R].LFAKYYDDTYPSVK.[E] | 1xHomocysteineMMTS [K4] | 0,000266738 | 1872,85467 | Liver/ FAIMS/MMTS |
| P61971 | P61971 [50-63] | [K].AAIVEKLSSLPFQK.[I] | 1xHomocysteineMMTS [K6] | 0,00832764 | 1693,90156 | Liver/ FAIMS/MMTS |
| P61979 | P61979 [22-35] | [K].RPAEDMEEEQAFKR.[S] | 1xHomocysteineIAA [K13] | 4,32935E-05 | 1909,85311 | Brain/ No FAIMS/IAA |
| P61979 | P61979 [22-35] | [K].RPAEDMEEEQAFKR.[S] | 1xHomocysteineIAA [K13] | 4,32935E-05 | 1909,85311 | Brain/ FAIMS/IAA |
| P61979 | P61979 [22-35] | [K].RPAEDMEEEQAFKR.[S] | 1xHomocysteineMMTS [K13] | 0,00373588 | 1898,81936 | Liver/ FAIMS/MMTS |
| P61982 | P61982 [62-77] | [R].VISSIEQKTSADGNEK.[K] | 1xHomocysteineIAA [K8] | 4,32935E-05 | 1879,90658 | Liver/ FAIMS/IAA |
| P61982 | P61982 [143-162] | [K].RATVVESSEKAYSEAHEISK.[E] | 1xHomocysteineIAA [K10] | 4,32935E-05 | 2395,15581 | Liver/ FAIMS/IAA |
| P61982 | P61982 [144-162] | [R].ATVVESSEKAYSEAHEISK.[E] | 1xHomocysteineMMTS [K9] | 4,49774E-05 | 2228,02096 | Liver/ FAIMS/MMTS |
| P61982 | P61982 [62-77] | [R].VISSIEQKTSADGNEK.[K] | 1xHomocysteineMMTS [K8] | 4,49774E-05 | 1868,87284 | Liver/ FAIMS/MMTS |
| P61982 | P61982 [144-162] | [R].ATVVESSEKAYSEAHEISK.[E] | 1xHomocysteineMMTS [K9] | 4,49774E-05 | 2228,02096 | Brain/ No FAIMS/MMTS |
| P61982 | P61982 [20-42] | [R].YDDMAAAMKNVTELNEPLSNEER.[N] | 1xHomocysteineMMTS [K9] | 4,49774E-05 | 2803,20416 | Brain/ No FAIMS/MMTS |
| P61982 | P61982 [5-12] | [R].EQLVQKAR.[L] | 1xHomocysteineMMTS [K6] | 0,008514 | 1134,57581 | Brain/ No FAIMS/MMTS |
| P61982 | P61982 [133-143] | [R].YLAEVATGEKR.[A] | 1xHomocysteineMMTS [K10] | 6,70238E-05 | 1399,67083 | Brain/ No FAIMS/MMTS |
| P61982 | P61982 [153-172] | [K].AYSEAHEISKEHMQPTHPIR.[L] | 1xHomocysteineMMTS [K10] | 4,49774E-05 | 2524,15299 | Brain/ No FAIMS/MMTS |
| P61982 | P61982 [144-162] | [R].ATVVESSEKAYSEAHEISK.[E] | 1xHomocysteineMMTS [K9] | 4,49774E-05 | 2228,02096 | Brain/ FAIMS/MMTS |
| P61982 | P61982 [20-42] | [R].YDDMAAAMKNVTELNEPLSNEER.[N] | 1xHomocysteineMMTS [K9] | 4,49774E-05 | 2803,20416 | Brain/ FAIMS/MMTS |
| P61982 | P61982 [62-77] | [R].VISSIEQKTSADGNEK.[K] | 1xHomocysteineMMTS [K8] | 4,49774E-05 | 1868,87284 | Brain/ FAIMS/MMTS |
| P61982 | P61982 [5-12] | [R].EQLVQKAR.[L] | 1xHomocysteineMMTS [K6] | 0,008514 | 1134,57581 | Brain/ FAIMS/MMTS |
| P61982 | P61982 [133-143] | [R].YLAEVATGEKR.[A] | 1xHomocysteineMMTS [K10] | 6,70238E-05 | 1399,67083 | Brain/ FAIMS/MMTS |
| P61982 | P61982 [153-172] | [K].AYSEAHEISKEHMQPTHPIR.[L] | 1xHomocysteineMMTS [K10] | 4,49774E-05 | 2524,15299 | Brain/ FAIMS/MMTS |
| P62073 | P62073 [40-53] | [K].EAELSKGESVCLDR.[C] | 1xCarbamidomethyl [C11]; 1xHomocysteineIAA [K6] | 0,000318237 | 1766,80476 | Brain/ FAIMS/IAA |
| P62141 | P62141 [143-149] | [R].FNIKLWK.[T] | 1xHomocysteineIAA [K4] | 0,00445979 | 1122,61285 | Brain/ FAIMS/IAA |
| P62196 | P62196 [214-229] | [R].VSGSELVQKFIGEGAR.[M] | 1xHomocysteineIAA [K9] | 0,000258769 | 1850,94291 | Brain/ FAIMS/IAA |
| P62242 | P62242 [129-140] | [K].LTPEEEEILNKK.[R] | 1xHomocysteineIAA [K11] | 0,00033298 | 1616,82 | Liver/ FAIMS/IAA |
| P62242 | P62242 [126-139] | [K].GAKLTPEEEEILNK.[K] | 1xHomocysteineIAA [K3] | 9,61721E-05 | 1744,87858 | Brain/ FAIMS/IAA |
| P62242 | P62242 [155-170] | [K].NAKISSLLEEQFQQGK.[L] | 1xHomocysteineMMTS [K3] | 4,49774E-05 | 1982,96741 | Liver/ FAIMS/MMTS |
| P62242 | P62242 [126-139] | [K].GAKLTPEEEEILNK.[K] | 1xHomocysteineMMTS [K3] | 0,00461842 | 1733,84483 | Liver/ FAIMS/MMTS |
| P62245 | P62245 [79-88] | [R].FDVQLKDLEK.[W] | 1xHomocysteineIAA [K6] | 9,61721E-05 | 1408,71408 | Liver/ FAIMS/IAA |
| P62245 | P62245 [79-88] | [R].FDVQLKDLEK.[W] | 1xHomocysteineIAA [K6] | 9,61721E-05 | 1408,71408 | Brain/ FAIMS/IAA |
| P62245 | P62245 [79-88] | [R].FDVQLKDLEK.[W] | 1xHomocysteineMMTS [K6] | 0,0004709 | 1397,68033 | Liver/ No FAIMS/MMTS |
| P62245 | P62245 [58-68] | [R].AGKIVVNLTGR.[L] | 1xHomocysteineMMTS [K3] | 0,00832764 | 1290,70207 | Liver/ No FAIMS/MMTS |
| P62245 | P62245 [79-88] | [R].FDVQLKDLEK.[W] | 1xHomocysteineMMTS [K6] | 0,0004709 | 1397,68033 | Liver/ FAIMS/MMTS |
| P62245 | P62245 [79-88] | [R].FDVQLKDLEK.[W] | 1xHomocysteineMMTS [K6] | 0,0004709 | 1397,68033 | Brain/ FAIMS/MMTS |
| P62259 | P62259 [95-118] | [K].LICCDILDVLDKHLIPAANTGESK.[V] | 2xCarbamidomethyl [C3; C4]; 1xHomocysteineIAA [K12] | 4,32935E-05 | 2869,42565 | Liver/ FAIMS/IAA |
| P62259 | P62259 [142-153] | [R].KEAAENSLVAYK.[A] | 1xHomocysteineIAA [K1] | 4,32935E-05 | 1496,74135 | Brain/ No FAIMS/IAA |
| P62259 | P62259 [142-153] | [R].KEAAENSLVAYK.[A] | 1xHomocysteineIAA [K1] | 4,32935E-05 | 1496,74135 | Brain/ FAIMS/IAA |
| P62259 | P62259 [43-56] | [R].NLLSVAYKNVIGAR.[R] | 1xHomocysteineMMTS [K8] | 4,49774E-05 | 1680,89239 | Liver/ No FAIMS/MMTS |
| P62259 | P62259 [107-123] | [K].HLIPAANTGESKVFYYK.[M] | 1xHomocysteineMMTS [K12] | 4,49774E-05 | 2101,02453 | Liver/ No FAIMS/MMTS |
| P62259 | P62259 [142-153] | [R].KEAAENSLVAYK.[A] | 1xHomocysteineMMTS [K1] | 4,49774E-05 | 1485,70761 | Liver/ No FAIMS/MMTS |
| P62259 | P62259 [142-153] | [R].KEAAENSLVAYK.[A] | 1xHomocysteineMMTS [K1] | 4,49774E-05 | 1485,70761 | Liver/ FAIMS/MMTS |
| P62259 | P62259 [29-42] | [K].KVAGMDVELTVEER.[N] | 1xOxidation [M5]; 1xHomocysteineMMTS [K1] | 0,00230389 | 1754,81215 | Liver/ FAIMS/MMTS |

| | | | | | | |
|---|---|---|---|---|---|---|
| P62259 | P62259 [43-56] | [R].NLLSVAYKNVIGAR.[R] | 1xHomocysteineMMTS [K8] | 4,49774E-05 | 1680,89239 | Liver/ FAIMS/MMTS |
| P62259 | P62259 [62-78] | [R].IISSIEQKEENKGGEDK.[L] | 1xHomocysteineMMTS [K] | 4,49774E-05 | 2066,97328 | Liver/ FAIMS/MMTS |
| P62259 | P62259 [43-56] | [R].NLLSVAYKNVIGAR.[R] | 1xHomocysteineMMTS [K8] | 4,49774E-05 | 1680,89239 | Brain/ No FAIMS/MMTS |
| P62259 | P62259 [107-123] | [K].HLIPAANTGESKVFYYK.[M] | 1xHomocysteineMMTS [K12] | 4,49774E-05 | 2101,02453 | Brain/ No FAIMS/MMTS |
| P62259 | P62259 [142-153] | [R].KEAAENSLVAYK.[A] | 1xHomocysteineMMTS [K1] | 4,49774E-05 | 1485,70761 | Brain/ No FAIMS/MMTS |
| P62259 | P62259 [1-19] | [-].MDDREDLVYQAKLAEQAER.[Y] | 1xAcetyl [N-Term]; 1xHomocysteineMMTS [K12] | 4,49774E-05 | 2485,1156 | Brain/ No FAIMS/MMTS |
| P62259 | P62259 [20-29] | [R].YDEMVESMKK.[V] | 1xHomocysteineMMTS [K9] | 0,000192903 | 1422,57719 | Brain/ FAIMS/MMTS |
| P62259 | P62259 [43-56] | [R].NLLSVAYKNVIGAR.[R] | 1xHomocysteineMMTS [K8] | 4,49774E-05 | 1680,89239 | Brain/ FAIMS/MMTS |
| P62259 | P62259 [107-123] | [K].HLIPAANTGESKVFYYK.[M] | 1xHomocysteineMMTS [K12] | 4,49774E-05 | 2101,02453 | Brain/ FAIMS/MMTS |
| P62259 | P62259 [142-153] | [R].KEAAENSLVAYK.[A] | 1xHomocysteineMMTS [K1] | 4,49774E-05 | 1485,70761 | Brain/ FAIMS/MMTS |
| P62259 | P62259 [62-78] | [R].IISSIEQKEENKGGEDK.[L] | 1xHomocysteineMMTS [K] | 4,49774E-05 | 2066,97328 | Brain/ FAIMS/MMTS |
| P62259 | P62259 [1-19] | [-].MDDREDLVYQAKLAEQAER.[Y] | 1xAcetyl [N-Term]; 1xHomocysteineMMTS [K12] | 4,49774E-05 | 2485,1156 | Brain/ FAIMS/MMTS |
| P62267 | P62267 [108-119] | [R].KGHAVGDIPGVR.[F] | 1xHomocysteineIAA [K1] | 0,00332757 | 1379,72123 | Brain/ FAIMS/IAA |
| P62267 | P62267 [49-60] | [K].GIVLEKVGVEAK.[Q] | 1xHomocysteineMMTS [K6] | 0,000738471 | 1404,75892 | Liver/ No FAIMS/MMTS |
| P62267 | P62267 [38-54] | [K].ANPFGGASHAKGIVLEK.[V] | 1xHomocysteineMMTS [K11] | 0,000829616 | 1858,93023 | Liver/ FAIMS/MMTS |
| P62270 | P62270 [76-86] | [R].QYKIPDWFLNR.[Q] | 1xHomocysteineMMTS [K3] | 0,00107839 | 1642,78686 | Liver/ FAIMS/MMTS |
| P62274 | P62274 [45-54] | [R].QYAKDIGFIK.[L] | 1xHomocysteineMMTS [K4] | 0,00782025 | 1345,66429 | Liver/ FAIMS/MMTS |
| P62281 | P62281 [23-32] | [R].VLLGETGKEK.[L] | 1xHomocysteineIAA [K8] | 6,30912E-05 | 1247,6664 | Brain/ FAIMS/IAA |
| P62281 | P62281 [36-45] | [R].YYKNIGLGFK.[T] | 1xHomocysteineMMTS [K3] | 0,000564742 | 1365,66938 | Liver/ FAIMS/MMTS |
| P62301 | P62301 [65-73] | [R].FVTGNKILR.[I] | 1xHomocysteineIAA [K6] | 0,000177335 | 1221,67724 | Brain/ FAIMS/IAA |
| P62301 | P62301 [79-94] | [K].GLAPDLPEDLYHLIKK.[A] | 1xHomocysteineMMTS [K15] | 0,00399941 | 1985,02347 | Liver/ FAIMS/MMTS |
| P62301 | P62301 [28-39] | [K].LTSDDVKEQIYK.[L] | 1xHomocysteineMMTS [K7] | 0,00631419 | 1601,75496 | Liver/ FAIMS/MMTS |
| P62305 | P62305 [10-23] | [K].VQKVMVQPINLIFR.[Y] | 1xHomocysteineIAA [K3] | 0,000800823 | 1859,03939 | Brain/ FAIMS/IAA |
| P62317 | P62317 [1-18] | [-].MSLLNKPKSEMTPEELQK.[R] | 1xMet-loss+Acetyl [N-Term]; 1xHomocysteineIAA [K8] | 0,000258769 | 2188,09882 | Brain/ FAIMS/IAA |
| P62331 | P62331 [56-69] | [K].NVKFNVWDVGGQDK.[I] | 1xHomocysteineIAA [K3] | 6,30912E-05 | 1779,84828 | Brain/ FAIMS/IAA |
| P62484; Q8CBW3 | P62484 [108-120]; Q8CBW3 [108-120] | [R].EIGILTTNKNTSR.[T] | 1xHomocysteineIAA [K9] | 4,32935E-05 | 1620,83738 | Brain/ FAIMS/IAA |
| P62627 | P62627 [1-10] | [-].MAEVEETLKR.[L] | 1xMet-loss+Acetyl [N-Term]; 1xHomocysteineIAA [K9] | 9,61721E-05 | 1290,63583 | Brain/ No FAIMS/IAA |
| P62627 | P62627 [1-10] | [-].MAEVEETLKR.[L] | 1xMet-loss+Acetyl [N-Term]; 1xHomocysteineIAA [K9] | 9,61721E-05 | 1290,63583 | Brain/ FAIMS/IAA |
| P62631 | P62631 [147-165] | [K].QLIVGVNKMDSTEPAYSEK.[R] | 1xHomocysteineIAA [K8] | 4,32935E-05 | 2283,09954 | Brain/ FAIMS/IAA |
| P62702 | P62702 [101-108] | [R].LIYDTKGR.[F] | 1xHomocysteineIAA [K6] | 9,61721E-05 | 1139,58775 | Liver/ FAIMS/IAA |
| P62702 | P62702 [231-240] | [K].GNKPWISLPR.[G] | 1xHomocysteineMMTS [K3] | 0,00191513 | 1330,67586 | Liver/ No FAIMS/MMTS |
| P62702 | P62702 [114-122] | [R].ITPEEAKYK.[L] | 1xHomocysteineMMTS [K7] | 0,00166208 | 1241,59046 | Liver/ FAIMS/MMTS |
| P62715; P63330 | P62715 [71-89]; P63330 [71-89] | [R].IGGKSPDTNYLFMGDYVDR.[G] | 1xHomocysteineIAA [K4] | 4,32935E-05 | 2322,05293 | Brain/ FAIMS/IAA |
| P62717 | P62717 [75-83] | [R].VKNFGIWLR.[Y] | 1xHomocysteineIAA [K2] | 0,000741565 | 1306,70887 | Brain/ FAIMS/IAA |
| P62737; P68134 | P62737 [216-240]; P68134 [216-240] | [K].EKLCYVALDFENEMATAASSSSLEK.[S] | 1xCarbamidomethyl [C4]; 1xHomocysteineIAA [K2] | 4,32935E-05 | 2967,34204 | Liver/ FAIMS/IAA |
| P62737; P68134 | P62737 [64-86]; P68134 [64-86] | [K].RGILTLKYPIEHGIITNWDDMEK.[I] | 1xOxidation [M21]; 1xHomocysteineIAA [K7] | 4,32935E-05 | 2932,46956 | Brain/ FAIMS/IAA |
| P62737; P68134 | P62737 [65-86]; P68134 [65-86] | [R].GILTLKYPIEHGIITNWDDMEK.[I] | 1xOxidation [M20]; 1xHomocysteineMMTS [K6] | 4,49774E-05 | 2765,33471 | Liver/ No FAIMS/MMTS |
| P62737; P68134 | P62737 [98-118]; P68134 [98-118] | [R].VAPEEHPTLLTEAPLNPKANR.[E] | 1xHomocysteineMMTS [K18] | 4,49774E-05 | 2460,23737 | Liver/ No FAIMS/MMTS |
| P62737; P68134 | P62737 [65-86]; P68134 [65-86] | [R].GILTLKYPIEHGIITNWDDMEK.[I] | 1xOxidation [M20]; 1xHomocysteineMMTS [K6] | 4,49774E-05 | 2765,33471 | Liver/ FAIMS/MMTS |
| P62737; P68134 | P62737 [65-86]; P68134 [65-86] | [R].GILTLKYPIEHGIITNWDDMEK.[I] | 1xHomocysteineMMTS [K6] | 4,49774E-05 | 2749,33979 | Liver/ FAIMS/MMTS |
| P62737; P68134 | P62737 [98-118]; P68134 [98-118] | [R].VAPEEHPTLLTEAPLNPKANR.[E] | 1xHomocysteineMMTS [K18] | 4,49774E-05 | 2460,23737 | Liver/ FAIMS/MMTS |
| P62737; P68134 | P62737 [65-86]; P68134 [65-86] | [R].GILTLKYPIEHGIITNWDDMEK.[I] | 1xHomocysteineMMTS [K6] | 4,49774E-05 | 2749,33979 | Brain/ No FAIMS/MMTS |
| P62737; P68134 | P62737 [98-118]; P68134 [98-118] | [R].VAPEEHPTLLTEAPLNPKANR.[E] | 1xHomocysteineMMTS [K18] | 4,49774E-05 | 2460,23737 | Brain/ No FAIMS/MMTS |
| P62737; P68134 | P62737 [65-86]; P68134 [65-86] | [R].GILTLKYPIEHGIITNWDDMEK.[I] | 1xOxidation [M20]; 1xHomocysteineMMTS [K6] | 4,49774E-05 | 2765,33471 | Brain/ FAIMS/MMTS |
| P62737; P68134 | P62737 [65-86]; P68134 [65-86] | [R].GILTLKYPIEHGIITNWDDMEK.[I] | 1xHomocysteineMMTS [K6] | 4,49774E-05 | 2749,33979 | Brain/ FAIMS/MMTS |
| P62746 | P62746 [151-164] | [R].IQAYDYLECSAKTK.[E] | 1xCarbamidomethyl [C9]; 1xHomocysteineIAA [K12] | 4,32935E-05 | 1863,86155 | Brain/ FAIMS/IAA |
| P62748 | P62748 [155-171] | [R].QMDTNNDGKLSLEEFIK.[G] | 1xHomocysteineIAA [K9] | 4,32935E-05 | 2155,99983 | Brain/ FAIMS/IAA |
| P62748 | P62748 [164-174] | [K].LSLEEFIKGAK.[S] | 1xHomocysteineMMTS [K8] | 0,000507453 | 1397,71672 | Brain/ FAIMS/MMTS |
| P62751 | P62751 [79-89] | [K].FPLTTESAMKK.[I] | 1xHomocysteineIAA [K10] | 0,00219809 | 1426,70688 | Liver/ FAIMS/IAA |
| P62751 | P62751 [79-89] | [K].FPLTTESAMKK.[I] | 1xHomocysteineIAA [K10] | 0,00219809 | 1426,70688 | Brain/ FAIMS/IAA |
| P62751 | P62751 [8-18] | [K].EAPAPPKAEAK.[A] | 1xHomocysteineMMTS [K7] | 0,000530332 | 1271,61225 | Liver/ No FAIMS/MMTS |
| P62751 | P62751 [8-18] | [K].EAPAPPKAEAK.[A] | 1xHomocysteineMMTS [K7] | 0,000530332 | 1271,61225 | Liver/ FAIMS/MMTS |
| P62754 | P62754 [32-51] | [R].MATEVAADALGEEWKGYVVR.[I] | 1xHomocysteineIAA [K15] | 4,32935E-05 | 2369,12643 | Brain/ No FAIMS/IAA |
| P62754 | P62754 [32-51] | [R].MATEVAADALGEEWKGYVVR.[I] | 1xHomocysteineIAA [K15] | 4,32935E-05 | 2369,12643 | Brain/ FAIMS/IAA |
| P62761 | P62761 [4-18] | [K].QNSKLAPEVMEDLVK.[S] | 1xHomocysteineIAA [K4] | 4,32935E-05 | 1874,93505 | Brain/ FAIMS/IAA |
| P62761 | P62761 [4-18] | [K].QNSKLAPEVMEDLVK.[S] | 1xHomocysteineMMTS [K4] | 4,49774E-05 | 1863,9013 | Brain/ FAIMS/MMTS |
| P62761 | P62761 [161-172] | [K].NKDDQITLDEFK.[E] | 1xHomocysteineMMTS [K2] | 0,00277223 | 1628,72947 | Brain/ FAIMS/MMTS |
| P62761 | P62761 [8-28] | [K].LAPEVMEDLVKSTEFNEHELK.[Q] | 1xHomocysteineMMTS [K11] | 4,49774E-05 | 2621,22957 | Brain/ FAIMS/MMTS |
| P62761 | P62761 [19-32] | [K].STEFNEHELKQWYK.[G] | 1xHomocysteineMMTS [K10] | 4,49774E-05 | 2001,88334 | Brain/ FAIMS/MMTS |
| P62774 | P62774 [86-94] | [K].LLLSKGADK.[T] | 1xHomocysteineIAA [K5] | 0,000289025 | 1118,62381 | Liver/ FAIMS/IAA |
| P62774 | P62774 [95-114] | [K].TVKGPDGLTALEATDNQAIK.[A] | 1xHomocysteineIAA [K3] | 9,61721E-05 | 2216,12272 | Brain/ No FAIMS/IAA |
| P62806 | P62806 [57-68] | [R].GVLKVFLENVIR.[D] | 1xHomocysteineIAA [K4] | 4,32935E-05 | 1560,89304 | Liver/ FAIMS/IAA |
| P62806 | P62806 [79-92] | [K].RKTVTAMDVVYALK.[R] | 1xHomocysteineIAA [K2] | 4,32935E-05 | 1768,94482 | Brain/ No FAIMS/IAA |

| | | | | | | |
|---|---|---|---|---|---|---|
| P62806 | P62806 [79-92] | [K].RKTVTAMDVVYALK.[R] | 1xHomocysteineIAA [K2] | 4,32935E-05 | 1768,94482 | Brain/ FAIMS/IAA |
| P62806 | P62806 [81-93] | [K].TVTAMDVVYALKR.[Q] | 1xHomocysteineMMTS [K12] | 4,49774E-05 | 1629,81612 | Liver/ No FAIMS/MMTS |
| P62806 | P62806 [69-79] | [R].DAVTYTEHAKR.[K] | 1xHomocysteineMMTS [K10] | 0,000764279 | 1453,65624 | Liver/ No FAIMS/MMTS |
| P62806 | P62806 [81-93] | [K].TVTAMDVVYALKR.[Q] | 1xOxidation [M5]; 1xHomocysteineMMTS [K12] | 0,0004709 | 1645,81103 | Liver/ FAIMS/MMTS |
| P62806 | P62806 [25-36] | [R].DNIQGITKPAIR.[R] | 1xHomocysteineMMTS [K8] | 4,49774E-05 | 1488,76613 | Liver/ FAIMS/MMTS |
| P62806 | P62806 [81-93] | [K].TVTAMDVVYALKR.[Q] | 1xHomocysteineMMTS [K12] | 4,49774E-05 | 1629,81612 | Liver/ FAIMS/MMTS |
| P62806 | P62806 [25-36] | [R].DNIQGITKPAIR.[R] | 1xHomocysteineMMTS [K8] | 4,49774E-05 | 1488,76613 | Brain/ No FAIMS/MMTS |
| P62806 | P62806 [81-93] | [K].TVTAMDVVYALKR.[Q] | 1xHomocysteineMMTS [K12] | 4,49774E-05 | 1629,81612 | Brain/ No FAIMS/MMTS |
| P62806 | P62806 [69-79] | [R].DAVTYTEHAKR.[K] | 1xHomocysteineMMTS [K10] | 0,000764279 | 1453,65624 | Brain/ No FAIMS/MMTS |
| P62806 | P62806 [80-92] | [R].KTVTAMDVVYALK.[R] | 1xHomocysteineMMTS [K1] | 0,00074635 | 1601,80997 | Brain/ No FAIMS/MMTS |
| P62806 | P62806 [25-36] | [R].DNIQGITKPAIR.[R] | 1xHomocysteineMMTS [K8] | 4,49774E-05 | 1488,76613 | Brain/ FAIMS/MMTS |
| P62806 | P62806 [81-93] | [K].TVTAMDVVYALKR.[Q] | 1xHomocysteineMMTS [K12] | 4,49774E-05 | 1629,81612 | Brain/ FAIMS/MMTS |
| P62806 | P62806 [69-79] | [R].DAVTYTEHAKR.[K] | 1xHomocysteineMMTS [K10] | 0,000764279 | 1453,65624 | Brain/ FAIMS/MMTS |
| P62814 | P62814 [413-430] | [R].KDHADVSNQLYACYAIGK.[D] | 1xCarbamidomethyl [C13]; 1xHomocysteineIAA [K1] | 4,32935E-05 | 2227,02705 | Brain/ FAIMS/IAA |
| P62814 | P62814 [458-471] | [K].FEKNFITQGPYENR.[T] | 1xHomocysteineMMTS [K3] | 0,000231655 | 1905,86221 | Brain/ FAIMS/MMTS |
| P62821 | P62821 [120-131] | [K].LLVGNKCDLTTK.[K] | 1xCarbamidomethyl [C7]; 1xHomocysteineIAA [K6] | 4,32935E-05 | 1535,79201 | Brain/ No FAIMS/IAA |
| P62821 | P62821 [120-131] | [K].LLVGNKCDLTTK.[K] | 1xCarbamidomethyl [C7]; 1xHomocysteineIAA [K6] | 4,32935E-05 | 1535,79201 | Brain/ FAIMS/IAA |
| P62821; Q9D1G1 | P62821 [14-30]; Q9D1G1 [11-27] | [K].LLLIGDSGVGKSCLLLR.[F] | 1xCarbamidomethyl [C13]; 1xHomocysteineIAA [K11] | 4,32935E-05 | 1988,10311 | Brain/ No FAIMS/IAA |
| P62821; Q9D1G1 | P62821 [14-30]; Q9D1G1 [11-27] | [K].LLLIGDSGVGKSCLLLR.[F] | 1xCarbamidomethyl [C13]; 1xHomocysteineIAA [K11] | 4,32935E-05 | 1988,10311 | Brain/ FAIMS/IAA |
| P62821; Q9D1G1 | P62821 [59-72]; Q9D1G1 [56-69] | [K].TIKLQIWDTAGQER.[F] | 1xHomocysteineMMTS [K3] | 0,00110986 | 1821,8986 | Brain/ FAIMS/MMTS |
| P62823 | P62823 [17-32] | [R].FGQKDSSDQNFDYMFK.[L] | 1xHomocysteineIAA [K4] | 0,000932144 | 2130,88955 | Brain/ FAIMS/IAA |
| P62827 | P62827 [57-71] | [R].GPIKFNVWDTAGQEK.[F] | 1xHomocysteineIAA [K4] | 4,32935E-05 | 1863,90579 | Brain/ FAIMS/IAA |
| P62827 | P62827 [57-71] | [R].GPIKFNVWDTAGQEK.[F] | 1xHomocysteineMMTS [K4] | 0,0004709 | 1852,87205 | Liver/ FAIMS/MMTS |
| P62827 | P62827 [57-71] | [R].GPIKFNVWDTAGQEK.[F] | 1xHomocysteineMMTS [K4] | 0,0004709 | 1852,87205 | Brain/ FAIMS/MMTS |
| P62830 | P62830 [52-67] | [R].LPAAGVGDMVMATVKK.[G] | 1xHomocysteineIAA [K15] | 4,32935E-05 | 1761,90599 | Liver/ FAIMS/IAA |
| P62830 | P62830 [52-67] | [R].LPAAGVGDMVMATVKK.[G] | 1xHomocysteineIAA [K15] | 4,32935E-05 | 1761,90599 | Brain/ FAIMS/IAA |
| P62830 | P62830 [75-83] | [K].KVHPAVVIR.[Q] | 1xHomocysteineMMTS [K1] | 0,000133296 | 1181,66457 | Liver/ No FAIMS/MMTS |
| P62830 | P62830 [52-67] | [R].LPAAGVGDMVMATVKK.[G] | 1xHomocysteineMMTS [K15] | 0,00194807 | 1750,87225 | Liver/ FAIMS/MMTS |
| P62830 | P62830 [75-83] | [K].KVHPAVVIR.[Q] | 1xHomocysteineMMTS [K1] | 0,000133296 | 1181,66457 | Liver/ FAIMS/MMTS |
| P62830 | P62830 [52-67] | [R].LPAAGVGDMVMATVKK.[G] | 1xHomocysteineMMTS [K15] | 0,00194807 | 1750,87225 | Brain/ FAIMS/MMTS |
| P62843 | P62843 [1-8] | [-].MAEVEQKK.[K] | 1xMet-loss+Acetyl [N-Term]; 1xHomocysteineMMTS [K7] | 0,000781536 | 1036,48018 | Liver/ FAIMS/MMTS |
| P62849 | P62849 [33-41] | [K].ATVPKTEIR.[E] | 1xHomocysteineIAA [K5] | 0,000538793 | 1188,64052 | Liver/ No FAIMS/IAA |
| P62849 | P62849 [33-41] | [K].ATVPKTEIR.[E] | 1xHomocysteineIAA [K5] | 0,000538793 | 1188,64052 | Liver/ FAIMS/IAA |
| P62849 | P62849 [33-41] | [K].ATVPKTEIR.[E] | 1xHomocysteineMMTS [K5] | 0,00545792 | 1177,60678 | Liver/ No FAIMS/MMTS |
| P62849 | P62849 [69-84] | [K].TTGFGMIYDSLDYAKK.[N] | 1xHomocysteineMMTS [K15] | 6,70238E-05 | 1972,88532 | Liver/ FAIMS/MMTS |
| P62852 | P62852 [61-76] | [K].EVPNYKLITPAVVSER.[L] | 1xHomocysteineIAA [K6] | 0,00012645 | 1989,04737 | Brain/ FAIMS/IAA |
| P62852 | P62852 [86-98] | [R].AALQELLSKGLIK.[L] | 1xHomocysteineMMTS [K9] | 4,49774E-05 | 1546,86953 | Liver/ No FAIMS/MMTS |
| P62852 | P62852 [112-125] | [R].NTKGGDAPAAGEDA.[-] | 1xHomocysteineMMTS [K3] | 0,00663464 | 1436,57805 | Liver/ No FAIMS/MMTS |
| P62852 | P62852 [86-98] | [R].AALQELLSKGLIK.[L] | 1xHomocysteineMMTS [K9] | 4,49774E-05 | 1546,86953 | Liver/ FAIMS/MMTS |
| P62852 | P62852 [61-76] | [K].EVPNYKLITPAVVSER.[L] | 1xHomocysteineMMTS [K6] | 0,00466355 | 1978,01363 | Liver/ FAIMS/MMTS |
| P62852 | P62852 [112-125] | [R].NTKGGDAPAAGEDA.[-] | 1xHomocysteineMMTS [K3] | 0,00663464 | 1436,57805 | Liver/ FAIMS/MMTS |
| P62852 | P62852 [42-52] | [R].DKLNNLVLFDK.[A] | 1xHomocysteineMMTS [K2] | 0,008514 | 1481,74908 | Liver/ FAIMS/MMTS |
| P62858 | P62858 [6-13] | [R].VQPIKLAR.[V] | 1xHomocysteineIAA [K5] | 0,000584831 | 1098,64521 | Liver/ FAIMS/IAA |
| P62858 | P62858 [6-13] | [R].VQPIKLAR.[V] | 1xHomocysteineIAA [K5] | 0,000584831 | 1098,64521 | Brain/ FAIMS/IAA |
| P62862 | P62862 [42-52] | [R].FVNVVPTFGKK.[K] | 1xHomocysteineMMTS [K10] | 0,000599431 | 1398,72722 | Liver/ No FAIMS/MMTS |
| P62862 | P62862 [42-52] | [R].FVNVVPTFGKK.[K] | 1xHomocysteineMMTS [K10] | 0,000599431 | 1398,72722 | Liver/ FAIMS/MMTS |
| P62874 | P62874 [9-19] | [R].QEAEQLKNQIR.[D] | 1xHomocysteineIAA [K7] | 4,32935E-05 | 1530,7693 | Brain/ FAIMS/IAA |
| P62874 | P62874 [9-19] | [R].QEAEQLKNQIR.[D] | 1xHomocysteineMMTS [K7] | 0,000929608 | 1519,73556 | Brain/ FAIMS/MMTS |
| P62874; P62880 | P62874 [79-96]; P62880 [79-96] | [K].LIIWDSYTTNKVHAIPLR.[S] | 1xHomocysteineIAA [K11] | 4,32935E-05 | 2314,23763 | Brain/ No FAIMS/IAA |
| P62874; P62880 | P62874 [79-96]; P62880 [79-96] | [K].LIIWDSYTTNKVHAIPLR.[S] | 1xHomocysteineIAA [K11] | 4,32935E-05 | 2314,23763 | Brain/ FAIMS/IAA |
| P62878 | P62878 [1-20] | [-].MAAAMDVDTPSGTNSGAGKK.[R] | 1xMet-loss+Acetyl [N-Term]; 1xHomocysteineIAA [K20] | 4,32935E-05 | 1993,89536 | Brain/ FAIMS/IAA |
| P62880 | P62880 [23-42] | [R].KACGDSTLTQITAGLDPVGR.[I] | 1xCarbamidomethyl [C3]; 1xHomocysteineIAA [K1] | 4,32935E-05 | 2234,09038 | Brain/ No FAIMS/IAA |
| P62889 | P62889 [27-36] | [K].YVLGYKQTLK.[M] | 1xHomocysteineIAA [K6] | 0,0035595 | 1386,74498 | Liver/ FAIMS/IAA |
| P62889 | P62889 [43-56] | [K].AKLVILANNCPALR.[K] | 1xCarbamidomethyl [C10]; 1xHomocysteineIAA [K2] | 0,00012645 | 1726,94549 | Brain/ FAIMS/IAA |
| P62897 | P62897 [40-54] | [R].KTGQAAGFSYTDANK.[N] | 1xHomocysteineIAA [K1] | 4,32935E-05 | 1732,79591 | Brain/ No FAIMS/IAA |
| P62897 | P62897 [40-54] | [R].KTGQAAGFSYTDANK.[N] | 1xHomocysteineIAA [K1] | 4,32935E-05 | 1732,79591 | Brain/ FAIMS/IAA |
| P62897 | P62897 [93-101] | [R].ADLIAYLKK.[A] | 1xHomocysteineMMTS [K8] | 0,0031519 | 1197,63701 | Liver/ No FAIMS/MMTS |
| P62897 | P62897 [93-101] | [R].ADLIAYLKK.[A] | 1xHomocysteineMMTS [K8] | 0,0031519 | 1197,63701 | Liver/ FAIMS/MMTS |
| P62897 | P62897 [27-39] | [K].HKTGPNLHGLFGR.[K] | 1xHomocysteineMMTS [K2] | 4,49774E-05 | 1596,7886 | Liver/ FAIMS/MMTS |
| P62897 | P62897 [40-54] | [R].KTGQAAGFSYTDANK.[N] | 1xHomocysteineMMTS [K1] | 4,49774E-05 | 1721,76217 | Liver/ FAIMS/MMTS |
| P62897 | P62897 [40-54] | [R].KTGQAAGFSYTDANK.[N] | 1xHomocysteineMMTS [K1] | 4,49774E-05 | 1721,76217 | Brain/ FAIMS/MMTS |
| P62900 | P62900 [52-63] | [K].FAMKEMGTPDVR.[I] | 1xHomocysteineMMTS [K4] | 0,000530332 | 1544,67282 | Liver/ FAIMS/MMTS |
| P62908 | P62908 [11-27] | [K].FVADGIFKAELNEFLTR.[E] | 1xHomocysteineIAA [K8] | 9,61721E-05 | 2144,08449 | Brain/ FAIMS/IAA |
| P62911 | P62911 [94-108] | [K].SYCAEIAHNVSSKNR.[K] | 1xCarbamidomethyl [C3]; 1xHomocysteineIAA [K13] | 0,000960518 | 1909,86434 | Brain/ FAIMS/IAA |
| P62918 | P62918 [129-145] | [R].ASGNYATVISHNPETKK.[T] | 1xHomocysteineIAA [K16] | 0,000595249 | 1990,9651 | Liver/ FAIMS/IAA |
| P62918 | P62918 [55-64] | [R].GAPLAKVVFR.[D] | 1xHomocysteineIAA [K6] | 4,32935E-05 | 1231,69797 | Brain/ No FAIMS/IAA |
| P62918 | P62918 [55-64] | [R].GAPLAKVVFR.[D] | 1xHomocysteineIAA [K6] | 4,32935E-05 | 1231,69797 | Brain/ FAIMS/IAA |
| P62918 | P62918 [55-64] | [R].GAPLAKVVFR.[D] | 1xHomocysteineMMTS [K6] | 6,70238E-05 | 1220,66423 | Liver/ No FAIMS/MMTS |

| | | | | | | |
|---|---|---|---|---|---|---|
| P62918 | P62918 [43-54] | [K].GIVKDIIHDPGR.[G] | 1xHomocysteineMMTS [K4] | 0,000266738 | 1482,75556 | Liver/ No FAIMS/MMTS |
| P62918 | P62918 [55-64] | [R].GAPLAKVVFR.[D] | 1xHomocysteineMMTS [K6] | 6,70238E-05 | 1220,66423 | Liver/ FAIMS/MMTS |
| P62918 | P62918 [129-145] | [R].ASGNYATVISHNPETKK.[T] | 1xHomocysteineMMTS [K16] | 6,70238E-05 | 1979,93136 | Liver/ FAIMS/MMTS |
| P62960 | P62960 [255-277] | [R].EDGNEEDKENQGDETQGQQPPQR.[R] | 1xHomocysteineMMTS [K8] | 0,00818121 | 2791,1166 | Liver/ No FAIMS/MMTS |
| P62960 | P62960 [52-62] | [K].VIATKVLGTVK.[W] | 1xHomocysteineMMTS [K5] | 0,000846174 | 1291,74763 | Liver/ FAIMS/MMTS |
| P62962 | P62962 [27-54] | [K].DSPSVWAAVPGKTFVSITPAEVGVLVGK.[D] | 1xHomocysteineIAA [K12] | 4,32935E-05 | 2985,5754 | Brain/ FAIMS/IAA |
| P62962 | P62962 [39-56] | [K].TFVSITPAEVGVLVGKDR.[S] | 1xHomocysteineMMTS [K16] | 0,000211896 | 2051,06639 | Liver/ No FAIMS/MMTS |
| P62962 | P62962 [39-56] | [K].TFVSITPAEVGVLVGKDR.[S] | 1xHomocysteineMMTS [K16] | 0,000211896 | 2051,06639 | Liver/ FAIMS/MMTS |
| P62962 | P62962 [90-108] | [R].TKSTGGAPTFNVTVTMTAK.[T] | 1xHomocysteineMMTS [K2] | 0,00704418 | 2074,99699 | Liver/ FAIMS/MMTS |
| P62983 | P62983 [43-54] | [R].LIFAGKQLEDGR.[T] | 1xHomocysteineIAA [K6] | 4,32935E-05 | 1520,78897 | Liver/ FAIMS/IAA |
| P62983 | P62983 [43-54] | [R].LIFAGKQLEDGR.[T] | 1xHomocysteineIAA [K6] | 4,32935E-05 | 1520,78897 | Brain/ No FAIMS/IAA |
| P62983 | P62983 [43-54] | [R].LIFAGKQLEDGR.[T] | 1xHomocysteineIAA [K6] | 4,32935E-05 | 1520,78897 | Brain/ FAIMS/IAA |
| P62983 | P62983 [55-72] | [R].TLSDYNIQKESTLHLVLR.[L] | 1xHomocysteineMMTS [K9] | 4,49774E-05 | 2293,1679 | Liver/ No FAIMS/MMTS |
| P62983 | P62983 [43-54] | [R].LIFAGKQLEDGR.[T] | 1xHomocysteineMMTS [K6] | 4,49774E-05 | 1509,75523 | Liver/ No FAIMS/MMTS |
| P62983 | P62983 [1-11] | [-G].MQIFVKTLTGK.[T] | 1xHomocysteineMMTS [K6] | 4,49774E-05 | 1428,74116 | Liver/ No FAIMS/MMTS |
| P62983 | P62983 [7-27] | [K].TLTGKTITLEVEPSDTIENVK.[A] | 1xHomocysteineMMTS [K5] | 4,49774E-05 | 2451,2357 | Liver/ No FAIMS/MMTS |
| P62983 | P62983 [12-29] | [K].TITLEVEPSDTIENVKAK.[I] | 1xHomocysteineMMTS [K16] | 4,49774E-05 | 2150,07193 | Liver/ No FAIMS/MMTS |
| P62983 | P62983 [55-72] | [R].TLSDYNIQKESTLHLVLR.[L] | 1xHomocysteineMMTS [K9] | 4,49774E-05 | 2293,1679 | Liver/ FAIMS/MMTS |
| P62983 | P62983 [43-54] | [R].LIFAGKQLEDGR.[T] | 1xHomocysteineMMTS [K6] | 4,49774E-05 | 1509,75523 | Liver/ FAIMS/MMTS |
| P62983 | P62983 [1-11] | [-G].MQIFVKTLTGK.[T] | 1xHomocysteineMMTS [K6] | 4,49774E-05 | 1428,74116 | Liver/ FAIMS/MMTS |
| P62983 | P62983 [7-27] | [K].TLTGKTITLEVEPSDTIENVK.[A] | 1xHomocysteineMMTS [K5] | 4,49774E-05 | 2451,2357 | Liver/ FAIMS/MMTS |
| P62983 | P62983 [30-42] | [K].IQDKEGIPPDQQR.[L] | 1xHomocysteineMMTS [K4] | 0,000738471 | 1686,7938 | Liver/ FAIMS/MMTS |
| P62983 | P62983 [12-29] | [K].TITLEVEPSDTIENVKAK.[I] | 1xHomocysteineMMTS [K16] | 4,49774E-05 | 2150,07193 | Liver/ FAIMS/MMTS |
| P62983 | P62983 [43-54] | [R].LIFAGKQLEDGR.[T] | 1xHomocysteineMMTS [K6] | 4,49774E-05 | 1509,75523 | Brain/ No FAIMS/MMTS |
| P62983 | P62983 [1-11] | [-G].MQIFVKTLTGK.[T] | 1xHomocysteineMMTS [K6] | 4,49774E-05 | 1428,74116 | Brain/ No FAIMS/MMTS |
| P62983 | P62983 [7-27] | [K].TLTGKTITLEVEPSDTIENVK.[A] | 1xHomocysteineMMTS [K5] | 4,49774E-05 | 2451,2357 | Brain/ No FAIMS/MMTS |
| P62983 | P62983 [12-29] | [K].TITLEVEPSDTIENVKAK.[I] | 1xHomocysteineMMTS [K16] | 4,49774E-05 | 2150,07193 | Brain/ No FAIMS/MMTS |
| P62983 | P62983 [55-72] | [R].TLSDYNIQKESTLHLVLR.[L] | 1xHomocysteineMMTS [K9] | 4,49774E-05 | 2293,1679 | Brain/ FAIMS/MMTS |
| P62983 | P62983 [43-54] | [R].LIFAGKQLEDGR.[T] | 1xHomocysteineMMTS [K6] | 4,49774E-05 | 1509,75523 | Brain/ FAIMS/MMTS |
| P62983 | P62983 [1-11] | [-G].MQIFVKTLTGK.[T] | 1xHomocysteineMMTS [K6] | 4,49774E-05 | 1428,74116 | Brain/ FAIMS/MMTS |
| P62983 | P62983 [7-27] | [K].TLTGKTITLEVEPSDTIENVK.[A] | 1xHomocysteineMMTS [K5] | 4,49774E-05 | 2451,2357 | Brain/ FAIMS/MMTS |
| P62983 | P62983 [30-42] | [K].IQDKEGIPPDQQR.[L] | 1xHomocysteineMMTS [K4] | 0,000738471 | 1686,7938 | Brain/ FAIMS/MMTS |
| P62983 | P62983 [12-29] | [K].TITLEVEPSDTIENVKAK.[I] | 1xHomocysteineMMTS [K16] | 4,49774E-05 | 2150,07193 | Brain/ FAIMS/MMTS |
| P63001 | P63001 [148-163] | [K].EIGAVKYLECSALTQR.[G] | 1xCarbamidomethyl [C10]; 1xHomocysteineIAA [K6] | 4,32935E-05 | 2011,99396 | Brain/ FAIMS/IAA |
| P63011 | P63011 [179-202] | [R].LVDVICEKMSESLDTADPAVTGAK.[Q] | 1xCarbamidomethyl [C6]; 1xHomocysteineIAA [K8] | 4,32935E-05 | 2723,29363 | Brain/ FAIMS/IAA |
| P63011 | P63011 [9-24] | [R].YGQKESSDQNFDYMFK.[I] | 1xHomocysteineMMTS [K4] | 4,49774E-05 | 2149,86637 | Brain/ FAIMS/MMTS |
| P63011; P35276 | P63011 [25-41]; P35276 [25-41] | [K].LLLIGNSSVGKTSFLFR.[Y] | 1xHomocysteineIAA [K11] | 4,32935E-05 | 2026,11539 | Brain/ No FAIMS/IAA |
| P63011; P35276 | P63011 [25-41]; P35276 [25-41] | [K].LLLIGNSSVGKTSFLFR.[Y] | 1xHomocysteineIAA [K11] | 4,32935E-05 | 2026,11539 | Brain/ FAIMS/IAA |
| P63011; P62823 | P63011 [168-178]; P62823 [176-186] | [K].DNINVKQTFER.[L] | 1xHomocysteineIAA [K6] | 0,000273428 | 1537,74275 | Brain/ No FAIMS/IAA |
| P63011; P62823 | P63011 [168-178]; P62823 [176-186] | [K].DNINVKQTFER.[L] | 1xHomocysteineIAA [K6] | 0,000273428 | 1537,74275 | Brain/ FAIMS/IAA |
| P63017 | P63017 [584-601] | [K].NQTAEKEEFEHQQKELEK.[V] | 1xHomocysteineIAA [K14] | 4,32935E-05 | 2419,11943 | Liver/ FAIMS/IAA |
| P63017 | P63017 [417-447] | [R].NTTIPTKQTQTFTTYSDNQPGVLIQVYEGER.[A] | 1xHomocysteineIAA [K7] | 4,32935E-05 | 3703,79083 | Brain/ FAIMS/IAA |
| P63017 | P63017 [129-155] | [K].EIAEAYLGKTVTNAVVTVPAYFNDSQR.[Q] | 1xHomocysteineMMTS [K9] | 0,00239411 | 3119,51763 | Liver/ No FAIMS/MMTS |
| P63017 | P63017 [312-322] | [R].GTLDPVEKALR.[D] | 1xHomocysteineMMTS [K8] | 0,000173718 | 1361,69157 | Liver/ No FAIMS/MMTS |
| P63017 | P63017 [50-71] | [R].LIGDAAKNQVAMNPTNTVFDAK.[R] | 1xHomocysteineMMTS [K7] | 4,49774E-05 | 2481,19346 | Liver/ No FAIMS/MMTS |
| P63017 | P63017 [584-597] | [K].NQTAEKEEFEHQQK.[E] | 1xHomocysteineMMTS [K6] | 4,49774E-05 | 1908,82147 | Liver/ No FAIMS/MMTS |
| P63017 | P63017 [510-517] | [R].LSKEDIER.[M] | 1xHomocysteineMMTS [K3] | 0,000929608 | 1152,53876 | Liver/ No FAIMS/MMTS |
| P63017 | P63017 [326-342] | [K].LDKSQIHDIVLVGGSTR.[I] | 1xHomocysteineMMTS [K3] | 4,49774E-05 | 2001,02559 | Liver/ No FAIMS/MMTS |
| P63017 | P63017 [584-601] | [K].NQTAEKEEFEHQQKELEK.[V] | 1xHomocysteineMMTS [K14] | 0,00732281 | 2408,08568 | Liver/ No FAIMS/MMTS |
| P63017 | P63017 [237-247] | [R].MVNHFIAEFKR.[K] | 1xHomocysteineMMTS [K10] | 0,000282457 | 1554,73781 | Liver/ No FAIMS/MMTS |
| P63017 | P63017 [50-71] | [R].LIGDAAKNQVAMNPTNTVFDAK.[R] | 1xHomocysteineMMTS [K7] | 4,49774E-05 | 2481,19346 | Liver/ FAIMS/MMTS |
| P63017 | P63017 [103-112] | [K].VQVEYKGETK.[S] | 1xHomocysteineMMTS [K6] | 0,000530332 | 1343,63338 | Liver/ FAIMS/MMTS |
| P63017 | P63017 [536-550] | [K].VSSKNSLESYAFNMK.[A] | 1xHomocysteineMMTS [K4] | 4,49774E-05 | 1867,8387 | Liver/ FAIMS/MMTS |
| P63017 | P63017 [156-171] | [R].QATKDAGTIAGLNVLR.[I] | 1xHomocysteineMMTS [K4] | 0,00138774 | 1790,92515 | Liver/ FAIMS/MMTS |
| P63017 | P63017 [510-517] | [R].LSKEDIER.[M] | 1xHomocysteineMMTS [K3] | 0,000929608 | 1152,53876 | Liver/ FAIMS/MMTS |
| P63017 | P63017 [326-342] | [K].LDKSQIHDIVLVGGSTR.[I] | 1xHomocysteineMMTS [K3] | 4,49774E-05 | 2001,02559 | Liver/ FAIMS/MMTS |
| P63017 | P63017 [50-71] | [R].LIGDAAKNQVAMNPTNTVFDAK.[R] | 1xOxidation [M12]; 1xHomocysteineMMTS [K7] | 4,49774E-05 | 2497,18838 | Liver/ FAIMS/MMTS |
| P63017 | P63017 [312-322] | [R].GTLDPVEKALR.[D] | 1xHomocysteineMMTS [K8] | 0,000173718 | 1361,69157 | Liver/ FAIMS/MMTS |
| P63017 | P63017 [518-531] | [R].MVQEAEKYKAEDEK.[Q] | 1xHomocysteineMMTS [K7] | 0,00166208 | 1860,81763 | Liver/ FAIMS/MMTS |
| P63017 | P63017 [584-597] | [K].NQTAEKEEFEHQQK.[E] | 1xHomocysteineMMTS [K6] | 4,49774E-05 | 1908,82147 | Liver/ FAIMS/MMTS |
| P63017 | P63017 [109-126] | [K].GETKSFYPEEVSSMVLTK.[M] | 1xHomocysteineMMTS [K4] | 0,000322194 | 2195,00689 | Liver/ FAIMS/MMTS |
| P63017 | P63017 [127-137] | [K].MKEIAEAYLGK.[T] | 1xHomocysteineMMTS [K2] | 0,000282457 | 1415,67314 | Liver/ FAIMS/MMTS |
| P63017 | P63017 [57-72] | [K].NQVAMNPTNTVFDAKR.[L] | 1xHomocysteineMMTS [K15] | 4,49774E-05 | 1968,90885 | Liver/ FAIMS/MMTS |
| P63017 | P63017 [113-128] | [K].SFYPEEVSSMVLTKMK.[E] | 1xHomocysteineMMTS [K14] | 0,000152766 | 2038,93564 | Liver/ FAIMS/MMTS |
| P63017 | P63017 [57-72] | [K].NQVAMNPTNTVFDAKR.[L] | 1xHomocysteineMMTS [K15] | 4,49774E-05 | 1968,90885 | Brain/ No FAIMS/MMTS |
| P63017 | P63017 [113-128] | [K].SFYPEEVSSMVLTKMK.[E] | 1xHomocysteineMMTS [K14] | 0,000152766 | 2038,93564 | Brain/ No FAIMS/MMTS |
| P63017 | P63017 [312-322] | [R].GTLDPVEKALR.[D] | 1xHomocysteineMMTS [K8] | 0,000173718 | 1361,69157 | Brain/ FAIMS/MMTS |
| P63017 | P63017 [518-531] | [R].MVQEAEKYKAEDEK.[Q] | 1xHomocysteineMMTS [K7] | 0,00166208 | 1860,81763 | Brain/ FAIMS/MMTS |
| P63017 | P63017 [50-71] | [R].LIGDAAKNQVAMNPTNTVFDAK.[R] | 1xHomocysteineMMTS [K7] | 4,49774E-05 | 2481,19346 | Brain/ FAIMS/MMTS |
| P63017 | P63017 [103-112] | [K].VQVEYKGETK.[S] | 1xHomocysteineMMTS [K6] | 0,000530332 | 1343,63338 | Brain/ FAIMS/MMTS |
| P63017 | P63017 [534-550] | [R].DKVSSKNSLESYAFNMK.[A] | 1xHomocysteineMMTS [K6] | 0,00140305 | 2110,96061 | Brain/ FAIMS/MMTS |

| | | | | | | |
|---|---|---|---|---|---|---|
| P63017 | P63017 [584-597] | [K].NQTAEKEEFEHQQK.[E] | 1xHomocysteineMMTS [K6] | 4,49774E-05 | 1908,82147 | Brain/ FAIMS/MMTS |
| P63017 | P63017 [536-550] | [K].VSSKNSLESYAFNMK.[A] | 1xHomocysteineMMTS [K4] | 4,49774E-05 | 1867,8387 | Brain/ FAIMS/MMTS |
| P63017 | P63017 [510-517] | [R].LSKEDIER.[M] | 1xHomocysteineMMTS [K3] | 0,000929608 | 1152,53876 | Brain/ FAIMS/MMTS |
| P63017 | P63017 [326-342] | [K].LDKSQIHDIVLVGGSTR.[I] | 1xHomocysteineMMTS [K3] | 4,49774E-05 | 2001,02559 | Brain/ FAIMS/MMTS |
| P63017 | P63017 [57-72] | [K].NQVAMNPTNTVFDAKR.[L] | 1xHomocysteineMMTS [K15] | 4,49774E-05 | 1968,90885 | Brain/ FAIMS/MMTS |
| P63017 | P63017 [113-128] | [K].SFYPEEVSSMVLTKMK.[E] | 1xHomocysteineMMTS [K14] | 0,000152766 | 2038,93564 | Brain/ FAIMS/MMTS |
| P63017 | P63017 [237-247] | [R].MVNHFIAEFKR.[K] | 1xHomocysteineMMTS [K10] | 0,000282457 | 1554,73781 | Brain/ FAIMS/MMTS |
| P63017; P17156 | P63017 [498-507]; P17156 [501-510] | [K].ENKITITNDK.[G] | 1xHomocysteineIAA [K3] | 9,61721E-05 | 1349,67294 | Liver/ FAIMS/IAA |
| P63017; P17156 | P63017 [349-361]; P17156 [352-364] | [K].LLQDFFNGKELNK.[S] | 1xHomocysteineIAA [K9] | 4,32935E-05 | 1739,87852 | Brain/ FAIMS/IAA |
| P63017; P17156 | P63017 [349-361]; P17156 [352-364] | [K].LLQDFFNGKELNK.[S] | 1xHomocysteineMMTS [K9] | 4,49774E-05 | 1728,84477 | Liver/ FAIMS/MMTS |
| P63017; P17156 | P63017 [346-357]; P17156 [349-360] | [K].IQKLLQDFFNGK.[E] | 1xHomocysteineMMTS [K3] | 0,0033721 | 1613,81783 | Liver/ FAIMS/MMTS |
| P63017; P17156 | P63017 [172-188]; P17156 [173-189] | [R].IINEPTAAAIAYGLDKK.[VG] | 1xHomocysteineMMTS [K16] | 4,49774E-05 | 1951,00273 | Brain/ No FAIMS/MMTS |
| P63017; P17156 | P63017 [349-361]; P17156 [352-364] | [K].LLQDFFNGKELNK.[S] | 1xHomocysteineMMTS [K9] | 4,49774E-05 | 1728,84477 | Brain/ FAIMS/MMTS |
| P63017; P17156 | P63017 [172-188]; P17156 [173-189] | [R].IINEPTAAAIAYGLDKK.[VG] | 1xHomocysteineMMTS [K16] | 4,49774E-05 | 1951,00273 | Brain/ FAIMS/MMTS |
| P63024; P63044 | P63024 [44-70]; P63044 [57-83] | [R].DQKLSELDDRADALQAGASQFETSAAK.[L] | 1xHomocysteineIAA [K3] | 4,32935E-05 | 3039,43238 | Brain/ FAIMS/IAA |
| P63024; P63044 | P63024 [54-72]; P63044 [67-85] | [R].ADALQAGASQFETSAAKLK.[R] | 1xHomocysteineMMTS [K17] | 4,49774E-05 | 2069,99944 | Brain/ No FAIMS/MMTS |
| P63024; P63044; Q62442 | P63024 [44-53]; P63044 [57-66]; Q62442 [59-68] | [R].DQKLSELDDR.[A] | 1xHomocysteineIAA [K3] | 4,32935E-05 | 1392,64237 | Brain/ No FAIMS/IAA |
| P63024; P63044; Q62442 | P63024 [44-53]; P63044 [57-66]; Q62442 [59-68] | [R].DQKLSELDDR.[A] | 1xHomocysteineIAA [K3] | 4,32935E-05 | 1392,64237 | Brain/ FAIMS/IAA |
| P63024; P63044; Q62442 | P63024 [35-43]; P63044 [48-56]; Q62442 [50-58] | [R].VNVDKVLER.[D] | 1xHomocysteineMMTS [K5] | 0,000211896 | 1234,62824 | Brain/ No FAIMS/MMTS |
| P63024; P63044; Q62442 | P63024 [44-53]; P63044 [57-66]; Q62442 [59-68] | [R].DQKLSELDDR.[A] | 1xHomocysteineMMTS [K3] | 4,49774E-05 | 1381,60863 | Brain/ No FAIMS/MMTS |
| P63024; P63044; Q62442 | P63024 [35-43]; P63044 [48-56]; Q62442 [50-58] | [R].VNVDKVLER.[D] | 1xHomocysteineMMTS [K5] | 0,000211896 | 1234,62824 | Brain/ FAIMS/MMTS |
| P63024; P63044; Q62442 | P63024 [44-53]; P63044 [57-66]; Q62442 [59-68] | [R].DQKLSELDDR.[A] | 1xHomocysteineMMTS [K3] | 4,49774E-05 | 1381,60863 | Brain/ FAIMS/MMTS |
| P63028 | P63028 [6-21] | [R].DLISHDELFSDIYKIR.[E] | 1xHomocysteineIAA [K14] | 4,32935E-05 | 2138,05867 | Liver/ No FAIMS/IAA |
| P63028 | P63028 [6-21] | [R].DLISHDELFSDIYKIR.[E] | 1xHomocysteineIAA [K14] | 4,32935E-05 | 2138,05867 | Liver/ FAIMS/IAA |
| P63028 | P63028 [6-21] | [R].DLISHDELFSDIYKIR.[E] | 1xHomocysteineIAA [K14] | 4,32935E-05 | 2138,05867 | Brain/ No FAIMS/IAA |
| P63028 | P63028 [6-21] | [R].DLISHDELFSDIYKIR.[E] | 1xHomocysteineIAA [K14] | 4,32935E-05 | 2138,05867 | Brain/ FAIMS/IAA |
| P63028 | P63028 [101-110] | [K].GKLEEQKPER.[V] | 1xHomocysteineMMTS [K2] | 0,00980048 | 1376,66608 | Liver/ FAIMS/MMTS |
| P63028 | P63028 [6-21] | [R].DLISHDELFSDIYKIR.[E] | 1xHomocysteineMMTS [K14] | 0,00107839 | 2127,02492 | Liver/ FAIMS/MMTS |
| P63028 | P63028 [6-21] | [R].DLISHDELFSDIYKIR.[E] | 1xHomocysteineMMTS [K14] | 0,00107839 | 2127,02492 | Brain/ FAIMS/MMTS |
| P63030 | P63030 [98-109] | [R].LINYEMSKRPSA.[-] | 1xHomocysteineIAA [K8] | 0,00146914 | 1582,77161 | Brain/ FAIMS/IAA |
| P63038 | P63038 [250-268] | [K].KISSVQSIVPALEIANAHR.[K] | 1xHomocysteineIAA [K1] | 4,32935E-05 | 2207,1965 | Liver/ FAIMS/IAA |
| P63038 | P63038 [250-268] | [K].KISSVQSIVPALEIANAHR.[K] | 1xHomocysteineIAA [K1] | 4,32935E-05 | 2207,1965 | Brain/ FAIMS/IAA |
| P63038 | P63038 [345-359] | [K].VGEVIVTKDDAMLLK.[G] | 1xOxidation [M12]; 1xHomocysteineMMTS [K8] | 4,49774E-05 | 1809,91589 | Liver/ No FAIMS/MMTS |
| P63038 | P63038 [397-417] | [K].LSDGVAVLKVGGTSDVEVNEK.[K] | 1xHomocysteineMMTS [K9] | 0,00115861 | 2279,12576 | Liver/ No FAIMS/MMTS |
| P63038 | P63038 [474-493] | [K].IPAMTIAKNAGVEGSLIVEK.[I] | 1xHomocysteineMMTS [K8] | 4,49774E-05 | 2204,14874 | Liver/ No FAIMS/MMTS |
| P63038 | P63038 [345-359] | [K].VGEVIVTKDDAMLLK.[G] | 1xHomocysteineMMTS [K8] | 4,49774E-05 | 1793,92097 | Liver/ No FAIMS/MMTS |
| P63038 | P63038 [76-87] | [K].DGVTVAKSIDLK.[D] | 1xHomocysteineMMTS [K7] | 4,49774E-05 | 1408,71745 | Liver/ No FAIMS/MMTS |
| P63038 | P63038 [196-205] | [R].KGVITVKDGK.[T] | 1xHomocysteineMMTS [K7] | 0,00255629 | 1207,65373 | Liver/ No FAIMS/MMTS |
| P63038 | P63038 [463-470] | [K].IGIEIIKR.[A] | 1xHomocysteineMMTS [K7] | 0,000530332 | 1104,62678 | Liver/ No FAIMS/MMTS |
| P63038 | P63038 [353-361] | [K].DDAMLLKGK.[G] | 1xHomocysteineMMTS [K7] | 0,000530332 | 1153,5414 | Liver/ No FAIMS/MMTS |
| P63038 | P63038 [517-526] | [K].GIIDPTKVVR.[T] | 1xHomocysteineMMTS [K7] | 0,000152766 | 1260,68027 | Liver/ No FAIMS/MMTS |
| P63038 | P63038 [92-121] | [K].NIGAKLVQDVANNTNEEAGDGTTTATVLAR.[S] | 1xHomocysteineMMTS [K5] | 4,49774E-05 | 3206,54161 | Liver/ No FAIMS/MMTS |
| P63038 | P63038 [126-133] | [K].EGFEKISK.[G] | 1xHomocysteineMMTS [K5] | 0,00363897 | 1100,51148 | Liver/ No FAIMS/MMTS |
| P63038 | P63038 [122-130] | [R].SIAKEGFEK.[I] | 1xHomocysteineMMTS [K4] | 0,00252783 | 1171,54859 | Liver/ No FAIMS/MMTS |
| P63038 | P63038 [29-37] | [K].DVKFGADAR.[A] | 1xHomocysteineMMTS [K3] | 0,008514 | 1141,51288 | Liver/ No FAIMS/MMTS |
| P63038 | P63038 [471-481] | [R].ALKIPAMTIAK.[N] | 1xHomocysteineMMTS [K3] | 8,93346E-05 | 1319,72478 | Liver/ No FAIMS/MMTS |
| P63038 | P63038 [394-405] | [R].LAKLSDGVAVLK.[V] | 1xHomocysteineMMTS [K3] | 4,49774E-05 | 1376,764 | Liver/ No FAIMS/MMTS |
| P63038 | P63038 [73-82] | [K].VTKDGVTVAK.[S] | 1xHomocysteineMMTS [K3] | 4,49774E-05 | 1180,60644 | Liver/ No FAIMS/MMTS |
| P63038 | P63038 [131-141] | [K].ISKGANPVEIR.[R] | 1xHomocysteineMMTS [K3] | 0,000282457 | 1346,6919 | Liver/ No FAIMS/MMTS |
| P63038 | P63038 [371-389] | [R].IQEITEQLDITTSEYEKEK.[L] | 1xHomocysteineMMTS [K17] | 4,49774E-05 | 2460,15203 | Liver/ No FAIMS/MMTS |
| P63038 | P63038 [206-221] | [K].TLNDELEIIEGMKFDR.[G] | 1xHomocysteineMMTS [K13] | 4,49774E-05 | 2085,96536 | Liver/ No FAIMS/MMTS |
| P63038 | P63038 [406-418] | [K].VGGTSDVEVNEKK.[D] | 1xHomocysteineMMTS [K12] | 4,49774E-05 | 1524,70325 | Liver/ No FAIMS/MMTS |
| P63038 | P63038 [61-75] | [R].TVIIEQSWGSPKVTK.[D] | 1xHomocysteineMMTS [K12] | 4,49774E-05 | 1835,9394 | Liver/ No FAIMS/MMTS |
| P63038 | P63038 [181-192] | [K].DIGNIISDAMKK.[V] | 1xHomocysteineMMTS [K11] | 0,000152766 | 1467,70042 | Liver/ No FAIMS/MMTS |
| P63038 | P63038 [250-268] | [K].KISSVQSIVPALEIANAHR.[K] | 1xHomocysteineMMTS [K1] | 4,49774E-05 | 2196,16275 | Liver/ No FAIMS/MMTS |
| P63038 | P63038 [474-493] | [K].IPAMTIAKNAGVEGSLIVEK.[I] | 1xOxidation [M4]; 1xHomocysteineMMTS [K8] | 0,000946421 | 2220,14366 | Liver/ FAIMS/MMTS |
| P63038 | P63038 [345-359] | [K].VGEVIVTKDDAMLLK.[G] | 1xOxidation [M12]; 1xHomocysteineMMTS [K8] | 4,49774E-05 | 1809,91589 | Liver/ FAIMS/MMTS |

| | | | | | | |
|---|---|---|---|---|---|---|
| P63038 | P63038 [293-309] | [K].VGLQVVAVKAPGFGDNR.[K] | 1xHomocysteineMMTS [K9] | 4,49774E-05 | 1889,97243 | Liver/ FAIMS/MMTS |
| P63038 | P63038 [463-470] | [K].IGIEIIKR.[A] | 1xHomocysteineMMTS [K7] | 0,000530332 | 1104,62678 | Liver/ FAIMS/MMTS |
| P63038 | P63038 [353-361] | [K].DDAMLLKGK.[G] | 1xHomocysteineMMTS [K7] | 0,000530332 | 1153,5414 | Liver/ FAIMS/MMTS |
| P63038 | P63038 [517-526] | [K].GIIDPTKVVR.[T] | 1xHomocysteineMMTS [K7] | 0,000152766 | 1260,68027 | Liver/ FAIMS/MMTS |
| P63038 | P63038 [203-218] | [K].DGKTLNDELEIIEGMK.[F] | 1xHomocysteineMMTS [K3] | 4,49774E-05 | 1967,91226 | Liver/ FAIMS/MMTS |
| P63038 | P63038 [471-481] | [R].ALKIPAMTIAK.[N] | 1xHomocysteineMMTS [K3] | 8,93346E-05 | 1319,72478 | Liver/ FAIMS/MMTS |
| P63038 | P63038 [394-405] | [R].LAKLSDGVAVLK.[V] | 1xHomocysteineMMTS [K3] | 4,49774E-05 | 1376,764 | Liver/ FAIMS/MMTS |
| P63038 | P63038 [73-82] | [K].VTKDGVTVAK.[S] | 1xHomocysteineMMTS [K3] | 4,49774E-05 | 1180,60644 | Liver/ FAIMS/MMTS |
| P63038 | P63038 [131-141] | [K].ISKGANPVEIR.[R] | 1xHomocysteineMMTS [K3] | 0,000282457 | 1346,6919 | Liver/ FAIMS/MMTS |
| P63038 | P63038 [206-221] | [K].TLNDELEIIEGMKFDR.[G] | 1xHomocysteineMMTS [K13] | 4,49774E-05 | 2085,96536 | Liver/ FAIMS/MMTS |
| P63038 | P63038 [406-418] | [K].VGGTSDVEVNEKK.[D] | 1xHomocysteineMMTS [K12] | 4,49774E-05 | 1524,70325 | Liver/ FAIMS/MMTS |
| P63038 | P63038 [61-75] | [R].TVIIEQSWGSPKVTK.[D] | 1xHomocysteineMMTS [K12] | 4,49774E-05 | 1835,9394 | Liver/ FAIMS/MMTS |
| P63038 | P63038 [181-192] | [K].DIGNIISDAMKK.[V] | 1xHomocysteineMMTS [K11] | 0,000152766 | 1467,70042 | Liver/ FAIMS/MMTS |
| P63038 | P63038 [250-268] | [K].KISSVQSIVPALEIANAHR.[K] | 1xHomocysteineMMTS [K1] | 4,49774E-05 | 2196,16275 | Liver/ FAIMS/MMTS |
| P63038 | P63038 [206-221] | [K].TLNDELEIIEGMKFDR.[G] | 1xOxidation [M12]; 1xHomocysteineMMTS [K13] | 0,000250158 | 2101,96027 | Liver/ FAIMS/MMTS |
| P63038 | P63038 [181-192] | [K].DIGNIISDAMKK.[V] | 1xOxidation [M10]; 1xHomocysteineMMTS [K11] | 0,00651765 | 1483,69533 | Liver/ FAIMS/MMTS |
| P63038 | P63038 [345-359] | [K].VGEVIVTKDDAMLLK.[G] | 1xHomocysteineMMTS [K8] | 4,49774E-05 | 1793,92097 | Liver/ FAIMS/MMTS |
| P63038 | P63038 [76-87] | [K].DGVTVAKSIDLK.[D] | 1xHomocysteineMMTS [K7] | 4,49774E-05 | 1408,71745 | Liver/ FAIMS/MMTS |
| P63038 | P63038 [196-205] | [R].KGVITVKDGK.[T] | 1xHomocysteineMMTS [K7] | 0,00255629 | 1207,65373 | Liver/ FAIMS/MMTS |
| P63038 | P63038 [197-205] | [K].GVITVKDGK.[T] | 1xHomocysteineMMTS [K6] | 0,000173718 | 1079,55876 | Liver/ FAIMS/MMTS |
| P63038 | P63038 [92-121] | [K].NIGAKLVQDVANNTNEEAGDGTTTATVLAR.[S] | 1xHomocysteineMMTS [K5] | 4,49774E-05 | 3206,54161 | Liver/ FAIMS/MMTS |
| P63038 | P63038 [126-133] | [K].EGFEKISK.[G] | 1xHomocysteineMMTS [K5] | 0,00363897 | 1100,51148 | Liver/ FAIMS/MMTS |
| P63038 | P63038 [83-91] | [K].SIDLKDKYK.[N] | 1xHomocysteineMMTS [K5] | 0,00820834 | 1272,63266 | Liver/ FAIMS/MMTS |
| P63038 | P63038 [311-344] | [K].NQLKDMAIATGGAVFGEEGLNLNLEDVQAHDLGK.[V] | 1xHomocysteineMMTS [K4] | 6,70238E-05 | 3730,78734 | Liver/ FAIMS/MMTS |
| P63038 | P63038 [122-130] | [R].SIAKEGFEK.[I] | 1xHomocysteineMMTS [K4] | 0,00252783 | 1171,54859 | Liver/ FAIMS/MMTS |
| P63038 | P63038 [29-37] | [K].DVKFGADAR.[A] | 1xHomocysteineMMTS [K3] | 0,008514 | 1141,51288 | Liver/ FAIMS/MMTS |
| P63038 | P63038 [291-301] | [R].LKVGLQVVAVK.[A] | 1xHomocysteineMMTS [K2] | 6,70238E-05 | 1316,77926 | Liver/ FAIMS/MMTS |
| P63038 | P63038 [371-389] | [R].IQEITEQLDITTSEYEKEK.[L] | 1xHomocysteineMMTS [K17] | 4,49774E-05 | 2460,15203 | Liver/ FAIMS/MMTS |
| P63038 | P63038 [196-202] | [R].KGVITVK.[D] | 1xHomocysteineMMTS [K1] | 0,00905461 | 907,51036 | Liver/ FAIMS/MMTS |
| P63038 | P63038 [345-359] | [K].VGEVIVTKDDAMLLK.[G] | 1xHomocysteineMMTS [K8] | 4,49774E-05 | 1793,92097 | Brain/ No FAIMS/MMTS |
| P63038 | P63038 [463-470] | [K].IGIEIIKR.[A] | 1xHomocysteineMMTS [K7] | 0,000530332 | 1104,62678 | Brain/ No FAIMS/MMTS |
| P63038 | P63038 [517-526] | [K].GIIDPTKVVR.[T] | 1xHomocysteineMMTS [K7] | 0,000152766 | 1260,68027 | Brain/ No FAIMS/MMTS |
| P63038 | P63038 [471-481] | [R].ALKIPAMTIAK.[N] | 1xHomocysteineMMTS [K3] | 8,93346E-05 | 1319,72478 | Brain/ No FAIMS/MMTS |
| P63038 | P63038 [206-221] | [K].TLNDELEIIEGMKFDR.[G] | 1xHomocysteineMMTS [K13] | 4,49774E-05 | 2085,96536 | Brain/ No FAIMS/MMTS |
| P63038 | P63038 [345-359] | [K].VGEVIVTKDDAMLLK.[G] | 1xHomocysteineMMTS [K8] | 4,49774E-05 | 1793,92097 | Brain/ FAIMS/MMTS |
| P63038 | P63038 [76-87] | [K].DGVTVAKSIDLK.[D] | 1xHomocysteineMMTS [K7] | 4,49774E-05 | 1408,71745 | Brain/ FAIMS/MMTS |
| P63038 | P63038 [353-361] | [K].DDAMLLKGK.[G] | 1xHomocysteineMMTS [K7] | 0,000530332 | 1153,5414 | Brain/ FAIMS/MMTS |
| P63038 | P63038 [517-526] | [K].GIIDPTKVVR.[T] | 1xHomocysteineMMTS [K7] | 0,000152766 | 1260,68027 | Brain/ FAIMS/MMTS |
| P63038 | P63038 [122-130] | [R].SIAKEGFEK.[I] | 1xHomocysteineMMTS [K4] | 0,00252783 | 1171,54859 | Brain/ FAIMS/MMTS |
| P63038 | P63038 [203-218] | [K].DGKTLNDELEIIEGMK.[F] | 1xHomocysteineMMTS [K3] | 4,49774E-05 | 1967,91226 | Brain/ FAIMS/MMTS |
| P63038 | P63038 [471-481] | [R].ALKIPAMTIAK.[N] | 1xHomocysteineMMTS [K3] | 8,93346E-05 | 1319,72478 | Brain/ FAIMS/MMTS |
| P63038 | P63038 [73-82] | [K].VTKDGVTVAK.[S] | 1xHomocysteineMMTS [K3] | 4,49774E-05 | 1180,60644 | Brain/ FAIMS/MMTS |
| P63038 | P63038 [131-141] | [K].ISKGANPVEIR.[R] | 1xHomocysteineMMTS [K3] | 0,000282457 | 1346,6919 | Brain/ FAIMS/MMTS |
| P63038 | P63038 [206-221] | [K].TLNDELEIIEGMKFDR.[G] | 1xHomocysteineMMTS [K13] | 4,49774E-05 | 2085,96536 | Brain/ FAIMS/MMTS |
| P63038 | P63038 [406-418] | [K].VGGTSDVEVNEKK.[D] | 1xHomocysteineMMTS [K12] | 4,49774E-05 | 1524,70325 | Brain/ FAIMS/MMTS |
| P63040 | P63040 [1-14] | [-].MEFVMKQALGGATK.[D] | 1xAcetyl [N-Term]; 1xHomocysteineIAA [K6] | 4,32935E-05 | 1726,83249 | Brain/ No FAIMS/IAA |
| P63040 | P63040 [1-14] | [-].MEFVMKQALGGATK.[D] | 1xAcetyl [N-Term]; 1xHomocysteineIAA [K6] | 4,32935E-05 | 1726,83249 | Brain/ FAIMS/IAA |
| P63040 | P63040 [19-32] | [K].MLGGDEEKDPDAAK.[K] | 1xHomocysteineMMTS [K8] | 0,000599431 | 1638,6808 | Brain/ FAIMS/MMTS |
| P63054 | P63054 [25-41] | [K].KVQEEFDIDMDAPETER.[A] | 1xHomocysteineIAA [K1] | 4,32935E-05 | 2225,96892 | Brain/ No FAIMS/IAA |
| P63054 | P63054 [25-41] | [K].KVQEEFDIDMDAPETER.[A] | 1xHomocysteineIAA [K1] | 4,32935E-05 | 2225,96892 | Brain/ FAIMS/IAA |
| P63085 | P63085 [341-351] | [K].LKELIFEETAR.[F] | 1xHomocysteineIAA [K2] | 0,00251002 | 1522,79339 | Brain/ FAIMS/IAA |
| P63087; P62137 | P63087 [235-246]; P62137 [235-246] | [K].FLHKHDLDLICR.[A] | 1xCarbamidomethyl [C11]; 1xHomocysteineIAA [K4] | 0,00493325 | 1740,86724 | Brain/ FAIMS/IAA |
| P63087; P62137; P62141 | P63087 [247-261]; P62137 [247-261]; P62141 [246-260] | [R].AHQVVEDGYEFFAKR.[Q] | 1xHomocysteineIAA [K14] | 4,32935E-05 | 1969,92251 | Brain/ FAIMS/IAA |
| P63087; P62141 | P63087 [112-122]; P62141 [111-121] | [K].IKYPENFFLLR.[G] | 1xHomocysteineIAA [K2] | 9,61721E-05 | 1613,85085 | Brain/ FAIMS/IAA |
| P63094 | P63094 [202-216] | [R].VLTSGIFETKFQVDK.[V] | 1xHomocysteineIAA [K10] | 0,000258769 | 1885,97281 | Brain/ FAIMS/IAA |
| P63094; Q9DC51; Q8CGK7; P18872; P08752; B2RSH2 | P63094 [43-58]; Q9DC51 [36-51]; Q8CGK7 [45-60]; P18872 [36-51]; P08752 [36-51]; B2RSH2 [36-51] | [RK].LLLLGAGESGKSTIVK.[Q] | 1xHomocysteineIAA [K11] | 4,32935E-05 | 1759,99863 | Brain/ No FAIMS/IAA |
| P63094; Q9DC51; Q8CGK7; P18872; P08752; B2RSH2 | P63094 [43-58]; Q9DC51 [36-51]; Q8CGK7 [45-60]; P18872 [36-51]; P08752 [36-51]; B2RSH2 [36-51] | [RK].LLLLGAGESGKSTIVK.[Q] | 1xHomocysteineIAA [K11] | 4,32935E-05 | 1759,99863 | Brain/ FAIMS/IAA |
| P63101 | P63101 [140-158] | [K].GIVDQSQQAYQEAFEISKK.[E] | 1xHomocysteineIAA [K] | 4,32935E-05 | 2343,12854 | Liver/ FAIMS/IAA |
| P63101 | P63101 [139-157] | [K].KGIVDQSQQAYQEAFEISK.[K] | 1xHomocysteineIAA [K1] | 4,32935E-05 | 2343,12854 | Brain/ No FAIMS/IAA |
| P63101 | P63101 [139-157] | [K].KGIVDQSQQAYQEAFEISK.[K] | 1xHomocysteineIAA [K1] | 4,32935E-05 | 2343,12854 | Brain/ FAIMS/IAA |
| P63101 | P63101 [128-139] | [R].YLAEVAAGDDKK.[G] | 1xHomocysteineMMTS [K] | 4,49774E-05 | 1442,66541 | Liver/ No FAIMS/MMTS |
| P63101 | P63101 [139-157] | [K].KGIVDQSQQAYQEAFEISK.[K] | 1xHomocysteineMMTS [K1] | 4,49774E-05 | 2332,09479 | Liver/ FAIMS/MMTS |

| | | | | | | |
|---|---|---|---|---|---|---|
| P63101 | P63101 [61-74] | [R].VVSSIEQKTEGAEK.[K] | 1xHomocysteineMMTS [K8] | 4,49774E-05 | 1667,79788 | Liver/ FAIMS/MMTS |
| P63101 | P63101 [84-91] | [R].EKIETELR.[D] | 1xHomocysteineMMTS [K2] | 0,00264931 | 1180,57006 | Liver/ FAIMS/MMTS |
| P63101 | P63101 [140-158] | [K].GIVDQSQQAYQEAFEISKK.[E] | 1xHomocysteineMMTS [K18] | 4,49774E-05 | 2332,09479 | Liver/ FAIMS/MMTS |
| P63101 | P63101 [128-139] | [R].YLAEVAAGDDKK.[G] | 1xHomocysteineMMTS [K] | 4,49774E-05 | 1442,66541 | Liver/ FAIMS/MMTS |
| P63101 | P63101 [1-9] | [-].MDKNELVQK.[A] | 1xAcetyl [N-Term]; 1xHomocysteineMMTS [K3] | 4,49774E-05 | 1309,59489 | Liver/ FAIMS/MMTS |
| P63101 | P63101 [61-74] | [R].VVSSIEQKTEGAEK.[K] | 1xHomocysteineMMTS [K8] | 4,49774E-05 | 1667,79788 | Brain/ No FAIMS/MMTS |
| P63101 | P63101 [84-91] | [R].EKIETELR.[D] | 1xHomocysteineMMTS [K2] | 0,00264931 | 1180,57006 | Brain/ No FAIMS/MMTS |
| P63101 | P63101 [140-158] | [K].GIVDQSQQAYQEAFEISKK.[E] | 1xHomocysteineMMTS [K18] | 4,49774E-05 | 2332,09479 | Brain/ No FAIMS/MMTS |
| P63101 | P63101 [104-120] | [K].FLIPNASQPESKVFYLK.[M] | 1xHomocysteineMMTS [K12] | 4,49774E-05 | 2144,09188 | Brain/ No FAIMS/MMTS |
| P63101 | P63101 [139-157] | [K].KGIVDQSQQAYQEAFEISK.[K] | 1xHomocysteineMMTS [K1] | 4,49774E-05 | 2332,09479 | Brain/ No FAIMS/MMTS |
| P63101 | P63101 [128-139] | [R].YLAEVAAGDDKK.[G] | 1xHomocysteineMMTS [K] | 4,49774E-05 | 1442,66541 | Brain/ No FAIMS/MMTS |
| P63101 | P63101 [1-11] | [-].MDKNELVQKAK.[L] | 1xAcetyl [N-Term]; 1xHomocysteineMMTS [K9] | 4,49774E-05 | 1508,72697 | Brain/ No FAIMS/MMTS |
| P63101 | P63101 [1-9] | [-].MDKNELVQK.[A] | 1xAcetyl [N-Term]; 1xHomocysteineMMTS [K3] | 4,49774E-05 | 1309,59489 | Brain/ No FAIMS/MMTS |
| P63101 | P63101 [61-75] | [R].VVSSIEQKTEGAEKK.[Q] | 1xHomocysteineMMTS [K8] | 4,49774E-05 | 1795,89285 | Brain/ FAIMS/MMTS |
| P63101 | P63101 [61-74] | [R].VVSSIEQKTEGAEK.[K] | 1xHomocysteineMMTS [K8] | 4,49774E-05 | 1667,79788 | Brain/ FAIMS/MMTS |
| P63101 | P63101 [84-91] | [R].EKIETELR.[D] | 1xHomocysteineMMTS [K2] | 0,00264931 | 1180,57006 | Brain/ FAIMS/MMTS |
| P63101 | P63101 [140-158] | [K].GIVDQSQQAYQEAFEISKK.[E] | 1xHomocysteineMMTS [K18] | 4,49774E-05 | 2332,09479 | Brain/ FAIMS/MMTS |
| P63101 | P63101 [128-157] | [R].YLAEVAAGDDKKGIVDQSQQAYQEAFEISK.[K] | 1xHomocysteineMMTS [K12] | 0,000152766 | 3464,63485 | Brain/ FAIMS/MMTS |
| P63101 | P63101 [104-120] | [K].FLIPNASQPESKVFYLK.[M] | 1xHomocysteineMMTS [K12] | 4,49774E-05 | 2144,09188 | Brain/ FAIMS/MMTS |
| P63101 | P63101 [139-157] | [K].KGIVDQSQQAYQEAFEISK.[K] | 1xHomocysteineMMTS [K1] | 4,49774E-05 | 2332,09479 | Brain/ FAIMS/MMTS |
| P63101 | P63101 [128-139] | [R].YLAEVAAGDDKK.[G] | 1xHomocysteineMMTS [K] | 4,49774E-05 | 1442,66541 | Brain/ FAIMS/MMTS |
| P63101 | P63101 [1-11] | [-].MDKNELVQKAK.[L] | 1xAcetyl [N-Term]; 1xHomocysteineMMTS [K9] | 4,49774E-05 | 1508,72697 | Brain/ FAIMS/MMTS |
| P63101 | P63101 [1-9] | [-].MDKNELVQK.[A] | 1xAcetyl [N-Term]; 1xHomocysteineMMTS [K3] | 4,49774E-05 | 1309,59489 | Brain/ FAIMS/MMTS |
| P63158 | P63158 [30-43] | [K].KHPDASVNFSEFSK.[K] | 1xHomocysteineIAA [K1] | 4,32935E-05 | 1766,81664 | Brain/ No FAIMS/IAA |
| P63158 | P63158 [30-43] | [K].KHPDASVNFSEFSK.[K] | 1xHomocysteineIAA [K1] | 4,32935E-05 | 1766,81664 | Brain/ FAIMS/IAA |
| P63158 | P63158 [113-127] | [K].IKGEHPGLSIGDVAK.[K] | 1xHomocysteineMMTS [K2] | 0,000530332 | 1683,85567 | Brain/ FAIMS/MMTS |
| P63158; P30681 | P63158 [155-163]; P30681 [155-163] | [K].YEKDIAAYR.[A] | 1xHomocysteineIAA [K3] | 0,000273428 | 1302,6147 | Liver/ FAIMS/IAA |
| P63158; P30681 | P63158 [155-163]; P30681 [155-163] | [K].YEKDIAAYR.[A] | 1xHomocysteineIAA [K3] | 0,000273428 | 1302,6147 | Brain/ No FAIMS/IAA |
| P63158; P30681 | P63158 [155-163]; P30681 [155-163] | [K].YEKDIAAYR.[A] | 1xHomocysteineIAA [K3] | 0,000273428 | 1302,6147 | Brain/ FAIMS/IAA |
| P63168 | P63168 [10-36] | [K].NADMSEEMQQDSVECATQALEKYNIEK.[D] | 1xCarbamidomethyl [C15]; 1xHomocysteineIAA [K22] | 4,32935E-05 | 3335,41707 | Brain/ FAIMS/IAA |
| P63213 | P63213 [33-62] | [K].AAADLMAYCEAHAKEDPLLTPVPASENPFR.[E] | 1xCarbamidomethyl [C9]; 1xHomocysteineIAA [K14] | 4,32935E-05 | 3458,61776 | Brain/ No FAIMS/IAA |
| P63213 | P63213 [33-62] | [K].AAADLMAYCEAHAKEDPLLTPVPASENPFR.[E] | 1xCarbamidomethyl [C9]; 1xHomocysteineIAA [K14] | 4,32935E-05 | 3458,61776 | Brain/ FAIMS/IAA |
| P63213 | P63213 [14-27] | [R].KLVEQLKMEANIDR.[I] | 1xHomocysteineMMTS [K7] | 4,49774E-05 | 1849,93327 | Brain/ FAIMS/MMTS |
| P63242 | P63242 [110-126] | [R].EDLRLPEGDLGKEIEQK.[Y] | 1xHomocysteineIAA [K12] | 4,32935E-05 | 2143,06996 | Brain/ FAIMS/IAA |
| P63260 | P63260 [1-28] | [-].MEEEIAALVIDNGSGMCKAGFAGDDAPR.[A] | 1xCarbamidomethyl [C17]; 1xMet-loss+Acetyl [N-Term]; 1xHomocysteineIAA [K18] | 4,32935E-05 | 3009,33868 | Brain/ No FAIMS/IAA |
| P63276 | P63276 [24-33] | [R].LGNDFHTNKR.[V] | 1xHomocysteineMMTS [K9] | 0,0076359 | 1364,6198 | Liver/ FAIMS/MMTS |
| P63318 | P63318 [639-649] | [R].SGENFDKFFTR.[A] | 1xHomocysteineIAA [K7] | 0,00012645 | 1521,67909 | Brain/ FAIMS/IAA |
| P63318; P20444 | P63318 [182-199]; P20444 [182-199] | [RK].NLIPMDPNGLSDPYVKLK.[L] | 1xHomocysteineIAA [K16] | 0,00052325 | 2188,11407 | Brain/ FAIMS/IAA |
| P63321 | P63321 [146-161] | [R].ADQWNVNYVETSAKTR.[A] | 1xHomocysteineIAA [K14] | 4,32935E-05 | 2055,95526 | Brain/ No FAIMS/IAA |
| P63321 | P63321 [146-161] | [R].ADQWNVNYVETSAKTR.[A] | 1xHomocysteineIAA [K14] | 4,32935E-05 | 2055,95526 | Brain/ FAIMS/IAA |
| P63323 | P63323 [122-131] | [K].DVIEEYFKCK.[K] | 1xCarbamidomethyl [C9]; 1xHomocysteineIAA [K8] | 9,61721E-05 | 1504,68106 | Brain/ FAIMS/IAA |
| P63323 | P63323 [117-129] | [K].ESQAKDVIEEYFK.[C] | 1xHomocysteineMMTS [K5] | 0,00182026 | 1748,78698 | Liver/ No FAIMS/MMTS |
| P63323 | P63323 [117-129] | [K].ESQAKDVIEEYFK.[C] | 1xHomocysteineMMTS [K5] | 0,00182026 | 1748,78698 | Liver/ FAIMS/MMTS |
| P63328 | P63328 [415-441] | [R].EESESVLTLKGLTPTGMLPSGVLSGGK.[Q] | 1xHomocysteineIAA [K10] | 4,32935E-05 | 2861,46347 | Brain/ FAIMS/IAA |
| P63328 | P63328 [467-478] | [K].ITSFEEAKGLDR.[I] | 1xHomocysteineMMTS [K8] | 0,000507453 | 1528,71343 | Brain/ FAIMS/MMTS |
| P67778 | P67778 [198-207] | [R].FVVEKAEQQK.[K] | 1xHomocysteineIAA [K5] | 0,00282663 | 1379,69876 | Liver/ FAIMS/IAA |
| P67778 | P67778 [118-133] | [R].VLPSITTEILKSVVAR.[F] | 1xHomocysteineIAA [K11] | 4,32935E-05 | 1900,09359 | Brain/ FAIMS/IAA |
| P67778 | P67778 [1-11] | [-].MAAKVFESIGK.[F] | 1xMet-loss+Acetyl [N-Term]; 1xHomocysteineMMTS [K4] | 0,00191513 | 1254,62209 | Liver/ FAIMS/MMTS |
| P67778 | P67778 [198-207] | [R].FVVEKAEQQK.[K] | 1xHomocysteineMMTS [K5] | 0,00794311 | 1368,66502 | Liver/ FAIMS/MMTS |
| P67778 | P67778 [208-219] | [K].KAAIISAEGDSK.[A] | 1xHomocysteineMMTS [K1] | 0,00885916 | 1352,65485 | Liver/ FAIMS/MMTS |
| P67778 | P67778 [240-253] | [R].KLEAAEDIAYQLSR.[S] | 1xHomocysteineMMTS [K1] | 4,49774E-05 | 1769,85607 | Liver/ FAIMS/MMTS |
| P68037 | P68037 [123-133] | [R].ADLAEEYSKDR.[K] | 1xHomocysteineIAA [K9] | 4,32935E-05 | 1470,65293 | Brain/ FAIMS/IAA |
| P68040 | P68040 [265-280] | [K].IIVDELKQEVISTSSK.[A] | 1xHomocysteineIAA [K7] | 0,00263986 | 1963,04162 | Liver/ FAIMS/IAA |
| P68040 | P68040 [246-264] | [R].YWLCAATGPSIKIWDLEGK.[I] | 1xCarbamidomethyl [C4]; 1xHomocysteineIAA [K12] | 0,00012645 | 2382,16209 | Brain/ FAIMS/IAA |
| P68040 | P68040 [265-280] | [K].IIVDELKQEVISTSSK.[A] | 1xHomocysteineMMTS [K7] | 4,49774E-05 | 1952,00788 | Liver/ FAIMS/MMTS |
| P68134 | P68134 [293-314] | [R].KDLYANNVMSGGTTMYPGIADR.[M] | 1xHomocysteineIAA [K1] | 4,32935E-05 | 2548,16289 | Liver/ No FAIMS/IAA |
| P68134 | P68134 [293-314] | [R].KDLYANNVMSGGTTMYPGIADR.[M] | 1xHomocysteineIAA [K1] | 4,32935E-05 | 2548,16289 | Liver/ FAIMS/IAA |
| P68134 | P68134 [71-97] | [K].YPIEHGIITNWDDMEKIWHHTFYNELR.[V] | 1xHomocysteineMMTS [K16] | 4,49774E-05 | 3620,65481 | Liver/ No FAIMS/MMTS |
| P68134 | P68134 [293-314] | [R].KDLYANNVMSGGTTMYPGIADR.[M] | 1xHomocysteineMMTS [K1] | 4,49774E-05 | 2537,12915 | Liver/ No FAIMS/MMTS |
| P68134 | P68134 [293-314] | [R].KDLYANNVMSGGTTMYPGIADR.[M] | 1xHomocysteineMMTS [K1] | 4,49774E-05 | 2537,12915 | Liver/ FAIMS/MMTS |
| P68134 | P68134 [293-314] | [R].KDLYANNVMSGGTTMYPGIADR.[M] | 2xOxidation [M9; M15]; 1xHomocysteineMMTS [K1] | 4,49774E-05 | 2569,11898 | Liver/ FAIMS/MMTS |

| | | | | | | |
|---|---|---|---|---|---|---|
| P68134 | P68134 [71-97] | [K].YPIEHGIITNWDDMEKIWHHTFYNELR.[V] | 1xOxidation [M14]; 1xHomocysteineMMTS [K16] | 4,49774E-05 | 3636,64973 | Liver/ FAIMS/MMTS |
| P68134 | P68134 [293-314] | [R].KDLYANNVMSGGTTMYPGIADR.[M] | 1xOxidation [M]; 1xHomocysteineMMTS [K1] | 4,49774E-05 | 2553,12406 | Liver/ FAIMS/MMTS |
| P68134 | P68134 [71-97] | [K].YPIEHGIITNWDDMEKIWHHTFYNELR.[V] | 1xHomocysteineMMTS [K16] | 4,49774E-05 | 3620,65481 | Liver/ FAIMS/MMTS |
| P68134 | P68134 [71-97] | [K].YPIEHGIITNWDDMEKIWHHTFYNELR.[V] | 1xHomocysteineMMTS [K16] | 4,49774E-05 | 3620,65481 | Brain/ No FAIMS/MMTS |
| P68134 | P68134 [71-97] | [K].YPIEHGIITNWDDMEKIWHHTFYNELR.[V] | 1xHomocysteineMMTS [K16] | 4,49774E-05 | 3620,65481 | Brain/ FAIMS/MMTS |
| P68254 | P68254 [140-158] | [K].QTIENSQGAYQEAFDISKK.[E] | 1xHomocysteineIAA [K18] | 4,32935E-05 | 2331,09215 | Brain/ FAIMS/IAA |
| P68254 | P68254 [76-83] | [K].LQLIKDYR.[E] | 1xHomocysteineMMTS [K5] | 0,00818121 | 1211,62751 | Liver/ FAIMS/MMTS |
| P68254 | P68254 [139-157] | [R].KQTIENSQGAYQEAFDISK.[K] | 1xHomocysteineMMTS [K1] | 4,49774E-05 | 2320,05841 | Liver/ FAIMS/MMTS |
| P68254 | P68254 [42-55] | [R].NLLSVAYKNVVGGR.[R] | 1xHomocysteineMMTS [K8] | 4,49774E-05 | 1652,86109 | Brain/ No FAIMS/MMTS |
| P68254 | P68254 [76-83] | [K].LQLIKDYR.[E] | 1xHomocysteineMMTS [K5] | 0,00818121 | 1211,62751 | Brain/ No FAIMS/MMTS |
| P68254 | P68254 [42-55] | [R].NLLSVAYKNVVGGR.[R] | 1xHomocysteineMMTS [K8] | 4,49774E-05 | 1652,86109 | Brain/ FAIMS/MMTS |
| P68254 | P68254 [61-75] | [R].VISSIEQKTDTSDKK.[L] | 1xHomocysteineMMTS [K8] | 0,000530332 | 1841,89833 | Brain/ FAIMS/MMTS |
| P68254 | P68254 [76-83] | [K].LQLIKDYR.[E] | 1xHomocysteineMMTS [K5] | 0,00818121 | 1211,62751 | Brain/ FAIMS/MMTS |
| P68254 | P68254 [139-157] | [R].KQTIENSQGAYQEAFDISK.[K] | 1xHomocysteineMMTS [K1] | 4,49774E-05 | 2320,05841 | Brain/ FAIMS/MMTS |
| P68254 | P68254 [1-9] | [-].MEKTELIQK.[A] | 1xAcetyl [N-Term]; 1xHomocysteineMMTS [K3] | 4,49774E-05 | 1324,63094 | Brain/ FAIMS/MMTS |
| P68254; O70456; Q9CQV8; P63101 | P68254 [10-18]; O70456 [10-18]; Q9CQV8 [12-20]; P63101 [10-18] | [K].AKLAEQAER.[Y] | 1xHomocysteineIAA [K2] | 0,00243432 | 1189,59938 | Brain/ No FAIMS/IAA |
| P68254; O70456; Q9CQV8; P63101 | P68254 [10-18]; O70456 [10-18]; Q9CQV8 [12-20]; P63101 [10-18] | [K].AKLAEQAER.[Y] | 1xHomocysteineIAA [K2] | 0,00243432 | 1189,59938 | Brain/ FAIMS/IAA |
| P68254; O70456; Q9CQV8; P63101 | P68254 [10-18]; O70456 [10-18]; Q9CQV8 [12-20]; P63101 [10-18] | [K].AKLAEQAER.[Y] | 1xHomocysteineMMTS [K2] | 0,000706375 | 1178,56564 | Liver/ FAIMS/MMTS |
| P68254; O70456; Q9CQV8; P63101 | P68254 [10-18]; O70456 [10-18]; Q9CQV8 [12-20]; P63101 [10-18] | [K].AKLAEQAER.[Y] | 1xHomocysteineMMTS [K2] | 0,000706375 | 1178,56564 | Brain/ No FAIMS/MMTS |
| P68254; O70456; Q9CQV8; P63101 | P68254 [10-18]; O70456 [10-18]; Q9CQV8 [12-20]; P63101 [10-18] | [K].AKLAEQAER.[Y] | 1xHomocysteineMMTS [K2] | 0,000706375 | 1178,56564 | Brain/ FAIMS/MMTS |
| P68254; Q9CQV8 | P68254 [121-127]; Q9CQV8 [123-129] | [K].MKGDYFR.[Y] | 1xHomocysteineIAA [K2] | 0,00347063 | 1090,48085 | Brain/ No FAIMS/IAA |
| P68254; Q9CQV8 | P68254 [121-127]; Q9CQV8 [123-129] | [K].MKGDYFR.[Y] | 1xHomocysteineIAA [K2] | 0,00347063 | 1090,48085 | Brain/ FAIMS/IAA |
| P68254; Q9CQV8; P63101 | P68254 [158-167]; Q9CQV8 [160-169]; P63101 [158-167] | [K].KEMQPTHPIR.[L] | 1xHomocysteineIAA [K1] | 4,32935E-05 | 1410,69805 | Brain/ No FAIMS/IAA |
| P68254; Q9CQV8; P63101 | P68254 [158-167]; Q9CQV8 [160-169]; P63101 [158-167] | [K].KEMQPTHPIR.[L] | 1xHomocysteineIAA [K1] | 4,32935E-05 | 1410,69805 | Brain/ FAIMS/IAA |
| P68254; Q9CQV8; P63101 | P68254 [158-167]; Q9CQV8 [160-169]; P63101 [158-167] | [K].KEMQPTHPIR.[L] | 1xHomocysteineMMTS [K1] | 4,49774E-05 | 1399,66431 | Liver/ FAIMS/MMTS |
| P68254; Q9CQV8; P63101 | P68254 [158-167]; Q9CQV8 [160-169]; P63101 [158-167] | [K].KEMQPTHPIR.[L] | 1xHomocysteineMMTS [K1] | 4,49774E-05 | 1399,66431 | Brain/ No FAIMS/MMTS |
| P68254; Q9CQV8; P63101 | P68254 [158-167]; Q9CQV8 [160-169]; P63101 [158-167] | [K].KEMQPTHPIR.[L] | 1xHomocysteineMMTS [K1] | 4,49774E-05 | 1399,66431 | Brain/ FAIMS/MMTS |
| P68368 | P68368 [340-370] | [R].SIQFVDWCPTGFKVGINYQPPTVVPGGDLAK.[V] | 1xCarbamidomethyl [C8]; 1xHomocysteineIAA [K13] | 4,32935E-05 | 3564,76541 | Brain/ No FAIMS/IAA |
| P68368 | P68368 [340-370] | [R].SIQFVDWCPTGFKVGINYQPPTVVPGGDLAK.[V] | 1xCarbamidomethyl [C8]; 1xHomocysteineIAA [K13] | 4,32935E-05 | 3564,76541 | Brain/ FAIMS/IAA |
| P68368 | P68368 [327-338] | [K].DVNAAIAAIKTK.[R] | 1xHomocysteineMMTS [K10] | 6,70238E-05 | 1377,72287 | Brain/ No FAIMS/MMTS |
| P68368 | P68368 [106-121] | [R].GHYTIGKEIIDPVLDR.[I] | 1xHomocysteineMMTS [K7] | 4,49774E-05 | 1988,99323 | Brain/ FAIMS/MMTS |
| P68368 | P68368 [327-338] | [K].DVNAAIAAIKTK.[R] | 1xHomocysteineMMTS [K10] | 6,70238E-05 | 1377,72287 | Brain/ FAIMS/MMTS |
| P68369; P68373 | P68369 [106-121]; P68373 [106-121] | [R].GHYTIGKEIIDLVLDR.[I] | 1xHomocysteineIAA [K7] | 4,32935E-05 | 2016,05827 | Brain/ No FAIMS/IAA |
| P68369; P68373 | P68369 [106-121]; P68373 [106-121] | [R].GHYTIGKEIIDLVLDR.[I] | 1xHomocysteineIAA [K7] | 4,32935E-05 | 2016,05827 | Brain/ FAIMS/IAA |
| P68369; P68373; P68368 | P68369 [309-320]; P68373 [309-320]; P68368 [309-320] | [R].HGKYMACCLLYR.[G] | 2xCarbamidomethyl [C7; C8]; 1xHomocysteineIAA [K3] | 4,32935E-05 | 1745,77427 | Brain/ No FAIMS/IAA |
| P68369; P68373; P68368 | P68369 [309-320]; P68373 [309-320]; P68368 [309-320] | [R].HGKYMACCLLYR.[G] | 2xCarbamidomethyl [C7; C8]; 1xHomocysteineIAA [K3] | 4,32935E-05 | 1745,77427 | Brain/ FAIMS/IAA |
| P68372 | P68372 [47-62] | [R].INVYYNEATGGKYVPR.[A] | 1xHomocysteineIAA [K12] | 4,32935E-05 | 2017,98002 | Brain/ No FAIMS/IAA |
| P68372 | P68372 [47-62] | [R].INVYYNEATGGKYVPR.[A] | 1xHomocysteineIAA [K12] | 4,32935E-05 | 2017,98002 | Brain/ FAIMS/IAA |
| P68372 | P68372 [47-62] | [R].INVYYNEATGGKYVPR.[A] | 1xHomocysteineMMTS [K12] | 4,49774E-05 | 2006,94628 | Brain/ FAIMS/MMTS |
| P68404 | P68404 [584-601] | [R].LGCGPEGERDIKEHAFFR.[Y] | 1xCarbamidomethyl [C3]; 1xHomocysteineIAA [K12] | 4,32935E-05 | 2292,06483 | Brain/ FAIMS/IAA |

| | | | | | | |
|---|---|---|---|---|---|---|
| P68404 | P68404 [602-610] | [R].YIDWEKLER.[K] | 1xHomocysteineMMTS [K6] | 0,00207528 | 1414,64937 | Brain/ FAIMS/MMTS |
| P68404; P20444 | P68404 [43-62]; P20444 [43-62] | [R].FFKQPTFCSHCTDFIWGFGK.[Q] | 2xCarbamidomethyl [C8; C11]; 1xHomocysteineIAA [K3] | 4,32935E-05 | 2684,18832 | Brain/ FAIMS/IAA |
| P68510 | P68510 [144-162] | [K].NSVVEASEAAYKEAFEISK.[E] | 1xHomocysteineIAA [K12] | 4,32935E-05 | 2246,06454 | Brain/ No FAIMS/IAA |
| P68510 | P68510 [144-162] | [K].NSVVEASEAAYKEAFEISK.[E] | 1xHomocysteineIAA [K12] | 4,32935E-05 | 2246,06454 | Brain/ FAIMS/IAA |
| P68510 | P68510 [20-28] | [R].YDDMASAMK.[A] | 1xOxidation [M8]; 1xHomocysteineMMTS [K9] | 0,00783297 | 1210,42471 | Brain/ FAIMS/MMTS |
| P68510 | P68510 [20-42] | [R].YDDMASAMKAVTELNEPLSNEDR.[N] | 1xHomocysteineMMTS [K9] | 4,49774E-05 | 2762,17761 | Brain/ FAIMS/MMTS |
| P68510 | P68510 [62-77] | [R].VISSIEQKTMADGNEK.[K] | 1xHomocysteineMMTS [K8] | 0,00156537 | 1912,88129 | Brain/ FAIMS/MMTS |
| P68510 | P68510 [144-162] | [K].NSVVEASEAAYKEAFEISK.[E] | 1xHomocysteineMMTS [K12] | 0,00125756 | 2235,0308 | Brain/ FAIMS/MMTS |
| P68510 | P68510 [143-155] | [K].KNSVVEASEAAYK.[E] | 1xHomocysteineMMTS [K1] | 4,49774E-05 | 1558,72399 | Brain/ FAIMS/MMTS |
| P68510; O70456; P61982; P63101 | P68510 [121-132]; O70456 [118-129]; P61982 [121-132]; P63101 [116-127] | [KR].VFYLKMKGDYYR.[Y] | 2xHomocysteineIAA [K5; K7] | 6,30912E-05 | 1930,90124 | Brain/ FAIMS/IAA |
| P68510; O70456; P61982; P63101 | P68510 [126-132]; O70456 [123-129]; P61982 [126-132]; P63101 [121-127] | [K].MKGDYYR.[Y] | 1xHomocysteineMMTS [K2] | 0,00883288 | 1095,44202 | Liver/ FAIMS/MMTS |
| P68510; O70456; P61982; P63101 | P68510 [126-132]; O70456 [123-129]; P61982 [126-132]; P63101 [121-127] | [K].MKGDYYR.[Y] | 1xHomocysteineMMTS [K2] | 0,00883288 | 1095,44202 | Brain/ No FAIMS/MMTS |
| P68510; O70456; P61982; P63101 | P68510 [126-132]; O70456 [123-129]; P61982 [126-132]; P63101 [121-127] | [K].MKGDYYR.[Y] | 1xHomocysteineMMTS [K2] | 0,00883288 | 1095,44202 | Brain/ FAIMS/MMTS |
| P68510; P61982; Q9CQV8; P63101 | P68510 [43-56]; P61982 [43-56]; Q9CQV8 [44-57]; P63101 [42-55] | [R].NLLSVAYKNVVGAR.[R] | 1xHomocysteineIAA [K8] | 4,32935E-05 | 1677,91049 | Liver/ No FAIMS/IAA |
| P68510; P61982; Q9CQV8; P63101 | P68510 [43-56]; P61982 [43-56]; Q9CQV8 [44-57]; P63101 [42-55] | [R].NLLSVAYKNVVGAR.[R] | 1xHomocysteineIAA [K8] | 4,32935E-05 | 1677,91049 | Liver/ FAIMS/IAA |
| P68510; P61982; Q9CQV8; P63101 | P68510 [43-56]; P61982 [43-56]; Q9CQV8 [44-57]; P63101 [42-55] | [R].NLLSVAYKNVVGAR.[R] | 1xHomocysteineIAA [K8] | 4,32935E-05 | 1677,91049 | Brain/ No FAIMS/IAA |
| P68510; P61982; Q9CQV8; P63101 | P68510 [43-56]; P61982 [43-56]; Q9CQV8 [44-57]; P63101 [42-55] | [R].NLLSVAYKNVVGAR.[R] | 1xHomocysteineIAA [K8] | 4,32935E-05 | 1677,91049 | Brain/ FAIMS/IAA |
| P68510; P61982; Q9CQV8; P63101 | P68510 [43-56]; P61982 [43-56]; Q9CQV8 [44-57]; P63101 [42-55] | [R].NLLSVAYKNVVGAR.[R] | 1xHomocysteineMMTS [K8] | 4,49774E-05 | 1666,87674 | Liver/ No FAIMS/MMTS |
| P68510; P61982; Q9CQV8; P63101 | P68510 [43-56]; P61982 [43-56]; Q9CQV8 [44-57]; P63101 [42-55] | [R].NLLSVAYKNVVGAR.[R] | 1xHomocysteineMMTS [K8] | 4,49774E-05 | 1666,87674 | Liver/ FAIMS/MMTS |
| P68510; P61982; Q9CQV8; P63101 | P68510 [43-56]; P61982 [43-56]; Q9CQV8 [44-57]; P63101 [42-55] | [R].NLLSVAYKNVVGAR.[R] | 1xHomocysteineMMTS [K8] | 4,49774E-05 | 1666,87674 | Brain/ No FAIMS/MMTS |
| P68510; P61982; Q9CQV8; P63101 | P68510 [43-56]; P61982 [43-56]; Q9CQV8 [44-57]; P63101 [42-55] | [R].NLLSVAYKNVVGAR.[R] | 1xHomocysteineMMTS [K8] | 4,49774E-05 | 1666,87674 | Brain/ FAIMS/MMTS |
| P68510; P68254; O70456; P61982; Q9CQV8; P63101 | P68510 [121-127]; P68254 [116-122]; O70456 [118-124]; P61982 [121-127]; Q9CQV8 [118-124]; P63101 [116-122] | [KR].VFYLKMK.[G] | 1xOxidation [M6]; 1xHomocysteineIAA [K5] | 9,61721E-05 | 1118,57368 | Brain/ No FAIMS/IAA |
| P68510; P68254; O70456; P61982; Q9CQV8; P63101 | P68510 [121-127]; P68254 [116-122]; O70456 [118-124]; P61982 [121-127]; Q9CQV8 [118-124]; P63101 [116-122] | [KR].VFYLKMK.[G] | 1xOxidation [M6]; 1xHomocysteineIAA [K5] | 9,61721E-05 | 1118,57368 | Brain/ FAIMS/IAA |
| P70168 | P70168 [860-870] | [K].TLATWATKELR.[K] | 1xHomocysteineIAA [K8] | 9,61721E-05 | 1463,76751 | Brain/ FAIMS/IAA |
| P70202 | P70202 [31-48] | [K].LFLVQTVQQASKEDIPGR.[G] | 1xHomocysteineIAA [K12] | 4,32935E-05 | 2203,15396 | Brain/ FAIMS/IAA |
| P70290 | P70290 [16-30] | [R].TALSDLYLEHLLQKR.[N] | 1xHomocysteineIAA [K14] | 0,000289025 | 1974,04771 | Brain/ FAIMS/IAA |
| P70296 | P70296 [149-155] | [K].FKVETFR.[K] | 1xHomocysteineIAA [K2] | 0,00253458 | 1100,55573 | Liver/ FAIMS/IAA |
| P70296 | P70296 [156-179] | [R].KKYNLGAPVAGTCYQAEWDDYVPK.[L] | 1xCarbamidomethyl [C13]; 1xHomocysteineIAA [K2] | 4,32935E-05 | 2947,37532 | Brain/ FAIMS/IAA |
| P70296 | P70296 [149-155] | [K].FKVETFR.[K] | 1xHomocysteineMMTS [K2] | 0,00545792 | 1089,52198 | Liver/ No FAIMS/MMTS |
| P70296 | P70296 [149-155] | [K].FKVETFR.[K] | 1xHomocysteineMMTS [K2] | 0,00545792 | 1089,52198 | Liver/ FAIMS/MMTS |
| P70296 | P70296 [180-187] | [K].LYEQLSGK.[-] | 1xHomocysteineMMTS [K8] | 0,0004709 | 1100,51148 | Liver/ FAIMS/MMTS |
| P70296 | P70296 [180-187] | [K].LYEQLSGK.[-] | 1xHomocysteineMMTS [K8] | 0,0004709 | 1100,51148 | Brain/ FAIMS/MMTS |
| P70296 | P70296 [149-155] | [K].FKVETFR.[K] | 1xHomocysteineMMTS [K2] | 0,00545792 | 1089,52198 | Brain/ FAIMS/MMTS |

| | | | | | | |
|---|---|---|---|---|---|---|
| P70297 | P70297 [155-170] | [K].ASPALVAKDPGTVATK.[K] | 1xHomocysteineIAA [K8] | 4,32935E-05 | 1699,90473 | Brain/ FAIMS/IAA |
| P70297; O88811 | P70297 [353-363]; O88811 [343-353] | [R].KHSELSELNVK.[V] | 1xHomocysteineIAA [K1] | 0,00230865 | 1457,74169 | Brain/ FAIMS/IAA |
| P70333; O35737 | P70333 [180-188]; O35737 [180-188] | [R].YIEIFKSSR.[A] | 1xHomocysteineIAA [K6] | 0,000800823 | 1316,66673 | Brain/ FAIMS/IAA |
| P70349 | P70349 [1-21] | [-].MADEIAKAQVAQPGGDTIFGK.[I] | 1xMet-loss+Acetyl [N-Term]; 1xHomocysteineIAA [K7] | 4,32935E-05 | 2232,09651 | Liver/ FAIMS/IAA |
| P70349 | P70349 [58-82] | [K].KHISQISVADDDDESLLGHLMIVGK.[K] | 1xHomocysteineIAA [K1] | 4,32935E-05 | 2894,43865 | Brain/ No FAIMS/IAA |
| P70349 | P70349 [58-82] | [K].KHISQISVADDDDESLLGHLMIVGK.[K] | 1xHomocysteineIAA [K1] | 4,32935E-05 | 2894,43865 | Brain/ FAIMS/IAA |
| P70349 | P70349 [1-21] | [-].MADEIAKAQVAQPGGDTIFGK.[I] | 1xMet-loss+Acetyl [N-Term]; 1xHomocysteineMMTS [K7] | 4,49774E-05 | 2221,06276 | Liver/ No FAIMS/MMTS |
| P70349 | P70349 [26-37] | [K].EIPAKIIFEDDR.[C] | 1xHomocysteineMMTS [K5] | 4,49774E-05 | 1608,77603 | Liver/ No FAIMS/MMTS |
| P70349 | P70349 [1-21] | [-].MADEIAKAQVAQPGGDTIFGK.[I] | 1xMet-loss+Acetyl [N-Term]; 1xHomocysteineMMTS [K7] | 4,49774E-05 | 2221,06276 | Liver/ FAIMS/MMTS |
| P70349 | P70349 [26-37] | [K].EIPAKIIFEDDR.[C] | 1xHomocysteineMMTS [K5] | 4,49774E-05 | 1608,77603 | Liver/ FAIMS/MMTS |
| P70349 | P70349 [8-24] | [K].AQVAQPGGDTIFGKIIR.[K] | 1xHomocysteineMMTS [K14] | 4,49774E-05 | 1933,99865 | Liver/ FAIMS/MMTS |
| P70349 | P70349 [1-21] | [-].MADEIAKAQVAQPGGDTIFGK.[I] | 1xMet-loss+Acetyl [N-Term]; 1xHomocysteineMMTS [K7] | 4,49774E-05 | 2221,06276 | Brain/ No FAIMS/MMTS |
| P70349 | P70349 [1-21] | [-].MADEIAKAQVAQPGGDTIFGK.[I] | 1xMet-loss+Acetyl [N-Term]; 1xHomocysteineMMTS [K7] | 4,49774E-05 | 2221,06276 | Brain/ FAIMS/MMTS |
| P70349 | P70349 [26-37] | [K].EIPAKIIFEDDR.[C] | 1xHomocysteineMMTS [K5] | 4,49774E-05 | 1608,77603 | Brain/ FAIMS/MMTS |
| P70349 | P70349 [8-24] | [K].AQVAQPGGDTIFGKIIR.[K] | 1xHomocysteineMMTS [K14] | 4,49774E-05 | 1933,99865 | Brain/ FAIMS/MMTS |
| P70404 | P70404 [137-157] | [R].TSLDLYANVIHCKSLPGVVTR.[H] | 1xCarbamidomethyl [C12]; 1xHomocysteineIAA [K13] | 4,32935E-05 | 2517,29522 | Brain/ FAIMS/IAA |
| P70414 | P70414 [54-71] | [K].KGVILPIWEPQDPSFGDK.[I] | 1xHomocysteineIAA [K1] | 4,32935E-05 | 2200,1107 | Brain/ FAIMS/IAA |
| P70441 | P70441 [117-143] | [K].SEQAEPPAAADTHEAGDQNEAEKSHLR.[E] | 1xHomocysteineIAA [K23] | 4,32935E-05 | 3062,35044 | Brain/ FAIMS/IAA |
| P70441 | P70441 [223-235] | [K].LLVVDKETDEFFK.[K] | 1xHomocysteineMMTS [K6] | 0,0024085 | 1745,84886 | Liver/ FAIMS/MMTS |
| P70670 | P70670 [2086-2114] | [K].SPASDTYIVFGEAKIEDLSQQAQLAAAEK.[F] | 1xHomocysteineIAA [K14] | 4,32935E-05 | 3254,58855 | Brain/ FAIMS/IAA |
| P70670 | P70670 [2100-2116] | [K].IEDLSQQAQLAAAEKFK.[V] | 1xHomocysteineMMTS [K15] | 4,49774E-05 | 2053,00927 | Liver/ No FAIMS/MMTS |
| P70670 | P70670 [2100-2116] | [K].IEDLSQQAQLAAAEKFK.[V] | 1xHomocysteineMMTS [K15] | 4,49774E-05 | 2053,00927 | Liver/ FAIMS/MMTS |
| P70691; Q63886; Q6ZQM8; Q64435; Q62452 | P70691 [429-437]; Q63886 [431-439]; Q6ZQM8 [427-435]; Q64435 [427-435]; Q62452 [424-432] | [K].TVINNKSYK.[E] | 1xHomocysteineMMTS [K6] | 0,00210568 | 1229,60169 | Liver/ FAIMS/MMTS |
| P70691; Q63886; Q6ZQM8; Q64435; Q62452 | P70691 [348-365]; Q63886 [350-367]; Q6ZQM8 [346-363]; Q64435 [346-363]; Q62452 [343-360] | [K].NTILVKWLPQNDLLGHPK.[T] | 1xHomocysteineMMTS [K6] | 0,00272554 | 2249,19333 | Liver/ FAIMS/MMTS |
| P70691; Q63886; Q6ZQM8; Q64435; Q62452 | P70691 [354-367]; Q63886 [356-369]; Q6ZQM8 [352-365]; Q64435 [352-365]; Q62452 [349-362] | [K].WLPQNDLLGHPKTR.[A] | 1xHomocysteineMMTS [K12] | 0,0004709 | 1837,92 | Liver/ FAIMS/MMTS |
| P70691; Q63886; Q6ZQM8; Q64435; Q62452 | P70691 [318-328]; Q63886 [320-330]; Q6ZQM8 [316-326]; Q64435 [316-326]; Q62452 [313-323] | [K].KAMEIAEALGR.[I] | 1xHomocysteineMMTS [K1] | 0,000231655 | 1351,65307 | Liver/ FAIMS/MMTS |
| P70694 | P70694 [156-170] | [K].DAGLAKSIGVSNFNR.[R] | 1xHomocysteineMMTS [K6] | 0,00359453 | 1711,82544 | Liver/ FAIMS/MMTS |
| P80313 | P80313 [154-166] | [K].MLEKCAMTALSSK.[L] | 1xCarbamidomethyl [C5]; 1xHomocysteineIAA [K4] | 9,61721E-05 | 1643,76237 | Brain/ FAIMS/IAA |
| P80314 | P80314 [428-441] | [R].TPGKEAVAMESFAK.[A] | 1xHomocysteineIAA [K4] | 0,00012645 | 1639,78184 | Brain/ FAIMS/IAA |
| P80315 | P80315 [528-539] | [R].SILKIDDVVNTR.[-] | 1xHomocysteineIAA [K4] | 0,00012645 | 1546,82575 | Brain/ FAIMS/IAA |
| P83917 | P83917 [151-167] | [K].EANVKCPQVVISFYEER.[L] | 1xCarbamidomethyl [C6]; 1xHomocysteineIAA [K5] | 4,32935E-05 | 2242,0631 | Brain/ FAIMS/IAA |
| P83940 | P83940 [21-33] | [K].LISSDGHEFIVKR.[E] | 1xHomocysteineIAA [K12] | 4,32935E-05 | 1674,8632 | Brain/ FAIMS/IAA |
| P84075 | P84075 [155-171] | [R].QMDTNNDGKLSLEEFIR.[G] | 1xHomocysteineIAA [K9] | 4,32935E-05 | 2184,00598 | Brain/ No FAIMS/IAA |
| P84075 | P84075 [155-171] | [R].QMDTNNDGKLSLEEFIR.[G] | 1xHomocysteineIAA [K9] | 4,32935E-05 | 2184,00598 | Brain/ FAIMS/IAA |
| P84075 | P84075 [155-171] | [R].QMDTNNDGKLSLEEFIR.[G] | 1xHomocysteineMMTS [K9] | 0,000133296 | 2172,97224 | Brain/ FAIMS/MMTS |
| P84075; P62748 | P84075 [51-70]; P62748 [51-70] | [K].IYANFFPYGDASKFAEHVFR.[T] | 1xHomocysteineIAA [K13] | 4,32935E-05 | 2553,20198 | Brain/ No FAIMS/IAA |
| P84075; P62748 | P84075 [51-70]; P62748 [51-70] | [K].IYANFFPYGDASKFAEHVFR.[T] | 1xHomocysteineIAA [K13] | 4,32935E-05 | 2553,20198 | Brain/ FAIMS/IAA |
| P84086 | P84086 [99-122] | [K].KAIPAGCGDEEEEEEESILDTVLK.[Y] | 1xCarbamidomethyl [C7]; 1xHomocysteineIAA [K1] | 4,32935E-05 | 2835,29105 | Brain/ No FAIMS/IAA |
| P84086 | P84086 [99-122] | [K].KAIPAGCGDEEEEEEESILDTVLK.[Y] | 1xCarbamidomethyl [C7]; 1xHomocysteineIAA [K1] | 4,32935E-05 | 2835,29105 | Brain/ FAIMS/IAA |
| P84086; P63040 | P84086 [7-18]; P63040 [7-18] | [K].QALGGATKDMGK.[M] | 1xHomocysteineIAA [K8] | 4,32935E-05 | 1350,65043 | Brain/ No FAIMS/IAA |
| P84086; P63040 | P84086 [7-18]; P63040 [7-18] | [K].QALGGATKDMGK.[M] | 1xHomocysteineIAA [K8] | 4,32935E-05 | 1350,65043 | Brain/ FAIMS/IAA |
| P84086; P63040 | P84086 [7-18]; P63040 [7-18] | [K].QALGGATKDMGK.[M] | 1xHomocysteineMMTS [K8] | 0,000152766 | 1339,61669 | Brain/ FAIMS/MMTS |
| P84089 | P84089 [1-17] | [-].MSHTILLVQPTKRPEGR.[T] | 1xMet-loss+Acetyl [N-Term]; 1xHomocysteineIAA [K12] | 0,00410529 | 2048,10695 | Brain/ FAIMS/IAA |
| P84091 | P84091 [140-162] | [K].SQHQTKEEQSQITSQVTGQIGWR.[R] | 1xHomocysteineIAA [K6] | 4,32935E-05 | 2830,35368 | Brain/ FAIMS/IAA |
| P84096 | P84096 [129-147] | [R].LKEQGQAPITPQQGQALAK.[Q] | 1xHomocysteineIAA [K2] | 6,30912E-05 | 2180,14921 | Liver/ FAIMS/IAA |
| P84096 | P84096 [129-147] | [R].LKEQGQAPITPQQGQALAK.[Q] | 1xHomocysteineIAA [K2] | 6,30912E-05 | 2180,14921 | Brain/ FAIMS/IAA |

| | | | | | | |
|---|---|---|---|---|---|---|
| P84099 | P84099 [137-146] | [R].ILMEHIHKLK.[A] | 1xHomocysteineIAA [K8] | 0,000210768 | 1435,79122 | Brain/ No FAIMS/IAA |
| P84099 | P84099 [137-146] | [R].ILMEHIHKLK.[A] | 1xHomocysteineIAA [K8] | 0,000210768 | 1435,79122 | Brain/ FAIMS/IAA |
| P84244 | P84244 [74-84] | [R].EIAQDFKTDLR.[F] | 1xHomocysteineMMTS [K7] | 0,00107839 | 1498,70286 | Liver/ FAIMS/MMTS |
| P84244 | P84244 [118-129] | [R].VTIMPKDIQLAR.[R] | 1xHomocysteineMMTS [K6] | 4,49774E-05 | 1547,81064 | Liver/ FAIMS/MMTS |
| P84244 | P84244 [74-84] | [R].EIAQDFKTDLR.[F] | 1xHomocysteineMMTS [K7] | 0,00107839 | 1498,70286 | Brain/ FAIMS/MMTS |
| P84244 | P84244 [118-129] | [R].VTIMPKDIQLAR.[R] | 1xHomocysteineMMTS [K6] | 4,49774E-05 | 1547,81064 | Brain/ FAIMS/MMTS |
| P84244 | P84244 [55-64] | [R].YQKSTELLIR.[K] | 1xHomocysteineMMTS [K3] | 0,000530332 | 1413,72287 | Brain/ FAIMS/MMTS |
| P84244; P84228 | P84244 [118-129]; P84228 [118-129] | [R].VTIMPKDIQLAR.[R] | 1xHomocysteineIAA [K6] | 4,32935E-05 | 1558,84438 | Brain/ No FAIMS/IAA |
| P84244; P84228 | P84244 [118-129]; P84228 [118-129] | [R].VTIMPKDIQLAR.[R] | 1xHomocysteineIAA [K6] | 4,32935E-05 | 1558,84438 | Brain/ FAIMS/IAA |
| P97300 | P97300 [245-271] | [R].SENKNEGQDAMMYCKSVGYPHPEWIW R.[K] | 1xCarbamidomethyl [C14]; 1xHomocysteineIAA [K15] | 4,32935E-05 | 3486,51225 | Brain/ FAIMS/IAA |
| P97315 | P97315 [42-59] | [K].KNLDSTTVAVHGEEIYCK.[S] | 1xCarbamidomethyl [C17]; 1xHomocysteineIAA [K1] | 4,32935E-05 | 2238,05293 | Brain/ FAIMS/IAA |
| P97315 | P97315 [70-91] | [K].GYGYGQGAGTLSTDKGESLGIK.[H] | 1xHomocysteineMMTS [K15] | 0,000530332 | 2322,07406 | Brain/ FAIMS/MMTS |
| P97315; P97314 | P97315 [169-178]; P97314 [169-178] | [K].GCYAKNFGPK.[G] | 1xCarbamidomethyl [C2]; 1xHomocysteineIAA [K5] | 0,000228034 | 1315,59219 | Brain/ No FAIMS/IAA |
| P97315; P97314 | P97315 [169-178]; P97314 [169-178] | [K].GCYAKNFGPK.[G] | 1xCarbamidomethyl [C2]; 1xHomocysteineIAA [K5] | 0,000228034 | 1315,59219 | Brain/ FAIMS/IAA |
| P97328 | P97328 [142-152] | [R].NASEQVKMLQR.[I] | 1xHomocysteineIAA [K7] | 0,000243434 | 1477,72499 | Liver/ FAIMS/IAA |
| P97328 | P97328 [142-152] | [R].NASEQVKMLQR.[I] | 1xHomocysteineMMTS [K7] | 0,00074635 | 1466,69125 | Liver/ FAIMS/MMTS |
| P97343 | P97343 [370-387] | [R].GQVFVEYANAGDSKAAQK.[L] | 1xHomocysteineMMTS [K14] | 0,00452171 | 2045,94192 | Brain/ FAIMS/MMTS |
| P97351 | P97351 [241-252] | [K].AAGDETGAKVER.[A] | 1xHomocysteineMMTS [K9] | 0,00275712 | 1366,60896 | Liver/ FAIMS/MMTS |
| P97352 | P97352 [79-88] | [R].LIGELAKEVR.[K] | 1xHomocysteineIAA [K7] | 4,32935E-05 | 1301,72458 | Liver/ FAIMS/IAA |
| P97352 | P97352 [79-88] | [R].LIGELAKEVR.[K] | 1xHomocysteineIAA [K7] | 4,32935E-05 | 1301,72458 | Brain/ FAIMS/IAA |
| P97352 | P97352 [79-88] | [R].LIGELAKEVR.[K] | 1xHomocysteineMMTS [K7] | 8,93346E-05 | 1290,69084 | Liver/ FAIMS/MMTS |
| P97370 | P97370 [270-277] | [R].VSFKVTAR.[A] | 1xHomocysteineIAA [K4] | 0,000305143 | 1081,58228 | Brain/ No FAIMS/IAA |
| P97370 | P97370 [270-277] | [R].VSFKVTAR.[A] | 1xHomocysteineIAA [K4] | 0,000305143 | 1081,58228 | Brain/ FAIMS/IAA |
| P97371 | P97371 [6-18] | [R].VHPEAQAKVDVFR.[E] | 1xHomocysteineMMTS [K8] | 0,000282457 | 1658,81414 | Liver/ FAIMS/MMTS |
| P97371 | P97371 [188-198] | [K].AAKQPHVGDYR.[Q] | 1xHomocysteineMMTS [K3] | 0,00460899 | 1404,6511 | Liver/ FAIMS/MMTS |
| P97371 | P97371 [36-58] | [K].KISELDAFLKEPALNEANLSNLK.[A] | 1xHomocysteineMMTS [K1] | 4,49774E-05 | 2720,39975 | Liver/ FAIMS/MMTS |
| P97384 | P97384 [370-384] | [R].LGTDESKFNAILCSR.[S] | 1xCarbamidomethyl [C13]; 1xHomocysteineIAA [K7] | 0,000624482 | 1884,89424 | Brain/ FAIMS/IAA |
| P97427 | P97427 [486-496] | [R].SKVFGLHSVSR.[G] | 1xHomocysteineIAA [K2] | 4,32935E-05 | 1390,72598 | Brain/ FAIMS/IAA |
| P97449 | P97449 [141-158] | [R].TLDGTPAPNIDKTELVER.[T] | 1xHomocysteineMMTS [K12] | 0,00386874 | 2132,03622 | Liver/ FAIMS/MMTS |
| P97450 | P97450 [80-94] | [K].QMYGKGEMDTFPTFK.[F] | 1xHomocysteineIAA [K5] | 4,32935E-05 | 1953,85435 | Liver/ FAIMS/IAA |
| P97450 | P97450 [80-94] | [K].QMYGKGEMDTFPTFK.[F] | 1xHomocysteineIAA [K5] | 4,32935E-05 | 1953,85435 | Brain/ FAIMS/IAA |
| P97450 | P97450 [42-48] | [K].LFVDKIR.[E] | 1xHomocysteineMMTS [K5] | 0,00991259 | 1053,55837 | Liver/ No FAIMS/MMTS |
| P97450 | P97450 [95-108] | [K].FDDPKFEVIDKPQS.[-] | 1xHomocysteineMMTS [K5] | 0,000764279 | 1827,82918 | Liver/ No FAIMS/MMTS |
| P97450 | P97450 [80-94] | [K].QMYGKGEMDTFPTFK.[F] | 1xHomocysteineMMTS [K5] | 0,000250158 | 1942,82061 | Liver/ FAIMS/MMTS |
| P97450 | P97450 [95-108] | [K].FDDPKFEVIDKPQS.[-] | 1xHomocysteineMMTS [K5] | 0,000764279 | 1827,82918 | Liver/ FAIMS/MMTS |
| P97450 | P97450 [42-48] | [K].LFVDKIR.[E] | 1xHomocysteineMMTS [K5] | 0,00991259 | 1053,55837 | Liver/ FAIMS/MMTS |
| P97450 | P97450 [80-94] | [K].QMYGKGEMDTFPTFK.[F] | 1xHomocysteineMMTS [K5] | 0,000250158 | 1942,82061 | Brain/ FAIMS/MMTS |
| P97457 | P97457 [155-169] | [K].NICYVITHGDAKDQE.[-] | 1xCarbamidomethyl [C3]; 1xHomocysteineIAA [K12] | 4,32935E-05 | 1936,85277 | Liver/ No FAIMS/IAA |
| P97457 | P97457 [155-169] | [K].NICYVITHGDAKDQE.[-] | 1xCarbamidomethyl [C3]; 1xHomocysteineIAA [K12] | 4,32935E-05 | 1936,85277 | Liver/ FAIMS/IAA |
| P97457 | P97457 [117-130] | [K].KQFLEELLTTQCDR.[F] | 1xMethylthio [C12]; 1xHomocysteineMMTS [K1] | 0,000266738 | 1932,86862 | Liver/ No FAIMS/MMTS |
| P97457 | P97457 [106-116] | [K].VLDPEGKGTIK.[K] | 1xHomocysteineMMTS [K7] | 0,000192903 | 1319,66977 | Liver/ No FAIMS/MMTS |
| P97457 | P97457 [42-51] | [R].DGIIDKEDLR.[D] | 1xHomocysteineMMTS [K6] | 4,49774E-05 | 1336,62355 | Liver/ No FAIMS/MMTS |
| P97457 | P97457 [60-72] | [R].LNVKNEELDAMMK.[E] | 1xHomocysteineMMTS [K4] | 4,49774E-05 | 1697,77293 | Liver/ No FAIMS/MMTS |
| P97457 | P97457 [91-105] | [K].LKGADPEDVITGAFK.[V] | 1xHomocysteineMMTS [K2] | 4,49774E-05 | 1723,83935 | Liver/ No FAIMS/MMTS |
| P97457 | P97457 [60-72] | [R].LNVKNEELDAMMK.[E] | 2xOxidation [M11; M12]; 1xHomocysteineMMTS [K4] | 0,000152766 | 1729,76276 | Liver/ FAIMS/MMTS |
| P97457 | P97457 [60-72] | [R].LNVKNEELDAMMK.[E] | 1xOxidation [M]; 1xHomocysteineMMTS [K4] | 4,49774E-05 | 1713,76785 | Liver/ FAIMS/MMTS |
| P97457 | P97457 [117-130] | [K].KQFLEELLTTQCDR.[F] | 1xMethylthio [C12]; 1xHomocysteineMMTS [K1] | 0,000266738 | 1932,86862 | Liver/ FAIMS/MMTS |
| P97457 | P97457 [106-116] | [K].VLDPEGKGTIK.[K] | 1xHomocysteineMMTS [K7] | 0,000192903 | 1319,66977 | Liver/ FAIMS/MMTS |
| P97457 | P97457 [42-51] | [R].DGIIDKEDLR.[D] | 1xHomocysteineMMTS [K6] | 4,49774E-05 | 1336,62355 | Liver/ FAIMS/MMTS |
| P97457 | P97457 [60-72] | [R].LNVKNEELDAMMK.[E] | 1xHomocysteineMMTS [K4] | 4,49774E-05 | 1697,77293 | Liver/ FAIMS/MMTS |
| P97457 | P97457 [91-105] | [K].LKGADPEDVITGAFK.[V] | 1xHomocysteineMMTS [K2] | 4,49774E-05 | 1723,83935 | Liver/ FAIMS/MMTS |
| P97493 | P97493 [148-162] | [K].FVGIKDEDQLEAFLK.[K] | 1xHomocysteineIAA [K5] | 9,61721E-05 | 1925,96773 | Brain/ FAIMS/IAA |
| P97493 | P97493 [148-162] | [K].FVGIKDEDQLEAFLK.[K] | 1xHomocysteineMMTS [K5] | 0,000173718 | 1914,93398 | Liver/ FAIMS/MMTS |
| P97496 | P97496 [590-601] | [K].EKPIDLQNFGLR.[T] | 1xHomocysteineMMTS [K2] | 0,00677626 | 1592,79234 | Brain/ No FAIMS/MMTS |
| P97765 | P97765 [57-64] | [R].VIFLSKGK.[D] | 1xHomocysteineIAA [K6] | 0,00128643 | 1065,61251 | Brain/ No FAIMS/IAA |
| P97797 | P97797 [216-238] | [R].VVLNSMDVNSKVICEVAHITLDR.[S] | 1xCarbamidomethyl [C14]; 1xHomocysteineIAA [K11] | 4,32935E-05 | 2786,39977 | Brain/ No FAIMS/IAA |
| P97797 | P97797 [216-238] | [R].VVLNSMDVNSKVICEVAHITLDR.[S] | 1xCarbamidomethyl [C14]; 1xHomocysteineIAA [K11] | 4,32935E-05 | 2786,39977 | Brain/ FAIMS/IAA |
| P97807 | P97807 [59-71] | [K].VPTDKYYGAQTVR.[S] | 1xHomocysteineIAA [K5] | 6,30912E-05 | 1671,81592 | Liver/ FAIMS/IAA |
| P97807 | P97807 [442-464] | [R].INKLMNESLMLVTALNPHIGYDK.[A] | 1xHomocysteineIAA [K3] | 4,32935E-05 | 2788,41944 | Brain/ FAIMS/IAA |
| P97807 | P97807 [211-220] | [K].LHDALSAKSK.[E] | 1xHomocysteineMMTS [K8] | 0,00804194 | 1232,61259 | Liver/ No FAIMS/MMTS |
| P97807 | P97807 [170-180] | [K].KPVHPNDHVNK.[S] | 1xHomocysteineMMTS [K1] | 0,00409028 | 1447,6933 | Liver/ No FAIMS/MMTS |
| P97807 | P97807 [72-84] | [R].STMNFKIGGATER.[M] | 1xOxidation [M3]; 1xHomocysteineMMTS [K6] | 4,49774E-05 | 1590,70729 | Liver/ FAIMS/MMTS |
| P97807 | P97807 [59-71] | [K].VPTDKYYGAQTVR.[S] | 1xHomocysteineMMTS [K5] | 0,000706375 | 1660,78217 | Liver/ FAIMS/MMTS |
| P97807 | P97807 [99-119] | [R].AAAEVNQEYGLDPKIASAIMK.[A] | 1xHomocysteineMMTS [K14] | 0,000133296 | 2382,1502 | Liver/ FAIMS/MMTS |
| P97807 | P97807 [49-63] | [R].VEFDTFGELKVPTDK.[Y] | 1xHomocysteineMMTS [K10] | 0,000133296 | 1887,8867 | Liver/ FAIMS/MMTS |
| P97952 | P97952 [63-72] | [K].GTEEFVKILR.[Y] | 1xHomocysteineIAA [K7] | 0,00071373 | 1365,7195 | Brain/ No FAIMS/IAA |

| | | | | | | |
|---|---|---|---|---|---|---|
| P97952 | P97952 [63-72] | [K].GTEEFVKILR.[Y] | 1xHomocysteineIAA [K7] | 0,00071373 | 1365,7195 | Brain/ FAIMS/IAA |
| P99024 | P99024 [363-380] | [K].MAVTFIGNSTAIQELFKR.[I] | 1xHomocysteineIAA [K17] | 4,32935E-05 | 2200,12531 | Brain/ No FAIMS/IAA |
| P99024 | P99024 [363-380] | [K].MAVTFIGNSTAIQELFKR.[I] | 1xHomocysteineIAA [K17] | 4,32935E-05 | 2200,12531 | Brain/ FAIMS/IAA |
| P99026 | P99026 [197-211] | [R].EVLEKQPVLSQTEAR.[E] | 1xHomocysteineIAA [K5] | 0,00434736 | 1900,97969 | Brain/ FAIMS/IAA |
| P99027 | P99027 [62-98] | [K].LASVPAGGAVAVSAAPGSAAPAAGSAPAAAEEKKDEK.[K] | 1xHomocysteineIAA [K] | 4,32935E-05 | 3448,73767 | Liver/ No FAIMS/IAA |
| P99027 | P99027 [62-98] | [K].LASVPAGGAVAVSAAPGSAAPAAGSAPAAAEEKKDEK.[K] | 1xHomocysteineIAA [K] | 4,32935E-05 | 3448,73767 | Liver/ FAIMS/IAA |
| P99027 | P99027 [62-98] | [K].LASVPAGGAVAVSAAPGSAAPAAGSAPAAAEEKKDEK.[K] | 1xHomocysteineIAA [K] | 4,32935E-05 | 3448,73767 | Brain/ FAIMS/IAA |
| P99027 | P99027 [42-61] | [K].VISELNGKNIEDVIAQGVGK.[L] | 1xHomocysteineMMTS [K8] | 4,49774E-05 | 2246,15191 | Liver/ No FAIMS/MMTS |
| P99027 | P99027 [62-98] | [K].LASVPAGGAVAVSAAPGSAAPAAGSAPAAAEEKKDEK.[K] | 1xHomocysteineMMTS [K33] | 4,49774E-05 | 3437,70393 | Liver/ No FAIMS/MMTS |
| P99027 | P99027 [25-41] | [K].KILDSVGIEADDDRLNK.[V] | 1xHomocysteineMMTS [K1] | 4,49774E-05 | 2064,01 | Liver/ No FAIMS/MMTS |
| P99027 | P99027 [62-98] | [K].LASVPAGGAVAVSAAPGSAAPAAGSAPAAAEEKKDEK.[K] | 1xHomocysteineMMTS [K33] | 4,49774E-05 | 3437,70393 | Liver/ FAIMS/MMTS |
| P99027 | P99027 [42-61] | [K].VISELNGKNIEDVIAQGVGK.[L] | 1xHomocysteineMMTS [K8] | 4,49774E-05 | 2246,15191 | Liver/ FAIMS/MMTS |
| P99027 | P99027 [3-24] | [R].YVASYLLAALGGNSSPSAKDIK.[K] | 1xHomocysteineMMTS [K19] | 4,49774E-05 | 2388,19378 | Liver/ FAIMS/MMTS |
| P99027 | P99027 [25-41] | [K].KILDSVGIEADDDRLNK.[V] | 1xHomocysteineMMTS [K1] | 4,49774E-05 | 2064,01 | Liver/ FAIMS/MMTS |
| P99028 | P99028 [33-43] | [R].EHCEQLEKCVK.[A] | 2xCarbamidomethyl [C3; C9]; 1xHomocysteineIAA [K8] | 0,000584831 | 1633,71311 | Liver/ FAIMS/IAA |
| P99029 | P99029 [72-81] | [K].VNLAELFKGK.[K] | 1xHomocysteineIAA [K8] | 4,32935E-05 | 1292,70312 | Liver/ FAIMS/IAA |
| P99029 | P99029 [82-98] | [K].KGVLFGVPGAFTPGCSK.[T] | 1xCarbamidomethyl [C15]; 1xHomocysteineIAA [K1] | 4,32935E-05 | 1895,95064 | Brain/ FAIMS/IAA |
| P99029 | P99029 [72-81] | [K].VNLAELFKGK.[K] | 1xHomocysteineMMTS [K8] | 0,000231655 | 1281,66938 | Liver/ No FAIMS/MMTS |
| P99029 | P99029 [54-71] | [K].VGDAIPSVEVFEGEPGKK.[V] | 1xHomocysteineMMTS [K17] | 4,49774E-05 | 2020,97182 | Liver/ No FAIMS/MMTS |
| P99029 | P99029 [99-114] | [K].THLPGFVEQAGALKAK.[G] | 1xHomocysteineMMTS [K14] | 0,0018995 | 1829,94007 | Liver/ No FAIMS/MMTS |
| P99029 | P99029 [72-81] | [K].VNLAELFKGK.[K] | 1xHomocysteineMMTS [K8] | 0,000231655 | 1281,66938 | Liver/ FAIMS/MMTS |
| P99029 | P99029 [54-71] | [K].VGDAIPSVEVFEGEPGKK.[V] | 1xHomocysteineMMTS [K17] | 4,49774E-05 | 2020,97182 | Liver/ FAIMS/MMTS |
| P99029 | P99029 [72-81] | [K].VNLAELFKGK.[K] | 1xHomocysteineMMTS [K8] | 0,000231655 | 1281,66938 | Brain/ No FAIMS/MMTS |
| P99029 | P99029 [54-71] | [K].VGDAIPSVEVFEGEPGKK.[V] | 1xHomocysteineMMTS [K17] | 4,49774E-05 | 2020,97182 | Brain/ No FAIMS/MMTS |
| P99029 | P99029 [71-79] | [K].KVNLAELFK.[G] | 1xHomocysteineMMTS [K1] | 4,49774E-05 | 1224,64791 | Brain/ No FAIMS/MMTS |
| P99029 | P99029 [72-81] | [K].VNLAELFKGK.[K] | 1xHomocysteineMMTS [K8] | 0,000231655 | 1281,66938 | Brain/ FAIMS/MMTS |
| P99029 | P99029 [54-71] | [K].VGDAIPSVEVFEGEPGKK.[V] | 1xHomocysteineMMTS [K17] | 4,49774E-05 | 2020,97182 | Brain/ FAIMS/MMTS |
| P99029 | P99029 [99-114] | [K].THLPGFVEQAGALKAK.[G] | 1xHomocysteineMMTS [K14] | 0,0018995 | 1829,94007 | Brain/ FAIMS/MMTS |
| P99029 | P99029 [71-79] | [K].KVNLAELFK.[G] | 1xHomocysteineMMTS [K1] | 4,49774E-05 | 1224,64791 | Brain/ FAIMS/MMTS |
| Q00493 | Q00493 [434-455] | [K].KVAVPFSPAVGVDFELESFSER.[K] | 1xHomocysteineIAA [K1] | 0,00071373 | 2584,2752 | Brain/ FAIMS/IAA |
| Q00623 | Q00623 [93-106] | [R].DFWDNLEKETDWVR.[Q] | 1xHomocysteineIAA [K8] | 4,32935E-05 | 2026,89635 | Liver/ FAIMS/IAA |
| Q00623 | Q00623 [93-106] | [R].DFWDNLEKETDWVR.[Q] | 1xHomocysteineIAA [K8] | 4,32935E-05 | 2026,89635 | Brain/ FAIMS/IAA |
| Q00623 | Q00623 [201-216] | [R].LAELKSNPTLNEYHTR.[A] | 1xHomocysteineMMTS [K5] | 4,49774E-05 | 2048,98921 | Liver/ No FAIMS/MMTS |
| Q00623 | Q00623 [140-154] | [R].QKVAPLGAELQESAR.[Q] | 1xHomocysteineMMTS [K2] | 4,49774E-05 | 1759,88295 | Liver/ No FAIMS/MMTS |
| Q00623 | Q00623 [118-129] | [K].QKVQPYLDEFQK.[K] | 1xHomocysteineMMTS [K2] | 4,49774E-05 | 1685,80257 | Liver/ No FAIMS/MMTS |
| Q00623 | Q00623 [34-46] | [K].VKDFANVYVDAVK.[D] | 1xHomocysteineMMTS [K2] | 0,0004709 | 1630,79676 | Liver/ No FAIMS/MMTS |
| Q00623 | Q00623 [131-139] | [K].WKEDVELYR.[Q] | 1xHomocysteineMMTS [K2] | 0,000152766 | 1400,63372 | Liver/ No FAIMS/MMTS |
| Q00623 | Q00623 [36-50] | [K].DFANVYVDAVKDSGR.[D] | 1xHomocysteineMMTS [K11] | 6,70238E-05 | 1818,81493 | Liver/ No FAIMS/MMTS |
| Q00623 | Q00623 [120-130] | [K].VQPYLDEFQKK.[W] | 1xHomocysteineMMTS [K10] | 0,000152766 | 1557,744 | Liver/ No FAIMS/MMTS |
| Q00623 | Q00623 [112-119] | [K].DLEEVKQK.[V] | 1xHomocysteineMMTS [K6] | 0,00239411 | 1151,54351 | Liver/ FAIMS/MMTS |
| Q00623 | Q00623 [118-129] | [K].QKVQPYLDEFQK.[K] | 1xHomocysteineMMTS [K2] | 4,49774E-05 | 1685,80257 | Liver/ FAIMS/MMTS |
| Q00623 | Q00623 [34-46] | [K].VKDFANVYVDAVK.[D] | 1xHomocysteineMMTS [K2] | 0,0004709 | 1630,79676 | Liver/ FAIMS/MMTS |
| Q00623 | Q00623 [131-139] | [K].WKEDVELYR.[Q] | 1xHomocysteineMMTS [K2] | 0,000152766 | 1400,63372 | Liver/ FAIMS/MMTS |
| Q00623 | Q00623 [120-130] | [K].VQPYLDEFQKK.[W] | 1xHomocysteineMMTS [K10] | 0,000152766 | 1557,744 | Liver/ FAIMS/MMTS |
| Q00623 | Q00623 [201-216] | [R].LAELKSNPTLNEYHTR.[A] | 1xHomocysteineMMTS [K5] | 4,49774E-05 | 2048,98921 | Liver/ FAIMS/MMTS |
| Q00623 | Q00623 [219-227] | [K].THLKTLGEK.[A] | 1xHomocysteineMMTS [K4] | 0,00721311 | 1189,60678 | Liver/ FAIMS/MMTS |
| Q00623 | Q00623 [140-154] | [R].QKVAPLGAELQESAR.[Q] | 1xHomocysteineMMTS [K2] | 4,49774E-05 | 1759,88295 | Liver/ FAIMS/MMTS |
| Q00623 | Q00623 [155-163] | [R].QKLQELQGR.[L] | 1xHomocysteineMMTS [K2] | 0,00631419 | 1262,63439 | Liver/ FAIMS/MMTS |
| Q00623 | Q00623 [36-50] | [K].DFANVYVDAVKDSGR.[D] | 1xHomocysteineMMTS [K11] | 6,70238E-05 | 1818,81493 | Liver/ FAIMS/MMTS |
| Q00897 | Q00897 [309-328] | [R].LSISGNYNLKTLMSPLGITR.[I] | 1xOxidation [M13]; 1xHomocysteineMMTS [K10] | 0,00109439 | 2357,20257 | Liver/ No FAIMS/MMTS |
| Q00897 | Q00897 [309-328] | [R].LSISGNYNLKTLMSPLGITR.[I] | 1xHomocysteineMMTS [K10] | 0,000547722 | 2341,20765 | Liver/ FAIMS/MMTS |
| Q00897 | Q00897 [309-328] | [R].LSISGNYNLKTLMSPLGITR.[I] | 1xOxidation [M13]; 1xHomocysteineMMTS [K10] | 0,00109439 | 2357,20257 | Liver/ FAIMS/MMTS |
| Q00898 | Q00898 [214-235] | [K].KPFDPENTKQAEFHVDESTTVK.[V] | 1xHomocysteineIAA [K9] | 9,61721E-05 | 2721,28247 | Liver/ FAIMS/IAA |
| Q00898 | Q00898 [329-354] | [R].IFNSGADLSGITEENAPLKLSQAVHK.[A] | 1xHomocysteineMMTS [K19] | 4,49774E-05 | 2902,44374 | Liver/ No FAIMS/MMTS |
| Q00898 | Q00898 [214-222] | [K].KPFDPENTK.[Q] | 1xHomocysteineMMTS [K1] | 0,00171109 | 1238,55441 | Liver/ FAIMS/MMTS |
| Q00898 | Q00898 [329-354] | [R].IFNSGADLSGITEENAPLKLSQAVHK.[A] | 1xHomocysteineMMTS [K19] | 4,49774E-05 | 2902,44374 | Liver/ FAIMS/MMTS |
| Q01279 | Q01279 [742-750] | [K].IPVAIKELR.[E] | 1xHomocysteineIAA [K6] | 0,00156934 | 1212,71329 | Liver/ FAIMS/IAA |
| Q01279 | Q01279 [742-750] | [K].IPVAIKELR.[E] | 1xHomocysteineMMTS [K6] | 0,000250158 | 1201,67955 | Liver/ FAIMS/MMTS |
| Q01768 | Q01768 [43-56] | [R].ASEEHLKQHYIDLK.[D] | 1xHomocysteineIAA [K7] | 0,000273428 | 1884,92726 | Brain/ FAIMS/IAA |
| Q01768; P15532 | Q01768 [89-105]; P15532 [89-105] | [R].VMLGETNPADSKPGTIR.[G] | 1xHomocysteineIAA [K12] | 4,32935E-05 | 1959,96266 | Liver/ FAIMS/IAA |
| Q01768; P15532 | Q01768 [89-105]; P15532 [89-105] | [R].VMLGETNPADSKPGTIR.[G] | 1xHomocysteineIAA [K12] | 4,32935E-05 | 1959,96266 | Brain/ FAIMS/IAA |
| Q01768; P15532 | Q01768 [7-18]; P15532 [7-18] | [R].TFIAIKPDGVQR.[G] | 1xHomocysteineMMTS [K6] | 6,70238E-05 | 1507,77597 | Liver/ No FAIMS/MMTS |
| Q01768; P15532 | Q01768 [89-105]; P15532 [89-105] | [R].VMLGETNPADSKPGTIR.[G] | 1xHomocysteineMMTS [K12] | 4,49774E-05 | 1948,92891 | Liver/ No FAIMS/MMTS |
| Q01768; P15532 | Q01768 [89-105]; P15532 [89-105] | [R].VMLGETNPADSKPGTIR.[G] | 1xHomocysteineMMTS [K12] | 4,49774E-05 | 1948,92891 | Liver/ FAIMS/MMTS |
| Q01768; P15532 | Q01768 [89-105]; P15532 [89-105] | [R].VMLGETNPADSKPGTIR.[G] | 1xOxidation [M2]; 1xHomocysteineMMTS [K12] | 4,49774E-05 | 1964,92383 | Liver/ FAIMS/MMTS |

| | | | | | | |
|---|---|---|---|---|---|---|
| Q01768; P15532 | Q01768 [19-27]; P15532 [19-27] | [R].GLVGEIIKR.[F] | 1xHomocysteineMMTS [K8] | 0,00222764 | 1147,6326 | Liver/ FAIMS/MMTS |
| Q01768; P15532 | Q01768 [7-18]; P15532 [7-18] | [R].TFIAIKPDGVQR.[G] | 1xHomocysteineMMTS [K6] | 6,70238E-05 | 1507,77597 | Liver/ FAIMS/MMTS |
| Q01768; P15532 | Q01768 [89-105]; P15532 [89-105] | [R].VMLGETNPADSKPGTIR.[G] | 1xHomocysteineMMTS [K12] | 4,49774E-05 | 1948,92891 | Brain/ FAIMS/MMTS |
| Q01853 | Q01853 [240-256] | [R].GILLYGPPGTGKTLIAR.[A] | 1xHomocysteineIAA [K12] | 4,32935E-05 | 1901,06771 | Brain/ No FAIMS/IAA |
| Q01853 | Q01853 [240-256] | [R].GILLYGPPGTGKTLIAR.[A] | 1xHomocysteineIAA [K12] | 4,32935E-05 | 1901,06771 | Brain/ FAIMS/IAA |
| Q02053 | Q02053 [369-384] | [R].SPPSVKQNSLDEDLIR.[K] | 1xHomocysteineIAA [K6] | 4,32935E-05 | 1971,98042 | Brain/ FAIMS/IAA |
| Q02248 | Q02248 [125-151] | [R].LAEPSQMLKHAVVNLINYQDDAELATR.[A] | 1xHomocysteineIAA [K9] | 4,32935E-05 | 3213,60309 | Brain/ FAIMS/IAA |
| Q02819 | Q02819 [69-80] | [R].EKLQAANAEDIK.[S] | 1xHomocysteineIAA [K2] | 0,00012645 | 1503,74717 | Liver/ FAIMS/IAA |
| Q02819 | Q02819 [69-80] | [R].EKLQAANAEDIK.[S] | 1xHomocysteineIAA [K2] | 0,00012645 | 1503,74717 | Brain/ FAIMS/IAA |
| Q03265 | Q03265 [428-441] | [K].QVAGTMKLELAQYR.[E] | 1xHomocysteineIAA [K7] | 4,32935E-05 | 1781,90369 | Brain/ FAIMS/IAA |
| Q03265 | Q03265 [172-182] | [R].VGLKAPGIIPR.[I] | 1xHomocysteineMMTS [K4] | 8,93346E-05 | 1283,73264 | Liver/ FAIMS/MMTS |
| Q03265 | Q03265 [532-541] | [K].ISEQSDAKLK.[E] | 1xHomocysteineMMTS [K8] | 0,000738471 | 1281,61773 | Liver/ FAIMS/MMTS |
| Q03265 | Q03265 [187-204] | [R].EPMQTGIKAVDSLVPIGR.[G] | 1xHomocysteineMMTS [K8] | 4,49774E-05 | 2074,04936 | Liver/ FAIMS/MMTS |
| Q03265 | Q03265 [532-541] | [K].ISEQSDAKLK.[E] | 1xHomocysteineMMTS [K8] | 0,000738471 | 1281,61773 | Brain/ No FAIMS/MMTS |
| Q03265 | Q03265 [532-541] | [K].ISEQSDAKLK.[E] | 1xHomocysteineMMTS [K8] | 0,000738471 | 1281,61773 | Brain/ FAIMS/MMTS |
| Q03265 | Q03265 [187-204] | [R].EPMQTGIKAVDSLVPIGR.[G] | 1xHomocysteineMMTS [K8] | 4,49774E-05 | 2074,04936 | Brain/ FAIMS/MMTS |
| Q03265 | Q03265 [172-182] | [R].VGLKAPGIIPR.[I] | 1xHomocysteineMMTS [K4] | 8,93346E-05 | 1283,73264 | Brain/ FAIMS/MMTS |
| Q03265 | Q03265 [219-231] | [K].TSIAIDTIINQKR.[F] | 1xHomocysteineMMTS [K12] | 6,70238E-05 | 1635,85567 | Brain/ FAIMS/MMTS |
| Q03265 | Q03265 [150-167] | [R].VVDALGNAIDGKGPIGSK.[T] | 1xHomocysteineMMTS [K12] | 4,49774E-05 | 1873,95103 | Brain/ FAIMS/MMTS |
| Q03517 | Q03517 [477-492] | [R].ANQIPKVAWIPDVESR.[Q] | 1xHomocysteineIAA [K6] | 4,32935E-05 | 1997,02731 | Brain/ FAIMS/IAA |
| Q03958 | Q03958 [59-73] | [K].LLGPVLVKQELGEAR.[A] | 1xHomocysteineIAA [K8] | 4,32935E-05 | 1796,00987 | Brain/ FAIMS/IAA |
| Q03958 | Q03958 [59-73] | [K].LLGPVLVKQELGEAR.[A] | 1xHomocysteineMMTS [K8] | 0,00182026 | 1784,97612 | Brain/ FAIMS/MMTS |
| Q04447 | Q04447 [152-172] | [R].RAIEKLAVEALSSLDGDLSGR.[Y] | 1xHomocysteineIAA [K5] | 4,32935E-05 | 2374,23948 | Brain/ No FAIMS/IAA |
| Q04447 | Q04447 [152-172] | [R].RAIEKLAVEALSSLDGDLSGR.[Y] | 1xHomocysteineIAA [K5] | 4,32935E-05 | 2374,23948 | Brain/ FAIMS/IAA |
| Q04447 | Q04447 [1-13] | [-].MPFSNSHNTQKLR.[F] | 1xMet-loss [N-Term]; 1xHomocysteineMMTS [K11] | 4,49774E-05 | 1591,74679 | Brain/ No FAIMS/MMTS |
| Q04447 | Q04447 [216-236] | [R].GIWHNDNKTFLVWINEEDHLR.[V] | 1xHomocysteineMMTS [K8] | 4,49774E-05 | 2799,313 | Brain/ No FAIMS/MMTS |
| Q04447 | Q04447 [293-304] | [R].AGVHIKLPHLGK.[H] | 1xHomocysteineMMTS [K6] | 0,000564742 | 1432,79156 | Brain/ No FAIMS/MMTS |
| Q04447 | Q04447 [299-307] | [K].LPHLGKHEK.[F] | 1xHomocysteineMMTS [K6] | 0,0020903 | 1221,62309 | Brain/ No FAIMS/MMTS |
| Q04447 | Q04447 [237-247] | [R].VISMQKGGNMK.[E] | 1xHomocysteineMMTS [K6] | 4,49774E-05 | 1355,63023 | Brain/ No FAIMS/MMTS |
| Q04447 | Q04447 [243-252] | [K].GGNMKEVFTR.[F] | 1xHomocysteineMMTS [K5] | 0,000530332 | 1301,57991 | Brain/ No FAIMS/MMTS |
| Q04447 | Q04447 [152-172] | [R].RAIEKLAVEALSSLDGDLSGR.[Y] | 1xHomocysteineMMTS [K5] | 4,49774E-05 | 2363,20574 | Brain/ No FAIMS/MMTS |
| Q04447 | Q04447 [153-172] | [R].AIEKLAVEALSSLDGDLSGR.[Y] | 1xHomocysteineMMTS [K4] | 4,49774E-05 | 2207,10463 | Brain/ No FAIMS/MMTS |
| Q04447 | Q04447 [14-43] | [R].FPAEDEFPDLSSHNNHMAKVLTPELYAELR.[A] | 1xHomocysteineMMTS [K19] | 4,49774E-05 | 3633,68108 | Brain/ No FAIMS/MMTS |
| Q04447 | Q04447 [367-381] | [R].LEQGQAIDDLMPAQK.[-] | 1xHomocysteineMMTS [K15] | 4,49774E-05 | 1819,8387 | Brain/ No FAIMS/MMTS |
| Q04447 | Q04447 [97-130] | [R].HGGYQPSDEHKTDLNPDNLQGGDDLDPNYVLSSR.[V] | 1xHomocysteineMMTS [K11] | 4,49774E-05 | 3916,71372 | Brain/ No FAIMS/MMTS |
| Q04447 | Q04447 [1-13] | [-].MPFSNSHNTQKLR.[F] | 1xMet-loss [N-Term]; 1xHomocysteineMMTS [K11] | 4,49774E-05 | 1591,74679 | Brain/ FAIMS/MMTS |
| Q04447 | Q04447 [216-236] | [R].GIWHNDNKTFLVWINEEDHLR.[V] | 1xHomocysteineMMTS [K8] | 4,49774E-05 | 2799,313 | Brain/ FAIMS/MMTS |
| Q04447 | Q04447 [293-304] | [R].AGVHIKLPHLGK.[H] | 1xHomocysteineMMTS [K6] | 0,000564742 | 1432,79156 | Brain/ FAIMS/MMTS |
| Q04447 | Q04447 [308-314] | [K].FSEVLKR.[L] | 1xHomocysteineMMTS [K6] | 0,00179098 | 1041,52198 | Brain/ FAIMS/MMTS |
| Q04447 | Q04447 [237-247] | [R].VISMQKGGNMK.[E] | 1xHomocysteineMMTS [K6] | 4,49774E-05 | 1355,63023 | Brain/ FAIMS/MMTS |
| Q04447 | Q04447 [243-252] | [K].GGNMKEVFTR.[F] | 1xHomocysteineMMTS [K5] | 0,000530332 | 1301,57991 | Brain/ FAIMS/MMTS |
| Q04447 | Q04447 [152-172] | [R].RAIEKLAVEALSSLDGDLSGR.[Y] | 1xHomocysteineMMTS [K5] | 4,49774E-05 | 2363,20574 | Brain/ FAIMS/MMTS |
| Q04447 | Q04447 [153-172] | [R].AIEKLAVEALSSLDGDLSGR.[Y] | 1xHomocysteineMMTS [K4] | 4,49774E-05 | 2207,10463 | Brain/ FAIMS/MMTS |
| Q04447 | Q04447 [305-314] | [K].HEKFSEVLKR.[L] | 1xHomocysteineMMTS [K3] | 6,70238E-05 | 1435,71845 | Brain/ FAIMS/MMTS |
| Q04447 | Q04447 [14-43] | [R].FPAEDEFPDLSSHNNHMAKVLTPELYAELR.[A] | 1xHomocysteineMMTS [K19] | 4,49774E-05 | 3633,68108 | Brain/ FAIMS/MMTS |
| Q04447 | Q04447 [367-381] | [R].LEQGQAIDDLMPAQK.[-] | 1xHomocysteineMMTS [K15] | 4,49774E-05 | 1819,8387 | Brain/ FAIMS/MMTS |
| Q04447 | Q04447 [97-130] | [R].HGGYQPSDEHKTDLNPDNLQGGDDLDPNYVLSSR.[V] | 1xHomocysteineMMTS [K11] | 4,49774E-05 | 3916,71372 | Brain/ FAIMS/MMTS |
| Q04736; P05480; P39688 | Q04736 [432-446]; P05480 [426-440]; P39688 [428-442] | [K].FPIKWTAPEAALYGR.[F] | 1xHomocysteineIAA [K4] | 0,00137306 | 1893,968 | Brain/ FAIMS/IAA |
| Q05186 | Q05186 [169-182] | [R].FKASDLDGDLTATR.[E] | 1xHomocysteineIAA [K2] | 0,000177335 | 1683,80066 | Brain/ FAIMS/IAA |
| Q05816 | Q05816 [82-109] | [R].KTETVCTFQDGALVQHQQWDGKESTITR.[K] | 1xCarbamidomethyl [C6]; 1xHomocysteineIAA [K22] | 4,32935E-05 | 3437,62126 | Brain/ FAIMS/IAA |
| Q06185 | Q06185 [60-71] | [R].ELAEAQDDSILK.[-] | 1xHomocysteineIAA [K12] | 4,32935E-05 | 1505,7152 | Liver/ FAIMS/IAA |
| Q06185 | Q06185 [1-15] | [-].MVPPVQVSPLIKFGR.[Y] | 1xMet-loss [N-Term]; 1xHomocysteineIAA [K12] | 4,32935E-05 | 1710,97236 | Brain/ No FAIMS/IAA |
| Q06185 | Q06185 [1-15] | [-].MVPPVQVSPLIKFGR.[Y] | 1xMet-loss [N-Term]; 1xHomocysteineIAA [K12] | 4,32935E-05 | 1710,97236 | Brain/ FAIMS/IAA |
| Q06185 | Q06185 [1-15] | [-].MVPPVQVSPLIKFGR.[Y] | 1xMet-loss [N-Term]; 1xHomocysteineMMTS [K12] | 4,49774E-05 | 1699,93861 | Liver/ No FAIMS/MMTS |
| Q06185 | Q06185 [1-15] | [-].MVPPVQVSPLIKFGR.[Y] | 1xMet-loss [N-Term]; 1xHomocysteineMMTS [K12] | 4,49774E-05 | 1699,93861 | Liver/ FAIMS/MMTS |
| Q06185 | Q06185 [60-71] | [R].ELAEAQDDSILK.[-] | 1xHomocysteineMMTS [K12] | 4,49774E-05 | 1494,68146 | Liver/ FAIMS/MMTS |
| Q06185 | Q06185 [1-15] | [-].MVPPVQVSPLIKFGR.[Y] | 1xMet-loss [N-Term]; 1xHomocysteineMMTS [K12] | 4,49774E-05 | 1699,93861 | Brain/ FAIMS/MMTS |
| Q06890 | Q06890 [40-53] | [R].YINKEIQNAVQGVK.[H] | 1xHomocysteineMMTS [K4] | 0,000282457 | 1766,89279 | Liver/ FAIMS/MMTS |
| Q07076 | Q07076 [400-416] | [R].LYYSMKGAGTDDSTLVR.[I] | 1xHomocysteineIAA [K6] | 4,32935E-05 | 2050,95724 | Brain/ FAIMS/IAA |
| Q07076 | Q07076 [347-360] | [R].SFPQLKATMEAYSR.[M] | 1xHomocysteineMMTS [K6] | 0,00833937 | 1791,82266 | Brain/ FAIMS/MMTS |
| Q08331 | Q08331 [245-255] | [R].KSVMSLAEAGK.[L] | 1xHomocysteineIAA [K1] | 4,32935E-05 | 1294,64937 | Brain/ No FAIMS/IAA |
| Q08331 | Q08331 [245-255] | [R].KSVMSLAEAGK.[L] | 1xHomocysteineIAA [K1] | 4,32935E-05 | 1294,64937 | Brain/ FAIMS/IAA |
| Q08331 | Q08331 [231-244] | [K].NKKEMNIQQLTTYR.[K] | 2xHomocysteineMMTS [K2; K3] | 0,00760806 | 2092,94689 | Brain/ No FAIMS/MMTS |
| Q08331 | Q08331 [40-52] | [K].ELENFFQELEKAR.[K] | 1xHomocysteineMMTS [K11] | 0,000133296 | 1815,84042 | Brain/ No FAIMS/MMTS |

| | | | | | | |
|---|---|---|---|---|---|---|
| Q08331 | Q08331 [143-161] | [K].ANRPYDEPKLQEYTQTILR.[M] | 1xHomocysteineMMTS [K9] | 0,000282457 | 2498,21664 | Brain/ FAIMS/MMTS |
| Q08331 | Q08331 [54-67] | [K].GSGMMSKSDNFGEK.[M] | 1xHomocysteineMMTS [K7] | 0,000530332 | 1637,64264 | Brain/ FAIMS/MMTS |
| Q08331 | Q08331 [40-52] | [K].ELENFFQELEKAR.[K] | 1xHomocysteineMMTS [K11] | 0,000133296 | 1815,84042 | Brain/ FAIMS/MMTS |
| Q08331 | Q08331 [246-258] | [K].SVMSLAEAGKLYR.[K] | 1xHomocysteineMMTS [K10] | 4,49774E-05 | 1587,76917 | Brain/ FAIMS/MMTS |
| Q08331 | Q08331 [233-244] | [K].KEMNIQQLTTYR.[K] | 1xHomocysteineMMTS [K1] | 0,000152766 | 1687,79644 | Brain/ FAIMS/MMTS |
| Q11011 | Q11011 [852-864] | [R].LIKLSVEGFAVDK.[M] | 1xHomocysteineIAA [K3] | 0,00559513 | 1592,87164 | Brain/ FAIMS/IAA |
| Q2PFD7 | Q2PFD7 [757-783] | [R].IGSTTNPFLDIPHDPNAAVYKSGFLAR.[K] | 1xHomocysteineIAA [K21] | 0,00948001 | 3075,53567 | Brain/ FAIMS/IAA |
| Q30KP6 | Q30KP6 [21-32] | [K].KSYPEYGSLDLR.[K] | 1xHomocysteineIAA [K1] | 0,00814371 | 1601,76282 | Liver/ FAIMS/IAA |
| Q3TCN2 | Q3TCN2 [189-204] | [R].LTLLQLKGLEDSYEGR.[L] | 1xHomocysteineIAA [K7] | 0,000876609 | 2009,0372 | Brain/ FAIMS/IAA |
| Q3TTY5;<br>Q6NXH9;<br>P50446;<br>Q8BGZ7;<br>P07744;<br>Q3UV17;<br>Q922U2;<br>P19246 | Q3TTY5 [503-511];<br>Q6NXH9 [435-443];<br>P50446 [456-464];<br>Q8BGZ7 [454-462];<br>P07744 [450-458];<br>Q3UV17 [471-479];<br>Q922U2 [466-474];<br>P19246 [402-410] | [R].KLLEGEECR.[MLI] | 1xCarbamidomethyl [C8];<br>1xHomocysteineIAA [K1] | 0,00286445 | 1307,60823 | Brain/ No FAIMS/IAA |
| Q3TTY5;<br>Q6NXH9;<br>P50446;<br>Q8BGZ7;<br>P07744;<br>Q3UV17;<br>Q922U2;<br>P19246 | Q3TTY5 [503-511];<br>Q6NXH9 [435-443];<br>P50446 [456-464];<br>Q8BGZ7 [454-462];<br>P07744 [450-458];<br>Q3UV17 [471-479];<br>Q922U2 [466-474];<br>P19246 [402-410] | [R].KLLEGEECR.[MLI] | 1xCarbamidomethyl [C8];<br>1xHomocysteineIAA [K1] | 0,00286445 | 1307,60823 | Brain/ FAIMS/IAA |
| Q3TTY5;<br>Q9R0H5;<br>Q6NXH9;<br>Q6IFZ9;<br>P50446;<br>P04104;<br>Q6IME9;<br>Q8BGZ7;<br>Q8VED5;<br>P07744;<br>P11679;<br>Q6IFZ6;<br>Q3UV17;<br>Q9DCV7;<br>Q922U2 | Q3TTY5 [210-218];<br>Q9R0H5 [142-150];<br>Q6NXH9 [142-150];<br>Q6IFZ9 [117-125];<br>P50446 [163-171];<br>P04104 [199-207];<br>Q6IME9 [145-153];<br>Q8BGZ7 [161-169];<br>Q8VED5 [150-158];<br>P07744 [157-165];<br>P11679 [108-116];<br>Q6IFZ6 [178-186];<br>Q3UV17 [176-184];<br>Q9DCV7 [96-104];<br>Q922U2 [173-181] | [K].FASFIDKVR.[F] | 1xHomocysteineMMTS [K7] | 0,00175939 | 1245,61186 | Liver/ No FAIMS/MMTS |
| Q3TTY5;<br>Q9R0H5;<br>Q6NXH9;<br>Q6IFZ9;<br>P50446;<br>P04104;<br>Q6IME9;<br>Q8BGZ7;<br>Q8VED5;<br>P07744;<br>P11679;<br>Q6IFZ6;<br>Q3UV17;<br>Q9DCV7;<br>Q922U2 | Q3TTY5 [210-218];<br>Q9R0H5 [142-150];<br>Q6NXH9 [142-150];<br>Q6IFZ9 [117-125];<br>P50446 [163-171];<br>P04104 [199-207];<br>Q6IME9 [145-153];<br>Q8BGZ7 [161-169];<br>Q8VED5 [150-158];<br>P07744 [157-165];<br>P11679 [108-116];<br>Q6IFZ6 [178-186];<br>Q3UV17 [176-184];<br>Q9DCV7 [96-104];<br>Q922U2 [173-181] | [K].FASFIDKVR.[F] | 1xHomocysteineMMTS [K7] | 0,00175939 | 1245,61186 | Liver/ FAIMS/MMTS |
| Q3TXX4 | Q3TXX4 [272-283] | [R].KYIEDAIGESAK.[L] | 1xHomocysteineIAA [K1] | 4,32935E-05 | 1497,72537 | Brain/ No FAIMS/IAA |
| Q3U0V1 | Q3U0V1 [192-206] | [R].SVSLTGAPESVQKAK.[M] | 1xHomocysteineIAA [K13] | 9,61721E-05 | 1675,86835 | Brain/ FAIMS/IAA |
| Q3U0V1 | Q3U0V1 [357-369] | [R].IQFKQDDGTGPEK.[I] | 1xHomocysteineMMTS [K4] | 4,49774E-05 | 1625,7298 | Liver/ FAIMS/MMTS |
| Q3UBX0 | Q3UBX0 [207-217] | [R].SSGSHLEAKVR.[G] | 1xHomocysteineMMTS [K9] | 0,00740868 | 1333,63512 | Liver/ FAIMS/MMTS |
| Q3UES3 | Q3UES3 [380-388] | [R].KQICELLLR.[K] | 1xCarbamidomethyl [C4];<br>1xHomocysteineIAA [K1] | 0,00340596 | 1346,72829 | Liver/ FAIMS/IAA |
| Q3UGC7 | Q3UGC7 [110-122] | [K].VLTPEEQLADKLR.[L] | 1xHomocysteineMMTS [K11] | 4,49774E-05 | 1674,85534 | Liver/ No FAIMS/MMTS |
| Q3UGC7 | Q3UGC7 [110-122] | [K].VLTPEEQLADKLR.[L] | 1xHomocysteineMMTS [K11] | 4,49774E-05 | 1674,85534 | Liver/ FAIMS/MMTS |
| Q3UHL1 | Q3UHL1 [69-78] | [K].NEIGILKMVK.[H] | 1xHomocysteineIAA [K7] | 4,32935E-05 | 1318,72214 | Brain/ FAIMS/IAA |
| Q3UI43 | Q3UI43 [122-137] | [R].TNALNVSQKMVEMFVR.[T] | 1xHomocysteineIAA [K9] | 0,00687906 | 2041,00275 | Brain/ FAIMS/IAA |
| Q3UL36 | Q3UL36 [165-176] | [R].IMEKQLLEELER.[Q] | 1xHomocysteineIAA [K4] | 0,000595249 | 1704,8659 | Brain/ FAIMS/IAA |
| Q3ULJ0 | Q3ULJ0 [123-139] | [K].GIDEGPDGLKLISDIIR.[E] | 1xHomocysteineIAA [K10] | 4,32935E-05 | 1985,0372 | Brain/ FAIMS/IAA |
| Q3UNH4 | Q3UNH4 [384-396] | [R].VSLVKTETLSSGK.[E] | 1xHomocysteineIAA [K5] | 4,32935E-05 | 1522,81452 | Brain/ FAIMS/IAA |
| Q3UNH4 | Q3UNH4 [384-396] | [R].VSLVKTETLSSGK.[E] | 1xHomocysteineMMTS [K5] | 0,000617167 | 1511,78078 | Brain/ FAIMS/MMTS |
| Q3UPI1 | Q3UPI1 [225-239] | [R].LLADSEVASILPISK.[S] | 1xHomocysteineMMTS [K15] | 0,00312202 | 1718,90671 | Brain/ No FAIMS/MMTS |
| Q3UPI1 | Q3UPI1 [225-239] | [R].LLADSEVASILPISK.[S] | 1xHomocysteineMMTS [K15] | 0,00312202 | 1718,90671 | Brain/ FAIMS/MMTS |
| Q3URK3 | Q3URK3 [1932-1944] | [K].SLNKPNHGFDINK.[I] | 1xHomocysteineMMTS [K4] | 0,00550312 | 1646,77776 | Liver/ FAIMS/MMTS |
| Q3UZ18 | Q3UZ18 [844-856] | [K].ISSLFNILQHILK.[K] | 1xHomocysteineMMTS [K13] | 0,00576292 | 1688,92263 | Brain/ No FAIMS/MMTS |
| Q3UZP4 | Q3UZP4 [55-73] | [K].EQLEKQMATSGPPTAGGLR.[W] | 1xHomocysteineIAA [K5] | 0,00883279 | 2145,0427 | Brain/ FAIMS/IAA |
| Q3V132;<br>P48962;<br>P51881 | Q3V132 [94-107];<br>P48962 [81-94]; P51881<br>[81-94] | [R].YFPTQALNFAFKDK.[Y] | 1xHomocysteineIAA [K12] | 4,32935E-05 | 1863,90982 | Brain/ FAIMS/IAA |
| Q4KMM3 | Q4KMM3 [89-105] | [R].MSFQKPKGTIEYTVESR.[D] | 1xHomocysteineIAA [K7] | 4,32935E-05 | 2175,05729 | Brain/ FAIMS/IAA |
| Q56A07 | Q56A07 [99-115] | [R].VEFSGNPSKYDVSVTLK.[N] | 1xHomocysteineIAA [K9] | 4,32935E-05 | 2044,00557 | Brain/ FAIMS/IAA |
| Q5DTL9 | Q5DTL9 [390-408] | [R].SIATLMTDEVFHDVAYKAK.[D] | 1xHomocysteineIAA [K17] | 4,32935E-05 | 2313,12536 | Brain/ FAIMS/IAA |
| Q5FW60 | Q5FW60 [28-48] | [R].NFNVEKINGEWYTIMLATDKR.[E] | 1xHomocysteineIAA [K6] | 4,32935E-05 | 2716,32217 | Liver/ No FAIMS/IAA |

| | | | | | | |
|---|---|---|---|---|---|---|
| Q5FW60 | Q5FW60 [28-48] | [R].NFNVEKINGEWYTIMLATDKR.[E] | 1xHomocysteineIAA [K6] | 4,32935E-05 | 2716,32217 | Liver/ FAIMS/IAA |
| Q5FWK3 | Q5FWK3 [405-421] | [K].DAAITLKAINPINTFTK.[F] | 1xHomocysteineIAA [K7] | 0,00490791 | 2005,07867 | Brain/ FAIMS/IAA |
| Q5GH66 | Q5GH66 [514-526] | [K].LAGSKADLAAQGK.[E] | 1xHomocysteineIAA [K5] | 0,00662819 | 1403,73112 | Liver/ FAIMS/IAA |
| Q5M8N0 | Q5M8N0 [40-51] | [K].LLTGSSYKVEVK.[I] | 1xHomocysteineIAA [K8] | 4,32935E-05 | 1497,79814 | Brain/ FAIMS/IAA |
| Q5QNQ9 | Q5QNQ9 [1243-1269] | [K].GAKGHQGYLGEMGIPGEPGPPGTPGPK.[G] | 1xOxidation [M12]; 1xHomocysteineIAA [K27] | 0,00619157 | 2776,31814 | Liver/ No FAIMS/IAA |
| Q5SQX6 | Q5SQX6 [771-785] | [R].ISAAMYKSLDQAISR.[F] | 1xHomocysteineIAA [K7] | 4,32935E-05 | 1827,90916 | Brain/ FAIMS/IAA |
| Q5SRX1 | Q5SRX1 [130-146] | [R].SSPDLTGVVHIYEELKR.[R] | 1xHomocysteineIAA [K16] | 4,32935E-05 | 2117,06956 | Brain/ FAIMS/IAA |
| Q5SYD0 | Q5SYD0 [804-814] | [K].VAAMEMLKGQR.[A] | 1xHomocysteineIAA [K8] | 0,0027732 | 1407,69052 | Brain/ FAIMS/IAA |
| Q5XG73 | Q5XG73 [444-456] | [R].LHKLETLTASQAK.[L] | 1xHomocysteineMMTS [K3] | 0,00995323 | 1602,83421 | Liver/ FAIMS/MMTS |
| Q60597 | Q60597 [901-914] | [R].LLFCTGKVYYDLTR.[E] | 1xCarbamidomethyl [C4]; 1xHomocysteineIAA [K7] | 4,32935E-05 | 1922,9503 | Brain/ FAIMS/IAA |
| Q60598 | Q60598 [153-168] | [K].DYSSGFGGKYGVQADR.[V] | 1xHomocysteineIAA [K9] | 4,32935E-05 | 1880,82319 | Brain/ FAIMS/IAA |
| Q60605 | Q60605 [80-94] | [K].NKDQGTYEDYVEGLR.[V] | 1xHomocysteineIAA [K2] | 4,32935E-05 | 1960,87053 | Liver/ FAIMS/IAA |
| Q60605 | Q60605 [80-94] | [K].NKDQGTYEDYVEGLR.[V] | 1xHomocysteineIAA [K2] | 4,32935E-05 | 1960,87053 | Brain/ FAIMS/IAA |
| Q60605 | Q60605 [80-94] | [K].NKDQGTYEDYVEGLR.[V] | 1xHomocysteineMMTS [K2] | 4,49774E-05 | 1949,83679 | Liver/ FAIMS/MMTS |
| Q60605 | Q60605 [51-63] | [K].VLGNPKSDEMNVK.[V] | 1xHomocysteineMMTS [K6] | 0,000192903 | 1593,74335 | Liver/ FAIMS/MMTS |
| Q60605 | Q60605 [95-110] | [R].VFDKEGNGTVMGAEIR.[H] | 1xHomocysteineMMTS [K4] | 0,000599431 | 1885,8605 | Liver/ FAIMS/MMTS |
| Q60605 | Q60605 [51-63] | [K].VLGNPKSDEMNVK.[V] | 1xHomocysteineMMTS [K6] | 0,000192903 | 1593,74335 | Brain/ FAIMS/MMTS |
| Q60605 | Q60605 [80-94] | [K].NKDQGTYEDYVEGLR.[V] | 1xHomocysteineMMTS [K2] | 4,49774E-05 | 1949,83679 | Brain/ FAIMS/MMTS |
| Q60625 | Q60625 [592-604] | [K].LFSCEVDGKPEPR.[V] | 1xCarbamidomethyl [C4]; 1xHomocysteineIAA [K9] | 4,32935E-05 | 1707,7829 | Brain/ FAIMS/IAA |
| Q60631 | Q60631 [7-20] | [K].YDFKATADDELSFK.[R] | 1xHomocysteineIAA [K4] | 4,32935E-05 | 1823,81564 | Brain/ FAIMS/IAA |
| Q60634 | Q60634 [156-169] | [K].DVYDKVDYLSSLGK.[T] | 1xHomocysteineIAA [K5] | 9,61721E-05 | 1775,85203 | Brain/ FAIMS/IAA |
| Q60668 | Q60668 [184-199] | [K].IFVGGLSPDTPEEKIR.[E] | 1xHomocysteineIAA [K14] | 4,32935E-05 | 1931,98952 | Brain/ FAIMS/IAA |
| Q60771 | Q60771 [190-207] | [R].FYYSSGSSSPTHAKSAHV.[-] | 1xHomocysteineIAA [K14] | 4,32935E-05 | 2086,92871 | Brain/ No FAIMS/IAA |
| Q60771 | Q60771 [190-207] | [R].FYYSSGSSSPTHAKSAHV.[-] | 1xHomocysteineIAA [K14] | 4,32935E-05 | 2086,92871 | Brain/ FAIMS/IAA |
| Q60829 | Q60829 [108-139] | [R].ELGYPQEDDEEDEDEEEDEEEDSQAEVLKGSR.[G] | 1xHomocysteineIAA [K29] | 4,32935E-05 | 3916,54687 | Brain/ FAIMS/IAA |
| Q60864 | Q60864 [207-223] | [K].KEPKPEPMEEDLPENKK.[Q] | 1xHomocysteineIAA [K] | 4,32935E-05 | 2212,06243 | Liver/ FAIMS/IAA |
| Q60864 | Q60864 [207-222] | [K].KEPKPEPMEEDLPENK.[K] | 1xHomocysteineIAA [K] | 4,32935E-05 | 2083,96747 | Brain/ FAIMS/IAA |
| Q60864 | Q60864 [154-169] | [R].ELIEQLQNKPSDLGTK.[L] | 1xHomocysteineMMTS [K9] | 0,000266738 | 1975,98272 | Liver/ FAIMS/MMTS |
| Q60864 | Q60864 [524-533] | [K].NPVIAQKIQK.[L] | 1xHomocysteineMMTS [K7] | 0,00503864 | 1301,70682 | Liver/ FAIMS/MMTS |
| Q60864 | Q60864 [531-543] | [K].IQKLMDVGLIAIR.[-] | 1xHomocysteineMMTS [K3] | 4,49774E-05 | 1632,89979 | Liver/ FAIMS/MMTS |
| Q60864 | Q60864 [154-169] | [R].ELIEQLQNKPSDLGTK.[L] | 1xHomocysteineMMTS [K9] | 0,000266738 | 1975,98272 | Brain/ FAIMS/MMTS |
| Q60864 | Q60864 [524-533] | [K].NPVIAQKIQK.[L] | 1xHomocysteineMMTS [K7] | 0,00503864 | 1301,70682 | Brain/ FAIMS/MMTS |
| Q60864 | Q60864 [531-543] | [K].IQKLMDVGLIAIR.[-] | 1xHomocysteineMMTS [K3] | 4,49774E-05 | 1632,89979 | Brain/ FAIMS/MMTS |
| Q60870 | Q60870 [144-158] | [R].HESQVDSVVKDVKDK.[A] | 1xHomocysteineIAA [K13] | 0,00399809 | 1886,92765 | Brain/ FAIMS/IAA |
| Q60902 | Q60902 [466-479] | [R].QKCQDETQTISSLK.[T] | 1xCarbamidomethyl [C3]; 1xHomocysteineIAA [K2] | 0,00012645 | 1839,85752 | Brain/ FAIMS/IAA |
| Q60930 | Q60930 [237-268] | [K].YQLDPTASISAKVNNSSLIGVGYTQTLRPGVK.[L] | 1xHomocysteineIAA [K12] | 4,32935E-05 | 3551,85264 | Brain/ FAIMS/IAA |
| Q60930 | Q60930 [231-248] | [R].FGIAAKYQLDPTASISAK.[V] | 1xHomocysteineMMTS [K6] | 4,49774E-05 | 2044,02419 | Brain/ FAIMS/MMTS |
| Q60931 | Q60931 [257-274] | [K].LTLSALIDGKNFNAGGHK.[V] | 1xHomocysteineIAA [K10] | 4,32935E-05 | 2030,04877 | Brain/ FAIMS/IAA |
| Q60931 | Q60931 [16-28] | [K].DVFNKGYGFGMVK.[I] | 1xHomocysteineMMTS [K5] | 0,00166208 | 1624,73205 | Brain/ FAIMS/MMTS |
| Q60932 | Q60932 [107-122] | [R].GLKLTFDSSFSPNTGK.[K] | 1xHomocysteineIAA [K3] | 4,32935E-05 | 1872,91602 | Brain/ No FAIMS/IAA |
| Q60932 | Q60932 [107-122] | [R].GLKLTFDSSFSPNTGK.[K] | 1xHomocysteineIAA [K3] | 4,32935E-05 | 1872,91602 | Brain/ FAIMS/IAA |
| Q60932 | Q60932 [29-41] | [R].DVFTKGYGFGLIK.[L] | 1xHomocysteineMMTS [K5] | 0,00137282 | 1607,79603 | Brain/ No FAIMS/MMTS |
| Q60932 | Q60932 [29-41] | [R].DVFTKGYGFGLIK.[L] | 1xHomocysteineMMTS [K5] | 0,00137282 | 1607,79603 | Brain/ FAIMS/MMTS |
| Q60932 | Q60932 [107-122] | [R].GLKLTFDSSFSPNTGK.[K] | 1xHomocysteineMMTS [K3] | 0,000302483 | 1861,88228 | Brain/ FAIMS/MMTS |
| Q60932 | Q60932 [110-123] | [K].LTFDSSFSPNTGKK.[N] | 1xHomocysteineMMTS [K13] | 4,49774E-05 | 1691,77675 | Brain/ FAIMS/MMTS |
| Q61016 | Q61016 [28-44] | [K].VSKASSDLMGYCEQHAR.[N] | 1xCarbamidomethyl [C12]; 1xHomocysteineIAA [K3] | 4,32935E-05 | 2112,92595 | Brain/ FAIMS/IAA |
| Q61081 | Q61081 [110-122] | [K].EKNMPWNVDTLSK.[D] | 1xHomocysteineIAA [K2] | 4,32935E-05 | 1735,8142 | Brain/ FAIMS/IAA |
| Q61166 | Q61166 [169-181] | [R].TTAAPKAGPGMVR.[K] | 1xHomocysteineIAA [K6] | 0,000177335 | 1430,72426 | Brain/ FAIMS/IAA |
| Q61166; Q8R001 | Q61166 [215-222]; Q8R001 [265-272] | [R].DFYFGKLR.[NE] | 1xHomocysteineIAA [K6] | 0,000258769 | 1219,59284 | Brain/ FAIMS/IAA |
| Q61171 | Q61171 [92-109] | [R].KEGGLGPLNIPLLADVTK.[S] | 1xHomocysteineIAA [K1] | 4,32935E-05 | 2009,10997 | Liver/ FAIMS/IAA |
| Q61171 | Q61171 [67-91] | [R].KLGCEVLGVSVDSQFTHLAWINTPR.[K] | 1xCarbamidomethyl [C4]; 1xHomocysteineIAA [K1] | 4,32935E-05 | 3001,50226 | Brain/ FAIMS/IAA |
| Q61171 | Q61171 [11-29] | [K].SAPDFTATAVVDGAFKEIK.[L] | 1xHomocysteineMMTS [K16] | 0,000266738 | 2130,02459 | Liver/ No FAIMS/MMTS |
| Q61171 | Q61171 [92-109] | [R].KEGGLGPLNIPLLADVTK.[S] | 1xHomocysteineMMTS [K1] | 4,49774E-05 | 1998,07623 | Liver/ No FAIMS/MMTS |
| Q61171 | Q61171 [128-139] | [R].GLFIIDAKGVLR.[Q] | 1xHomocysteineMMTS [K8] | 0,000211896 | 1464,80654 | Liver/ FAIMS/MMTS |
| Q61171 | Q61171 [92-109] | [R].KEGGLGPLNIPLLADVTK.[S] | 1xHomocysteineMMTS [K1] | 4,49774E-05 | 1998,07623 | Liver/ FAIMS/MMTS |
| Q61171 | Q61171 [11-29] | [K].SAPDFTATAVVDGAFKEIK.[L] | 1xHomocysteineMMTS [K16] | 0,000266738 | 2130,02459 | Liver/ FAIMS/MMTS |
| Q61171 | Q61171 [110-127] | [K].SLSQNYGVLKNDEGIAYR.[G] | 1xHomocysteineMMTS [K10] | 4,49774E-05 | 2190,0318 | Liver/ FAIMS/MMTS |
| Q61171 | Q61171 [128-139] | [R].GLFIIDAKGVLR.[Q] | 1xHomocysteineMMTS [K8] | 0,000211896 | 1464,80654 | Brain/ FAIMS/MMTS |
| Q61171 | Q61171 [110-127] | [K].SLSQNYGVLKNDEGIAYR.[G] | 1xHomocysteineMMTS [K10] | 4,49774E-05 | 2190,0318 | Brain/ FAIMS/MMTS |
| Q61171 | Q61171 [92-109] | [R].KEGGLGPLNIPLLADVTK.[S] | 1xHomocysteineMMTS [K1] | 4,49774E-05 | 1998,07623 | Brain/ FAIMS/MMTS |
| Q61176 | Q61176 [285-308] | [K].TAEEVKSTVNTAVALTLACFGTQR.[E] | 1xCarbamidomethyl [C19]; 1xHomocysteineIAA [K6] | 4,32935E-05 | 2741,35967 | Liver/ FAIMS/IAA |
| Q61176 | Q61176 [49-71] | [R].DHGDLAFVDVPNDSSFQIVKNPR.[S] | 1xHomocysteineMMTS [K20] | 4,49774E-05 | 2733,27594 | Liver/ No FAIMS/MMTS |
| Q61176 | Q61176 [309-323] | [R].EGNHKPGTDYLKPPK.[-] | 1xHomocysteineMMTS [K] | 0,000599431 | 1843,88295 | Liver/ No FAIMS/MMTS |
| Q61176 | Q61176 [267-290] | [K].TGLLSGLDIMEVNPTLGKTAEEVK.[S] | 1xOxidation [M10]; 1xHomocysteineMMTS [K18] | 4,49774E-05 | 2694,33985 | Liver/ No FAIMS/MMTS |
| Q61176 | Q61176 [1-17] | [-].MSSKPKSLEIIGAPFSK.[G] | 1xMet-loss+Acetyl [N-Term]; 1xHomocysteineMMTS [K6] | 0,000250158 | 1893,98127 | Liver/ No FAIMS/MMTS |
| Q61176 | Q61176 [22-32] | [R].GGVEKGPAALR.[K] | 1xHomocysteineMMTS [K5] | 4,49774E-05 | 1217,61292 | Liver/ No FAIMS/MMTS |
| Q61176 | Q61176 [72-89] | [R].SVGKANEELAGVVAEVQK.[N] | 1xHomocysteineMMTS [K4] | 4,49774E-05 | 1990,99362 | Liver/ No FAIMS/MMTS |
| Q61176 | Q61176 [40-48] | [K].LKETEYDVR.[D] | 1xHomocysteineMMTS [K2] | 0,000192903 | 1315,60208 | Liver/ No FAIMS/MMTS |
| Q61176 | Q61176 [267-290] | [K].TGLLSGLDIMEVNPTLGKTAEEVK.[S] | 1xHomocysteineMMTS [K18] | 4,49774E-05 | 2678,34494 | Liver/ No FAIMS/MMTS |
| Q61176 | Q61176 [181-196] | [R].DVDPGEHYIIKTLGIK.[Y] | 1xHomocysteineMMTS [K11] | 4,49774E-05 | 1960,98708 | Liver/ No FAIMS/MMTS |

| | | | | | | |
|---|---|---|---|---|---|---|
| Q61176 | Q61176 [7-21] | [K].SLEIIGAPFSKGQPR.[G] | 1xHomocysteineMMTS [K11] | 4,49774E-05 | 1762,89787 | Liver/ No FAIMS/MMTS |
| Q61176 | Q61176 [33-39] | [R].KAGLLEK.[L] | 1xHomocysteineMMTS [K1] | 0,00161608 | 921,48962 | Liver/ No FAIMS/MMTS |
| Q61176 | Q61176 [197-210] | [K].YFSMTEVDKLGIGK.[V] | 1xOxidation [M4]; 1xHomocysteineMMTS [K9] | 4,49774E-05 | 1766,81618 | Liver/ FAIMS/MMTS |
| Q61176 | Q61176 [1-17] | [-].MSSKPKSLEIIGAPFSK.[G] | 1xMet-loss+Acetyl [N-Term]; 1xHomocysteineMMTS [K6] | 0,000250158 | 1893,98127 | Liver/ FAIMS/MMTS |
| Q61176 | Q61176 [192-205] | [K].TLGIKYFSMTEVDK.[L] | 1xHomocysteineMMTS [K5] | 8,93346E-05 | 1794,84748 | Liver/ FAIMS/MMTS |
| Q61176 | Q61176 [22-32] | [R].GGVEKGPAALR.[K] | 1xHomocysteineMMTS [K5] | 4,49774E-05 | 1217,61292 | Liver/ FAIMS/MMTS |
| Q61176 | Q61176 [72-89] | [R].SVGKANEELAGVVAEVQK.[N] | 1xHomocysteineMMTS [K4] | 4,49774E-05 | 1990,99362 | Liver/ FAIMS/MMTS |
| Q61176 | Q61176 [49-71] | [R].DHGDLAFVDVPNDSSFQIVKNPR.[S] | 1xHomocysteineMMTS [K20] | 4,49774E-05 | 2733,27594 | Liver/ FAIMS/MMTS |
| Q61176 | Q61176 [40-48] | [K].LKETEYDVR.[D] | 1xHomocysteineMMTS [K2] | 0,000192903 | 1315,60208 | Liver/ FAIMS/MMTS |
| Q61176 | Q61176 [181-196] | [R].DVDPGEHYIIKTLGIK.[Y] | 1xHomocysteineMMTS [K11] | 4,49774E-05 | 1960,98708 | Liver/ FAIMS/MMTS |
| Q61176 | Q61176 [7-21] | [K].SLEIIGAPFSKGQPR.[G] | 1xHomocysteineMMTS [K11] | 4,49774E-05 | 1762,89787 | Liver/ FAIMS/MMTS |
| Q61176 | Q61176 [192-205] | [K].TLGIKYFSMTEVDK.[L] | 1xOxidation [M9]; 1xHomocysteineMMTS [K5] | 0,000152766 | 1810,84239 | Liver/ FAIMS/MMTS |
| Q61176 | Q61176 [206-222] | [K].LGIGKVMEETFSYLLGR.[K] | 1xOxidation [M7]; 1xHomocysteineMMTS [K5] | 4,49774E-05 | 2092,02757 | Liver/ FAIMS/MMTS |
| Q61176 | Q61176 [267-290] | [K].TGLLSGLDIMEVNPTLGKTAEEVK.[S] | 1xOxidation [M10]; 1xHomocysteineMMTS [K18] | 4,49774E-05 | 2694,33985 | Liver/ FAIMS/MMTS |
| Q61176 | Q61176 [197-210] | [K].YFSMTEVDKLGIGK.[V] | 1xHomocysteineMMTS [K9] | 4,49774E-05 | 1750,82126 | Liver/ FAIMS/MMTS |
| Q61176 | Q61176 [267-290] | [K].TGLLSGLDIMEVNPTLGKTAEEVK.[S] | 1xHomocysteineMMTS [K18] | 4,49774E-05 | 2678,34494 | Liver/ FAIMS/MMTS |
| Q61176 | Q61176 [309-323] | [R].EGNHKPGTDYLKPPK.[-] | 1xHomocysteineMMTS [K] | 0,000599431 | 1843,88295 | Liver/ FAIMS/MMTS |
| Q61206 | Q61206 [129-142] | [R].QPQAKIIVLGLLPR.[G] | 1xHomocysteineIAA [K5] | 0,00918796 | 1720,03021 | Brain/ No FAIMS/IAA |
| Q61207 | Q61207 [447-458] | [K].KLVLYLEHNLEK.[N] | 1xHomocysteineIAA [K1] | 4,32935E-05 | 1672,90909 | Brain/ FAIMS/IAA |
| Q61282 | Q61282 [552-565] | [R].TPCVGDKDSSPGVR.[T] | 1xCarbamidomethyl [C3]; 1xHomocysteineIAA [K7] | 0,00012645 | 1648,74177 | Brain/ FAIMS/IAA |
| Q61292 | Q61292 [1741-1752] | [R].DLLQAAQDKLQR.[L] | 1xHomocysteineIAA [K9] | 4,32935E-05 | 1572,81625 | Brain/ FAIMS/IAA |
| Q61316 | Q61316 [54-68] | [K].SQVISNAKNTVQGFK.[R] | 1xHomocysteineIAA [K8] | 4,32935E-05 | 1794,91669 | Brain/ FAIMS/IAA |
| Q61330 | Q61330 [816-824] | [K].VAPAKVWAK.[G] | 1xHomocysteineIAA [K5] | 0,00314066 | 1143,63431 | Brain/ FAIMS/IAA |
| Q61335 | Q61335 [214-224] | [K].QSEGLTKEYDR.[L] | 1xHomocysteineIAA [K7] | 0,000318237 | 1499,67948 | Liver/ FAIMS/IAA |
| Q61335 | Q61335 [225-237] | [R].LLEEHAKLQASVR.[G] | 1xHomocysteineIAA [K7] | 4,32935E-05 | 1667,88975 | Brain/ FAIMS/IAA |
| Q61335 | Q61335 [204-220] | [K].AENEALAMQKQSEGLTK.[E] | 1xOxidation [M8]; 1xHomocysteineMMTS [K10] | 4,49774E-05 | 2026,92422 | Liver/ FAIMS/MMTS |
| Q61335 | Q61335 [189-198] | [R].KLKDELASTK.[K] | 1xHomocysteineMMTS [K3] | 0,00213323 | 1295,66977 | Liver/ FAIMS/MMTS |
| Q61361 | Q61361 [454-474] | [R].FKDLEALEEEKEQEDLWVWPR.[E] | 1xHomocysteineIAA [K] | 4,32935E-05 | 2863,36072 | Brain/ No FAIMS/IAA |
| Q61361 | Q61361 [454-474] | [R].FKDLEALEEEKEQEDLWVWPR.[E] | 1xHomocysteineIAA [K] | 4,32935E-05 | 2863,36072 | Brain/ FAIMS/IAA |
| Q61361 | Q61361 [582-593] | [R].ELETPSEEKSGR.[T] | 1xHomocysteineMMTS [K9] | 0,00325546 | 1524,66687 | Brain/ FAIMS/MMTS |
| Q61414; P19001; Q9QWL7 | Q61414 [109-117]; P19001 [94-102]; Q9QWL7 [95-103] | [R].LASYLDKVR.[A] | 1xHomocysteineIAA [K7] | 0,00111602 | 1238,65617 | Brain/ FAIMS/IAA |
| Q61425 | Q61425 [237-249] | [R].GDASKEDIDTAMK.[L] | 1xHomocysteineIAA [K5] | 0,00012645 | 1554,67743 | Liver/ FAIMS/IAA |
| Q61425 | Q61425 [180-192] | [K].LVEVIKTPMTSQK.[T] | 1xHomocysteineMMTS [K6] | 4,49774E-05 | 1636,84708 | Liver/ FAIMS/MMTS |
| Q61425 | Q61425 [180-192] | [K].LVEVIKTPMTSQK.[T] | 1xOxidation [M9]; 1xHomocysteineMMTS [K6] | 0,000152766 | 1652,842 | Liver/ FAIMS/MMTS |
| Q61425 | Q61425 [237-249] | [R].GDASKEDIDTAMK.[L] | 1xOxidation [M12]; 1xHomocysteineMMTS [K5] | 0,000437433 | 1559,63861 | Liver/ FAIMS/MMTS |
| Q61425 | Q61425 [237-249] | [R].GDASKEDIDTAMK.[L] | 1xHomocysteineMMTS [K5] | 4,49774E-05 | 1543,64369 | Liver/ FAIMS/MMTS |
| Q61490 | Q61490 [375-404] | [R].SSPSFSSLHYQDAGNYVCETALQEVEGLK K.[R] | 1xCarbamidomethyl [C18]; 1xHomocysteineIAA [K29] | 4,32935E-05 | 3518,62026 | Brain/ FAIMS/IAA |
| Q61548 | Q61548 [101-125] | [R].NTLFNLSNFLDKSGSHGYDMSTFIR.[R] | 1xHomocysteineIAA [K12] | 4,32935E-05 | 3038,4135 | Brain/ FAIMS/IAA |
| Q61553 | Q61553 [69-82] | [R].YLAADKDGNVTCER.[E] | 1xCarbamidomethyl [C12]; 1xHomocysteineIAA [K6] | 4,32935E-05 | 1785,78944 | Brain/ FAIMS/IAA |
| Q61598 | Q61598 [265-276] | [K].VVGVKSEGEIAR.[C] | 1xHomocysteineIAA [K5] | 4,32935E-05 | 1417,74677 | Brain/ No FAIMS/IAA |
| Q61598 | Q61598 [265-276] | [K].VVGVKSEGEIAR.[C] | 1xHomocysteineIAA [K5] | 4,32935E-05 | 1417,74677 | Brain/ FAIMS/IAA |
| Q61598 | Q61598 [265-276] | [K].VVGVKSEGEIAR.[C] | 1xHomocysteineMMTS [K5] | 0,000863044 | 1406,71303 | Liver/ FAIMS/MMTS |
| Q61644 | Q61644 [121-130] | [K].QIMGGFKETK.[E] | 1xHomocysteineIAA [K7] | 4,32935E-05 | 1312,6388 | Brain/ No FAIMS/IAA |
| Q61644 | Q61644 [121-130] | [K].QIMGGFKETK.[E] | 1xHomocysteineIAA [K7] | 4,32935E-05 | 1312,6388 | Brain/ FAIMS/IAA |
| Q61644 | Q61644 [244-251] | [K].EVLLDIKR.[H] | 1xHomocysteineMMTS [K7] | 0,000530332 | 1148,61661 | Brain/ No FAIMS/MMTS |
| Q61644 | Q61644 [66-79] | [R].QLIEKGPQYGSLER.[A] | 1xHomocysteineMMTS [K5] | 4,49774E-05 | 1780,87205 | Brain/ FAIMS/MMTS |
| Q61644 | Q61644 [171-186] | [R].EMNSKTEQSVTPEQQK.[K] | 1xHomocysteineMMTS [K5] | 0,000653201 | 2026,88784 | Brain/ FAIMS/MMTS |
| Q61644 | Q61644 [49-62] | [K].IEKAYAQQLTDWAK.[R] | 1xHomocysteineMMTS [K3] | 4,49774E-05 | 1827,8768 | Brain/ FAIMS/MMTS |
| Q61644 | Q61644 [80-99] | [R].AWGAMMTEADKVSELHQEVK.[N] | 1xHomocysteineMMTS [K11] | 8,93346E-05 | 2423,08622 | Brain/ FAIMS/MMTS |
| Q61644 | Q61644 [80-109] | [R].AWGAMMTEADKVSELHQEVKNSLLNE DLEK.[V] | 1xHomocysteineMMTS [K11] | 0,00995323 | 3578,66339 | Brain/ FAIMS/MMTS |
| Q61644 | Q61644 [319-341] | [K].KAEGATLSNATGAVESTSQAGDR.[G] | 1xHomocysteineMMTS [K1] | 4,49774E-05 | 2384,08166 | Brain/ FAIMS/MMTS |
| Q61646 | Q61646 [103-119] | [R].IIGGSMDAKGSFPWQAK.[M] | 1xHomocysteineIAA [K9] | 4,32935E-05 | 1966,95136 | Liver/ FAIMS/IAA |
| Q61646 | Q61646 [103-119] | [R].IIGGSMDAKGSFPWQAK.[M] | 1xHomocysteineIAA [K9] | 4,32935E-05 | 1966,95136 | Brain/ FAIMS/IAA |
| Q61646 | Q61646 [333-346] | [R].ATDLKDWVQETMAK.[N] | 1xHomocysteineMMTS [K5] | 4,49774E-05 | 1798,81724 | Liver/ No FAIMS/MMTS |
| Q61646 | Q61646 [103-119] | [R].IIGGSMDAKGSFPWQAK.[M] | 1xOxidation [M6]; 1xHomocysteineMMTS [K9] | 0,000706375 | 1971,91254 | Liver/ FAIMS/MMTS |
| Q61646 | Q61646 [103-119] | [R].IIGGSMDAKGSFPWQAK.[M] | 1xHomocysteineMMTS [K9] | 0,000173718 | 1955,91762 | Liver/ FAIMS/MMTS |
| Q61646 | Q61646 [333-346] | [R].ATDLKDWVQETMAK.[N] | 1xHomocysteineMMTS [K5] | 4,49774E-05 | 1798,81724 | Liver/ FAIMS/MMTS |
| Q61696; P63017; P16627; P17156 | Q61696 [501-509]; P63017 [501-509]; P16627 [503-511]; P17156 [504-512] | [K].ITITNDKGR.[L] | 1xHomocysteineIAA [K7] | 6,30912E-05 | 1191,61503 | Liver/ FAIMS/IAA |
| Q61696; P63017; P16627; P17156 | Q61696 [501-509]; P63017 [501-509]; P16627 [503-511]; P17156 [504-512] | [K].ITITNDKGR.[L] | 1xHomocysteineIAA [K7] | 6,30912E-05 | 1191,61503 | Brain/ FAIMS/IAA |

| | | | | | | |
|---|---|---|---|---|---|---|
| Q61696; P63017; P16627; P17156 | Q61696 [501-509]; P63017 [501-509]; P16627 [503-511]; P17156 [504-512] | [K].ITITNDKGR.[L] | 1xHomocysteineMMTS [K7] | 6,70238E-05 | 1180,58129 | Liver/ No FAIMS/MMTS |
| Q61696; P63017; P16627; P17156 | Q61696 [501-509]; P63017 [501-509]; P16627 [503-511]; P17156 [504-512] | [K].ITITNDKGR.[L] | 1xHomocysteineMMTS [K7] | 6,70238E-05 | 1180,58129 | Liver/ FAIMS/MMTS |
| Q61696; P63017; P16627; P17156 | Q61696 [501-509]; P63017 [501-509]; P16627 [503-511]; P17156 [504-512] | [K].ITITNDKGR.[L] | 1xHomocysteineMMTS [K7] | 6,70238E-05 | 1180,58129 | Brain/ No FAIMS/MMTS |
| Q61696; P63017; P16627; P17156 | Q61696 [501-509]; P63017 [501-509]; P16627 [503-511]; P17156 [504-512] | [K].ITITNDKGR.[L] | 1xHomocysteineMMTS [K7] | 6,70238E-05 | 1180,58129 | Brain/ FAIMS/MMTS |
| Q61699 | Q61699 [220-234] | [K].LKVLGTAFDPFLGGK.[N] | 1xHomocysteineIAA [K2] | 0,00012645 | 1736,94039 | Brain/ FAIMS/IAA |
| Q61699; Q61316; P48722 | Q61699 [622-630]; Q61316 [605-613]; P48722 [607-615] | [K].MIMQDKLEK.[E] | 1xHomocysteineIAA [K6] | 0,000273428 | 1309,63128 | Brain/ FAIMS/IAA |
| Q61735 | Q61735 [45-59] | [R].NVEAQSTEEMFVKWK.[L] | 1xHomocysteineIAA [K13] | 4,32935E-05 | 1999,92521 | Brain/ FAIMS/IAA |
| Q61753 | Q61753 [381-394] | [R].EASKQADVNLVNAK.[L] | 1xHomocysteineIAA [K4] | 0,00927232 | 1660,83229 | Brain/ FAIMS/IAA |
| Q61768; P28738; P33175 | Q61768 [241-256]; P28738 [242-257]; P33175 [242-257] | [K].TGAEGAVLDEAKNINK.[S] | 1xHomocysteineIAA [K12] | 4,32935E-05 | 1803,89054 | Brain/ FAIMS/IAA |
| Q61792 | Q61792 [74-85] | [R].LKQQSELQSQVR.[Y] | 1xHomocysteineIAA [K2] | 4,32935E-05 | 1617,83771 | Liver/ FAIMS/IAA |
| Q61792 | Q61792 [112-121] | [K].KTQDQISNIK.[Y] | 1xHomocysteineIAA [K1] | 4,32935E-05 | 1348,68893 | Brain/ No FAIMS/IAA |
| Q61792 | Q61792 [112-121] | [K].KTQDQISNIK.[Y] | 1xHomocysteineIAA [K1] | 4,32935E-05 | 1348,68893 | Brain/ FAIMS/IAA |
| Q61792 | Q61792 [74-85] | [R].LKQQSELQSQVR.[Y] | 1xHomocysteineMMTS [K2] | 4,49774E-05 | 1606,80397 | Liver/ FAIMS/MMTS |
| Q61792 | Q61792 [74-85] | [R].LKQQSELQSQVR.[Y] | 1xHomocysteineMMTS [K2] | 4,49774E-05 | 1606,80397 | Brain/ FAIMS/MMTS |
| Q61838 | Q61838 [899-913] | [K].LTEVPALVHKDTVVK.[S] | 1xHomocysteineIAA [K10] | 0,000114608 | 1823,00953 | Liver/ FAIMS/IAA |
| Q61838 | Q61838 [1176-1188] | [R].SELLESLNKDAVK.[E] | 1xHomocysteineMMTS [K9] | 0,00185311 | 1608,79715 | Liver/ FAIMS/MMTS |
| Q61838 | Q61838 [1014-1026] | [K].SKAINYLISGYQR.[Q] | 1xHomocysteineMMTS [K2] | 0,00166208 | 1675,82946 | Liver/ FAIMS/MMTS |
| Q61879 | Q61879 [1397-1407] | [K].LLKDVEALSQR.[L] | 1xHomocysteineIAA [K3] | 0,000243434 | 1445,77807 | Brain/ FAIMS/IAA |
| Q61885 | Q61885 [95-108] | [R].GRTELLKETISEGK.[V] | 1xHomocysteineIAA [K7] | 4,32935E-05 | 1734,90546 | Brain/ No FAIMS/IAA |
| Q61885 | Q61885 [95-108] | [R].GRTELLKETISEGK.[V] | 1xHomocysteineIAA [K7] | 4,32935E-05 | 1734,90546 | Brain/ FAIMS/IAA |
| Q61885 | Q61885 [102-112] | [K].ETISEGKVTLR.[I] | 1xHomocysteineMMTS [K7] | 4,49774E-05 | 1395,69705 | Brain/ FAIMS/MMTS |
| Q61885 | Q61885 [81-94] | [R].NGKDQDAEQAPEYR.[G] | 1xHomocysteineMMTS [K3] | 0,00294218 | 1783,73741 | Brain/ FAIMS/MMTS |
| Q61937 | Q61937 [238-248] | [K].GPSSVEDIKAK.[M] | 1xHomocysteineIAA [K] | 6,30912E-05 | 1304,65148 | Liver/ FAIMS/IAA |
| Q61937 | Q61937 [238-248] | [K].GPSSVEDIKAK.[M] | 1xHomocysteineIAA [K] | 6,30912E-05 | 1304,65148 | Brain/ No FAIMS/IAA |
| Q61937 | Q61937 [238-248] | [K].GPSSVEDIKAK.[M] | 1xHomocysteineIAA [K] | 6,30912E-05 | 1304,65148 | Brain/ FAIMS/IAA |
| Q61990; P60335; P57722 | Q61990 [18-31]; P60335 [18-31]; P57722 [50-63] | [R].LLMHGKEVGSIIGK.[K] | 1xHomocysteineIAA [K6] | 4,32935E-05 | 1655,89714 | Liver/ FAIMS/IAA |
| Q61990; P60335; P57722 | Q61990 [18-31]; P60335 [18-31]; P57722 [50-63] | [R].LLMHGKEVGSIIGK.[K] | 1xHomocysteineIAA [K6] | 4,32935E-05 | 1655,89714 | Brain/ FAIMS/IAA |
| Q61990; P60335; P57722 | Q61990 [18-31]; P60335 [18-31]; P57722 [50-63] | [R].LLMHGKEVGSIIGK.[K] | 1xHomocysteineMMTS [K6] | 0,00230389 | 1644,8634 | Liver/ FAIMS/MMTS |
| Q61990; P60335; P57722 | Q61990 [18-31]; P60335 [18-31]; P57722 [50-63] | [R].LLMHGKEVGSIIGK.[K] | 1xHomocysteineMMTS [K6] | 0,00230389 | 1644,8634 | Brain/ FAIMS/MMTS |
| Q62048 | Q62048 [89-101] | [K].ISEEEELDTKLTR.[I] | 1xHomocysteineIAA [K10] | 4,32935E-05 | 1736,83711 | Brain/ FAIMS/IAA |
| Q62059 | Q62059 [215-228] | [R].EGCYGDMMGKEGVR.[T] | 1xCarbamidomethyl [C3]; 1xHomocysteineIAA [K10] | 4,32935E-05 | 1762,70156 | Brain/ FAIMS/IAA |
| Q62188 | Q62188 [401-423] | [R].IAVGSDSDLVIWDPDALKIVSAK.[N] | 1xHomocysteineIAA [K18] | 0,00012645 | 2586,34836 | Brain/ FAIMS/IAA |
| Q62189 | Q62189 [77-89] | [R].SMQGFPFYDKPMR.[I] | 1xHomocysteineIAA [K10] | 8,35381E-05 | 1777,78588 | Brain/ FAIMS/IAA |
| Q62261 | Q62261 [329-348] | [R].KFANSLVGVQQQLQAFNTYR.[T] | 1xHomocysteineIAA [K1] | 4,32935E-05 | 2486,26089 | Brain/ FAIMS/IAA |
| Q62261 | Q62261 [1246-1256] | [R].IQEKVDSIDDR.[H] | 1xHomocysteineMMTS [K4] | 0,000530332 | 1480,67704 | Brain/ FAIMS/MMTS |
| Q62261; P15508 | Q62261 [119-127]; P15508 [119-127] | [K].ALQFLKEQR.[V] | 1xHomocysteineIAA [K6] | 0,00033298 | 1306,69362 | Brain/ FAIMS/IAA |
| Q62277 | Q62277 [164-179] | [K].GLSDVKMATDPENIIK.[E] | 1xHomocysteineIAA [K6] | 4,32935E-05 | 1904,94561 | Brain/ No FAIMS/IAA |
| Q62277 | Q62277 [164-179] | [K].GLSDVKMATDPENIIK.[E] | 1xHomocysteineIAA [K6] | 4,32935E-05 | 1904,94561 | Brain/ FAIMS/IAA |
| Q62277 | Q62277 [164-179] | [K].GLSDVKMATDPENIIK.[E] | 1xHomocysteineMMTS [K6] | 4,49774E-05 | 1893,91187 | Brain/ No FAIMS/MMTS |
| Q62277 | Q62277 [21-30] | [R].VVKEPLGFVK.[V] | 1xHomocysteineMMTS [K3] | 0,00818121 | 1278,69486 | Brain/ No FAIMS/MMTS |
| Q62277 | Q62277 [164-179] | [K].GLSDVKMATDPENIIK.[E] | 1xHomocysteineMMTS [K6] | 4,49774E-05 | 1893,91187 | Brain/ FAIMS/MMTS |
| Q62283 | Q62283 [114-122] | [R].HEIKDTFLR.[T] | 1xHomocysteineIAA [K4] | 0,000318237 | 1332,67288 | Brain/ No FAIMS/IAA |
| Q62283 | Q62283 [114-122] | [R].HEIKDTFLR.[T] | 1xHomocysteineIAA [K4] | 0,000318237 | 1332,67288 | Brain/ FAIMS/IAA |
| Q62318 | Q62318 [298-309] | [K].MAILQIMKELNK.[R] | 1xHomocysteineIAA [K8] | 0,00932126 | 1605,8525 | Brain/ FAIMS/IAA |
| Q62376 | Q62376 [110-120] | [R].VNYDTTESKLR.[R] | 1xHomocysteineIAA [K9] | 0,00012645 | 1499,71587 | Brain/ FAIMS/IAA |
| Q62418 | Q62418 [291-307] | [R].SPFLQKQLTQPETSYGR.[E] | 1xHomocysteineIAA [K6] | 4,32935E-05 | 2154,06481 | Brain/ FAIMS/IAA |
| Q62420 | Q62420 [137-152] | [K].QNFIDPLQNLHDKDLR.[E] | 1xHomocysteineIAA [K13] | 4,32935E-05 | 2140,0604 | Brain/ FAIMS/IAA |
| Q62420 | Q62420 [21-38] | [K].VGGAEGTKLDDDFKEMER.[K] | 1xHomocysteineMMTS [K8] | 4,49774E-05 | 2159,9406 | Brain/ FAIMS/MMTS |
| Q62420 | Q62420 [17-28] | [K].VSEKVGGAEGTK.[L] | 1xHomocysteineMMTS [K4] | 0,000152766 | 1324,62355 | Brain/ FAIMS/MMTS |
| Q62420 | Q62420 [137-152] | [K].QNFIDPLQNLHDKDLR.[E] | 1xHomocysteineMMTS [K13] | 4,49774E-05 | 2129,02665 | Brain/ FAIMS/MMTS |
| Q62420; Q62419 | Q62420 [21-34]; Q62419 [21-34] | [K].VGGAEGTKLDDDFK.[ED] | 1xHomocysteineIAA [K8] | 6,30912E-05 | 1625,74756 | Brain/ FAIMS/IAA |
| Q62420; Q62421; Q62419 | Q62420 [1-8]; Q62421 [1-8]; Q62419 [1-8] | [-].MSVAGLKK.[Q] | 1xMet-loss+Acetyl [N-Term]; 1xHomocysteineIAA [K7] | 0,00256222 | 918,50771 | Brain/ FAIMS/IAA |

| | | | | | | |
|---|---|---|---|---|---|---|
| Q62420; Q62421; Q62419 | Q62420 [1-8]; Q62421 [1-8]; Q62419 [1-8] | [-].MSVAGLKK.[Q] | 1xMet-loss+Acetyl [N-Term]; 1xHomocysteineMMTS [K7] | 0,000929608 | 907,47397 | Brain/ FAIMS/MMTS |
| Q62425 | Q62425 [49-64] | [K].NNPEPWNKLGPNEQYK.[F] | 1xHomocysteineIAA [K8] | 4,32935E-05 | 2101,976 | Liver/ FAIMS/IAA |
| Q62425 | Q62425 [49-64] | [K].NNPEPWNKLGPNEQYK.[F] | 1xHomocysteineIAA [K8] | 4,32935E-05 | 2101,976 | Brain/ FAIMS/IAA |
| Q62425 | Q62425 [49-64] | [K].NNPEPWNKLGPNEQYK.[F] | 1xHomocysteineMMTS [K8] | 4,49774E-05 | 2090,94226 | Liver/ FAIMS/MMTS |
| Q62425 | Q62425 [65-76] | [K].FYSVNVDYSKLK.[K] | 1xHomocysteineMMTS [K10] | 4,49774E-05 | 1625,77021 | Liver/ FAIMS/MMTS |
| Q62425 | Q62425 [48-64] | [R].KNNPEPWNKLGPNEQYK.[F] | 1xHomocysteineMMTS [K9] | 0,000192903 | 2219,03722 | Liver/ FAIMS/MMTS |
| Q62425 | Q62425 [57-74] | [K].LGPNEQYKFYSVNVDYSK.[L] | 1xHomocysteineMMTS [K8] | 4,49774E-05 | 2314,05187 | Liver/ FAIMS/MMTS |
| Q62425 | Q62425 [57-74] | [K].LGPNEQYKFYSVNVDYSK.[L] | 1xHomocysteineMMTS [K8] | 4,49774E-05 | 2314,05187 | Brain/ No FAIMS/MMTS |
| Q62425 | Q62425 [65-76] | [K].FYSVNVDYSKLK.[K] | 1xHomocysteineMMTS [K10] | 4,49774E-05 | 1625,77021 | Brain/ No FAIMS/MMTS |
| Q62425 | Q62425 [48-64] | [R].KNNPEPWNKLGPNEQYK.[F] | 1xHomocysteineMMTS [K9] | 0,000192903 | 2219,03722 | Brain/ FAIMS/MMTS |
| Q62425 | Q62425 [49-64] | [K].NNPEPWNKLGPNEQYK.[F] | 1xHomocysteineMMTS [K8] | 4,49774E-05 | 2090,94226 | Brain/ FAIMS/MMTS |
| Q62425 | Q62425 [57-74] | [K].LGPNEQYKFYSVNVDYSK.[L] | 1xHomocysteineMMTS [K8] | 4,49774E-05 | 2314,05187 | Brain/ FAIMS/MMTS |
| Q62425 | Q62425 [65-76] | [K].FYSVNVDYSKLK.[K] | 1xHomocysteineMMTS [K10] | 4,49774E-05 | 1625,77021 | Brain/ FAIMS/MMTS |
| Q62426 | Q62426 [35-44] | [K].FDVFKAISFK.[R] | 1xHomocysteineIAA [K5] | 4,32935E-05 | 1375,70787 | Liver/ FAIMS/IAA |
| Q62426 | Q62426 [35-44] | [K].FDVFKAISFK.[R] | 1xHomocysteineIAA [K5] | 4,32935E-05 | 1375,70787 | Brain/ FAIMS/IAA |
| Q62426 | Q62426 [35-44] | [K].FDVFKAISFK.[R] | 1xHomocysteineMMTS [K5] | 0,0060387 | 1364,67413 | Liver/ FAIMS/MMTS |
| Q62426 | Q62426 [69-89] | [R].VFQPLPHENKPLTLSSYQTNK.[E] | 1xHomocysteineMMTS [K10] | 0,000133296 | 2604,29489 | Liver/ FAIMS/MMTS |
| Q62442 | Q62442 [69-87] | [R].ADALQAGASQFESSAAKLK.[R] | 1xHomocysteineIAA [K17] | 4,32935E-05 | 2067,01753 | Brain/ FAIMS/IAA |
| Q62443 | Q62443 [70-80] | [K].ETILSQKETIR.[E] | 1xHomocysteineIAA [K7] | 0,0050458 | 1491,78355 | Brain/ FAIMS/IAA |
| Q62465 | Q62465 [380-390] | [K].VADAMKQMQEK.[K] | 1xHomocysteineIAA [K6] | 4,32935E-05 | 1452,66437 | Brain/ FAIMS/IAA |
| Q63810 | Q63810 [148-165] | [R].ISFEEFCAVVGGLDIHKK.[M] | 1xCarbamidomethyl [C7]; 1xHomocysteineIAA [K17] | 4,32935E-05 | 2223,09367 | Brain/ No FAIMS/IAA |
| Q63810 | Q63810 [148-165] | [R].ISFEEFCAVVGGLDIHKK.[M] | 1xCarbamidomethyl [C7]; 1xHomocysteineIAA [K17] | 4,32935E-05 | 2223,09367 | Brain/ FAIMS/IAA |
| Q63810 | Q63810 [136-147] | [K].TIINADKDGDGR.[I] | 1xHomocysteineMMTS [K7] | 4,49774E-05 | 1437,64607 | Brain/ FAIMS/MMTS |
| Q63810 | Q63810 [126-147] | [K].DTQLQQIVDKTIINADKDGDGR.[I] | 1xHomocysteineMMTS [K10] | 4,49774E-05 | 2606,25488 | Brain/ FAIMS/MMTS |
| Q63836 | Q63836 [399-419] | [R].LYATTSLYSDWDKQFYPDLIR.[E] | 1xHomocysteineIAA [K13] | 4,32935E-05 | 2769,32288 | Liver/ FAIMS/IAA |
| Q63836 | Q63836 [277-289] | [R].FYKNEEGTWSVEK.[V] | 1xHomocysteineMMTS [K3] | 4,49774E-05 | 1779,77167 | Liver/ No FAIMS/MMTS |
| Q63836 | Q63836 [399-419] | [R].LYATTSLYSDWDKQFYPDLIR.[E] | 1xHomocysteineMMTS [K13] | 4,49774E-05 | 2758,28914 | Liver/ No FAIMS/MMTS |
| Q63836 | Q63836 [277-289] | [R].FYKNEEGTWSVEK.[V] | 1xHomocysteineMMTS [K3] | 4,49774E-05 | 1779,77167 | Liver/ FAIMS/MMTS |
| Q63836 | Q63836 [175-195] | [K].GTWEKPGGASPMGYDFWYQPR.[H] | 1xHomocysteineMMTS [K5] | 4,49774E-05 | 2593,10973 | Liver/ FAIMS/MMTS |
| Q63836 | Q63836 [280-296] | [K].NEEGTWSVEKVIQVPSK.[F] | 1xHomocysteineMMTS [K10] | 4,49774E-05 | 2093,00419 | Liver/ FAIMS/MMTS |
| Q63836; P17563 | Q63836 [35-62]; P17563 [35-62] | [R].NTGTEAPDYLATVDVDPKSPQYSQVIHR.[L] | 1xHomocysteineIAA [K18] | 4,32935E-05 | 3275,56373 | Liver/ FAIMS/IAA |
| Q63836; P17563 | Q63836 [118-130]; P17563 [118-130] | [K].LHKVIEASEIQAK.[C] | 1xHomocysteineMMTS [K3] | 4,49774E-05 | 1628,84986 | Liver/ No FAIMS/MMTS |
| Q63836; P17563 | Q63836 [92-103]; P17563 [92-103] | [R].NKLILPGLISSR.[I] | 1xHomocysteineMMTS [K2] | 0,000173718 | 1473,828 | Liver/ No FAIMS/MMTS |
| Q63836; P17563 | Q63836 [35-62]; P17563 [35-62] | [R].NTGTEAPDYLATVDVDPKSPQYSQVIHR.[L] | 1xHomocysteineMMTS [K18] | 4,49774E-05 | 3264,52999 | Liver/ No FAIMS/MMTS |
| Q63836; P17563 | Q63836 [383-398]; P17563 [383-398] | [R].IPGGPQMIQLSLDGKR.[L] | 1xHomocysteineMMTS [K15] | 4,49774E-05 | 1872,94926 | Liver/ No FAIMS/MMTS |
| Q63836; P17563 | Q63836 [438-460]; P17563 [438-460] | [K].LNPNFLVDFGKEPLGPALAHELR.[Y] | 1xHomocysteineMMTS [K11] | 4,49774E-05 | 2710,38437 | Liver/ No FAIMS/MMTS |
| Q63836; P17563 | Q63836 [334-344]; P17563 [334-344] | [R].QYDISNPQKPR.[L] | 1xHomocysteineMMTS [K9] | 0,000266738 | 1508,69844 | Liver/ FAIMS/MMTS |
| Q63836; P17563 | Q63836 [92-103]; P17563 [92-103] | [R].NKLILPGLISSR.[I] | 1xHomocysteineMMTS [K2] | 0,000173718 | 1473,828 | Liver/ FAIMS/MMTS |
| Q63836; P17563 | Q63836 [35-62]; P17563 [35-62] | [R].NTGTEAPDYLATVDVDPKSPQYSQVIHR.[L] | 1xHomocysteineMMTS [K18] | 4,49774E-05 | 3264,52999 | Liver/ FAIMS/MMTS |
| Q63836; P17563 | Q63836 [118-130]; P17563 [118-130] | [K].LHKVIEASEIQAK.[C] | 1xHomocysteineMMTS [K3] | 4,49774E-05 | 1628,84986 | Liver/ FAIMS/MMTS |
| Q63836; P17563 | Q63836 [383-398]; P17563 [383-398] | [R].IPGGPQMIQLSLDGKR.[L] | 1xHomocysteineMMTS [K15] | 4,49774E-05 | 1872,94926 | Liver/ FAIMS/MMTS |
| Q63836; P17563 | Q63836 [438-460]; P17563 [438-460] | [K].LNPNFLVDFGKEPLGPALAHELR.[Y] | 1xHomocysteineMMTS [K11] | 4,49774E-05 | 2710,38437 | Liver/ FAIMS/MMTS |
| Q63880 | Q63880 [112-119] | [R].FTLNEKMK.[I] | 1xHomocysteineIAA [K6] | 0,00258601 | 1184,58023 | Liver/ FAIMS/IAA |
| Q63880 | Q63880 [112-119] | [R].FTLNEKMK.[I] | 1xHomocysteineMMTS [K6] | 0,00459586 | 1173,54648 | Liver/ FAIMS/MMTS |
| Q63880; Q8VCU1 | Q63880 [50-58]; Q8VCU1 [50-58] | [R].QVGVKDTDR.[M] | 1xHomocysteineMMTS [K5] | 0,00461842 | 1180,5449 | Liver/ No FAIMS/MMTS |
| Q63880; Q8VCU1 | Q63880 [451-457]; Q8VCU1 [451-457] | [K].FKPAWVK.[A] | 1xHomocysteineMMTS [K2] | 0,00570776 | 1038,52634 | Liver/ No FAIMS/MMTS |
| Q63880; Q8VCU1 | Q63880 [554-560]; Q8VCU1 [554-560] | [R].KIQEWHR.[E] | 1xHomocysteineMMTS [K1] | 0,000599431 | 1159,54993 | Liver/ No FAIMS/MMTS |
| Q63880; Q8VCU1 | Q63880 [50-58]; Q8VCU1 [50-58] | [R].QVGVKDTDR.[M] | 1xHomocysteineMMTS [K5] | 0,00461842 | 1180,5449 | Liver/ FAIMS/MMTS |
| Q63880; Q8VCU1 | Q63880 [554-560]; Q8VCU1 [554-560] | [R].KIQEWHR.[E] | 1xHomocysteineMMTS [K1] | 0,000599431 | 1159,54993 | Liver/ FAIMS/MMTS |
| Q63912 | Q63912 [117-132] | [K].SDTAYQWNLKYLDVSK.[N] | 1xHomocysteineIAA [K10] | 4,32935E-05 | 2105,00082 | Brain/ No FAIMS/IAA |
| Q63912 | Q63912 [117-132] | [K].SDTAYQWNLKYLDVSK.[N] | 1xHomocysteineIAA [K10] | 4,32935E-05 | 2105,00082 | Brain/ FAIMS/IAA |
| Q640R3 | Q640R3 [50-68] | [K].SALLSVQYSSTSSDKPVVK.[W] | 1xHomocysteineIAA [K15] | 4,32935E-05 | 2170,10601 | Brain/ FAIMS/IAA |
| Q64105 | Q64105 [50-63] | [R].QLKEELGAQQPDLK.[V] | 1xHomocysteineIAA [K3] | 0,00410529 | 1770,90546 | Brain/ FAIMS/IAA |
| Q64105 | Q64105 [50-63] | [R].QLKEELGAQQPDLK.[V] | 1xHomocysteineMMTS [K3] | 0,00109439 | 1759,87172 | Liver/ FAIMS/MMTS |
| Q64288 | Q64288 [109-128] | [R].KEDSDAMDWNEADALEFGER.[L] | 1xHomocysteineIAA [K1] | 4,32935E-05 | 2502,01839 | Brain/ FAIMS/IAA |
| Q64288 | Q64288 [63-77] | [K].LQFDHWNVVLDKPGK.[V] | 1xHomocysteineMMTS [K12] | 0,000829616 | 1958,96153 | Brain/ FAIMS/MMTS |
| Q64324 | Q64324 [1-13] | [-].MAPLGLKAVVGEK.[I] | 1xMet-loss [N-Term]; 1xHomocysteineMMTS [K7] | 0,00624351 | 1344,73779 | Brain/ No FAIMS/MMTS |
| Q64332 | Q64332 [116-132] | [K].VLLVVDEPHTDWAKCFR.[G] | 1xCarbamidomethyl [C15]; 1xHomocysteineIAA [K14] | 4,32935E-05 | 2259,1049 | Brain/ FAIMS/IAA |
| Q64374 | Q64374 [284-299] | [K].ITGLGVKGIAPYSYAG.[-] | 1xHomocysteineIAA [K7] | 4,32935E-05 | 1740,89892 | Liver/ No FAIMS/IAA |

| | | | | | | |
|---|---|---|---|---|---|---|
| Q64374 | Q64374 [284-299] | [K].ITGLGVKGIAPYSYAG.[-] | 1xHomocysteineIAA [K7] | 4,32935E-05 | 1740,89892 | Liver/ FAIMS/IAA |
| Q64374 | Q64374 [275-290] | [R].QPDAGNIFKITGLGVK.[G] | 1xHomocysteineMMTS [K9] | 0,000361246 | 1820,93974 | Liver/ No FAIMS/MMTS |
| Q64374 | Q64374 [284-299] | [K].ITGLGVKGIAPYSYAG.[-] | 1xHomocysteineMMTS [K7] | 4,49774E-05 | 1729,86517 | Liver/ No FAIMS/MMTS |
| Q64374 | Q64374 [102-112] | [R].FNDGKVDPAGR.[Y] | 1xHomocysteineMMTS [K5] | 4,49774E-05 | 1338,59292 | Liver/ No FAIMS/MMTS |
| Q64374 | Q64374 [235-244] | [R].LQTVKLPVDK.[T] | 1xHomocysteineMMTS [K5] | 4,49774E-05 | 1303,71124 | Liver/ No FAIMS/MMTS |
| Q64374 | Q64374 [130-145] | [R].HQGSLYSLFPDHSVKK.[Y] | 1xHomocysteineMMTS [K15] | 4,49774E-05 | 2005,96226 | Liver/ No FAIMS/MMTS |
| Q64374 | Q64374 [196-213] | [K].MEKDEQIPDGMCIDAEGK.[L] | 1xMethylthio [C12];<br>1xHomocysteineMMTS [K3] | 4,49774E-05 | 2217,86631 | Liver/ FAIMS/MMTS |
| Q64374 | Q64374 [275-290] | [R].QPDAGNIFKITGLGVK.[G] | 1xHomocysteineMMTS [K9] | 0,000361246 | 1820,93974 | Liver/ FAIMS/MMTS |
| Q64374 | Q64374 [284-299] | [K].ITGLGVKGIAPYSYAG.[-] | 1xHomocysteineMMTS [K7] | 4,49774E-05 | 1729,86517 | Liver/ FAIMS/MMTS |
| Q64374 | Q64374 [102-112] | [R].FNDGKVDPAGR.[Y] | 1xHomocysteineMMTS [K5] | 4,49774E-05 | 1338,59292 | Liver/ FAIMS/MMTS |
| Q64374 | Q64374 [235-244] | [R].LQTVKLPVDK.[T] | 1xHomocysteineMMTS [K5] | 4,49774E-05 | 1303,71124 | Liver/ FAIMS/MMTS |
| Q64374 | Q64374 [130-145] | [R].HQGSLYSLFPDHSVKK.[Y] | 1xHomocysteineMMTS [K15] | 4,49774E-05 | 2005,96226 | Liver/ FAIMS/MMTS |
| Q64374 | Q64374 [145-163] | [K].KYFDQVDISNGLDWSLDHK.[I] | 1xHomocysteineMMTS [K1] | 4,49774E-05 | 2443,10569 | Liver/ FAIMS/MMTS |
| Q64433 | Q64433 [21-36] | [R].SAAETVTKGGIMLPEK.[S] | 1xHomocysteineIAA [K8] | 4,32935E-05 | 1805,91358 | Liver/ FAIMS/IAA |
| Q64433 | Q64433 [21-36] | [R].SAAETVTKGGIMLPEK.[S] | 1xHomocysteineIAA [K8] | 4,32935E-05 | 1805,91358 | Brain/ No FAIMS/IAA |
| Q64433 | Q64433 [21-36] | [R].SAAETVTKGGIMLPEK.[S] | 1xHomocysteineIAA [K8] | 4,32935E-05 | 1805,91358 | Brain/ FAIMS/IAA |
| Q64433 | Q64433 [29-40] | [K].GGIMLPEKSQGK.[V] | 1xHomocysteineMMTS [K8] | 0,000266738 | 1407,67929 | Liver/ No FAIMS/MMTS |
| Q64433 | Q64433 [21-36] | [R].SAAETVTKGGIMLPEK.[S] | 1xHomocysteineMMTS [K8] | 4,49774E-05 | 1794,87984 | Liver/ No FAIMS/MMTS |
| Q64433 | Q64433 [93-102] | [R].DSDILGKYVD.[-] | 1xHomocysteineMMTS [K7] | 0,0017273 | 1287,55955 | Liver/ No FAIMS/MMTS |
| Q64433 | Q64433 [81-92] | [K].VVLDDKDYFLFR.[D] | 1xHomocysteineMMTS [K6] | 4,49774E-05 | 1692,81241 | Liver/ No FAIMS/MMTS |
| Q64433 | Q64433 [37-54] | [K].SQGKVLQATVVAVGSGGK.[G] | 1xHomocysteineMMTS [K4] | 4,49774E-05 | 1848,96701 | Liver/ No FAIMS/MMTS |
| Q64433 | Q64433 [67-80] | [K].VGDKVLLPEYGGTK.[V] | 1xHomocysteineMMTS [K4] | 4,49774E-05 | 1638,82298 | Liver/ No FAIMS/MMTS |
| Q64433 | Q64433 [55-66] | [K].GKSGEIEPVSVK.[V] | 1xHomocysteineMMTS [K2] | 4,49774E-05 | 1392,68615 | Liver/ No FAIMS/MMTS |
| Q64433 | Q64433 [41-56] | [K].VLQATVVAVGSGGKGK.[S] | 1xHomocysteineMMTS [K14] | 4,49774E-05 | 1633,87641 | Liver/ No FAIMS/MMTS |
| Q64433 | Q64433 [57-70] | [K].SGEIEPVSVKVGDK.[V] | 1xHomocysteineMMTS [K10] | 4,49774E-05 | 1606,7815 | Liver/ No FAIMS/MMTS |
| Q64433 | Q64433 [71-92] | [K].VLLPEYGGTKVVLDDKDYFLFR.[D] | 1xHomocysteineMMTS [K10] | 4,49774E-05 | 2750,39321 | Liver/ No FAIMS/MMTS |
| Q64433 | Q64433 [8-15] | [R].KFLPLFDR.[V] | 1xHomocysteineMMTS [K1] | 0,00570776 | 1198,61113 | Liver/ No FAIMS/MMTS |
| Q64433 | Q64433 [21-36] | [R].SAAETVTKGGIMLPEK.[S] | 1xHomocysteineMMTS [K8] | 4,49774E-05 | 1794,87984 | Liver/ FAIMS/MMTS |
| Q64433 | Q64433 [81-92] | [K].VVLDDKDYFLFR.[D] | 1xHomocysteineMMTS [K6] | 4,49774E-05 | 1692,81241 | Liver/ FAIMS/MMTS |
| Q64433 | Q64433 [37-54] | [K].SQGKVLQATVVAVGSGGK.[G] | 1xHomocysteineMMTS [K4] | 4,49774E-05 | 1848,96701 | Liver/ FAIMS/MMTS |
| Q64433 | Q64433 [67-80] | [K].VGDKVLLPEYGGTK.[V] | 1xHomocysteineMMTS [K4] | 4,49774E-05 | 1638,82298 | Liver/ FAIMS/MMTS |
| Q64433 | Q64433 [55-66] | [K].GKSGEIEPVSVK.[V] | 1xHomocysteineMMTS [K2] | 4,49774E-05 | 1392,68615 | Liver/ FAIMS/MMTS |
| Q64433 | Q64433 [57-70] | [K].SGEIEPVSVKVGDK.[V] | 1xHomocysteineMMTS [K10] | 4,49774E-05 | 1606,7815 | Liver/ FAIMS/MMTS |
| Q64433 | Q64433 [71-92] | [K].VLLPEYGGTKVVLDDKDYFLFR.[D] | 1xHomocysteineMMTS [K10] | 4,49774E-05 | 2750,39321 | Liver/ FAIMS/MMTS |
| Q64433 | Q64433 [8-15] | [R].KFLPLFDR.[V] | 1xHomocysteineMMTS [K1] | 0,00570776 | 1198,61113 | Liver/ FAIMS/MMTS |
| Q64433 | Q64433 [21-36] | [R].SAAETVTKGGIMLPEK.[S] | 1xOxidation [M12];<br>1xHomocysteineMMTS [K8] | 0,000133296 | 1810,87475 | Liver/ FAIMS/MMTS |
| Q64433 | Q64433 [29-40] | [K].GGIMLPEKSQGK.[V] | 1xHomocysteineMMTS [K8] | 0,000266738 | 1407,67929 | Liver/ FAIMS/MMTS |
| Q64433 | Q64433 [93-102] | [R].DSDILGKYVD.[-] | 1xHomocysteineMMTS [K7] | 0,0017273 | 1287,55955 | Liver/ FAIMS/MMTS |
| Q64433 | Q64433 [41-56] | [K].VLQATVVAVGSGGKGK.[S] | 1xHomocysteineMMTS [K14] | 4,49774E-05 | 1633,87641 | Liver/ FAIMS/MMTS |
| Q64433 | Q64433 [29-40] | [K].GGIMLPEKSQGK.[V] | 1xHomocysteineMMTS [K8] | 0,000266738 | 1407,67929 | Brain/ No FAIMS/MMTS |
| Q64433 | Q64433 [67-80] | [K].VGDKVLLPEYGGTK.[V] | 1xHomocysteineMMTS [K4] | 4,49774E-05 | 1638,82298 | Brain/ No FAIMS/MMTS |
| Q64433 | Q64433 [81-92] | [K].VVLDDKDYFLFR.[D] | 1xHomocysteineMMTS [K6] | 4,49774E-05 | 1692,81241 | Brain/ FAIMS/MMTS |
| Q64433 | Q64433 [55-66] | [K].GKSGEIEPVSVK.[V] | 1xHomocysteineMMTS [K2] | 4,49774E-05 | 1392,68615 | Brain/ FAIMS/MMTS |
| Q64433 | Q64433 [71-92] | [K].VLLPEYGGTKVVLDDKDYFLFR.[D] | 1xHomocysteineMMTS [K10] | 4,49774E-05 | 2750,39321 | Brain/ FAIMS/MMTS |
| Q64442 | Q64442 [64-78] | [K].KPMVLGHEAAGTVTK.[V] | 1xHomocysteineIAA [K1] | 4,32935E-05 | 1712,88222 | Liver/ FAIMS/IAA |
| Q64442 | Q64442 [1-21] | [-].MAAPAKGENLSLVVHGPGDIR.[L] | 1xMet-loss+Acetyl [N-Term];<br>1xHomocysteineMMTS [K6] | 4,49774E-05 | 2206,11072 | Liver/ No FAIMS/MMTS |
| Q64442 | Q64442 [331-340] | [K].AVEAFETAKK.[G] | 1xHomocysteineMMTS [K9] | 0,000507453 | 1256,60136 | Liver/ No FAIMS/MMTS |
| Q64442 | Q64442 [79-91] | [K].VGELVKHLKPGDR.[V] | 1xHomocysteineMMTS [K6] | 0,000173718 | 1610,85053 | Liver/ No FAIMS/MMTS |
| Q64442 | Q64442 [315-325] | [K].TLNVKPLVTHR.[F] | 1xHomocysteineMMTS [K5] | 0,000152766 | 1440,78139 | Liver/ No FAIMS/MMTS |
| Q64442 | Q64442 [291-299] | [R].EVDIKGVFR.[Y] | 1xHomocysteineMMTS [K5] | 0,00356631 | 1225,60678 | Liver/ No FAIMS/MMTS |
| Q64442 | Q64442 [326-339] | [R].FPLEKAVEAFETAK.[K] | 1xHomocysteineMMTS [K5] | 6,70238E-05 | 1742,84919 | Liver/ No FAIMS/MMTS |
| Q64442 | Q64442 [64-78] | [K].KPMVLGHEAAGTVTK.[V] | 1xHomocysteineMMTS [K1] | 4,49774E-05 | 1701,84848 | Liver/ No FAIMS/MMTS |
| Q64442 | Q64442 [64-78] | [K].KPMVLGHEAAGTVTK.[V] | 1xOxidation [M3];<br>1xHomocysteineMMTS [K1] | 4,49774E-05 | 1717,84339 | Liver/ FAIMS/MMTS |
| Q64442 | Q64442 [1-21] | [-].MAAPAKGENLSLVVHGPGDIR.[L] | 1xMet-loss+Acetyl [N-Term];<br>1xHomocysteineMMTS [K6] | 4,49774E-05 | 2206,11072 | Liver/ FAIMS/MMTS |
| Q64442 | Q64442 [315-325] | [K].TLNVKPLVTHR.[F] | 1xHomocysteineMMTS [K5] | 0,000152766 | 1440,78139 | Liver/ FAIMS/MMTS |
| Q64442 | Q64442 [64-78] | [K].KPMVLGHEAAGTVTK.[V] | 1xHomocysteineMMTS [K1] | 4,49774E-05 | 1701,84848 | Liver/ FAIMS/MMTS |
| Q64442 | Q64442 [331-340] | [K].AVEAFETAKK.[G] | 1xHomocysteineMMTS [K9] | 0,000507453 | 1256,60136 | Liver/ FAIMS/MMTS |
| Q64442 | Q64442 [57-78] | [R].IGDFVVKKPMVLGHEAAGTVTK.[V] | 1xHomocysteineMMTS [K7] | 4,49774E-05 | 2460,28115 | Liver/ FAIMS/MMTS |
| Q64442 | Q64442 [291-299] | [R].EVDIKGVFR.[Y] | 1xHomocysteineMMTS [K5] | 0,00356631 | 1225,60678 | Liver/ FAIMS/MMTS |
| Q64442 | Q64442 [326-339] | [R].FPLEKAVEAFETAK.[K] | 1xHomocysteineMMTS [K5] | 6,70238E-05 | 1742,84919 | Liver/ FAIMS/MMTS |
| Q64442 | Q64442 [215-235] | [K].EVGADFTIQVGKETPQEIASK.[V] | 1xHomocysteineMMTS [K12] | 4,49774E-05 | 2410,16287 | Liver/ FAIMS/MMTS |
| Q64471 | Q64471 [198-208] | [R].VEAAVGKDLFR.[E] | 1xHomocysteineMMTS [K7] | 4,49774E-05 | 1367,681 | Liver/ FAIMS/MMTS |
| Q64478 | Q64478 [1-12] | [-].MPEPAKSAPAPK.[K] | 1xMet-loss [N-Term];<br>1xHomocysteineIAA [K6] | 4,32935E-05 | 1266,65108 | Liver/ No FAIMS/IAA |
| Q64478 | Q64478 [1-12] | [-].MPEPAKSAPAPK.[K] | 1xMet-loss [N-Term];<br>1xHomocysteineIAA [K6] | 4,32935E-05 | 1266,65108 | Brain/ No FAIMS/IAA |
| Q64520 | Q64520 [138-148] | [R].NTETEESLAKR.[L] | 1xHomocysteineIAA [K10] | 0,000861826 | 1451,67948 | Brain/ FAIMS/IAA |
| Q64521 | Q64521 [442-463] | [R].SMAEDTVDAAVKFHNLNAGPSR.[T] | 1xHomocysteineIAA [K12] | 0,000228034 | 2504,16567 | Brain/ FAIMS/IAA |
| Q64523 | Q64523 [90-100] | [R].NDEELNKLLGK.[V] | 1xHomocysteineIAA [K7] | 9,61721E-05 | 1446,7257 | Brain/ No FAIMS/IAA |
| Q64523 | Q64523 [90-100] | [R].NDEELNKLLGK.[V] | 1xHomocysteineIAA [K7] | 9,61721E-05 | 1446,7257 | Brain/ FAIMS/IAA |
| Q64523 | Q64523 [90-100] | [R].NDEELNKLLGK.[V] | 1xHomocysteineMMTS [K7] | 0,000250158 | 1435,69196 | Liver/ No FAIMS/MMTS |
| Q64523 | Q64523 [90-100] | [R].NDEELNKLLGK.[V] | 1xHomocysteineMMTS [K7] | 0,000250158 | 1435,69196 | Liver/ FAIMS/MMTS |
| Q64523 | Q64523 [90-100] | [R].NDEELNKLLGK.[V] | 1xHomocysteineMMTS [K7] | 0,000250158 | 1435,69196 | Brain/ FAIMS/MMTS |
| Q64523;<br>Q8BFU2 | Q64523 [101-120];<br>Q8BFU2 [101-120] | [KR].VTIAQGGVLPNIQAVLLPKK.[T] | 1xHomocysteineIAA [K19] | 4,32935E-05 | 2233,31007 | Brain/ FAIMS/IAA |
| Q64525 | Q64525 [1-12] | [-].MPDPAKSAPAPK.[K] | 1xMet-loss [N-Term];<br>1xHomocysteineIAA [K6] | 4,32935E-05 | 1252,63543 | Brain/ FAIMS/IAA |

| | | | | | | |
|---|---|---|---|---|---|---|
| Q64525; | Q64525 [36-47]; | | | | | |
| Q64478 | Q64478 [36-47] | [K].ESYSVYVYKVLK.[Q] | 1xHomocysteineIAA [K9] | 9,61721E-05 | 1651,84001 | Liver/ FAIMS/IAA |
| Q64525; | Q64525 [35-44]; | | | | | |
| Q64478 | Q64478 [35-44] | [R].KESYSVYVYK.[V] | 1xHomocysteineIAA [K1] | 4,32935E-05 | 1439,68753 | Brain/ No FAIMS/IAA |
| Q64525; | Q64525 [35-44]; | | | | | |
| Q64478 | Q64478 [35-44] | [R].KESYSVYVYK.[V] | 1xHomocysteineIAA [K1] | 4,32935E-05 | 1439,68753 | Brain/ FAIMS/IAA |
| Q64525; | Q64525 [101-117]; | | | | | |
| Q64478; | Q64478 [101-117]; | | | | | |
| Q9D2U9 | Q9D2U9 [101-117] | [R].LLLPGELAKHAVSEGTK.[A] | 1xHomocysteineIAA [K9] | 4,32935E-05 | 1937,05246 | Liver/ FAIMS/IAA |
| Q64525; | Q64525 [48-73]; | | | | | |
| Q64478; | Q64478 [48-73]; | | 1xOxidation [M13]; | | | |
| Q9D2U9 | Q9D2U9 [48-73] | [K].QVHPDTGISSKAMGIMNSFVNDIFER.[I] | 1xHomocysteineIAA [K11] | 4,32935E-05 | 3083,43834 | Brain/ FAIMS/IAA |
| Q64525; | Q64525 [36-47]; | | | | | |
| Q8CGP1 | Q8CGP1 [36-47] | [K].ESYSVYVYKVLK.[Q] | 1xHomocysteineMMTS [K9] | 0,00109439 | 1640,80626 | Liver/ FAIMS/MMTS |
| Q64525; | Q64525 [35-44]; | | | | | |
| Q8CGP1 | Q8CGP1 [35-44] | [R].KESYSVYVYK.[V] | 1xHomocysteineMMTS [K1] | 4,49774E-05 | 1428,65379 | Liver/ FAIMS/MMTS |
| Q64525; | Q64525 [101-117]; | | | | | |
| Q8CGP1; | Q8CGP1 [101-117]; | | | | | |
| Q9D2U9 | Q9D2U9 [101-117] | [R].LLLPGELAKHAVSEGTK.[A] | 1xHomocysteineMMTS [K9] | 4,49774E-05 | 1926,01872 | Liver/ No FAIMS/MMTS |
| Q64525; | Q64525 [110-121]; | | | | | |
| Q8CGP1; | Q8CGP1 [110-121]; | | | | | |
| Q9D2U9 | Q9D2U9 [110-121] | [K].HAVSEGTKAVTK.[Y] | 1xHomocysteineMMTS [K8] | 4,49774E-05 | 1390,68173 | Liver/ No FAIMS/MMTS |
| Q64525; | Q64525 [45-58]; | | | | | |
| Q8CGP1; | Q8CGP1 [45-58]; | | | | | |
| Q9D2U9 | Q9D2U9 [45-58] | [K].VLKQVHPDTGISSK.[A] | 1xHomocysteineMMTS [K3] | 4,49774E-05 | 1671,85567 | Liver/ No FAIMS/MMTS |
| Q64525; | Q64525 [101-117]; | | | | | |
| Q8CGP1; | Q8CGP1 [101-117]; | | | | | |
| Q9D2U9 | Q9D2U9 [101-117] | [R].LLLPGELAKHAVSEGTK.[A] | 1xHomocysteineMMTS [K9] | 4,49774E-05 | 1926,01872 | Liver/ FAIMS/MMTS |
| Q64525; | Q64525 [110-121]; | | | | | |
| Q8CGP1; | Q8CGP1 [110-121]; | | | | | |
| Q9D2U9 | Q9D2U9 [110-121] | [K].HAVSEGTKAVTK.[Y] | 1xHomocysteineMMTS [K8] | 4,49774E-05 | 1390,68173 | Liver/ FAIMS/MMTS |
| Q64525; | Q64525 [45-58]; | | | | | |
| Q8CGP1; | Q8CGP1 [45-58]; | | | | | |
| Q9D2U9 | Q9D2U9 [45-58] | [K].VLKQVHPDTGISSK.[A] | 1xHomocysteineMMTS [K3] | 4,49774E-05 | 1671,85567 | Liver/ FAIMS/MMTS |
| Q64525; | Q64525 [101-117]; | | | | | |
| Q8CGP1; | Q8CGP1 [101-117]; | | | | | |
| Q9D2U9 | Q9D2U9 [101-117] | [R].LLLPGELAKHAVSEGTK.[A] | 1xHomocysteineMMTS [K9] | 4,49774E-05 | 1926,01872 | Brain/ No FAIMS/MMTS |
| Q64525; | Q64525 [110-121]; | | | | | |
| Q8CGP1; | Q8CGP1 [110-121]; | | | | | |
| Q9D2U9 | Q9D2U9 [110-121] | [K].HAVSEGTKAVTK.[Y] | 1xHomocysteineMMTS [K8] | 4,49774E-05 | 1390,68173 | Brain/ No FAIMS/MMTS |
| Q64525; | Q64525 [101-117]; | | | | | |
| Q8CGP1; | Q8CGP1 [101-117]; | | | | | |
| Q9D2U9 | Q9D2U9 [101-117] | [R].LLLPGELAKHAVSEGTK.[A] | 1xHomocysteineMMTS [K9] | 4,49774E-05 | 1926,01872 | Brain/ FAIMS/MMTS |
| Q64525; | Q64525 [110-121]; | | | | | |
| Q8CGP1; | Q8CGP1 [110-121]; | | | | | |
| Q9D2U9 | Q9D2U9 [110-121] | [K].HAVSEGTKAVTK.[Y] | 1xHomocysteineMMTS [K8] | 4,49774E-05 | 1390,68173 | Brain/ FAIMS/MMTS |
| Q64525; | Q64525 [45-58]; | | | | | |
| Q8CGP1; | Q8CGP1 [45-58]; | | | | | |
| Q9D2U9 | Q9D2U9 [45-58] | [K].VLKQVHPDTGISSK.[A] | 1xHomocysteineMMTS [K3] | 4,49774E-05 | 1671,85567 | Brain/ FAIMS/MMTS |
| Q68ED7 | Q68ED7 [43-57] | [R].LQLQKSQYLQLGPSR.[G] | 1xHomocysteineIAA [K5] | 0,00323447 | 1933,03239 | Brain/ FAIMS/IAA |
| | | | 1xCarbamidomethyl [C4]; | | | |
| Q68EF4 | Q68EF4 [14-35] | [R].LPLCLLLSLYGSWVPSSLGKPK.[G] | 1xHomocysteineIAA [K20] | 0,00518879 | 2602,41355 | Brain/ No FAIMS/IAA |
| Q68FD5 | Q68FD5 [367-382] | [R].KFNALFAQGNYSEAAK.[V] | 1xHomocysteineIAA [K1] | 4,32935E-05 | 1932,92726 | Brain/ FAIMS/IAA |
| Q68FD5 | Q68FD5 [84-96] | [K].AGKTLQIFNIEMK.[S] | 1xHomocysteineMMTS [K3] | 0,0004709 | 1655,83177 | Brain/ FAIMS/MMTS |
| Q6GQX6 | Q6GQX6 [540-546] | [R].LPSDKLK.[A] | 1xHomocysteineIAA [K5] | 0,00158131 | 974,53393 | Brain/ No FAIMS/IAA |
| Q6IRU2 | Q6IRU2 [13-27] | [R].KIQALQQQADDAEDR.[A] | 1xHomocysteineIAA [K1] | 4,32935E-05 | 1902,89741 | Liver/ No FAIMS/IAA |
| Q6IRU2 | Q6IRU2 [13-27] | [R].KIQALQQQADDAEDR.[A] | 1xHomocysteineIAA [K1] | 4,32935E-05 | 1902,89741 | Liver/ FAIMS/IAA |
| Q6IRU2 | Q6IRU2 [13-27] | [R].KIQALQQQADDAEDR.[A] | 1xHomocysteineIAA [K1] | 4,32935E-05 | 1902,89741 | Brain/ FAIMS/IAA |
| Q6IRU2 | Q6IRU2 [13-27] | [R].KIQALQQQADDAEDR.[A] | 1xHomocysteineMMTS [K1] | 4,49774E-05 | 1891,86367 | Liver/ FAIMS/MMTS |
| Q6IRU2 | Q6IRU2 [43-54] | [R].EKAEGDAAALNR.[R] | 1xHomocysteineMMTS [K2] | 0,00363897 | 1407,63551 | Liver/ FAIMS/MMTS |
| Q6IRU2 | Q6IRU2 [216-228] | [K].TIDDLEEKLAQAK.[E] | 1xHomocysteineMMTS [K8] | 0,0061262 | 1636,79207 | Brain/ FAIMS/MMTS |
| Q6IRU2 | Q6IRU2 [13-27] | [R].KIQALQQQADDAEDR.[A] | 1xHomocysteineMMTS [K1] | 4,49774E-05 | 1891,86367 | Brain/ FAIMS/MMTS |
| Q6IRU5 | Q6IRU5 [114-129] | [R].LQELDAASKVTEQEWR.[E] | 1xHomocysteineIAA [K9] | 4,32935E-05 | 2077,00188 | Brain/ No FAIMS/IAA |
| Q6IRU5 | Q6IRU5 [114-129] | [R].LQELDAASKVTEQEWR.[E] | 1xHomocysteineIAA [K9] | 4,32935E-05 | 2077,00188 | Brain/ FAIMS/IAA |
| Q6IRU5 | Q6IRU5 [114-129] | [R].LQELDAASKVTEQEWR.[E] | 1xHomocysteineMMTS [K9] | 4,49774E-05 | 2065,96814 | Brain/ FAIMS/MMTS |
| Q6IRU5 | Q6IRU5 [182-194] | [K].ESKEETPGTEWEK.[V] | 1xHomocysteineMMTS [K3] | 0,00115861 | 1712,71421 | Brain/ FAIMS/MMTS |
| Q6P069 | Q6P069 [117-135] | [R].SGTVDPQELQKALTTMGFR.[L] | 1xHomocysteineIAA [K11] | 4,32935E-05 | 2253,10021 | Brain/ No FAIMS/IAA |
| Q6P069 | Q6P069 [117-135] | [R].SGTVDPQELQKALTTMGFR.[L] | 1xHomocysteineIAA [K11] | 4,32935E-05 | 2253,10021 | Brain/ FAIMS/IAA |
| Q6P8X1 | Q6P8X1 [54-68] | [K].SSLPNFKQNEFSVVR.[Q] | 1xHomocysteineIAA [K7] | 0,000243434 | 1925,95381 | Brain/ FAIMS/IAA |
| Q6PDL0 | Q6PDL0 [51-67] | [K].LPSGKNILVFGEDGSGK.[T] | 1xHomocysteineIAA [K5] | 0,000177335 | 1891,95822 | Brain/ FAIMS/IAA |
| | | | 1xCarbamidomethyl [C15]; | | | |
| Q6PDY2 | Q6PDY2 [211-229] | [R].VVEPIRPKEASGSACDLPR.[E] | 1xHomocysteineIAA [K8] | 0,00496744 | 2255,1271 | Brain/ FAIMS/IAA |
| Q6PER3 | Q6PER3 [114-130] | [K].FFDANYDGKDYNPLLAR.[Q] | 1xHomocysteineIAA [K9] | 4,32935E-05 | 2193,00696 | Brain/ FAIMS/IAA |
| Q6PHN9 | Q6PHN9 [11-27] | [K].LLIIGDSGVGKSSLLLR.[F] | 1xHomocysteineIAA [K11] | 4,32935E-05 | 1915,10449 | Brain/ FAIMS/IAA |
| Q6PHS9 | Q6PHS9 [192-203] | [R].LDFIEDPNFKNK.[V] | 1xHomocysteineIAA [K10] | 0,00049316 | 1653,79412 | Brain/ FAIMS/IAA |
| Q6PHZ8; | Q6PHZ8 [93-109]; | | 1xCarbamidomethyl [C6]; | | | |
| Q9JJ57 | Q9JJ57 [70-86] | [R].GFKNECPSGVVNEETFK.[EQ] | 1xHomocysteineIAA [K3] | 0,00847886 | 2115,9474 | Brain/ FAIMS/IAA |
| Q6PIC6 | Q6PIC6 [466-477] | [K].KVAEIPFNSTNK.[Y] | 1xHomocysteineIAA [K1] | 4,32935E-05 | 1521,77299 | Brain/ No FAIMS/IAA |
| Q6PIC6 | Q6PIC6 [466-477] | [K].KVAEIPFNSTNK.[Y] | 1xHomocysteineIAA [K1] | 4,32935E-05 | 1521,77299 | Brain/ FAIMS/IAA |
| Q6PIC6 | Q6PIC6 [493-503] | [R].YLLVMKGAPER.[I] | 1xHomocysteineMMTS [K6] | 4,49774E-05 | 1439,72076 | Brain/ FAIMS/MMTS |
| Q6PIC6 | Q6PIC6 [425-435] | [K].GGQDNIPVLKR.[D] | 1xHomocysteineMMTS [K10] | 0,00164853 | 1359,68715 | Brain/ FAIMS/MMTS |

| | | | | | | |
|---|---|---|---|---|---|---|
| Q6PIE5 | Q6PIE5 [203-225] | [R].IISSHGCKVDNSSLTGESEPQTR.[S] | 1xCarbamidomethyl [C7]; 1xHomocysteineIAA [K8] | 4,32935E-05 | 2676,2352 | Brain/ No FAIMS/IAA |
| Q6PIE5 | Q6PIE5 [203-225] | [R].IISSHGCKVDNSSLTGESEPQTR.[S] | 1xCarbamidomethyl [C7]; 1xHomocysteineIAA [K8] | 4,32935E-05 | 2676,2352 | Brain/ FAIMS/IAA |
| Q6PIE5 | Q6PIE5 [429-442] | [R].AVFKAGQENISVSK.[R] | 1xHomocysteineMMTS [K4] | 4,49774E-05 | 1640,81347 | Brain/ FAIMS/MMTS |
| Q6PIE5 | Q6PIE5 [565-579] | [R].GFKFDTDELNFPTEK.[L] | 1xHomocysteineMMTS [K3] | 0,00233521 | 1950,86121 | Brain/ FAIMS/MMTS |
| Q6Q477 | Q6Q477 [589-600] | [R].MFSKGASEIMLR.[R] | 1xHomocysteineIAA [K4] | 0,000429391 | 1543,74295 | Brain/ FAIMS/IAA |
| Q6Q477; G5E829 | Q6Q477 [234-248]; G5E829 [238-252] | [K].IDESSLTGESDHVKK.[TS] | 1xHomocysteineIAA [K14] | 0,000414013 | 1818,85382 | Brain/ FAIMS/IAA |
| Q6WKZ7 | Q6WKZ7 [462-471] | [K].GDIVTVHEKK.[E] | 1xHomocysteineIAA [K9] | 0,00258601 | 1299,67255 | Liver/ No FAIMS/IAA |
| Q6WKZ7 | Q6WKZ7 [462-471] | [K].GDIVTVHEKK.[E] | 1xHomocysteineIAA [K9] | 0,00258601 | 1299,67255 | Liver/ FAIMS/IAA |
| Q6X893 | Q6X893 [89-104] | [R].KYVFFLDPCNLDLINR.[K] | 1xCarbamidomethyl [C9]; 1xHomocysteineIAA [K1] | 0,000318237 | 2201,08819 | Brain/ FAIMS/IAA |
| Q6ZQ38 | Q6ZQ38 [352-364] | [R].AAAKCLDAVVSTR.[H] | 1xCarbamidomethyl [C5]; 1xHomocysteineIAA [K4] | 9,61721E-05 | 1535,76686 | Brain/ FAIMS/IAA |
| Q6ZWN5 | Q6ZWN5 [92-101] | [K].MKLDYILGLK.[I] | 1xHomocysteineIAA [K2] | 0,00012645 | 1367,74254 | Brain/ FAIMS/IAA |
| Q6ZWN5 | Q6ZWN5 [117-127] | [K].LGLAKSIHHAR.[V] | 1xHomocysteineMMTS [K5] | 0,00510827 | 1365,72421 | Liver/ No FAIMS/MMTS |
| Q6ZWN5 | Q6ZWN5 [84-93] | [R].IGVLDEGKMK.[L] | 1xHomocysteineMMTS [K8] | 0,00180518 | 1252,60981 | Liver/ FAIMS/MMTS |
| Q6ZWN5 | Q6ZWN5 [110-121] | [R].LQTQVFKLGLAK.[S] | 1xHomocysteineMMTS [K7] | 0,00750949 | 1508,83275 | Liver/ FAIMS/MMTS |
| Q6ZWN5 | Q6ZWN5 [151-162] | [R].LDSQKHIDFSLR.[S] | 1xHomocysteineMMTS [K5] | 0,00344412 | 1621,78251 | Liver/ FAIMS/MMTS |
| Q6ZWV3 | Q6ZWV3 [99-110] | [R].INKMLSCAGADR.[L] | 1xCarbamidomethyl [C7]; 1xHomocysteineIAA [K3] | 9,61721E-05 | 1509,69706 | Liver/ FAIMS/IAA |
| Q6ZWV3 | Q6ZWV3 [99-110] | [R].INKMLSCAGADR.[L] | 1xCarbamidomethyl [C7]; 1xHomocysteineIAA [K3] | 9,61721E-05 | 1509,69706 | Brain/ FAIMS/IAA |
| Q6ZWV3 | Q6ZWV3 [117-128] | [R].GAFGKPQGTVAR.[V] | 1xHomocysteineMMTS [K5] | 0,000266738 | 1351,66094 | Liver/ FAIMS/MMTS |
| Q6ZWV3 | Q6ZWV3 [176-189] | [K].FNADEFEDMVAEKR.[L] | 1xHomocysteineMMTS [K13] | 0,000829616 | 1863,77102 | Liver/ FAIMS/MMTS |
| Q6ZWV7 | Q6ZWV7 [36-46] | [K].VTGGAASKLSK.[I] | 1xHomocysteineMMTS [K8] | 0,00367611 | 1181,60169 | Liver/ FAIMS/MMTS |
| Q71M36 | Q71M36 [399-411] | [R].CNTQDYIWHKGMR.[C] | 1xCarbamidomethyl [C1]; 1xHomocysteineIAA [K10] | 0,00146914 | 1882,81455 | Brain/ FAIMS/IAA |
| Q76MZ3 | Q76MZ3 [547-566] | [K].IGPILDNSTLQSEVKPILEK.[L] | 1xHomocysteineIAA [K15] | 4,32935E-05 | 2368,27922 | Brain/ FAIMS/IAA |
| Q76MZ3; Q7TNP2 | Q76MZ3 [34-46]; Q7TNP2 [46-58] | [K].KLSTIALALGVER.[T] | 1xHomocysteineIAA [K1] | 0,00107087 | 1544,88287 | Brain/ FAIMS/IAA |
| Q78IK2 | Q78IK2 [1-17] | [-].MAGAESDGQFQFTGIKK.[Y] | 1xMet-loss [N-Term]; 1xHomocysteineIAA [K16] | 0,000902509 | 1857,87997 | Brain/ FAIMS/IAA |
| Q78JT3 | Q78JT3 [156-175] | [R].TGKPNPDQLLKELPFPLNTR.[S] | 1xHomocysteineMMTS [K11] | 0,00230389 | 2441,26795 | Liver/ FAIMS/MMTS |
| Q78ZA7 | Q78ZA7 [67-76] | [R].INALKQLQVR.[C] | 1xHomocysteineIAA [K5] | 4,32935E-05 | 1356,77801 | Brain/ FAIMS/IAA |
| Q7TMF3 | Q7TMF3 [1-9] | [-].MELVEVLKR.[G] | 1xAcetyl [N-Term]; 1xHomocysteineIAA [K8] | 4,32935E-05 | 1332,7014 | Brain/ No FAIMS/IAA |
| Q7TMF3 | Q7TMF3 [1-9] | [-].MELVEVLKR.[G] | 1xAcetyl [N-Term]; 1xHomocysteineIAA [K8] | 4,32935E-05 | 1332,7014 | Brain/ FAIMS/IAA |
| Q7TMM9 | Q7TMM9 [47-62] | [R].INVYYNEAAGNKYVPR.[A] | 1xHomocysteineIAA [K12] | 4,32935E-05 | 2044,99092 | Brain/ No FAIMS/IAA |
| Q7TMM9 | Q7TMM9 [47-62] | [R].INVYYNEAAGNKYVPR.[A] | 1xHomocysteineIAA [K12] | 4,32935E-05 | 2044,99092 | Brain/ FAIMS/IAA |
| Q7TMM9 | Q7TMM9 [47-62] | [R].INVYYNEAAGNKYVPR.[A] | 1xHomocysteineMMTS [K12] | 4,49774E-05 | 2033,95718 | Brain/ No FAIMS/MMTS |
| Q7TMM9 | Q7TMM9 [47-62] | [R].INVYYNEAAGNKYVPR.[A] | 1xHomocysteineMMTS [K12] | 4,49774E-05 | 2033,95718 | Brain/ FAIMS/MMTS |
| Q7TMM9; P68372; Q9CWF2 | Q7TMM9 [360-379]; P68372 [360-379]; Q9CWF2 [360-379] | [R].GLKMSATFIGNSTAIQELFK.[R] | 1xHomocysteineIAA [K3] | 4,32935E-05 | 2330,1883 | Brain/ FAIMS/IAA |
| Q7TMM9; P68372; Q9CWF2 | Q7TMM9 [363-380]; P68372 [363-380]; Q9CWF2 [363-380] | [K].MSATFIGNSTAIQELFKR.[I] | 1xOxidation [M1]; 1xHomocysteineMMTS [K17] | 0,00635755 | 2193,05009 | Liver/ FAIMS/MMTS |
| Q7TMM9; P68372; Q9CWF2 | Q7TMM9 [363-380]; P68372 [363-380]; Q9CWF2 [363-380] | [K].MSATFIGNSTAIQELFKR.[I] | 1xOxidation [M1]; 1xHomocysteineMMTS [K17] | 0,00635755 | 2193,05009 | Brain/ FAIMS/MMTS |
| Q7TMM9; P68372; Q9CWF2 | Q7TMM9 [360-379]; P68372 [360-379]; Q9CWF2 [360-379] | [R].GLKMSATFIGNSTAIQELFK.[R] | 1xHomocysteineMMTS [K3] | 4,49774E-05 | 2319,15456 | Brain/ FAIMS/MMTS |
| Q7TMM9; P68372; Q9CWF2 | Q7TMM9 [363-380]; P68372 [363-380]; Q9CWF2 [363-380] | [K].MSATFIGNSTAIQELFKR.[I] | 1xHomocysteineMMTS [K17] | 0,000530332 | 2177,05518 | Brain/ FAIMS/MMTS |
| Q7TMM9; P99024; P68372; Q9CWF2 | Q7TMM9 [321-336]; P99024 [321-336]; P68372 [321-336]; Q9CWF2 [321-336] | [R].MSMKEVDEQMLNVQNK.[N] | 2xOxidation [M3; M10]; 1xHomocysteineIAA [K4] | 4,32935E-05 | 2129,93341 | Brain/ FAIMS/IAA |
| Q7TMM9; P99024; P68372; Q9CWF2 | Q7TMM9 [321-336]; P99024 [321-336]; P68372 [321-336]; Q9CWF2 [321-336] | [R].MSMKEVDEQMLNVQNK.[N] | 1xHomocysteineMMTS [K4] | 4,49774E-05 | 2086,90983 | Brain/ No FAIMS/MMTS |
| Q7TMM9; P99024; P68372; Q9CWF2 | Q7TMM9 [321-336]; P99024 [321-336]; P68372 [321-336]; Q9CWF2 [321-336] | [R].MSMKEVDEQMLNVQNK.[N] | 1xHomocysteineMMTS [K4] | 4,49774E-05 | 2086,90983 | Brain/ FAIMS/MMTS |
| Q7TMM9; P99024; P68372; Q9CWF2 | Q7TMM9 [325-350]; P99024 [325-350]; P68372 [325-350]; Q9CWF2 [325-350] | [K].EVDEQMLNVQNKNSSYFVEWIPNNVK.[ T] | 1xHomocysteineMMTS [K12] | 4,49774E-05 | 3287,51698 | Brain/ FAIMS/MMTS |
| Q7TMM9; Q9CWF2 | Q7TMM9 [122-154]; Q9CWF2 [122-154] | [R].KESESCDCLQGFQLTHSLGGGTGSGMGT LLISK.[I] | 2xCarbamidomethyl [C6; C8]; 1xHomocysteineIAA [K1] | 4,32935E-05 | 3629,67026 | Brain/ No FAIMS/IAA |
| Q7TMM9; Q9CWF2 | Q7TMM9 [122-154]; Q9CWF2 [122-154] | [R].KESESCDCLQGFQLTHSLGGGTGSGMGT LLISK.[I] | 2xCarbamidomethyl [C6; C8]; 1xHomocysteineIAA [K1] | 4,32935E-05 | 3629,67026 | Brain/ FAIMS/IAA |

| | | | | | | |
|---|---|---|---|---|---|---|
| Q7TMM9; Q9D6F9; P99024; P68372; Q9CWF2 | Q7TMM9 [337-359]; Q9D6F9 [337-359]; P99024 [337-359]; P68372 [337-359]; Q9CWF2 [337-359] | [K].NSSYFVEWIPNNVKTAVCDIPPR.[G] | 1xCarbamidomethyl [C18]; 1xHomocysteineIAA [K14] | 4,32935E-05 | 2880,38074 | Brain/ No FAIMS/IAA |
| Q7TMM9; Q9D6F9; P99024; P68372; Q9CWF2 | Q7TMM9 [337-359]; Q9D6F9 [337-359]; P99024 [337-359]; P68372 [337-359]; Q9CWF2 [337-359] | [K].NSSYFVEWIPNNVKTAVCDIPPR.[G] | 1xCarbamidomethyl [C18]; 1xHomocysteineIAA [K14] | 4,32935E-05 | 2880,38074 | Brain/ FAIMS/IAA |
| Q7TMM9; Q9D6F9; Q9ERD7; A2AQ07; P99024; P68372; Q9CWF2; Q922F4 | Q7TMM9 [252-262]; Q9D6F9 [252-262]; Q9ERD7 [252-262]; A2AQ07 [252-262]; P99024 [252-262]; P68372 [252-262]; Q9CWF2 [252-262]; Q922F4 [252-262] | [R].KLAVNMVPFPR.[L] | 1xHomocysteineIAA [K1] | 4,32935E-05 | 1445,77557 | Brain/ No FAIMS/IAA |
| Q7TMM9; Q9D6F9; Q9ERD7; A2AQ07; P99024; P68372; Q9CWF2; Q922F4 | Q7TMM9 [252-262]; Q9D6F9 [252-262]; Q9ERD7 [252-262]; A2AQ07 [252-262]; P99024 [252-262]; P68372 [252-262]; Q9CWF2 [252-262]; Q922F4 [252-262] | [R].KLAVNMVPFPR.[L] | 1xHomocysteineIAA [K1] | 4,32935E-05 | 1445,77557 | Brain/ FAIMS/IAA |
| Q7TMM9; Q9D6F9; Q9ERD7; P99024; P68372; Q9CWF2; Q922F4 | Q7TMM9 [252-262]; Q9D6F9 [252-262]; Q9ERD7 [252-262]; P99024 [252-262]; P68372 [252-262]; Q9CWF2 [252-262]; Q922F4 [252-262] | [R].KLAVNMVPFPR.[L] | 1xHomocysteineMMTS [K1] | 0,000456432 | 1434,74183 | Brain/ No FAIMS/MMTS |
| Q7TMM9; Q9D6F9; Q9ERD7; P99024; P68372; Q9CWF2; Q922F4 | Q7TMM9 [252-262]; Q9D6F9 [252-262]; Q9ERD7 [252-262]; P99024 [252-262]; P68372 [252-262]; Q9CWF2 [252-262]; Q922F4 [252-262] | [R].KLAVNMVPFPR.[L] | 1xHomocysteineMMTS [K1] | 0,000456432 | 1434,74183 | Brain/ FAIMS/MMTS |
| Q7TNP2 | Q7TNP2 [170-179] | [R].ASNAVKAEIR.[Q] | 1xHomocysteineIAA [K6] | 9,61721E-05 | 1232,64158 | Brain/ FAIMS/IAA |
| Q7TQD2 | Q7TQD2 [89-103] | [K].NVTVTDVDIVFSKIK.[G] | 1xHomocysteineIAA [K13] | 4,32935E-05 | 1851,98846 | Brain/ No FAIMS/IAA |
| Q7TQD2 | Q7TQD2 [89-103] | [K].NVTVTDVDIVFSKIK.[G] | 1xHomocysteineIAA [K13] | 4,32935E-05 | 1851,98846 | Brain/ FAIMS/IAA |
| Q7TQD2 | Q7TQD2 [147-164] | [R].APVISGVTKAVSSPTVSR.[L] | 1xHomocysteineMMTS [K9] | 4,49774E-05 | 1919,00888 | Brain/ No FAIMS/MMTS |
| Q7TQD2 | Q7TQD2 [89-103] | [K].NVTVTDVDIVFSKIK.[G] | 1xHomocysteineMMTS [K13] | 4,49774E-05 | 1840,95472 | Brain/ No FAIMS/MMTS |
| Q7TQD2 | Q7TQD2 [147-164] | [R].APVISGVTKAVSSPTVSR.[L] | 1xHomocysteineMMTS [K9] | 4,49774E-05 | 1919,00888 | Brain/ FAIMS/MMTS |
| Q7TQD2 | Q7TQD2 [15-25] | [K].TPPKSPGDPAR.[A] | 1xHomocysteineMMTS [K4] | 0,00924953 | 1285,60275 | Brain/ FAIMS/MMTS |
| Q7TQD2 | Q7TQD2 [192-213] | [R].VDLVDESGYVPGYKHAGTYDQK.[V] | 1xHomocysteineMMTS [K14] | 4,49774E-05 | 2604,1745 | Brain/ FAIMS/MMTS |
| Q7TQD2 | Q7TQD2 [89-103] | [K].NVTVTDVDIVFSKIK.[G] | 1xHomocysteineMMTS [K13] | 4,49774E-05 | 1840,95472 | Brain/ FAIMS/MMTS |
| Q7TQF7 | Q7TQF7 [62-76] | [R].GYLAAIKGMQEASMK.[L] | 1xHomocysteineIAA [K7] | 4,32935E-05 | 1771,85396 | Brain/ FAIMS/IAA |
| Q7TQF7 | Q7TQF7 [1-11] | [-].MADIKTGIFAK.[N] | 1xMet-loss+Acetyl [N-Term]; 1xHomocysteineMMTS [K5] | 0,0025135 | 1268,63774 | Brain/ FAIMS/MMTS |
| Q7TQF7 | Q7TQF7 [62-76] | [R].GYLAAIKGMQEASMK.[L] | 1xHomocysteineMMTS [K7] | 4,49774E-05 | 1760,82022 | Brain/ FAIMS/MMTS |
| Q7TQF7 | Q7TQF7 [669-683] | [R].DLATYKGLFPENFTR.[R] | 1xHomocysteineMMTS [K6] | 0,000231655 | 1934,91392 | Brain/ FAIMS/MMTS |
| Q7TQF7 | Q7TQF7 [28-47] | [K].LGKADETKDEQFEEYVQNFK.[R] | 1xHomocysteineMMTS [K3] | 0,00593578 | 2581,15852 | Brain/ FAIMS/MMTS |
| Q7TQI3 | Q7TQI3 [257-271] | [K].VYLLYRPGHYDILYK.[-] | 1xHomocysteineIAA [K15] | 4,32935E-05 | 2087,07828 | Brain/ No FAIMS/IAA |
| Q7TQI3 | Q7TQI3 [257-271] | [K].VYLLYRPGHYDILYK.[-] | 1xHomocysteineIAA [K15] | 4,32935E-05 | 2087,07828 | Brain/ FAIMS/IAA |
| Q7TQI3 | Q7TQI3 [186-198] | [R].ESKFFEHFIEGGR.[T] | 1xHomocysteineMMTS [K3] | 0,000282457 | 1745,77742 | Brain/ FAIMS/MMTS |
| Q7TSJ2 | Q7TSJ2 [696-716] | [K].NQSPVVPASTKDQSFPTPAPR.[K] | 1xHomocysteineIAA [K11] | 4,32935E-05 | 2398,18197 | Brain/ FAIMS/IAA |
| Q80SW1; Q68FL4 | Q80SW1 [268-284]; Q68FL4 [351-367] | [K].AGKLCVPAMNVNDSVTK.[Q] | 1xCarbamidomethyl [C5]; 1xHomocysteineIAA [K3] | 0,00263986 | 1977,95546 | Brain/ FAIMS/IAA |
| Q80SZ7 | Q80SZ7 [1-12] | [-].MSGSSSVAAMKK.[V] | 1xMet-loss+Acetyl [N-Term]; 1xHomocysteineIAA [K12] | 0,00286445 | 1268,59733 | Brain/ FAIMS/IAA |
| Q80TB8 | Q80TB8 [230-239] | [R].NADYVQEVKR.[I] | 1xHomocysteineIAA [K9] | 0,0027602 | 1395,66852 | Brain/ FAIMS/IAA |
| Q80TJ1 | Q80TJ1 [20-32] | [R].GKDVLGSAASGAR.[L] | 1xHomocysteineIAA [K2] | 0,0014138 | 1362,67942 | Brain/ FAIMS/IAA |
| Q80TL4 | Q80TL4 [356-369] | [K].SQEKALLPTEQESR.[Y] | 1xHomocysteineIAA [K4] | 0,000273428 | 1789,87489 | Brain/ FAIMS/IAA |
| Q80U72 | Q80U72 [1092-1102] | [K].APGEKLGISIR.[G] | 1xHomocysteineIAA [K5] | 0,00777414 | 1314,71983 | Brain/ FAIMS/IAA |
| Q80UK0; P58774 | Q80UK0 [419-425]; P58774 [227-233] | [K].LLEEKLK.[SE] | 1xHomocysteineIAA [K5] | 0,00399809 | 1046,59144 | Liver/ FAIMS/IAA |
| Q80UW2 | Q80UW2 [82-95] | [R].WKELVDGAPLWLLK.[C] | 1xHomocysteineIAA [K2] | 0,000210768 | 1841,99824 | Brain/ No FAIMS/IAA |
| Q80UW2 | Q80UW2 [82-95] | [R].WKELVDGAPLWLLK.[C] | 1xHomocysteineIAA [K2] | 0,000210768 | 1841,99824 | Brain/ FAIMS/IAA |
| Q80V26 | Q80V26 [98-113] | [R].EGADDKMTSGDVLSNR.[K] | 1xHomocysteineIAA [K6] | 0,00540576 | 1868,8113 | Brain/ FAIMS/IAA |
| Q80VP1 | Q80VP1 [92-107] | [R].VSQQCKENMYAVQTLK.[D] | 1xCarbamidomethyl [C5]; 1xHomocysteineIAA [K6] | 4,32935E-05 | 2100,98749 | Brain/ FAIMS/IAA |
| Q80X56 | Q80X56 [188-197] | [K].DAISAYKDNK.[I] | 1xHomocysteineIAA [K7] | 0,00818027 | 1298,60453 | Brain/ FAIMS/IAA |
| Q80X90 | Q80X90 [2005-2016] | [R].AKVYGQGLSEGR.[T] | 1xHomocysteineMMTS [K2] | 0,00534072 | 1427,67698 | Liver/ FAIMS/MMTS |
| Q80YN3 | Q80YN3 [369-380] | [K].SDKANFTPQETR.[G] | 1xHomocysteineIAA [K3] | 9,61721E-05 | 1567,71693 | Brain/ FAIMS/IAA |
| Q80Z24 | Q80Z24 [213-233] | [R].VIVNFAPTIQEIKSGTVTPGR.[S] | 1xHomocysteineIAA [K13] | 4,32935E-05 | 2401,29079 | Brain/ No FAIMS/IAA |
| Q80ZJ1 | Q80ZJ1 [106-117] | [R].YEKVPVILVGNK.[V] | 1xHomocysteineIAA [K3] | 6,30912E-05 | 1532,85051 | Brain/ FAIMS/IAA |
| Q810U3 | Q810U3 [553-569] | [K].LTVSWLKDDEPLYIGNR.[M] | 1xHomocysteineIAA [K7] | 4,32935E-05 | 2193,10086 | Brain/ FAIMS/IAA |
| Q810U3 | Q810U3 [335-355] | [K].AAPYWLDEPKNLILAPGEDGR.[L] | 1xHomocysteineMMTS [K10] | 0,00939394 | 2488,19993 | Brain/ FAIMS/MMTS |
| Q810U4 | Q810U4 [542-553] | [R].IIKQPEYAVVQR.[G] | 1xHomocysteineIAA [K3] | 4,32935E-05 | 1617,87812 | Brain/ FAIMS/IAA |

| | | | | | | |
|---|---|---|---|---|---|---|
| Q811F1 | Q811F1 [63-76] | [R].KLLSSLQYNKNLLK.[Y] | 1xHomocysteineMMTS [K10] 1xCarbamidomethyl [C5]; | 0,000738471 | 1825,00742 | Brain/ FAIMS/MMTS |
| Q8BFR5 | Q8BFR5 [286-300] | [K].KGDECELLGHNKNIR.[T] | 1xHomocysteineIAA [K12] | 4,32935E-05 | 1956,93784 | Brain/ FAIMS/IAA |
| Q8BFR5 | Q8BFR5 [91-102] | [K].KYEEIDNAPEER.[A] | 1xHomocysteineMMTS [K1] | 0,000192903 | 1655,70398 | Liver/ FAIMS/MMTS |
| Q8BFU2 | Q8BFU2 [90-100] | [R].NDEELNKLLGR.[V] | 1xHomocysteineIAA [K7] | 4,32935E-05 | 1474,73185 | Brain/ No FAIMS/IAA |
| Q8BFU2 | Q8BFU2 [90-100] | [R].NDEELNKLLGR.[V] | 1xHomocysteineIAA [K7] | 4,32935E-05 | 1474,73185 | Brain/ FAIMS/IAA |
| Q8BFU2 | Q8BFU2 [90-100] | [R].NDEELNKLLGR.[V] | 1xHomocysteineMMTS [K7] | 4,49774E-05 | 1463,69811 | Liver/ FAIMS/MMTS |
| Q8BFU2 | Q8BFU2 [90-100] | [R].NDEELNKLLGR.[V] | 1xHomocysteineMMTS [K7] | 4,49774E-05 | 1463,69811 | Brain/ FAIMS/MMTS |
| Q8BFU3 | Q8BFU3 [348-359] | [R].AWKAEILSLESR.[K] | 1xHomocysteineIAA [K3] 1xCarbamidomethyl [C13]; | 0,000177335 | 1576,81519 | Brain/ FAIMS/IAA |
| Q8BFZ3 | Q8BFZ3 [41-62] | [R].HQGVMVGMGQKDCYVGDEAQSK.[R] | 1xHomocysteineIAA [K11] | 0,000273428 | 2598,12037 | Brain/ No FAIMS/IAA |
| Q8BFZ3; P60710; P62737; P68134; P63260 | Q8BFZ3 [328-336]; P60710 [327-335]; P62737 [329-337]; P68134 [329-337]; P63260 [327-335] | [K].IKIIAPPER.[K] | 1xHomocysteineIAA [K2] | 0,00012645 | 1210,69764 | Liver/ FAIMS/IAA |
| Q8BFZ3; P60710; P62737; P68134; P63260 | Q8BFZ3 [328-337]; P60710 [327-336]; P62737 [329-338]; P68134 [329-338]; P63260 [327-336] | [K].IKIIAPPERK.[Y] | 1xHomocysteineIAA [K2] | 4,32935E-05 | 1338,7926 | Brain/ FAIMS/IAA |
| Q8BFZ3; P60710; P62737; P68134; P63260 | Q8BFZ3 [328-336]; P60710 [327-335]; P62737 [329-337]; P68134 [329-337]; P63260 [327-335] | [K].IKIIAPPER.[K] | 1xHomocysteineMMTS [K2] | 0,000929608 | 1199,6639 | Liver/ No FAIMS/MMTS |
| Q8BFZ3; P60710; P62737; P68134; P63260 | Q8BFZ3 [328-336]; P60710 [327-335]; P62737 [329-337]; P68134 [329-337]; P63260 [327-335] | [K].IKIIAPPER.[K] | 1xHomocysteineMMTS [K2] | 0,000929608 | 1199,6639 | Liver/ FAIMS/MMTS |
| Q8BFZ3; P60710; P62737; P68134; P63260 | Q8BFZ3 [185-197]; P60710 [184-196]; P62737 [186-198]; P68134 [186-198]; P63260 [184-196] | [R].DLTDYLMKILTER.[G] | 1xHomocysteineMMTS [K8] | 0,00760806 | 1773,85837 | Brain/ No FAIMS/MMTS |
| Q8BFZ3; P60710; P62737; P68134; P63260 | Q8BFZ3 [328-336]; P60710 [327-335]; P62737 [329-337]; P68134 [329-337]; P63260 [327-335] | [K].IKIIAPPER.[K] | 1xHomocysteineMMTS [K2] | 0,000929608 | 1199,6639 | Brain/ No FAIMS/MMTS |
| Q8BFZ3; P60710; P62737; P68134; P63260 | Q8BFZ3 [328-336]; P60710 [327-335]; P62737 [329-337]; P68134 [329-337]; P63260 [327-335] | [K].IKIIAPPER.[K] | 1xHomocysteineMMTS [K2] | 0,000929608 | 1199,6639 | Brain/ FAIMS/MMTS |
| Q8BFZ9; Q91X78 | Q8BFZ9 [193-201]; Q91X78 [195-203] | [K].LLIAAQKQK.[V] | 1xHomocysteineIAA [K7] | 0,000318237 | 1186,69764 | Brain/ FAIMS/IAA |
| Q8BG05 | Q8BG05 [128-143] | [K].IFVGGIKEDTEEYNLR.[D] | 1xHomocysteineIAA [K7] | 4,32935E-05 | 2057,00082 | Brain/ No FAIMS/IAA |
| Q8BG05 | Q8BG05 [128-143] | [K].IFVGGIKEDTEEYNLR.[D] | 1xHomocysteineIAA [K7] | 4,32935E-05 | 2057,00082 | Brain/ FAIMS/IAA |
| Q8BG05 | Q8BG05 [149-161] | [K].YGKIETIEVMEDR.[Q] | 1xHomocysteineMMTS [K3] | 0,000530332 | 1745,79069 | Liver/ FAIMS/MMTS |
| Q8BG05 | Q8BG05 [149-161] | [K].YGKIETIEVMEDR.[Q] | 1xHomocysteineMMTS [K3] | 0,000530332 | 1745,79069 | Brain/ FAIMS/MMTS |
| Q8BG39 | Q8BG39 [462-476] | [K].MYFKHVLFEDTFFDK.[C] | 1xHomocysteineIAA [K4] | 4,32935E-05 | 2140,98708 | Brain/ FAIMS/IAA |
| Q8BGN8 | Q8BGN8 [210-226] | [R].YLSDPMEKHSSSYNQGR.[Y] | 1xHomocysteineIAA [K8] | 4,32935E-05 | 2172,94371 | Brain/ No FAIMS/IAA |
| Q8BGN8 | Q8BGN8 [210-226] | [R].YLSDPMEKHSSSYNQGR.[Y] | 1xHomocysteineIAA [K8] | 4,32935E-05 | 2172,94371 | Brain/ FAIMS/IAA |
| Q8BGQ7 | Q8BGQ7 [913-930] | [R].GLKASEWVQQVSGLMDGK.[G] | 1xHomocysteineIAA [K3] | 4,32935E-05 | 2107,03107 | Brain/ FAIMS/IAA |
| Q8BGR9 | Q8BGR9 [30-42] | [K].QFLKTLTGVLPER.[Q] | 1xHomocysteineIAA [K4] | 0,00913964 | 1675,91999 | Brain/ FAIMS/IAA |
| Q8BGU5 | Q8BGU5 [280-301] | [R].SLAEANNLSFPLEPLSRERAHK.[L] | 1xHomocysteineMMTS [K22] | 0,00125756 | 2642,31775 | Brain/ No FAIMS/MMTS |
| Q8BGY2; P63242 | Q8BGY2 [35-47]; P63242 [35-47] | [K].GRPCKIVEMSTSK.[T] | 1xCarbamidomethyl [C4]; 1xHomocysteineIAA [K5] | 4,32935E-05 | 1666,80734 | Liver/ FAIMS/IAA |
| Q8BGY2; P63242 | Q8BGY2 [68-86]; P63242 [68-86] | [K].KYEDICPSTHNMDVPNIKR.[N] | 1xCarbamidomethyl [C6]; 1xHomocysteineIAA [K18] | 4,32935E-05 | 2491,15266 | Brain/ FAIMS/IAA |
| Q8BGZ1 | Q8BGZ1 [4-28] | [K].NNSKLAPEVLEDLVQNTEFSEQELK.[Q] | 1xHomocysteineIAA [K4] | 4,32935E-05 | 3048,48302 | Brain/ FAIMS/IAA |
| Q8BGZ1; P62761 | Q8BGZ1 [55-70]; P62761 [55-70] | [K].FFPYGDASKFAQHAFR.[T] | 1xHomocysteineIAA [K9] | 4,32935E-05 | 2062,95923 | Brain/ No FAIMS/IAA |
| Q8BGZ1; P62761 | Q8BGZ1 [55-70]; P62761 [55-70] | [K].FFPYGDASKFAQHAFR.[T] | 1xHomocysteineIAA [K9] | 4,32935E-05 | 2062,95923 | Brain/ FAIMS/IAA |
| Q8BGZ1; P62761 | Q8BGZ1 [55-70]; P62761 [55-70] | [K].FFPYGDASKFAQHAFR.[T] | 1xHomocysteineMMTS [K9] | 0,000400718 | 2051,92548 | Brain/ FAIMS/MMTS |
| Q8BH95 | Q8BH95 [198-211] | [R].ISAQDAKQAGLVSK.[I] | 1xHomocysteineIAA [K7] | 4,32935E-05 | 1589,83157 | Liver/ FAIMS/IAA |
| Q8BH95 | Q8BH95 [198-211] | [R].ISAQDAKQAGLVSK.[I] | 1xHomocysteineIAA [K7] | 4,32935E-05 | 1589,83157 | Brain/ FAIMS/IAA |
| Q8BH95 | Q8BH95 [198-211] | [R].ISAQDAKQAGLVSK.[I] | 1xHomocysteineMMTS [K7] | 4,49774E-05 | 1578,79782 | Liver/ No FAIMS/MMTS |
| Q8BH95 | Q8BH95 [182-197] | [R].AVGKSLAMEMVLTGDR.[I] | 1xHomocysteineMMTS [K4] 1xOxidation [M]; | 0,000437433 | 1840,87879 | Liver/ No FAIMS/MMTS |
| Q8BH95 | Q8BH95 [182-197] | [R].AVGKSLAMEMVLTGDR.[I] | 1xHomocysteineMMTS [K4] | 4,49774E-05 | 1856,87371 | Liver/ FAIMS/MMTS |
| Q8BH95 | Q8BH95 [198-211] | [R].ISAQDAKQAGLVSK.[I] | 1xHomocysteineMMTS [K7] 2xOxidation [M8; M10]; | 4,49774E-05 | 1578,79782 | Liver/ FAIMS/MMTS |
| Q8BH95 | Q8BH95 [182-197] | [R].AVGKSLAMEMVLTGDR.[I] | 1xHomocysteineMMTS [K4] | 0,000282457 | 1872,86862 | Liver/ FAIMS/MMTS |
| Q8BH95 | Q8BH95 [93-106] | [K].AFAAGADIKEMQNR.[T] | 1xHomocysteineMMTS [K9] | 4,49774E-05 | 1684,76039 | Liver/ FAIMS/MMTS |
| Q8BH95 | Q8BH95 [182-197] | [R].AVGKSLAMEMVLTGDR.[I] | 1xHomocysteineMMTS [K4] | 0,000437433 | 1840,87879 | Liver/ FAIMS/MMTS |
| Q8BHN3 | Q8BHN3 [268-278] | [R].LKVTEGGEPYR.[L] | 1xHomocysteineIAA [K2] | 0,00156934 | 1422,70458 | Liver/ FAIMS/IAA |

| | | | | | | |
|---|---|---|---|---|---|---|
| Q8BHN3 | Q8BHN3 [446-460] | [R].FPQPLNMLEHLASKR.[R] | 1xHomocysteineIAA [K14] | 4,32935E-05 | 1954,99898 | Brain/ FAIMS/IAA |
| Q8BI22 | Q8BI22 [1049-1064] | [R].FLSSPQFSHSLPVFTK.[R] | 1xHomocysteineIAA [K16] | 0,000318237 | 1995,99969 | Liver/ No FAIMS/IAA |
| Q8BI22 | Q8BI22 [1049-1064] | [R].FLSSPQFSHSLPVFTK.[R] | 1xHomocysteineIAA [K16] | 0,000318237 | 1995,99969 | Liver/ FAIMS/IAA |
| Q8BJI1 | Q8BJI1 [35-49] | [K].QSVLNVAGETGGKQK.[V] | 1xHomocysteineIAA [K13] | 0,000258769 | 1689,85884 | Brain/ FAIMS/IAA |
| Q8BJU0 | Q8BJU0 [152-165] | [R].AIGIDPGYSKAYGR.[M] | 1xHomocysteineIAA [K10] | 4,32935E-05 | 1641,80535 | Brain/ FAIMS/IAA |
| Q8BK48 | Q8BK48 [367-378] | [K].TLPAVLKSTALK.[M] | 1xHomocysteineMMTS [K7] | 0,000211896 | 1404,7953 | Liver/ FAIMS/MMTS |
| Q8BK64 | Q8BK64 [322-336] | [R].YYFEGIKQTFGYGAR.[L] | 1xHomocysteineIAA [K7] | 0,00097531 | 1973,92144 | Brain/ FAIMS/IAA |
| Q8BKX1 | Q8BKX1 [380-398] | [R].VKAIFSHAAGDNSTLLSFK.[E] | 1xHomocysteineIAA [K2] | 4,32935E-05 | 2180,11685 | Brain/ FAIMS/IAA |
| Q8BKZ9 | Q8BKZ9 [219-227] | [K].LVELKQMGK.[I] | 1xHomocysteineIAA [K5] | 4,32935E-05 | 1219,65373 | Brain/ FAIMS/IAA |
| Q8BL66 | Q8BL66 [187-202] | [R].AAAEQKVTHLTEDLNK.[Q] | 1xHomocysteineIAA [K6] | 9,61721E-05 | 1941,96985 | Brain/ FAIMS/IAA |
| Q8BLK3 | Q8BLK3 [234-253] | [R].QASLKCEASAVPAPDFEWYR.[D] | 1xCarbamidomethyl [C6]; 1xHomocysteineIAA [K5] | 4,32935E-05 | 2499,14314 | Brain/ No FAIMS/IAA |
| Q8BLK3 | Q8BLK3 [234-253] | [R].QASLKCEASAVPAPDFEWYR.[D] | 1xCarbamidomethyl [C6]; 1xHomocysteineIAA [K5] | 4,32935E-05 | 2499,14314 | Brain/ FAIMS/IAA |
| Q8BLQ9 | Q8BLQ9 [180-191] | [R].KTFTVSSTLDFR.[V] | 1xHomocysteineIAA [K1] | 4,32935E-05 | 1575,78355 | Brain/ No FAIMS/IAA |
| Q8BLQ9 | Q8BLQ9 [180-191] | [R].KTFTVSSTLDFR.[V] | 1xHomocysteineIAA [K1] | 4,32935E-05 | 1575,78355 | Brain/ FAIMS/IAA |
| Q8BMF4 | Q8BMF4 [364-378] | [K].GIDLTQVKGTGPEGR.[I] | 1xHomocysteineIAA [K8] | 4,32935E-05 | 1701,85884 | Brain/ FAIMS/IAA |
| Q8BMK4 | Q8BMK4 [307-319] | [R].ELVSLKQEQQAFK.[Q] | 1xHomocysteineIAA [K6] | 0,00012645 | 1721,88908 | Brain/ FAIMS/IAA |
| Q8BMS1 | Q8BMS1 [387-399] | [K].TLLKDTTVTGLGR.[G] | 1xHomocysteineIAA [K4] | 0,000698725 | 1548,8414 | Brain/ FAIMS/IAA |
| Q8BN59 | Q8BN59 [288-297] | [K].AVLIGMKPPK.[K] | 1xOxidation [M6]; 1xHomocysteineIAA [K7] | 0,00777414 | 1243,69011 | Liver/ FAIMS/IAA |
| Q8BNY6 | Q8BNY6 [103-118] | [R].WAFKLYDLDNDGYITR.[N] | 1xHomocysteineIAA [K4] | 4,32935E-05 | 2164,0168 | Brain/ FAIMS/IAA |
| Q8BP47 | Q8BP47 [66-75] | [R].WDVISKSQMK.[N] | 1xHomocysteineIAA [K6] | 0,00467963 | 1395,67592 | Brain/ FAIMS/IAA |
| Q8BP67 | Q8BP67 [1-12] | [-].MKVELCSFSGYK.[I] | 1xCarbamidomethyl [C6]; 1xHomocysteineIAA [K2] | 0,00621607 | 1622,73753 | Liver/ FAIMS/IAA |
| Q8BP67 | Q8BP67 [28-43] | [K].VFQFLNAKCESAFLSK.[R] | 1xCarbamidomethyl [C9]; 1xHomocysteineIAA [K8] | 4,32935E-05 | 2063,00888 | Brain/ FAIMS/IAA |
| Q8BP67 | Q8BP67 [125-135] | [K].TAMAAAKAPTK.[A] | 1xHomocysteineMMTS [K7] | 0,000192903 | 1223,5945 | Liver/ FAIMS/MMTS |
| Q8BP67 | Q8BP67 [81-94] | [R].AITGASLADIMAKR.[N] | 1xHomocysteineMMTS [K13] | 0,0004709 | 1580,79571 | Liver/ FAIMS/MMTS |
| Q8BP92 | Q8BP92 [285-308] | [K].KLSEEEILENQDLFLTSEATDYGR.[Q] | 1xHomocysteineIAA [K1] | 4,32935E-05 | 2974,39862 | Brain/ FAIMS/IAA |
| Q8BSY0 | Q8BSY0 [114-121] | [K].VLLGLKER.[S] | 1xHomocysteineIAA [K6] | 0,000800823 | 1101,64488 | Brain/ FAIMS/IAA |
| Q8BU31; P61226; Q80ZJ1 | Q8BU31 [42-68]; P61226 [42-68]; Q80ZJ1 [42-68] | [R].KEIEVDSSPSVLEILDTAGTEQFASMR.[D] | 1xHomocysteineIAA [K1] | 4,32935E-05 | 3126,49696 | Brain/ FAIMS/IAA |
| Q8BVE3 | Q8BVE3 [108-117] | [R].VSIFFDYAKR.[S] | 1xHomocysteineIAA [K9] | 0,00049316 | 1419,70893 | Brain/ FAIMS/IAA |
| Q8BVI4 | Q8BVI4 [136-164] | [K].AALDGTPGMIGYGMAKGAVHQLCQSLAGK.[N] | 1xCarbamidomethyl [C23]; 1xHomocysteineIAA [K16] | 0,000960518 | 3076,48351 | Brain/ FAIMS/IAA |
| Q8BVP1 | Q8BVP1 [371-388] | [R].KGNLPKPPGDLFSLIGLK.[I] | 2xHomocysteineMMTS [K6; K18] | 0,00624351 | 2220,14115 | Liver/ FAIMS/MMTS |
| Q8BWF0 | Q8BWF0 [252-267] | [K].AKEVGEVLCTDPLVSK.[I] | 1xCarbamidomethyl [C9]; 1xHomocysteineIAA [K2] | 0,00012645 | 1918,96126 | Brain/ FAIMS/IAA |
| Q8BWR2 | Q8BWR2 [159-173] | [K].NFGADTTKIFYIGLR.[G] | 1xHomocysteineIAA [K8] | 0,00115802 | 1889,95783 | Brain/ FAIMS/IAA |
| Q8BWT1 | Q8BWT1 [270-278] | [K].KHNFTPLAR.[V] | 1xHomocysteineIAA [K1] | 0,00012645 | 1257,65209 | Liver/ No FAIMS/IAA |
| Q8BWT1 | Q8BWT1 [134-143] | [R].FGTKFGLDLK.[L] | 1xHomocysteineIAA [K4] | 0,000177335 | 1299,67657 | Liver/ FAIMS/IAA |
| Q8BWT1 | Q8BWT1 [159-176] | [K].LPMGMTAENLAAKYNISR.[E] | 1xOxidation [M]; 1xHomocysteineMMTS [K13] | 4,49774E-05 | 2159,0116 | Liver/ No FAIMS/MMTS |
| Q8BWT1 | Q8BWT1 [134-143] | [R].FGTKFGLDLK.[L] | 1xHomocysteineMMTS [K4] | 0,00074635 | 1288,64283 | Liver/ No FAIMS/MMTS |
| Q8BWT1 | Q8BWT1 [242-270] | [K].DGTVTAGNASGVSDGAGAVIIASEDAVKK.[H] | 1xHomocysteineMMTS [K28] | 4,49774E-05 | 2823,3499 | Liver/ No FAIMS/MMTS |
| Q8BWT1 | Q8BWT1 [270-278] | [K].KHNFTPLAR.[V] | 1xHomocysteineMMTS [K1] | 0,00579131 | 1246,61834 | Liver/ No FAIMS/MMTS |
| Q8BWT1 | Q8BWT1 [159-176] | [K].LPMGMTAENLAAKYNISR.[E] | 1xOxidation [M]; 1xHomocysteineMMTS [K13] | 4,49774E-05 | 2159,0116 | Liver/ FAIMS/MMTS |
| Q8BWT1 | Q8BWT1 [134-143] | [R].FGTKFGLDLK.[L] | 1xHomocysteineMMTS [K4] | 0,00074635 | 1288,64283 | Liver/ FAIMS/MMTS |
| Q8BWT1 | Q8BWT1 [190-209] | [R].WKAANEAGYFNEEMAPIEVK.[T] | 1xHomocysteineMMTS [K2] | 4,49774E-05 | 2460,10325 | Liver/ FAIMS/MMTS |
| Q8BWT1 | Q8BWT1 [213-234] | [K].GKQTMQVDEHARPQTTLEQLQK.[L] | 1xOxidation [M5]; 1xHomocysteineMMTS [K2] | 0,00524294 | 2745,31168 | Liver/ FAIMS/MMTS |
| Q8BWT1 | Q8BWT1 [190-209] | [R].WKAANEAGYFNEEMAPIEVK.[T] | 1xOxidation [M14]; 1xHomocysteineMMTS [K2] | 4,49774E-05 | 2476,09816 | Liver/ FAIMS/MMTS |
| Q8BWT1 | Q8BWT1 [192-211] | [K].AANEAGYFNEEMAPIEVKTK.[K] | 1xOxidation [M12]; 1xHomocysteineMMTS [K18] | 0,000846174 | 2391,06653 | Liver/ FAIMS/MMTS |
| Q8BWT1 | Q8BWT1 [333-360] | [K].ALDLDPSKTNVSGGAIALGHPLGGSGSR.[I] | 1xHomocysteineMMTS [K8] | 4,49774E-05 | 2810,39237 | Liver/ FAIMS/MMTS |
| Q8BWT1 | Q8BWT1 [242-270] | [K].DGTVTAGNASGVSDGAGAVIIASEDAVKK.[H] | 1xHomocysteineMMTS [K28] | 4,49774E-05 | 2823,3499 | Liver/ FAIMS/MMTS |
| Q8BWT1 | Q8BWT1 [215-240] | [K].QTMQVDEHARPQTTLEQLQKLPSVFK.[K] | 1xHomocysteineMMTS [K20] | 0,00479272 | 3215,60098 | Liver/ FAIMS/MMTS |
| Q8BWT1 | Q8BWT1 [213-234] | [K].GKQTMQVDEHARPQTTLEQLQK.[L] | 1xHomocysteineMMTS [K2] | 0,00669057 | 2729,31676 | Liver/ FAIMS/MMTS |
| Q8BWT1 | Q8BWT1 [192-211] | [K].AANEAGYFNEEMAPIEVKTK.[K] | 1xHomocysteineMMTS [K18] | 4,49774E-05 | 2375,07161 | Liver/ FAIMS/MMTS |
| Q8BWT1 | Q8BWT1 [144-171] | [K].LEDTLWAGLTDQHVKLPMGMTAENLAAK.[Y] | 1xHomocysteineMMTS [K15] | 4,49774E-05 | 3216,55601 | Liver/ FAIMS/MMTS |
| Q8BWT1 | Q8BWT1 [159-176] | [K].LPMGMTAENLAAKYNISR.[E] | 1xHomocysteineMMTS [K13] | 0,000192903 | 2143,01668 | Liver/ FAIMS/MMTS |
| Q8BWT1 | Q8BWT1 [241-269] | [K].KDGTVTAGNASGVSDGAGAVIIASEDAVK.[K] | 1xHomocysteineMMTS [K1] | 4,49774E-05 | 2823,3499 | Liver/ FAIMS/MMTS |
| Q8BYI9 | Q8BYI9 [812-827] | [R].DKEEDMLEVLLDATKR.[H] | 1xHomocysteineIAA [K] | 4,32935E-05 | 2079,00967 | Brain/ No FAIMS/IAA |
| Q8BYI9 | Q8BYI9 [812-827] | [R].DKEEDMLEVLLDATKR.[H] | 1xHomocysteineIAA [K] | 4,32935E-05 | 2079,00967 | Brain/ FAIMS/IAA |
| Q8BYM5 | Q8BYM5 [333-346] | [K].SAKELVEQDIQPAR.[Y] | 1xHomocysteineIAA [K3] | 0,00012645 | 1757,88506 | Brain/ FAIMS/IAA |
| Q8C0E2 | Q8C0E2 [284-294] | [R].YFKQQEVVLWR.[K] | 1xHomocysteineIAA [K3] | 0,00124346 | 1669,85191 | Brain/ FAIMS/IAA |
| Q8C0M9 | Q8C0M9 [195-213] | [R].GNLAYATSTGGIVNKMVGR.[V] | 1xHomocysteineIAA [K15] | 4,32935E-05 | 2083,0423 | Brain/ FAIMS/IAA |
| Q8C0M9 | Q8C0M9 [142-160] | [K].FAEDMGIPQVPVEKLITER.[T] | 1xHomocysteineMMTS [K14] | 0,000250158 | 2335,14947 | Brain/ FAIMS/MMTS |
| Q8C196 | Q8C196 [804-814] | [R].TFEESFQKALR.[M] | 1xHomocysteineMMTS [K8] | 4,49774E-05 | 1518,70795 | Liver/ FAIMS/MMTS |
| Q8C196 | Q8C196 [113-127] | [R].DELGLNKYMESDGIK.[V] | 1xHomocysteineMMTS [K7] | 4,49774E-05 | 1874,83328 | Liver/ FAIMS/MMTS |
| Q8C196 | Q8C196 [1472-1481] | [K].LFAEAVQKSR.[T] | 1xHomocysteineMMTS [K8] | 0,00235089 | 1311,65479 | Liver/ FAIMS/MMTS |

| | | | | | | |
|---|---|---|---|---|---|---|
| Q8C196 | Q8C196 [870-880] | [R].LTSIDKWFLYK.[M] | 1xHomocysteineMMTS [K6] | 0,00107839 | 1576,79022 | Liver/ FAIMS/MMTS |
| Q8C196 | Q8C196 [788-803] | [R].IGSSMKSVGEVMAIGR.[T] | 1xHomocysteineMMTS [K6] | 4,49774E-05 | 1784,85258 | Liver/ FAIMS/MMTS |
| Q8C196 | Q8C196 [890-905] | [K].TLKGLNSDSVTEETLR.[K] | 1xHomocysteineMMTS [K3] | 0,000282457 | 1925,93069 | Liver/ FAIMS/MMTS |
| Q8C196 | Q8C196 [1158-1174] | [R].VSQEHPVVLTKFVEGAR.[E] | 1xHomocysteineMMTS [K11] | 0,000863044 | 2059,04633 | Liver/ FAIMS/MMTS |
| Q8C196 | Q8C196 [148-167] | [K].SLGQWLQEEKVPAIYGVDTR.[M] | 1xHomocysteineMMTS [K10] | 0,0004709 | 2452,19993 | Liver/ FAIMS/MMTS |
| Q8C1B7 | Q8C1B7 [55-79] | [K].STLMDTLFNTKFESDPATHNEPGVR.[L] | 1xHomocysteineIAA [K11] | 4,32935E-05 | 2981,37678 | Brain/ FAIMS/IAA |
| Q8C419 | Q8C419 [896-910] | [R].TSMLQKSLSVIASAK.[E] | 1xHomocysteineIAA [K6] | 0,00520029 | 1737,92375 | Brain/ FAIMS/IAA |
| Q8C854 | Q8C854 [104-112] | [K].WQAIKDLMR.[E] | 1xHomocysteineIAA [K5] | 0,000787268 | 1334,67077 | Brain/ FAIMS/IAA |
| Q8C886 | Q8C886 [566-585] | [R].EGKPSSLPPPPDEEAPIWNK.[T] | 2xHomocysteineIAA [K3; K20] | 0,0043712 | 2536,18467 | Brain/ No FAIMS/IAA |
| Q8C8R3 | Q8C8R3 [3556-3576] | [R].IENPNSLQDQSHALLKYWLER.[D] | 1xHomocysteineIAA [K16] | 4,32935E-05 | 2728,35116 | Brain/ FAIMS/IAA |
| Q8C8T8 | Q8C8T8 [103-119] | [K].GDGAALQEMTSQINQKK.[C] | 1xHomocysteineIAA [K16] | 4,32935E-05 | 1992,94773 | Brain/ FAIMS/IAA |
| Q8C8V1 | Q8C8V1 [250-261] | [K].FTTVYNLKAHMK.[G] | 1xOxidation [M11]; 1xHomocysteineIAA [K8] | 0,00033298 | 1642,80799 | Brain/ No FAIMS/IAA |
| Q8C8V1 | Q8C8V1 [250-261] | [K].FTTVYNLKAHMK.[G] | 1xOxidation [M11]; 1xHomocysteineIAA [K8] | 0,00033298 | 1642,80799 | Brain/ FAIMS/IAA |
| Q8CAQ8 | Q8CAQ8 [284-296] | [R].RKAVDEAADALLK.[A] | 1xHomocysteineIAA [K2] | 4,32935E-05 | 1573,83665 | Brain/ FAIMS/IAA |
| Q8CAQ8 | Q8CAQ8 [443-456] | [R].TFDSAVAKALEHHR.[S] | 1xHomocysteineMMTS [K8] | 0,000133296 | 1744,82577 | Liver/ No FAIMS/MMTS |
| Q8CAQ8 | Q8CAQ8 [499-515] | [K].VQEQELKYEFEQGLSEK.[L] | 1xHomocysteineMMTS [K7] | 0,00117575 | 2247,0308 | Liver/ No FAIMS/MMTS |
| Q8CAQ8 | Q8CAQ8 [123-132] | [K].LPVAQSQKTK.[G] | 1xHomocysteineMMTS [K8] | 0,00509539 | 1262,65954 | Liver/ FAIMS/MMTS |
| Q8CAQ8 | Q8CAQ8 [443-456] | [R].TFDSAVAKALEHHR.[S] | 1xHomocysteineMMTS [K8] | 0,000133296 | 1744,82577 | Liver/ FAIMS/MMTS |
| Q8CAQ8 | Q8CAQ8 [719-728] | [R].VAQDWLKEAR.[M] | 1xHomocysteineMMTS [K7] | 0,000530332 | 1378,6606 | Liver/ FAIMS/MMTS |
| Q8CAQ8 | Q8CAQ8 [499-515] | [K].VQEQELKYEFEQGLSEK.[L] | 1xHomocysteineMMTS [K7] | 0,00117575 | 2247,0308 | Liver/ FAIMS/MMTS |
| Q8CAQ8 | Q8CAQ8 [205-221] | [R].LAQQEKQEQVEMESLAK.[S] | 1xHomocysteineMMTS [K6] | 4,49774E-05 | 2152,00829 | Brain/ FAIMS/MMTS |
| Q8CAY6 | Q8CAY6 [195-210] | [K].AGHFDKEIVPVLVSSR.[K] | 1xHomocysteineIAA [K6] | 6,30912E-05 | 1928,00584 | Liver/ FAIMS/IAA |
| Q8CAY6 | Q8CAY6 [195-210] | [K].AGHFDKEIVPVLVSSR.[K] | 1xHomocysteineIAA [K6] | 6,30912E-05 | 1928,00584 | Brain/ FAIMS/IAA |
| Q8CAY6 | Q8CAY6 [273-279] | [R].MLKPLAR.[I] | 1xHomocysteineMMTS [K3] | 0,00180518 | 991,52496 | Liver/ No FAIMS/MMTS |
| Q8CAY6 | Q8CAY6 [195-210] | [K].AGHFDKEIVPVLVSSR.[K] | 1xHomocysteineMMTS [K6] | 4,49774E-05 | 1916,9721 | Liver/ FAIMS/MMTS |
| Q8CAY6 | Q8CAY6 [176-188] | [R].EAQDKVAVLSQNR.[A] | 1xHomocysteineMMTS [K5] | 0,000152766 | 1620,78324 | Liver/ FAIMS/MMTS |
| Q8CAY6 | Q8CAY6 [211-223] | [R].KGLTEVKIDEFPR.[H] | 1xHomocysteineMMTS [K7] | 0,00271031 | 1694,86042 | Liver/ FAIMS/MMTS |
| Q8CC35 | Q8CC35 [495-511] | [R].QFGEKAWAPPASSMADR.[S] | 1xHomocysteineIAA [K5] | 0,0075237 | 2022,91604 | Brain/ FAIMS/IAA |
| Q8CCT4 | Q8CCT4 [44-60] | [R].QGKLEVEGGPGEQAQQK.[G] | 1xHomocysteineIAA [K3] | 0,000243434 | 1956,94436 | Brain/ FAIMS/IAA |
| Q8CCT4; Q8R0A5 | Q8CCT4 [95-107]; Q8R0A5 [95-107] | [R].AAEKRPAEDYVPR.[K] | 1xHomocysteineIAA [K4] | 4,32935E-05 | 1675,82206 | Brain/ FAIMS/IAA |
| Q8CCT4; Q8R0A5 | Q8CCT4 [95-107]; Q8R0A5 [95-107] | [R].AAEKRPAEDYVPR.[K] | 1xHomocysteineMMTS [K4] | 0,0025135 | 1664,78832 | Brain/ FAIMS/MMTS |
| Q8CDN6 | Q8CDN6 [1-23] | [-].MVGVKPVGSDPDFQPELSGAGSR.[L] | 1xMet-loss [N-Term]; 1xHomocysteineIAA [K5] | 4,32935E-05 | 2373,15033 | Brain/ FAIMS/IAA |
| Q8CDN6 | Q8CDN6 [1-23] | [-].MVGVKPVGSDPDFQPELSGAGSR.[L] | 1xMet-loss [N-Term]; 1xHomocysteineMMTS [K5] | 0,00255629 | 2362,11659 | Liver/ FAIMS/MMTS |
| Q8CGP1 | Q8CGP1 [1-12] | [-].MPEPAKSAPAPK.[K] | 1xMet-loss [N-Term]; 1xHomocysteineMMTS [K6] | 0,000173718 | 1255,61734 | Liver/ No FAIMS/MMTS |
| Q8CGP1 | Q8CGP1 [1-12] | [-].MPEPAKSAPAPK.[K] | 1xMet-loss [N-Term]; 1xHomocysteineMMTS [K6] | 0,000173718 | 1255,61734 | Liver/ FAIMS/MMTS |
| Q8CGP1 | Q8CGP1 [1-12] | [-].MPEPAKSAPAPK.[K] | 1xMet-loss [N-Term]; 1xHomocysteineMMTS [K6] | 0,000173718 | 1255,61734 | Brain/ No FAIMS/MMTS |
| Q8CGP1 | Q8CGP1 [1-12] | [-].MPEPAKSAPAPK.[K] | 1xMet-loss [N-Term]; 1xHomocysteineMMTS [K6] | 0,000173718 | 1255,61734 | Brain/ FAIMS/MMTS |
| Q8CHC4 | Q8CHC4 [568-578] | [K].LAGIQEFQDKR.[S] | 1xHomocysteineIAA [K10] | 4,32935E-05 | 1478,74202 | Brain/ FAIMS/IAA |
| Q8CHH5 | Q8CHH5 [989-1005] | [K].GSGEPQPDLQLTKSLEK.[T] | 2xHomocysteineMMTS [K13; K17] | 0,00708682 | 2152,97454 | Liver/ No FAIMS/MMTS |
| Q8CHH5 | Q8CHH5 [989-1005] | [K].GSGEPQPDLQLTKSLEK.[T] | 2xHomocysteineMMTS [K13; K17] | 0,00708682 | 2152,97454 | Liver/ FAIMS/MMTS |
| Q8CHH9 | Q8CHH9 [179-193] | [K].VNIIPIIAKADTISK.[S] | 1xHomocysteineIAA [K9] | 4,32935E-05 | 1770,01937 | Brain/ No FAIMS/IAA |
| Q8CHH9 | Q8CHH9 [179-193] | [K].VNIIPIIAKADTISK.[S] | 1xHomocysteineIAA [K9] | 4,32935E-05 | 1770,01937 | Brain/ FAIMS/IAA |
| Q8CHH9; Q8C1B7 | Q8CHH9 [174-187]; Q8C1B7 [171-184] | [K].KLDSKVNIIPIIAK.[A] | 1xHomocysteineIAA [K5] | 4,32935E-05 | 1726,02954 | Brain/ No FAIMS/IAA |
| Q8CHH9; Q8C1B7 | Q8CHH9 [174-187]; Q8C1B7 [171-184] | [K].KLDSKVNIIPIIAK.[A] | 1xHomocysteineIAA [K5] | 4,32935E-05 | 1726,02954 | Brain/ FAIMS/IAA |
| Q8CHP8 | Q8CHP8 [167-183] | [R].AVVVGFDPHFSYMKLTK.[A] | 1xHomocysteineIAA [K14] | 0,000960518 | 2113,06091 | Brain/ FAIMS/IAA |
| Q8CHT0 | Q8CHT0 [124-138] | [R].AQVFLKAADMLSGPR.[R] | 1xHomocysteineMMTS [K6] | 0,000530332 | 1766,87503 | Liver/ FAIMS/MMTS |
| Q8CI43; Q60605 | Q8CI43 [94-112]; Q60605 [38-56] | [R].ALGQNPTNAEVLKVLGNPK.[NS] | 1xHomocysteineIAA [K13] | 4,32935E-05 | 2137,1434 | Brain/ No FAIMS/IAA |
| Q8CI43; Q60605 | Q8CI43 [94-112]; Q60605 [38-56] | [R].ALGQNPTNAEVLKVLGNPK.[NS] | 1xHomocysteineIAA [K13] | 4,32935E-05 | 2137,1434 | Brain/ FAIMS/IAA |
| Q8CI94 | Q8CI94 [508-521] | [R].IGEGFLTDLSQLKK.[L] | 1xHomocysteineIAA [K13] | 4,32935E-05 | 1722,90948 | Brain/ FAIMS/IAA |
| Q8JZK9; P54869 | Q8JZK9 [90-102]; P54869 [127-139] | [R].LEVGTETIIDKSK.[SA] | 1xHomocysteineMMTS [K11] | 0,0047171 | 1595,80191 | Liver/ FAIMS/MMTS |
| Q8JZR6; Q5DTL9 | Q8JZR6 [139-152]; Q5DTL9 [140-153] | [R].WLKFEEDVEDGGER.[W] | 1xHomocysteineIAA [K3] | 4,32935E-05 | 1882,8276 | Brain/ FAIMS/IAA |
| Q8JZS0 | Q8JZS0 [188-199] | [R].YTPKVLEEMEAR.[F] | 1xHomocysteineIAA [K4] | 0,000273428 | 1639,78184 | Brain/ FAIMS/IAA |
| Q8JZS0 | Q8JZS0 [188-199] | [R].YTPKVLEEMEAR.[F] | 1xHomocysteineMMTS [K4] | 0,000437433 | 1628,7481 | Brain/ FAIMS/MMTS |
| Q8JZW4 | Q8JZW4 [33-42] | [R].NLLDKDMFSK.[S] | 1xHomocysteineIAA [K5] | 0,000595249 | 1384,65993 | Brain/ FAIMS/IAA |
| Q8JZW4; Q9DC53; Q1RLL3 | Q8JZW4 [192-205]; Q9DC53 [186-199]; Q1RLL3 [163-176] | [K].DFFGKSDPFLVFYR.[S] | 1xHomocysteineIAA [K5] | 0,00818027 | 1911,90982 | Brain/ FAIMS/IAA |
| Q8K021 | Q8K021 [53-78] | [K].MPNVPNTQPAIMKPTEEHPAYTQITK.[E] | 1xHomocysteineIAA [K13] | 4,32935E-05 | 3110,51077 | Brain/ FAIMS/IAA |
| Q8K0E8 | Q8K0E8 [358-366] | [K].YQVSVNKYK.[G] | 1xHomocysteineIAA [K7] | 0,000273428 | 1302,65108 | Liver/ FAIMS/IAA |
| Q8K0T0 | Q8K0T0 [349-364] | [K].GSVSEDELIAAIKEAK.[G] | 1xHomocysteineIAA [K13] | 4,32935E-05 | 1833,92625 | Brain/ FAIMS/IAA |
| Q8K0U4 | Q8K0U4 [358-372] | [K].LLCKIFGEDFIEQFK.[I] | 1xCarbamidomethyl [C3]; 1xHomocysteineIAA [K4] | 4,32935E-05 | 2061,01838 | Brain/ FAIMS/IAA |

| | | | | | | |
|---|---|---|---|---|---|---|
| Q8K183 | Q8K183 [276-292] | [K].AEAGEGQKPSPAQLELR.[M] | 1xHomocysteineIAA [K8] | 4,32935E-05 | 1954,9651 | Liver/ FAIMS/IAA |
| Q8K183 | Q8K183 [138-160] | [R].DKVVPVADIITPNQFEAELLSGR.[K] | 1xHomocysteineIAA [K2] | 4,32935E-05 | 2685,39163 | Brain/ FAIMS/IAA |
| Q8K183 | Q8K183 [276-292] | [K].AEAGEGQKPSPAQLELR.[M] | 1xHomocysteineMMTS [K8] | 4,49774E-05 | 1943,93136 | Liver/ FAIMS/MMTS |
| Q8K1M6 | Q8K1M6 [61-76] | [R].RPLILQLVHVSPEDKR.[K] | 1xHomocysteineIAA [K15] | 4,32935E-05 | 2074,15899 | Brain/ FAIMS/IAA |
| Q8K1Z0 | Q8K1Z0 [162-171] | [K].LVQLGQAEKR.[K] | 1xHomocysteineIAA [K9] | 0,00033298 | 1315,71508 | Liver/ FAIMS/IAA |
| Q8K1Z0 | Q8K1Z0 [162-171] | [K].LVQLGQAEKR.[K] | 1xHomocysteineIAA [K9] | 0,00033298 | 1315,71508 | Brain/ FAIMS/IAA |
| Q8K1Z0 | Q8K1Z0 [46-76] | [R].SSEEQKQQPPHSSSQQHSETQGPEFSRPPPR.[Y] | 1xHomocysteineMMTS [K6] | 0,000302483 | 3677,64558 | Liver/ FAIMS/MMTS |
| Q8K1Z0 | Q8K1Z0 [152-170] | [R].LNQVLEEEQKLVQLGQAEK.[R] | 1xHomocysteineMMTS [K10] | 0,000133296 | 2359,19959 | Liver/ FAIMS/MMTS |
| Q8K221 | Q8K221 [217-232] | [K].TMEDTLMTVKQYEAAR.[L] | 1xHomocysteineIAA [K10] | 4,32935E-05 | 2060,94496 | Brain/ FAIMS/IAA |
| Q8K2B3 | Q8K2B3 [513-527] | [R].LNMQKSMQNHAAVFR.[V] | 1xHomocysteineIAA [K5] | 4,32935E-05 | 1948,93025 | Brain/ FAIMS/IAA |
| Q8K2T1 | Q8K2T1 [46-56] | [K].ELKQQGAEVVR.[G] | 1xHomocysteineIAA [K3] | 0,00107087 | 1430,74202 | Brain/ FAIMS/IAA |
| Q8K3C3 | Q8K3C3 [12-22] | [R].LKQNLEEQLDR.[L] | 1xHomocysteineIAA [K2] | 0,00324696 | 1559,78462 | Brain/ FAIMS/IAA |
| Q8K3J1 | Q8K3J1 [43-59] | [K].EQESEVDMKSATDNAAR.[I] | 1xHomocysteineIAA [K9] | 4,32935E-05 | 2054,87536 | Brain/ FAIMS/IAA |
| Q8K4F5 | Q8K4F5 [226-245] | [R].LNLDTLAQHLDKIMTFPQQR.[E] | 1xHomocysteineIAA [K12] | 0,000538793 | 2556,30612 | Brain/ FAIMS/IAA |
| Q8K4Z3 | Q8K4Z3 [264-280] | [K].KYQLNLPSYPDTECVYR.[L] | 1xCarbamidomethyl [C14]; 1xHomocysteineIAA [K1] | 4,32935E-05 | 2320,07366 | Brain/ FAIMS/IAA |
| Q8K596 | Q8K596 [645-658] | [R].IAEMGKPVLGENNR.[L] | 1xHomocysteineIAA [K6] | 4,32935E-05 | 1701,84109 | Brain/ FAIMS/IAA |
| Q8K596; P70414 | Q8K596 [250-258]; P70414 [253-261] | [R].LLFYKYVYK.[R] | 1xHomocysteineIAA [K5] | 0,000305143 | 1410,74901 | Brain/ FAIMS/IAA |
| Q8QZR3 | Q8QZR3 [365-377] | [R].ETLPAVLKNTAAR.[M] | 1xHomocysteineMMTS [K8] | 0,00206087 | 1546,80799 | Liver/ No FAIMS/MMTS |
| Q8QZR3 | Q8QZR3 [295-307] | [K].SKQEILAINQVFK.[M] | 1xHomocysteineMMTS [K2] | 0,000152766 | 1680,88116 | Liver/ No FAIMS/MMTS |
| Q8QZR3 | Q8QZR3 [295-307] | [K].SKQEILAINQVFK.[M] | 1xHomocysteineMMTS [K2] | 0,000152766 | 1680,88116 | Liver/ FAIMS/MMTS |
| Q8QZT1 | Q8QZT1 [241-255] | [K].GKPDVVVKEDEEYKR.[V] | 1xHomocysteineIAA [K] | 4,32935E-05 | 1964,9746 | Liver/ No FAIMS/IAA |
| Q8QZT1 | Q8QZT1 [241-255] | [K].GKPDVVVKEDEEYKR.[V] | 1xHomocysteineIAA [K] | 4,32935E-05 | 1964,9746 | Liver/ FAIMS/IAA |
| Q8QZT1 | Q8QZT1 [241-255] | [K].GKPDVVVKEDEEYKR.[V] | 1xHomocysteineIAA [K] | 4,32935E-05 | 1964,9746 | Brain/ No FAIMS/IAA |
| Q8QZT1 | Q8QZT1 [241-255] | [K].GKPDVVVKEDEEYKR.[V] | 1xHomocysteineIAA [K] | 4,32935E-05 | 1964,9746 | Brain/ FAIMS/IAA |
| Q8QZT1 | Q8QZT1 [163-178] | [R].GATPYGGVKLEDLIVK.[D] | 1xHomocysteineMMTS [K9] | 6,70238E-05 | 1822,94415 | Liver/ No FAIMS/MMTS |
| Q8QZT1 | Q8QZT1 [172-187] | [K].LEDLIVKDGLTDVYNK.[I] | 1xHomocysteineMMTS [K7] | 4,49774E-05 | 1997,99223 | Liver/ No FAIMS/MMTS |
| Q8QZT1 | Q8QZT1 [256-263] | [R].VDFSKVPK.[L] | 1xHomocysteineMMTS [K5] | 0,00677626 | 1082,5373 | Liver/ No FAIMS/MMTS |
| Q8QZT1 | Q8QZT1 [76-84] | [K].AGIPKEEVK.[E] | 1xHomocysteineMMTS [K5] | 0,00419691 | 1133,56933 | Liver/ No FAIMS/MMTS |
| Q8QZT1 | Q8QZT1 [301-308] | [R].LNVKPLAR.[I] | 1xHomocysteineMMTS [K4] | 0,0046895 | 1073,59582 | Liver/ No FAIMS/MMTS |
| Q8QZT1 | Q8QZT1 [81-102] | [K].EEVKEVYMGNVIQGGEGQAPTR.[Q] | 1xHomocysteineMMTS [K4] | 4,49774E-05 | 2554,17345 | Liver/ No FAIMS/MMTS |
| Q8QZT1 | Q8QZT1 [64-80] | [K].LGTAAIQGAIEKAGIPK.[E] | 1xHomocysteineMMTS [K12] | 4,49774E-05 | 1800,97104 | Liver/ No FAIMS/MMTS |
| Q8QZT1 | Q8QZT1 [241-255] | [K].GKPDVVVKEDEEYKR.[V] | 1xHomocysteineMMTS [K] | 4,49774E-05 | 1953,94086 | Liver/ No FAIMS/MMTS |
| Q8QZT1 | Q8QZT1 [163-178] | [R].GATPYGGVKLEDLIVK.[D] | 1xHomocysteineMMTS [K9] | 6,70238E-05 | 1822,94415 | Liver/ FAIMS/MMTS |
| Q8QZT1 | Q8QZT1 [172-187] | [K].LEDLIVKDGLTDVYNK.[I] | 1xHomocysteineMMTS [K7] | 4,49774E-05 | 1997,99223 | Liver/ FAIMS/MMTS |
| Q8QZT1 | Q8QZT1 [64-80] | [K].LGTAAIQGAIEKAGIPK.[E] | 1xHomocysteineMMTS [K12] | 4,49774E-05 | 1800,97104 | Liver/ FAIMS/MMTS |
| Q8QZT1 | Q8QZT1 [241-255] | [K].GKPDVVVKEDEEYKR.[V] | 1xHomocysteineMMTS [K] | 4,49774E-05 | 1953,94086 | Liver/ FAIMS/MMTS |
| Q8QZT1 | Q8QZT1 [363-390] | [K].MLEIDPQKVNIHGGAVSLGHPIGMSGAR.[I] | 1xHomocysteineMMTS [K8] | 0,000282457 | 3047,50458 | Liver/ FAIMS/MMTS |
| Q8QZT1 | Q8QZT1 [76-84] | [K].AGIPKEEVK.[E] | 1xHomocysteineMMTS [K5] | 0,00419691 | 1133,56933 | Liver/ FAIMS/MMTS |
| Q8QZT1 | Q8QZT1 [301-308] | [R].LNVKPLAR.[I] | 1xHomocysteineMMTS [K4] | 0,0046895 | 1073,59582 | Liver/ FAIMS/MMTS |
| Q8QZT1 | Q8QZT1 [81-102] | [K].EEVKEVYMGNVIQGGEGQAPTR.[Q] | 1xHomocysteineMMTS [K4] | 4,49774E-05 | 2554,17345 | Liver/ FAIMS/MMTS |
| Q8QZT1 | Q8QZT1 [219-227] | [R].SKEAWDAGK.[F] | 1xHomocysteineMMTS [K2] | 0,00367611 | 1154,49689 | Liver/ FAIMS/MMTS |
| Q8QZT1 | Q8QZT1 [163-178] | [R].GATPYGGVKLEDLIVK.[D] | 1xHomocysteineMMTS [K9] | 6,70238E-05 | 1822,94415 | Brain/ FAIMS/MMTS |
| Q8QZT1 | Q8QZT1 [172-187] | [K].LEDLIVKDGLTDVYNK.[I] | 1xHomocysteineMMTS [K7] | 4,49774E-05 | 1997,99223 | Brain/ FAIMS/MMTS |
| Q8R001 | Q8R001 [235-262] | [R].SDKDLETQVIQLNEQVHSLKLALEGVEK.[E] | 1xHomocysteineIAA [K20] | 4,32935E-05 | 3337,7308 | Brain/ FAIMS/IAA |
| Q8R0A5 | Q8R0A5 [46-67] | [R].QGKPEEEAKPDEQGQDEGKPEK.[Q] | 1xHomocysteineIAA [K] | 4,32935E-05 | 2627,18896 | Brain/ No FAIMS/IAA |
| Q8R0A5 | Q8R0A5 [46-67] | [R].QGKPEEEAKPDEQGQDEGKPEK.[Q] | 1xHomocysteineIAA [K] | 4,32935E-05 | 2627,18896 | Brain/ FAIMS/IAA |
| Q8R0A7 | Q8R0A7 [106-120] | [K].IFSGGEDLDQEEKAK.[F] | 1xHomocysteineIAA [K13] | 4,32935E-05 | 1839,84292 | Brain/ FAIMS/IAA |
| Q8R0F8 | Q8R0F8 [29-38] | [R].NYADHVKEMR.[S] | 1xHomocysteineIAA [K7] | 0,00163516 | 1436,64093 | Brain/ FAIMS/IAA |
| Q8R164 | Q8R164 [117-126] | [R].DAKDAVDLMK.[A] | 1xHomocysteineIAA [K3] | 0,00297155 | 1279,60208 | Brain/ FAIMS/IAA |
| Q8R1I1 | Q8R1I1 [35-54] | [R].AFDQGADAIYEHINEGKLWK.[H] | 1xHomocysteineIAA [K17] | 4,32935E-05 | 2479,17107 | Brain/ FAIMS/IAA |
| Q8R1I1 | Q8R1I1 [35-54] | [R].AFDQGADAIYEHINEGKLWK.[H] | 1xHomocysteineMMTS [K17] | 0,000456432 | 2468,13733 | Liver/ FAIMS/MMTS |
| Q8R1Q8 | Q8R1Q8 [485-501] | [K].KSGQKPVLSDVHAELDR.[I] | 1xHomocysteineIAA [K5] | 4,32935E-05 | 2053,0495 | Brain/ FAIMS/IAA |
| Q8R2E6 | Q8R2E6 [1-22] | [-].MSEILFFSPQPLFSHMMNKNSR.[L] | 1xMet-loss+Acetyl [N-Term]; 1xHomocysteineMMTS [K19] | 0,00586177 | 2715,25502 | Liver/ FAIMS/MMTS |
| Q8R317 | Q8R317 [60-74] | [R].FKSHIDQLVLIFAGK.[I] | 1xHomocysteineIAA [K2] | 4,32935E-05 | 1890,0306 | Liver/ FAIMS/IAA |
| Q8R317 | Q8R317 [60-74] | [R].FKSHIDQLVLIFAGK.[I] | 1xHomocysteineIAA [K2] | 4,32935E-05 | 1890,0306 | Brain/ FAIMS/IAA |
| Q8R326 | Q8R326 [127-142] | [R].TLAEIAKAELDGTILK.[S] | 1xHomocysteineIAA [K7] | 0,00107087 | 1860,01468 | Brain/ FAIMS/IAA |
| Q8R366 | Q8R366 [376-385] | [R].HIAMEKVASR.[T] | 1xHomocysteineIAA [K6] | 4,32935E-05 | 1315,66094 | Brain/ No FAIMS/IAA |
| Q8R366 | Q8R366 [376-385] | [R].HIAMEKVASR.[T] | 1xHomocysteineIAA [K6] | 4,32935E-05 | 1315,66094 | Brain/ FAIMS/IAA |
| Q8R366 | Q8R366 [376-385] | [R].HIAMEKVASR.[T] | 1xHomocysteineMMTS [K6] | 0,00115861 | 1304,62719 | Brain/ FAIMS/MMTS |
| Q8R3V5 | Q8R3V5 [19-32] | [R].AVQFTEEKFGQAEK.[T] | 1xHomocysteineIAA [K8] | 9,61721E-05 | 1785,84761 | Brain/ FAIMS/IAA |
| Q8R404 | Q8R404 [76-82] | [K].IKFPNFR.[D] | 1xHomocysteineIAA [K2] | 0,000668843 | 1095,5768 | Brain/ No FAIMS/IAA |
| Q8R404 | Q8R404 [76-82] | [K].IKFPNFR.[D] | 1xHomocysteineIAA [K2] | 0,000668843 | 1095,5768 | Brain/ FAIMS/IAA |
| Q8R570 | Q8R570 [271-285] | [R].FIGKPDVAYQLISAK.[M] | 1xHomocysteineIAA [K4] | 0,00299711 | 1823,97242 | Brain/ FAIMS/IAA |
| Q8R5H6 | Q8R5H6 [183-195] | [K].NLDRPHEPEKVPR.[A] | 1xHomocysteineIAA [K10] | 0,00012645 | 1760,88606 | Brain/ FAIMS/IAA |
| Q8R5I6; P10649; P15626; Q80W21 | Q8R5I6 [53-78]; P10649 [53-78]; P15626 [53-78]; Q80W21 [53-78] | [K].LGLDFPNLPYLIDGSHKITQSNAILR.[Y] | 1xHomocysteineIAA [K17] | 4,32935E-05 | 3069,619 | Liver/ FAIMS/IAA |
| Q8R5I6; P10649; P15626; Q80W21 | Q8R5I6 [53-78]; P10649 [53-78]; P15626 [53-78]; Q80W21 [53-78] | [K].LGLDFPNLPYLIDGSHKITQSNAILR.[Y] | 1xHomocysteineIAA [K17] | 4,32935E-05 | 3069,619 | Brain/ FAIMS/IAA |
| Q8R5I6; P10649; P15626; Q80W21 | Q8R5I6 [51-69]; P10649 [51-69]; P15626 [51-69]; Q80W21 [51-69] | [K].FKLGLDFPNLPYLIDGSHK.[I] | 1xHomocysteineMMTS [K2] | 4,49774E-05 | 2337,17701 | Liver/ No FAIMS/MMTS |

| | | | | | | |
|---|---|---|---|---|---|---|
| Q8R5I6; P10649; P15626; Q80W21 | Q8R5I6 [51-69]; P10649 [51-69]; P15626 [51-69]; Q80W21 [51-69] | [K].FKLGLDFPNLPYLIDGSHK.[I] | 1xHomocysteineMMTS [K2] | 4,49774E-05 | 2337,17701 | Liver/ FAIMS/MMTS |
| Q8R5I6; P10649; P15626; Q80W21 | Q8R5I6 [53-78]; P10649 [53-78]; P15626 [53-78]; Q80W21 [53-78] | [K].LGLDFPNLPYLIDGSHKITQSNAILR.[Y] | 1xHomocysteineMMTS [K17] | 0,000173718 | 3058,58526 | Liver/ FAIMS/MMTS |
| Q8R5M8 | Q8R5M8 [191-211] | [K].GKSEVEEWSDMYTVTSQLMLK.[V] | 1xHomocysteineIAA [K2] | 0,000210768 | 2635,20884 | Brain/ FAIMS/IAA |
| Q8VBW5 | Q8VBW5 [717-728] | [K].KKTGNMSSESTK.[T] | 1xHomocysteineMMTS [K2] | 0,00531384 | 1460,6542 | Liver/ FAIMS/MMTS |
| Q8VBY2 | Q8VBY2 [130-143] | [K].LQSEIGKGAYGVVR.[L] | 1xHomocysteineIAA [K7] | 0,00608856 | 1650,8632 | Brain/ FAIMS/IAA |
| Q8VC26 | Q8VC26 [148-155] | [R].KLLLLSHK.[A] | 1xHomocysteineMMTS [K1] | 0,000929608 | 1114,64752 | Brain/ No FAIMS/MMTS |
| Q8VC30 | Q8VC30 [523-535] | [K].AVKSAEAAAEATK.[N] | 1xHomocysteineMMTS [K3] | 6,70238E-05 | 1409,67631 | Liver/ No FAIMS/MMTS |
| Q8VC30 | Q8VC30 [485-493] | [K].YGKAAPGDR.[T] | 1xHomocysteineMMTS [K3] | 0,008514 | 1097,48666 | Liver/ No FAIMS/MMTS |
| Q8VC30 | Q8VC30 [511-525] | [K].SPGASLLPVLTKAVK.[S] | 1xHomocysteineMMTS [K12] | 0,000133296 | 1643,9223 | Liver/ No FAIMS/MMTS |
| Q8VC30 | Q8VC30 [485-493] | [K].YGKAAPGDR.[T] | 1xHomocysteineMMTS [K3] | 0,008514 | 1097,48666 | Liver/ FAIMS/MMTS |
| Q8VC30 | Q8VC30 [523-535] | [K].AVKSAEAAAEATK.[N] | 1xHomocysteineMMTS [K3] | 6,70238E-05 | 1409,67631 | Liver/ FAIMS/MMTS |
| Q8VC30 | Q8VC30 [511-525] | [K].SPGASLLPVLTKAVK.[S] | 1xHomocysteineMMTS [K12] | 0,000133296 | 1643,9223 | Liver/ FAIMS/MMTS |
| Q8VC30 | Q8VC30 [526-543] | [K].SAEAAAEATKNMEAGAGR.[A] | 1xHomocysteineMMTS [K10] | 4,49774E-05 | 1897,82009 | Liver/ FAIMS/MMTS |
| Q8VCC2 | Q8VCC2 [287-299] | [R].QKTEDELLEVSLK.[M] | 1xHomocysteineMMTS [K2] | 0,000231655 | 1694,83393 | Liver/ FAIMS/MMTS |
| Q8VCC2; Q91WU0 | Q8VCC2 [481-496]; Q91WU0 [481-496] | [KR].EGASEEEINLSKMVMK.[F] | 1xHomocysteineMMTS [K12] | 0,00242157 | 1957,87377 | Liver/ FAIMS/MMTS |
| Q8VCH0 | Q8VCH0 [78-102] | [K].LKPEQLGDISVGNVLQPGAGAIMAR.[I] | 1xHomocysteineMMTS [K2] | 4,49774E-05 | 2697,38847 | Liver/ FAIMS/MMTS |
| Q8VCH0; Q921H8 | Q8VCH0 [291-305]; Q921H8 [291-305] | [R].SKAEELGLPILGVLR.[S] | 1xHomocysteineMMTS [K2] | 0,00125756 | 1757,96522 | Liver/ FAIMS/MMTS |
| Q8VCT4 | Q8VCT4 [300-312] | [K].LNLFKLDLLGNPK.[E] | 1xHomocysteineMMTS [K5] | 8,93346E-05 | 1647,89608 | Liver/ No FAIMS/MMTS |
| Q8VCT4 | Q8VCT4 [330-351] | [K].APEEILAEKSFSTVPYIVGINK.[Q] | 1xHomocysteineMMTS [K9] | 0,00664737 | 2568,30881 | Liver/ FAIMS/MMTS |
| Q8VCT4 | Q8VCT4 [375-388] | [K].TANSLLWKSYPTLK.[I] | 1xHomocysteineMMTS [K8] | 0,000192903 | 1784,90737 | Liver/ FAIMS/MMTS |
| Q8VCT4 | Q8VCT4 [383-400] | [K].SYPTLKISENMIPVVAEK.[Y] | 1xHomocysteineMMTS [K6] | 0,00151645 | 2182,09564 | Liver/ FAIMS/MMTS |
| Q8VCT4 | Q8VCT4 [300-312] | [K].LNLFKLDLLGNPK.[E] | 1xHomocysteineMMTS [K5] | 8,93346E-05 | 1647,89608 | Liver/ FAIMS/MMTS |
| Q8VCT4 | Q8VCT4 [522-536] | [K].EGYLKIGASTQAAQR.[L] | 1xHomocysteineMMTS [K5] | 0,00531384 | 1755,85165 | Liver/ FAIMS/MMTS |
| Q8VCT4 | Q8VCT4 [537-549] | [R].LKDKEVSFWAELR.[A] | 1xHomocysteineMMTS [K4] | 0,00386874 | 1783,88697 | Liver/ FAIMS/MMTS |
| Q8VCT4 | Q8VCT4 [371-382] | [K].LDQKTANSLLWK.[S] | 1xHomocysteineMMTS [K4] | 0,000724331 | 1579,7971 | Liver/ FAIMS/MMTS |
| Q8VCT4 | Q8VCT4 [481-496] | [K].DGASEEETNLSKMVMK.[F] | 1xHomocysteineMMTS [K12] | 0,0039709 | 1931,82173 | Liver/ FAIMS/MMTS |
| Q8VCT4 | Q8VCT4 [389-410] | [K].ISENMIPVVAEKYLGGTDDLTK.[K] | 1xHomocysteineMMTS [K12] | 4,49774E-05 | 2556,23941 | Liver/ FAIMS/MMTS |
| Q8VCU1 | Q8VCU1 [355-367] | [K].SLNILDKLEHLSR.[E] | 1xHomocysteineMMTS [K7] | 4,49774E-05 | 1700,88222 | Liver/ No FAIMS/MMTS |
| Q8VCU1 | Q8VCU1 [355-367] | [K].SLNILDKLEHLSR.[E] | 1xHomocysteineMMTS [K7] | 4,49774E-05 | 1700,88222 | Liver/ FAIMS/MMTS |
| Q8VCW8 | Q8VCW8 [532-547] | [K].HPQVQEAQVVGVKDER.[M] | 1xHomocysteineIAA [K13] | 8,35381E-05 | 1992,99198 | Liver/ FAIMS/IAA |
| Q8VCW8 | Q8VCW8 [479-490] | [K].TFETVGQDKWYR.[T] | 1xHomocysteineMMTS [K9] | 0,000211896 | 1692,75087 | Liver/ FAIMS/MMTS |
| Q8VCW8 | Q8VCW8 [200-209] | [K].AQPGALKSER.[L] | 1xHomocysteineMMTS [K7] | 0,00567876 | 1219,59219 | Liver/ FAIMS/MMTS |
| Q8VCW8 | Q8VCW8 [100-111] | [R].LNFAQLKEEVDK.[A] | 1xHomocysteineMMTS [K7] | 4,49774E-05 | 1596,77603 | Liver/ FAIMS/MMTS |
| Q8VCW8 | Q8VCW8 [585-602] | [R].YIVFVEGYPLTISGKIQK.[F] | 1xHomocysteineMMTS [K15] | 0,00797002 | 2218,16504 | Liver/ FAIMS/MMTS |
| Q8VD04 | Q8VD04 [241-253] | [R].LQTEKETLFNDSR.[N] | 1xHomocysteineIAA [K5] | 0,00251002 | 1754,83777 | Brain/ FAIMS/IAA |
| Q8VD37 | Q8VD37 [636-653] | [K].KVSEQKPQATYYNVDMLK.[Y] | 1xHomocysteineIAA [K1] | 4,32935E-05 | 2316,13626 | Brain/ FAIMS/IAA |
| Q8VDN2 | Q8VDN2 [178-198] | [K].MSINAEDVVVGDLVEVKGGDR.[I] | 1xHomocysteineIAA [K17] | 4,32935E-05 | 2376,15337 | Brain/ FAIMS/IAA |
| Q8VDN2 | Q8VDN2 [178-198] | [K].MSINAEDVVVGDLVEVKGGDR.[I] | 1xHomocysteineMMTS [K17] | 0,000282457 | 2365,11963 | Brain/ FAIMS/MMTS |
| Q8VDN2; Q64436 | Q8VDN2 [503-513]; Q64436 [512-522] | [KR].HLLVMKGAPER.[IV] | 1xHomocysteineIAA [K6] | 4,32935E-05 | 1424,75009 | Brain/ No FAIMS/IAA |
| Q8VDN2; Q64436 | Q8VDN2 [503-513]; Q64436 [512-522] | [KR].HLLVMKGAPER.[IV] | 1xHomocysteineIAA [K6] | 4,32935E-05 | 1424,75009 | Brain/ FAIMS/IAA |
| Q8VDN2; Q6PIC6 | Q8VDN2 [841-848]; Q6PIC6 [831-838] | [KR].TDKLVNER.[L] | 1xHomocysteineIAA [K3] | 0,0021859 | 1148,57283 | Brain/ No FAIMS/IAA |
| Q8VDN2; Q6PIC6 | Q8VDN2 [841-848]; Q6PIC6 [831-838] | [KR].TDKLVNER.[L] | 1xHomocysteineIAA [K3] | 0,0021859 | 1148,57283 | Brain/ FAIMS/IAA |
| Q8VDN2; Q6PIC6 | Q8VDN2 [841-848]; Q6PIC6 [831-838] | [KR].TDKLVNER.[L] | 1xHomocysteineMMTS [K3] | 0,000400718 | 1137,53909 | Brain/ No FAIMS/MMTS |
| Q8VDN2; Q6PIC6 | Q8VDN2 [841-848]; Q6PIC6 [831-838] | [KR].TDKLVNER.[L] | 1xHomocysteineMMTS [K3] | 0,000400718 | 1137,53909 | Brain/ FAIMS/MMTS |
| Q8VDN2; Q6PIC6; Q6PIE5 | Q8VDN2 [942-952]; Q6PIC6 [932-942]; Q6PIE5 [939-949] | [R].NSVFQQGMKNK.[I] | 1xHomocysteineIAA [K9] | 4,32935E-05 | 1454,68788 | Brain/ No FAIMS/IAA |
| Q8VDN2; Q6PIC6; Q6PIE5 | Q8VDN2 [942-952]; Q6PIC6 [932-942]; Q6PIE5 [939-949] | [R].NSVFQQGMKNK.[I] | 1xHomocysteineIAA [K9] | 4,32935E-05 | 1454,68788 | Brain/ FAIMS/IAA |
| Q8VDN2; Q6PIC6; Q6PIE5 | Q8VDN2 [626-647]; Q6PIC6 [616-637]; Q6PIE5 [623-644] | [K].AIAKGVGIISEGNETVEDIAAR.[L] | 1xHomocysteineMMTS [K4] | 4,49774E-05 | 2376,18976 | Brain/ No FAIMS/MMTS |
| Q8VDN2; Q6PIC6; Q6PIE5 | Q8VDN2 [626-647]; Q6PIC6 [616-637]; Q6PIE5 [623-644] | [K].AIAKGVGIISEGNETVEDIAAR.[L] | 1xHomocysteineMMTS [K4] | 4,49774E-05 | 2376,18976 | Brain/ FAIMS/MMTS |
| Q8VDN2; Q6PIC6; Q6PIE5 | Q8VDN2 [708-727]; Q6PIC6 [698-717]; Q6PIE5 [705-724] | [R].QGAIVAVTGDGVNDSPALKK.[A] | 1xHomocysteineMMTS [K19] | 4,49774E-05 | 2103,05729 | Brain/ FAIMS/MMTS |
| Q8VDN2; Q9WV27; Q64436; Q6PIC6; Q6PIE5 | Q8VDN2 [767-774]; Q9WV27 [776-783]; Q64436 [776-783]; Q6PIC6 [757-764]; Q6PIE5 [764-771] | [R].LIFDNLKK.[ST] | 1xHomocysteineMMTS [K7] | 0,00869467 | 1153,6108 | Brain/ FAIMS/MMTS |

| | | | | | | |
|---|---|---|---|---|---|---|
| Q8VDN2; | Q8VDN2 [767-774]; | | | | | |
| Q9WV27; | Q9WV27 [776-783]; | | | | | |
| Q64436; | Q64436 [776-783]; | | | | | |
| Q9Z1W8; | Q9Z1W8 [778-785]; | | | | | |
| Q6PIC6; | Q6PIC6 [757-764]; | | | | | |
| Q6PIE5 | Q6PIE5 [764-771] | [R].LIFDNLKK.[ST] | 1xHomocysteineIAA [K] | 9,61721E-05 | 1164,64454 | Brain/ No FAIMS/IAA |
| Q8VDN2; | Q8VDN2 [767-774]; | | | | | |
| Q9WV27; | Q9WV27 [776-783]; | | | | | |
| Q64436; | Q64436 [776-783]; | | | | | |
| Q9Z1W8; | Q9Z1W8 [778-785]; | | | | | |
| Q6PIC6; | Q6PIC6 [757-764]; | | | | | |
| Q6PIE5 | Q6PIE5 [764-771] | [R].LIFDNLKK.[ST] | 1xHomocysteineIAA [K] | 9,61721E-05 | 1164,64454 | Brain/ FAIMS/IAA |
| Q8VDN2; | Q8VDN2 [157-173]; | | | | | |
| Q9WV27; | Q9WV27 [167-183]; | | | | | |
| Q6PIC6 | Q6PIC6 [147-163] | [K].IMESFKNMVPQQALVIR.[NDE] | 1xHomocysteineIAA [K6] | 4,32935E-05 | 2178,1232 | Brain/ FAIMS/IAA |
| Q8VDN2; | Q8VDN2 [157-173]; | | | | | |
| Q9WV27; | Q9WV27 [167-183]; | | | | | |
| Q6PIC6 | Q6PIC6 [147-163] | [K].IMESFKNMVPQQALVIR.[NDE] | 1xHomocysteineMMTS [K6] | 0,000361246 | 2167,08945 | Brain/ FAIMS/MMTS |
| Q8VDN2; | Q8VDN2 [354-377]; | | | | | |
| Q9WV27; | Q9WV27 [364-387]; | | | | | |
| Q6PIC6; | Q6PIC6 [344-367]; | | 2xCarbamidomethyl [C3; C21]; | | | |
| Q6PIE5 | Q6PIE5 [352-375] | [R].KNCLVKNLEAVETLGSTSTICSDK.[T] | 1xHomocysteineIAA [K6] | 4,32935E-05 | 2841,37909 | Brain/ FAIMS/IAA |
| Q8VDN2; | Q8VDN2 [608-625]; | | | | | |
| Q9WV27; | Q9WV27 [617-634]; | | | | | |
| Q6PIC6; | Q6PIC6 [598-615]; | | | | | |
| Q6PIE5 | Q6PIE5 [605-622] | [R].SAGIKVIMVTGDHPITAK.[A] | 1xHomocysteineMMTS [K5] | 4,49774E-05 | 2001,03298 | Brain/ FAIMS/MMTS |
| Q8VDN2; | Q8VDN2 [613-629]; | | | | | |
| Q9WV27; | Q9WV27 [622-638]; | | | | | |
| Q6PIC6; | Q6PIC6 [603-619]; | | | | | |
| Q6PIE5 | Q6PIE5 [610-626] | [K].VIMVTGDHPITAKAIAK.[GS] | 1xHomocysteineMMTS [K13] | 0,000282457 | 1928,01661 | Brain/ FAIMS/MMTS |
| Q8VDN2; | Q8VDN2 [608-625]; | | | | | |
| Q9WV27; | Q9WV27 [617-634]; | | | | | |
| Q9Z1W8; | Q9Z1W8 [619-636]; | | | | | |
| Q6PIC6; | Q6PIC6 [598-615]; | | | | | |
| Q6PIE5 | Q6PIE5 [605-622] | [R].SAGIKVIMVTGDHPITAK.[A] | 1xHomocysteineIAA [K5] | 4,32935E-05 | 2012,06673 | Brain/ No FAIMS/IAA |
| Q8VDN2; | Q8VDN2 [608-625]; | | | | | |
| Q9WV27; | Q9WV27 [617-634]; | | | | | |
| Q9Z1W8; | Q9Z1W8 [619-636]; | | | | | |
| Q6PIC6; | Q6PIC6 [598-615]; | | | | | |
| Q6PIE5 | Q6PIE5 [605-622] | [R].SAGIKVIMVTGDHPITAK.[A] | 1xHomocysteineIAA [K5] | 4,32935E-05 | 2012,06673 | Brain/ FAIMS/IAA |
| Q8VDQ8 | Q8VDQ8 [126-136] | [K].KHPEPFFALAK.[E] | 1xHomocysteineIAA [K1] | 4,32935E-05 | 1458,75622 | Brain/ No FAIMS/IAA |
| Q8VDQ8 | Q8VDQ8 [126-136] | [K].KHPEPFFALAK.[E] | 1xHomocysteineIAA [K1] | 4,32935E-05 | 1458,75622 | Brain/ FAIMS/IAA |
| Q8VDQ8 | Q8VDQ8 [43-57] | [R].NLFTQTLGLGSQKER.[L] | 1xHomocysteineMMTS [K13] | 4,49774E-05 | 1854,92006 | Brain/ No FAIMS/MMTS |
| | | | 1xCarbamidomethyl [C10]; | | | |
| Q8VE33 | Q8VE33 [66-78] | [R].LVIAEKGLACEER.[D] | 1xHomocysteineIAA [K6] | 0,000273428 | 1661,83494 | Brain/ FAIMS/IAA |
| Q8VEH3 | Q8VEH3 [133-146] | [R].DLAGALDEKELIEK.[M] | 1xHomocysteineIAA [K9] | 4,32935E-05 | 1717,86768 | Brain/ FAIMS/IAA |
| Q8VEK0 | Q8VEK0 [9-26] | [K].DEVDGGPAGPPGGAAKTR.[R] | 1xHomocysteineIAA [K16] | 0,00459966 | 1825,84974 | Brain/ FAIMS/IAA |
| Q8VEM8 | Q8VEM8 [205-221] | [K].EEGLNAFYKGVAPLWMR.[Q] | 1xHomocysteineIAA [K9] | 4,32935E-05 | 2155,04633 | Brain/ FAIMS/IAA |
| Q8VEM8 | Q8VEM8 [205-221] | [K].EEGLNAFYKGVAPLWMR.[Q] | 1xHomocysteineMMTS [K9] | 0,00750949 | 2144,01258 | Brain/ FAIMS/MMTS |
| Q8VHW2 | Q8VHW2 [76-88] | [K].KDPGGLTHSGLWR.[I] | 1xHomocysteineIAA [K1] | 0,00507124 | 1597,79037 | Brain/ FAIMS/IAA |
| Q8VIJ6 | Q8VIJ6 [335-350] | [R].ALAEIAKAELDDTPMR.[G] | 1xHomocysteineIAA [K7] | 4,32935E-05 | 1917,94086 | Brain/ FAIMS/IAA |
| Q91V12 | Q91V12 [170-184] | [K].NVDKVLEVPPIVYLR.[Q] | 1xHomocysteineIAA [K4] | 9,61721E-05 | 1928,06738 | Brain/ FAIMS/IAA |
| Q91V14 | Q91V14 [896-906] | [K].KDLTTFLYHLR.[I] | 1xHomocysteineIAA [K1] | 4,32935E-05 | 1580,82536 | Brain/ No FAIMS/IAA |
| Q91V14 | Q91V14 [896-906] | [K].KDLTTFLYHLR.[I] | 1xHomocysteineIAA [K1] | 4,32935E-05 | 1580,82536 | Brain/ FAIMS/IAA |
| Q91V41 | Q91V41 [52-61] | [R].IIEVSGQKIK.[L] | 1xHomocysteineIAA [K8] | 4,32935E-05 | 1288,72933 | Brain/ No FAIMS/IAA |
| Q91V41 | Q91V41 [52-61] | [R].IIEVSGQKIK.[L] | 1xHomocysteineIAA [K8] | 4,32935E-05 | 1288,72933 | Brain/ FAIMS/IAA |
| Q91V61 | Q91V61 [72-91] | [K].YVYDSAFHPDTGEKVVLIGR.[M] | 1xHomocysteineIAA [K14] | 0,000114608 | 2440,19656 | Brain/ FAIMS/IAA |
| Q91V76 | Q91V76 [190-211] | [R].QTLEEHYGDKPVGMGGTFIVQK.[G] | 1xHomocysteineIAA [K10] | 0,00192453 | 2608,25342 | Liver/ FAIMS/IAA |
| Q91V76 | Q91V76 [59-75] | [R].IAEVGGVPYLLPLVNKK.[K] | 1xHomocysteineMMTS [K16] | 4,49774E-05 | 1973,09624 | Liver/ No FAIMS/MMTS |
| Q91V76 | Q91V76 [59-75] | [R].IAEVGGVPYLLPLVNKK.[K] | 1xHomocysteineMMTS [K16] | 4,49774E-05 | 1973,09624 | Liver/ FAIMS/MMTS |
| Q91V76 | Q91V76 [77-88] | [K].VYDLNEIAKVIK.[L] | 1xHomocysteineMMTS [K9] | 0,0004709 | 1567,82225 | Liver/ FAIMS/MMTS |
| Q91V76 | Q91V76 [170-177] | [K].VIEVQAKR.[R] | 1xHomocysteineMMTS [K7] | 0,00154915 | 1105,58565 | Liver/ FAIMS/MMTS |
| Q91V76 | Q91V76 [303-315] | [R].IDQPKETHAFGRD.[-] | 1xHomocysteineMMTS [K5] | 0,000599431 | 1676,75194 | Liver/ FAIMS/MMTS |
| Q91V92 | Q91V92 [977-985] | [R].VQILKDFVK.[Q] | 1xHomocysteineIAA [K5] | 0,00936942 | 1263,71296 | Brain/ FAIMS/IAA |
| Q91VD9 | Q91VD9 [309-325] | [R].NEKGLLTYTSWEDALSR.[V] | 1xHomocysteineIAA [K3] | 0,000114608 | 2157,02809 | Brain/ FAIMS/IAA |
| Q91VD9 | Q91VD9 [309-325] | [R].NEKGLLTYTSWEDALSR.[V] | 1xHomocysteineMMTS [K3] | 0,00140305 | 2145,99435 | Liver/ FAIMS/MMTS |
| Q91VN4 | Q91VN4 [127-141] | [K].KFQQEQLAVQDEMVR.[V] | 1xHomocysteineIAA [K1] | 4,32935E-05 | 2022,97356 | Brain/ FAIMS/IAA |
| Q91VR2 | Q91VR2 [90-100] | [K].KHLIIGVSSDR.[G] | 1xHomocysteineIAA [K1] | 4,32935E-05 | 1398,75219 | Brain/ No FAIMS/IAA |
| Q91VR2 | Q91VR2 [90-100] | [K].KHLIIGVSSDR.[G] | 1xHomocysteineIAA [K1] | 4,32935E-05 | 1398,75219 | Brain/ FAIMS/IAA |
| Q91VR2 | Q91VR2 [90-100] | [K].KHLIIGVSSDR.[G] | 1xHomocysteineMMTS [K1] | 0,00576292 | 1387,71845 | Liver/ FAIMS/MMTS |
| Q91VR2 | Q91VR2 [263-277] | [K].NASDMIDKLTLTFNR.[T] | 1xHomocysteineMMTS [K8] | 0,000706375 | 1901,8918 | Brain/ FAIMS/MMTS |
| Q91VR2 | Q91VR2 [90-100] | [K].KHLIIGVSSDR.[G] | 1xHomocysteineMMTS [K1] | 0,00576292 | 1387,71845 | Brain/ FAIMS/MMTS |
| Q91VR5 | Q91VR5 [163-173] | [K].FGFGFGGTGKK.[S] | 1xHomocysteineIAA [K11] | 0,00943155 | 1276,6143 | Brain/ FAIMS/IAA |
| Q91VR7 | Q91VR7 [50-65] | [K].TKFLVPDHVNMSELVK.[I] | 1xHomocysteineIAA [K2] | 4,32935E-05 | 2031,04018 | Brain/ FAIMS/IAA |
| Q91VR7; | Q91VR7 [43-51]; | | | | | |
| Q9CQV6 | Q9CQV6 [43-51] | [K].QLPVLDKTK.[F] | 1xHomocysteineIAA [K7] | 0,000305143 | 1215,67657 | Brain/ FAIMS/IAA |
| Q91VR8 | Q91VR8 [47-59] | [R].LATLNEKLTALER.[R] | 1xHomocysteineIAA [K7] | 0,00012645 | 1645,89417 | Brain/ FAIMS/IAA |
| | | | 1xCarbamidomethyl [C12]; | | | |
| Q91VS7 | Q91VS7 [39-55] | [R].ITNKVFANPEDCAGFGK.[G] | 1xHomocysteineIAA [K4] | 0,00187392 | 2041,94701 | Liver/ FAIMS/IAA |
| | | | 1xMethylthio [C12]; | | | |
| Q91VS7 | Q91VS7 [39-55] | [R].ITNKVFANPEDCAGFGK.[G] | 1xHomocysteineMMTS [K4] | 4,49774E-05 | 2019,87952 | Liver/ FAIMS/MMTS |
| Q91VZ6 | Q91VZ6 [13-26] | [K].LNEQHQLILSKLLR.[E] | 1xHomocysteineIAA [K11] | 4,32935E-05 | 1879,05821 | Brain/ FAIMS/IAA |

| | | | | | | |
|---|---|---|---|---|---|---|
| Q91WD5 | Q91WD5 [123-138] | [K].LIEYKTYLQALPYFDR.[L] | 1xHomocysteineIAA [K5] | 4,32935E-05 | 2207,12054 | Brain/ FAIMS/IAA |
| Q91WG0; Q8BK48; Q8QZR3 | Q91WG0 [537-547]; Q8BK48 [535-545]; Q8QZR3 [534-544] | [R].LQFWTKTLPQK.[I] | 1xHomocysteineMMTS [K6] | 0,000530332 | 1552,80145 | Liver/ FAIMS/MMTS |
| Q91WJ8 | Q91WJ8 [391-405] | [K].GGETIKSISQQSGAR.[I] | 1xHomocysteineIAA [K6] | 0,000414013 | 1692,83336 | Brain/ FAIMS/IAA |
| Q91WS0 | Q91WS0 [90-105] | [K].HNEETGDNVGPLIIKK.[K] | 1xHomocysteineIAA [K15] | 4,32935E-05 | 1937,97494 | Brain/ No FAIMS/IAA |
| Q91WS0 | Q91WS0 [90-105] | [K].HNEETGDNVGPLIIKK.[K] | 1xHomocysteineIAA [K15] | 4,32935E-05 | 1937,97494 | Brain/ FAIMS/IAA |
| Q91WS0 | Q91WS0 [43-55] | [K].AMVNLQIQKDNPK.[V] | 1xHomocysteineMMTS [K9] | 0,00575155 | 1661,81718 | Liver/ FAIMS/MMTS |
| Q91WS0 | Q91WS0 [52-68] | [K].DNPKVVHAFDMEDLGDK.[A] | 1xHomocysteineMMTS [K4] | 8,93346E-05 | 2092,91366 | Liver/ FAIMS/MMTS |
| Q91WS0 | Q91WS0 [52-68] | [K].DNPKVVHAFDMEDLGDK.[A] | 1xHomocysteineMMTS [K4] | 8,93346E-05 | 2092,91366 | Brain/ FAIMS/MMTS |
| Q91WU0 | Q91WU0 [300-312] | [K].LNLYKLSLQGDTK.[N] | 1xHomocysteineIAA [K5] | 0,00012645 | 1666,88327 | Liver/ FAIMS/IAA |
| Q91WU0 | Q91WU0 [537-551] | [R].LKEEEVTFWTQSLAK.[K] | 1xHomocysteineMMTS [K2] | 4,49774E-05 | 1971,95545 | Liver/ No FAIMS/MMTS |
| Q91WU0 | Q91WU0 [313-332] | [K].NSDQFVTSVLDGVVLPKDPK.[E] | 1xHomocysteineMMTS [K17] | 4,49774E-05 | 2321,15158 | Liver/ No FAIMS/MMTS |
| Q91WU0 | Q91WU0 [300-312] | [K].LNLYKLSLQGDTK.[N] | 1xHomocysteineMMTS [K5] | 0,000266738 | 1655,84952 | Liver/ FAIMS/MMTS |
| Q91WU0 | Q91WU0 [518-536] | [K].YDQKEGYLHIGGTTQQAQR.[L] | 1xHomocysteineMMTS [K4] | 0,000530332 | 2356,08087 | Liver/ FAIMS/MMTS |
| Q91WU0 | Q91WU0 [537-551] | [R].LKEEEVTFWTQSLAK.[K] | 1xHomocysteineMMTS [K2] | 4,49774E-05 | 1971,95545 | Liver/ FAIMS/MMTS |
| Q91WU0 | Q91WU0 [313-332] | [K].NSDQFVTSVLDGVVLPKDPK.[E] | 1xHomocysteineMMTS [K17] | 4,49774E-05 | 2321,15158 | Liver/ FAIMS/MMTS |
| Q91X72 | Q91X72 [166-174] | [R].KWFWDFATR.[T] | 1xHomocysteineIAA [K1] | 0,00223964 | 1430,6674 | Liver/ FAIMS/IAA |
| Q91X72 | Q91X72 [385-409] | [K].SGAQATWTEVSWPHEKVDGALCLDK.[S] | 1xCarbamidomethyl [C22]; 1xHomocysteineIAA [K16] | 4,32935E-05 | 2959,3713 | Brain/ FAIMS/IAA |
| Q91X72 | Q91X72 [103-114] | [R].GPDSVFLIKEDK.[V] | 1xHomocysteineMMTS [K9] | 0,000530332 | 1510,72801 | Liver/ No FAIMS/MMTS |
| Q91X72 | Q91X72 [103-114] | [R].GPDSVFLIKEDK.[V] | 1xHomocysteineMMTS [K9] | 0,000530332 | 1510,72801 | Liver/ FAIMS/MMTS |
| Q91X72 | Q91X72 [90-102] | [R].WKNPITSVDAAFR.[G] | 1xHomocysteineMMTS [K2] | 0,00842005 | 1667,80324 | Liver/ FAIMS/MMTS |
| Q91X83 | Q91X83 [230-250] | [K].AVVPAKYLDEDTVYHLQPSGR.[F] | 1xHomocysteineIAA [K6] | 4,32935E-05 | 2532,25513 | Liver/ FAIMS/IAA |
| Q91X83 | Q91X83 [230-250] | [K].AVVPAKYLDEDTVYHLQPSGR.[F] | 1xHomocysteineMMTS [K6] | 4,49774E-05 | 2521,22139 | Liver/ FAIMS/MMTS |
| Q91X83 | Q91X83 [364-374] | [R].DLDLKKPIYQK.[T] | 1xHomocysteineMMTS [K5] | 0,00191513 | 1523,79603 | Liver/ FAIMS/MMTS |
| Q91X83 | Q91X83 [225-235] | [K].EQVIKAVVPAK.[Y] | 1xHomocysteineMMTS [K5] | 0,001242 | 1344,73779 | Liver/ FAIMS/MMTS |
| Q91X83; Q3THS6 | Q91X83 [301-308]; Q3THS6 [300-307] | [R].WVAKSLVK.[AG] | 1xHomocysteineIAA [K4] | 0,00222614 | 1104,62341 | Liver/ No FAIMS/IAA |
| Q91X83; Q3THS6 | Q91X83 [301-308]; Q3THS6 [300-307] | [R].WVAKSLVK.[AG] | 1xHomocysteineIAA [K4] | 0,00222614 | 1104,62341 | Liver/ FAIMS/IAA |
| Q91X97 | Q91X97 [161-171] | [R].DGKLSLEEFIR.[G] | 1xHomocysteineMMTS [K3] | 0,000322194 | 1469,7127 | Brain/ FAIMS/MMTS |
| Q91X97; P84075; P62748 | Q91X97 [131-148]; P84075 [131-148]; P62748 [131-148] | [K].MVSSVMKMPEDESTPEKR.[T] | 1xHomocysteineIAA [K7] | 4,32935E-05 | 2255,01747 | Brain/ FAIMS/IAA |
| Q91X97; P84075; P62748 | Q91X97 [131-147]; P84075 [131-147]; P62748 [131-147] | [K].MVSSVMKMPEDESTPEK.[R] | 1xHomocysteineMMTS [K7] | 0,00980048 | 2087,88262 | Brain/ FAIMS/MMTS |
| Q91X97; P84075; P62748 | Q91X97 [172-181]; P84075 [172-181]; P62748 [172-181] | [RK].GAKSDPSIVR.[L] | 1xHomocysteineMMTS [K3] | 0,00384072 | 1192,58129 | Brain/ FAIMS/MMTS |
| Q91X97; P84075; P62748 | Q91X97 [138-148]; P84075 [138-148]; P62748 [138-148] | [K].MPEDESTPEKR.[T] | 1xHomocysteineMMTS [K10] | 0,000266738 | 1481,60691 | Brain/ FAIMS/MMTS |
| Q91XD4 | Q91XD4 [307-320] | [R].LGLDSLAPFDPKER.[I] | 1xHomocysteineIAA [K12] | 9,61721E-05 | 1731,87343 | Liver/ FAIMS/IAA |
| Q91XD4 | Q91XD4 [307-320] | [R].LGLDSLAPFDPKER.[I] | 1xHomocysteineMMTS [K12] | 0,000530332 | 1720,83969 | Liver/ FAIMS/MMTS |
| Q91XE0 | Q91XE0 [170-192] | [K].LSSLDVVHAALVNKFWLFGGNER.[S] | 1xHomocysteineMMTS [K14] | 0,00461842 | 2735,37962 | Liver/ No FAIMS/MMTS |
| Q91XE0 | Q91XE0 [170-192] | [K].LSSLDVVHAALVNKFWLFGGNER.[S] | 1xHomocysteineMMTS [K14] | 0,00461842 | 2735,37962 | Brain/ FAIMS/MMTS |
| Q91XM9; Q811D0; P70175 | Q91XM9 [751-764]; Q811D0 [804-817]; P70175 [748-761] | [RK].GKHCILDVSGNAIK.[R] | 1xCarbamidomethyl [C4]; 1xHomocysteineIAA [K2] | 0,00631319 | 1685,84617 | Brain/ FAIMS/IAA |
| Q91XV3 | Q91XV3 [147-181] | [K].KTEAPAAAGPEAKSDAAPAASDSKPSSAEPAPSSK.[E] | 2xHomocysteineIAA [K13; K] | 4,32935E-05 | 3627,69013 | Brain/ No FAIMS/IAA |
| Q91XV3 | Q91XV3 [147-181] | [K].KTEAPAAAGPEAKSDAAPAASDSKPSSAEPAPSSK.[E] | 2xHomocysteineIAA [K13; K] | 4,32935E-05 | 3627,69013 | Brain/ FAIMS/IAA |
| Q91XV3 | Q91XV3 [25-52] | [K].KAEGAGTEEEGTPKESEPQAAADATEVK.[E] | 1xHomocysteineMMTS [K14] | 4,49774E-05 | 2993,33504 | Brain/ No FAIMS/MMTS |
| Q91XV3 | Q91XV3 [61-72] | [K].DAADGEAKAEEK.[E] | 1xHomocysteineMMTS [K8] | 4,49774E-05 | 1396,57191 | Brain/ No FAIMS/MMTS |
| Q91XV3 | Q91XV3 [82-91] | [K].EEAPKAEPEK.[S] | 1xHomocysteineMMTS [K5] | 0,00106681 | 1290,57045 | Brain/ No FAIMS/MMTS |
| Q91XV3 | Q91XV3 [87-123] | [K].AEPEKSEGAAEEQPEPAPAPEQEAAAPGPAAGGEAPK.[A] | 1xHomocysteineMMTS [K5] | 4,49774E-05 | 3728,66906 | Brain/ No FAIMS/MMTS |
| Q91XV3 | Q91XV3 [196-226] | [K].APAPAAPAAAEPQAEAPAAAASSEQSVAVKE.[-] | 1xHomocysteineMMTS [K30] | 4,49774E-05 | 3021,42921 | Brain/ No FAIMS/MMTS |
| Q91XV3 | Q91XV3 [124-147] | [K].AGEASAESTGAADGAAPEEGEAKK.[T] | 1xHomocysteineMMTS [K23] | 4,49774E-05 | 2367,00749 | Brain/ No FAIMS/MMTS |
| Q91XV3 | Q91XV3 [160-195] | [K].SDAAPAASDSKPSSAEPAPSSKETPAASEAPSSAAK.[A] | 1xHomocysteineMMTS [K22] | 4,49774E-05 | 3518,58975 | Brain/ No FAIMS/MMTS |
| Q91XV3 | Q91XV3 [39-60] | [K].ESEPQAAADATEVKESTEEKPK.[D] | 1xHomocysteineMMTS [K14] | 4,49774E-05 | 2537,13817 | Brain/ No FAIMS/MMTS |
| Q91XV3 | Q91XV3 [26-52] | [K].AEGAGTEEEGTPKESEPQAAADATEVK.[E] | 1xHomocysteineMMTS [K13] | 4,49774E-05 | 2865,24007 | Brain/ No FAIMS/MMTS |
| Q91XV3 | Q91XV3 [147-181] | [K].KTEAPAAAGPEAKSDAAPAASDSKPSSAEPAPSSK.[E] | 1xHomocysteineMMTS [K13] | 4,49774E-05 | 3442,61009 | Brain/ No FAIMS/MMTS |
| Q91XV3 | Q91XV3 [160-181] | [K].SDAAPAASDSKPSSAEPAPSSK.[E] | 1xHomocysteineMMTS [K11] | 4,49774E-05 | 2220,97474 | Brain/ No FAIMS/MMTS |
| Q91XV3 | Q91XV3 [147-159] | [K].KTEAPAAAGPEAK.[S] | 1xHomocysteineMMTS [K1] | 4,49774E-05 | 1403,66575 | Brain/ No FAIMS/MMTS |
| Q91XV3 | Q91XV3 [25-38] | [K].KAEGAGTEEEGTPK.[E] | 1xHomocysteineMMTS [K1] | 4,49774E-05 | 1566,67743 | Brain/ No FAIMS/MMTS |
| Q91XV3 | Q91XV3 [53-68] | [K].ESTEEKPKDAADGEAK.[A] | 1xHomocysteineMMTS [K] | 4,49774E-05 | 1867,80482 | Brain/ No FAIMS/MMTS |
| Q91XV3 | Q91XV3 [73-86] | [K].EADKAAAAKEEAPK.[A] | 1xHomocysteineMMTS [K] | 0,00239411 | 1591,74545 | Brain/ No FAIMS/MMTS |
| Q91XV3 | Q91XV3 [25-52] | [K].KAEGAGTEEEGTPKESEPQAAADATEVK.[E] | 1xHomocysteineMMTS [K14] | 4,49774E-05 | 2993,33504 | Brain/ FAIMS/MMTS |
| Q91XV3 | Q91XV3 [11-20] | [K].GYNVNDEKAK.[D] | 1xHomocysteineMMTS [K8] | 0,00924953 | 1300,56603 | Brain/ FAIMS/MMTS |
| Q91XV3 | Q91XV3 [61-72] | [K].DAADGEAKAEEK.[E] | 1xHomocysteineMMTS [K8] | 4,49774E-05 | 1396,57191 | Brain/ FAIMS/MMTS |
| Q91XV3 | Q91XV3 [61-76] | [K].DAADGEAKAEEKEADK.[A] | 1xHomocysteineMMTS [K8] | 0,00327 | 1839,77352 | Brain/ FAIMS/MMTS |

| | | | | | | |
|---|---|---|---|---|---|---|
| Q91XV3 | Q91XV3 [82-123] | [K].EEAPKAEPEKSEGAAEEQPEPAPAPEQEAAAPGPAAGGEAPK.[A] | 1xHomocysteineMMTS [K5] | 4,49774E-05 | 4282,93909 | Brain/ FAIMS/MMTS |
| Q91XV3 | Q91XV3 [82-91] | [K].EEAPKAEPEK.[S] | 1xHomocysteineMMTS [K5] | 0,00106681 | 1290,57045 | Brain/ FAIMS/MMTS |
| Q91XV3 | Q91XV3 [87-123] | [K].AEPEKSEGAAEEQPEPAPAPEQEAAAPGPAAGGEAPK.[A] | 1xHomocysteineMMTS [K5] | 4,49774E-05 | 3728,66906 | Brain/ FAIMS/MMTS |
| Q91XV3 | Q91XV3 [124-147] | [K].AGEASAESTGAADGAAPEEGEAKK.[T] | 1xHomocysteineMMTS [K23] | 4,49774E-05 | 2367,00749 | Brain/ FAIMS/MMTS |
| Q91XV3 | Q91XV3 [160-195] | [K].SDAAPAASDSKPSSAEPAPSSKETPAASEAPSSAAK.[A] | 1xHomocysteineMMTS [K22] | 4,49774E-05 | 3518,58975 | Brain/ FAIMS/MMTS |
| Q91XV3 | Q91XV3 [182-225] | [K].ETPAASEAPSSAAKAPAPAAPAAAEPQAEAPAAAASSEQSVAVK.[E] | 1xHomocysteineMMTS [K14] | 4,49774E-05 | 4190,00163 | Brain/ FAIMS/MMTS |
| Q91XV3 | Q91XV3 [39-60] | [K].ESEPQAAADATEVKESTEEKPK.[D] | 1xHomocysteineMMTS [K14] | 4,49774E-05 | 2537,13817 | Brain/ FAIMS/MMTS |
| Q91XV3 | Q91XV3 [26-52] | [K].AEGAGTEEEGTPKESEPQAAADATEVK.[E] | 1xHomocysteineMMTS [K13] | 4,49774E-05 | 2865,24007 | Brain/ FAIMS/MMTS |
| Q91XV3 | Q91XV3 [147-181] | [K].KTEAPAAAGPEAKSDAAPAASDSKPSSAEPAPSSK.[E] | 1xHomocysteineMMTS [K13] | 4,49774E-05 | 3442,61009 | Brain/ FAIMS/MMTS |
| Q91XV3 | Q91XV3 [148-181] | [K].TEAPAAAGPEAKSDAAPAASDSKPSSAEPAPSSK.[E] | 1xHomocysteineMMTS [K12] | 4,49774E-05 | 3314,51512 | Brain/ FAIMS/MMTS |
| Q91XV3 | Q91XV3 [160-181] | [K].SDAAPAASDSKPSSAEPAPSSK.[E] | 1xHomocysteineMMTS [K11] | 4,49774E-05 | 2220,97474 | Brain/ FAIMS/MMTS |
| Q91XV3 | Q91XV3 [147-159] | [K].KTEAPAAAGPEAK.[S] | 1xHomocysteineMMTS [K1] | 4,49774E-05 | 1403,66575 | Brain/ FAIMS/MMTS |
| Q91XV3 | Q91XV3 [25-38] | [K].KAEGAGTEEEGTPK.[E] | 1xHomocysteineMMTS [K1] | 4,49774E-05 | 1566,67743 | Brain/ FAIMS/MMTS |
| Q91XV3 | Q91XV3 [53-68] | [K].ESTEEKPKDAADGEAK.[A] | 1xHomocysteineMMTS [K] | 4,49774E-05 | 1867,80482 | Brain/ FAIMS/MMTS |
| Q91XV3 | Q91XV3 [73-86] | [K].EADKAAAAKEEAPK.[A] | 1xHomocysteineMMTS [K] | 0,00239411 | 1591,74545 | Brain/ FAIMS/MMTS |
| Q91XV3 | Q91XV3 [53-72] | [K].ESTEEKPKDAADGEAKAEEK.[E] | 1xHomocysteineMMTS [K] | 4,49774E-05 | 2325,02208 | Brain/ FAIMS/MMTS |
| Q91Y97 | Q91Y97 [14-22] | [K].KELSEIAQR.[I] | 1xHomocysteineIAA [K1] | 4,32935E-05 | 1247,64125 | Liver/ No FAIMS/IAA |
| Q91Y97 | Q91Y97 [14-22] | [K].KELSEIAQR.[I] | 1xHomocysteineIAA [K1] | 4,32935E-05 | 1247,64125 | Liver/ FAIMS/IAA |
| Q91Y97 | Q91Y97 [5-14] | [R].FPALTPEQKK.[E] | 1xHomocysteineMMTS [K9] | 4,49774E-05 | 1321,66429 | Liver/ No FAIMS/MMTS |
| Q91Y97 | Q91Y97 [323-331] | [K].ATQEAFMKR.[A] | 1xHomocysteineMMTS [K8] | 0,0004709 | 1244,55845 | Liver/ No FAIMS/MMTS |
| Q91Y97 | Q91Y97 [141-149] | [K].DGVDFGKWR.[A] | 1xHomocysteineMMTS [K7] | 0,000706375 | 1242,53942 | Liver/ No FAIMS/MMTS |
| Q91Y97 | Q91Y97 [102-121] | [K].GIVVGIKLDQGGAPLAGTNK.[E] | 1xHomocysteineMMTS [K7] | 4,49774E-05 | 2071,10384 | Liver/ No FAIMS/MMTS |
| Q91Y97 | Q91Y97 [23-43] | [R].IVANGKGILAADESVGTMGNR.[L] | 1xHomocysteineMMTS [K6] | 0,000980684 | 2236,08827 | Liver/ No FAIMS/MMTS |
| Q91Y97 | Q91Y97 [100-108] | [K].EKGIVVGIK.[L] | 1xHomocysteineMMTS [K2] | 0,00101551 | 1105,6108 | Liver/ No FAIMS/MMTS |
| Q91Y97 | Q91Y97 [47-57] | [R].IKVENTEENRR.[Q] | 1xHomocysteineMMTS [K2] | 4,49774E-05 | 1550,74137 | Liver/ No FAIMS/MMTS |
| Q91Y97 | Q91Y97 [47-56] | [R].IKVENTEENR.[R] | 1xHomocysteineMMTS [K2] | 0,000266738 | 1394,64026 | Liver/ No FAIMS/MMTS |
| Q91Y97 | Q91Y97 [109-134] | [K].LDQGGAPLAGTNKETTIQGLDGLSER.[C] | 1xHomocysteineMMTS [K13] | 4,49774E-05 | 2804,35532 | Liver/ No FAIMS/MMTS |
| Q91Y97 | Q91Y97 [14-22] | [K].KELSEIAQR.[I] | 1xHomocysteineMMTS [K1] | 6,70238E-05 | 1236,6075 | Liver/ No FAIMS/MMTS |
| Q91Y97 | Q91Y97 [243-259] | [K].KYTPEQVAMATVTALHR.[T] | 1xHomocysteineMMTS [K1] | 4,49774E-05 | 2079,0184 | Liver/ No FAIMS/MMTS |
| Q91Y97 | Q91Y97 [322-330] | [K].KATQEAFMK.[R] | 1xHomocysteineMMTS [K1] | 0,000829616 | 1216,5523 | Liver/ No FAIMS/MMTS |
| Q91Y97 | Q91Y97 [5-14] | [R].FPALTPEQKK.[E] | 1xHomocysteineMMTS [K9] | 4,49774E-05 | 1321,66429 | Liver/ FAIMS/MMTS |
| Q91Y97 | Q91Y97 [102-121] | [K].GIVVGIKLDQGGAPLAGTNK.[E] | 1xHomocysteineMMTS [K7] | 4,49774E-05 | 2071,10384 | Liver/ FAIMS/MMTS |
| Q91Y97 | Q91Y97 [100-108] | [K].EKGIVVGIK.[L] | 1xHomocysteineMMTS [K2] | 0,00101551 | 1105,6108 | Liver/ FAIMS/MMTS |
| Q91Y97 | Q91Y97 [47-57] | [R].IKVENTEENRR.[Q] | 1xHomocysteineMMTS [K2] | 4,49774E-05 | 1550,74137 | Liver/ FAIMS/MMTS |
| Q91Y97 | Q91Y97 [305-321] | [R].ALQASALAAWGGKAANK.[K] | 1xHomocysteineMMTS [K13] | 4,49774E-05 | 1790,90402 | Liver/ FAIMS/MMTS |
| Q91Y97 | Q91Y97 [109-134] | [K].LDQGGAPLAGTNKETTIQGLDGLSER.[C] | 1xHomocysteineMMTS [K13] | 4,49774E-05 | 2804,35532 | Liver/ FAIMS/MMTS |
| Q91Y97 | Q91Y97 [14-22] | [K].KELSEIAQR.[I] | 1xHomocysteineMMTS [K1] | 6,70238E-05 | 1236,6075 | Liver/ FAIMS/MMTS |
| Q91Y97 | Q91Y97 [243-259] | [K].KYTPEQVAMATVTALHR.[T] | 1xHomocysteineMMTS [K1] | 4,49774E-05 | 2079,0184 | Liver/ FAIMS/MMTS |
| Q91Y97 | Q91Y97 [140-147] | [K].KDGVDFGK.[W] | 1xHomocysteineMMTS [K1] | 0,00927584 | 1028,45396 | Liver/ FAIMS/MMTS |
| Q91Y97 | Q91Y97 [243-259] | [K].KYTPEQVAMATVTALHR.[T] | 1xOxidation [M9]; 1xHomocysteineMMTS [K1] | 4,49774E-05 | 2095,01331 | Liver/ FAIMS/MMTS |
| Q91Y97 | Q91Y97 [322-330] | [K].KATQEAFMK.[R] | 1xOxidation [M8]; 1xHomocysteineMMTS [K1] | 0,00567876 | 1232,54721 | Liver/ FAIMS/MMTS |
| Q91Y97 | Q91Y97 [323-331] | [K].ATQEAFMKR.[A] | 1xHomocysteineMMTS [K8] | 0,0004709 | 1244,55845 | Liver/ FAIMS/MMTS |
| Q91Y97 | Q91Y97 [141-149] | [K].DGVDFGKWR.[A] | 1xHomocysteineMMTS [K7] | 0,000706375 | 1242,53942 | Liver/ FAIMS/MMTS |
| Q91Y97 | Q91Y97 [47-56] | [R].IKVENTEENR.[R] | 1xHomocysteineMMTS [K2] | 0,000266738 | 1394,64026 | Liver/ FAIMS/MMTS |
| Q91Y97 | Q91Y97 [322-330] | [K].KATQEAFMK.[R] | 1xHomocysteineMMTS [K1] | 0,000829616 | 1216,5523 | Liver/ FAIMS/MMTS |
| Q91YT0 | Q91YT0 [72-86] | [K].GPDWILGEMKTSGLR.[G] | 1xHomocysteineIAA [K10] | 4,32935E-05 | 1833,8986 | Brain/ FAIMS/IAA |
| Q91Z53 | Q91Z53 [77-95] | [R].VISTLSVGVDHLALDEIKK.[R] | 1xHomocysteineMMTS [K18] | 0,00275712 | 2200,17159 | Liver/ FAIMS/MMTS |
| Q91ZA3 | Q91ZA3 [254-264] | [R].LLIEKFIDNPR.[H] | 1xHomocysteineMMTS [K5] | 0,000361246 | 1520,79637 | Liver/ FAIMS/MMTS |
| Q91ZX7 | Q91ZX7 [2001-2012] | [R].YVVISQGLDKPR.[A] | 1xHomocysteineIAA [K10] | 4,32935E-05 | 1548,82027 | Brain/ FAIMS/IAA |
| Q91ZX7 | Q91ZX7 [2001-2012] | [R].YVVISQGLDKPR.[A] | 1xHomocysteineMMTS [K10] | 0,00275712 | 1537,78653 | Liver/ FAIMS/MMTS |
| Q91ZZ3 | Q91ZZ3 [7-21] | [K].GLSMAKEGVVAAAEK.[T] | 1xHomocysteineIAA [K6] | 4,32935E-05 | 1634,82404 | Brain/ No FAIMS/IAA |
| Q91ZZ3 | Q91ZZ3 [7-21] | [K].GLSMAKEGVVAAAEK.[T] | 1xHomocysteineIAA [K6] | 4,32935E-05 | 1634,82404 | Brain/ FAIMS/IAA |
| Q91ZZ3 | Q91ZZ3 [44-59] | [K].TSGVVQGVASVAEKTK.[E] | 1xHomocysteineMMTS [K14] | 4,49774E-05 | 1723,87172 | Brain/ No FAIMS/MMTS |
| Q91ZZ3 | Q91ZZ3 [7-21] | [K].GLSMAKEGVVAAAEK.[T] | 1xHomocysteineMMTS [K6] | 4,49774E-05 | 1623,7903 | Brain/ FAIMS/MMTS |
| Q91ZZ3 | Q91ZZ3 [44-59] | [K].TSGVVQGVASVAEKTK.[E] | 1xHomocysteineMMTS [K14] | 4,49774E-05 | 1723,87172 | Brain/ FAIMS/MMTS |
| Q91ZZ3; O55042 | Q91ZZ3 [33-43]; O55042 [33-43] | [K].TKEGVLYVGSK.[T] | 1xHomocysteineIAA [K2] | 4,32935E-05 | 1354,70351 | Brain/ No FAIMS/IAA |
| Q91ZZ3; O55042 | Q91ZZ3 [33-43]; O55042 [33-43] | [K].TKEGVLYVGSK.[T] | 1xHomocysteineIAA [K2] | 4,32935E-05 | 1354,70351 | Brain/ FAIMS/IAA |
| Q91ZZ3; O55042 | Q91ZZ3 [13-23]; O55042 [13-23] | [K].EGVVAAAEKTK.[Q] | 1xHomocysteineMMTS [K9] | 0,00074635 | 1265,62282 | Brain/ No FAIMS/MMTS |
| Q91ZZ3; Q9Z0F7 | Q91ZZ3 [24-34]; Q9Z0F7 [24-34] | [K].QGVTEAAEKTK.[E] | 1xHomocysteineIAA [K9] | 4,32935E-05 | 1335,65729 | Brain/ No FAIMS/IAA |
| Q91ZZ3; Q9Z0F7 | Q91ZZ3 [24-34]; Q9Z0F7 [24-34] | [K].QGVTEAAEKTK.[E] | 1xHomocysteineIAA [K9] | 4,32935E-05 | 1335,65729 | Brain/ FAIMS/IAA |
| Q91ZZ3; Q9Z0F7 | Q91ZZ3 [22-32]; Q9Z0F7 [22-32] | [K].TKQGVTEAAEK.[T] | 1xHomocysteineMMTS [K2] | 0,000173718 | 1324,62355 | Brain/ No FAIMS/MMTS |
| Q91ZZ3; Q9Z0F7 | Q91ZZ3 [24-34]; Q9Z0F7 [24-34] | [K].QGVTEAAEKTK.[E] | 1xHomocysteineMMTS [K9] | 0,000530332 | 1324,62355 | Brain/ FAIMS/MMTS |
| Q91ZZ3; Q9Z0F7 | Q91ZZ3 [22-32]; Q9Z0F7 [22-32] | [K].TKQGVTEAAEK.[T] | 1xHomocysteineMMTS [K2] | 0,000173718 | 1324,62355 | Brain/ FAIMS/MMTS |

| | | | | | | |
|---|---|---|---|---|---|---|
| Q921G7 | Q921G7 [88-98] | [K].QLAAEQGKDIR.[V] | 1xHomocysteineMMTS [K8] | 0,0031519 | 1391,67698 | Liver/ FAIMS/MMTS |
| Q921G7 | Q921G7 [146-154] | [R].FAILTEKHR.[I] | 1xHomocysteineMMTS [K7] | 0,000282457 | 1277,64931 | Liver/ FAIMS/MMTS |
| Q921G7 | Q921G7 [344-357] | [K].HHPSIQPTLEGGKR.[I] | 1xHomocysteineMMTS [K13] | 0,001242 | 1719,84175 | Liver/ FAIMS/MMTS |
| Q921G7 | Q921G7 [405-425] | [K].SGSLAAESIFKQLTSENLQSK.[T] | 1xHomocysteineMMTS [K11] | 4,49774E-05 | 2401,17377 | Liver/ FAIMS/MMTS |
| Q921H8 | Q921H8 [78-102] | [R].LKPEQLGDISVGNVLEPGAGAVMAR.[I] | 1xHomocysteineMMTS [K2] | 0,000738471 | 2684,35684 | Liver/ FAIMS/MMTS |
| Q921I1 | Q921I1 [252-264] | [R].KPVDQYEDCYLAR.[I] | 1xCarbamidomethyl [C9]; 1xHomocysteineIAA [K1] | 4,32935E-05 | 1830,81493 | Liver/ FAIMS/IAA |
| Q921I1 | Q921I1 [252-264] | [R].KPVDQYEDCYLAR.[I] | 1xCarbamidomethyl [C9]; 1xHomocysteineIAA [K1] | 4,32935E-05 | 1830,81493 | Brain/ FAIMS/IAA |
| Q921I1 | Q921I1 [642-650] | [K].STTKDLLFR.[D] | 1xHomocysteineMMTS [K4] | 0,00219832 | 1243,61734 | Liver/ No FAIMS/MMTS |
| Q921I1 | Q921I1 [298-310] | [K].SKDFQLFSSPLGK.[D] | 1xHomocysteineMMTS [K2] | 0,000211896 | 1616,78111 | Liver/ No FAIMS/MMTS |
| Q921I1 | Q921I1 [123-135] | [K].GTDFQLNQLEGKK.[S] | 1xHomocysteineMMTS [K12] | 0,000211896 | 1640,77709 | Liver/ No FAIMS/MMTS |
| Q921I1 | Q921I1 [555-572] | [K].HQTVLDNTEGKNPAEWAK.[N] | 1xHomocysteineMMTS [K11] | 4,49774E-05 | 2201,0114 | Liver/ No FAIMS/MMTS |
| Q921I1 | Q921I1 [573-588] | [K].NLKQEDFELLCPDGTR.[K] | 1xMethylthio [C11]; 1xHomocysteineMMTS [K3] | 0,00714378 | 2086,90646 | Liver/ FAIMS/MMTS |
| Q921I1 | Q921I1 [311-323] | [K].DLLFKDSAFGLLR.[V] | 1xHomocysteineMMTS [K5] | 4,49774E-05 | 1657,84405 | Liver/ FAIMS/MMTS |
| Q921I1 | Q921I1 [298-310] | [K].SKDFQLFSSPLGK.[D] | 1xHomocysteineMMTS [K2] | 0,000211896 | 1616,78111 | Liver/ FAIMS/MMTS |
| Q921I1 | Q921I1 [122-134] | [K].KGTDFQLNQLEGK.[K] | 1xHomocysteineMMTS [K1] | 0,000152766 | 1640,77709 | Liver/ FAIMS/MMTS |
| Q921I1 | Q921I1 [288-297] | [K].VAQEHFGKGK.[S] | 1xHomocysteineMMTS [K8] | 0,00225753 | 1263,59727 | Liver/ FAIMS/MMTS |
| Q921I1 | Q921I1 [548-572] | [K].GDVAFVKHQTVLDNTEGKNPAEWAK.[N] | 1xHomocysteineMMTS [K7] | 4,49774E-05 | 2917,39712 | Liver/ FAIMS/MMTS |
| Q921I1 | Q921I1 [642-650] | [K].STTKDLLFR.[D] | 1xHomocysteineMMTS [K4] | 0,00219832 | 1243,61734 | Liver/ FAIMS/MMTS |
| Q921I1 | Q921I1 [298-315] | [K].SKDFQLFSSPLGKDLLFK.[D] | 1xHomocysteineMMTS [K13] | 0,00760806 | 2233,13956 | Liver/ FAIMS/MMTS |
| Q921I1 | Q921I1 [123-135] | [K].GTDFQLNQLEGKK.[S] | 1xHomocysteineMMTS [K12] | 0,000211896 | 1640,77709 | Liver/ FAIMS/MMTS |
| Q921I1 | Q921I1 [456-469] | [K].ASDTSITWNNLKGK.[K] | 1xHomocysteineMMTS [K12] | 0,00439215 | 1697,79855 | Liver/ FAIMS/MMTS |
| Q921I1 | Q921I1 [555-572] | [K].HQTVLDNTEGKNPAEWAK.[N] | 1xHomocysteineMMTS [K11] | 4,49774E-05 | 2201,0114 | Liver/ FAIMS/MMTS |
| Q921I1 | Q921I1 [216-236] | [K].DGGGDVAFVKHTTIFEVLPEK.[A] | 1xHomocysteineMMTS [K10] | 0,00210568 | 2422,17813 | Liver/ FAIMS/MMTS |
| Q921I1 | Q921I1 [298-310] | [K].SKDFQLFSSPLGK.[D] | 1xHomocysteineMMTS [K2] | 0,000211896 | 1616,78111 | Brain/ FAIMS/MMTS |
| Q921M7 | Q921M7 [187-207] | [R].MSLFYAEATPMLKTLSDATTK.[F] | 1xHomocysteineIAA [K13] | 4,32935E-05 | 2493,20738 | Brain/ FAIMS/IAA |
| Q921M7; Q8BHZ0 | Q921M7 [279-286]; Q8BHZ0 [278-285] | [K].IDMKGCIK.[V] | 1xCarbamidomethyl [C6]; 1xHomocysteineIAA [K4] | 0,00162159 | 1138,54173 | Brain/ FAIMS/IAA |
| Q922B1 | Q922B1 [85-97] | [K].VDLSTSTDWKEAK.[S] | 1xHomocysteineIAA [K10] | 0,00391874 | 1653,77886 | Brain/ FAIMS/IAA |
| Q922F4 | Q922F4 [360-379] | [R].GLKMASTFIGNSTAIQELFK.[R] | 1xHomocysteineIAA [K3] | 6,30912E-05 | 2330,1883 | Brain/ FAIMS/IAA |
| Q922J6 | Q922J6 [155-172] | [K].ESSEQVQPTCPKELPGHK.[N] | 1xCarbamidomethyl [C10]; 1xHomocysteineIAA [K12] | 4,32935E-05 | 2225,03253 | Brain/ No FAIMS/IAA |
| Q922J6 | Q922J6 [155-172] | [K].ESSEQVQPTCPKELPGHK.[N] | 1xCarbamidomethyl [C10]; 1xHomocysteineIAA [K12] | 4,32935E-05 | 2225,03253 | Brain/ FAIMS/IAA |
| Q922Q1 | Q922Q1 [179-189] | [R].LVQFDTSMKGR.[T] | 1xHomocysteineIAA [K9] | 0,000698725 | 1455,70828 | Brain/ FAIMS/IAA |
| Q922R8 | Q922R8 [232-241] | [R].YGIKGFPTIK.[I] | 1xHomocysteineIAA [K4] | 0,00399809 | 1297,69731 | Liver/ FAIMS/IAA |
| Q922R8 | Q922R8 [242-256] | [K].IFQKGESPVDYDGGR.[T] | 1xHomocysteineIAA [K4] | 0,00124346 | 1841,84867 | Brain/ FAIMS/IAA |
| Q922R8 | Q922R8 [68-77] | [K].AATALKDVVK.[V] | 1xHomocysteineMMTS [K6] | 0,000879882 | 1178,62718 | Liver/ FAIMS/MMTS |
| Q922R8 | Q922R8 [369-386] | [K].FALLKGSFSEQGINEFLR.[E] | 1xHomocysteineMMTS [K5] | 4,49774E-05 | 2219,09876 | Liver/ FAIMS/MMTS |
| Q922R8 | Q922R8 [242-256] | [K].IFQKGESPVDYDGGR.[T] | 1xHomocysteineMMTS [K4] | 0,000437433 | 1830,81493 | Liver/ FAIMS/MMTS |
| Q923D2 | Q923D2 [135-145] | [R].MHKILQESGLK.[Y] | 1xHomocysteineIAA [K3] | 0,00012645 | 1457,76032 | Liver/ FAIMS/IAA |
| Q923D2 | Q923D2 [93-105] | [R].NIVTAMKAHGVDK.[V] | 1xHomocysteineMMTS [K7] | 0,00427174 | 1546,75385 | Liver/ FAIMS/MMTS |
| Q923D2 | Q923D2 [175-187] | [R].VISKHDLGHFMLR.[C] | 1xHomocysteineMMTS [K4] | 4,49774E-05 | 1715,85423 | Liver/ FAIMS/MMTS |
| Q93092 | Q93092 [259-269] | [K].LLGELLKDNSK.[L] | 1xHomocysteineIAA [K7] | 0,00012645 | 1403,75628 | Liver/ FAIMS/IAA |
| Q93092 | Q93092 [205-219] | [K].SYEPQEDPGVKSVTK.[I] | 1xHomocysteineIAA [K11] | 4,32935E-05 | 1837,86365 | Brain/ FAIMS/IAA |
| Q93092 | Q93092 [287-295] | [K].IHLDEKAFR.[W] | 1xHomocysteineMMTS [K6] | 0,00483466 | 1291,62857 | Liver/ No FAIMS/MMTS |
| Q93092 | Q93092 [270-286] | [K].LAPALSVKAAQTSDSEK.[I] | 1xHomocysteineMMTS [K8] | 4,49774E-05 | 1878,92996 | Liver/ FAIMS/MMTS |
| Q93092 | Q93092 [259-269] | [K].LLGELLKDNSK.[L] | 1xHomocysteineMMTS [K7] | 0,00510827 | 1392,72253 | Liver/ FAIMS/MMTS |
| Q93092 | Q93092 [315-324] | [K].FAADAIKLER.[M] | 1xHomocysteineMMTS [K7] | 0,00171109 | 1296,64389 | Liver/ FAIMS/MMTS |
| Q93092 | Q93092 [287-295] | [K].IHLDEKAFR.[W] | 1xHomocysteineMMTS [K6] | 0,00483466 | 1291,62857 | Liver/ FAIMS/MMTS |
| Q93092 | Q93092 [125-136] | [R].LIELYKEAGVGK.[D] | 1xHomocysteineMMTS [K6] | 0,00501021 | 1482,76948 | Liver/ FAIMS/MMTS |
| Q93092 | Q93092 [205-219] | [K].SYEPQEDPGVKSVTK.[I] | 1xHomocysteineMMTS [K11] | 0,000192903 | 1826,82991 | Liver/ FAIMS/MMTS |
| Q99020 | Q99020 [233-250] | [K].VAQPKEVYQQQQYGSGGR.[G] | 1xHomocysteineIAA [K5] | 4,32935E-05 | 2197,04548 | Liver/ FAIMS/IAA |
| Q99020 | Q99020 [117-131] | [R].GFGFILFKDSSSVEK.[V] | 1xHomocysteineIAA [K8] | 4,32935E-05 | 1834,9044 | Brain/ FAIMS/IAA |
| Q99020 | Q99020 [117-131] | [R].GFGFILFKDSSSVEK.[V] | 1xHomocysteineMMTS [K8] | 0,0018995 | 1823,87065 | Liver/ FAIMS/MMTS |
| Q99020 | Q99020 [233-250] | [K].VAQPKEVYQQQQYGSGGR.[G] | 1xHomocysteineMMTS [K5] | 0,00593578 | 2186,01173 | Liver/ FAIMS/MMTS |
| Q99020 | Q99020 [161-176] | [K].IFVGGLNPEATEEKIR.[E] | 1xHomocysteineMMTS [K14] | 0,0004709 | 1935,96668 | Liver/ FAIMS/MMTS |
| Q99J08 | Q99J08 [5-18] | [R].VGDLSPKQEEALAK.[F] | 1xHomocysteineIAA [K7] | 9,61721E-05 | 1658,8418 | Liver/ FAIMS/IAA |
| Q99J08 | Q99J08 [5-18] | [R].VGDLSPKQEEALAK.[F] | 1xHomocysteineMMTS [K7] | 0,000599431 | 1647,80805 | Liver/ No FAIMS/MMTS |
| Q99J08 | Q99J08 [110-122] | [K].GLLFSASKQDLLR.[T] | 1xHomocysteineMMTS [K8] | 4,49774E-05 | 1610,83929 | Liver/ FAIMS/MMTS |
| Q99J08 | Q99J08 [46-57] | [R].SFDLQKSEAMLR.[K] | 1xHomocysteineMMTS [K6] | 0,000250158 | 1587,73278 | Liver/ FAIMS/MMTS |
| Q99J08 | Q99J08 [389-403] | [K].AAEEKMNQQGADTPK.[-] | 1xHomocysteineMMTS [K5] | 0,000507453 | 1780,76627 | Liver/ FAIMS/MMTS |
| Q99J85 | Q99J85 [258-270] | [R].QEVEKELNALQGR.[V] | 1xHomocysteineIAA [K5] | 0,000289025 | 1687,84319 | Brain/ FAIMS/IAA |
| Q99J99 | Q99J99 [9-24] | [R].ALVSAQWVAEALKAPR.[S] | 1xHomocysteineIAA [K13] | 9,61721E-05 | 1884,01601 | Brain/ FAIMS/IAA |
| Q99JB2 | Q99JB2 [141-148] | [K].LSLDKVFR.[E] | 1xHomocysteineIAA [K5] | 0,000318237 | 1151,62414 | Brain/ FAIMS/IAA |
| Q99JI6; P62835 | Q99JI6 [150-163]; P62835 [150-163] | [K].SKINVNEIFYDLVR.[Q] | 1xHomocysteineIAA [K2] | 4,32935E-05 | 1883,96839 | Brain/ No FAIMS/IAA |
| Q99JI6; P62835 | Q99JI6 [150-163]; P62835 [150-163] | [K].SKINVNEIFYDLVR.[Q] | 1xHomocysteineIAA [K2] | 4,32935E-05 | 1883,96839 | Brain/ FAIMS/IAA |
| Q99JP6 | Q99JP6 [245-261] | [R].AGEKEATSQSVEQLEAR.[V] | 1xHomocysteineIAA [K4] | 4,32935E-05 | 2006,94476 | Brain/ FAIMS/IAA |
| Q99JP6; Q9Z2Y3; Q9QWW1 | Q99JP6 [73-84]; Q9Z2Y3 [70-81]; Q9QWW1 [70-81] | [K].TSQKFGQWADSR.[A] | 1xHomocysteineIAA [K4] | 0,00012645 | 1584,72235 | Brain/ FAIMS/IAA |
| Q99JP7 | Q99JP7 [45-63] | [R].KDEDAFLGDPDTDPDSFLK.[S] | 1xHomocysteineIAA [K1] | 0,00124346 | 2299,00708 | Brain/ FAIMS/IAA |
| Q99JW2 | Q99JW2 [1-17] | [-].MTTKDPESEHPSVTLFR.[Q] | 1xMet-loss+Acetyl [N-Term]; 1xHomocysteineIAA [K4] | 4,32935E-05 | 2059,97533 | Brain/ FAIMS/IAA |
| Q99JX3 | Q99JX3 [212-228] | [K].KISLPGQMTGTPITPLK.[D] | 1xHomocysteineIAA [K1] | 0,0014138 | 1956,06567 | Brain/ FAIMS/IAA |

| | | | | | | |
|---|---|---|---|---|---|---|
| Q99JY9 | Q99JY9 [92-102] | [R].FMEQVIFKYLR.[A] | 1xHomocysteineIAA [K8] | 0,000318237 | 1647,83857 | Brain/ FAIMS/IAA |
| Q99K51 | Q99K51 [1-23] | [-].MDEMATTQISKDELDELKEAFAK.[V] | 1xMet-loss [N-Term]; 1xHomocysteineIAA [K11] | 4,32935E-05 | 2686,25862 | Liver/ No FAIMS/IAA |
| Q99K51 | Q99K51 [1-23] | [-].MDEMATTQISKDELDELKEAFAK.[V] | 1xMet-loss [N-Term]; 1xHomocysteineIAA [K11] | 4,32935E-05 | 2686,25862 | Brain/ No FAIMS/IAA |
| Q99K51 | Q99K51 [531-545] | [R].TLSEAGKSTSIQSFK.[D] | 1xHomocysteineMMTS [K7] | 4,49774E-05 | 1746,84008 | Liver/ FAIMS/MMTS |
| Q99K85 | Q99K85 [118-133] | [K].FGTVNIVHPKLGSYTK.[I] | 1xHomocysteineIAA [K10] | 8,35381E-05 | 1935,01568 | Brain/ FAIMS/IAA |
| Q99KB8 | Q99KB8 [118-130] | [K].LEPGLKVYGGDDR.[I] | 1xHomocysteineIAA [K6] | 9,61721E-05 | 1592,77372 | Brain/ FAIMS/IAA |
| Q99KB8 | Q99KB8 [118-130] | [K].LEPGLKVYGGDDR.[I] | 1xHomocysteineMMTS [K6] | 0,000192903 | 1581,73997 | Liver/ FAIMS/MMTS |
| Q99KB8 | Q99KB8 [234-250] | [R].HVEPGNAAIQEKLAWAK.[E] | 1xHomocysteineMMTS [K12] | 0,000738471 | 2025,00446 | Liver/ FAIMS/MMTS |
| Q99KF1 | Q99KF1 [159-167] | [K].DKLSELQLR.[V] | 1xHomocysteineMMTS [K2] | 0,00074635 | 1264,6388 | Liver/ FAIMS/MMTS |
| Q99KI0 | Q99KI0 [396-409] | [R].SAAVAKQALAHGLK.[C] | 1xHomocysteineIAA [K6] | 4,32935E-05 | 1538,84716 | Liver/ No FAIMS/IAA |
| Q99KI0 | Q99KI0 [396-409] | [R].SAAVAKQALAHGLK.[C] | 1xHomocysteineIAA [K6] | 4,32935E-05 | 1538,84716 | Brain/ No FAIMS/IAA |
| Q99KI0 | Q99KI0 [396-409] | [R].SAAVAKQALAHGLK.[C] | 1xHomocysteineIAA [K6] | 4,32935E-05 | 1538,84716 | Brain/ FAIMS/IAA |
| Q99KI0 | Q99KI0 [45-56] | [R].YDLLEKNINIVR.[K] | 1xHomocysteineMMTS [K6] | 0,000133296 | 1652,84986 | Brain/ FAIMS/MMTS |
| Q99KI0 | Q99KI0 [143-160] | [R].AKDINQEVYNFLATAGAK.[Y] | 1xHomocysteineMMTS [K2] | 0,000231655 | 2116,02017 | Brain/ FAIMS/MMTS |
| Q99KI0 | Q99KI0 [30-44] | [R].AKVAMSHFEPSEYIR.[Y] | 1xHomocysteineMMTS [K2] | 0,000507453 | 1927,88632 | Brain/ FAIMS/MMTS |
| Q99KI0 | Q99KI0 [507-520] | [K].FNPETDFLTGKDGK.[K] | 1xHomocysteineMMTS [K11] | 0,000266738 | 1731,77167 | Brain/ FAIMS/MMTS |
| Q99KJ8 | Q99KJ8 [157-176] | [K].LLGPDAAINLADPDGALAKR.[L] | 1xHomocysteineIAA [K19] | 4,32935E-05 | 2165,13831 | Brain/ No FAIMS/IAA |
| Q99KJ8 | Q99KJ8 [157-176] | [K].LLGPDAAINLADPDGALAKR.[L] | 1xHomocysteineIAA [K19] | 4,32935E-05 | 2165,13831 | Brain/ FAIMS/IAA |
| Q99KP3 | Q99KP3 [186-197] | [R].VLKEIDGFVLNR.[L] | 1xHomocysteineIAA [K3] | 0,000348152 | 1576,85157 | Brain/ FAIMS/IAA |
| Q99KR3 | Q99KR3 [230-242] | [R].DNFEKSFTVTELR.[T] | 1xHomocysteineMMTS [K5] | 8,93346E-05 | 1748,79822 | Liver/ FAIMS/MMTS |
| Q99KR3 | Q99KR3 [190-209] | [K].IKANIIYPGHGPVIHNAEAK.[I] | 1xHomocysteineMMTS [K2] | 4,49774E-05 | 2305,19439 | Liver/ FAIMS/MMTS |
| Q99KR7 | Q99KR7 [61-72] | [R].VVLELKADVVPK.[T] | 1xHomocysteineMMTS [K6] | 0,00351971 | 1472,82152 | Liver/ FAIMS/MMTS |
| Q99L13 | Q99L13 [76-94] | [K].EFKEAGEQVASSPAEVAEK.[A] | 1xHomocysteineIAA [K3] | 0,0027602 | 2180,01759 | Liver/ FAIMS/IAA |
| Q99L13 | Q99L13 [238-249] | [K].LLAKILNMSSGR.[C] | 1xHomocysteineIAA [K4] | 0,000624482 | 1476,80252 | Brain/ FAIMS/IAA |
| Q99L13 | Q99L13 [320-330] | [K].KDFSSVFQYLR.[E] | 1xHomocysteineMMTS [K1] | 0,00461842 | 1552,72868 | Liver/ No FAIMS/MMTS |
| Q99L13 | Q99L13 [76-94] | [K].EFKEAGEQVASSPAEVAEK.[A] | 1xHomocysteineMMTS [K3] | 0,000133296 | 2168,98385 | Liver/ FAIMS/MMTS |
| Q99L13 | Q99L13 [125-144] | [K].GSLLIDSSTIDPSVSKELAK.[E] | 1xHomocysteineMMTS [K16] | 0,000250158 | 2223,1247 | Liver/ FAIMS/MMTS |
| Q99L13 | Q99L13 [284-310] | [K].DLGLAQDSATSTKTPILLGSLAHQIYR.[M] | 1xHomocysteineMMTS [K13] | 4,49774E-05 | 3032,55435 | Liver/ FAIMS/MMTS |
| Q99L13 | Q99L13 [320-330] | [K].KDFSSVFQYLR.[E] | 1xHomocysteineMMTS [K1] | 0,00461842 | 1552,72868 | Liver/ FAIMS/MMTS |
| Q99L13 | Q99L13 [124-140] | [K].KGSLLIDSSTIDPSVSK.[E] | 1xHomocysteineMMTS [K1] | 4,49774E-05 | 1909,96093 | Liver/ FAIMS/MMTS |
| Q99L47 | Q99L47 [117-131] | [K].KGAAIEALNDGELQK.[A] | 1xHomocysteineIAA [K1] | 4,32935E-05 | 1730,87416 | Brain/ No FAIMS/IAA |
| Q99L47 | Q99L47 [117-131] | [K].KGAAIEALNDGELQK.[A] | 1xHomocysteineIAA [K1] | 4,32935E-05 | 1730,87416 | Brain/ FAIMS/IAA |
| Q99L47 | Q99L47 [132-145] | [K].AIDLFTDAIKLNPR.[L] | 1xHomocysteineMMTS [K10] | 0,00214892 | 1749,90262 | Liver/ FAIMS/MMTS |
| Q99LC5 | Q99LC5 [127-146] | [K].LNVAPVSDIIEIKSPDTFVR.[T] | 1xHomocysteineIAA [K13] | 4,32935E-05 | 2387,26391 | Liver/ FAIMS/IAA |
| Q99LC5 | Q99LC5 [227-249] | [K].SGENFKLLYDLADQLHAAVGASR.[A] | 1xHomocysteineIAA [K6] | 4,32935E-05 | 2649,30896 | Brain/ FAIMS/IAA |
| Q99LC5 | Q99LC5 [70-85] | [K].VAGVAKVLVAQHDAYK.[G] | 1xHomocysteineMMTS [K6] | 4,49774E-05 | 1831,95572 | Liver/ No FAIMS/MMTS |
| Q99LC5 | Q99LC5 [123-139] | [R].VAAKLNVAPVSDIIEIK.[S] | 1xHomocysteineMMTS [K4] | 4,49774E-05 | 1943,07042 | Liver/ No FAIMS/MMTS |
| Q99LC5 | Q99LC5 [204-216] | [K].LTKSDRPELTGAK.[V] | 1xHomocysteineMMTS [K3] | 4,49774E-05 | 1578,79782 | Liver/ No FAIMS/MMTS |
| Q99LC5 | Q99LC5 [188-206] | [K].APSSSSVGISEWLDQKLTK.[S] | 1xHomocysteineMMTS [K16] | 4,49774E-05 | 2196,06752 | Liver/ No FAIMS/MMTS |
| Q99LC5 | Q99LC5 [127-146] | [K].LNVAPVSDIIEIKSPDTFVR.[T] | 1xHomocysteineMMTS [K13] | 4,49774E-05 | 2376,23016 | Liver/ No FAIMS/MMTS |
| Q99LC5 | Q99LC5 [76-101] | [K].VLVAQHDAYKGLLPEELTPLILETQK.[Q] | 1xHomocysteineMMTS [K10] | 4,49774E-05 | 3081,63629 | Liver/ No FAIMS/MMTS |
| Q99LC5 | Q99LC5 [322-332] | [K].VVPEMTEILKK.[K] | 1xHomocysteineMMTS [K10] | 0,000798864 | 1449,75139 | Liver/ No FAIMS/MMTS |
| Q99LC5 | Q99LC5 [207-223] | [K].SDRPELTGAKVVVSGGR.[G] | 1xHomocysteineMMTS [K10] | 4,49774E-05 | 1890,95243 | Liver/ No FAIMS/MMTS |
| Q99LC5 | Q99LC5 [123-139] | [R].VAAKLNVAPVSDIIEIK.[S] | 1xHomocysteineMMTS [K4] | 4,49774E-05 | 1943,07042 | Liver/ FAIMS/MMTS |
| Q99LC5 | Q99LC5 [204-216] | [K].LTKSDRPELTGAK.[V] | 1xHomocysteineMMTS [K3] | 4,49774E-05 | 1578,79782 | Liver/ FAIMS/MMTS |
| Q99LC5 | Q99LC5 [188-206] | [K].APSSSSVGISEWLDQKLTK.[S] | 1xHomocysteineMMTS [K16] | 4,49774E-05 | 2196,06752 | Liver/ FAIMS/MMTS |
| Q99LC5 | Q99LC5 [127-146] | [K].LNVAPVSDIIEIKSPDTFVR.[T] | 1xHomocysteineMMTS [K13] | 4,49774E-05 | 2376,23016 | Liver/ FAIMS/MMTS |
| Q99LC5 | Q99LC5 [322-332] | [K].VVPEMTEILKK.[K] | 1xHomocysteineMMTS [K10] | 0,000798864 | 1449,75139 | Liver/ FAIMS/MMTS |
| Q99LC5 | Q99LC5 [207-223] | [K].SDRPELTGAKVVVSGGR.[G] | 1xHomocysteineMMTS [K10] | 4,49774E-05 | 1890,95243 | Liver/ FAIMS/MMTS |
| Q99LC5 | Q99LC5 [70-85] | [K].VAGVAKVLVAQHDAYK.[G] | 1xHomocysteineMMTS [K6] | 4,49774E-05 | 1831,95572 | Liver/ FAIMS/MMTS |
| Q99LC5 | Q99LC5 [170-203] | [R].GTSFEAAATSGGSASSEKAPSSSSVGISE WLDQK.[L] | 1xHomocysteineMMTS [K18] | 0,000266738 | 3479,55772 | Liver/ FAIMS/MMTS |
| Q99LC5 | Q99LC5 [76-101] | [K].VLVAQHDAYKGLLPEELTPLILETQK.[Q] | 1xHomocysteineMMTS [K10] | 4,49774E-05 | 3081,63629 | Liver/ FAIMS/MMTS |
| Q99LC5 | Q99LC5 [123-139] | [R].VAAKLNVAPVSDIIEIK.[S] | 1xHomocysteineMMTS [K4] | 4,49774E-05 | 1943,07042 | Brain/ FAIMS/MMTS |
| Q99LC5 | Q99LC5 [188-206] | [K].APSSSSVGISEWLDQKLTK.[S] | 1xHomocysteineMMTS [K16] | 4,49774E-05 | 2196,06752 | Brain/ FAIMS/MMTS |
| Q99LD4 | Q99LD4 [56-68] | [R].VEALKMALSFVQR.[T] | 1xHomocysteineIAA [K5] | 0,000318237 | 1665,88149 | Brain/ FAIMS/IAA |
| Q99LI8 | Q99LI8 [203-210] | [K].FGIEKEVR.[V] | 1xHomocysteineIAA [K5] | 0,000258769 | 1151,58775 | Brain/ FAIMS/IAA |
| Q99LX0 | Q99LX0 [33-48] | [K].VTVAGLAGKDPVQCSR.[D] | 1xCarbamidomethyl [C14]; 1xHomocysteineIAA [K9] | 4,32935E-05 | 1831,91531 | Liver/ FAIMS/IAA |
| Q99LX0 | Q99LX0 [99-122] | [R].KGLIAAICAGPTALLAHEVGFGCK.[V] | 2xCarbamidomethyl [C8; C23]; 1xHomocysteineIAA [K1] | 4,32935E-05 | 2628,34588 | Brain/ FAIMS/IAA |
| Q99LX0 | Q99LX0 [123-132] | [K].VTTHPLAKDK.[M] | 1xHomocysteineMMTS [K8] | 0,00255629 | 1272,64389 | Liver/ No FAIMS/MMTS |
| Q99LX0 | Q99LX0 [183-189] | [K].APLVLKD.[-] | 1xHomocysteineMMTS [K6] | 0,00427174 | 918,47872 | Liver/ No FAIMS/MMTS |
| Q99LX0 | Q99LX0 [29-41] | [R].AGIKVTVAGLAGK.[D] | 1xHomocysteineMMTS [K4] | 0,00575155 | 1347,74869 | Liver/ No FAIMS/MMTS |
| Q99LX0 | Q99LX0 [183-189] | [K].APLVLKD.[-] | 1xHomocysteineMMTS [K6] | 0,00427174 | 918,47872 | Liver/ FAIMS/MMTS |
| Q99LX0 | Q99LX0 [90-98] | [K].EILKEQESR.[K] | 1xHomocysteineMMTS [K4] | 0,0074518 | 1294,61298 | Liver/ FAIMS/MMTS |
| Q99LX0 | Q99LX0 [146-156] | [R].VEKDGLILTSR.[G] | 1xHomocysteineMMTS [K3] | 0,00221411 | 1393,71778 | Liver/ FAIMS/MMTS |
| Q99LX0 | Q99LX0 [63-93] | [K].TQGPYDVVVLPGGNLGAQNLSESPMVK EILK.[E] | 1xHomocysteineMMTS [K27] | 4,49774E-05 | 3416,72624 | Liver/ FAIMS/MMTS |
| Q99LY9 | Q99LY9 [17-32] | [R].HFMFLSAEQPYKNAAR.[C] | 1xHomocysteineIAA [K12] | 0,00012645 | 2083,98406 | Brain/ FAIMS/IAA |
| Q99MP8 | Q99MP8 [458-465] | [R].LSDLLKEK.[Q] | 1xHomocysteineMMTS [K6] | 0,00191513 | 1108,57408 | Brain/ FAIMS/MMTS |
| Q99N28 | Q99N28 [120-137] | [K].SLVTVLGIPQKPIITGYK.[S] | 1xHomocysteineIAA [K11] | 4,32935E-05 | 2101,20896 | Brain/ No FAIMS/IAA |
| Q99N28 | Q99N28 [120-137] | [K].SLVTVLGIPQKPIITGYK.[S] | 1xHomocysteineIAA [K11] | 4,32935E-05 | 2101,20896 | Brain/ FAIMS/IAA |
| Q99P30 | Q99P30 [13-22] | [R].NNLIDDAKAR.[L] | 1xHomocysteineIAA [K8] | 4,32935E-05 | 1303,64231 | Liver/ FAIMS/IAA |
| Q99P30 | Q99P30 [13-22] | [R].NNLIDDAKAR.[L] | 1xHomocysteineMMTS [K8] | 0,000231655 | 1292,60857 | Liver/ FAIMS/MMTS |
| Q99P72 | Q99P72 [1128-1144] | [R].HQAQIDHYLGLANKSVK.[D] | 1xHomocysteineIAA [K14] | 4,32935E-05 | 2096,07057 | Brain/ No FAIMS/IAA |
| Q99P72 | Q99P72 [1128-1144] | [R].HQAQIDHYLGLANKSVK.[D] | 1xHomocysteineIAA [K14] | 4,32935E-05 | 2096,07057 | Brain/ FAIMS/IAA |

| | | | | | | |
|---|---|---|---|---|---|---|
| Q99PJ0 | Q99PJ0 [230-257] | [K].GTGVPVGQKGTLQCEASAVPSAEFQWFK.[D] | 1xCarbamidomethyl [C14]; 1xHomocysteineIAA [K9] | 4,32935E-05 | 3153,51322 | Brain/ No FAIMS/IAA |
| Q99PJ0 | Q99PJ0 [230-257] | [K].GTGVPVGQKGTLQCEASAVPSAEFQWFK.[D] | 1xCarbamidomethyl [C14]; 1xHomocysteineIAA [K9] | 4,32935E-05 | 3153,51322 | Brain/ FAIMS/IAA |
| Q99PL5 | Q99PL5 [1189-1201] | [R].LKELESQVSCLEK.[E] | 1xCarbamidomethyl [C10]; 1xHomocysteineIAA [K2] | 4,32935E-05 | 1736,85573 | Liver/ FAIMS/IAA |
| Q99PL5 | Q99PL5 [1561-1570] | [R].AAIKLQELLK.[T] | 1xHomocysteineMMTS [K4] | 0,000192903 | 1289,73198 | Liver/ No FAIMS/MMTS |
| Q99PL5 | Q99PL5 [1010-1029] | [R].DALNQATSQVESKQNTELAK.[L] | 1xHomocysteineMMTS [K13] | 4,49774E-05 | 2338,10133 | Liver/ No FAIMS/MMTS |
| Q99PL5 | Q99PL5 [971-982] | [R].DHVKEVQQLQGK.[I] | 1xHomocysteineMMTS [K4] | 0,00750949 | 1571,76686 | Liver/ FAIMS/MMTS |
| Q99PL5 | Q99PL5 [1383-1394] | [K].LKGELESSDQVR.[E] | 1xHomocysteineMMTS [K2] | 0,000564742 | 1523,71924 | Liver/ FAIMS/MMTS |
| Q99PL5 | Q99PL5 [1062-1077] | [K].AATFEKQVLQLQASHK.[E] | 1xHomocysteineMMTS [K6] | 0,00718635 | 1961,99356 | Liver/ FAIMS/MMTS |
| Q99PL5 | Q99PL5 [1244-1254] | [R].SLTQAKEESEK.[Q] | 1xHomocysteineMMTS [K6] | 6,70238E-05 | 1412,63959 | Liver/ FAIMS/MMTS |
| Q99PL5 | Q99PL5 [1250-1264] | [K].EESEKQLHLAEAQTK.[E] | 1xHomocysteineMMTS [K5] | 0,00501021 | 1903,88882 | Liver/ FAIMS/MMTS |
| Q99PL5 | Q99PL5 [1561-1570] | [R].AAIKLQELLK.[T] | 1xHomocysteineMMTS [K4] | 0,000192903 | 1289,73198 | Liver/ FAIMS/MMTS |
| Q99PL5 | Q99PL5 [1504-1515] | [R].QKLTAEFEEAQR.[T] | 1xHomocysteineMMTS [K2] | 0,00650587 | 1612,74579 | Liver/ FAIMS/MMTS |
| Q99PL5 | Q99PL5 [1357-1368] | [K].AKVGAAEEELHK.[S] | 1xHomocysteineMMTS [K2] | 4,49774E-05 | 1444,6923 | Liver/ FAIMS/MMTS |
| Q99PL5 | Q99PL5 [1010-1029] | [R].DALNQATSQVESKQNTELAK.[L] | 1xHomocysteineMMTS [K13] | 4,49774E-05 | 2338,10133 | Liver/ FAIMS/MMTS |
| Q99PL5 | Q99PL5 [813-829] | [K].ASMVQSQEAPKQDAPAK.[K] | 1xHomocysteineMMTS [K11] | 4,49774E-05 | 1948,89253 | Liver/ FAIMS/MMTS |
| Q99PL5 | Q99PL5 [1332-1346] | [R].TILAETEGMLKDLQK.[S] | 1xHomocysteineMMTS [K11] | 0,000547722 | 1852,9217 | Liver/ FAIMS/MMTS |
| Q99PL5 | Q99PL5 [1529-1545] | [R].AAGPLESSGKEEITQLK.[E] | 1xHomocysteineMMTS [K10] | 4,49774E-05 | 1920,94052 | Liver/ FAIMS/MMTS |
| Q99PL5 | Q99PL5 [1359-1370] | [K].VGAAEEELHKSR.[V] | 1xHomocysteineMMTS [K10] | 4,49774E-05 | 1488,69336 | Liver/ FAIMS/MMTS |
| Q99PT1 | Q99PT1 [34-50] | [K].SIQEIQELDKDDESLRK.[Y] | 1xHomocysteineIAA [K10] | 4,32935E-05 | 2220,08125 | Brain/ FAIMS/IAA |
| Q99PT1 | Q99PT1 [153-172] | [R].AEEYEFLTPMEEAPKGMLAR.[G] | 1xHomocysteineMMTS [K15] | 4,49774E-05 | 2475,10629 | Liver/ FAIMS/MMTS |
| Q99PT1 | Q99PT1 [153-172] | [R].AEEYEFLTPMEEAPKGMLAR.[G] | 1xHomocysteineMMTS [K15] | 4,49774E-05 | 2475,10629 | Brain/ FAIMS/MMTS |
| Q9CPP6 | Q9CPP6 [67-86] | [K].KLEALLQGGEVEEVILQAEK.[E] | 1xHomocysteineIAA [K1] | 4,32935E-05 | 2370,25849 | Brain/ No FAIMS/IAA |
| Q9CPP6 | Q9CPP6 [67-86] | [K].KLEALLQGGEVEEVILQAEK.[E] | 1xHomocysteineIAA [K1] | 4,32935E-05 | 2370,25849 | Brain/ FAIMS/IAA |
| Q9CPQ1 | Q9CPQ1 [47-55] | [R].KKAYAEFYR.[N] | 1xHomocysteineIAA [K2] | 0,00012645 | 1349,66707 | Brain/ No FAIMS/IAA |
| Q9CPQ1 | Q9CPQ1 [47-55] | [R].KKAYAEFYR.[N] | 1xHomocysteineIAA [K2] | 0,00012645 | 1349,66707 | Brain/ FAIMS/IAA |
| Q9CPQ1 | Q9CPQ1 [56-67] | [R].NYDSMKDFEEMR.[K] | 1xHomocysteineMMTS [K6] | 0,00263388 | 1727,65321 | Brain/ FAIMS/MMTS |
| Q9CPQ8 | Q9CPQ8 [56-66] | [K].IIQSAKTGSFK.[H] | 1xHomocysteineIAA [K6] | 0,000114608 | 1353,7195 | Liver/ FAIMS/IAA |
| Q9CPQ8 | Q9CPQ8 [56-66] | [K].IIQSAKTGSFK.[H] | 1xHomocysteineIAA [K6] | 0,000114608 | 1353,7195 | Brain/ FAIMS/IAA |
| Q9CPR4 | Q9CPR4 [154-168] | [K].EQIVPKPEEEVAQKK.[K] | 1xHomocysteineIAA [K14] | 9,61721E-05 | 1926,00009 | Liver/ FAIMS/IAA |
| Q9CPR4 | Q9CPR4 [154-168] | [K].EQIVPKPEEEVAQKK.[K] | 1xHomocysteineIAA [K14] | 9,61721E-05 | 1926,00009 | Brain/ FAIMS/IAA |
| Q9CPR4 | Q9CPR4 [97-121] | [K].NAESNAELKGLDVDSLVIEHIQVNK.[A] | 1xHomocysteineMMTS [K9] | 4,49774E-05 | 2898,43357 | Liver/ FAIMS/MMTS |
| Q9CPR4 | Q9CPR4 [87-105] | [K].SAEFLLHMLKNAESNAELK.[G] | 1xHomocysteineMMTS [K10] | 0,000530332 | 2308,11342 | Liver/ FAIMS/MMTS |
| Q9CPU0 | Q9CPU0 [142-151] | [R].FEELGVKFVK.[K] | 1xHomocysteineIAA [K7] | 4,32935E-05 | 1369,71843 | Liver/ FAIMS/IAA |
| Q9CPU0 | Q9CPU0 [142-151] | [R].FEELGVKFVK.[K] | 1xHomocysteineIAA [K7] | 4,32935E-05 | 1369,71843 | Brain/ No FAIMS/IAA |
| Q9CPU0 | Q9CPU0 [142-151] | [R].FEELGVKFVK.[K] | 1xHomocysteineIAA [K7] | 4,32935E-05 | 1369,71843 | Brain/ FAIMS/IAA |
| Q9CPU0 | Q9CPU0 [141-151] | [K].RFEELGVKFVK.[K] | 1xHomocysteineMMTS [K8] | 0,00110986 | 1514,7858 | Liver/ No FAIMS/MMTS |
| Q9CPU0 | Q9CPU0 [142-151] | [R].FEELGVKFVK.[K] | 1xHomocysteineMMTS [K7] | 4,49774E-05 | 1358,68469 | Liver/ No FAIMS/MMTS |
| Q9CPU0 | Q9CPU0 [149-157] | [K].FVKKPDDGK.[M] | 1xHomocysteineMMTS [K3] | 0,00456651 | 1196,58023 | Liver/ No FAIMS/MMTS |
| Q9CPU0 | Q9CPU0 [86-95] | [K].SEKTAWTFSR.[K] | 1xHomocysteineMMTS [K3] | 0,000530332 | 1375,61332 | Liver/ No FAIMS/MMTS |
| Q9CPU0 | Q9CPU0 [44-51] | [K].KSLDFYTR.[V] | 1xHomocysteineMMTS [K1] | 0,00359453 | 1192,54893 | Liver/ No FAIMS/MMTS |
| Q9CPU0 | Q9CPU0 [142-151] | [R].FEELGVKFVK.[K] | 1xHomocysteineMMTS [K7] | 4,49774E-05 | 1358,68469 | Liver/ FAIMS/MMTS |
| Q9CPU0 | Q9CPU0 [86-95] | [K].SEKTAWTFSR.[K] | 1xHomocysteineMMTS [K3] | 0,000530332 | 1375,61332 | Liver/ FAIMS/MMTS |
| Q9CPU0 | Q9CPU0 [44-51] | [K].KSLDFYTR.[V] | 1xHomocysteineMMTS [K1] | 0,00359453 | 1192,54893 | Liver/ FAIMS/MMTS |
| Q9CPU0 | Q9CPU0 [141-151] | [K].RFEELGVKFVK.[K] | 1xHomocysteineMMTS [K8] | 0,00110986 | 1514,7858 | Liver/ FAIMS/MMTS |
| Q9CPU0 | Q9CPU0 [84-95] | [K].DKSEKTAWTFSR.[K] | 1xHomocysteineMMTS [K5] | 0,00137282 | 1618,73522 | Liver/ FAIMS/MMTS |
| Q9CPU0 | Q9CPU0 [68-85] | [K].FSLYFLAYEDKNDIPKDK.[S] | 1xHomocysteineMMTS [K16] | 0,000635569 | 2369,11922 | Liver/ FAIMS/MMTS |
| Q9CPW0 | Q9CPW0 [1183-1196] | [R].VQFNHIAPLKAALR.[Q] | 1xHomocysteineIAA [K10] | 0,000243434 | 1751,97375 | Brain/ FAIMS/IAA |
| Q9CPW2 | Q9CPW2 [75-87] | [R].GKVGDNVLYLAQR.[H] | 1xHomocysteineIAA [K2] | 0,00473945 | 1606,83699 | Brain/ FAIMS/IAA |
| Q9CPW4 | Q9CPW4 [88-100] | [K].ANDIEKAVQSLDK.[N] | 1xHomocysteineIAA [K6] | 9,61721E-05 | 1604,79485 | Brain/ FAIMS/IAA |
| Q9CPW5 | Q9CPW5 [23-32] | [R].LLASKSLLNR.[Y] | 1xHomocysteineIAA [K5] | 0,000318237 | 1288,74057 | Liver/ FAIMS/IAA |
| Q9CPW5 | Q9CPW5 [23-32] | [R].LLASKSLLNR.[Y] | 1xHomocysteineMMTS [K5] | 4,49774E-05 | 1277,70682 | Liver/ No FAIMS/MMTS |
| Q9CPW5 | Q9CPW5 [23-32] | [R].LLASKSLLNR.[Y] | 1xHomocysteineMMTS [K5] | 4,49774E-05 | 1277,70682 | Liver/ FAIMS/MMTS |
| Q9CPY7 | Q9CPY7 [105-122] | [R].SAGVDDQENWHEGKENIR.[A] | 1xHomocysteineIAA [K14] | 4,32935E-05 | 2257,98908 | Liver/ FAIMS/IAA |
| Q9CPY7 | Q9CPY7 [105-122] | [R].SAGVDDQENWHEGKENIR.[A] | 1xHomocysteineIAA [K14] | 4,32935E-05 | 2257,98908 | Brain/ FAIMS/IAA |
| Q9CPY7 | Q9CPY7 [215-224] | [R].FAEIIEKNLK.[S] | 1xHomocysteineMMTS [K7] | 4,49774E-05 | 1367,70616 | Liver/ No FAIMS/MMTS |
| Q9CPY7 | Q9CPY7 [343-351] | [K].ANKPGDVVR.[A] | 1xHomocysteineMMTS [K3] | 0,00134077 | 1118,54451 | Liver/ No FAIMS/MMTS |
| Q9CPY7 | Q9CPY7 [105-122] | [R].SAGVDDQENWHEGKENIR.[A] | 1xHomocysteineMMTS [K14] | 4,49774E-05 | 2246,95534 | Liver/ No FAIMS/MMTS |
| Q9CPY7 | Q9CPY7 [215-224] | [R].FAEIIEKNLK.[S] | 1xHomocysteineMMTS [K7] | 4,49774E-05 | 1367,70616 | Liver/ FAIMS/MMTS |
| Q9CPY7 | Q9CPY7 [343-351] | [K].ANKPGDVVR.[A] | 1xHomocysteineMMTS [K3] | 0,00134077 | 1118,54451 | Liver/ FAIMS/MMTS |
| Q9CPY7 | Q9CPY7 [354-368] | [R].NGKTIQVDNTDAEGR.[L] | 1xHomocysteineMMTS [K3] | 0,000530332 | 1780,79526 | Liver/ FAIMS/MMTS |
| Q9CPY7 | Q9CPY7 [44-61] | [K].DKDDDLPQFTSAGESFNK.[L] | 1xHomocysteineMMTS [K2] | 0,0031519 | 2176,91616 | Liver/ FAIMS/MMTS |
| Q9CPY7 | Q9CPY7 [105-122] | [R].SAGVDDQENWHEGKENIR.[A] | 1xHomocysteineMMTS [K14] | 4,49774E-05 | 2246,95534 | Liver/ FAIMS/MMTS |
| Q9CPY7 | Q9CPY7 [177-200] | [K].LHGSGDLEAWEKGVLFASGQNLAR.[H] | 1xHomocysteineMMTS [K12] | 4,49774E-05 | 2718,31266 | Liver/ FAIMS/MMTS |
| Q9CQ19; Q3THE2 | Q9CQ19 [46-63]; Q3THE2 [46-63] | [R].DGFIDKEDLHDMLASLGK.[N] | 1xHomocysteineIAA [K6] | 4,32935E-05 | 2178,02057 | Brain/ FAIMS/IAA |
| Q9CQ62 | Q9CQ62 [92-106] | [R].NIDVLKATAEEISSK.[T] | 1xHomocysteineIAA [K6] | 9,61721E-05 | 1791,91569 | Liver/ FAIMS/IAA |
| Q9CQ62 | Q9CQ62 [92-106] | [R].NIDVLKATAEEISSK.[T] | 1xHomocysteineMMTS [K6] | 4,49774E-05 | 1780,88195 | Liver/ No FAIMS/MMTS |
| Q9CQ62 | Q9CQ62 [317-330] | [K].VTKEEWDIIEGLIR.[K] | 1xHomocysteineMMTS [K3] | 0,0017273 | 1863,93432 | Liver/ No FAIMS/MMTS |
| Q9CQ62 | Q9CQ62 [215-230] | [K].SGVEAMNKSLAAEWGR.[Y] | 1xHomocysteineMMTS [K8] | 0,00150009 | 1868,84518 | Liver/ FAIMS/MMTS |
| Q9CQ62 | Q9CQ62 [92-106] | [R].NIDVLKATAEEISSK.[T] | 1xHomocysteineMMTS [K6] | 4,49774E-05 | 1780,88195 | Liver/ FAIMS/MMTS |
| Q9CQ62 | Q9CQ62 [317-330] | [K].VTKEEWDIIEGLIR.[K] | 1xHomocysteineMMTS [K3] | 0,0017273 | 1863,93432 | Liver/ FAIMS/MMTS |
| Q9CQ62 | Q9CQ62 [98-110] | [K].ATAEEISSKTGNK.[V] | 1xHomocysteineMMTS [K9] | 0,000564742 | 1498,6876 | Liver/ FAIMS/MMTS |
| Q9CQ62 | Q9CQ62 [235-246] | [R].FNIIQPGPIKTK.[G] | 1xHomocysteineMMTS [K10] | 0,00074635 | 1518,8171 | Liver/ FAIMS/MMTS |
| Q9CQ69 | Q9CQ69 [26-40] | [R].AFPSYFSKGIPNVLR.[R] | 1xHomocysteineIAA [K8] | 4,32935E-05 | 1869,968 | Brain/ FAIMS/IAA |
| Q9CQ75 | Q9CQ75 [69-89] | [R].YAFGQEKTVSLNNLSADEVTR.[A] | 1xHomocysteineIAA [K7] | 4,32935E-05 | 2516,20858 | Brain/ FAIMS/IAA |
| Q9CQ92 | Q9CQ92 [96-112] | [R].GLLQTEPQNNQAKELER.[L] | 1xHomocysteineIAA [K13] | 4,32935E-05 | 2142,06079 | Brain/ No FAIMS/IAA |
| Q9CQ92 | Q9CQ92 [96-112] | [R].GLLQTEPQNNQAKELER.[L] | 1xHomocysteineIAA [K13] | 4,32935E-05 | 2142,06079 | Brain/ FAIMS/IAA |

| | | | | | | |
|---|---|---|---|---|---|---|
| Q9CQ92 | Q9CQ92 [20-32] | [R].KFQSEQAAGSVSK.[S] | 1xHomocysteineMMTS [K1] | 0,00312202 | 1529,70867 | Liver/ FAIMS/MMTS |
| Q9CQ92 | Q9CQ92 [20-32] | [R].KFQSEQAAGSVSK.[S] | 1xHomocysteineMMTS [K1] | 0,00312202 | 1529,70867 | Brain/ FAIMS/MMTS |
| Q9CQA3 | Q9CQA3 [233-244] | [R].LAKLQDPFSVYR.[C] | 1xHomocysteineIAA [K3] | 4,32935E-05 | 1610,83592 | Liver/ FAIMS/IAA |
| Q9CQA3 | Q9CQA3 [233-244] | [R].LAKLQDPFSVYR.[C] | 1xHomocysteineIAA [K3] | 4,32935E-05 | 1610,83592 | Brain/ FAIMS/IAA |
| Q9CQA3 | Q9CQA3 [233-244] | [R].LAKLQDPFSVYR.[C] | 1xHomocysteineMMTS [K3] | 0,00487938 | 1599,80218 | Liver/ FAIMS/MMTS |
| Q9CQA3 | Q9CQA3 [119-128] | [R].IDTDLSKVSK.[I] | 1xHomocysteineMMTS [K7] | 0,00843159 | 1268,62248 | Liver/ FAIMS/MMTS |
| Q9CQA3 | Q9CQA3 [233-244] | [R].LAKLQDPFSVYR.[C] | 1xHomocysteineMMTS [K3] | 0,00487938 | 1599,80218 | Brain/ FAIMS/MMTS |
| Q9CQC7 | Q9CQC7 [115-123] | [R].LIQEGKLDR.[K] | 1xHomocysteineIAA [K6] | 0,000595249 | 1245,66198 | Brain/ FAIMS/IAA |
| Q9CQH3 | Q9CQH3 [124-135] | [R].NFYDGPEKNYEK.[T] | 1xHomocysteineIAA [K8] | 4,32935E-05 | 1677,72135 | Brain/ FAIMS/IAA |
| Q9CQH3 | Q9CQH3 [136-150] | [K].TLAILQIESEKAELR.[L] | 1xHomocysteineMMTS [K11] | 4,49774E-05 | 1876,98708 | Liver/ FAIMS/MMTS |
| Q9CQH3 | Q9CQH3 [136-150] | [K].TLAILQIESEKAELR.[L] | 1xHomocysteineMMTS [K11] | 4,49774E-05 | 1876,98708 | Brain/ FAIMS/MMTS |
| Q9CQH7; Q64152 | Q9CQH7 [6-14]; Q64152 [53-61] | [K].LAKLQAQVR.[I] | 1xHomocysteineMMTS [K3] | 0,00509539 | 1189,65439 | Liver/ FAIMS/MMTS |
| Q9CQI3 | Q9CQI3 [42-64] | [R].LVVLDEELEGVSPDELKDELPER.[Q] | 1xHomocysteineIAA [K17] | 4,32935E-05 | 2797,38118 | Brain/ No FAIMS/IAA |
| Q9CQI6 | Q9CQI6 [64-73] | [R].FTTGDAMSKR.[S] | 1xHomocysteineIAA [K9] | 9,61721E-05 | 1287,58202 | Brain/ FAIMS/IAA |
| Q9CQJ6 | Q9CQJ6 [115-122] | [K].VTIAKIPR.[A] | 1xHomocysteineIAA [K5] | 0,00114487 | 1071,63431 | Brain/ FAIMS/IAA |
| Q9CQK7 | Q9CQK7 [111-121] | [K].LNEIVDQIKTR.[R] | 1xHomocysteineIAA [K9] | 0,0014138 | 1502,79954 | Brain/ FAIMS/IAA |
| Q9CQM5 | Q9CQM5 [36-54] | [K].DTEGKSWCPDCVEAEPVIR.[E] | 2xCarbamidomethyl [C8; C11]; 1xHomocysteineIAA [K5] | 9,61721E-05 | 2422,04719 | Brain/ FAIMS/IAA |
| Q9CQQ7 | Q9CQQ7 [132-144] | [K].IAQLEEVKQSSMK.[Q] | 1xHomocysteineIAA [K8] | 4,32935E-05 | 1664,8346 | Liver/ FAIMS/IAA |
| Q9CQQ7 | Q9CQQ7 [209-221] | [R].RKEEEHMIDWVEK.[H] | 1xHomocysteineIAA [K2] | 4,32935E-05 | 1902,88368 | Brain/ FAIMS/IAA |
| Q9CQQ7 | Q9CQQ7 [155-163] | [K].AQQALVQKR.[H] | 1xHomocysteineMMTS [K8] | 0,00150009 | 1204,62891 | Liver/ No FAIMS/MMTS |
| Q9CQQ7 | Q9CQQ7 [155-163] | [K].AQQALVQKR.[H] | 1xHomocysteineMMTS [K8] | 0,00150009 | 1204,62891 | Liver/ FAIMS/MMTS |
| Q9CQQ7 | Q9CQQ7 [132-144] | [K].IAQLEEVKQSSMK.[Q] | 1xHomocysteineMMTS [K8] | 4,49774E-05 | 1653,80086 | Liver/ FAIMS/MMTS |
| Q9CQQ7 | Q9CQQ7 [127-139] | [K].LNEEKIAQLEEVK.[Q] | 1xHomocysteineMMTS [K5] | 0,00110986 | 1705,84992 | Liver/ FAIMS/MMTS |
| Q9CQQ7 | Q9CQQ7 [155-163] | [K].AQQALVQKR.[H] | 1xHomocysteineMMTS [K8] | 0,00150009 | 1204,62891 | Brain/ FAIMS/MMTS |
| Q9CQQ7 | Q9CQQ7 [132-144] | [K].IAQLEEVKQSSMK.[Q] | 1xHomocysteineMMTS [K8] | 4,49774E-05 | 1653,80086 | Brain/ FAIMS/MMTS |
| Q9CQQ7 | Q9CQQ7 [140-154] | [K].QSSMKQIQDAIDMEK.[A] | 1xHomocysteineMMTS [K5] | 0,000738471 | 1914,8428 | Brain/ FAIMS/MMTS |
| Q9CQR2 | Q9CQR2 [72-81] | [R].LAKADGIVSK.[N] | 1xHomocysteineIAA [K3] | 0,00012645 | 1175,64527 | Liver/ No FAIMS/IAA |
| Q9CQR2 | Q9CQR2 [72-81] | [R].LAKADGIVSK.[N] | 1xHomocysteineIAA [K3] | 0,00012645 | 1175,64527 | Liver/ FAIMS/IAA |
| Q9CQR2 | Q9CQR2 [72-81] | [R].LAKADGIVSK.[N] | 1xHomocysteineIAA [K3] | 0,00012645 | 1175,64527 | Brain/ FAIMS/IAA |
| Q9CQR2 | Q9CQR2 [72-81] | [R].LAKADGIVSK.[N] | 1xHomocysteineMMTS [K3] | 0,00669057 | 1164,61153 | Liver/ No FAIMS/MMTS |
| Q9CQR2 | Q9CQR2 [72-81] | [R].LAKADGIVSK.[N] | 1xHomocysteineMMTS [K3] | 0,00669057 | 1164,61153 | Liver/ FAIMS/MMTS |
| Q9CQR2 | Q9CQR2 [23-41] | [R].IIAAKDHASIQMNVAEVDR.[T] | 1xHomocysteineMMTS [K5] | 0,00180518 | 2244,09335 | Liver/ FAIMS/MMTS |
| Q9CQR4 | Q9CQR4 [38-51] | [K].LICEMKVEEQHTNK.[L] | 1xCarbamidomethyl [C3]; 1xHomocysteineIAA [K6] | 4,32935E-05 | 1932,89761 | Brain/ FAIMS/IAA |
| Q9CQV6 | Q9CQV6 [52-68] | [K].FLVPDHVNMSELIKIIR.[R] | 1xHomocysteineIAA [K14] | 4,32935E-05 | 2198,18243 | Brain/ FAIMS/IAA |
| Q9CQV8 | Q9CQV8 [63-73] | [R].VISSIEQKTER.[N] | 1xHomocysteineIAA [K8] | 4,32935E-05 | 1463,75225 | Liver/ FAIMS/IAA |
| Q9CQV8 | Q9CQV8 [141-160] | [K].QTTVSNSQQAYQEAFEISKK.[E] | 1xHomocysteineIAA [K19] | 4,32935E-05 | 2461,16638 | Brain/ No FAIMS/IAA |
| Q9CQV8 | Q9CQV8 [141-160] | [K].QTTVSNSQQAYQEAFEISKK.[E] | 1xHomocysteineIAA [K19] | 4,32935E-05 | 2461,16638 | Brain/ FAIMS/IAA |
| Q9CQV8 | Q9CQV8 [63-73] | [R].VISSIEQKTER.[N] | 1xHomocysteineMMTS [K8] | 4,49774E-05 | 1452,71851 | Liver/ No FAIMS/MMTS |
| Q9CQV8 | Q9CQV8 [141-160] | [K].QTTVSNSQQAYQEAFEISKK.[E] | 1xHomocysteineMMTS [K19] | 0,00605191 | 2450,13263 | Liver/ No FAIMS/MMTS |
| Q9CQV8 | Q9CQV8 [21-43] | [R].YDDMAAAMKAVTEQGHELSNEER.[N] | 1xHomocysteineMMTS [K9] | 4,49774E-05 | 2758,15755 | Liver/ FAIMS/MMTS |
| Q9CQV8 | Q9CQV8 [63-73] | [R].VISSIEQKTER.[N] | 1xHomocysteineMMTS [K8] | 4,49774E-05 | 1452,71851 | Liver/ FAIMS/MMTS |
| Q9CQV8 | Q9CQV8 [63-73] | [R].VISSIEQKTER.[N] | 1xHomocysteineMMTS [K8] | 4,49774E-05 | 1452,71851 | Brain/ No FAIMS/MMTS |
| Q9CQV8 | Q9CQV8 [1-11] | [-].MTMDKSELVQK.[A] | 1xMet-loss+Acetyl [N-Term]; 1xHomocysteineMMTS [K5] | 4,49774E-05 | 1383,63167 | Brain/ FAIMS/MMTS |
| Q9CQV8 | Q9CQV8 [21-43] | [R].YDDMAAAMKAVTEQGHELSNEER.[N] | 1xHomocysteineMMTS [K9] | 4,49774E-05 | 2758,15755 | Brain/ FAIMS/MMTS |
| Q9CQV8 | Q9CQV8 [63-73] | [R].VISSIEQKTER.[N] | 1xHomocysteineMMTS [K8] | 4,49774E-05 | 1452,71851 | Brain/ FAIMS/MMTS |
| Q9CQV8 | Q9CQV8 [130-159] | [R].YLSEVASGENKQTTVSNSQQAYQEAFEISK.[K] | 1xHomocysteineMMTS [K11] | 4,49774E-05 | 3499,59919 | Brain/ FAIMS/MMTS |
| Q9CQW1 | Q9CQW1 [142-163] | [K].VQAELDETKIILHNTMESLLER.[G] | 1xHomocysteineIAA [K9] | 4,32935E-05 | 2756,39573 | Brain/ FAIMS/IAA |
| Q9CQW2 | Q9CQW2 [63-77] | [K].GNVTIKIWDIGGQPR.[F] | 1xHomocysteineIAA [K6] | 0,000177335 | 1827,95341 | Brain/ FAIMS/IAA |
| Q9CQX2 | Q9CQX2 [89-103] | [K].QYYIGDVHPSDLKPK.[G] | 1xHomocysteineIAA [K13] | 4,32935E-05 | 1933,94766 | Liver/ FAIMS/IAA |
| Q9CQX2 | Q9CQX2 [89-103] | [K].QYYIGDVHPSDLKPK.[G] | 1xHomocysteineIAA [K13] | 4,32935E-05 | 1933,94766 | Brain/ FAIMS/IAA |
| Q9CQX2 | Q9CQX2 [1-24] | [-].MATPEASGSGEKVEGSEPSVTYYR.[L] | 1xMet-loss+Acetyl [N-Term]; 1xHomocysteineMMTS [K12] | 4,49774E-05 | 2606,13851 | Liver/ No FAIMS/MMTS |
| Q9CQX2 | Q9CQX2 [1-24] | [-].MATPEASGSGEKVEGSEPSVTYYR.[L] | 1xMet-loss+Acetyl [N-Term]; 1xHomocysteineMMTS [K12] | 4,49774E-05 | 2606,13851 | Liver/ FAIMS/MMTS |
| Q9CQX2 | Q9CQX2 [89-103] | [K].QYYIGDVHPSDLKPK.[G] | 1xHomocysteineMMTS [K13] | 0,0046895 | 1922,91392 | Liver/ FAIMS/MMTS |
| Q9CQX2 | Q9CQX2 [85-103] | [R].EMLKQYYIGDVHPSDLKPK.[G] | 1xHomocysteineMMTS [K4] | 4,49774E-05 | 2424,17602 | Liver/ FAIMS/MMTS |
| Q9CQZ5 | Q9CQZ5 [81-90] | [R].VVDLLVIKGK.[M] | 1xHomocysteineIAA [K8] | 0,0027602 | 1257,75991 | Brain/ FAIMS/IAA |
| Q9CQZ6 | Q9CQZ6 [30-41] | [K].IEGTPLETVQKK.[L] | 1xHomocysteineIAA [K11] | 4,32935E-05 | 1516,80395 | Brain/ No FAIMS/IAA |
| Q9CQZ6 | Q9CQZ6 [30-41] | [K].IEGTPLETVQKK.[L] | 1xHomocysteineIAA [K11] | 4,32935E-05 | 1516,80395 | Brain/ FAIMS/IAA |
| Q9CQZ6 | Q9CQZ6 [27-40] | [R].QWKIEGTPLETVQK.[K] | 1xHomocysteineMMTS [K3] | 0,00110986 | 1819,9081 | Liver/ FAIMS/MMTS |
| Q9CQZ6 | Q9CQZ6 [27-40] | [R].QWKIEGTPLETVQK.[K] | 1xHomocysteineMMTS [K3] | 0,00110986 | 1819,9081 | Brain/ FAIMS/MMTS |
| Q9CR21 | Q9CR21 [89-97] | [K].LYDKIDPEK.[L] | 1xHomocysteineIAA [K4] | 4,32935E-05 | 1294,63476 | Liver/ FAIMS/IAA |
| Q9CR21 | Q9CR21 [138-152] | [K].LMCPQEIVDYIADKK.[D] | 1xCarbamidomethyl [C3]; 1xHomocysteineIAA [K14] | 4,32935E-05 | 1996,95407 | Brain/ No FAIMS/IAA |
| Q9CR21 | Q9CR21 [138-152] | [K].LMCPQEIVDYIADKK.[D] | 1xCarbamidomethyl [C3]; 1xHomocysteineIAA [K14] | 4,32935E-05 | 1996,95407 | Brain/ FAIMS/IAA |
| Q9CR21 | Q9CR21 [83-92] | [R].VLYVLKLYDK.[I] | 1xHomocysteineMMTS [K6] | 0,000152766 | 1416,76294 | Liver/ FAIMS/MMTS |
| Q9CR21 | Q9CR21 [83-92] | [R].VLYVLKLYDK.[I] | 1xHomocysteineMMTS [K6] | 0,000152766 | 1416,76294 | Brain/ No FAIMS/MMTS |
| Q9CR21 | Q9CR21 [83-92] | [R].VLYVLKLYDK.[I] | 1xHomocysteineMMTS [K6] | 0,000152766 | 1416,76294 | Brain/ FAIMS/MMTS |
| Q9CR51 | Q9CR51 [59-78] | [K].EAAALGSHGSCSSEVEKETR.[E] | 1xCarbamidomethyl [C11]; 1xHomocysteineIAA [K17] | 4,32935E-05 | 2279,00268 | Brain/ No FAIMS/IAA |
| Q9CR51 | Q9CR51 [59-78] | [K].EAAALGSHGSCSSEVEKETR.[E] | 1xCarbamidomethyl [C11]; 1xHomocysteineIAA [K17] | 4,32935E-05 | 2279,00268 | Brain/ FAIMS/IAA |
| Q9CR51 | Q9CR51 [35-48] | [K].QAKEEAQAEIEQYR.[L] | 1xHomocysteineMMTS [K3] | 0,00343014 | 1855,83131 | Liver/ FAIMS/MMTS |
| Q9CR51 | Q9CR51 [35-48] | [K].QAKEEAQAEIEQYR.[L] | 1xHomocysteineMMTS [K3] | 0,00343014 | 1855,83131 | Brain/ FAIMS/MMTS |

| | | | | | | |
|---|---|---|---|---|---|---|
| Q9CR57 | Q9CR57 [148-162] | [K].AAIAAAAAAAAAKAK.[V] | 1xHomocysteineMMTS [K13] | 4,49774E-05 | 1403,74975 | Liver/ No FAIMS/MMTS |
| Q9CR57 | Q9CR57 [148-162] | [K].AAIAAAAAAAAAKAK.[V] | 1xHomocysteineMMTS [K13] | 4,49774E-05 | 1403,74975 | Liver/ FAIMS/MMTS |
| Q9CR62 | Q9CR62 [49-59] | [R].MQLSGEGAKTR.[E] | 1xHomocysteineIAA [K9] | 9,61721E-05 | 1351,64568 | Brain/ FAIMS/IAA |
| Q9CR68 | Q9CR68 [251-274] | [R].KGPAPLNLEVPAYEFTSDDVVVVG.[-] | 1xHomocysteineIAA [K1] | 4,32935E-05 | 2690,33819 | Brain/ No FAIMS/IAA |
| Q9CR68 | Q9CR68 [251-274] | [R].KGPAPLNLEVPAYEFTSDDVVVVG.[-] | 1xHomocysteineIAA [K1] | 4,32935E-05 | 2690,33819 | Brain/ FAIMS/IAA |
| Q9CR68 | Q9CR68 [94-104] | [R].AEVLDSTKSSK.[E] | 1xHomocysteineMMTS [K8] | 0,000437433 | 1327,62321 | Liver/ No FAIMS/MMTS |
| Q9CR68 | Q9CR68 [93-104] | [R].RAEVLDSTKSSK.[E] | 1xHomocysteineMMTS [K9] | 0,0004709 | 1483,72432 | Liver/ FAIMS/MMTS |
| Q9CR68 | Q9CR68 [156-168] | [K].LSDIPEGKNMAFK.[W] | 1xHomocysteineMMTS [K8] | 4,49774E-05 | 1612,75318 | Liver/ FAIMS/MMTS |
| Q9CR68 | Q9CR68 [94-104] | [R].AEVLDSTKSSK.[E] | 1xHomocysteineMMTS [K8] | 0,000437433 | 1327,62321 | Liver/ FAIMS/MMTS |
| Q9CR68 | Q9CR68 [152-163] | [K].IEIKLSDIPEGK.[N] | 1xHomocysteineMMTS [K4] | 0,000192903 | 1504,77496 | Liver/ FAIMS/MMTS |
| Q9CR68 | Q9CR68 [94-104] | [R].AEVLDSTKSSK.[E] | 1xHomocysteineMMTS [K8] | 0,000437433 | 1327,62321 | Brain/ No FAIMS/MMTS |
| Q9CR68 | Q9CR68 [94-104] | [R].AEVLDSTKSSK.[E] | 1xHomocysteineMMTS [K8] | 0,000437433 | 1327,62321 | Brain/ FAIMS/MMTS |
| Q9CR95 | Q9CR95 [149-160] | [K].LDLGFKEGQTIK.[L] | 1xHomocysteineIAA [K6] | 9,61721E-05 | 1522,79339 | Brain/ FAIMS/IAA |
| Q9CRB6 | Q9CRB6 [49-63] | [K].AVTGTDVDIVFSKVK.[A] | 1xHomocysteineIAA [K13] | 4,32935E-05 | 1752,92005 | Brain/ No FAIMS/IAA |
| Q9CRB6 | Q9CRB6 [49-63] | [K].AVTGTDVDIVFSKVK.[A] | 1xHomocysteineIAA [K13] | 4,32935E-05 | 1752,92005 | Brain/ FAIMS/IAA |
| Q9CRB9 | Q9CRB9 [49-64] | [R].YSSVYGASVSDEDLKR.[R] | 1xHomocysteineIAA [K15] | 4,32935E-05 | 1949,89093 | Brain/ FAIMS/IAA |
| Q9CRC3 | Q9CRC3 [17-33] | [K].EPETAPPAAGPVATDPK.[G] | 1xHomocysteineIAA [K17] | 0,00709499 | 1821,86874 | Brain/ No FAIMS/IAA |
| Q9CVB6 | Q9CVB6 [180-189] | [K].VFMQEFKEGR.[R] | 1xHomocysteineIAA [K7] | 6,30912E-05 | 1444,67117 | Brain/ FAIMS/IAA |
| Q9CVB6 | Q9CVB6 [158-170] | [K].KDRVTVVFSTVFK.[D] | 1xHomocysteineMMTS [K1] | 0,00805184 | 1688,88624 | Liver/ FAIMS/MMTS |
| Q9CWF2 | Q9CWF2 [47-62] | [R].INVYYNEATGNKYVPR.[A] | 1xHomocysteineIAA [K12] | 4,32935E-05 | 2075,00149 | Brain/ No FAIMS/IAA |
| Q9CWF2 | Q9CWF2 [47-62] | [R].INVYYNEATGNKYVPR.[A] | 1xHomocysteineIAA [K12] | 4,32935E-05 | 2075,00149 | Brain/ FAIMS/IAA |
| Q9CWH1 | Q9CWH1 [428-434] | [K].EIKPNIR.[-] | 1xHomocysteineMMTS [K3] | 0,00524294 | 1032,53288 | Liver/ No FAIMS/MMTS |
| Q9CWH1 | Q9CWH1 [428-434] | [K].EIKPNIR.[-] | 1xHomocysteineMMTS [K3] | 0,00524294 | 1032,53288 | Liver/ FAIMS/MMTS |
| Q9CWJ9 | Q9CWJ9 [65-79] | [R].VKTLHPAVHAGILAR.[N] | 1xHomocysteineIAA [K2] | 4,32935E-05 | 1757,0003 | Brain/ FAIMS/IAA |
| Q9CWK8 | Q9CWK8 [113-126] | [R].IESKSISAPVIFDR.[S] | 1xHomocysteineIAA [K4] | 0,000177335 | 1735,90473 | Brain/ FAIMS/IAA |
| Q9CWS0 | Q9CWS0 [252-267] | [K].LKDHLLIPVSNSEMEK.[V] | 1xHomocysteineIAA [K2] | 4,32935E-05 | 2027,03001 | Brain/ FAIMS/IAA |
| Q9CWZ7 | Q9CWZ7 [152-162] | [R].QAVELLGKASR.[L] | 1xHomocysteineIAA [K8] | 4,32935E-05 | 1345,72564 | Brain/ FAIMS/IAA |
| Q9CXW3 | Q9CXW3 [164-175] | [R].WDYLTQVEKECK.[E] | 1xCarbamidomethyl [C11]; 1xHomocysteineIAA [K9] | 9,61721E-05 | 1772,79822 | Brain/ FAIMS/IAA |
| Q9CXW4 | Q9CXW4 [36-52] | [R].AAKVLEQLTGQTPVFSK.[A] | 1xHomocysteineIAA [K3] | 4,32935E-05 | 1991,06302 | Liver/ FAIMS/IAA |
| Q9CXW4 | Q9CXW4 [36-52] | [R].AAKVLEQLTGQTPVFSK.[A] | 1xHomocysteineIAA [K3] | 4,32935E-05 | 1991,06302 | Brain/ FAIMS/IAA |
| Q9CXW4 | Q9CXW4 [36-52] | [R].AAKVLEQLTGQTPVFSK.[A] | 1xHomocysteineMMTS [K3] | 4,49774E-05 | 1980,02928 | Liver/ FAIMS/MMTS |
| Q9CXW4 | Q9CXW4 [39-54] | [K].VLEQLTGQTPVFSKAR.[Y] | 1xHomocysteineMMTS [K14] | 4,49774E-05 | 1936,99831 | Liver/ FAIMS/MMTS |
| Q9CXW4 | Q9CXW4 [79-88] | [K].AEEILEKGLK.[V] | 1xHomocysteineMMTS [K7] | 0,00531384 | 1292,65887 | Liver/ FAIMS/MMTS |
| Q9CYZ2 | Q9CYZ2 [81-95] | [R].LGLSTLGELKQNLSR.[S] | 1xHomocysteineIAA [K10] | 4,32935E-05 | 1802,97929 | Brain/ No FAIMS/IAA |
| Q9CYZ2 | Q9CYZ2 [81-95] | [R].LGLSTLGELKQNLSR.[S] | 1xHomocysteineIAA [K10] | 4,32935E-05 | 1802,97929 | Brain/ FAIMS/IAA |
| Q9CYZ2 | Q9CYZ2 [81-95] | [R].LGLSTLGELKQNLSR.[S] | 1xHomocysteineMMTS [K10] | 0,00434837 | 1791,94555 | Liver/ FAIMS/MMTS |
| Q9CZ13 | Q9CZ13 [143-165] | [K].VVELLADIVQNSSLEDSQIEKER.[D] | 1xHomocysteineIAA [K21] | 4,32935E-05 | 2788,40331 | Liver/ FAIMS/IAA |
| Q9CZ13 | Q9CZ13 [103-126] | [K].NRPGNALEKEVESIGAHLNAYSTR.[E] | 1xHomocysteineIAA [K9] | 4,32935E-05 | 2800,3795 | Brain/ FAIMS/IAA |
| Q9CZ13 | Q9CZ13 [103-126] | [K].NRPGNALEKEVESIGAHLNAYSTR.[E] | 1xHomocysteineMMTS [K9] | 4,49774E-05 | 2789,34576 | Liver/ FAIMS/MMTS |
| Q9CZ13 | Q9CZ13 [143-165] | [K].VVELLADIVQNSSLEDSQIEKER.[D] | 1xHomocysteineMMTS [K21] | 0,000152766 | 2777,36957 | Liver/ FAIMS/MMTS |
| Q9CZ44 | Q9CZ44 [112-124] | [R].KKSPNELVDDLFK.[G] | 1xHomocysteineIAA [K2] | 0,00306243 | 1706,87818 | Brain/ FAIMS/IAA |
| Q9CZC8 | Q9CZC8 [303-315] | [R].SIFKPFIFVDDVK.[L] | 1xHomocysteineIAA [K4] | 4,32935E-05 | 1728,90294 | Brain/ FAIMS/IAA |
| Q9CZM2 | Q9CZM2 [78-93] | [K].GATYGKPVHHGVNQLK.[F] | 1xHomocysteineMMTS [K6] | 4,49774E-05 | 1868,92582 | Liver/ FAIMS/MMTS |
| Q9CZM2 | Q9CZM2 [55-63] | [K].AKQGYVIYR.[I] | 1xHomocysteineMMTS [K2] | 0,00482082 | 1260,62276 | Liver/ FAIMS/MMTS |
| Q9CZT8 | Q9CZT8 [9-24] | [K].TGIKDASDQNFDYMFK.[L] | 1xHomocysteineIAA [K4] | 0,0020894 | 2053,89939 | Brain/ FAIMS/IAA |
| Q9CZU6 | Q9CZU6 [74-92] | [R].GMKGLVYETSVLDPDEGIR.[F] | 1xHomocysteineIAA [K3] | 4,32935E-05 | 2253,08898 | Brain/ No FAIMS/IAA |
| Q9CZU6 | Q9CZU6 [74-92] | [R].GMKGLVYETSVLDPDEGIR.[F] | 1xHomocysteineIAA [K3] | 4,32935E-05 | 2253,08898 | Brain/ FAIMS/IAA |
| Q9CZU6 | Q9CZU6 [74-92] | [R].GMKGLVYETSVLDPDEGIR.[F] | 1xHomocysteineMMTS [K3] | 4,49774E-05 | 2242,05524 | Brain/ FAIMS/MMTS |
| Q9CZX8 | Q9CZX8 [42-56] | [K].HKELAPYDENWFYTR.[A] | 1xHomocysteineIAA [K2] | 4,32935E-05 | 2142,97018 | Brain/ FAIMS/IAA |
| Q9CZX8 | Q9CZX8 [1-16] | [-].MPGVTVKDVNQQEFVR.[A] | 1xMet-loss [N-Term]; 1xHomocysteineMMTS [K7] | 4,49774E-05 | 1878,92006 | Liver/ FAIMS/MMTS |
| Q9CZX8 | Q9CZX8 [30-41] | [K].VPEWVDTVKLAK.[H] | 1xHomocysteineMMTS [K9] | 0,000266738 | 1547,79603 | Liver/ FAIMS/MMTS |
| Q9CZX8 | Q9CZX8 [42-56] | [K].HKELAPYDENWFYTR.[A] | 1xHomocysteineMMTS [K2] | 4,49774E-05 | 2131,93644 | Liver/ FAIMS/MMTS |
| Q9CZX8 | Q9CZX8 [28-38] | [K].LKVPEWVDTVK.[L] | 1xHomocysteineMMTS [K2] | 0,000530332 | 1476,75892 | Liver/ FAIMS/MMTS |
| Q9CZX8 | Q9CZX8 [134-144] | [R].IAGQVAAANKK.[H] | 1xHomocysteineMMTS [K10] | 0,00074635 | 1233,64422 | Liver/ FAIMS/MMTS |
| Q9CZY3 | Q9CZY3 [64-74] | [R].IYSLKIECGPK.[Y] | 1xCarbamidomethyl [C8]; 1xHomocysteineIAA [K5] | 9,61721E-05 | 1481,74908 | Brain/ FAIMS/IAA |
| Q9D051 | Q9D051 [53-71] | [K].VFLLGEEVAQYDGAYKVSR.[G] | 1xHomocysteineIAA [K16] | 4,32935E-05 | 2318,14854 | Brain/ FAIMS/IAA |
| Q9D051 | Q9D051 [337-354] | [K].VLEDNSVPQVKDIIFAVK.[K] | 1xHomocysteineMMTS [K11] | 0,000738471 | 2177,13447 | Liver/ FAIMS/MMTS |
| Q9D051 | Q9D051 [337-354] | [K].VLEDNSVPQVKDIIFAVK.[K] | 1xHomocysteineMMTS [K11] | 0,000738471 | 2177,13447 | Brain/ FAIMS/MMTS |
| Q9D0E1 | Q9D0E1 [371-381] | [R].INEILSNALKR.[G] | 1xHomocysteineIAA [K10] | 0,00247317 | 1444,79406 | Brain/ FAIMS/IAA |
| Q9D0F3 | Q9D0F3 [349-358] | [R].IHLEIKQLNR.[Q] | 1xHomocysteineIAA [K6] | 4,32935E-05 | 1437,79948 | Liver/ FAIMS/IAA |
| Q9D0F3 | Q9D0F3 [349-358] | [R].IHLEIKQLNR.[Q] | 1xHomocysteineMMTS [K6] | 0,00586177 | 1426,76574 | Liver/ No FAIMS/MMTS |
| Q9D0F3 | Q9D0F3 [349-358] | [R].IHLEIKQLNR.[Q] | 1xHomocysteineMMTS [K6] | 0,00586177 | 1426,76574 | Liver/ FAIMS/MMTS |
| Q9D0J8 | Q9D0J8 [81-92] | [R].TAEEEDEADPKR.[Q] | 1xHomocysteineIAA [K11] | 4,32935E-05 | 1563,65914 | Liver/ No FAIMS/IAA |
| Q9D0J8 | Q9D0J8 [81-92] | [R].TAEEEDEADPKR.[Q] | 1xHomocysteineIAA [K11] | 4,32935E-05 | 1563,65914 | Liver/ FAIMS/IAA |
| Q9D0J8 | Q9D0J8 [81-92] | [R].TAEEEDEADPKR.[Q] | 1xHomocysteineIAA [K11] | 4,32935E-05 | 1563,65914 | Brain/ No FAIMS/IAA |
| Q9D0J8 | Q9D0J8 [81-92] | [R].TAEEEDEADPKR.[Q] | 1xHomocysteineIAA [K11] | 4,32935E-05 | 1563,65914 | Brain/ FAIMS/IAA |
| Q9D0J8 | Q9D0J8 [81-92] | [R].TAEEEDEADPKR.[Q] | 1xHomocysteineMMTS [K11] | 6,70238E-05 | 1552,6254 | Liver/ No FAIMS/MMTS |
| Q9D0J8 | Q9D0J8 [22-30] | [K].DKVEEKAGR.[K] | 1xHomocysteineMMTS [K] | 0,00718635 | 1194,56055 | Liver/ No FAIMS/MMTS |
| Q9D0J8 | Q9D0J8 [1-15] | [-].MSEKSVEAAAELSAK.[D] | 1xMet-loss+Acetyl [N-Term]; 1xHomocysteineMMTS [K4] | 4,49774E-05 | 1624,75568 | Liver/ FAIMS/MMTS |
| Q9D0J8 | Q9D0J8 [81-92] | [R].TAEEEDEADPKR.[Q] | 1xHomocysteineMMTS [K11] | 6,70238E-05 | 1552,6254 | Liver/ FAIMS/MMTS |
| Q9D0J8 | Q9D0J8 [5-18] | [K].SVEAAAELSAKDLK.[E] | 1xHomocysteineMMTS [K11] | 4,49774E-05 | 1594,7815 | Liver/ FAIMS/MMTS |
| Q9D0J8 | Q9D0J8 [1-15] | [-].MSEKSVEAAAELSAK.[D] | 1xMet-loss+Acetyl [N-Term]; 1xHomocysteineMMTS [K4] | 4,49774E-05 | 1624,75568 | Brain/ No FAIMS/MMTS |
| Q9D0J8 | Q9D0J8 [5-18] | [K].SVEAAAELSAKDLK.[E] | 1xHomocysteineMMTS [K11] | 4,49774E-05 | 1594,7815 | Brain/ No FAIMS/MMTS |
| Q9D0J8 | Q9D0J8 [22-30] | [K].DKVEEKAGR.[K] | 1xHomocysteineMMTS [K] | 0,00718635 | 1194,56055 | Brain/ No FAIMS/MMTS |
| Q9D0J8 | Q9D0J8 [1-15] | [-].MSEKSVEAAAELSAK.[D] | 1xMet-loss+Acetyl [N-Term]; 1xHomocysteineMMTS [K4] | 4,49774E-05 | 1624,75568 | Brain/ FAIMS/MMTS |
| Q9D0J8 | Q9D0J8 [81-92] | [R].TAEEEDEADPKR.[Q] | 1xHomocysteineMMTS [K11] | 6,70238E-05 | 1552,6254 | Brain/ FAIMS/MMTS |

| | | | | | | |
|---|---|---|---|---|---|---|
| Q9D0J8 | Q9D0J8 [5-18] | [K].SVEAAAELSAKDLK.[E] | 1xHomocysteineMMTS [K11] | 4,49774E-05 | 1594,7815 | Brain/ FAIMS/MMTS |
| Q9D0K2 | Q9D0K2 [419-434] | [K].MVKGMGGAMDLVSSSK.[T] | 1xHomocysteineIAA [K3] | 9,61721E-05 | 1771,82094 | Brain/ FAIMS/IAA |
| Q9D0M3 | Q9D0M3 [171-186] | [K].LSDYFPKPYPNPEAAR.[A] | 1xHomocysteineIAA [K7] | 4,32935E-05 | 2038,96912 | Brain/ FAIMS/IAA |
| Q9D0M5 | Q9D0M5 [5-31] | [R].KAVIKNADMSEDMQQDAVDCATQAMEK.[Y] | 1xCarbamidomethyl [C20]; 1xHomocysteineIAA [K5] | 4,32935E-05 | 3201,39892 | Brain/ FAIMS/IAA |
| Q9D0M5; P63168 | Q9D0M5 [49-60]; P63168 [49-60] | [K].KYNPTWHCIVGR.[N] | 1xCarbamidomethyl [C8]; 1xHomocysteineIAA [K1] | 4,32935E-05 | 1704,80973 | Brain/ No FAIMS/IAA |
| Q9D0M5; P63168 | Q9D0M5 [49-60]; P63168 [49-60] | [K].KYNPTWHCIVGR.[N] | 1xCarbamidomethyl [C8]; 1xHomocysteineIAA [K1] | 4,32935E-05 | 1704,80973 | Brain/ FAIMS/IAA |
| Q9D0S9 | Q9D0S9 [43-58] | [K].AQKAAPGGASPTIFSR.[I] | 1xHomocysteineIAA [K3] | 4,32935E-05 | 1732,87991 | Liver/ FAIMS/IAA |
| Q9D0S9 | Q9D0S9 [43-58] | [K].AQKAAPGGASPTIFSR.[I] | 1xHomocysteineIAA [K3] | 4,32935E-05 | 1732,87991 | Brain/ FAIMS/IAA |
| Q9D0S9 | Q9D0S9 [100-120] | [R].ISQAEEDDQQLLGHLLLVAKK.[I] | 1xHomocysteineMMTS [K20] | 4,49774E-05 | 2511,29456 | Liver/ No FAIMS/MMTS |
| Q9D0S9 | Q9D0S9 [100-120] | [R].ISQAEEDDQQLLGHLLLVAKK.[I] | 1xHomocysteineMMTS [K20] | 4,49774E-05 | 2511,29456 | Liver/ FAIMS/MMTS |
| Q9D0S9 | Q9D0S9 [121-132] | [K].IAQAQGLKDGYR.[L] | 1xHomocysteineMMTS [K8] | 4,49774E-05 | 1482,71918 | Liver/ FAIMS/MMTS |
| Q9D0W5 | Q9D0W5 [56-72] | [R].IIKDFMIQGGDPTGTGR.[G] | 1xHomocysteineIAA [K3] | 0,000595249 | 1979,96774 | Brain/ FAIMS/IAA |
| Q9D164 | Q9D164 [70-94] | [R].APGDEEAQVENLITTNAAEPQKAEN.[-] | 1xHomocysteineIAA [K22] | 4,32935E-05 | 2813,28941 | Brain/ FAIMS/IAA |
| Q9D172 | Q9D172 [187-212] | [K].GVEVTVGHEQEEGGKWPYAGTAEAIK.[A] | 1xHomocysteineIAA [K15] | 4,32935E-05 | 2916,38325 | Brain/ FAIMS/IAA |
| Q9D1A2 | Q9D1A2 [400-413] | [R].ALKTVFGVEPDLTR.[E] | 1xHomocysteineIAA [K3] | 0,000177335 | 1719,90982 | Brain/ FAIMS/IAA |
| Q9D1C8 | Q9D1C8 [1-25] | [-].MFHGIPATPGVGAPGNKPELYEEVK.[L] | 1xMet-loss [N-Term]; 1xHomocysteineIAA [K17] | 0,0008162 | 2681,3392 | Liver/ FAIMS/IAA |
| Q9D1E6 | Q9D1E6 [149-164] | [R].LSEEKAQASAISVGSR.[C] | 1xHomocysteineIAA [K5] | 8,35381E-05 | 1806,90144 | Brain/ FAIMS/IAA |
| Q9D1G1 | Q9D1G1 [154-171] | [K].NATNVEQAFMTMAAEIKK.[R] | 1xHomocysteineIAA [K17] | 9,61721E-05 | 2171,02936 | Brain/ FAIMS/IAA |
| Q9D1K2 | Q9D1K2 [33-53] | [R].HPNFLVVEKDTTINEIEDTFR.[Q] | 1xHomocysteineIAA [K9] | 4,32935E-05 | 2691,30829 | Brain/ FAIMS/IAA |
| Q9D1K2 | Q9D1K2 [33-53] | [R].HPNFLVVEKDTTINEIEDTFR.[Q] | 1xHomocysteineMMTS [K9] | 0,000599431 | 2680,27455 | Brain/ FAIMS/MMTS |
| Q9D1K2 | Q9D1K2 [95-107] | [K].EHPYDAAKDSILR.[R] | 1xHomocysteineMMTS [K8] | 0,0025135 | 1677,77234 | Brain/ FAIMS/MMTS |
| Q9D1K2 | Q9D1K2 [109-119] | [R].AKGMFTAEDLR.[-] | 1xHomocysteineMMTS [K2] | 0,00452171 | 1401,63234 | Brain/ FAIMS/MMTS |
| Q9D1P4 | Q9D1P4 [47-63] | [R].TTDFSDFLSIVGCTKGR.[H] | 1xCarbamidomethyl [C13]; 1xHomocysteineIAA [K15] | 0,00263986 | 2077,96814 | Brain/ FAIMS/IAA |
| Q9D1R9 | Q9D1R9 [77-88] | [R].AYGGSMCAKCVR.[D] | 2xCarbamidomethyl [C7; C10]; 1xHomocysteineIAA [K9] | 0,000960518 | 1533,64292 | Brain/ FAIMS/IAA |
| Q9D1R9 | Q9D1R9 [11-21] | [R].LSYNTASNKTR.[L] | 1xHomocysteineMMTS [K9] | 0,00127368 | 1417,65624 | Liver/ FAIMS/MMTS |
| Q9D1R9 | Q9D1R9 [94-105] | [R].AFLIEEQKIVVK.[V] | 1xHomocysteineMMTS [K8] | 4,49774E-05 | 1579,85863 | Liver/ FAIMS/MMTS |
| Q9D2G2 | Q9D2G2 [347-356] | [R].TINELGEKAR.[K] | 1xHomocysteineIAA [K8] | 9,61721E-05 | 1304,66271 | Brain/ FAIMS/IAA |
| Q9D2G2 | Q9D2G2 [347-356] | [R].TINELGEKAR.[K] | 1xHomocysteineMMTS [K8] | 0,000361246 | 1293,62897 | Liver/ FAIMS/MMTS |
| Q9D2G2 | Q9D2G2 [347-356] | [R].TINELGEKAR.[K] | 1xHomocysteineMMTS [K8] | 0,000361246 | 1293,62897 | Brain/ FAIMS/MMTS |
| Q9D2M8 | Q9D2M8 [62-72] | [R].IYSLKVECGSK.[Y] | 1xCarbamidomethyl [C8]; 1xHomocysteineIAA [K5] | 4,32935E-05 | 1457,7127 | Brain/ FAIMS/IAA |
| Q9D2P8 | Q9D2P8 [98-106] | [K].SPLMPAKPR.[S] | 1xHomocysteineIAA [K7] | 0,000595249 | 1170,6122 | Brain/ FAIMS/IAA |
| Q9D312 | Q9D312 [253-265] | [R].YEVLAQKNLQEAK.[E] | 1xHomocysteineIAA [K7] | 0,00371384 | 1707,87343 | Brain/ No FAIMS/IAA |
| Q9D394 | Q9D394 [379-387] | [K].MLEKDVCEK.[Q] | 1xCarbamidomethyl [C7]; 1xHomocysteineIAA [K4] | 0,000584831 | 1325,58981 | Brain/ FAIMS/IAA |
| Q9D3D9 | Q9D3D9 [157-168] | [R].IEANEALVKALE.[-] | 1xHomocysteineIAA [K9] | 4,32935E-05 | 1473,76176 | Liver/ FAIMS/IAA |
| Q9D3D9 | Q9D3D9 [157-168] | [R].IEANEALVKALE.[-] | 1xHomocysteineIAA [K9] | 4,32935E-05 | 1473,76176 | Brain/ No FAIMS/IAA |
| Q9D3D9 | Q9D3D9 [157-168] | [R].IEANEALVKALE.[-] | 1xHomocysteineIAA [K9] | 4,32935E-05 | 1473,76176 | Brain/ FAIMS/IAA |
| Q9D3D9 | Q9D3D9 [132-150] | [R].ANLEKAQSELSGAADEAAR.[A] | 1xHomocysteineMMTS [K5] | 4,49774E-05 | 2093,95903 | Liver/ No FAIMS/MMTS |
| Q9D3D9 | Q9D3D9 [157-168] | [R].IEANEALVKALE.[-] | 1xHomocysteineMMTS [K9] | 4,49774E-05 | 1462,72801 | Liver/ FAIMS/MMTS |
| Q9D3D9 | Q9D3D9 [132-150] | [R].ANLEKAQSELSGAADEAAR.[A] | 1xHomocysteineMMTS [K5] | 4,49774E-05 | 2093,95903 | Liver/ FAIMS/MMTS |
| Q9D3D9 | Q9D3D9 [157-168] | [R].IEANEALVKALE.[-] | 1xHomocysteineMMTS [K9] | 4,49774E-05 | 1462,72801 | Brain/ No FAIMS/MMTS |
| Q9D3D9 | Q9D3D9 [157-168] | [R].IEANEALVKALE.[-] | 1xHomocysteineMMTS [K9] | 4,49774E-05 | 1462,72801 | Brain/ FAIMS/MMTS |
| Q9D3D9 | Q9D3D9 [132-150] | [R].ANLEKAQSELSGAADEAAR.[A] | 1xHomocysteineMMTS [K5] | 4,49774E-05 | 2093,95903 | Brain/ FAIMS/MMTS |
| Q9D4B2 | Q9D4B2 [197-210] | [K].EYLEKLLLDEDLIK.[G] | 2xHomocysteineIAA [K5; K14] | 0,00727166 | 2082,04974 | Brain/ FAIMS/IAA |
| Q9D4D4; P40142 | Q9D4D4 [143-150]; P40142 [141-148] | [K].YFDKASYR.[V] | 1xHomocysteineIAA [K4] | 6,30912E-05 | 1223,55137 | Brain/ No FAIMS/IAA |
| Q9D4D4; P40142 | Q9D4D4 [143-150]; P40142 [141-148] | [K].YFDKASYR.[V] | 1xHomocysteineIAA [K4] | 6,30912E-05 | 1223,55137 | Brain/ FAIMS/IAA |
| Q9D4J1; Q9D8Y0 | Q9D4J1 [188-196]; Q9D8Y0 [189-197] | [RK].GAKNFFEAK.[AV] | 1xHomocysteineIAA [K3] | 0,00626664 | 1185,5721 | Brain/ No FAIMS/IAA |
| Q9D4J1; Q9D8Y0 | Q9D4J1 [188-196]; Q9D8Y0 [189-197] | [RK].GAKNFFEAK.[AV] | 1xHomocysteineIAA [K3] | 0,00626664 | 1185,5721 | Brain/ FAIMS/IAA |
| Q9D6F9 | Q9D6F9 [360-379] | [R].GLKMAATFIGNSTAIQELFK.[R] | 1xHomocysteineIAA [K3] | 4,32935E-05 | 2314,19338 | Brain/ No FAIMS/IAA |
| Q9D6F9 | Q9D6F9 [360-379] | [R].GLKMAATFIGNSTAIQELFK.[R] | 1xHomocysteineIAA [K3] | 4,32935E-05 | 2314,19338 | Brain/ FAIMS/IAA |
| Q9D6F9 | Q9D6F9 [360-379] | [R].GLKMAATFIGNSTAIQELFK.[R] | 1xOxidation [M4]; 1xHomocysteineMMTS [K3] | 0,00148443 | 2319,15456 | Brain/ FAIMS/MMTS |
| Q9D6F9 | Q9D6F9 [321-336] | [R].MSMKEVDEQMLSVQSK.[N] | 1xHomocysteineMMTS [K4] | 0,00545792 | 2032,88804 | Brain/ FAIMS/MMTS |
| Q9D6F9; P99024; P68372 | Q9D6F9 [122-156]; P99024 [122-156]; P68372 [122-156] | [R].KEAESCDCLQGFQLTHSLGGGTGSGMGTLLISKIR.[E] | 2xCarbamidomethyl [C6; C8]; 1xHomocysteineIAA [K33] | 4,32935E-05 | 3882,86052 | Brain/ FAIMS/IAA |
| Q9D6F9; Q9ERD7; P68372; Q922F4 | Q9D6F9 [283-306]; Q9ERD7 [283-306]; P68372 [283-306]; Q922F4 [283-306] | [R].ALTVPELTQQMFDAKNMMAACDPR.[H] | 1xCarbamidomethyl [C21]; 1xHomocysteineIAA [K15] | 4,32935E-05 | 2912,32317 | Brain/ No FAIMS/IAA |
| Q9D6F9; Q9ERD7; P68372; Q922F4 | Q9D6F9 [283-306]; Q9ERD7 [283-306]; P68372 [283-306]; Q922F4 [283-306] | [R].ALTVPELTQQMFDAKNMMAACDPR.[H] | 1xCarbamidomethyl [C21]; 1xHomocysteineIAA [K15] | 4,32935E-05 | 2912,32317 | Brain/ FAIMS/IAA |
| Q9D6J5 | Q9D6J5 [171-186] | [R].GGDPTKEPEPVVHYDI.[-] | 1xHomocysteineIAA [K6] | 4,32935E-05 | 1926,8902 | Liver/ FAIMS/IAA |
| Q9D6J5 | Q9D6J5 [171-186] | [R].GGDPTKEPEPVVHYDI.[-] | 1xHomocysteineIAA [K6] | 4,32935E-05 | 1926,8902 | Brain/ FAIMS/IAA |
| Q9D6J6 | Q9D6J6 [42-61] | [R].DTPENNPDTPFDFTPENYKR.[I] | 1xHomocysteineIAA [K19] | 4,32935E-05 | 2571,10926 | Brain/ FAIMS/IAA |
| Q9D6J6 | Q9D6J6 [42-61] | [R].DTPENNPDTPFDFTPENYKR.[I] | 1xHomocysteineMMTS [K19] | 4,49774E-05 | 2560,07551 | Brain/ FAIMS/MMTS |
| Q9D6K5 | Q9D6K5 [47-58] | [R].IKEDGAAAQDGR.[L] | 1xHomocysteineMMTS [K2] | 0,00854191 | 1393,61986 | Liver/ FAIMS/MMTS |

| | | | | | | |
|---|---|---|---|---|---|---|
| Q9D6M3 | Q9D6M3 [45-58] | [R].MYASMSDCLIKTIR.[S] | 1xCarbamidomethyl [C8]; 1xHomocysteineIAA [K11] | 4,32935E-05 | 1862,86314 | Brain/ FAIMS/IAA |
| Q9D6R2 | Q9D6R2 [116-134] | [R].KTFDLYANVRPCVSIEGYK.[T] | 1xCarbamidomethyl [C12]; 1xHomocysteineIAA [K1] | 4,32935E-05 | 2434,18936 | Brain/ FAIMS/IAA |
| Q9D6R2 | Q9D6R2 [170-178] | [K].LITEEASKR.[I] | 1xHomocysteineMMTS [K8] | 0,00271031 | 1209,5966 | Liver/ FAIMS/MMTS |
| Q9D6R2 | Q9D6R2 [53-66] | [K].IFDAAKAPIQWEER.[N] | 1xHomocysteineMMTS [K6] | 4,49774E-05 | 1836,87714 | Liver/ FAIMS/MMTS |
| Q9D6R2 | Q9D6R2 [170-178] | [K].LITEEASKR.[I] | 1xHomocysteineMMTS [K8] | 0,00271031 | 1209,5966 | Brain/ FAIMS/MMTS |
| Q9D6R2 | Q9D6R2 [53-66] | [K].IFDAAKAPIQWEER.[N] | 1xHomocysteineMMTS [K6] | 4,49774E-05 | 1836,87714 | Brain/ FAIMS/MMTS |
| Q9D6Y7 | Q9D6Y7 [180-194] | [K].VLSKHNFGPITTDIR.[E] | 1xHomocysteineIAA [K4] | 4,32935E-05 | 1871,97963 | Liver/ FAIMS/IAA |
| Q9D6Y7 | Q9D6Y7 [180-194] | [K].VLSKHNFGPITTDIR.[E] | 1xHomocysteineMMTS [K4] | 4,49774E-05 | 1860,94588 | Liver/ FAIMS/MMTS |
| Q9D6Y7 | Q9D6Y7 [39-54] | [R].TEPIPVTAKHHVSGNR.[T] | 1xHomocysteineMMTS [K9] | 4,49774E-05 | 1905,9422 | Liver/ FAIMS/MMTS |
| Q9D819 | Q9D819 [64-74] | [K].DPLNPIKQDVK.[K] | 1xHomocysteineIAA [K7] | 0,000741565 | 1440,75153 | Brain/ FAIMS/IAA |
| Q9D819 | Q9D819 [222-233] | [K].STHDYWKALVTK.[K] | 1xHomocysteineMMTS [K7] | 0,000173718 | 1611,7658 | Liver/ No FAIMS/MMTS |
| Q9D819 | Q9D819 [212-221] | [K].NKDFAVDIIK.[S] | 1xHomocysteineMMTS [K2] | 0,000530332 | 1325,6592 | Liver/ No FAIMS/MMTS |
| Q9D819 | Q9D819 [178-191] | [R].LKPGYLEATVDWFR.[R] | 1xHomocysteineMMTS [K2] | 4,49774E-05 | 1857,90262 | Liver/ No FAIMS/MMTS |
| Q9D819 | Q9D819 [53-63] | [R].WSNAKMEIATK.[D] | 1xOxidation [M6]; 1xHomocysteineMMTS [K5] | 0,00980048 | 1457,65855 | Liver/ FAIMS/MMTS |
| Q9D819 | Q9D819 [222-233] | [K].STHDYWKALVTK.[K] | 1xHomocysteineMMTS [K7] | 0,000173718 | 1611,7658 | Liver/ FAIMS/MMTS |
| Q9D819 | Q9D819 [64-74] | [K].DPLNPIKQDVK.[K] | 1xHomocysteineMMTS [K7] | 0,00338636 | 1429,71778 | Liver/ FAIMS/MMTS |
| Q9D819 | Q9D819 [58-70] | [K].MEIATKDPLNPIK.[Q] | 1xHomocysteineMMTS [K6] | 0,000282457 | 1632,81578 | Liver/ FAIMS/MMTS |
| Q9D819 | Q9D819 [195-211] | [K].VPDGKPENEFAFNAEFK.[N] | 1xHomocysteineMMTS [K5] | 0,0004709 | 2101,93577 | Liver/ FAIMS/MMTS |
| Q9D819 | Q9D819 [23-41] | [K].NEKGQYISPFHDVPIYADK.[D] | 1xHomocysteineMMTS [K3] | 0,00732281 | 2384,10496 | Liver/ FAIMS/MMTS |
| Q9D819 | Q9D819 [212-221] | [K].NKDFAVDIIK.[S] | 1xHomocysteineMMTS [K2] | 0,000530332 | 1325,6592 | Liver/ FAIMS/MMTS |
| Q9D819 | Q9D819 [178-191] | [R].LKPGYLEATVDWFR.[R] | 1xHomocysteineMMTS [K2] | 4,49774E-05 | 1857,90262 | Liver/ FAIMS/MMTS |
| Q9D855 | Q9D855 [19-29] | [R].KWYYNAAGFNK.[L] | 1xHomocysteineIAA [K1] | 4,32935E-05 | 1535,70999 | Brain/ FAIMS/IAA |
| Q9D855 | Q9D855 [5-18] | [R].SAVSASSKWLDGFR.[K] | 1xHomocysteineMMTS [K8] | 0,00115861 | 1673,77742 | Liver/ FAIMS/MMTS |
| Q9D855 | Q9D855 [84-100] | [K].YEEDKFYLEPYLKEVIR.[E] | 1xHomocysteineMMTS [K13] | 4,49774E-05 | 2397,15052 | Liver/ FAIMS/MMTS |
| Q9D855 | Q9D855 [84-96] | [K].YEEDKFYLEPYLK.[E] | 1xHomocysteineMMTS [K5] | 0,00448015 | 1899,85434 | Brain/ FAIMS/MMTS |
| Q9D855 | Q9D855 [84-100] | [K].YEEDKFYLEPYLKEVIR.[E] | 1xHomocysteineMMTS [K13] | 4,49774E-05 | 2397,15052 | Brain/ FAIMS/MMTS |
| Q9D892 | Q9D892 [95-110] | [K].LKPEGLHQLLAGFEDK.[S] | 1xHomocysteineIAA [K2] | 0,000194827 | 1969,02116 | Brain/ FAIMS/IAA |
| Q9D8B3 | Q9D8B3 [108-117] | [K].NMGYAAKAMK.[A] | 1xHomocysteineIAA [K7] | 0,000177335 | 1258,5741 | Brain/ FAIMS/IAA |
| Q9D8B4 | Q9D8B4 [82-104] | [R].EKPDDPLNYFIGGCAGGLTLGAR.[T] | 1xCarbamidomethyl [C14]; 1xHomocysteineIAA [K2] | 4,32935E-05 | 2595,23302 | Brain/ FAIMS/IAA |
| Q9D8B7 | Q9D8B7 [92-101] | [R].TDVFGKTSLR.[I] | 1xHomocysteineIAA [K6] | 0,00012645 | 1297,6569 | Brain/ FAIMS/IAA |
| Q9D8E6 | Q9D8E6 [292-300] | [R].ILKSPEIQR.[A] | 1xHomocysteineIAA [K3] | 0,00193838 | 1257,69837 | Liver/ FAIMS/IAA |
| Q9D8E6 | Q9D8E6 [292-300] | [R].ILKSPEIQR.[A] | 1xHomocysteineIAA [K3] | 0,00193838 | 1257,69837 | Brain/ FAIMS/IAA |
| Q9D8E6 | Q9D8E6 [15-29] | [K].GESSGKNVTLPAVFK.[A] | 1xHomocysteineMMTS [K6] | 0,00331216 | 1696,83969 | Liver/ FAIMS/MMTS |
| Q9D8E6 | Q9D8E6 [353-364] | [K].KLEAAATALATK.[S] | 1xHomocysteineMMTS [K1] | 0,000211896 | 1350,71197 | Liver/ FAIMS/MMTS |
| Q9D8N0 | Q9D8N0 [209-219] | [K].LCEKMAQFDAK.[K] | 1xCarbamidomethyl [C2]; 1xHomocysteineIAA [K4] | 0,00103362 | 1514,68002 | Brain/ FAIMS/IAA |
| Q9D8Y0 | Q9D8Y0 [1-12] | [-].MATDELASKLSR.[R] | 1xMet-loss+Acetyl [N-Term]; 1xHomocysteineIAA [K9] | 4,32935E-05 | 1406,6944 | Liver/ FAIMS/IAA |
| Q9D8Y0 | Q9D8Y0 [135-151] | [K].SMIQEVDEDFDSKLSFR.[E] | 1xHomocysteineIAA [K13] | 4,32935E-05 | 2219,99474 | Liver/ FAIMS/IAA |
| Q9D8Y0 | Q9D8Y0 [1-12] | [-].MATDELASKLSR.[R] | 1xMet-loss+Acetyl [N-Term]; 1xHomocysteineMMTS [K9] | 4,49774E-05 | 1395,66066 | Liver/ FAIMS/MMTS |
| Q9D8Y0 | Q9D8Y0 [79-91] | [R].VFNPYTEFKEFSR.[K] | 1xHomocysteineMMTS [K9] | 4,49774E-05 | 1826,82404 | Brain/ No FAIMS/MMTS |
| Q9D8Y0 | Q9D8Y0 [1-12] | [-].MATDELASKLSR.[R] | 1xMet-loss+Acetyl [N-Term]; 1xHomocysteineMMTS [K9] | 4,49774E-05 | 1395,66066 | Brain/ FAIMS/MMTS |
| Q9D8Y0 | Q9D8Y0 [79-91] | [R].VFNPYTEFKEFSR.[K] | 1xHomocysteineMMTS [K9] | 4,49774E-05 | 1826,82404 | Brain/ FAIMS/MMTS |
| Q9D8Y0 | Q9D8Y0 [192-206] | [K].NFFEAKVQAINVSSR.[F] | 1xHomocysteineMMTS [K6] | 8,93346E-05 | 1872,9095 | Brain/ FAIMS/MMTS |
| Q9DAK9 | Q9DAK9 [86-98] | [R].KIHVYGYSMGYGR.[A] | 1xHomocysteineIAA [K1] | 0,000595249 | 1704,79849 | Brain/ FAIMS/IAA |
| Q9DAS9 | Q9DAS9 [32-48] | [K].VSKASADLMSYCEEHAR.[S] | 1xCarbamidomethyl [C12]; 1xHomocysteineIAA [K3] | 4,32935E-05 | 2127,92562 | Brain/ FAIMS/IAA |
| Q9DAW9 | Q9DAW9 [252-265] | [K].VASQKGMSVYGLGR.[Q] | 1xHomocysteineIAA [K5] | 0,000348152 | 1626,80906 | Brain/ FAIMS/IAA |
| Q9DB05 | Q9DB05 [223-239] | [K].LAVQKYEELFPAFSDSR.[E] | 1xHomocysteineIAA [K5] | 4,32935E-05 | 2174,05867 | Brain/ FAIMS/IAA |
| Q9DB05; P28663 | Q9DB05 [141-163]; P28663 [141-163] | [K].AIAHYEQSADYYKGEESNSSANK.[C] | 1xHomocysteineIAA [K13] | 4,32935E-05 | 2736,18421 | Brain/ No FAIMS/IAA |
| Q9DB05; P28663 | Q9DB05 [141-163]; P28663 [141-163] | [K].AIAHYEQSADYYKGEESNSSANK.[C] | 1xHomocysteineIAA [K13] | 4,32935E-05 | 2736,18421 | Brain/ FAIMS/IAA |
| Q9DB15 | Q9DB15 [137-145] | [R].LTEAKPVDK.[V] | 1xHomocysteineMMTS [K5] | 0,00794311 | 1163,57989 | Liver/ FAIMS/MMTS |
| Q9DB20 | Q9DB20 [173-188] | [K].LEIKTDPSIMGGMIVR.[I] | 1xHomocysteineIAA [K4] | 4,32935E-05 | 1933,99079 | Brain/ FAIMS/IAA |
| Q9DB20 | Q9DB20 [173-188] | [K].LEIKTDPSIMGGMIVR.[I] | 1xHomocysteineMMTS [K4] | 0,00213323 | 1922,95704 | Brain/ FAIMS/MMTS |
| Q9DB77 | Q9DB77 [197-217] | [R].MGKITSEELHYFVQNHFTSAR.[M] | 1xHomocysteineIAA [K3] | 4,32935E-05 | 2669,2599 | Brain/ FAIMS/IAA |
| Q9DBB8 | Q9DBB8 [323-333] | [K].AIGVTFPQDKR.[-] | 1xHomocysteineMMTS [K10] | 0,000152766 | 1394,6919 | Liver/ FAIMS/MMTS |
| Q9DBG3 | Q9DBG3 [843-867] | [K].DIPNENELQFQIKECHLNADTVSSK.[L] | 1xCarbamidomethyl [C15]; 1xHomocysteineIAA [K13] | 4,32935E-05 | 3103,44592 | Brain/ FAIMS/IAA |
| Q9DBH5 | Q9DBH5 [303-316] | [K].SPKDNVDDPTGNFR.[N] | 1xHomocysteineMMTS [K3] | 0,00650587 | 1724,73668 | Liver/ FAIMS/MMTS |
| Q9DBJ1 | Q9DBJ1 [107-116] | [K].HGEAQVKIWR.[R] | 1xHomocysteineIAA [K7] | 4,32935E-05 | 1397,71066 | Brain/ FAIMS/IAA |
| Q9DBJ1 | Q9DBJ1 [107-116] | [K].HGEAQVKIWR.[R] | 1xHomocysteineMMTS [K7] | 0,000530332 | 1386,67692 | Liver/ No FAIMS/MMTS |
| Q9DBJ1 | Q9DBJ1 [107-116] | [K].HGEAQVKIWR.[R] | 1xHomocysteineMMTS [K7] | 0,000530332 | 1386,67692 | Liver/ FAIMS/MMTS |
| Q9DBJ1 | Q9DBJ1 [107-116] | [K].HGEAQVKIWR.[R] | 1xHomocysteineMMTS [K7] | 0,000530332 | 1386,67692 | Brain/ FAIMS/MMTS |
| Q9DBJ1 | Q9DBJ1 [22-40] | [R].FSGWYDADLSPAGHEEAKR.[G] | 1xHomocysteineMMTS [K18] | 4,49774E-05 | 2298,99066 | Brain/ FAIMS/MMTS |
| Q9DBJ1 | Q9DBJ1 [163-179] | [R].ALPFWNEEIVPQIKEGK.[R] | 1xHomocysteineMMTS [K14] | 4,49774E-05 | 2161,08204 | Brain/ FAIMS/MMTS |
| Q9DBJ1 | Q9DBJ1 [242-253] | [K].AMEAVAAQGKVK.[K] | 1xHomocysteineMMTS [K10] | 0,00109439 | 1365,66872 | Brain/ FAIMS/MMTS |
| Q9DBJ1; O70250 | Q9DBJ1 [91-106]; O70250 [91-106] | [R].HYGGLTGLNKAETAAK.[H] | 1xHomocysteineIAA [K10] | 4,32935E-05 | 1804,90104 | Brain/ No FAIMS/IAA |
| Q9DBJ1; O70250 | Q9DBJ1 [91-106]; O70250 [91-106] | [R].HYGGLTGLNKAETAAK.[H] | 1xHomocysteineIAA [K10] | 4,32935E-05 | 1804,90104 | Brain/ FAIMS/IAA |
| Q9DBJ1; O70250 | Q9DBJ1 [91-106]; O70250 [91-106] | [R].HYGGLTGLNKAETAAK.[H] | 1xHomocysteineMMTS [K10] | 4,49774E-05 | 1793,8673 | Liver/ FAIMS/MMTS |
| Q9DBP5 | Q9DBP5 [152-171] | [R].IQTYLESTKPIIDLYEEMGK.[V] | 1xHomocysteineIAA [K9] | 4,32935E-05 | 2545,25644 | Brain/ No FAIMS/IAA |
| Q9DBP5 | Q9DBP5 [152-171] | [R].IQTYLESTKPIIDLYEEMGK.[V] | 1xHomocysteineIAA [K9] | 4,32935E-05 | 2545,25644 | Brain/ FAIMS/IAA |

| | | | | | | |
|---|---|---|---|---|---|---|
| Q9DBP5 | Q9DBP5 [23-39] | [R].IVEKYGYTHLSAGELLR.[D] | 1xHomocysteineMMTS [K4] | 4,49774E-05 | 2112,06164 | Liver/ FAIMS/MMTS |
| Q9DBP5 | Q9DBP5 [43-55] | [R].KNPDSQYGELIEK.[Y] | 1xHomocysteineMMTS [K1] | 0,000530332 | 1683,77167 | Liver/ FAIMS/MMTS |
| Q9DC07 | Q9DC07 [113-128] | [R].TQEQISNVKYHEDFEK.[T] | 1xHomocysteineIAA [K9] | 4,32935E-05 | 2168,99171 | Brain/ FAIMS/IAA |
| Q9DC51; P08752; B2RSH2 | Q9DC51 [33-46]; P08752 [33-46]; B2RSH2 [33-46] | [KR].EVKLLLLGAGESGK.[S] | 1xHomocysteineIAA [K3] | 6,30912E-05 | 1587,87745 | Brain/ No FAIMS/IAA |
| Q9DC51; P08752; B2RSH2 | Q9DC51 [33-46]; P08752 [33-46]; B2RSH2 [33-46] | [KR].EVKLLLLGAGESGK.[S] | 1xHomocysteineIAA [K3] | 6,30912E-05 | 1587,87745 | Brain/ FAIMS/IAA |
| Q9DC51; P08752; B2RSH2 | Q9DC51 [33-46]; P08752 [33-46]; B2RSH2 [33-46] | [KR].EVKLLLLGAGESGK.[S] | 1xHomocysteineMMTS [K3] | 8,93346E-05 | 1576,84371 | Brain/ FAIMS/MMTS |
| Q9DC51; P18872; P08752; B2RSH2 | Q9DC51 [179-192]; P18872 [180-193]; P08752 [180-193]; B2RSH2 [179-192] | [R].VKTTGIVETHFTFK.[END] | 1xHomocysteineIAA [K2] | 4,32935E-05 | 1781,92547 | Brain/ No FAIMS/IAA |
| Q9DC51; P18872; P08752; B2RSH2 | Q9DC51 [179-192]; P18872 [180-193]; P08752 [180-193]; B2RSH2 [179-192] | [R].VKTTGIVETHFTFK.[END] | 1xHomocysteineIAA [K2] | 4,32935E-05 | 1781,92547 | Brain/ FAIMS/IAA |
| Q9DC51; Q8CGK7; Q6R0H7; P18872; P08752; B2RSH2 | Q9DC51 [36-51]; Q8CGK7 [45-60]; Q6R0H7 [782-797]; P18872 [36-51]; P08752 [36-51]; B2RSH2 [36-51] | [RK].LLLLGAGESGKSTIVK.[Q] | 1xHomocysteineMMTS [K11] | 4,49774E-05 | 1748,96489 | Brain/ No FAIMS/MMTS |
| Q9DC51; Q8CGK7; Q6R0H7; P18872; P08752; B2RSH2 | Q9DC51 [36-51]; Q8CGK7 [45-60]; Q6R0H7 [782-797]; P18872 [36-51]; P08752 [36-51]; B2RSH2 [36-51] | [RK].LLLLGAGESGKSTIVK.[Q] | 1xHomocysteineMMTS [K11] | 4,49774E-05 | 1748,96489 | Brain/ FAIMS/MMTS |
| Q9DC69 | Q9DC69 [118-132] | [R].KAVQHSNVVINLIGR.[E] | 1xHomocysteineIAA [K1] | 6,30912E-05 | 1822,0116 | Brain/ FAIMS/IAA |
| Q9DCJ5 | Q9DCJ5 [1-19] | [-].MPGIVELPTLEELKVEEVK.[V] | 1xMet-loss [N-Term]; 1xHomocysteineIAA [K14] | 4,32935E-05 | 2196,1832 | Brain/ FAIMS/IAA |
| Q9DCL9 | Q9DCL9 [31-46] | [R].VLLQSKDQITAGNAAR.[K] | 1xHomocysteineIAA [K6] | 0,000741565 | 1858,98036 | Brain/ FAIMS/IAA |
| Q9DCM0 | Q9DCM0 [61-71] | [R].DAQLIKELGLK.[L] | 1xHomocysteineMMTS [K6] | 0,00294218 | 1390,74327 | Liver/ FAIMS/MMTS |
| Q9DCS9 | Q9DCS9 [95-109] | [R].DFKVDQEIMNIIQER.[L] | 1xHomocysteineIAA [K3] | 4,32935E-05 | 2051,98887 | Liver/ FAIMS/IAA |
| Q9DCS9 | Q9DCS9 [1-16] | [-].MPDSWDKDVYPEPPSR.[T] | 1xMet-loss [N-Term]; 1xHomocysteineIAA [K7] | 4,32935E-05 | 1961,8698 | Brain/ No FAIMS/IAA |
| Q9DCS9 | Q9DCS9 [1-16] | [-].MPDSWDKDVYPEPPSR.[T] | 1xMet-loss [N-Term]; 1xHomocysteineIAA [K7] | 4,32935E-05 | 1961,8698 | Brain/ FAIMS/IAA |
| Q9DCS9 | Q9DCS9 [95-109] | [R].DFKVDQEIMNIIQER.[L] | 1xOxidation [M9]; 1xHomocysteineMMTS [K3] | 0,00669057 | 2056,95004 | Liver/ FAIMS/MMTS |
| Q9DCS9 | Q9DCS9 [128-137] | [K].ELEQFTKVTK.[A] | 1xHomocysteineMMTS [K7] | 0,00359453 | 1385,68033 | Liver/ FAIMS/MMTS |
| Q9DCT2 | Q9DCT2 [231-245] | [R].KFDLNSPWEAFPAYR.[Q] | 1xHomocysteineIAA [K1] | 4,32935E-05 | 2014,94799 | Liver/ FAIMS/IAA |
| Q9DCT2 | Q9DCT2 [231-245] | [R].KFDLNSPWEAFPAYR.[Q] | 1xHomocysteineIAA [K1] | 4,32935E-05 | 2014,94799 | Brain/ No FAIMS/IAA |
| Q9DCT2 | Q9DCT2 [231-245] | [R].KFDLNSPWEAFPAYR.[Q] | 1xHomocysteineIAA [K1] | 4,32935E-05 | 2014,94799 | Brain/ FAIMS/IAA |
| Q9DCT2 | Q9DCT2 [231-245] | [R].KFDLNSPWEAFPAYR.[Q] | 1xHomocysteineMMTS [K1] | 0,00185311 | 2003,91425 | Liver/ FAIMS/MMTS |
| Q9DCT2 | Q9DCT2 [231-245] | [R].KFDLNSPWEAFPAYR.[Q] | 1xHomocysteineMMTS [K1] | 0,00185311 | 2003,91425 | Brain/ FAIMS/MMTS |
| Q9DCT8 | Q9DCT8 [66-97] | [K].GVNIGGAGSYIYEKPQTEAPQVTGPIEVPVVR.[T] | 1xHomocysteineIAA [K14] | 4,32935E-05 | 3499,78898 | Brain/ FAIMS/IAA |
| Q9DCU2 | Q9DCU2 [1-11] | [-].MAEFPSKVSTR.[T] | 1xMet-loss+Acetyl [N-Term]; 1xHomocysteineIAA [K7] | 9,61721E-05 | 1337,65181 | Brain/ FAIMS/IAA |
| Q9DCU2 | Q9DCU2 [1-11] | [-].MAEFPSKVSTR.[T] | 1xMet-loss+Acetyl [N-Term]; 1xHomocysteineMMTS [K7] | 0,000738471 | 1326,61807 | Brain/ FAIMS/MMTS |
| Q9DCW4 | Q9DCW4 [234-248] | [R].SAGVKVETTEDLVAK.[L] | 1xHomocysteineIAA [K5] | 4,32935E-05 | 1720,87858 | Liver/ FAIMS/IAA |
| Q9DCW4 | Q9DCW4 [13-21] | [R].VIDFAVKIR.[V] | 1xHomocysteineMMTS [K7] | 0,000689517 | 1223,6639 | Liver/ No FAIMS/MMTS |
| Q9DCW4 | Q9DCW4 [234-248] | [R].SAGVKVETTEDLVAK.[L] | 1xHomocysteineMMTS [K5] | 4,49774E-05 | 1709,84483 | Liver/ No FAIMS/MMTS |
| Q9DCW4 | Q9DCW4 [206-221] | [K].IEVVKAGDLGVDLTSK.[V] | 1xHomocysteineMMTS [K5] | 4,49774E-05 | 1806,93398 | Liver/ No FAIMS/MMTS |
| Q9DCW4 | Q9DCW4 [115-124] | [K].EKVDLLFLGK.[Q] | 1xHomocysteineMMTS [K2] | 0,000250158 | 1324,70034 | Liver/ No FAIMS/MMTS |
| Q9DCW4 | Q9DCW4 [175-186] | [R].LKLPAVVTADLR.[L] | 1xHomocysteineMMTS [K2] | 4,49774E-05 | 1458,8171 | Liver/ No FAIMS/MMTS |
| Q9DCW4 | Q9DCW4 [239-250] | [K].VETTEDLVAKLK.[E] | 1xHomocysteineMMTS [K10] | 4,49774E-05 | 1508,76988 | Liver/ No FAIMS/MMTS |
| Q9DCW4 | Q9DCW4 [13-21] | [R].VIDFAVKIR.[V] | 1xHomocysteineMMTS [K7] | 0,000689517 | 1223,6639 | Liver/ FAIMS/MMTS |
| Q9DCW4 | Q9DCW4 [22-35] | [R].VKPDKSGVVTDGVK.[H] | 1xHomocysteineMMTS [K5] | 4,49774E-05 | 1591,81822 | Liver/ FAIMS/MMTS |
| Q9DCW4 | Q9DCW4 [234-248] | [R].SAGVKVETTEDLVAK.[L] | 1xHomocysteineMMTS [K5] | 4,49774E-05 | 1709,84483 | Liver/ FAIMS/MMTS |
| Q9DCW4 | Q9DCW4 [206-221] | [K].IEVVKAGDLGVDLTSK.[V] | 1xHomocysteineMMTS [K5] | 4,49774E-05 | 1806,93398 | Liver/ FAIMS/MMTS |
| Q9DCW4 | Q9DCW4 [115-124] | [K].EKVDLLFLGK.[Q] | 1xHomocysteineMMTS [K2] | 0,000250158 | 1324,70034 | Liver/ FAIMS/MMTS |
| Q9DCW4 | Q9DCW4 [175-186] | [R].LKLPAVVTADLR.[L] | 1xHomocysteineMMTS [K2] | 4,49774E-05 | 1458,8171 | Liver/ FAIMS/MMTS |
| Q9DCW4 | Q9DCW4 [239-250] | [K].VETTEDLVAKLK.[E] | 1xHomocysteineMMTS [K10] | 4,49774E-05 | 1508,76988 | Liver/ FAIMS/MMTS |
| Q9DCW4 | Q9DCW4 [107-114] | [R].VLAKLAEK.[E] | 1xHomocysteineMMTS [K4] | 0,00365334 | 1034,57368 | Liver/ FAIMS/MMTS |
| Q9DCW4 | Q9DCW4 [211-233] | [K].AGDLGVDLTSKVSVISVEEPPQR.[S] | 1xHomocysteineMMTS [K11] | 4,49774E-05 | 2559,2793 | Liver/ FAIMS/MMTS |
| Q9DCX2 | Q9DCX2 [112-121] | [R].IQEYEKQLEK.[M] | 1xHomocysteineIAA [K6] | 4,32935E-05 | 1481,73046 | Liver/ FAIMS/IAA |
| Q9DCX2 | Q9DCX2 [112-121] | [R].IQEYEKQLEK.[M] | 1xHomocysteineIAA [K6] | 4,32935E-05 | 1481,73046 | Brain/ No FAIMS/IAA |
| Q9DCX2 | Q9DCX2 [112-121] | [R].IQEYEKQLEK.[M] | 1xHomocysteineIAA [K6] | 4,32935E-05 | 1481,73046 | Brain/ FAIMS/IAA |
| Q9DCX2 | Q9DCX2 [42-58] | [R].LASLSEKPPAIDWAYYR.[A] | 1xHomocysteineMMTS [K7] | 4,49774E-05 | 2143,03509 | Liver/ No FAIMS/MMTS |
| Q9DCX2 | Q9DCX2 [79-99] | [K].IPVPEDKYTALVDQEEKEDVK.[S] | 1xHomocysteineMMTS [K7] | 4,49774E-05 | 2608,25208 | Liver/ No FAIMS/MMTS |
| Q9DCX2 | Q9DCX2 [86-99] | [K].YTALVDQEEKEDVK.[S] | 1xHomocysteineMMTS [K10] | 4,49774E-05 | 1829,82958 | Liver/ No FAIMS/MMTS |
| Q9DCX2 | Q9DCX2 [42-58] | [R].LASLSEKPPAIDWAYYR.[A] | 1xHomocysteineMMTS [K7] | 4,49774E-05 | 2143,03509 | Liver/ FAIMS/MMTS |
| Q9DCX2 | Q9DCX2 [26-41] | [K].AIGNALKSWNETFHAR.[L] | 1xHomocysteineMMTS [K7] | 4,49774E-05 | 1977,9422 | Liver/ FAIMS/MMTS |
| Q9DCX2 | Q9DCX2 [112-121] | [R].IQEYEKQLEK.[M] | 1xHomocysteineMMTS [K6] | 0,000250158 | 1470,69671 | Liver/ FAIMS/MMTS |
| Q9DCX2 | Q9DCX2 [79-99] | [K].IPVPEDKYTALVDQEEKEDVK.[S] | 1xHomocysteineMMTS [K7] | 4,49774E-05 | 2608,25208 | Liver/ FAIMS/MMTS |

| | | | | | | |
|---|---|---|---|---|---|---|
| Q9DCX2 | Q9DCX2 [86-99] | [K].YTALVDQEEKEDVK.[S] | 1xHomocysteineMMTS [K10] | 4,49774E-05 | 1829,82958 | Liver/ FAIMS/MMTS |
| Q9DCX2 | Q9DCX2 [42-58] | [R].LASLSEKPPAIDWAYYR.[A] | 1xHomocysteineMMTS [K7] | 4,49774E-05 | 2143,03509 | Brain/ FAIMS/MMTS |
| Q9DCX2 | Q9DCX2 [26-41] | [K].AIGNALKSWNETFHAR.[L] | 1xHomocysteineMMTS [K7] | 4,49774E-05 | 1977,9422 | Brain/ FAIMS/MMTS |
| Q9DCX2 | Q9DCX2 [112-121] | [R].IQEYEKQLEK.[M] | 1xHomocysteineMMTS [K6] | 0,000250158 | 1470,69671 | Brain/ FAIMS/MMTS |
| Q9DCX2 | Q9DCX2 [59-72] | [R].ANVAKPGLVDDFEK.[K] | 1xHomocysteineMMTS [K5] | 0,00618488 | 1665,79749 | Brain/ FAIMS/MMTS |
| Q9EPJ9 | Q9EPJ9 [123-148] | [K].EWSLESSPAQNWTPPQPKTLQFTAHR.[A] | 1xHomocysteineIAA [K18] | 4,32935E-05 | 3210,54254 | Brain/ FAIMS/IAA |
| Q9EQ20 | Q9EQ20 [108-117] | [R].YQQLIKENLK.[E] | 1xHomocysteineIAA [K6] | 0,00434736 | 1450,77226 | Brain/ FAIMS/IAA |
| Q9EQ20 | Q9EQ20 [42-52] | [K].LFIDGKFVESK.[S] | 1xHomocysteineMMTS [K6] | 0,000133296 | 1445,71672 | Liver/ No FAIMS/MMTS |
| Q9EQ20 | Q9EQ20 [42-52] | [K].LFIDGKFVESK.[S] | 1xHomocysteineMMTS [K6] | 0,000133296 | 1445,71672 | Liver/ FAIMS/MMTS |
| Q9EQ20 | Q9EQ20 [108-117] | [R].YQQLIKENLK.[E] | 1xHomocysteineMMTS [K6] | 0,000829616 | 1439,73852 | Liver/ FAIMS/MMTS |
| Q9EQS3 | Q9EQS3 [17-29] | [R].YLEKSGVLDTLTK.[V] | 1xHomocysteineIAA [K4] | 0,000177335 | 1640,85638 | Brain/ FAIMS/IAA |
| Q9EQS3 | Q9EQS3 [17-29] | [R].YLEKSGVLDTLTK.[V] | 1xHomocysteineMMTS [K4] | 0,00889969 | 1629,82264 | Liver/ FAIMS/MMTS |
| Q9EQU5 | Q9EQU5 [150-166] | [K].VLSKEFHLNESGDPSSK.[S] | 1xHomocysteineIAA [K4] | 4,32935E-05 | 2047,97533 | Brain/ FAIMS/IAA |
| Q9ERD7 | Q9ERD7 [360-379] | [R].GLKMSSTFIGNSTAIQELFK.[R] | 1xHomocysteineIAA [K3] | 4,32935E-05 | 2346,18321 | Brain/ No FAIMS/IAA |
| Q9ERD7 | Q9ERD7 [360-379] | [R].GLKMSSTFIGNSTAIQELFK.[R] | 1xHomocysteineIAA [K3] | 4,32935E-05 | 2346,18321 | Brain/ FAIMS/IAA |
| Q9ERD7 | Q9ERD7 [321-336] | [R].MSMKEVDEQMLAIQSK.[N] | 1xHomocysteineMMTS [K4] | 0,00074635 | 2030,90877 | Brain/ FAIMS/MMTS |
| Q9ERD7 | Q9ERD7 [363-380] | [K].MSSTFIGNSTAIQELFKR.[I] | 1xHomocysteineMMTS [K17] | 0,000437433 | 2193,05009 | Brain/ FAIMS/MMTS |
| Q9ERD7 | Q9ERD7 [47-62] | [R].ISVYYNEASSHKYVPR.[A] | 1xHomocysteineMMTS [K12] | 4,49774E-05 | 2075,96774 | Brain/ FAIMS/MMTS |
| Q9ERL7; Q9CQI3 | Q9ERL7 [111-124]; Q9CQI3 [111-124] | [RK].LVQTAELTKVFEIR.[TN] | 1xHomocysteineIAA [K9] | 4,32935E-05 | 1820,99388 | Brain/ No FAIMS/IAA |
| Q9ERL7; Q9CQI3 | Q9ERL7 [111-124]; Q9CQI3 [111-124] | [RK].LVQTAELTKVFEIR.[TN] | 1xHomocysteineIAA [K9] | 4,32935E-05 | 1820,99388 | Brain/ FAIMS/IAA |
| Q9ERT9 | Q9ERT9 [136-152] | [K].SAESNPKTQEQCGVEPR.[T] | 1xCarbamidomethyl [C12]; 1xHomocysteineIAA [K7] | 0,00124346 | 2090,92298 | Brain/ FAIMS/IAA |
| Q9ES97 | Q9ES97 [827-837] | [R].VYKSVIQAVQK.[S] | 1xHomocysteineIAA [K3] | 4,32935E-05 | 1436,793 | Brain/ FAIMS/IAA |
| Q9ESM3 | Q9ESM3 [110-120] | [R].LDASLIIKNVR.[L] | 1xHomocysteineIAA [K8] | 0,000258769 | 1415,8039 | Brain/ No FAIMS/IAA |
| Q9JHI5 | Q9JHI5 [52-65] | [R].HTISKFLQENLAPK.[A] | 1xHomocysteineMMTS [K5] | 0,000829616 | 1788,91352 | Liver/ No FAIMS/MMTS |
| Q9JHI5 | Q9JHI5 [311-326] | [R].EAFGQKIGQFQLMQGK.[M] | 1xOxidation [M13]; 1xHomocysteineMMTS [K6] | 0,000530332 | 1988,93908 | Liver/ FAIMS/MMTS |
| Q9JHI5 | Q9JHI5 [311-326] | [R].EAFGQKIGQFQLMQGK.[M] | 1xHomocysteineMMTS [K6] | 0,000250158 | 1972,94417 | Liver/ FAIMS/MMTS |
| Q9JHI5 | Q9JHI5 [52-65] | [R].HTISKFLQENLAPK.[A] | 1xHomocysteineMMTS [K5] | 0,000829616 | 1788,91352 | Liver/ FAIMS/MMTS |
| Q9JHI5 | Q9JHI5 [66-79] | [K].AQEIDQTNDFKNLR.[E] | 1xHomocysteineMMTS [K11] | 0,000152766 | 1854,84729 | Liver/ FAIMS/MMTS |
| Q9JHU4 | Q9JHU4 [3758-3773] | [R].ILDDDTIITTLENLKR.[E] | 1xHomocysteineIAA [K15] | 4,32935E-05 | 2047,07398 | Brain/ FAIMS/IAA |
| Q9JHW2 | Q9JHW2 [61-74] | [K].IPGESTQKLSEVAK.[E] | 1xHomocysteineMMTS [K8] | 0,000250158 | 1649,8237 | Liver/ FAIMS/MMTS |
| Q9JIA1 | Q9JIA1 [457-470] | [K].VMKWGGSSFQDIQR.[M] | 1xHomocysteineIAA [K3] | 8,35381E-05 | 1812,85198 | Brain/ FAIMS/IAA |
| Q9JII6 | Q9JII6 [62-72] | [K].ESVGSGKAVPR.[E] | 1xHomocysteineIAA [K7] | 4,32935E-05 | 1260,6365 | Liver/ FAIMS/IAA |
| Q9JII6 | Q9JII6 [14-30] | [K].MPLIGLGTWKSEPGQVK.[A] | 1xHomocysteineIAA [K10] | 4,32935E-05 | 2015,04526 | Brain/ FAIMS/IAA |
| Q9JII6 | Q9JII6 [146-157] | [K].ALEVLVAKGLVK.[A] | 1xHomocysteineMMTS [K8] | 8,93346E-05 | 1402,81604 | Liver/ No FAIMS/MMTS |
| Q9JII6 | Q9JII6 [62-72] | [K].ESVGSGKAVPR.[E] | 1xHomocysteineMMTS [K7] | 4,49774E-05 | 1249,60275 | Liver/ FAIMS/MMTS |
| Q9JII6 | Q9JII6 [298-309] | [R].YIVPMITVDGKR.[V] | 1xOxidation [M5]; 1xHomocysteineMMTS [K11] | 0,00883288 | 1570,779 | Liver/ FAIMS/MMTS |
| Q9JII6 | Q9JII6 [146-157] | [K].ALEVLVAKGLVK.[A] | 1xHomocysteineMMTS [K8] | 8,93346E-05 | 1402,81604 | Liver/ FAIMS/MMTS |
| Q9JII6 | Q9JII6 [222-243] | [R].HPDEPVLLEEPVVLALAEKHGR.[S] | 1xHomocysteineMMTS [K19] | 4,49774E-05 | 2611,33709 | Liver/ FAIMS/MMTS |
| Q9JII6 | Q9JII6 [14-30] | [K].MPLIGLGTWKSEPGQVK.[A] | 1xHomocysteineMMTS [K10] | 0,000282457 | 2004,01152 | Liver/ FAIMS/MMTS |
| Q9JIS5 | Q9JIS5 [508-520] | [R].GGQYFNDKFIGLR.[L] | 1xHomocysteineIAA [K8] | 4,32935E-05 | 1688,82134 | Brain/ No FAIMS/IAA |
| Q9JIS5 | Q9JIS5 [508-520] | [R].GGQYFNDKFIGLR.[L] | 1xHomocysteineIAA [K8] | 4,32935E-05 | 1688,82134 | Brain/ FAIMS/IAA |
| Q9JIS8; Q91V14 | Q9JIS8 [878-893]; Q91V14 [881-896] | [R].IFTVAQMDDNSIQMKK.[D] | 1xHomocysteineIAA [K15] | 8,35381E-05 | 2042,97078 | Brain/ FAIMS/IAA |
| Q9JIW9; P63321 | Q9JIW9 [163-174]; P63321 [162-173] | [R].ANVDKVFFDLMR.[E] | 1xHomocysteineIAA [K5] | 9,61721E-05 | 1628,79234 | Brain/ FAIMS/IAA |
| Q9JIW9; P63321 | Q9JIW9 [163-174]; P63321 [162-173] | [R].ANVDKVFFDLMR.[E] | 1xHomocysteineMMTS [K5] | 0,0025135 | 1617,7586 | Brain/ FAIMS/MMTS |
| Q9JIY0 | Q9JIY0 [334-356] | [R].KAKDPPQSPPDSESEQLLLETER.[L] | 2xHomocysteineIAA [K1; K3] | 9,61721E-05 | 2942,38701 | Brain/ No FAIMS/IAA |
| Q9JIY0 | Q9JIY0 [334-356] | [R].KAKDPPQSPPDSESEQLLLETER.[L] | 2xHomocysteineIAA [K1; K3] | 9,61721E-05 | 2942,38701 | Brain/ FAIMS/IAA |
| Q9JJI8 | Q9JJI8 [4-16] | [R].KIEEIKDFLLTAR.[R] | 1xHomocysteineIAA [K6] | 0,00656758 | 1749,95677 | Liver/ FAIMS/IAA |
| Q9JJI8 | Q9JJI8 [4-16] | [R].KIEEIKDFLLTAR.[R] | 1xHomocysteineIAA [K6] | 0,00656758 | 1749,95677 | Brain/ FAIMS/IAA |
| Q9JJI8 | Q9JJI8 [4-16] | [R].KIEEIKDFLLTAR.[R] | 1xHomocysteineMMTS [K] | 0,00074635 | 1738,92302 | Liver/ FAIMS/MMTS |
| Q9JJV2 | Q9JJV2 [128-136] | [K].AYSMAKYLR.[D] | 1xHomocysteineIAA [K6] | 4,32935E-05 | 1276,61767 | Brain/ FAIMS/IAA |
| Q9JJV2 | Q9JJV2 [90-105] | [R].TKSQGGEPTYNVAVGR.[A] | 1xHomocysteineMMTS [K2] | 0,000152766 | 1826,85238 | Brain/ FAIMS/MMTS |
| Q9JKB1 | Q9JKB1 [135-145] | [R].AKFLENYDAIR.[V] | 1xHomocysteineIAA [K2] | 0,00271978 | 1513,74677 | Brain/ No FAIMS/IAA |
| Q9JKB1 | Q9JKB1 [135-145] | [R].AKFLENYDAIR.[V] | 1xHomocysteineIAA [K2] | 0,00271978 | 1513,74677 | Brain/ FAIMS/IAA |
| Q9JKB3 | Q9JKB3 [78-88] | [K].VLATKVLGTVK.[W] | 1xHomocysteineIAA [K5] | 0,000318237 | 1302,78137 | Liver/ FAIMS/IAA |
| Q9JKC6 | Q9JKC6 [42-61] | [K].KDTQAEKQEQAAAPGPAATK.[K] | 1xHomocysteineIAA [K] | 4,32935E-05 | 2214,08192 | Brain/ FAIMS/IAA |
| Q9JKC6 | Q9JKC6 [17-33] | [K].VPQATAEAKATPAADGK.[A] | 1xHomocysteineMMTS [K9] | 4,49774E-05 | 1788,86188 | Brain/ No FAIMS/MMTS |
| Q9JKC6 | Q9JKC6 [43-61] | [K].DTQAEKQEQAAAPGPAATK.[K] | 1xHomocysteineMMTS [K6] | 4,49774E-05 | 2074,95321 | Brain/ No FAIMS/MMTS |
| Q9JKC6 | Q9JKC6 [34-41] | [K].APLTKPVK.[K] | 1xHomocysteineMMTS [K5] | 0,00194807 | 1016,56312 | Brain/ No FAIMS/MMTS |
| Q9JKC6 | Q9JKC6 [49-62] | [K].QEQAAAPGPAATKK.[T] | 1xHomocysteineMMTS [K13] | 4,49774E-05 | 1530,74031 | Brain/ No FAIMS/MMTS |
| Q9JKC6 | Q9JKC6 [7-25] | [K].SASSPKPDTKVPQATAEAK.[A] | 1xHomocysteineMMTS [K] | 4,49774E-05 | 2076,01 | Brain/ No FAIMS/MMTS |
| Q9JKC6 | Q9JKC6 [17-33] | [K].VPQATAEAKATPAADGK.[A] | 1xHomocysteineMMTS [K9] | 4,49774E-05 | 1788,86188 | Brain/ FAIMS/MMTS |
| Q9JKC6 | Q9JKC6 [43-61] | [K].DTQAEKQEQAAAPGPAATK.[K] | 1xHomocysteineMMTS [K6] | 4,49774E-05 | 2074,95321 | Brain/ FAIMS/MMTS |
| Q9JKC6 | Q9JKC6 [43-62] | [K].DTQAEKQEQAAAPGPAATKK.[T] | 1xHomocysteineMMTS [K19] | 4,49774E-05 | 2203,04818 | Brain/ FAIMS/MMTS |
| Q9JKC6 | Q9JKC6 [49-62] | [K].QEQAAAPGPAATKK.[T] | 1xHomocysteineMMTS [K13] | 4,49774E-05 | 1530,74031 | Brain/ FAIMS/MMTS |
| Q9JKC6 | Q9JKC6 [7-25] | [K].SASSPKPDTKVPQATAEAK.[A] | 1xHomocysteineMMTS [K] | 4,49774E-05 | 2076,01 | Brain/ FAIMS/MMTS |
| Q9JKD3 | Q9JKD3 [178-196] | [R].GSGGSFSKAQEEWTTGAWK.[N] | 1xHomocysteineIAA [K8] | 9,61721E-05 | 2187,97639 | Brain/ FAIMS/IAA |
| Q9JKF6 | Q9JKF6 [189-199] | [R].LKGEAEYQEIR.[N] | 1xHomocysteineIAA [K2] | 0,00261348 | 1509,7366 | Brain/ FAIMS/IAA |
| Q9JKK7 | Q9JKK7 [14-30] | [K].NIDEDELLGKLSEEELK.[Q] | 1xHomocysteineIAA [K10] | 4,32935E-05 | 2148,03766 | Brain/ No FAIMS/IAA |
| Q9JKK7 | Q9JKK7 [14-30] | [K].NIDEDELLGKLSEEELK.[Q] | 1xHomocysteineIAA [K10] | 4,32935E-05 | 2148,03766 | Brain/ FAIMS/IAA |
| Q9JKK7 | Q9JKK7 [12-23] | [K].YKNIDEDELLGK.[L] | 1xHomocysteineMMTS [K2] | 0,000282457 | 1599,73931 | Brain/ FAIMS/MMTS |
| Q9JKL5 | Q9JKL5 [137-155] | [R].NVVEELLSGNPHIEKESAR.[S] | 1xHomocysteineIAA [K15] | 0,00314066 | 2295,13977 | Brain/ FAIMS/IAA |
| Q9JKR6 | Q9JKR6 [803-811] | [R].KLCQGLFFR.[V] | 1xCarbamidomethyl [C3]; 1xHomocysteineIAA [K1] | 0,000741565 | 1342,67586 | Brain/ FAIMS/IAA |

| | | | | | | |
|---|---|---|---|---|---|---|
| Q9JKR6 | Q9JKR6 [719-729] | [R].SVQKLEELTLR.[D] | 1xHomocysteineMMTS [K4] | 0,00166208 | 1478,77055 | Liver/ No FAIMS/MMTS |
| Q9JKR6 | Q9JKR6 [878-889] | [K].LPATEKPVLLSK.[D] | 1xHomocysteineMMTS [K6] | 0,000829616 | 1458,80587 | Liver/ FAIMS/MMTS |
| Q9JKR6 | Q9JKR6 [399-407] | [K].AVGKEELGK.[N] | 1xHomocysteineMMTS [K4] | 0,0093731 | 1093,53803 | Liver/ FAIMS/MMTS |
| Q9JL62 | Q9JL62 [1-16] | [-].MALLAEHLLKPLPADR.[Q] | 1xMet-loss+Acetyl [N-Term]; 1xHomocysteineIAA [K10] | 4,32935E-05 | 1873,03641 | Brain/ FAIMS/IAA |
| Q9JL62 | Q9JL62 [1-16] | [-].MALLAEHLLKPLPADR.[Q] | 1xMet-loss+Acetyl [N-Term]; 1xHomocysteineMMTS [K10] | 4,49774E-05 | 1862,00267 | Brain/ FAIMS/MMTS |
| Q9JLM8 | Q9JLM8 [162-177] | [R].AVSSLATAKGGPSEVR.[E] | 1xHomocysteineIAA [K9] | 9,61721E-05 | 1703,87449 | Brain/ FAIMS/IAA |
| Q9JLZ3 | Q9JLZ3 [173-186] | [R].VAASSAKMGLVETK.[L] | 1xHomocysteineIAA [K7] | 4,32935E-05 | 1565,80257 | Brain/ FAIMS/IAA |
| Q9JM63 | Q9JM63 [363-375] | [R].EQAEKEGSALSVR.[I] | 1xHomocysteineIAA [K5] | 0,00518879 | 1577,7588 | Brain/ FAIMS/IAA |
| Q9JM76 | Q9JM76 [1-25] | [-].MPAYHSSLMDPDTKLIGNMALLPLR.[S] | 1xMet-loss [N-Term]; 1xHomocysteineIAA [K14] | 4,32935E-05 | 2827,43034 | Brain/ FAIMS/IAA |
| Q9JMA1 | Q9JMA1 [294-307] | [R].LQEEITKQSPTLQR.[N] | 1xHomocysteineIAA [K7] | 0,00357163 | 1844,95347 | Brain/ FAIMS/IAA |
| Q9JMD3 | Q9JMD3 [93-107] | [R].KKWDSNVIETFDIAR.[L] | 1xHomocysteineMMTS [K2] | 4,49774E-05 | 1984,96193 | Liver/ FAIMS/MMTS |
| Q9JMD3 | Q9JMD3 [1-19] | [-].MEKPAASTEPQGSRPALGR.[E] | 1xAcetyl [N-Term]; 1xHomocysteineMMTS [K3] | 0,00386874 | 2188,03075 | Liver/ FAIMS/MMTS |
| Q9JMG7 | Q9JMG7 [190-202] | [R].NTTADLQKAGEGT.[-] | 1xHomocysteineIAA [K8] | 0,00263986 | 1479,6744 | Brain/ FAIMS/IAA |
| Q9QUI0 | Q9QUI0 [99-118] | [K].WTPEVKHFCPNVPIILVGNK.[K] | 1xCarbamidomethyl [C9]; 1xHomocysteineIAA [K6] | 8,35381E-05 | 2522,30467 | Brain/ FAIMS/IAA |
| Q9QUK6 | Q9QUK6 [434-446] | [K].RVTEFSAFLSLEK.[L] | 1xHomocysteineIAA [K13] | 0,00454947 | 1700,86762 | Liver/ FAIMS/IAA |
| Q9QUK6 | Q9QUK6 [434-446] | [K].RVTEFSAFLSLEK.[L] | 1xHomocysteineIAA [K13] | 0,00454947 | 1700,86762 | Brain/ FAIMS/IAA |
| Q9QUM9 | Q9QUM9 [44-54] | [R].GKDCAVIVTQK.[K] | 1xCarbamidomethyl [C4]; 1xHomocysteineIAA [K2] | 0,00500667 | 1392,69738 | Brain/ FAIMS/IAA |
| Q9QUP5 | Q9QUP5 [42-55] | [R].LLVEAEQAKVFSHR.[G] | 1xHomocysteineIAA [K9] | 4,32935E-05 | 1800,94251 | Brain/ No FAIMS/IAA |
| Q9QUP5 | Q9QUP5 [42-55] | [R].LLVEAEQAKVFSHR.[G] | 1xHomocysteineIAA [K9] | 4,32935E-05 | 1800,94251 | Brain/ FAIMS/IAA |
| Q9QUP5 | Q9QUP5 [42-55] | [R].LLVEAEQAKVFSHR.[G] | 1xHomocysteineMMTS [K9] | 0,000489972 | 1789,90877 | Brain/ FAIMS/MMTS |
| Q9QUR7 | Q9QUR7 [130-144] | [R].GQMQKPFEDASFALR.[T] | 1xHomocysteineIAA [K5] | 6,30912E-05 | 1898,88876 | Brain/ FAIMS/IAA |
| Q9QUR7 | Q9QUR7 [130-144] | [R].GQMQKPFEDASFALR.[T] | 1xHomocysteineMMTS [K5] | 0,00222764 | 1887,85502 | Brain/ No FAIMS/MMTS |
| Q9QUR7 | Q9QUR7 [130-144] | [R].GQMQKPFEDASFALR.[T] | 1xHomocysteineMMTS [K5] | 0,00222764 | 1887,85502 | Brain/ FAIMS/MMTS |
| Q9QXD6 | Q9QXD6 [207-230] | [K].GNIYSLNEGYAKDFDPAINEYLQR.[K] | 1xHomocysteineIAA [K12] | 4,32935E-05 | 2964,38325 | Liver/ FAIMS/IAA |
| Q9QXD6 | Q9QXD6 [207-230] | [K].GNIYSLNEGYAKDFDPAINEYLQR.[K] | 1xHomocysteineMMTS [K12] | 4,49774E-05 | 2953,3495 | Liver/ No FAIMS/MMTS |
| Q9QXD6 | Q9QXD6 [51-73] | [R].QAGIAQLYGIAGSTNVTGDQVKK.[L] | 1xHomocysteineMMTS [K22] | 4,49774E-05 | 2482,24285 | Liver/ No FAIMS/MMTS |
| Q9QXD6 | Q9QXD6 [292-314] | [K].AGGLATTGDKDILDIVPTEIHQK.[A] | 1xHomocysteineMMTS [K10] | 4,49774E-05 | 2555,28438 | Liver/ No FAIMS/MMTS |
| Q9QXD6 | Q9QXD6 [206-218] | [K].KGNIYSLNEGYAK.[D] | 1xHomocysteineMMTS [K1] | 4,49774E-05 | 1619,75562 | Liver/ No FAIMS/MMTS |
| Q9QXD6 | Q9QXD6 [144-158] | [K].STDEPSEKDALQPGR.[D] | 1xHomocysteineMMTS [K8] | 4,49774E-05 | 1792,78402 | Liver/ FAIMS/MMTS |
| Q9QXD6 | Q9QXD6 [206-218] | [K].KGNIYSLNEGYAK.[D] | 1xHomocysteineMMTS [K1] | 4,49774E-05 | 1619,75562 | Liver/ FAIMS/MMTS |
| Q9QXD6 | Q9QXD6 [73-87] | [K].KLDILSNDLVINMLK.[S] | 1xOxidation [M13]; 1xHomocysteineMMTS [K1] | 0,0004709 | 1908,00029 | Liver/ FAIMS/MMTS |
| Q9QXD6 | Q9QXD6 [51-73] | [R].QAGIAQLYGIAGSTNVTGDQVKK.[L] | 1xHomocysteineMMTS [K22] | 4,49774E-05 | 2482,24285 | Liver/ FAIMS/MMTS |
| Q9QXD6 | Q9QXD6 [207-230] | [K].GNIYSLNEGYAKDFDPAINEYLQR.[K] | 1xHomocysteineMMTS [K12] | 4,49774E-05 | 2953,3495 | Liver/ FAIMS/MMTS |
| Q9QXD6 | Q9QXD6 [292-314] | [K].AGGLATTGDKDILDIVPTEIHQK.[A] | 1xHomocysteineMMTS [K10] | 4,49774E-05 | 2555,28438 | Liver/ FAIMS/MMTS |
| Q9QXF8 | Q9QXF8 [231-240] | [R].DGSPGFSKFR.[L] | 1xHomocysteineMMTS [K8] | 0,00995323 | 1260,54999 | Liver/ No FAIMS/MMTS |
| Q9QXF8 | Q9QXF8 [192-201] | [K].NIYYKSDLTK.[D] | 1xHomocysteineMMTS [K5] | 0,0020903 | 1407,66468 | Liver/ FAIMS/MMTS |
| Q9QXL2 | Q9QXL2 [1603-1621] | [R].GGILKLWNVDTFVPVGEMR.[G] | 1xHomocysteineIAA [K5] | 0,00352487 | 2305,18315 | Brain/ FAIMS/IAA |
| Q9QXS1 | Q9QXS1 [2024-2039] | [R].RLEEQAALHKADIEER.[L] | 1xHomocysteineIAA [K10] | 4,32935E-05 | 2082,03966 | Brain/ FAIMS/IAA |
| Q9QXS6 | Q9QXS6 [271-291] | [K].KSESEVEEAAAIIAQRPDNPR.[E] | 1xHomocysteineIAA [K1] | 4,32935E-05 | 2484,21473 | Brain/ FAIMS/IAA |
| Q9QXZ6 | Q9QXZ6 [644-656] | [K].LTEKESECTDVCR.[S] | 2xCarbamidomethyl [C8; C12]; 1xHomocysteineIAA [K4] | 0,00892174 | 1800,7561 | Liver/ FAIMS/IAA |
| Q9QY76 | Q9QY76 [140-155] | [K].IIPTSASKTEAPAAAK.[S] | 1xHomocysteineIAA [K8] | 4,32935E-05 | 1729,9153 | Brain/ No FAIMS/IAA |
| Q9QY76 | Q9QY76 [140-155] | [K].IIPTSASKTEAPAAAK.[S] | 1xHomocysteineIAA [K8] | 4,32935E-05 | 1729,9153 | Brain/ FAIMS/IAA |
| Q9QY76 | Q9QY76 [1-17] | [-].MAKVEQVLSLEPQHELK.[F] | 1xMet-loss+Acetyl [N-Term]; 1xHomocysteineMMTS [K3] | 4,49774E-05 | 2053,04566 | Liver/ FAIMS/MMTS |
| Q9QYB1 | Q9QYB1 [207-219] | [R].NFDIPKGMTGIWR.[Y] | 1xHomocysteineIAA [K6] | 4,32935E-05 | 1708,82979 | Brain/ FAIMS/IAA |
| Q9QYB8 | Q9QYB8 [512-529] | [K].SAGPQSQLLASVIAEKSR.[S] | 1xHomocysteineIAA [K16] | 4,32935E-05 | 2016,05425 | Brain/ No FAIMS/IAA |
| Q9QYB8 | Q9QYB8 [512-529] | [K].SAGPQSQLLASVIAEKSR.[S] | 1xHomocysteineIAA [K16] | 4,32935E-05 | 2016,05425 | Brain/ FAIMS/IAA |
| Q9QYC0 | Q9QYC0 [532-548] | [R].SLVQGELVTASKAIIEK.[E] | 1xHomocysteineIAA [K12] | 4,32935E-05 | 1960,07834 | Brain/ No FAIMS/IAA |
| Q9QYC0 | Q9QYC0 [583-594] | [K].QKGSEENLDETR.[E] | 1xHomocysteineMMTS [K2] | 0,00995323 | 1568,66793 | Brain/ FAIMS/MMTS |
| Q9QYG0 | Q9QYG0 [1-25] | [-].MAELQEVQITEEKPLLPGQTPETAK.[E] | 1xMet-loss+Acetyl [N-Term]; 1xHomocysteineIAA [K13] | 4,32935E-05 | 2865,45501 | Liver/ FAIMS/IAA |
| Q9QYG0 | Q9QYG0 [1-25] | [-].MAELQEVQITEEKPLLPGQTPETAK.[E] | 1xMet-loss+Acetyl [N-Term]; 1xHomocysteineIAA [K13] | 4,32935E-05 | 2865,45501 | Brain/ No FAIMS/IAA |
| Q9QYG0 | Q9QYG0 [1-25] | [-].MAELQEVQITEEKPLLPGQTPETAK.[E] | 1xMet-loss+Acetyl [N-Term]; 1xHomocysteineIAA [K13] | 4,32935E-05 | 2865,45501 | Brain/ FAIMS/IAA |
| Q9QYG0 | Q9QYG0 [1-25] | [-].MAELQEVQITEEKPLLPGQTPETAK.[E] | 1xMet-loss+Acetyl [N-Term]; 1xHomocysteineMMTS [K13] | 4,49774E-05 | 2854,42127 | Liver/ FAIMS/MMTS |
| Q9QYJ0 | Q9QYJ0 [273-292] | [K].IGLVEALCGFQFTFKHLDAR.[Q] | 1xCarbamidomethyl [C8]; 1xHomocysteineIAA [K15] | 4,32935E-05 | 2496,25263 | Brain/ FAIMS/IAA |
| Q9QYR6 | Q9QYR6 [1287-1309] | [R].KHSPGEITGPGGHFMTSDSSLTK.[S] | 1xHomocysteineIAA [K1] | 4,32935E-05 | 2545,18098 | Brain/ FAIMS/IAA |
| Q9QYR6 | Q9QYR6 [1720-1736] | [R].SPWASDFKDFQEPLPQK.[G] | 1xHomocysteineMMTS [K8] | 0,00222764 | 2182,99362 | Brain/ FAIMS/MMTS |
| Q9QYR6 | Q9QYR6 [1531-1542] | [R].EKEGAALDQDNR.[A] | 1xHomocysteineMMTS [K2] | 0,00446656 | 1508,6468 | Brain/ FAIMS/MMTS |
| Q9QZ23 | Q9QZ23 [104-121] | [R].IEGVKSVFFGPDFITVTK.[E] | 1xHomocysteineIAA [K5] | 4,32935E-05 | 2158,12529 | Brain/ FAIMS/IAA |
| Q9QZ88 | Q9QZ88 [44-60] | [K].ESYDYLKTLAGDVHIVR.[G] | 1xHomocysteineIAA [K7] | 4,32935E-05 | 2153,06956 | Brain/ FAIMS/IAA |
| Q9QZD9 | Q9QZD9 [214-227] | [K].LFDSTTLEHQKTFR.[T] | 1xHomocysteineIAA [K11] | 0,000243434 | 1896,92726 | Brain/ FAIMS/IAA |
| Q9QZM0 | Q9QZM0 [65-79] | [R].FKSQTDQLVLIFAGK.[I] | 1xHomocysteineIAA [K2] | 9,61721E-05 | 1868,99388 | Brain/ FAIMS/IAA |
| Q9QZX7 | Q9QZX7 [41-51] | [R].NLFFKCELFQK.[T] | 1xCarbamidomethyl [C6]; 1xHomocysteineIAA [K5] | 0,00239255 | 1647,80218 | Brain/ FAIMS/IAA |
| Q9R069 | Q9R069 [498-511] | [K].SSSLMVKVTSALSR.[E] | 1xHomocysteineIAA [K7] | 0,0075237 | 1639,85059 | Brain/ FAIMS/IAA |
| Q9R0H0 | Q9R0H0 [511-525] | [K].SKEVAWNLTSVDLVR.[A] | 1xHomocysteineMMTS [K2] | 0,0077081 | 1879,94046 | Liver/ FAIMS/MMTS |
| Q9R0K7 | Q9R0K7 [752-767] | [K].HTLVKGIIDSTHTEQR.[Q] | 1xHomocysteineIAA [K5] | 4,32935E-05 | 2009,02328 | Brain/ FAIMS/IAA |
| Q9R0P5 | Q9R0P5 [70-81] | [K].HFVGMLPEKDCR.[Y] | 1xCarbamidomethyl [C11]; 1xHomocysteineIAA [K9] | 4,32935E-05 | 1662,75491 | Brain/ No FAIMS/IAA |

| | | | | | | |
|---|---|---|---|---|---|---|
| Q9R0P5 | Q9R0P5 [70-81] | [K].HFVGMLPEKDCR.[Y] | 1xCarbamidomethyl [C11]; 1xHomocysteineIAA [K9] | 4,32935E-05 | 1662,75491 | Brain/ FAIMS/IAA |
| Q9R0P5 | Q9R0P5 [82-95] | [R].YALYDASFETKESR.[K] | 1xHomocysteineMMTS [K11] | 0,000231655 | 1842,8037 | Liver/ FAIMS/MMTS |
| Q9R0P5; P45591; P18760 | Q9R0P5 [115-125]; P45591 [115-125]; P18760 [115-125] | [K].MIYASSKDAIK.[K] | 1xHomocysteineIAA [K7] | 4,32935E-05 | 1400,69123 | Liver/ FAIMS/IAA |
| Q9R0P5; P45591; P18760 | Q9R0P5 [115-125]; P45591 [115-125]; P18760 [115-125] | [K].MIYASSKDAIK.[K] | 1xOxidation [M1]; 1xHomocysteineIAA [K7] | 4,32935E-05 | 1416,68615 | Brain/ FAIMS/IAA |
| Q9R0P5; P45591; P18760 | Q9R0P5 [115-125]; P45591 [115-125]; P18760 [115-125] | [K].MIYASSKDAIK.[K] | 1xHomocysteineMMTS [K7] | 0,000152766 | 1389,65749 | Liver/ FAIMS/MMTS |
| Q9R0P5; P45591; P18760 | Q9R0P5 [115-125]; P45591 [115-125]; P18760 [115-125] | [K].MIYASSKDAIK.[K] | 1xHomocysteineMMTS [K7] | 0,000152766 | 1389,65749 | Brain/ No FAIMS/MMTS |
| Q9R0P5; P45591; P18760 | Q9R0P5 [115-125]; P45591 [115-125]; P18760 [115-125] | [K].MIYASSKDAIK.[K] | 1xHomocysteineMMTS [K7] | 0,000152766 | 1389,65749 | Brain/ FAIMS/MMTS |
| Q9R0P9 | Q9R0P9 [66-78] | [K].QIEELKGQEVSPK.[V] | 1xHomocysteineIAA [K6] | 4,32935E-05 | 1658,8418 | Brain/ No FAIMS/IAA |
| Q9R0P9 | Q9R0P9 [66-78] | [K].QIEELKGQEVSPK.[V] | 1xHomocysteineIAA [K6] | 4,32935E-05 | 1658,8418 | Brain/ FAIMS/IAA |
| Q9R0P9 | Q9R0P9 [1-15] | [-].MQLKPMEINPEMLNK.[V] | 1xMet-loss [N-Term]; 1xHomocysteineMMTS [K4] | 0,00466355 | 1847,88863 | Brain/ No FAIMS/MMTS |
| Q9R0P9 | Q9R0P9 [116-129] | [K].QFLSETEKLSPEDR.[A] | 1xHomocysteineMMTS [K8] | 4,49774E-05 | 1841,84081 | Brain/ No FAIMS/MMTS |
| Q9R0P9 | Q9R0P9 [72-83] | [K].GQEVSPKVYFMK.[Q] | 1xHomocysteineMMTS [K7] | 0,000231655 | 1575,7368 | Brain/ No FAIMS/MMTS |
| Q9R0P9 | Q9R0P9 [66-78] | [K].QIEELKGQEVSPK.[V] | 1xHomocysteineMMTS [K6] | 4,49774E-05 | 1647,80805 | Brain/ No FAIMS/MMTS |
| Q9R0P9 | Q9R0P9 [1-15] | [-].MQLKPMEINPEMLNK.[V] | 1xHomocysteineMMTS [K4] | 4,49774E-05 | 1978,92911 | Brain/ No FAIMS/MMTS |
| Q9R0P9 | Q9R0P9 [116-129] | [K].QFLSETEKLSPEDR.[A] | 1xHomocysteineMMTS [K8] | 4,49774E-05 | 1841,84081 | Brain/ FAIMS/MMTS |
| Q9R0P9 | Q9R0P9 [72-83] | [K].GQEVSPKVYFMK.[Q] | 1xHomocysteineMMTS [K7] | 0,000231655 | 1575,7368 | Brain/ FAIMS/MMTS |
| Q9R0P9 | Q9R0P9 [66-78] | [K].QIEELKGQEVSPK.[V] | 1xHomocysteineMMTS [K6] | 4,49774E-05 | 1647,80805 | Brain/ FAIMS/MMTS |
| Q9R0P9 | Q9R0P9 [16-27] | [K].VLAKLGVAGQWR.[F] | 1xHomocysteineMMTS [K4] | 4,49774E-05 | 1460,78647 | Brain/ FAIMS/MMTS |
| Q9R0P9 | Q9R0P9 [1-15] | [-].MQLKPMEINPEMLNK.[V] | 1xHomocysteineMMTS [K4] | 4,49774E-05 | 1978,92911 | Brain/ FAIMS/MMTS |
| Q9R0P9 | Q9R0P9 [65-78] | [K].KQIEELKGQEVSPK.[V] | 1xHomocysteineMMTS [K1] | 4,49774E-05 | 1775,90302 | Brain/ FAIMS/MMTS |
| Q9R0Q6 | Q9R0Q6 [104-118] | [K].WSPLENKFAVGSGAR.[L] | 1xHomocysteineIAA [K7] | 4,32935E-05 | 1792,87991 | Brain/ FAIMS/IAA |
| Q9R0Q7 | Q9R0Q7 [34-48] | [K].SKLTFSCLGGSDNFK.[H] | 1xCarbamidomethyl [C7]; 1xHomocysteineIAA [K2] | 4,32935E-05 | 1834,84623 | Brain/ FAIMS/IAA |
| Q9R0Y5 | Q9R0Y5 [10-27] | [K].IIFVVGGPGSGKGTQCEK.[I] | 1xCarbamidomethyl [C16]; 1xHomocysteineIAA [K12] | 4,32935E-05 | 2007,99904 | Brain/ No FAIMS/IAA |
| Q9R0Y5 | Q9R0Y5 [10-27] | [K].IIFVVGGPGSGKGTQCEK.[I] | 1xCarbamidomethyl [C16]; 1xHomocysteineIAA [K12] | 4,32935E-05 | 2007,99904 | Brain/ FAIMS/IAA |
| Q9R0Y5 | Q9R0Y5 [98-108] | [R].EVKQGEEFEQK.[I] | 1xHomocysteineMMTS [K3] | 0,00255629 | 1513,66614 | Liver/ FAIMS/MMTS |
| Q9R0Y5 | Q9R0Y5 [28-44] | [K].IVQKYGYTHLSTGDLLR.[A] | 1xHomocysteineMMTS [K4] | 4,49774E-05 | 2127,07254 | Brain/ FAIMS/MMTS |
| Q9R1P0 | Q9R1P0 [51-64] | [R].NIHKLLDEVFFSEK.[I] | 1xHomocysteineIAA [K4] | 0,00012645 | 1892,9575 | Brain/ FAIMS/IAA |
| Q9R1P4 | Q9R1P4 [108-122] | [R].LVSLIGSKTQIPTQR.[Y] | 1xHomocysteineIAA [K8] | 6,30912E-05 | 1815,01568 | Brain/ FAIMS/IAA |
| Q9R1P4 | Q9R1P4 [108-122] | [R].LVSLIGSKTQIPTQR.[Y] | 1xHomocysteineMMTS [K8] | 0,000929608 | 1803,98194 | Liver/ FAIMS/MMTS |
| Q9R1Q8 | Q9R1Q8 [169-196] | [R].QGQNVIGLQMGSNKGASQAGMTGYGMPR.[Q] | 1xHomocysteineIAA [K14] | 4,32935E-05 | 3012,39067 | Brain/ FAIMS/IAA |
| Q9R1Q8 | Q9R1Q8 [85-98] | [K].MAFKQMEQISQFLK.[A] | 1xHomocysteineMMTS [K4] | 4,49774E-05 | 1891,89371 | Brain/ FAIMS/MMTS |
| Q9R1Q8 | Q9R1Q8 [18-29] | [K].IEQKYDADLENK.[L] | 1xHomocysteineMMTS [K4] | 0,00225753 | 1628,72947 | Brain/ FAIMS/MMTS |
| Q9R1Q9 | Q9R1Q9 [449-463] | [R].FDDHKGPTITLTQIV.[-] | 1xHomocysteineIAA [K5] | 9,61721E-05 | 1858,93676 | Brain/ FAIMS/IAA |
| Q9R1T2 | Q9R1T2 [185-199] | [K].VAKVSQGVEDGPEAK.[R] | 1xHomocysteineIAA [K3] | 0,00421895 | 1687,83196 | Brain/ FAIMS/IAA |
| Q9R1T4 | Q9R1T4 [400-420] | [R].KAAAELLQSQGSQAGGSQTLK.[R] | 1xHomocysteineIAA [K1] | 4,32935E-05 | 2247,13977 | Brain/ No FAIMS/IAA |
| Q9R1T4 | Q9R1T4 [400-420] | [R].KAAAELLQSQGSQAGGSQTLK.[R] | 1xHomocysteineIAA [K1] | 4,32935E-05 | 2247,13977 | Brain/ FAIMS/IAA |
| Q9R1T4 | Q9R1T4 [401-421] | [K].AAAELLQSQGSQAGGSQTLKR.[D] | 1xHomocysteineMMTS [K20] | 4,49774E-05 | 2264,11217 | Brain/ FAIMS/MMTS |
| Q9R1T4 | Q9R1T4 [400-420] | [R].KAAAELLQSQGSQAGGSQTLK.[R] | 1xHomocysteineMMTS [K1] | 4,49774E-05 | 2236,10603 | Brain/ FAIMS/MMTS |
| Q9R1T4; Q8C1B7 | Q9R1T4 [310-327]; Q8C1B7 [309-326] | [K].DTDPDSKPFSLQETYEAK.[R] | 1xHomocysteineIAA [K7] | 0,00012645 | 2244,99652 | Brain/ FAIMS/IAA |
| Q9R1T4; Q8CHH9; Q8C1B7 | Q9R1T4 [164-172]; Q8CHH9 [166-174]; Q8C1B7 [163-171] | [K].SLDLVTMKK.[L] | 1xHomocysteineIAA [K] | 0,00656758 | 1208,63774 | Brain/ FAIMS/IAA |
| Q9R1V6 | Q9R1V6 [571-583] | [K].LNIEGTEKGNCGK.[D] | 1xCarbamidomethyl [C11]; 1xHomocysteineIAA [K8] | 4,32935E-05 | 1593,73595 | Brain/ FAIMS/IAA |
| Q9R1V7 | Q9R1V7 [616-626] | [K].AAGSDKFCYEK.[L] | 1xCarbamidomethyl [C8]; 1xHomocysteineIAA [K6] | 0,00117224 | 1449,61371 | Brain/ FAIMS/IAA |
| Q9R257 | Q9R257 [41-56] | [K].FATVEVTDKPVDEALR.[E] | 1xHomocysteineIAA [K9] | 4,32935E-05 | 1963,97935 | Liver/ FAIMS/IAA |
| Q9R257 | Q9R257 [41-56] | [K].FATVEVTDKPVDEALR.[E] | 1xHomocysteineIAA [K9] | 4,32935E-05 | 1963,97935 | Brain/ FAIMS/IAA |
| Q9R257 | Q9R257 [41-56] | [K].FATVEVTDKPVDEALR.[E] | 1xHomocysteineMMTS [K9] | 4,49774E-05 | 1952,94561 | Liver/ FAIMS/MMTS |
| Q9WTL7 | Q9WTL7 [183-192] | [R].FGALTAEKLR.[T] | 1xHomocysteineIAA [K8] | 6,30912E-05 | 1279,68272 | Brain/ FAIMS/IAA |
| Q9WTP6 | Q9WTP6 [52-63] | [R].AMVASGSELGKK.[L] | 1xHomocysteineMMTS [K11] | 4,49774E-05 | 1340,63709 | Liver/ FAIMS/MMTS |
| Q9WTP7 | Q9WTP7 [30-43] | [K].HFELKHLSSGDLLR.[Q] | 1xHomocysteineIAA [K5] | 0,000289025 | 1825,93776 | Liver/ FAIMS/IAA |
| Q9WTP7 | Q9WTP7 [10-26] | [R].AVIMGAPGSGKGTVSSR.[I] | 1xHomocysteineIAA [K11] | 4,32935E-05 | 1748,8782 | Brain/ No FAIMS/IAA |
| Q9WTP7 | Q9WTP7 [30-43] | [K].HFELKHLSSGDLLR.[Q] | 1xHomocysteineMMTS [K5] | 4,49774E-05 | 1814,90402 | Liver/ No FAIMS/MMTS |
| Q9WTP7 | Q9WTP7 [30-43] | [K].HFELKHLSSGDLLR.[Q] | 1xHomocysteineMMTS [K5] | 4,49774E-05 | 1814,90402 | Liver/ FAIMS/MMTS |
| Q9WTP7 | Q9WTP7 [190-203] | [K].KGVLETFSGTETNK.[I] | 1xHomocysteineMMTS [K1] | 4,49774E-05 | 1673,78732 | Liver/ FAIMS/MMTS |
| Q9WTP7 | Q9WTP7 [58-73] | [K].TFIDQGKLIPDDVMTR.[L] | 1xHomocysteineMMTS [K7] | 0,000282457 | 2011,96496 | Liver/ FAIMS/MMTS |
| Q9WTQ5 | Q9WTQ5 [1506-1523] | [K].SNKIVQSVIQTAVDQFAR.[T] | 1xHomocysteineIAA [K3] | 9,61721E-05 | 2178,13356 | Brain/ FAIMS/IAA |
| Q9WTT4 | Q9WTT4 [35-48] | [K].QAKEEAQMEVEQYR.[R] | 1xHomocysteineIAA [K3] | 4,32935E-05 | 1912,85277 | Brain/ No FAIMS/IAA |
| Q9WTT4 | Q9WTT4 [35-48] | [K].QAKEEAQMEVEQYR.[R] | 1xHomocysteineIAA [K3] | 4,32935E-05 | 1912,85277 | Brain/ FAIMS/IAA |
| Q9WTT4 | Q9WTT4 [35-48] | [K].QAKEEAQMEVEQYR.[R] | 1xHomocysteineMMTS [K3] | 0,00207528 | 1901,81903 | Brain/ FAIMS/MMTS |
| Q9WTX5 | Q9WTX5 [114-128] | [K].GLLDVTCKTVANMIK.[G] | 1xCarbamidomethyl [C7]; 1xHomocysteineIAA [K8] | 4,32935E-05 | 1836,93802 | Brain/ No FAIMS/IAA |
| Q9WTX5 | Q9WTX5 [114-128] | [K].GLLDVTCKTVANMIK.[G] | 1xCarbamidomethyl [C7]; 1xHomocysteineIAA [K8] | 4,32935E-05 | 1836,93802 | Brain/ FAIMS/IAA |

| | | | | | | |
|---|---|---|---|---|---|---|
| Q9WU78 | Q9WU78 [687-698] | [R].FQNKCSDIVFAR.[K] | 1xCarbamidomethyl [C5]; 1xHomocysteineIAA [K4] | 4,32935E-05 | 1658,77776 | Brain/ FAIMS/IAA |
| Q9WU78 | Q9WU78 [323-334] | [R].VPDLKDLDPIGK.[A] | 1xHomocysteineMMTS [K5] | 0,00586177 | 1472,74875 | Liver/ FAIMS/MMTS |
| Q9WUA3 | Q9WUA3 [665-682] | [R].KNVLGHMQQGGAPSPFDR.[N] | 1xHomocysteineIAA [K1] | 4,32935E-05 | 2113,00659 | Brain/ FAIMS/IAA |
| Q9WUB3 | Q9WUB3 [18-30] | [R].GLAGVENVSELKK.[N] | 1xHomocysteineIAA [K12] | 0,00107087 | 1517,7992 | Brain/ FAIMS/IAA |
| Q9WUC3 | Q9WUC3 [71-81] | [K].MCASSCDFVKR.[H] | 2xCarbamidomethyl [C2; C6]; 1xHomocysteineIAA [K10] | 0,000876609 | 1534,62694 | Brain/ FAIMS/IAA |
| Q9WUM5 | Q9WUM5 [95-108] | [K].HLGLPVFNTVKEAK.[E] | 1xHomocysteineIAA [K11] | 4,32935E-05 | 1726,93089 | Brain/ FAIMS/IAA |
| Q9WUR9 | Q9WUR9 [46-60] | [K].TGTEVGDVAKQYLEK.[G] | 1xHomocysteineIAA [K10] | 0,000114608 | 1811,88439 | Brain/ FAIMS/IAA |
| Q9WV54 | Q9WV54 [366-385] | [K].LTVFTTLMDVTKGQFESHLR.[D] | 1xHomocysteineIAA [K12] | 0,000177335 | 2497,25778 | Brain/ FAIMS/IAA |
| Q9WV55 | Q9WV55 [115-125] | [K].EAKPDELMDSK.[L] | 1xHomocysteineIAA [K3] | 0,000318237 | 1436,63959 | Liver/ FAIMS/IAA |
| Q9WV55 | Q9WV55 [205-221] | [R].KVAHSDKPGSTSAVSFR.[D] | 1xHomocysteineIAA [K1] | 4,32935E-05 | 1947,97052 | Brain/ No FAIMS/IAA |
| Q9WV55 | Q9WV55 [205-221] | [R].KVAHSDKPGSTSAVSFR.[D] | 1xHomocysteineMMTS [K7] | 0,00446656 | 1936,93678 | Liver/ FAIMS/MMTS |
| Q9WV55 | Q9WV55 [205-221] | [R].KVAHSDKPGSTSAVSFR.[D] | 1xHomocysteineMMTS [K7] | 0,00446656 | 1936,93678 | Brain/ FAIMS/MMTS |
| Q9WV69 | Q9WV69 [379-394] | [R].VFAMSPEEFGKLALWK.[R] | 1xHomocysteineIAA [K11] | 4,32935E-05 | 2027,0129 | Brain/ FAIMS/IAA |
| Q9WV92 | Q9WV92 [646-668] | [R].KPTEFIGGVSSTTQSWVQKLETK.[T] | 1xHomocysteineIAA [K19] | 4,32935E-05 | 2725,38654 | Brain/ FAIMS/IAA |
| Q9WV92; O70318 | Q9WV92 [407-415]; O70318 [501-509] | [K].FLTLGSKFR.[Y] | 1xHomocysteineIAA [K7] | 0,00012645 | 1242,66634 | Brain/ FAIMS/IAA |
| Q9WV92; P48193 | Q9WV92 [345-351]; P48193 [438-444] | [K].VLKISYK.[R] | 1xHomocysteineIAA [K3] | 0,00584908 | 1024,58596 | Brain/ FAIMS/IAA |
| Q9WVE8 | Q9WVE8 [51-64] | [R].IEKAYAQQLTEWAR.[R] | 1xHomocysteineIAA [K3] | 4,32935E-05 | 1880,93234 | Brain/ FAIMS/IAA |
| Q9WVL0 | Q9WVL0 [134-152] | [K].VITSGFNALEKILQSTAGK.[Y] | 1xHomocysteineIAA [K11] | 4,32935E-05 | 2151,14781 | Liver/ FAIMS/IAA |
| Q9WVL0 | Q9WVL0 [88-96] | [R].LLPQDPQKR.[A] | 1xHomocysteineMMTS [K8] | 0,00365334 | 1257,64422 | Liver/ No FAIMS/MMTS |
| Q9WVL0 | Q9WVL0 [176-191] | [R].FKVDLSPYPTISHINK.[E] | 1xHomocysteineMMTS [K2] | 4,49774E-05 | 2022,01872 | Liver/ No FAIMS/MMTS |
| Q9WVL0 | Q9WVL0 [176-191] | [R].FKVDLSPYPTISHINK.[E] | 1xHomocysteineMMTS [K2] | 4,49774E-05 | 2022,01872 | Liver/ FAIMS/MMTS |
| Q9WVL0 | Q9WVL0 [88-96] | [R].LLPQDPQKR.[A] | 1xHomocysteineMMTS [K8] | 0,00365334 | 1257,64422 | Liver/ FAIMS/MMTS |
| Q9WVL0 | Q9WVL0 [41-63] | [K].DGGQQFTEEFQTLNPMKQVPALK.[I] | 1xHomocysteineMMTS [K17] | 4,49774E-05 | 2769,30447 | Liver/ FAIMS/MMTS |
| Q9Z0E0 | Q9Z0E0 [479-492] | [R].EILIKEGAPSLLCK.[Y] | 1xCarbamidomethyl [C13]; 1xHomocysteineIAA [K5] | 4,32935E-05 | 1744,93359 | Brain/ FAIMS/IAA |
| Q9Z0F7 | Q9Z0F7 [44-58] | [K].TKENVVQSVTSVAEK.[T] | 1xHomocysteineIAA [K2] | 4,32935E-05 | 1792,91094 | Brain/ FAIMS/IAA |
| Q9Z0P4 | Q9Z0P4 [315-324] | [K].KVLGLQDTIK.[A] | 1xHomocysteineIAA [K1] | 9,61721E-05 | 1288,72933 | Brain/ FAIMS/IAA |
| Q9Z0P4 | Q9Z0P4 [91-109] | [R].LEKEIDVLEFGESAPAASK.[E] | 1xHomocysteineMMTS [K3] | 6,70238E-05 | 2196,05628 | Brain/ FAIMS/MMTS |
| Q9Z0P4; Q8BR92 | Q9Z0P4 [15-22]; Q8BR92 [11-18] | [R].LQAIAEKR.[RK] | 1xHomocysteineIAA [K7] | 0,0015926 | 1102,60374 | Brain/ No FAIMS/IAA |
| Q9Z0S1 | Q9Z0S1 [218-232] | [R].VGGAGNKIIQLIEGK.[A] | 1xHomocysteineIAA [K7] | 9,61721E-05 | 1670,9258 | Brain/ FAIMS/IAA |
| Q9Z0Y1 | Q9Z0Y1 [149-163] | [K].ALLEGYNKTTMLLSK.[Q] | 1xHomocysteineIAA [K8] | 0,000162511 | 1855,96562 | Brain/ FAIMS/IAA |
| Q9Z130 | Q9Z130 [71-85] | [R].GFGFVLFKDAASVDK.[V] | 1xHomocysteineIAA [K8] | 8,35381E-05 | 1774,88327 | Brain/ FAIMS/IAA |
| Q9Z140 | Q9Z140 [193-208] | [R].TEVVKNNLNPSWEPFR.[L] | 1xHomocysteineIAA [K5] | 0,00012645 | 2104,02803 | Brain/ FAIMS/IAA |
| Q9Z172; P61957 | Q9Z172 [36-44]; P61957 [37-45] | [R].HTPLSKLMK.[A] | 1xHomocysteineIAA [K6] | 0,000114608 | 1228,65406 | Brain/ FAIMS/IAA |
| Q9Z172; P61957 | Q9Z172 [21-34]; P61957 [22-35] | [K].VAGQDGSVVQFKIK.[R] | 1xHomocysteineMMTS [K12] | 4,49774E-05 | 1638,83421 | Liver/ FAIMS/MMTS |
| Q9Z1B3 | Q9Z1B3 [582-596] | [K].SPVEFVEYNKMQLSR.[I] | 1xHomocysteineIAA [K10] | 4,32935E-05 | 2000,95684 | Brain/ FAIMS/IAA |
| Q9Z1F9 | Q9Z1F9 [265-274] | [R].YLLTMDKLWR.[K] | 1xHomocysteineIAA [K7] | 4,32935E-05 | 1512,77015 | Brain/ FAIMS/IAA |
| Q9Z1G4 | Q9Z1G4 [74-92] | [R].KANIPIMDTGENPEVPFPR.[D] | 1xHomocysteineIAA [K1] | 4,32935E-05 | 2299,12095 | Brain/ FAIMS/IAA |
| Q9Z1P6 | Q9Z1P6 [81-93] | [K].AAESSAMAATEKK.[A] | 1xHomocysteineIAA [K12] | 0,00173159 | 1468,67704 | Brain/ FAIMS/IAA |
| Q9Z1S5 | Q9Z1S5 [281-301] | [R].KTPWGIIEVENLNHCEFALLR.[D] | 1xCarbamidomethyl [C15]; 1xHomocysteineIAA [K1] | 4,32935E-05 | 2713,35889 | Brain/ FAIMS/IAA |
| Q9Z1Z0 | Q9Z1Z0 [895-904] | [K].LDLEVTDSKK.[E] | 1xHomocysteineMMTS [K9] | 0,00409028 | 1310,63305 | Liver/ No FAIMS/MMTS |
| Q9Z204 | Q9Z204 [206-217] | [K].QKVDSLLESLEK.[I] | 1xHomocysteineIAA [K2] | 4,32935E-05 | 1562,80943 | Brain/ FAIMS/IAA |
| Q9Z204 | Q9Z204 [241-252] | [K].SEEEQSSASVKK.[D] | 1xHomocysteineMMTS [K11] | 0,00924953 | 1471,64032 | Liver/ FAIMS/MMTS |
| Q9Z218 | Q9Z218 [360-371] | [K].KHEDESEAWLHR.[Q] | 1xHomocysteineIAA [K1] | 4,32935E-05 | 1710,76528 | Brain/ No FAIMS/IAA |
| Q9Z218 | Q9Z218 [360-371] | [K].KHEDESEAWLHR.[Q] | 1xHomocysteineIAA [K1] | 4,32935E-05 | 1710,76528 | Brain/ FAIMS/IAA |
| Q9Z2H5 | Q9Z2H5 [479-491] | [K].IKELKPEQETTPR.[H] | 1xHomocysteineIAA [K2] | 0,000318237 | 1742,91054 | Brain/ FAIMS/IAA |
| Q9Z2I0 | Q9Z2I0 [592-605] | [K].TGEEKYIEESAASK.[R] | 1xHomocysteineIAA [K5] | 9,61721E-05 | 1715,77926 | Brain/ FAIMS/IAA |
| Q9Z2I0 | Q9Z2I0 [257-273] | [K].LELAKFLQDTIEEMALK.[N] | 1xHomocysteineMMTS [K5] | 4,49774E-05 | 2155,08475 | Liver/ No FAIMS/MMTS |
| Q9Z2I0 | Q9Z2I0 [597-606] | [K].YIEESAASKR.[L] | 1xHomocysteineMMTS [K9] | 0,00550312 | 1316,59733 | Liver/ FAIMS/MMTS |
| Q9Z2I0 | Q9Z2I0 [451-462] | [K].VAEVEGEKVDNK.[A] | 1xHomocysteineMMTS [K8] | 0,00804194 | 1479,68179 | Liver/ FAIMS/MMTS |
| Q9Z2I0 | Q9Z2I0 [592-605] | [K].TGEEKYIEESAASK.[R] | 1xHomocysteineMMTS [K5] | 0,00110986 | 1704,74551 | Liver/ FAIMS/MMTS |
| Q9Z2I0 | Q9Z2I0 [257-273] | [K].LELAKFLQDTIEEMALK.[N] | 1xHomocysteineMMTS [K5] | 4,49774E-05 | 2155,08475 | Brain/ No FAIMS/MMTS |
| Q9Z2I8 | Q9Z2I8 [76-84] | [R].LNAKEIVLK.[A] | 1xHomocysteineMMTS [K4] | 0,00563517 | 1190,66356 | Liver/ No FAIMS/MMTS |
| Q9Z2I8 | Q9Z2I8 [408-425] | [K].SSGLPITSAVDLEDAAKK.[A] | 1xHomocysteineMMTS [K17] | 4,49774E-05 | 1964,96674 | Liver/ FAIMS/MMTS |
| Q9Z2I8 | Q9Z2I8 [60-74] | [R].FFVANTAKEALEAAK.[R] | 1xHomocysteineMMTS [K8] | 4,49774E-05 | 1772,87099 | Liver/ FAIMS/MMTS |
| Q9Z2I8 | Q9Z2I8 [76-84] | [R].LNAKEIVLK.[A] | 1xHomocysteineMMTS [K4] | 0,00563517 | 1190,66356 | Liver/ FAIMS/MMTS |
| Q9Z2I8 | Q9Z2I8 [394-424] | [R].LEGTNVQEAQNILKSSGLPITSAVDLEDAAK.[K] | 1xHomocysteineMMTS [K14] | 4,49774E-05 | 3374,6818 | Liver/ FAIMS/MMTS |
| Q9Z2I8 | Q9Z2I8 [48-56] | [K].KLMSEHGVR.[V] | 1xHomocysteineMMTS [K1] | 0,000507453 | 1219,57443 | Liver/ FAIMS/MMTS |
| Q9Z2I9 | Q9Z2I9 [121-139] | [K].IVFSPEEAKAVSSQMIGQK.[L] | 1xHomocysteineIAA [K9] | 4,32935E-05 | 2223,1148 | Brain/ No FAIMS/IAA |
| Q9Z2I9 | Q9Z2I9 [121-139] | [K].IVFSPEEAKAVSSQMIGQK.[L] | 1xHomocysteineIAA [K9] | 4,32935E-05 | 2223,1148 | Brain/ FAIMS/IAA |
| Q9Z2I9 | Q9Z2I9 [439-451] | [K].MVVKLSEIVTLAK.[E] | 1xHomocysteineMMTS [K4] | 0,0055918 | 1593,87765 | Liver/ FAIMS/MMTS |
| Q9Z2I9 | Q9Z2I9 [413-426] | [R].VDDAKALIADSGLK.[I] | 1xHomocysteineMMTS [K5] | 0,00869467 | 1578,78659 | Brain/ FAIMS/MMTS |
| Q9Z2Q6 | Q9Z2Q6 [58-73] | [K].STLVHSLFLTDLYKDR.[K] | 1xHomocysteineIAA [K14] | 4,32935E-05 | 2082,06884 | Brain/ FAIMS/IAA |
| Q9Z2U0 | Q9Z2U0 [110-117] | [R].YIASLKQR.[Y] | 1xHomocysteineIAA [K6] | 4,32935E-05 | 1152,61939 | Brain/ FAIMS/IAA |
| Q9Z2U1 | Q9Z2U1 [197-209] | [K].SSLIILKQVMEEK.[L] | 1xHomocysteineIAA [K7] | 4,32935E-05 | 1691,90704 | Brain/ FAIMS/IAA |
| Q9Z2X1 | Q9Z2X1 [219-230] | [R].YIGIVKQAGLDR.[M] | 1xHomocysteineIAA [K6] | 0,000243434 | 1506,80971 | Brain/ FAIMS/IAA |
| Q9Z2X1 | Q9Z2X1 [219-230] | [R].YIGIVKQAGLDR.[M] | 1xHomocysteineMMTS [K6] | 0,00109439 | 1495,77597 | Liver/ FAIMS/MMTS |
| Q9Z2Y3 | Q9Z2Y3 [336-348] | [R].SNLKTLLEILDGK.[I] | 1xHomocysteineIAA [K4] | 4,32935E-05 | 1617,88802 | Brain/ No FAIMS/IAA |
| Q9Z2Y3 | Q9Z2Y3 [336-348] | [R].SNLKTLLEILDGK.[I] | 1xHomocysteineIAA [K4] | 4,32935E-05 | 1617,88802 | Brain/ FAIMS/IAA |
| Q9Z2Y8 | Q9Z2Y8 [64-81] | [R].TFGENYVQELLEKASNPK.[I] | 1xHomocysteineIAA [K13] | 0,000988467 | 2241,08561 | Brain/ FAIMS/IAA |

| Master Protein Accessions |
|---|
| A2A432 |
| A2ABU4 |
| A2AEY4 |
| A2AJI0 |
| A2AS89 |
| A2ASS6 |
| A6X935 |
| B2RSH2 |
| B2RXS4 |
| B5X0G2 |
| B9EKR1 |
| D3YVF0 |
| D3Z6Q9 |
| D3Z7P3 |
| E9PUL5 |
| E9PV24 |
| E9PVX6 |
| E9Q5G3 |
| E9Q912 |
| F8VPU2 |
| G3X982 |
| G5E829 |
| G5E8K5 |
| O08532 |
| O08539 |
| O08553 |
| O08585 |
| O08599 |
| O08677 |
| O08709 |
| O08749 |
| O08756 |
| O08795 |
| O08917 |
| O09061 |
| O09111 |
| O09173 |
| O35136 |
| O35143 |
| O35215 |
| O35226 |
| O35295 |
| O35381 |
| O35405 |
| O35459 |
| O35490 |
| O35526 |
| O35633 |

| |
|---|
| O35639 |
| O35658 |
| O35685 |
| O35737 |
| O35943 |
| O54901 |
| O54983 |
| O54991 |
| O55013 |
| O55022 |
| O55042 |
| O55091 |
| O55100 |
| O55126 |
| O55131 |
| O55142 |
| O70161 |
| O70251 |
| O70318 |
| O70400 |
| O70435 |
| O70439 |
| O70443 |
| O70493 |
| O70591 |
| O88342 |
| O88343 |
| O88448 |
| O88485 |
| O88531 |
| O88569 |
| O88587 |
| O88643 |
| O88685 |
| O88696 |
| O88737 |
| O88844 |
| O88935 |
| O88952 |
| O88998 |
| O89023 |
| O89053 |
| O89112 |
| P00329 |
| P00405 |
| P00493 |
| P00920 |
| P01831 |
| P01837 |
| P01872 |

| |
|---|
| P01878 |
| P01887 |
| P01898 |
| P01942 |
| P02088 |
| P02089 |
| P02468 |
| P03930 |
| P03995 |
| P04247 |
| P04370 |
| P04925 |
| P04939 |
| P05063 |
| P05064 |
| P05132 |
| P05201 |
| P05202 |
| P05214 |
| P05784 |
| P05977 |
| P06151 |
| P06728 |
| P06745 |
| P06837 |
| P07309 |
| P07310 |
| P07724 |
| P07758 |
| P07759 |
| P07901 |
| P08003 |
| P08113 |
| P08226 |
| P08228 |
| P08249 |
| P08551 |
| P08553 |
| P08752 |
| P09041 |
| P09103 |
| P09411 |
| P09528 |
| P09542 |
| P09671 |
| P09813 |
| P0DN34 |
| P0DP27 |
| P10126 |
| P10605 |

| |
|---|
| P10637 |
| P10639 |
| P10648 |
| P10649 |
| P10852 |
| P10922 |
| P11352 |
| P11404 |
| P11438 |
| P11499 |
| P11588 |
| P11589 |
| P11627 |
| P11679 |
| P11725 |
| P11798 |
| P11983 |
| P12023 |
| P12658 |
| P12710 |
| P12787 |
| P12960 |
| P12970 |
| P13020 |
| P13595 |
| P13707 |
| P14094 |
| P14115 |
| P14131 |
| P14148 |
| P14152 |
| P14206 |
| P14211 |
| P14231 |
| P14602 |
| P14824 |
| P14869 |
| P14873 |
| P15105 |
| P15116 |
| P15209 |
| P15532 |
| P16015 |
| P16125 |
| P16330 |
| P16331 |
| P16388 |
| P16460 |
| P16546 |
| P16627 |

| |
|---|
| P16858 |
| P17047 |
| P17156 |
| P17182 |
| P17183 |
| P17426 |
| P17427 |
| P17563 |
| P17665 |
| P17710 |
| P17717 |
| P17742 |
| P17751 |
| P18052 |
| P18242 |
| P18572 |
| P18760 |
| P18872 |
| P19096 |
| P19157 |
| P19246 |
| P19253 |
| P19536 |
| P19783 |
| P20029 |
| P20065 |
| P20108 |
| P20152 |
| P20357 |
| P20801 |
| P20917 |
| P21107 |
| P21278 |
| P21300 |
| P21550 |
| P21614 |
| P21995 |
| P22599 |
| P23242 |
| P23506 |
| P23819 |
| P23927 |
| P23953 |
| P24270 |
| P24369 |
| P24457 |
| P24472 |
| P24549 |
| P25444 |
| P25688 |

| |
|---|
| P26041 |
| P26043 |
| P26350 |
| P26443 |
| P26645 |
| P26883 |
| P27546 |
| P27573 |
| P27600 |
| P27659 |
| P27773 |
| P28352 |
| P28474 |
| P28652 |
| P28663 |
| P28665 |
| P28667 |
| P28740 |
| P28843 |
| P29391 |
| P29595 |
| P29758 |
| P30115 |
| P30275 |
| P30416 |
| P31001 |
| P31648 |
| P31650 |
| P31786 |
| P31938 |
| P32020 |
| P32037 |
| P32848 |
| P34022 |
| P34884 |
| P34927 |
| P35278 |
| P35282 |
| P35486 |
| P35505 |
| P35564 |
| P35700 |
| P35762 |
| P35802 |
| P35803 |
| P35980 |
| P36536 |
| P38060 |
| P38647 |
| P39053 |

| |
|---|
| P39054 |
| P40124 |
| P40142 |
| P40237 |
| P40240 |
| P40336 |
| P40936 |
| P41105 |
| P42125 |
| P42669 |
| P42932 |
| P43006 |
| P43024 |
| P43274 |
| P45376 |
| P45591 |
| P45878 |
| P45952 |
| P46096 |
| P46097 |
| P46425 |
| P46460 |
| P46471 |
| P46638 |
| P46660 |
| P47708 |
| P47738 |
| P47753 |
| P47757 |
| P47857 |
| P47867 |
| P47911 |
| P47915 |
| P47962 |
| P47963 |
| P48024 |
| P48036 |
| P48320 |
| P48455 |
| P48678 |
| P48722 |
| P48771 |
| P48774 |
| P48962 |
| P49312 |
| P49429 |
| P50114 |
| P50172 |
| P50247 |
| P50396 |

| |
|---|
| P50431 |
| P50516 |
| P50518 |
| P50571 |
| P50580 |
| P51150 |
| P51410 |
| P51859 |
| P51863 |
| P51881 |
| P51910 |
| P52196 |
| P52480 |
| P52503 |
| P52760 |
| P53395 |
| P53564 |
| P53810 |
| P53994 |
| P54071 |
| P54099 |
| P54227 |
| P54728 |
| P54869 |
| P55050 |
| P55088 |
| P55264 |
| P56135 |
| P56375 |
| P56376 |
| P56382 |
| P56391 |
| P56393 |
| P56395 |
| P56399 |
| P56480 |
| P56546 |
| P56564 |
| P56565 |
| P56959 |
| P57746 |
| P57759 |
| P57776 |
| P57780 |
| P58252 |
| P58281 |
| P58771 |
| P58774 |
| P59648 |
| P59999 |

| |
|---|
| P60202 |
| P60469 |
| P60521 |
| P60710 |
| P60755 |
| P60761 |
| P60764 |
| P60766 |
| P60867 |
| P60879 |
| P60904 |
| P61021 |
| P61027 |
| P61082 |
| P61087 |
| P61089 |
| P61164 |
| P61205 |
| P61255 |
| P61264 |
| P61294 |
| P61329 |
| P61358 |
| P61458 |
| P61514 |
| P61750 |
| P61922 |
| P61924 |
| P61971 |
| P61979 |
| P61982 |
| P62073 |
| P62141 |
| P62196 |
| P62242 |
| P62245 |
| P62259 |
| P62267 |
| P62270 |
| P62274 |
| P62281 |
| P62301 |
| P62305 |
| P62317 |
| P62331 |
| P62484 |
| P62627 |
| P62631 |
| P62702 |
| P62715 |

| |
|---|
| P62717 |
| P62737 |
| P62746 |
| P62748 |
| P62751 |
| P62754 |
| P62761 |
| P62774 |
| P62806 |
| P62814 |
| P62821 |
| P62823 |
| P62827 |
| P62830 |
| P62843 |
| P62849 |
| P62852 |
| P62858 |
| P62862 |
| P62874 |
| P62878 |
| P62880 |
| P62889 |
| P62897 |
| P62900 |
| P62908 |
| P62911 |
| P62918 |
| P62960 |
| P62962 |
| P62983 |
| P63001 |
| P63011 |
| P63017 |
| P63024 |
| P63028 |
| P63030 |
| P63038 |
| P63040 |
| P63054 |
| P63085 |
| P63087 |
| P63094 |
| P63101 |
| P63158 |
| P63168 |
| P63213 |
| P63242 |
| P63260 |
| P63276 |

| |
|---|
| P63318 |
| P63321 |
| P63323 |
| P63328 |
| P67778 |
| P68037 |
| P68040 |
| P68134 |
| P68254 |
| P68368 |
| P68369 |
| P68372 |
| P68404 |
| P68404 |
| P68510 |
| P70168 |
| P70202 |
| P70290 |
| P70296 |
| P70297 |
| P70297 |
| P70333 |
| P70349 |
| P70404 |
| P70414 |
| P70441 |
| P70670 |
| P70691 |
| P70694 |
| P80313 |
| P80314 |
| P80315 |
| P83917 |
| P83940 |
| P84075 |
| P84086 |
| P84089 |
| P84091 |
| P84096 |
| P84099 |
| P84244 |
| P97300 |
| P97315 |
| P97328 |
| P97343 |
| P97351 |
| P97352 |
| P97370 |
| P97371 |
| P97384 |

| |
|---|
| P97427 |
| P97449 |
| P97450 |
| P97457 |
| P97493 |
| P97496 |
| P97765 |
| P97797 |
| P97807 |
| P97952 |
| P99024 |
| P99026 |
| P99027 |
| P99028 |
| P99029 |
| Q00493 |
| Q00623 |
| Q00897 |
| Q00898 |
| Q01279 |
| Q01768 |
| Q01853 |
| Q02053 |
| Q02248 |
| Q02819 |
| Q03265 |
| Q03517 |
| Q03958 |
| Q04447 |
| Q04736 |
| Q05186 |
| Q05816 |
| Q06185 |
| Q06890 |
| Q07076 |
| Q08331 |
| Q11011 |
| Q2PFD7 |
| Q30KP6 |
| Q3TCN2 |
| Q3TTY5 |
| Q3TXX4 |
| Q3U0V1 |
| Q3UBX0 |
| Q3UES3 |
| Q3UGC7 |
| Q3UHL1 |
| Q3UI43 |
| Q3UL36 |
| Q3ULJ0 |

| |
|---|
| Q3UNH4 |
| Q3UPI1 |
| Q3URK3 |
| Q3UZ18 |
| Q3UZP4 |
| Q3V132 |
| Q4KMM3 |
| Q56A07 |
| Q5DTL9 |
| Q5FW60 |
| Q5FWK3 |
| Q5GH66 |
| Q5M8N0 |
| Q5QNQ9 |
| Q5SQX6 |
| Q5SRX1 |
| Q5SYD0 |
| Q5XG73 |
| Q60597 |
| Q60598 |
| Q60605 |
| Q60625 |
| Q60631 |
| Q60634 |
| Q60668 |
| Q60771 |
| Q60829 |
| Q60864 |
| Q60870 |
| Q60902 |
| Q60930 |
| Q60931 |
| Q60932 |
| Q61016 |
| Q61081 |
| Q61166 |
| Q61171 |
| Q61176 |
| Q61206 |
| Q61207 |
| Q61282 |
| Q61292 |
| Q61316 |
| Q61330 |
| Q61335 |
| Q61361 |
| Q61414 |
| Q61425 |
| Q61490 |
| Q61548 |

| |
|---|
| Q61553 |
| Q61598 |
| Q61644 |
| Q61646 |
| Q61696 |
| Q61699 |
| Q61735 |
| Q61753 |
| Q61768 |
| Q61792 |
| Q61838 |
| Q61879 |
| Q61885 |
| Q61937 |
| Q61990 |
| Q62048 |
| Q62059 |
| Q62188 |
| Q62189 |
| Q62261 |
| Q62277 |
| Q62283 |
| Q62318 |
| Q62376 |
| Q62418 |
| Q62420 |
| Q62425 |
| Q62426 |
| Q62442 |
| Q62443 |
| Q62465 |
| Q63810 |
| Q63836 |
| Q63880 |
| Q63912 |
| Q640R3 |
| Q64105 |
| Q64288 |
| Q64324 |
| Q64332 |
| Q64374 |
| Q64433 |
| Q64442 |
| Q64471 |
| Q64478 |
| Q64520 |
| Q64521 |
| Q64523 |
| Q64525 |
| Q68ED7 |

| |
|---|
| Q68EF4 |
| Q68FD5 |
| Q6GQX6 |
| Q6IRU2 |
| Q6IRU5 |
| Q6P069 |
| Q6P8X1 |
| Q6PDL0 |
| Q6PDY2 |
| Q6PER3 |
| Q6PHN9 |
| Q6PHS9 |
| Q6PHZ8 |
| Q6PIC6 |
| Q6PIE5 |
| Q6Q477 |
| Q6WKZ7 |
| Q6X893 |
| Q6ZQ38 |
| Q6ZWN5 |
| Q6ZWV3 |
| Q6ZWV7 |
| Q71M36 |
| Q76MZ3 |
| Q78IK2 |
| Q78JT3 |
| Q78ZA7 |
| Q7TMF3 |
| Q7TMM9 |
| Q7TNP2 |
| Q7TQD2 |
| Q7TQF7 |
| Q7TQI3 |
| Q7TSJ2 |
| Q80SW1 |
| Q80SZ7 |
| Q80TB8 |
| Q80TJ1 |
| Q80TL4 |
| Q80U72 |
| Q80UK0 |
| Q80UW2 |
| Q80V26 |
| Q80VP1 |
| Q80X56 |
| Q80X90 |
| Q80YN3 |
| Q80Z24 |
| Q80ZJ1 |
| Q810U3 |

| |
|---|
| Q810U4 |
| Q811F1 |
| Q8BFR5 |
| Q8BFU2 |
| Q8BFU3 |
| Q8BFZ3 |
| Q8BFZ9 |
| Q8BG05 |
| Q8BG39 |
| Q8BGN8 |
| Q8BGQ7 |
| Q8BGR9 |
| Q8BGU5 |
| Q8BGY2 |
| Q8BGZ1 |
| Q8BH95 |
| Q8BHN3 |
| Q8BI22 |
| Q8BJI1 |
| Q8BJU0 |
| Q8BK48 |
| Q8BK64 |
| Q8BKX1 |
| Q8BKZ9 |
| Q8BL66 |
| Q8BLK3 |
| Q8BLQ9 |
| Q8BMF4 |
| Q8BMK4 |
| Q8BMS1 |
| Q8BN59 |
| Q8BNY6 |
| Q8BP47 |
| Q8BP67 |
| Q8BP92 |
| Q8BSY0 |
| Q8BU31 |
| Q8BVE3 |
| Q8BVI4 |
| Q8BVP1 |
| Q8BWF0 |
| Q8BWR2 |
| Q8BWT1 |
| Q8BYI9 |
| Q8BYM5 |
| Q8C0E2 |
| Q8C0M9 |
| Q8C196 |
| Q8C1B7 |
| Q8C419 |

| |
|---|
| Q8C854 |
| Q8C886 |
| Q8C8R3 |
| Q8C8T8 |
| Q8C8V1 |
| Q8CAQ8 |
| Q8CAY6 |
| Q8CC35 |
| Q8CCT4 |
| Q8CDN6 |
| Q8CGP1 |
| Q8CHC4 |
| Q8CHH5 |
| Q8CHH9 |
| Q8CHP8 |
| Q8CHT0 |
| Q8CI43 |
| Q8CI94 |
| Q8JZK9 |
| Q8JZR6 |
| Q8JZS0 |
| Q8JZW4 |
| Q8K021 |
| Q8K0E8 |
| Q8K0T0 |
| Q8K0U4 |
| Q8K183 |
| Q8K1M6 |
| Q8K1Z0 |
| Q8K221 |
| Q8K2B3 |
| Q8K2T1 |
| Q8K3C3 |
| Q8K3J1 |
| Q8K4F5 |
| Q8K4Z3 |
| Q8K596 |
| Q8QZR3 |
| Q8QZT1 |
| Q8R001 |
| Q8R0A5 |
| Q8R0A7 |
| Q8R0F8 |
| Q8R164 |
| Q8R1I1 |
| Q8R1Q8 |
| Q8R2E6 |
| Q8R317 |
| Q8R326 |
| Q8R366 |

| |
|---|
| Q8R3V5 |
| Q8R404 |
| Q8R570 |
| Q8R5H6 |
| Q8R5I6 |
| Q8R5M8 |
| Q8VBW5 |
| Q8VBY2 |
| Q8VC26 |
| Q8VC30 |
| Q8VCC2 |
| Q8VCH0 |
| Q8VCT4 |
| Q8VCU1 |
| Q8VCW8 |
| Q8VD04 |
| Q8VD37 |
| Q8VDN2 |
| Q8VDQ8 |
| Q8VE33 |
| Q8VEH3 |
| Q8VEK0 |
| Q8VEM8 |
| Q8VHW2 |
| Q8VIJ6 |
| Q91V12 |
| Q91V14 |
| Q91V41 |
| Q91V61 |
| Q91V76 |
| Q91V92 |
| Q91VD9 |
| Q91VN4 |
| Q91VR2 |
| Q91VR5 |
| Q91VR7 |
| Q91VR8 |
| Q91VS7 |
| Q91VZ6 |
| Q91WD5 |
| Q91WG0 |
| Q91WJ8 |
| Q91WS0 |
| Q91WU0 |
| Q91X72 |
| Q91X83 |
| Q91X97 |
| Q91XD4 |
| Q91XE0 |
| Q91XM9 |

| |
|---|
| Q91XV3 |
| Q91Y97 |
| Q91YT0 |
| Q91Z53 |
| Q91ZA3 |
| Q91ZX7 |
| Q91ZZ3 |
| Q921G7 |
| Q921H8 |
| Q921I1 |
| Q921M7 |
| Q922B1 |
| Q922F4 |
| Q922J6 |
| Q922Q1 |
| Q922R8 |
| Q923D2 |
| Q93092 |
| Q99020 |
| Q99J08 |
| Q99J85 |
| Q99J99 |
| Q99JB2 |
| Q99JI6; |
| Q99JP6 |
| Q99JP7 |
| Q99JW2 |
| Q99JX3 |
| Q99JY9 |
| Q99K51 |
| Q99K85 |
| Q99KB8 |
| Q99KF1 |
| Q99KI0 |
| Q99KJ8 |
| Q99KP3 |
| Q99KR3 |
| Q99KR7 |
| Q99L13 |
| Q99L47 |
| Q99LC5 |
| Q99LD4 |
| Q99LI8 |
| Q99LX0 |
| Q99LY9 |
| Q99MP8 |
| Q99N28 |
| Q99P30 |
| Q99P72 |
| Q99PJ0 |

| |
|---|
| Q99PL5 |
| Q99PT1 |
| Q9CPP6 |
| Q9CPQ1 |
| Q9CPQ8 |
| Q9CPR4 |
| Q9CPU0 |
| Q9CPW0 |
| Q9CPW2 |
| Q9CPW4 |
| Q9CPW5 |
| Q9CPY7 |
| Q9CQ19 |
| Q9CQ62 |
| Q9CQ69 |
| Q9CQ75 |
| Q9CQ92 |
| Q9CQA3 |
| Q9CQC7 |
| Q9CQH3 |
| Q9CQH7 |
| Q9CQI3 |
| Q9CQI6 |
| Q9CQJ6 |
| Q9CQK7 |
| Q9CQM5 |
| Q9CQQ7 |
| Q9CQR2 |
| Q9CQR4 |
| Q9CQV6 |
| Q9CQV8 |
| Q9CQW1 |
| Q9CQW2 |
| Q9CQX2 |
| Q9CQZ5 |
| Q9CQZ6 |
| Q9CR21 |
| Q9CR51 |
| Q9CR57 |
| Q9CR62 |
| Q9CR68 |
| Q9CR95 |
| Q9CRB6 |
| Q9CRB9 |
| Q9CRC3 |
| Q9CVB6 |
| Q9CWF2 |
| Q9CWH1 |
| Q9CWJ9 |
| Q9CWK8 |

| |
|---|
| Q9CWS0 |
| Q9CWZ7 |
| Q9CXW3 |
| Q9CXW4 |
| Q9CYZ2 |
| Q9CZ13 |
| Q9CZ44 |
| Q9CZC8 |
| Q9CZM2 |
| Q9CZT8 |
| Q9CZU6 |
| Q9CZX8 |
| Q9CZY3 |
| Q9D051 |
| Q9D0E1 |
| Q9D0F3 |
| Q9D0J8 |
| Q9D0K2 |
| Q9D0M3 |
| Q9D0M5 |
| Q9D0S9 |
| Q9D0W5 |
| Q9D164 |
| Q9D172 |
| Q9D1A2 |
| Q9D1C8 |
| Q9D1E6 |
| Q9D1G1 |
| Q9D1K2 |
| Q9D1P4 |
| Q9D1R9 |
| Q9D2G2 |
| Q9D2M8 |
| Q9D2P8 |
| Q9D312 |
| Q9D394 |
| Q9D3D9 |
| Q9D4B2 |
| Q9D4D4 |
| Q9D4J1 |
| Q9D6F9 |
| Q9D6J5 |
| Q9D6J6 |
| Q9D6K5 |
| Q9D6M3 |
| Q9D6R2 |
| Q9D6Y7 |
| Q9D819 |
| Q9D855 |
| Q9D892 |

| |
|---|
| Q9D8B3 |
| Q9D8B4 |
| Q9D8B7 |
| Q9D8E6 |
| Q9D8N0 |
| Q9D8Y0 |
| Q9DAK9 |
| Q9DAS9 |
| Q9DAW9 |
| Q9DB05 |
| Q9DB15 |
| Q9DB20 |
| Q9DB77 |
| Q9DBB8 |
| Q9DBG3 |
| Q9DBH5 |
| Q9DBJ1 |
| Q9DBP5 |
| Q9DC07 |
| Q9DC51 |
| Q9DC69 |
| Q9DCJ5 |
| Q9DCL9 |
| Q9DCM0 |
| Q9DCS9 |
| Q9DCT2 |
| Q9DCT8 |
| Q9DCU2 |
| Q9DCW4 |
| Q9DCX2 |
| Q9EPJ9 |
| Q9EQ20 |
| Q9EQS3 |
| Q9EQU5 |
| Q9ERD7 |
| Q9ERL7 |
| Q9ERT9 |
| Q9ES97 |
| Q9ESM3 |
| Q9JHI5 |
| Q9JHU4 |
| Q9JHW2 |
| Q9JIA1 |
| Q9JII6 |
| Q9JIS5 |
| Q9JIS8 |
| Q9JIW9 |
| Q9JIY0 |
| Q9JJI8 |
| Q9JJV2 |

| |
|---|
| Q9JKB1 |
| Q9JKB3 |
| Q9JKC6 |
| Q9JKD3 |
| Q9JKF6 |
| Q9JKK7 |
| Q9JKL5 |
| Q9JKR6 |
| Q9JL62 |
| Q9JLM8 |
| Q9JLZ3 |
| Q9JM63 |
| Q9JM76 |
| Q9JMA1 |
| Q9JMD3 |
| Q9JMG7 |
| Q9QUI0 |
| Q9QUK6 |
| Q9QUM9 |
| Q9QUP5 |
| Q9QUR7 |
| Q9QXD6 |
| Q9QXF8 |
| Q9QXL2 |
| Q9QXS1 |
| Q9QXS6 |
| Q9QXZ6 |
| Q9QY76 |
| Q9QYB1 |
| Q9QYB8 |
| Q9QYC0 |
| Q9QYG0 |
| Q9QYJ0 |
| Q9QYR6 |
| Q9QZ23 |
| Q9QZ88 |
| Q9QZD9 |
| Q9QZM0 |
| Q9QZX7 |
| Q9R069 |
| Q9R0H0 |
| Q9R0K7 |
| Q9R0P5 |
| Q9R0P9 |
| Q9R0Q6 |
| Q9R0Q7 |
| Q9R0Y5 |
| Q9R1P0 |
| Q9R1P4 |
| Q9R1Q8 |

| |
|---|
| Q9R1Q9 |
| Q9R1T2 |
| Q9R1T4 |
| Q9R1V6 |
| Q9R1V7 |
| Q9R257 |
| Q9WTL7 |
| Q9WTP6 |
| Q9WTP7 |
| Q9WTQ5 |
| Q9WTT4 |
| Q9WTX5 |
| Q9WU78 |
| Q9WUA3 |
| Q9WUB3 |
| Q9WUC3 |
| Q9WUM5 |
| Q9WUR9 |
| Q9WV54 |
| Q9WV55 |
| Q9WV69 |
| Q9WV92 |
| Q9WVE8 |
| Q9WVL0 |
| Q9Z0E0 |
| Q9Z0F7 |
| Q9Z0P4 |
| Q9Z0S1 |
| Q9Z0Y1 |
| Q9Z130 |
| Q9Z140 |
| Q9Z172 |
| Q9Z1B3 |
| Q9Z1F9 |
| Q9Z1G4 |
| Q9Z1P6 |
| Q9Z1S5 |
| Q9Z1Z0 |
| Q9Z204 |
| Q9Z218 |
| Q9Z2H5 |
| Q9Z2I0 |
| Q9Z2I8 |
| Q9Z2I9 |
| Q9Z2Q6 |
| Q9Z2U0 |
| Q9Z2U1 |
| Q9Z2X1 |
| Q9Z2Y3 |
| Q9Z2Y8 |

| From | To | Species | David Gene Name |
|---|---|---|---|
| P16330 | 12799 | Mus musculus | 2',3'-cyclic nucleotide 3' phosphodiesterase(Cnp) |
| Q9CQ62 | 67460 | Mus musculus | 2,4-dienoyl CoA reductase 1, mitochondrial(Decr1) |
| Q6PDY2 | 211488 | Mus musculus | 2-aminoethanethiol dioxygenase(Ado) |
| Q9Z0S1 | 23827 | Mus musculus | 3'(2'), 5'-bisphosphate nucleotidase 1(Bpnt1) |
| Q80V26 | 242291 | Mus musculus | 3'(2'), 5'-bisphosphate nucleotidase 2(Bpnt2) |
| P38060 | 15356 | Mus musculus | 3-hydroxy-3-methylglutaryl-Coenzyme A lyase(Hmgcl) |
| Q8JZK9 | 208715 | Mus musculus | 3-hydroxy-3-methylglutaryl-Coenzyme A synthase 1(Hmgcs1) |
| P54869 | 15360 | Mus musculus | 3-hydroxy-3-methylglutaryl-Coenzyme A synthase 2(Hmgcs2) |
| Q78JT3 | 107766 | Mus musculus | 3-hydroxyanthranilate 3,4-dioxygenase(Haao) |
| Q99L13 | 58875 | Mus musculus | 3-hydroxyisobutyrate dehydrogenase(Hibadh) |
| Q9D0K2 | 67041 | Mus musculus | 3-oxoacid CoA transferase 1(Oxct1) |
| Q61753 | 236539 | Mus musculus | 3-phosphoglycerate dehydrogenase(Phgdh) |
| P61922 | 268860 | Mus musculus | 4-aminobutyrate aminotransferase(Abat) |
| P49429 | 15445 | Mus musculus | 4-hydroxyphenylpyruvic acid dioxygenase(Hpd) |
| Q9CWJ9 | 108147 | Mus musculus | 5-aminoimidazole-4-carboxamide ribonucleotide formyltransferase/IMP cyclohydrolase(Atic) |
| Q9R1V6 | 11496 | Mus musculus | a disintegrin and metallopeptidase domain 22(Adam22) |
| Q9R1V7 | 23792 | Mus musculus | a disintegrin and metallopeptidase domain 23(Adam23) |
| Q9WTQ5 | 83397 | Mus musculus | A kinase anchor protein 12(Akap12) |
| D3YVF0 | 238276 | Mus musculus | A kinase anchor protein 5(Akap5) |
| Q8K4F5 | 68758 | Mus musculus | abhydrolase domain containing 11(Abhd11) |
| P62484 | 329165 | Mus musculus | abl interactor 2(Abi2) |
| Q8BWT1 | 52538 | Mus musculus | acetyl-CoA acyltransferase 2(Acaa2) |
| Q8QZT1 | 110446 | Mus musculus | acetyl-Coenzyme A acetyltransferase 1(Acat1) |
| Q8CAY6 | 110460 | Mus musculus | acetyl-Coenzyme A acetyltransferase 2(Acat2) |
| Q921H8 | 113868 | Mus musculus | acetyl-Coenzyme A acyltransferase 1A(Acaa1a) |
| Q921H8 | 235674 | Mus musculus | acetyl-Coenzyme A acyltransferase 1B(Acaa1b) |
| Q8VCH0 | 235674 | Mus musculus | acetyl-Coenzyme A acyltransferase 1B(Acaa1b) |
| O35381 | 11737 | Mus musculus | acidic nuclear phosphoprotein 32 family member A(Anp32a) |
| Q99KI0 | 11429 | Mus musculus | aconitase 2, mitochondrial(Aco2) |
| P68134 | 11459 | Mus musculus | actin alpha 1, skeletal muscle(Acta1) |
| P62737 | 11475 | Mus musculus | actin alpha 2, smooth muscle, aorta(Acta2) |
| P60710 | 11475 | Mus musculus | actin alpha 2, smooth muscle, aorta(Acta2) |
| Q9R0Q6 | 56443 | Mus musculus | actin related protein 2/3 complex, subunit 1A(Arpc1a) |
| Q9CVB6 | 76709 | Mus musculus | actin related protein 2/3 complex, subunit 2(Arpc2) |
| Q9JM76 | 56378 | Mus musculus | actin related protein 2/3 complex, subunit 3(Arpc3) |

| | | | |
|---|---|---|---|
| P59999 | 68089 | Mus musculus | actin related protein 2/3 complex, subunit 4(Arpc4) |
| Q9CPW4 | 67771 | Mus musculus | actin related protein 2/3 complex, subunit 5(Arpc5) |
| P60710 | 11461 | Mus musculus | actin, beta(Actb) |
| Q8BFZ3 | 238880 | Mus musculus | actin, beta-like 2(Actbl2) |
| P63260 | 11465 | Mus musculus | actin, gamma, cytoplasmic 1(Actg1) |
| P60710 | 11465 | Mus musculus | actin, gamma, cytoplasmic 1(Actg1) |
| P57780 | 60595 | Mus musculus | actinin alpha 4(Actn4) |
| Q61490 | 11658 | Mus musculus | activated leukocyte cell adhesion molecule(Alcam) |
| Q8VCW8 | 264895 | Mus musculus | acyl-CoA synthetase family member 2(Acsf2) |
| Q9CQR4 | 66834 | Mus musculus | acyl-CoA thioesterase 13(Acot13) |
| Q91V12 | 70025 | Mus musculus | acyl-CoA thioesterase 7(Acot7) |
| Q5XG73 | 74159 | Mus musculus | acyl-Coenzyme A binding domain containing 5(Acbd5) |
| P45952 | 11364 | Mus musculus | acyl-Coenzyme A dehydrogenase, medium chain(Acadm) |
| Q9R0H0 | 11430 | Mus musculus | acyl-Coenzyme A oxidase 1, palmitoyl(Acox1) |
| P56376 | 66204 | Mus musculus | acylphosphatase 1(Acyp1) |
| P56375 | 75572 | Mus musculus | acylphosphatase 2, muscle type(Acyp2) |
| P17426 | 11771 | Mus musculus | adaptor-related protein complex 2, alpha 1 subunit(Ap2a1) |
| P17427 | 11772 | Mus musculus | adaptor-related protein complex 2, alpha 2 subunit(Ap2a2) |
| Q9DBG3 | 71770 | Mus musculus | adaptor-related protein complex 2, beta 1 subunit(Ap2b1) |
| P84091 | 11773 | Mus musculus | adaptor-related protein complex 2, mu 1 subunit(Ap2m1) |
| Q9QYC0 | 11518 | Mus musculus | adducin 1(Add1) |
| Q9QYB8 | 11519 | Mus musculus | adducin 2(Add2) |
| P55264 | 11534 | Mus musculus | adenosine kinase(Adk) |
| Q9R0Y5 | 11636 | Mus musculus | adenylate kinase 1(Ak1) |
| Q9WTP6 | 11637 | Mus musculus | adenylate kinase 2(Ak2) |
| Q9WTP7 | 56248 | Mus musculus | adenylate kinase 3(Ak3) |
| Q9WUR9 | 11639 | Mus musculus | adenylate kinase 4(Ak4) |
| Q8VEH3 | 68724 | Mus musculus | ADP-ribosylation factor-like 8A(Arl8a) |
| Q9CQW2 | 67166 | Mus musculus | ADP-ribosylation factor-like 8B(Arl8b) |
| Q61282 | 11595 | Mus musculus | aggrecan(Acan) |
| A2AS89 | 75986 | Mus musculus | agmatinase(Agmat) |
| Q8BK64 | 217737 | Mus musculus | AHA1, activator of heat shock protein ATPase 1(Ahsa1) |
| P97449 | 16790 | Mus musculus | alanyl aminopeptidase, membrane(Anpep) |
| Q8BGQ7 | 234734 | Mus musculus | alanyl-tRNA synthetase 1(Aars1) |
| P07724 | 11657 | Mus musculus | albumin(Alb) |

| | | | |
|---|---|---|---|
| P00329 | 11522 | Mus musculus | alcohol dehydrogenase 1 (class I)(Adh1) |
| P28474 | 11532 | Mus musculus | alcohol dehydrogenase 5 (class III), chi polypeptide(Adh5) |
| P47738 | 11669 | Mus musculus | aldehyde dehydrogenase 2, mitochondrial(Aldh2) |
| Q8CHT0 | 212647 | Mus musculus | aldehyde dehydrogenase 4 family, member A1(Aldh4a1) |
| P24549 | 11668 | Mus musculus | aldehyde dehydrogenase family 1, subfamily A1(Aldh1a1) |
| Q9EQ20 | 104776 | Mus musculus | aldehyde dehydrogenase family 6, subfamily A1(Aldh6a1) |
| G3X982 | 71724 | Mus musculus | aldehyde oxidase 3(Aox3) |
| Q8BWF0 | 214579 | Mus musculus | aldhehyde dehydrogenase family 5, subfamily A1(Aldh5a1) |
| P45376 | 11677 | Mus musculus | aldo-keto reductase family 1 member B(Akr1b1) |
| Q9JII6 | 58810 | Mus musculus | aldo-keto reductase family 1, member A1(Akr1a1) |
| P21300 | 11997 | Mus musculus | aldo-keto reductase family 1, member B7(Akr1b7) |
| P70694 | 83702 | Mus musculus | aldo-keto reductase family 1, member C6(Akr1c6) |
| P05064 | 11674 | Mus musculus | aldolase A, fructose-bisphosphate(Aldoa) |
| Q91Y97 | 230163 | Mus musculus | aldolase B, fructose-bisphosphate(Aldob) |
| P05063 | 11676 | Mus musculus | aldolase C, fructose-bisphosphate(Aldoc) |
| Q8BHN3 | 14376 | Mus musculus | alpha glucosidase 2 alpha neutral subunit(Ganab) |
| Q99JW2 | 109652 | Mus musculus | aminoacylase 1(Acy1) |
| Q11011 | 19155 | Mus musculus | aminopeptidase puromycin sensitive(Npepps) |
| Q7TQF7 | 218038 | Mus musculus | amphiphysin(Amph) |
| P12023 | 11820 | Mus musculus | amyloid beta precursor protein(App) |
| Q8C8R3 | 109676 | Mus musculus | ankyrin 2, brain(Ank2) |
| G5E8K5 | 11735 | Mus musculus | ankyrin 3, epithelial(Ank3) |
| Q6GQX6 | 75691 | Mus musculus | ankyrin repeat and sterile alpha motif domain containing 6(Anks6) |
| P97384 | 11744 | Mus musculus | annexin A11(Anxa11) |
| O35639 | 11745 | Mus musculus | annexin A3(Anxa3) |
| P48036 | 11747 | Mus musculus | annexin A5(Anxa5) |
| P14824 | 11749 | Mus musculus | annexin A6(Anxa6) |
| Q07076 | 11750 | Mus musculus | annexin A7(Anxa7) |
| E9PVX6 | 17345 | Mus musculus | antigen identified by monoclonal antibody Ki 67(Mki67) |
| Q00623 | 11806 | Mus musculus | apolipoprotein A-I(Apoa1) |
| P09813 | 11807 | Mus musculus | apolipoprotein A-II(Apoa2) |
| P06728 | 11808 | Mus musculus | apolipoprotein A-IV(Apoa4) |
| P51910 | 11815 | Mus musculus | apolipoprotein D(Apod) |
| P08226 | 11816 | Mus musculus | apolipoprotein E(Apoe) |
| P28352 | 11792 | Mus musculus | apurinic/apyrimidinic endonuclease 1(Apex1) |

| | | | |
|---|---|---|---|
| P55088 | 11829 | Mus musculus | aquaporin 4(Aqp4) |
| P61205 | 11842 | Mus musculus | ARF GTPase 3(Arf3) |
| P61750 | 11843 | Mus musculus | ARF GTPase 4(Arf4) |
| P62331 | 11845 | Mus musculus | ARF GTPase 6(Arf6) |
| Q9EPJ9 | 228998 | Mus musculus | ARF GTPase activating protein 1(Arfgap1) |
| Q8K221 | 76932 | Mus musculus | ARF interacting protein 2(Arfip2) |
| Q61176 | 11846 | Mus musculus | arginase, liver(Arg1) |
| Q3UL36 | 234023 | Mus musculus | arginine and glutamate rich 1(Arglu1) |
| P16460 | 11898 | Mus musculus | argininosuccinate synthetase 1(Ass1) |
| P61164 | 54130 | Mus musculus | ARP1 actin-related protein 1A, centractin alpha(Actr1a) |
| Q99JY9 | 74117 | Mus musculus | ARP3 actin-related protein 3(Actr3) |
| P34927 | 11889 | Mus musculus | asialoglycoprotein receptor 1(Asgr1) |
| Q8C0M9 | 66514 | Mus musculus | asparaginase like 1(Asrgl1) |
| Q8BP47 | 70223 | Mus musculus | asparaginyl-tRNA synthetase 1(Nars1) |
| Q8BSY0 | 65973 | Mus musculus | aspartate-beta-hydroxylase(Asph) |
| Q91V92 | 104112 | Mus musculus | ATP citrate lyase(Acly) |
| P03930 | 17706 | Mus musculus | ATP synthase 8, mitochondrial(mt-Atp8) |
| Q03265 | 11946 | Mus musculus | ATP synthase F1 subunit alpha(Atp5f1a) |
| P56480 | 11947 | Mus musculus | ATP synthase F1 subunit beta(Atp5f1b) |
| Q9D3D9 | 66043 | Mus musculus | ATP synthase F1 subunit delta(Atp5f1d) |
| P56382 | 67126 | Mus musculus | ATP synthase F1 subunit epsilon(Atp5f1e) |
| Q91VR2 | 11949 | Mus musculus | ATP synthase F1 subunit gamma(Atp5f1c) |
| O35143 | 11983 | Mus musculus | ATP synthase inhibitory factor subunit 1(Atp5if1) |
| Q06185 | 11958 | Mus musculus | ATP synthase membrane subunit e(Atp5me) |
| P56135 | 57423 | Mus musculus | ATP synthase membrane subunit f(Atp5mf) |
| Q9CPQ8 | 27425 | Mus musculus | ATP synthase membrane subunit g(Atp5mg) |
| Q78IK2 | 66477 | Mus musculus | ATP synthase membrane subunit k(Atp5mk) |
| Q9DCX2 | 71679 | Mus musculus | ATP synthase peripheral stalk subunit d(Atp5pd) |
| P97450 | 11957 | Mus musculus | ATP synthase peripheral stalk subunit F6(Atp5pf) |
| Q9DB20 | 28080 | Mus musculus | ATP synthase peripheral stalk subunit OSCP(Atp5po) |
| Q9CQQ7 | 11950 | Mus musculus | ATP synthase peripheral stalk-membrane subunit b(Atp5pb) |
| G5E829 | 67972 | Mus musculus | ATPase, Ca++ transporting, plasma membrane 1(Atp2b1) |
| Q9R0K7 | 11941 | Mus musculus | ATPase, Ca++ transporting, plasma membrane 2(Atp2b2) |
| Q6Q477 | 381290 | Mus musculus | ATPase, Ca++ transporting, plasma membrane 4(Atp2b4) |
| Q9R1Q9 | 54411 | Mus musculus | ATPase, H+ transporting, lysosomal accessory protein 1(Atp6ap1) |

| | | | |
|---|---|---|---|
| Q9Z1G4 | 11975 | Mus musculus | ATPase, H+ transporting, lysosomal V0 subunit A1(Atp6v0a1) |
| P51863 | 11972 | Mus musculus | ATPase, H+ transporting, lysosomal V0 subunit D1(Atp6v0d1) |
| P50516 | 11964 | Mus musculus | ATPase, H+ transporting, lysosomal V1 subunit A(Atp6v1a) |
| P62814 | 11966 | Mus musculus | ATPase, H+ transporting, lysosomal V1 subunit B2(Atp6v1b2) |
| P57746 | 73834 | Mus musculus | ATPase, H+ transporting, lysosomal V1 subunit D(Atp6v1d) |
| P50518 | 11973 | Mus musculus | ATPase, H+ transporting, lysosomal V1 subunit E1(Atp6v1e1) |
| Q9D1K2 | 66144 | Mus musculus | ATPase, H+ transporting, lysosomal V1 subunit F(Atp6v1f) |
| Q9CR51 | 66290 | Mus musculus | ATPase, H+ transporting, lysosomal V1 subunit G1(Atp6v1g1) |
| Q9WTT4 | 66237 | Mus musculus | ATPase, H+ transporting, lysosomal V1 subunit G2(Atp6v1g2) |
| Q8BVE3 | 108664 | Mus musculus | ATPase, H+ transporting, lysosomal V1 subunit H(Atp6v1h) |
| Q8VDN2 | 11928 | Mus musculus | ATPase, Na+/K+ transporting, alpha 1 polypeptide(Atp1a1) |
| Q6PIE5 | 98660 | Mus musculus | ATPase, Na+/K+ transporting, alpha 2 polypeptide(Atp1a2) |
| Q6PIC6 | 232975 | Mus musculus | ATPase, Na+/K+ transporting, alpha 3 polypeptide(Atp1a3) |
| P14094 | 11931 | Mus musculus | ATPase, Na+/K+ transporting, beta 1 polypeptide(Atp1b1) |
| P14231 | 11932 | Mus musculus | ATPase, Na+/K+ transporting, beta 2 polypeptide(Atp1b2) |
| P97370 | 11933 | Mus musculus | ATPase, Na+/K+ transporting, beta 3 polypeptide(Atp1b3) |
| Q9JLZ3 | 11992 | Mus musculus | AU RNA binding protein/enoyl-coenzyme A hydratase(Auh) |
| Q61335 | 27061 | Mus musculus | B cell receptor associated protein 31(Bcap31) |
| Q9R069 | 57278 | Mus musculus | basal cell adhesion molecule(Bcam) |
| Q9CQH7 | 70533 | Mus musculus | basic transcription factor 3-like 4(Btf3l4) |
| P18572 | 12215 | Mus musculus | basigin(Bsg) |
| O88737 | 12217 | Mus musculus | bassoon(Bsn) |
| P01887 | 12010 | Mus musculus | beta-2 microglobulin(B2m) |
| O35490 | 12116 | Mus musculus | betaine-homocysteine methyltransferase(Bhmt) |
| Q923D2 | 233016 | Mus musculus | biliverdin reductase B(Blvrb) |
| Q8R164 | 68021 | Mus musculus | biphenyl hydrolase like(Bphl) |
| Q8VBW5 | 70508 | Mus musculus | bobby sox HMG box containing(Bbx) |
| Q91XV3 | 70350 | Mus musculus | brain abundant, membrane attached signal protein 1(Basp1) |
| Q80YN3 | 76960 | Mus musculus | brain enriched myelin associated protein 1(Bcas1) |
| Q8CI94 | 110078 | Mus musculus | brain glycogen phosphorylase(Pygb) |
| Q8BKX1 | 108100 | Mus musculus | brain-specific angiogenesis inhibitor 1-associated protein 2(Baiap2) |
| Q99MP8 | 72399 | Mus musculus | BRCA1 associated protein(Brap) |
| Q8CHH5 | 210982 | Mus musculus | BRD4 interacting chromatin remodeling complex associated protein like(Bicral) |
| Q61361 | 12032 | Mus musculus | brevican(Bcan) |
| Q91VR8 | 101314 | Mus musculus | BRICK1, SCAR/WAVE actin-nucleating complex subunit(Brk1) |

| | | | |
|---|---|---|---|
| O08539 | 30948 | Mus musculus | bridging integrator 1(Bin1) |
| D3Z6Q9 | 668218 | Mus musculus | bridging integrator 2(Bin2) |
| Q3UI43 | 68251 | Mus musculus | BRISC and BRCA1 A complex member 1(Babam1) |
| Q80TJ1 | 27062 | Mus musculus | Ca2+-dependent secretion activator(Cadps) |
| P15116 | 12558 | Mus musculus | cadherin 2(Cdh2) |
| P12658 | 12307 | Mus musculus | calbindin 1(Calb1) |
| Q08331 | 12308 | Mus musculus | calbindin 2(Calb2) |
| Q6PHS9 | 56808 | Mus musculus | calcium channel, voltage-dependent, alpha 2/delta subunit 2(Cacna2d2) |
| O08532 | 12293 | Mus musculus | calcium channel, voltage-dependent, alpha2/delta subunit 1(Cacna2d1) |
| Q8VHW2 | 81905 | Mus musculus | calcium channel, voltage-dependent, gamma subunit 8(Cacng8) |
| P11798 | 12322 | Mus musculus | calcium/calmodulin-dependent protein kinase II alpha(Camk2a) |
| P28652 | 12323 | Mus musculus | calcium/calmodulin-dependent protein kinase II, beta(Camk2b) |
| Q8VBY2 | 55984 | Mus musculus | calcium/calmodulin-dependent protein kinase kinase 1, alpha(Camkk1) |
| Q9CXW3 | 12301 | Mus musculus | calcyclin binding protein(Cacybp) |
| P0DP27 | 12314 | Mus musculus | calmodulin 2(Calm2) |
| P35564 | 12330 | Mus musculus | calnexin(Canx) |
| Q9DAW9 | 71994 | Mus musculus | calponin 3, acidic(Cnn3) |
| P14211 | 12317 | Mus musculus | calreticulin(Calr) |
| Q3UHL1 | 235604 | Mus musculus | CaM kinase-like vesicle-associated(Camkv) |
| Q5M8N0 | 380686 | Mus musculus | cannabinoid receptor interacting protein 1(Cnrip1) |
| P47753 | 12340 | Mus musculus | capping actin protein of muscle Z-line subunit alpha 1(Capza1) |
| P47757 | 12345 | Mus musculus | capping actin protein of muscle Z-line subunit beta(Capzb) |
| Q8C196 | 227231 | Mus musculus | carbamoyl-phosphate synthetase 1(Cps1) |
| P00920 | 12349 | Mus musculus | carbonic anhydrase 2(Car2) |
| P16015 | 12350 | Mus musculus | carbonic anhydrase 3(Car3) |
| P23953 | 13884 | Mus musculus | carboxylesterase 1C(Ces1c) |
| Q8VCT4 | 104158 | Mus musculus | carboxylesterase 1D(Ces1d) |
| Q91WU0 | 234564 | Mus musculus | carboxylesterase 1F(Ces1f) |
| Q8VCC2 | 12623 | Mus musculus | carboxylesterase 1G(Ces1g) |
| Q8QZR3 | 102022 | Mus musculus | carboxylesterase 2A(Ces2a) |
| Q91WG0 | 234671 | Mus musculus | carboxylesterase 2C(Ces2c) |
| Q8BK48 | 234673 | Mus musculus | carboxylesterase 2E(Ces2e) |
| Q63880 | 382053 | Mus musculus | carboxylesterase 3A(Ces3a) |
| Q8VCU1 | 13909 | Mus musculus | carboxylesterase 3B(Ces3b) |
| Q00493 | 12876 | Mus musculus | carboxypeptidase E(Cpe) |

| | | | |
|---|---|---|---|
| O88696 | 53895 | Mus musculus | caseinolytic mitochondrial matrix peptidase proteolytic subunit(Clpp) |
| P24270 | 12359 | Mus musculus | catalase(Cat) |
| O88587 | 12846 | Mus musculus | catechol-O-methyltransferase(Comt) |
| Q02248 | 12387 | Mus musculus | catenin beta 1(Ctnnb1) |
| P10605 | 13030 | Mus musculus | cathepsin B(Ctsb) |
| P18242 | 13033 | Mus musculus | cathepsin D(Ctsd) |
| O54901 | 17470 | Mus musculus | CD200 molecule(Cd200) |
| Q61735 | 16423 | Mus musculus | CD47 antigen (Rh-related antigen, integrin-associated signal transducer)(Cd47) |
| P35762 | 12520 | Mus musculus | CD81 antigen(Cd81) |
| P40237 | 12521 | Mus musculus | CD82 antigen(Cd82) |
| P40240 | 12527 | Mus musculus | CD9 antigen(Cd9) |
| Q91WS0 | 52637 | Mus musculus | CDGSH iron sulfur domain 1(Cisd1) |
| Q8R5M8 | 54725 | Mus musculus | cell adhesion molecule 1(Cadm1) |
| Q8BLQ9 | 239857 | Mus musculus | cell adhesion molecule 2(Cadm2) |
| Q99N28 | 94332 | Mus musculus | cell adhesion molecule 3(Cadm3) |
| Q9JKC6 | 57754 | Mus musculus | cell cycle exit and neuronal differentiation 1(Cend1) |
| Q61081 | 12539 | Mus musculus | cell division cycle 37(Cdc37) |
| P60766 | 12540 | Mus musculus | cell division cycle 42(Cdc42) |
| Q8BI22 | 75216 | Mus musculus | centrosomal protein 128(Cep128) |
| P80314 | 12461 | Mus musculus | chaperonin containing TCP1 subunit 2(Cct2) |
| P80315 | 12464 | Mus musculus | chaperonin containing TCP1 subunit 4(Cct4) |
| P80313 | 12468 | Mus musculus | chaperonin containing TCP1 subunit 7(Cct7) |
| P42932 | 12469 | Mus musculus | chaperonin containing TCP1 subunit 8(Cct8) |
| Q9D8B3 | 75608 | Mus musculus | charged multivesicular body protein 4B(Chmp4b) |
| Q9QYB1 | 29876 | Mus musculus | chloride intracellular channel 4(Clic4) |
| Q71M36 | 29873 | Mus musculus | chondroitin sulfate proteoglycan 5(Cspg5) |
| P83917 | 12412 | Mus musculus | chromobox 1(Cbx1) |
| Q9CZU6 | 12974 | Mus musculus | citrate synthase(Cs) |
| Q68FD5 | 67300 | Mus musculus | clathrin heavy chain(Cltc) |
| O08585 | 12757 | Mus musculus | clathrin light chain A(Clta) |
| Q6IRU5 | 74325 | Mus musculus | clathrin light chain B(Cltb) |
| Q60771 | 18417 | Mus musculus | claudin 11(Cldn11) |
| Q06890 | 12759 | Mus musculus | clusterin(Clu) |
| Q9D1A2 | 66054 | Mus musculus | CNDP dipeptidase 2(Cndp2) |
| Q9CQI6 | 72042 | Mus musculus | coactosin like F-actin binding protein 1(Cotl1) |

| | | | |
|---|---|---|---|
| P61924 | 56447 | Mus musculus | coatomer protein complex, subunit zeta 1(Copz1) |
| Q8K1Z0 | 67914 | Mus musculus | coenzyme Q9(Coq9) |
| P18760 | 12631 | Mus musculus | cofilin 1, non-muscle(Cfl1) |
| P45591 | 12632 | Mus musculus | cofilin 2, muscle(Cfl2) |
| Q9CRB9 | 66075 | Mus musculus | coiled-coil-helix-coiled-coil-helix domain containing 3(Chchd3) |
| Q91VN4 | 66098 | Mus musculus | coiled-coil-helix-coiled-coil-helix domain containing 6(Chchd6) |
| Q5QNQ9 | 373864 | Mus musculus | collagen, type XXVII, alpha 1(Col27a1) |
| P97427 | 12933 | Mus musculus | collapsin response mediator protein 1(Crmp1) |
| O35658 | 12261 | Mus musculus | complement component 1, q subcomponent binding protein(C1qbp) |
| P63040 | 12889 | Mus musculus | complexin 1(Cplx1) |
| P84086 | 12890 | Mus musculus | complexin 2(Cplx2) |
| P12960 | 12805 | Mus musculus | contactin 1(Cntn1) |
| Q61330 | 21367 | Mus musculus | contactin 2(Cntn2) |
| O54991 | 53321 | Mus musculus | contactin associated protein-like 1(Cntnap1) |
| Q9CPW0 | 66797 | Mus musculus | contactin associated protein-like 2(Cntnap2) |
| Q8JZW4 | 240058 | Mus musculus | copine V(Cpne5) |
| Q9Z140 | 12891 | Mus musculus | copine VI(Cpne6) |
| O89053 | 12721 | Mus musculus | coronin, actin binding protein 1A(Coro1a) |
| Q60598 | 13043 | Mus musculus | cortactin(Cttn) |
| Q04447 | 12709 | Mus musculus | creatine kinase, brain(Ckb) |
| P30275 | 12716 | Mus musculus | creatine kinase, mitochondrial 1, ubiquitous(Ckmt1) |
| P07310 | 12715 | Mus musculus | creatine kinase, muscle(Ckm) |
| Q68ED7 | 382056 | Mus musculus | CREB regulated transcription coactivator 1(Crtc1) |
| P23927 | 12955 | Mus musculus | crystallin, alpha B(Cryab) |
| Q99KP3 | 68631 | Mus musculus | crystallin, lambda 1(Cryl1) |
| O54983 | 12971 | Mus musculus | crystallin, mu(Crym) |
| P56546 | 13017 | Mus musculus | C-terminal binding protein 2(Ctbp2) |
| A2A432 | 72584 | Mus musculus | cullin 4B(Cul4b) |
| Q6ZQ38 | 71902 | Mus musculus | cullin associated and neddylation disassociated 1(Cand1) |
| P53564 | 13047 | Mus musculus | cut-like homeobox 1(Cux1) |
| P40124 | 12331 | Mus musculus | cyclase associated actin cytoskeleton regulatory protein 1(Cap1) |
| Q8BGU5 | 67974 | Mus musculus | cyclin Y(Ccny) |
| Q921M7 | 223601 | Mus musculus | CYFIP related Rac1 interactor B(Cyrib) |
| Q62426 | 13014 | Mus musculus | cystatin B(Cstb) |
| P97315 | 13007 | Mus musculus | cysteine and glycine-rich protein 1(Csrp1) |

| | | | |
|---|---|---|---|
| Q9D1P4 | 66917 | Mus musculus | cysteine and histidine rich domain containing 1(Chordc1) |
| Q9DCT8 | 68337 | Mus musculus | cysteine rich protein 2(Crip2) |
| Q9DBP5 | 66588 | Mus musculus | cytidine/uridine monophosphate kinase 1(Cmpk1) |
| P56395 | 109672 | Mus musculus | cytochrome b5 type A (microsomal)(Cyb5a) |
| Q9CQX2 | 66427 | Mus musculus | cytochrome b5 type B(Cyb5b) |
| P00405 | 17709 | Mus musculus | cytochrome c oxidase II, mitochondrial(mt-Co2) |
| P19783 | 12857 | Mus musculus | cytochrome c oxidase subunit 4I1(Cox4i1) |
| P12787 | 12858 | Mus musculus | cytochrome c oxidase subunit 5A(Cox5a) |
| P19536 | 12859 | Mus musculus | cytochrome c oxidase subunit 5B(Cox5b) |
| P43024 | 12861 | Mus musculus | cytochrome c oxidase subunit 6A1(Cox6a1) |
| Q9CPQ1 | 12864 | Mus musculus | cytochrome c oxidase subunit 6C(Cox6c) |
| P48771 | 12866 | Mus musculus | cytochrome c oxidase subunit 7A2(Cox7a2) |
| P56393 | 66142 | Mus musculus | cytochrome c oxidase subunit 7B(Cox7b) |
| P17665 | 12867 | Mus musculus | cytochrome c oxidase subunit 7C(Cox7c) |
| P56391 | 110323 | Mus musculus | cytochrome c oxidase, subunit 6B1(Cox6b1) |
| P62897 | 13063 | Mus musculus | cytochrome c, somatic(Cycs) |
| Q9D0M3 | 66445 | Mus musculus | cytochrome c-1(Cyc1) |
| P24457 | 545123 | Mus musculus | cytochrome P450, family 2, subfamily d, polypeptide 11(Cyp2d11) |
| Q5SQX6 | 76884 | Mus musculus | cytoplasmic FMR1 interacting protein 2(Cyfip2) |
| Q8BMK4 | 216197 | Mus musculus | cytoskeleton-associated protein 4(Ckap4) |
| O35215 | 13202 | Mus musculus | D-dopachrome tautomerase(Ddt) |
| Q91VR5 | 104721 | Mus musculus | DEAD box helicase 1(Ddx1) |
| Q30KP6 | 77673 | Mus musculus | defensin beta 41(Defb41) |
| Q9WV69 | 13829 | Mus musculus | dematin actin binding protein(Dmtn) |
| Q9CQJ6 | 68184 | Mus musculus | density-regulated protein(Denr) |
| P31001 | 13346 | Mus musculus | desmin(Des) |
| Q9R0P5 | 56431 | Mus musculus | destrin(Dstn) |
| P31786 | 13167 | Mus musculus | diazepam binding inhibitor(Dbi) |
| Q9DBB8 | 71755 | Mus musculus | dihydrodiol dehydrogenase(Dhdh) |
| P53395 | 13171 | Mus musculus | dihydrolipoamide branched chain transacylase E2(Dbt) |
| O08749 | 13382 | Mus musculus | dihydrolipoamide dehydrogenase(Dld) |
| Q8BMF4 | 235339 | Mus musculus | dihydrolipoamide S-acetyltransferase(Dlat) |
| Q9D2G2 | 78920 | Mus musculus | dihydrolipoamide S-succinyltransferase(Dlst) |
| O08553 | 12934 | Mus musculus | dihydropyrimidinase-like 2(Dpysl2) |
| Q62188 | 22240 | Mus musculus | dihydropyrimidinase-like 3(Dpysl3) |

| | | | |
|---|---|---|---|
| Q9CWS0 | 69219 | Mus musculus | dimethylarginine dimethylaminohydrolase 1(Ddah1) |
| P28843 | 13482 | Mus musculus | dipeptidylpeptidase 4(Dpp4) |
| Q9Z218 | 13483 | Mus musculus | dipeptidylpeptidase 6(Dpp6) |
| Q91XM9 | 23859 | Mus musculus | discs large MAGUK scaffold protein 2(Dlg2) |
| Q9QYJ0 | 56445 | Mus musculus | DnaJ heat shock protein family (Hsp40) member A2(Dnaja2) |
| P60904 | 13002 | Mus musculus | DnaJ heat shock protein family (Hsp40) member C5(Dnajc5) |
| Q9JLM8 | 13175 | Mus musculus | doublecortin-like kinase 1(Dclk1) |
| Q9QXS6 | 56320 | Mus musculus | drebrin 1(Dbn1) |
| Q62418 | 13169 | Mus musculus | drebrin-like(Dbnl) |
| Q99KJ8 | 69654 | Mus musculus | dynactin 2(Dctn2) |
| Q9Z0Y1 | 53598 | Mus musculus | dynactin 3(Dctn3) |
| P39053 | 13429 | Mus musculus | dynamin 1(Dnm1) |
| Q8K1M6 | 74006 | Mus musculus | dynamin 1-like(Dnm1l) |
| P39054 | 13430 | Mus musculus | dynamin 2(Dnm2) |
| Q9JHU4 | 13424 | Mus musculus | dynein cytoplasmic 1 heavy chain 1(Dync1h1) |
| O88485 | 13426 | Mus musculus | dynein cytoplasmic 1 intermediate chain 1(Dync1i1) |
| Q8R1Q8 | 235661 | Mus musculus | dynein cytoplasmic 1 light intermediate chain 1(Dync1li1) |
| P63168 | 56455 | Mus musculus | dynein light chain LC8-type 1(Dynll1) |
| Q9D0M5 | 68097 | Mus musculus | dynein light chain LC8-type 2(Dynll2) |
| P62627 | 67068 | Mus musculus | dynein light chain roadblock-type 1(Dynlrb1) |
| Q6PDL0 | 234663 | Mus musculus | dynein, cytoplasmic 1 light intermediate chain 2(Dync1li2) |
| Q8BL66 | 216238 | Mus musculus | early endosome antigen 1(Eea1) |
| Q9D4J1 | 98363 | Mus musculus | EF hand domain containing 1(Efhd1) |
| Q9D8Y0 | 27984 | Mus musculus | EF hand domain containing 2(Efhd2) |
| Q99LC5 | 110842 | Mus musculus | electron transferring flavoprotein, alpha polypeptide(Etfa) |
| Q9DCW4 | 110826 | Mus musculus | electron transferring flavoprotein, beta polypeptide(Etfb) |
| Q921G7 | 66841 | Mus musculus | electron transferring flavoprotein, dehydrogenase(Etfdh) |
| P83940 | 67923 | Mus musculus | elongin C(Eloc) |
| P21995 | 13723 | Mus musculus | embigin(Emb) |
| P57759 | 67397 | Mus musculus | endoplasmic reticulum protein 29(Erp29) |
| P17182 | 13806 | Mus musculus | enolase 1, alpha non-neuron(Eno1) |
| P17182 | 433182 | Mus musculus | enolase 1B, retrotransposed(Eno1b) |
| P17183 | 13807 | Mus musculus | enolase 2, gamma neuronal(Eno2) |
| P21550 | 13808 | Mus musculus | enolase 3, beta muscle(Eno3) |
| O35459 | 51798 | Mus musculus | enoyl coenzyme A hydratase 1, peroxisomal(Ech1) |

| | | | |
|---|---|---|---|
| Q8BH95 | 93747 | Mus musculus | enoyl Coenzyme A hydratase, short chain, 1, mitochondrial(Echs1) |
| P42125 | 13177 | Mus musculus | enoyl-Coenzyme A delta isomerase 1(Eci1) |
| Q60902 | 13859 | Mus musculus | epidermal growth factor receptor pathway substrate 15-like 1(Eps15l1) |
| Q01279 | 13649 | Mus musculus | epidermal growth factor receptor(Egfr) |
| Q80VP1 | 13854 | Mus musculus | epsin 1(Epn1) |
| Q8BFZ9 | 244373 | Mus musculus | ER lipid raft associated 2(Erlin2) |
| P84089 | 13877 | Mus musculus | ERH mRNA splicing and mitosis factor(Erh) |
| Q9Z2H5 | 13821 | Mus musculus | erythrocyte membrane protein band 4.1 like 1(Epb41l1) |
| O70318 | 13822 | Mus musculus | erythrocyte membrane protein band 4.1 like 2(Epb41l2) |
| Q9WV92 | 13823 | Mus musculus | erythrocyte membrane protein band 4.1 like 3(Epb41l3) |
| Q9DCM0 | 66071 | Mus musculus | ethylmalonic encephalopathy 1(Ethe1) |
| P10126 | 13627 | Mus musculus | eukaryotic translation elongation factor 1 alpha 1(Eef1a1) |
| P62631 | 13628 | Mus musculus | eukaryotic translation elongation factor 1 alpha 2(Eef1a2) |
| O70251 | 55949 | Mus musculus | eukaryotic translation elongation factor 1 beta 2(Eef1b2) |
| P57776 | 66656 | Mus musculus | eukaryotic translation elongation factor 1 delta(Eef1d) |
| Q9D8N0 | 67160 | Mus musculus | eukaryotic translation elongation factor 1 gamma(Eef1g) |
| P58252 | 13629 | Mus musculus | eukaryotic translation elongation factor 2(Eef2) |
| P48024 | 20918 | Mus musculus | eukaryotic translation initiation factor 1(Eif1) |
| Q9QZD9 | 54709 | Mus musculus | eukaryotic translation initiation factor 3, subunit I(Eif3i) |
| Q3UGC7 | 78655 | Mus musculus | eukaryotic translation initiation factor 3, subunit J1(Eif3j1) |
| P63242 | 276770 | Mus musculus | eukaryotic translation initiation factor 5A(Eif5a) |
| Q8BGY2 | 208691 | Mus musculus | eukaryotic translation initiation factor 5A2(Eif5a2) |
| Q91WJ8 | 51886 | Mus musculus | far upstream element (FUSE) binding protein 1(Fubp1) |
| Q61553 | 14086 | Mus musculus | fascin actin-bundling protein 1(Fscn1) |
| P12710 | 14080 | Mus musculus | fatty acid binding protein 1, liver(Fabp1) |
| P55050 | 14079 | Mus musculus | fatty acid binding protein 2, intestinal(Fabp2) |
| P11404 | 14077 | Mus musculus | fatty acid binding protein 3, muscle and heart(Fabp3) |
| Q05816 | 16592 | Mus musculus | fatty acid binding protein 5, epidermal(Fabp5) |
| P19096 | 14104 | Mus musculus | fatty acid synthase(Fasn) |
| P62862 | 14109 | Mus musculus | FAU ubiquitin like and ribosomal protein S30 fusion(Fau) |
| Q80UW2 | 230904 | Mus musculus | F-box protein 2(Fbxo2) |
| F8VPU2 | 223254 | Mus musculus | FERM, ARH/RhoGEF and pleckstrin domain protein 1(Farp1) |
| Q9CPW2 | 68165 | Mus musculus | ferredoxin 2(Fdx2) |
| P09528 | 14319 | Mus musculus | ferritin heavy polypeptide 1(Fth1) |
| P29391 | 14325 | Mus musculus | ferritin light polypeptide 1(Ftl1) |

| | | | |
|---|---|---|---|
| E9PV24 | 14161 | Mus musculus | fibrinogen alpha chain(Fga) |
| Q8K0E8 | 110135 | Mus musculus | fibrinogen beta chain(Fgb) |
| P61329 | 14167 | Mus musculus | fibroblast growth factor 12(Fgf12) |
| Q80X90 | 286940 | Mus musculus | filamin, beta(Flnb) |
| Q9CQ92 | 66437 | Mus musculus | fission, mitochondrial 1(Fis1) |
| P26883 | 14225 | Mus musculus | FK506 binding protein 1a(Fkbp1a) |
| P45878 | 14227 | Mus musculus | FK506 binding protein 2(Fkbp2) |
| P30416 | 14228 | Mus musculus | FK506 binding protein 4(Fkbp4) |
| O08917 | 14251 | Mus musculus | flotillin 1(Flot1) |
| Q60634 | 14252 | Mus musculus | flotillin 2(Flot2) |
| Q91XD4 | 14317 | Mus musculus | formiminotransferase cyclodeaminase(Ftcd) |
| O35943 | 14297 | Mus musculus | frataxin(Fxn) |
| Q9QXD6 | 14121 | Mus musculus | fructose bisphosphatase 1(Fbp1) |
| P97807 | 14194 | Mus musculus | fumarate hydratase 1(Fh1) |
| Q8R0F8 | 68636 | Mus musculus | fumarylacetoacetate hydrolase domain containing 1(Fahd1) |
| P35505 | 14085 | Mus musculus | fumarylacetoacetate hydrolase(Fah) |
| P56959 | 233908 | Mus musculus | fused in sarcoma(Fus) |
| Q9D164 | 59095 | Mus musculus | FXYD domain-containing ion transport regulator 6(Fxyd6) |
| P59648 | 57780 | Mus musculus | FXYD domain-containing ion transport regulator 7(Fxyd7) |
| Q99LD4 | 209318 | Mus musculus | G protein pathway suppressor 1(Gps1) |
| B2RSH2 | 14677 | Mus musculus | G protein subunit alpha i1(Gnai1) |
| P08752 | 14678 | Mus musculus | G protein subunit alpha i2(Gnai2) |
| Q9DC51 | 14679 | Mus musculus | G protein subunit alpha i3(Gnai3) |
| Q80SZ7 | 14707 | Mus musculus | G protein subunit gamma 5(Gng5) |
| Q8C419 | 241263 | Mus musculus | G protein-coupled receptor 158(Gpr158) |
| Q3UNH4 | 26913 | Mus musculus | G protein-regulated inducer of neurite outgrowth 1(Gprin1) |
| P60521 | 93739 | Mus musculus | GABA type A receptor associated protein like 2(Gabarapl2) |
| P50571 | 14400 | Mus musculus | gamma-aminobutyric acid type A receptor subunit beta 1(Gabrb1) |
| Q99JP7 | 207182 | Mus musculus | gamma-glutamyltransferase 7(Ggt7) |
| Q8VE33 | 228858 | Mus musculus | ganglioside-induced differentiation-associated protein 1-like 1(Gdap1l1) |
| P23242 | 14609 | Mus musculus | gap junction protein, alpha 1(Gja1) |
| P50396 | 14567 | Mus musculus | GDP dissociation inhibitor 1(Gdi1) |
| Q61598 | 14569 | Mus musculus | GDP dissociation inhibitor 2(Gdi2) |
| P13020 | 227753 | Mus musculus | gelsolin(Gsn) |
| Q9CQI3 | 63985 | Mus musculus | glia maturation factor, beta(Gmfb) |

| | | | |
|---|---|---|---|
| Q9ERL7 | 63986 | Mus musculus | glia maturation factor, gamma(Gmfg) |
| P03995 | 14580 | Mus musculus | glial fibrillary acidic protein(Gfap) |
| P06745 | 14751 | Mus musculus | glucose-6-phosphate isomerase 1(Gpi1) |
| P26443 | 14661 | Mus musculus | glutamate dehydrogenase 1(Glud1) |
| P23819 | 14800 | Mus musculus | glutamate receptor, ionotropic, AMPA2 (alpha 2)(Gria2) |
| Q68EF4 | 268934 | Mus musculus | glutamate receptor, metabotropic 4(Grm4) |
| P15105 | 14645 | Mus musculus | glutamate-ammonia ligase(Glul) |
| P05202 | 14719 | Mus musculus | glutamatic-oxaloacetic transaminase 2, mitochondrial(Got2) |
| P48320 | 14417 | Mus musculus | glutamic acid decarboxylase 2(Gad2) |
| P05201 | 14718 | Mus musculus | glutamic-oxaloacetic transaminase 1, soluble(Got1) |
| D3Z7P3 | 14660 | Mus musculus | glutaminase(Gls) |
| Q9D172 | 28295 | Mus musculus | glutamine amidotransferase like class 1 domain containing 3A(Gatd3a) |
| P11352 | 14775 | Mus musculus | glutathione peroxidase 1(Gpx1) |
| P10648 | 14858 | Mus musculus | glutathione S-transferase, alpha 2 (Yc2)(Gsta2) |
| P30115 | 14859 | Mus musculus | glutathione S-transferase, alpha 3(Gsta3) |
| P24472 | 14860 | Mus musculus | glutathione S-transferase, alpha 4(Gsta4) |
| P10649 | 14862 | Mus musculus | glutathione S-transferase, mu 1(Gstm1) |
| Q8R5I6 | 14865 | Mus musculus | glutathione S-transferase, mu 4(Gstm4) |
| P48774 | 14866 | Mus musculus | glutathione S-transferase, mu 5(Gstm5) |
| P19157 | 14870 | Mus musculus | glutathione S-transferase, pi 1(Gstp1) |
| P19157 | 14869 | Mus musculus | glutathione S-transferase, pi 2(Gstp2) |
| P46425 | 14869 | Mus musculus | glutathione S-transferase, pi 2(Gstp2) |
| P19157 | 1E+08 | Mus musculus | glutathione S-transferase, pi, pseudogene(Gstp-ps) |
| Q64471 | 14871 | Mus musculus | glutathione S-transferase, theta 1(Gstt1) |
| Q9WVL0 | 14874 | Mus musculus | glutathione transferase zeta 1 (maleylacetoacetate isomerase)(Gstz1) |
| P16858 | 14433 | Mus musculus | glyceraldehyde-3-phosphate dehydrogenase(Gapdh) |
| Q64521 | 14571 | Mus musculus | glycerol phosphate dehydrogenase 2, mitochondrial(Gpd2) |
| P13707 | 14555 | Mus musculus | glycerol-3-phosphate dehydrogenase 1 (soluble)(Gpd1) |
| Q3ULJ0 | 333433 | Mus musculus | glycerol-3-phosphate dehydrogenase 1-like(Gpd1l) |
| Q9QXF8 | 14711 | Mus musculus | glycine N-methyltransferase(Gnmt) |
| Q91XE0 | 107146 | Mus musculus | glycine-N-acyltransferase(Glyat) |
| Q9JL62 | 56356 | Mus musculus | glycolipid transfer protein(Gltp) |
| P35802 | 234267 | Mus musculus | glycoprotein m6a(Gpm6a) |
| P35803 | 14758 | Mus musculus | glycoprotein m6b(Gpm6b) |
| Q9CPU0 | 109801 | Mus musculus | glyoxalase 1(Glo1) |

| | | | |
|---|---|---|---|
| Q91Z53 | 76238 | Mus musculus | glyoxylate reductase/hydroxypyruvate reductase(Grhpr) |
| P63094 | 14683 | Mus musculus | GNAS complex locus(Gnas) |
| Q3UPI1 | 68659 | Mus musculus | golgi associated kinase 1B(Gask1b) |
| Q99JX3 | 70231 | Mus musculus | golgi reassembly stacking protein 2(Gorasp2) |
| Q8VD04 | 54645 | Mus musculus | GRIP1 associated protein 1(Gripap1) |
| P06837 | 14432 | Mus musculus | growth associated protein 43(Gap43) |
| Q60631 | 14784 | Mus musculus | growth factor receptor bound protein 2(Grb2) |
| P62874 | 14688 | Mus musculus | guanine nucleotide binding protein (G protein), beta 1(Gnb1) |
| P62880 | 14693 | Mus musculus | guanine nucleotide binding protein (G protein), beta 2(Gnb2) |
| P62880 | 14696 | Mus musculus | guanine nucleotide binding protein (G protein), beta 4(Gnb4) |
| Q9DAS9 | 14701 | Mus musculus | guanine nucleotide binding protein (G protein), gamma 12(Gng12) |
| P63213 | 14702 | Mus musculus | guanine nucleotide binding protein (G protein), gamma 2(Gng2) |
| Q61016 | 14708 | Mus musculus | guanine nucleotide binding protein (G protein), gamma 7(Gng7) |
| P21278 | 14672 | Mus musculus | guanine nucleotide binding protein, alpha 11(Gna11) |
| P27600 | 14673 | Mus musculus | guanine nucleotide binding protein, alpha 12(Gna12) |
| P18872 | 14681 | Mus musculus | guanine nucleotide binding protein, alpha O(Gnao1) |
| O70443 | 14687 | Mus musculus | guanine nucleotide binding protein, alpha z subunit(Gnaz) |
| Q64520 | 14923 | Mus musculus | guanylate kinase 1(Guk1) |
| P10922 | 14958 | Mus musculus | H1.0 linker histone(H1f0) |
| P43274 | 50709 | Mus musculus | H1.4 linker histone, cluster member(H1f4) |
| Q64523 | 319176 | Mus musculus | H2A clustered histone 20(H2ac20) |
| Q8BFU2 | 319162 | Mus musculus | H2A clustered histone 25(H2ac25) |
| Q8CGP1 | 319184 | Mus musculus | H2B clustered histone 12(H2bc12) |
| Q64525 | 319189 | Mus musculus | H2B clustered histone 18(H2bc18) |
| Q64478 | 319182 | Mus musculus | H2B clustered histone 9(H2bc9) |
| P84244 | 15078 | Mus musculus | H3.3 histone A(H3f3a) |
| P84244 | 15081 | Mus musculus | H3.3 histone B(H3f3b) |
| P62806 | 326619 | Mus musculus | H4 clustered histone 1(H4c1) |
| P62806 | 319159 | Mus musculus | H4 clustered histone 11(H4c11) |
| P62806 | 319160 | Mus musculus | H4 clustered histone 12(H4c12) |
| P62806 | 97122 | Mus musculus | H4 clustered histone 14(H4c14) |
| P62806 | 1E+08 | Mus musculus | H4 clustered histone 17(H4c17) |
| P62806 | 319161 | Mus musculus | H4 clustered histone 18(H4c18) |
| P62806 | 326620 | Mus musculus | H4 clustered histone 2(H4c2) |
| P62806 | 319155 | Mus musculus | H4 clustered histone 3(H4c3) |

| | | | |
|---|---|---|---|
| P62806 | 319156 | Mus musculus | H4 clustered histone 4(H4c4) |
| P62806 | 319157 | Mus musculus | H4 clustered histone 6(H4c6) |
| P62806 | 69386 | Mus musculus | H4 clustered histone 8(H4c8) |
| P62806 | 319158 | Mus musculus | H4 clustered histone 9(H4c9) |
| P62806 | 320332 | Mus musculus | H4 histone 16(H4c16) |
| Q61646 | 15439 | Mus musculus | haptoglobin(Hp) |
| Q9JMG7 | 29877 | Mus musculus | HDGF like 3(Hdgfl3) |
| Q61699 | 15505 | Mus musculus | heat shock 105kDa/110kDa protein 1(Hsph1) |
| Q64433 | 15528 | Mus musculus | heat shock protein 1 (chaperonin 10)(Hspe1) |
| P63038 | 15510 | Mus musculus | heat shock protein 1 (chaperonin)(Hspd1) |
| P14602 | 15507 | Mus musculus | heat shock protein 1(Hspb1) |
| Q8K0U4 | 73442 | Mus musculus | heat shock protein 12A(Hspa12a) |
| Q61696 | 193740 | Mus musculus | heat shock protein 1A(Hspa1a) |
| P16627 | 15482 | Mus musculus | heat shock protein 1-like(Hspa1l) |
| P17156 | 15512 | Mus musculus | heat shock protein 2(Hspa2) |
| P48722 | 18415 | Mus musculus | heat shock protein 4 like(Hspa4l) |
| Q61316 | 15525 | Mus musculus | heat shock protein 4(Hspa4) |
| P20029 | 14828 | Mus musculus | heat shock protein 5(Hspa5) |
| P63017 | 15481 | Mus musculus | heat shock protein 8(Hspa8) |
| P38647 | 15526 | Mus musculus | heat shock protein 9(Hspa9) |
| P11499 | 15516 | Mus musculus | heat shock protein 90 alpha (cytosolic), class B member 1(Hsp90ab1) |
| P07901 | 15519 | Mus musculus | heat shock protein 90, alpha (cytosolic), class A member 1(Hsp90aa1) |
| P08113 | 22027 | Mus musculus | heat shock protein 90, beta (Grp94), member 1(Hsp90b1) |
| Q9R257 | 15199 | Mus musculus | heme binding protein 1(Hebp1) |
| P01942 | 15122 | Mus musculus | hemoglobin alpha, adult chain 1(Hba-a1) |
| P02088 | 15129 | Mus musculus | hemoglobin, beta adult major chain(Hbb-b1) |
| P02089 | 15130 | Mus musculus | hemoglobin, beta adult minor chain(Hbb-b2) |
| Q91X72 | 15458 | Mus musculus | hemopexin(Hpx) |
| P51859 | 15191 | Mus musculus | heparin binding growth factor(Hdgf) |
| Q640R3 | 72927 | Mus musculus | hepatocyte cell adhesion molecule(Hepacam) |
| Q99020 | 15384 | Mus musculus | heterogeneous nuclear ribonucleoprotein A/B(Hnrnpab) |
| P49312 | 15382 | Mus musculus | heterogeneous nuclear ribonucleoprotein A1(Hnrnpa1) |
| O88569 | 53379 | Mus musculus | heterogeneous nuclear ribonucleoprotein A2/B1(Hnrnpa2b1) |
| Q8BG05 | 229279 | Mus musculus | heterogeneous nuclear ribonucleoprotein A3(Hnrnpa3) |
| Q9Z204 | 15381 | Mus musculus | heterogeneous nuclear ribonucleoprotein C(Hnrnpc) |

| | | | |
|---|---|---|---|
| Q60668 | 11991 | Mus musculus | heterogeneous nuclear ribonucleoprotein D(Hnrnpd) |
| Q9Z130 | 50926 | Mus musculus | heterogeneous nuclear ribonucleoprotein D-like(Hnrnpdl) |
| Q9Z2X1 | 98758 | Mus musculus | heterogeneous nuclear ribonucleoprotein F(Hnrnpf) |
| O35737 | 59013 | Mus musculus | heterogeneous nuclear ribonucleoprotein H1(Hnrnph1) |
| P70333 | 56258 | Mus musculus | heterogeneous nuclear ribonucleoprotein H2(Hnrnph2) |
| P61979 | 15387 | Mus musculus | heterogeneous nuclear ribonucleoprotein K(Hnrnpk) |
| Q9D0E1 | 76936 | Mus musculus | heterogeneous nuclear ribonucleoprotein M(Hnrnpm) |
| P17710 | 15275 | Mus musculus | hexokinase 1(Hk1) |
| Q99LI8 | 15239 | Mus musculus | HGF-regulated tyrosine kinase substrate(Hgs) |
| P63158 | 15289 | Mus musculus | high mobility group box 1(Hmgb1) |
| P84075 | 15444 | Mus musculus | hippocalcin(Hpca) |
| P62748 | 53602 | Mus musculus | hippocalcin-like 1(Hpcal1) |
| P84075 | 53602 | Mus musculus | hippocalcin-like 1(Hpcal1) |
| Q8BGZ1 | 170638 | Mus musculus | hippocalcin-like 4(Hpcal4) |
| P70349 | 15254 | Mus musculus | histidine triad nucleotide binding protein 1(Hint1) |
| Q9D0S9 | 68917 | Mus musculus | histidine triad nucleotide binding protein 2(Hint2) |
| P01898 | 15007 | Mus musculus | histocompatibility 2, Q region locus 10(H2-Q10) |
| Q9Z2Y3 | 26556 | Mus musculus | homer scaffolding protein 1(Homer1) |
| Q99JP6 | 26558 | Mus musculus | homer scaffolding protein 3(Homer3) |
| O09173 | 15233 | Mus musculus | homogentisate 1, 2-dioxygenase(Hgd) |
| Q9QUP5 | 12950 | Mus musculus | hyaluronan and proteoglycan link protein 1(Hapln1) |
| Q9ESM3 | 73940 | Mus musculus | hyaluronan and proteoglycan link protein 2(Hapln2) |
| Q99KB8 | 14651 | Mus musculus | hydroxyacyl glutathione hydrolase(Hagh) |
| Q8BMS1 | 97212 | Mus musculus | hydroxyacyl-CoA dehydrogenase trifunctional multienzyme complex subunit alpha(Hadha) |
| Q61425 | 15107 | Mus musculus | hydroxyacyl-Coenzyme A dehydrogenase(Hadh) |
| O08756 | 15108 | Mus musculus | hydroxysteroid (17-beta) dehydrogenase 10(Hsd17b10) |
| P50172 | 15483 | Mus musculus | hydroxysteroid 11-beta dehydrogenase 1(Hsd11b1) |
| P00493 | 15452 | Mus musculus | hypoxanthine phosphoribosyltransferase 1(Hprt1) |
| Q9JKR6 | 12282 | Mus musculus | hypoxia up-regulated 1(Hyou1) |
| P01872 | 16019 | Mus musculus | immunoglobulin heavy constant mu(Ighm) |
| P01837 | 16071 | Mus musculus | immunoglobulin kappa constant(Igkc) |
| Q8R366 | 140559 | Mus musculus | immunoglobulin superfamily, member 8(Igsf8) |
| O55091 | 16210 | Mus musculus | impact, RWD domain protein(Impact) |
| P40936 | 21743 | Mus musculus | indolethylamine N-methyltransferase(Inmt) |
| Q8CAQ8 | 76614 | Mus musculus | inner membrane protein, mitochondrial(Immt) |

| | | | |
|---|---|---|---|
| Q9D892 | 16434 | Mus musculus | inosine triphosphatase (nucleoside triphosphate pyrophosphatase)(Itpa) |
| A6X935 | 16427 | Mus musculus | inter alpha-trypsin inhibitor, heavy chain 4(Itih4) |
| Q3UZ18 | 93697 | Mus musculus | interactor of little elongation complex ELL subunit 2(Ice2) |
| Q60625 | 15898 | Mus musculus | intercellular adhesion molecule 5, telencephalin(Icam5) |
| P46660 | 226180 | Mus musculus | internexin neuronal intermediate filament protein, alpha(Ina) |
| O88844 | 15926 | Mus musculus | isocitrate dehydrogenase 1 (NADP+), soluble(Idh1) |
| P54071 | 269951 | Mus musculus | isocitrate dehydrogenase 2 (NADP+), mitochondrial(Idh2) |
| Q9D6R2 | 67834 | Mus musculus | isocitrate dehydrogenase 3 (NAD+) alpha(Idh3a) |
| P70404 | 15929 | Mus musculus | isocitrate dehydrogenase 3 (NAD+), gamma(Idh3g) |
| Q9JHI5 | 56357 | Mus musculus | isovaleryl coenzyme A dehydrogenase(Ivd) |
| Q9D8B7 | 83964 | Mus musculus | junction adhesion molecule 3(Jam3) |
| P70168 | 16211 | Mus musculus | karyopherin subunit beta 1(Kpnb1) |
| Q61414 | 16665 | Mus musculus | keratin 15(Krt15) |
| P05784 | 16668 | Mus musculus | keratin 18(Krt18) |
| Q3TTY5 | 16681 | Mus musculus | keratin 2(Krt2) |
| Q9D312 | 66809 | Mus musculus | keratin 20(Krt20) |
| P11679 | 16691 | Mus musculus | keratin 8(Krt8) |
| P97328 | 16548 | Mus musculus | ketohexokinase(Khk) |
| Q3U0V1 | 16549 | Mus musculus | KH-type splicing regulatory protein(Khsrp) |
| Q9QXL2 | 16564 | Mus musculus | kinesin family member 21A(Kif21a) |
| E9Q5G3 | 71819 | Mus musculus | kinesin family member 23(Kif23) |
| P28740 | 16563 | Mus musculus | kinesin family member 2A(Kif2a) |
| Q61768 | 16573 | Mus musculus | kinesin family member 5B(Kif5b) |
| O88448 | 16594 | Mus musculus | kinesin light chain 2(Klc2) |
| O08677 | 16644 | Mus musculus | kininogen 1(Kng1) |
| Q6PHZ8 | 80334 | Mus musculus | Kv channel interacting protein 4(Kcnip4) |
| P11627 | 16728 | Mus musculus | L1 cell adhesion molecule(L1cam) |
| Q8BN59 | 67557 | Mus musculus | La ribonucleoprotein 6, translational regulator(Larp6) |
| Q99KR3 | 212442 | Mus musculus | lactamase, beta 2(Lactb2) |
| P06151 | 16828 | Mus musculus | lactate dehydrogenase A(Ldha) |
| P16125 | 16832 | Mus musculus | lactate dehydrogenase B(Ldhb) |
| P48678 | 16905 | Mus musculus | lamin A(Lmna) |
| Q61292 | 16779 | Mus musculus | laminin, beta 2(Lamb2) |
| Q61292 | 226519 | Mus musculus | laminin, gamma 1(Lamc1) |
| P02468 | 226519 | Mus musculus | laminin, gamma 1(Lamc1) |

| | | | |
|---|---|---|---|
| O89112 | 14768 | Mus musculus | LanC (bacterial lantibiotic synthetase component C)-like 1(Lancl1) |
| P70202 | 17035 | Mus musculus | latexin(Lxn) |
| Q9DBH5 | 66890 | Mus musculus | lectin, mannose-binding 2(Lman2) |
| Q9D0F3 | 70361 | Mus musculus | lectin, mannose-binding, 1(Lman1) |
| Q9CPY7 | 66988 | Mus musculus | leucine aminopeptidase 3(Lap3) |
| Q8K3C3 | 69151 | Mus musculus | leucine zipper and CTNNBIP1 domain containing(Lzic) |
| Q9Z2I0 | 56384 | Mus musculus | leucine zipper-EF-hand containing transmembrane protein 1(Letm1) |
| Q9JIA1 | 56839 | Mus musculus | leucine-rich repeat LGI family, member 1(Lgi1) |
| Q61792 | 16796 | Mus musculus | LIM and SH3 protein 1(Lasp1) |
| Q8BLK3 | 268890 | Mus musculus | limbic system-associated membrane protein(Lsamp) |
| Q8JZS0 | 108030 | Mus musculus | lin-7 homolog A, crumbs cell polarity complex component(Lin7a) |
| O88952 | 22343 | Mus musculus | lin-7 homolog C, crumbs cell polarity complex component(Lin7c) |
| Q91ZX7 | 16971 | Mus musculus | low density lipoprotein receptor-related protein 1(Lrp1) |
| Q9WUC3 | 23934 | Mus musculus | lymphocyte antigen 6 family member H(Ly6h) |
| Q9WTL7 | 26394 | Mus musculus | lysophospholipase 2(Lypla2) |
| P11438 | 16783 | Mus musculus | lysosomal-associated membrane protein 1(Lamp1) |
| P17047 | 16784 | Mus musculus | lysosomal-associated membrane protein 2(Lamp2) |
| P34884 | 17319 | Mus musculus | macrophage migration inhibitory factor (glycosylation-inhibiting factor)(Mif) |
| P11588 | 17840 | Mus musculus | major urinary protein 1(Mup1) |
| B5X0G2 | 1E+08 | Mus musculus | major urinary protein 17(Mup17) |
| P11589 | 17841 | Mus musculus | major urinary protein 2(Mup2) |
| Q5FW60 | 381530 | Mus musculus | major urinary protein 20(Mup20) |
| P04939 | 17842 | Mus musculus | major urinary protein 3(Mup3) |
| P14152 | 17449 | Mus musculus | malate dehydrogenase 1, NAD (soluble)(Mdh1) |
| P08249 | 17448 | Mus musculus | malate dehydrogenase 2, NAD (mitochondrial)(Mdh2) |
| P60755 | 320772 | Mus musculus | MAM domain containing glycosylphosphatidylinositol anchor 2(Mdga2) |
| A2AJI0 | 245877 | Mus musculus | MAP7 domain containing 1(Map7d1) |
| A2AEY4 | 320923 | Mus musculus | MAP7 domain containing 3(Map7d3) |
| P28667 | 17357 | Mus musculus | MARCKS-like 1(Marcksl1) |
| P70290 | 17524 | Mus musculus | membrane protein, palmitoylated(Mpp1) |
| Q99J99 | 246221 | Mus musculus | mercaptopyruvate sulfurtransferase(Mpst) |
| P20917 | 50994 | Mus musculus | metastasis associated gene 2(Mtag2) |
| Q91X83 | 11720 | Mus musculus | methionine adenosyltransferase 1A(Mat1a) |
| Q9D6Y7 | 110265 | Mus musculus | methionine sulfoxide reductase A(Msra) |
| Q91VS7 | 56615 | Mus musculus | microsomal glutathione S-transferase 1(Mgst1) |

| | | | |
|---|---|---|---|
| Q9QYR6 | 17754 | Mus musculus | microtubule-associated protein 1 A(Map1a) |
| Q91VR7 | 66734 | Mus musculus | microtubule-associated protein 1 light chain 3 alpha(Map1lc3a) |
| Q9CQV6 | 67443 | Mus musculus | microtubule-associated protein 1 light chain 3 beta(Map1lc3b) |
| P14873 | 17755 | Mus musculus | microtubule-associated protein 1B(Map1b) |
| P20357 | 17756 | Mus musculus | microtubule-associated protein 2(Map2) |
| P27546 | 17758 | Mus musculus | microtubule-associated protein 4(Map4) |
| Q7TSJ2 | 17760 | Mus musculus | microtubule-associated protein 6(Map6) |
| P10637 | 17762 | Mus musculus | microtubule-associated protein tau(Mapt) |
| Q61166 | 13589 | Mus musculus | microtubule-associated protein, RP/EB family, member 1(Mapre1) |
| Q8R001 | 212307 | Mus musculus | microtubule-associated protein, RP/EB family, member 2(Mapre2) |
| Q6PER3 | 100732 | Mus musculus | microtubule-associated protein, RP/EB family, member 3(Mapre3) |
| Q922Q1 | 67247 | Mus musculus | mitochondrial amidoxime reducing component 2(Mtarc2) |
| Q8R404 | 224904 | Mus musculus | mitochondrial contact site and cristae organizing system subunit 13(Micos13) |
| P63030 | 55951 | Mus musculus | mitochondrial pyruvate carrier 1(Mpc1) |
| Q9DB15 | 56282 | Mus musculus | mitochondrial ribosomal protein L12(Mrpl12) |
| P63085 | 26413 | Mus musculus | mitogen-activated protein kinase 1(Mapk1) |
| P31938 | 26395 | Mus musculus | mitogen-activated protein kinase kinase 1(Map2k1) |
| P26041 | 17698 | Mus musculus | moesin(Msn) |
| Q922B1 | 107227 | Mus musculus | mono-ADP ribosylhydrolase 1(Macrod1) |
| P28665 | 17836 | Mus musculus | murinoglobulin 1(Mug1) |
| Q9WUB3 | 19309 | Mus musculus | muscle glycogen phosphorylase(Pygm) |
| Q9EQS3 | 56309 | Mus musculus | MYC binding protein(Mycbp) |
| Q8C854 | 17876 | Mus musculus | myelin basic protein expression factor 2, repressor(Myef2) |
| P04370 | 17196 | Mus musculus | myelin basic protein(Mbp) |
| Q61885 | 17441 | Mus musculus | myelin oligodendrocyte glycoprotein(Mog) |
| P27573 | 17528 | Mus musculus | myelin protein zero(Mpz) |
| P20917 | 17136 | Mus musculus | myelin-associated glycoprotein(Mag) |
| Q9D2P8 | 17433 | Mus musculus | myelin-associated oligodendrocytic basic protein(Mobp) |
| P04247 | 17189 | Mus musculus | myoglobin(Mb) |
| A2ABU4 | 242702 | Mus musculus | myomesin family, member 3(Myom3) |
| Q5SYD0 | 338367 | Mus musculus | myosin ID(Myo1d) |
| P97457 | 17907 | Mus musculus | myosin light chain, phosphorylatable, fast skeletal muscle(Mylpf) |
| Q61879 | 77579 | Mus musculus | myosin, heavy polypeptide 10, non-muscle(Myh10) |
| P05977 | 17901 | Mus musculus | myosin, light polypeptide 1(Myl1) |
| P09542 | 17897 | Mus musculus | myosin, light polypeptide 3(Myl3) |

| | | | |
|---|---|---|---|
| Q60605 | 17904 | Mus musculus | myosin, light polypeptide 6, alkali, smooth muscle and non-muscle(Myl6) |
| Q8CI43 | 216459 | Mus musculus | myosin, light polypeptide 6B(Myl6b) |
| Q9CQ19 | 98932 | Mus musculus | myosin, light polypeptide 9, regulatory(Myl9) |
| P62774 | 14489 | Mus musculus | myotrophin(Mtpn) |
| P26645 | 17118 | Mus musculus | myristoylated alanine rich protein kinase C substrate(Marcks) |
| Q9WV54 | 11886 | Mus musculus | N-acylsphingosine amidohydrolase 1(Asah1) |
| Q8K4Z3 | 246703 | Mus musculus | NAD(P)HX epimerase(Naxe) |
| Q91VD9 | 227197 | Mus musculus | NADH:ubiquinone oxidoreductase core subunit S1(Ndufs1) |
| Q91WD5 | 226646 | Mus musculus | NADH:ubiquinone oxidoreductase core subunit S2(Ndufs2) |
| Q9DCT2 | 68349 | Mus musculus | NADH:ubiquinone oxidoreductase core subunit S3(Ndufs3) |
| Q99LY9 | 595136 | Mus musculus | NADH:ubiquinone oxidoreductase core subunit S5(Ndufs5) |
| P52503 | 407785 | Mus musculus | NADH:ubiquinone oxidoreductase core subunit S6(Ndufs6) |
| Q8K3J1 | 225887 | Mus musculus | NADH:ubiquinone oxidoreductase core subunit S8(Ndufs8) |
| Q91YT0 | 17995 | Mus musculus | NADH:ubiquinone oxidoreductase core subunit V1(Ndufv1) |
| Q9D6J6 | 72900 | Mus musculus | NADH:ubiquinone oxidoreductase core subunit V2(Ndufv2) |
| Q9D8B4 | 69875 | Mus musculus | NADH:ubiquinone oxidoreductase subunit A11(Ndufa11) |
| Q7TMF3 | 66414 | Mus musculus | NADH:ubiquinone oxidoreductase subunit A12(Ndufa12) |
| Q9CQ75 | 17991 | Mus musculus | NADH:ubiquinone oxidoreductase subunit A2(Ndufa2) |
| Q9CPP6 | 68202 | Mus musculus | NADH:ubiquinone oxidoreductase subunit A5(Ndufa5) |
| Q9CQZ5 | 67130 | Mus musculus | NADH:ubiquinone oxidoreductase subunit A6(Ndufa6) |
| Q9Z1P6 | 66416 | Mus musculus | NADH:ubiquinone oxidoreductase subunit A7(Ndufa7) |
| Q9DCJ5 | 68375 | Mus musculus | NADH:ubiquinone oxidoreductase subunit A8(Ndufa8) |
| Q9DC69 | 66108 | Mus musculus | NADH:ubiquinone oxidoreductase subunit A9(Ndufa9) |
| Q9CR21 | 70316 | Mus musculus | NADH:ubiquinone oxidoreductase subunit AB1(Ndufab1) |
| P0DN34 | 1,03E+08 | Mus musculus | NADH:ubiquinone oxidoreductase subunit B1(Ndufb1) |
| Q9DCS9 | 68342 | Mus musculus | NADH:ubiquinone oxidoreductase subunit B10(Ndufb10) |
| O09111 | 104130 | Mus musculus | NADH:ubiquinone oxidoreductase subunit B11(Ndufb11) |
| Q9CQZ6 | 66495 | Mus musculus | NADH:ubiquinone oxidoreductase subunit B3(Ndufb3) |
| Q9CQC7 | 68194 | Mus musculus | NADH:ubiquinone oxidoreductase subunit B4(Ndufb4) |
| Q9CQH3 | 66046 | Mus musculus | NADH:ubiquinone oxidoreductase subunit B5(Ndufb5) |
| Q9D6J5 | 67264 | Mus musculus | NADH:ubiquinone oxidoreductase subunit B8(Ndufb8) |
| P52503 | 623286 | Mus musculus | NADH:ubiquinone oxidoreductase subunit S6B(Ndufs6b) |
| P70670 | 17938 | Mus musculus | nascent polypeptide-associated complex alpha polypeptide(Naca) |
| Q62425 | 17992 | Mus musculus | Ndufa4, mitochondrial complex associated(Ndufa4) |
| Q9DC07 | 74103 | Mus musculus | nebulette(Nebl) |

| | | | |
|---|---|---|---|
| Q9CR95 | 67602 | Mus musculus | NECAP endocytosis associated 1(Necap1) |
| Q9JKF6 | 58235 | Mus musculus | nectin cell adhesion molecule 1(Nectin1) |
| Q9DB05 | 108124 | Mus musculus | N-ethylmaleimide sensitive fusion protein attachment protein alpha(Napa) |
| P28663 | 17957 | Mus musculus | N-ethylmaleimide sensitive fusion protein attachment protein beta(Napb) |
| Q9CWZ7 | 108123 | Mus musculus | N-ethylmaleimide sensitive fusion protein attachment protein gamma(Napg) |
| P46460 | 18195 | Mus musculus | N-ethylmaleimide sensitive fusion protein(Nsf) |
| P13595 | 17967 | Mus musculus | neural cell adhesion molecule 1(Ncam1) |
| O35136 | 17968 | Mus musculus | neural cell adhesion molecule 2(Ncam2) |
| P29595 | 18002 | Mus musculus | neural precursor cell expressed, developmentally down-regulated gene 8(Nedd8) |
| Q91X97 | 52589 | Mus musculus | neurocalcin delta(Ncald) |
| Q9Z0E0 | 26562 | Mus musculus | neurochondrin(Ncdn) |
| Q810U3 | 269116 | Mus musculus | neurofascin(Nfasc) |
| P19246 | 380684 | Mus musculus | neurofilament, heavy polypeptide(Nefh) |
| P08551 | 18039 | Mus musculus | neurofilament, light polypeptide(Nefl) |
| P08553 | 18040 | Mus musculus | neurofilament, medium polypeptide(Nefm) |
| P60761 | 64011 | Mus musculus | neurogranin(Nrgn) |
| Q8BYM5 | 245537 | Mus musculus | neuroligin 3(Nlgn3) |
| Q8BNY6 | 14299 | Mus musculus | neuronal calcium sensor 1(Ncs1) |
| Q810U4 | 319504 | Mus musculus | neuronal cell adhesion molecule(Nrcam) |
| Q80Z24 | 320840 | Mus musculus | neuronal growth regulator 1(Negr1) |
| Q62443 | 18164 | Mus musculus | neuronal pentraxin 1(Nptx1) |
| Q99J85 | 73340 | Mus musculus | neuronal pentraxin receptor(Nptxr) |
| P97300 | 20320 | Mus musculus | neuroplastin(Nptn) |
| Q99PJ0 | 235106 | Mus musculus | neurotrimin(Ntm) |
| P15209 | 18212 | Mus musculus | neurotrophic tyrosine kinase, receptor, type 2(Ntrk2) |
| Q9QZ23 | 56748 | Mus musculus | NFU1 iron-sulfur cluster scaffold(Nfu1) |
| P70441 | 26941 | Mus musculus | NHERF family PDZ scaffold protein 1(Nherf1) |
| O55126 | 14467 | Mus musculus | nipsnap homolog 2(Nipsnap2) |
| Q6WKZ7 | 329416 | Mus musculus | nitric oxide synthase trafficker(Nostrin) |
| Q9JHW2 | 52633 | Mus musculus | nitrilase family, member 2(Nit2) |
| P15532 | 18102 | Mus musculus | NME/NM23 nucleoside diphosphate kinase 1(Nme1) |
| Q01768 | 18103 | Mus musculus | NME/NM23 nucleoside diphosphate kinase 2(Nme2) |
| Q8K2T1 | 67824 | Mus musculus | NmrA-like family domain containing 1(Nmral1) |
| Q9QYG0 | 29811 | Mus musculus | N-myc downstream regulated gene 2(Ndrg2) |
| Q9CZ44 | 386649 | Mus musculus | NSFL1 (p97) cofactor (p47)(Nsfl1c) |

| | | | |
|---|---|---|---|
| P61971 | 68051 | Mus musculus | nuclear transport factor 2(Nutf2) |
| Q02819 | 18220 | Mus musculus | nucleobindin 1(Nucb1) |
| Q61937 | 18148 | Mus musculus | nucleophosmin 1(Npm1) |
| Q78ZA7 | 17955 | Mus musculus | nucleosome assembly protein 1-like 4(Nap1l4) |
| O35685 | 18221 | Mus musculus | nudC nuclear distribution protein(Nudc) |
| Q99P30 | 67528 | Mus musculus | nudix hydrolase 7(Nudt7) |
| O88998 | 56177 | Mus musculus | olfactomedin 1(Olfm1) |
| Q64288 | 18378 | Mus musculus | olfactory marker protein(Omp) |
| Q63912 | 18377 | Mus musculus | oligodendrocyte myelin glycoprotein(Omg) |
| P58281 | 74143 | Mus musculus | OPA1, mitochondrial dynamin like GTPase(Opa1) |
| P29758 | 18242 | Mus musculus | ornithine aminotransferase(Oat) |
| P11725 | 18416 | Mus musculus | ornithine transcarbamylase(Otc) |
| Q7TQI3 | 107260 | Mus musculus | OTU domain, ubiquitin aldehyde binding 1(Otub1) |
| Q9D4B2 | 74407 | Mus musculus | outer dynein arm complex subunit 4(Odad4) |
| Q4KMM3 | 170719 | Mus musculus | oxidation resistance 1(Oxr1) |
| Q60597 | 18293 | Mus musculus | oxoglutarate (alpha-ketoglutarate) dehydrogenase (lipoamide)(Ogdh) |
| O88643 | 18479 | Mus musculus | p21 (RAC1) activated kinase 1(Pak1) |
| O88531 | 19063 | Mus musculus | palmitoyl-protein thioesterase 1(Ppt1) |
| Q9Z0P4 | 18483 | Mus musculus | paralemmin(Palm) |
| Q8R326 | 66645 | Mus musculus | paraspeckle protein 1(Pspc1) |
| Q9D0J8 | 69202 | Mus musculus | parathymosin(Ptms) |
| Q99LX0 | 57320 | Mus musculus | Parkinson disease (autosomal recessive, early onset) 7(Park7) |
| P32848 | 19293 | Mus musculus | parvalbumin(Pvalb) |
| O70400 | 54132 | Mus musculus | PDZ and LIM domain 1 (elfin)(Pdlim1) |
| Q9QUR7 | 23988 | Mus musculus | peptidyl-prolyl cis/trans isomerase, NIMA-interacting 1(Pin1) |
| Q9D0W5 | 68816 | Mus musculus | peptidylprolyl isomerase (cyclophilin)-like 1(Ppil1) |
| P17742 | 268373 | Mus musculus | peptidylprolyl isomerase A(Ppia) |
| P24369 | 19035 | Mus musculus | peptidylprolyl isomerase B(Ppib) |
| Q99KR7 | 105675 | Mus musculus | peptidylprolyl isomerase F (cyclophilin F)(Ppif) |
| P35700 | 18477 | Mus musculus | peroxiredoxin 1(Prdx1) |
| Q61171 | 21672 | Mus musculus | peroxiredoxin 2(Prdx2) |
| P20108 | 11757 | Mus musculus | peroxiredoxin 3(Prdx3) |
| P99029 | 54683 | Mus musculus | peroxiredoxin 5(Prdx5) |
| O08709 | 11758 | Mus musculus | peroxiredoxin 6(Prdx6) |
| Q80TL4 | 230085 | Mus musculus | PHD finger protein 24(Phf24) |

| | | | |
|---|---|---|---|
| P16331 | 18478 | Mus musculus | phenylalanine hydroxylase(Pah) |
| P70296 | 23980 | Mus musculus | phosphatidylethanolamine binding protein 1(Pebp1) |
| P53810 | 18738 | Mus musculus | phosphatidylinositol transfer protein, alpha(Pitpna) |
| O70161 | 18717 | Mus musculus | phosphatidylinositol-4-phosphate 5-kinase, type 1 gamma(Pip5k1c) |
| P47857 | 18642 | Mus musculus | phosphofructokinase, muscle(Pfkm) |
| Q9WUA3 | 56421 | Mus musculus | phosphofructokinase, platelet(Pfkp) |
| P09411 | 18655 | Mus musculus | phosphoglycerate kinase 1(Pgk1) |
| P09041 | 18663 | Mus musculus | phosphoglycerate kinase 2(Pgk2) |
| Q9DBJ1 | 18648 | Mus musculus | phosphoglycerate mutase 1(Pgam1) |
| Q8CHP8 | 67078 | Mus musculus | phosphoglycolate phosphatase(Pgp) |
| Q9DAK9 | 75454 | Mus musculus | phosphohistidine phosphatase 1(Phpt1) |
| Q3TCN2 | 71772 | Mus musculus | phospholipase B domain containing 2(Plbd2) |
| Q9Z1B3 | 18795 | Mus musculus | phospholipase C, beta 1(Plcb1) |
| O35405 | 18807 | Mus musculus | phospholipase D family member 3(Pld3) |
| Q9DCL9 | 67054 | Mus musculus | phosphoribosylaminoimidazole carboxylase, phosphoribosylaminoribosylaminoimidazole, succinocarboxamide synthetase(Paics) |
| Q99K85 | 107272 | Mus musculus | phosphoserine aminotransferase 1(Psat1) |
| Q8BWR2 | 66193 | Mus musculus | PITH (C-terminal proteasome-interacting domain of thioredoxin-like) domain containing 1(Pithd1) |
| Q9DCU2 | 67801 | Mus musculus | plasma membrane proteolipid(Pllp) |
| Q99K51 | 102866 | Mus musculus | plastin 3 (T-isoform)(Pls3) |
| Q61206 | 18475 | Mus musculus | platelet-activating factor acetylhydrolase, isoform 1b, subunit 2(Pafah1b2) |
| Q2PFD7 | 234353 | Mus musculus | pleckstrin and Sec7 domain containing 3(Psd3) |
| Q8C886 | 231002 | Mus musculus | pleckstrin homology domain containing, family N member 1(Plekhn1) |
| Q9JIY0 | 67220 | Mus musculus | pleckstrin homology domain containing, family O member 1(Plekho1) |
| Q9QXS1 | 18810 | Mus musculus | plectin(Plec) |
| B2RXS4 | 140570 | Mus musculus | plexin B2(Plxnb2) |
| Q61990 | 18521 | Mus musculus | poly(rC) binding protein 2(Pcbp2) |
| P54099 | 18975 | Mus musculus | polymerase (DNA directed), gamma(Polg) |
| Q9JM63 | 16513 | Mus musculus | potassium inwardly-rectifying channel, subfamily J, member 10(Kcnj10) |
| P16388 | 16485 | Mus musculus | potassium voltage-gated channel, shaker-related subfamily, member 1(Kcna1) |
| P47915 | 1,15E+08 | Mus musculus | predicted gene 8940(Gm8940) |
| P63323 | 1,01E+08 | Mus musculus | predicted gene, 21571(Gm21571) |
| P45878 | 1,15E+08 | Mus musculus | predicted gene, 51400(Gm51400) |
| P47915 | 1,01E+08 | Mus musculus | predicted pseudogene 10913(Gm10913) |
| O70591 | 18637 | Mus musculus | prefoldin 2(Pfdn2) |
| Q03958 | 14976 | Mus musculus | prefoldin subunit 6(Pfdn6) |

| | | | |
|---|---|---|---|
| P04925 | 19122 | Mus musculus | prion protein(Prnp) |
| P62962 | 18643 | Mus musculus | profilin 1(Pfn1) |
| Q9JJV2 | 18645 | Mus musculus | profilin 2(Pfn2) |
| O55022 | 53328 | Mus musculus | progesterone receptor membrane component 1(Pgrmc1) |
| Q9WU78 | 18571 | Mus musculus | programmed cell death 6 interacting protein(Pdcd6ip) |
| P67778 | 18673 | Mus musculus | prohibitin 1(Phb1) |
| Q62048 | 18611 | Mus musculus | proliferation and apoptosis adaptor protein 15A(Pea15a) |
| P50580 | 18813 | Mus musculus | proliferation-associated 2G4(Pa2g4) |
| E9PUL5 | 69017 | Mus musculus | proline-rich transmembrane protein 2(Prrt2) |
| P09103 | 18453 | Mus musculus | prolyl 4-hydroxylase, beta polypeptide(P4hb) |
| Q91ZA3 | 110821 | Mus musculus | propionyl-Coenzyme A carboxylase, alpha polypeptide(Pcca) |
| Q61207 | 19156 | Mus musculus | prosaposin(Psap) |
| Q9R0Q7 | 56351 | Mus musculus | prostaglandin E synthase 3(Ptges3) |
| P62196 | 19184 | Mus musculus | protease (prosome, macropain) 26S subunit, ATPase 5(Psmc5) |
| P46471 | 19181 | Mus musculus | proteasome (prosome, macropain) 26S subunit, ATPase 2(Psmc2) |
| O88685 | 19182 | Mus musculus | proteasome (prosome, macropain) 26S subunit, ATPase 3(Psmc3) |
| O35226 | 19185 | Mus musculus | proteasome (prosome, macropain) 26S subunit, non-ATPase, 4(Psmd4) |
| P97371 | 19186 | Mus musculus | proteasome (prosome, macropain) activator subunit 1 (PA28 alpha)(Psme1) |
| O09061 | 19170 | Mus musculus | proteasome (prosome, macropain) subunit, beta type 1(Psmb1) |
| P99026 | 19172 | Mus musculus | proteasome (prosome, macropain) subunit, beta type 4(Psmb4) |
| Q9R1P4 | 26440 | Mus musculus | proteasome subunit alpha 1(Psma1) |
| O70435 | 19167 | Mus musculus | proteasome subunit alpha 3(Psma3) |
| Q9R1P0 | 26441 | Mus musculus | proteasome subunit alpha 4(Psma4) |
| Q9Z2U1 | 26442 | Mus musculus | proteasome subunit alpha 5(Psma5) |
| Q9QUM9 | 26443 | Mus musculus | proteasome subunit alpha 6(Psma6) |
| Q9Z2U0 | 26444 | Mus musculus | proteasome subunit alpha 7(Psma7) |
| P27773 | 14827 | Mus musculus | protein disulfide isomerase associated 3(Pdia3) |
| P08003 | 12304 | Mus musculus | protein disulfide isomerase associated 4(Pdia4) |
| Q922R8 | 71853 | Mus musculus | protein disulfide isomerase associated 6(Pdia6) |
| Q61644 | 23969 | Mus musculus | protein kinase C and casein kinase substrate in neurons 1(Pacsin1) |
| Q9WVE8 | 23970 | Mus musculus | protein kinase C and casein kinase substrate in neurons 2(Pacsin2) |
| O08795 | 19089 | Mus musculus | protein kinase C substrate 80K-H(Prkcsh) |
| P68404 | 18751 | Mus musculus | protein kinase C, beta(Prkcb) |
| P63318 | 18752 | Mus musculus | protein kinase C, gamma(Prkcg) |
| P05132 | 18747 | Mus musculus | protein kinase, cAMP dependent, catalytic, alpha(Prkaca) |

| | | | |
|---|---|---|---|
| P62141 | 19046 | Mus musculus | protein phosphatase 1 catalytic subunit beta(Ppp1cb) |
| P63087 | 19047 | Mus musculus | protein phosphatase 1 catalytic subunit gamma(Ppp1cc) |
| Q9ERT9 | 58200 | Mus musculus | protein phosphatase 1, regulatory inhibitor subunit 1A(Ppp1r1a) |
| Q60829 | 19049 | Mus musculus | protein phosphatase 1, regulatory inhibitor subunit 1B(Ppp1r1b) |
| P62715 | 19053 | Mus musculus | protein phosphatase 2 (formerly 2A), catalytic subunit, beta isoform(Ppp2cb) |
| Q76MZ3 | 51792 | Mus musculus | protein phosphatase 2, regulatory subunit A, alpha(Ppp2r1a) |
| Q7TNP2 | 73699 | Mus musculus | protein phosphatase 2, regulatory subunit A, beta(Ppp2r1b) |
| P63328 | 19055 | Mus musculus | protein phosphatase 3, catalytic subunit, alpha isoform(Ppp3ca) |
| P48455 | 19057 | Mus musculus | protein phosphatase 3, catalytic subunit, gamma isoform(Ppp3cc) |
| Q63810 | 19058 | Mus musculus | protein phosphatase 3, regulatory subunit B, alpha isoform (calcineurin B, type I)(Ppp3r1) |
| P18052 | 19262 | Mus musculus | protein tyrosine phosphatase receptor type A(Ptpra) |
| B9EKR1 | 19283 | Mus musculus | protein tyrosine phosphatase receptor type Z, polypeptide 1(Ptprz1) |
| P60469 | 76787 | Mus musculus | protein tyrosine phosphatase, receptor type, f polypeptide (PTPRF), interacting protein (liprin), alpha 3(Ppfia3) |
| P23506 | 18537 | Mus musculus | protein-L-isoaspartate (D-aspartate) O-methyltransferase 1(Pcmt1) |
| P60202 | 18823 | Mus musculus | proteolipid protein (myelin) 1(Plp1) |
| P26350 | 19231 | Mus musculus | prothymosin alpha(Ptma) |
| P61458 | 13180 | Mus musculus | pterin 4 alpha carbinolamine dehydratase/dimerization cofactor of hepatocyte nuclear factor 1 alpha (TCF1) 1(Pcbd1) |
| P42669 | 19290 | Mus musculus | purine rich element binding protein A(Pura) |
| O35295 | 19291 | Mus musculus | purine rich element binding protein B(Purb) |
| P63054 | 18546 | Mus musculus | Purkinje cell protein 4(Pcp4) |
| Q8K183 | 216134 | Mus musculus | pyridoxal (pyridoxine, vitamin B6) kinase(Pdxk) |
| Q9Z2Y8 | 114863 | Mus musculus | pyridoxal phosphate binding protein(Plpbp) |
| Q9D819 | 67895 | Mus musculus | pyrophosphatase (inorganic) 1(Ppa1) |
| Q9D051 | 68263 | Mus musculus | pyruvate dehydrogenase (lipoamide) beta(Pdhb) |
| Q8BKZ9 | 27402 | Mus musculus | pyruvate dehydrogenase complex, component X(Pdhx) |
| P35486 | 18597 | Mus musculus | pyruvate dehydrogenase E1 alpha 1(Pdha1) |
| P52480 | 18746 | Mus musculus | pyruvate kinase, muscle(Pkm) |
| Q61838 | 11287 | Mus musculus | PZP, alpha-2-macroglobulin like(Pzp) |
| Q8BVI4 | 110391 | Mus musculus | quinoid dihydropteridine reductase(Qdpr) |
| P61027 | 19325 | Mus musculus | RAB10, member RAS oncogene family(Rab10) |
| P46638 | 19326 | Mus musculus | RAB11B, member RAS oncogene family(Rab11b) |
| Q91V41 | 68365 | Mus musculus | RAB14, member RAS oncogene family(Rab14) |
| P62821 | 19324 | Mus musculus | RAB1A, member RAS oncogene family(Rab1a) |
| Q9D1G1 | 76308 | Mus musculus | RAB1B, member RAS oncogene family(Rab1b) |

| | | | |
|---|---|---|---|
| P35282 | 216344 | Mus musculus | RAB21, member RAS oncogene family(Rab21) |
| P53994 | 59021 | Mus musculus | RAB2A, member RAS oncogene family(Rab2a) |
| Q6PHN9 | 77407 | Mus musculus | RAB35, member RAS oncogene family(Rab35) |
| P63011 | 19339 | Mus musculus | RAB3A, member RAS oncogene family(Rab3a) |
| Q9CZT8 | 69908 | Mus musculus | RAB3B, member RAS oncogene family(Rab3b) |
| P62823 | 67295 | Mus musculus | RAB3C, member RAS oncogene family(Rab3c) |
| P61021 | 19344 | Mus musculus | RAB5B, member RAS oncogene family(Rab5b) |
| P35278 | 19345 | Mus musculus | RAB5C, member RAS oncogene family(Rab5c) |
| P61294 | 270192 | Mus musculus | RAB6B, member RAS oncogene family(Rab6b) |
| P51150 | 19349 | Mus musculus | RAB7, member RAS oncogene family(Rab7) |
| P47708 | 19894 | Mus musculus | rabphilin 3A(Rph3a) |
| P63001 | 19353 | Mus musculus | Rac family small GTPase 1(Rac1) |
| P60764 | 170758 | Mus musculus | Rac family small GTPase 3(Rac3) |
| P54728 | 19359 | Mus musculus | RAD23 homolog B, nucleotide excision repair protein(Rad23b) |
| P26043 | 19684 | Mus musculus | radixin(Rdx) |
| P34022 | 19385 | Mus musculus | RAN binding protein 1(Ranbp1) |
| P62827 | 19384 | Mus musculus | RAN, member RAS oncogene family(Ran) |
| E9Q912 | 229877 | Mus musculus | RAP1, GTP-GDP dissociation stimulator 1(Rap1gds1) |
| Q8BU31 | 72065 | Mus musculus | RAP2C, member of RAS oncogene family(Rap2c) |
| Q9QUI0 | 11848 | Mus musculus | ras homolog family member A(Rhoa) |
| P62746 | 11852 | Mus musculus | ras homolog family member B(Rhob) |
| P84096 | 56212 | Mus musculus | ras homolog family member G(Rhog) |
| Q99JI6 | 215449 | Mus musculus | RAS related protein 1b(Rap1b) |
| Q80ZJ1 | 76108 | Mus musculus | RAS related protein 2a(Rap2a) |
| P52760 | 15473 | Mus musculus | reactive intermediate imine deaminase A homolog(Rida) |
| Q60870 | 13476 | Mus musculus | receptor accessory protein 5(Reep5) |
| P68040 | 14694 | Mus musculus | receptor for activated C kinase 1(Rack1) |
| P24270 | 12285 | Mus musculus | recessive cataract(cac) |
| Q64374 | 19733 | Mus musculus | regucalcin(Rgn) |
| Q05186 | 19672 | Mus musculus | reticulocalbin 1(Rcn1) |
| Q8BP92 | 26611 | Mus musculus | reticulocalbin 2(Rcn2) |
| Q8K0T0 | 104001 | Mus musculus | reticulon 1(Rtn1) |
| Q9ES97 | 20168 | Mus musculus | reticulon 3(Rtn3) |
| Q99P72 | 68585 | Mus musculus | reticulon 4(Rtn4) |
| Q99PT1 | 192662 | Mus musculus | Rho GDP dissociation inhibitor alpha(Arhgdia) |

| | | | |
|---|---|---|---|
| Q5FWK3 | 228359 | Mus musculus | Rho GTPase activating protein 1(Arhgap1) |
| Q6ZWV3 | 110954 | Mus musculus | ribosomal protein L10(Rpl10) |
| Q9CXW4 | 67025 | Mus musculus | ribosomal protein L11(Rpl11) |
| P47963 | 270106 | Mus musculus | ribosomal protein L13(Rpl13) |
| P19253 | 22121 | Mus musculus | ribosomal protein L13A(Rpl13a) |
| Q9CR57 | 67115 | Mus musculus | ribosomal protein L14(Rpl14) |
| Q9CZM2 | 66480 | Mus musculus | ribosomal protein L15(Rpl15) |
| Q9CPR4 | 319195 | Mus musculus | ribosomal protein L17(Rpl17) |
| P35980 | 19899 | Mus musculus | ribosomal protein L18(Rpl18) |
| P62717 | 76808 | Mus musculus | ribosomal protein L18A(Rpl18a) |
| P84099 | 19921 | Mus musculus | ribosomal protein L19(Rpl19) |
| P62830 | 65019 | Mus musculus | ribosomal protein L23(Rpl23) |
| P62751 | 268449 | Mus musculus | ribosomal protein L23A(Rpl23a) |
| Q8BP67 | 68193 | Mus musculus | ribosomal protein L24(Rpl24) |
| P61255 | 19941 | Mus musculus | ribosomal protein L26(Rpl26) |
| P61358 | 19942 | Mus musculus | ribosomal protein L27(Rpl27) |
| P14115 | 26451 | Mus musculus | ribosomal protein L27A(Rpl27a) |
| P41105 | 19943 | Mus musculus | ribosomal protein L28(Rpl28) |
| P47915 | 19944 | Mus musculus | ribosomal protein L29(Rpl29) |
| P27659 | 27367 | Mus musculus | ribosomal protein L3(Rpl3) |
| P62889 | 19946 | Mus musculus | ribosomal protein L30(Rpl30) |
| P62900 | 114641 | Mus musculus | ribosomal protein L31(Rpl31) |
| P62911 | 19951 | Mus musculus | ribosomal protein L32(Rpl32) |
| Q9D1R9 | 68436 | Mus musculus | ribosomal protein L34(Rpl34) |
| Q6ZWV7 | 66489 | Mus musculus | ribosomal protein L35(Rpl35) |
| O55142 | 57808 | Mus musculus | ribosomal protein L35A(Rpl35a) |
| P61514 | 19981 | Mus musculus | ribosomal protein L37a(Rpl37a) |
| Q9JJI8 | 67671 | Mus musculus | ribosomal protein L38(Rpl38) |
| Q9D8E6 | 67891 | Mus musculus | ribosomal protein L4(Rpl4) |
| P47962 | 1,01E+08 | Mus musculus | ribosomal protein L5(Rpl5) |
| P47911 | 19988 | Mus musculus | ribosomal protein L6(Rpl6) |
| P14148 | 19989 | Mus musculus | ribosomal protein L7(Rpl7) |
| P12970 | 27176 | Mus musculus | ribosomal protein L7A(Rpl7a) |
| P62918 | 26961 | Mus musculus | ribosomal protein L8(Rpl8) |
| P51410 | 20005 | Mus musculus | ribosomal protein L9(Rpl9) |

| | | | |
|---|---|---|---|
| P14869 | 11837 | Mus musculus | ribosomal protein lateral stalk subunit P0(Rplp0) |
| P99027 | 67186 | Mus musculus | ribosomal protein lateral stalk subunit P2(Rplp2) |
| P62281 | 27207 | Mus musculus | ribosomal protein S11(Rps11) |
| P63323 | 20042 | Mus musculus | ribosomal protein S12(Rps12) |
| P63323 | 670832 | Mus musculus | ribosomal protein S12, pseudogene 10(Rps12-ps10) |
| P63323 | 434843 | Mus musculus | ribosomal protein S12, pseudogene 11(Rps12-ps11) |
| P63323 | 665463 | Mus musculus | ribosomal protein S12, pseudogene 12(Rps12-ps12) |
| P63323 | 665579 | Mus musculus | ribosomal protein S12, pseudogene 13(Rps12-ps13) |
| P63323 | 665611 | Mus musculus | ribosomal protein S12, pseudogene 14(Rps12-ps14) |
| P63323 | 665522 | Mus musculus | ribosomal protein S12, pseudogene 15(Rps12-ps15) |
| P63323 | 626048 | Mus musculus | ribosomal protein S12, pseudogene 16(Rps12-ps16) |
| P63323 | 1E+08 | Mus musculus | ribosomal protein S12, pseudogene 18(Rps12-ps18) |
| P63323 | 671641 | Mus musculus | ribosomal protein S12, pseudogene 24(Rps12-ps24) |
| P62301 | 68052 | Mus musculus | ribosomal protein S13(Rps13) |
| P62843 | 20054 | Mus musculus | ribosomal protein S15(Rps15) |
| P62245 | 267019 | Mus musculus | ribosomal protein S15A(Rps15a) |
| P62245 | 1E+08 | Mus musculus | ribosomal protein S15A, pseudogene 5(Rps15a-ps5) |
| P14131 | 20055 | Mus musculus | ribosomal protein S16(Rps16) |
| P63276 | 20068 | Mus musculus | ribosomal protein S17(Rps17) |
| P62270 | 20084 | Mus musculus | ribosomal protein S18(Rps18) |
| Q9CZX8 | 20085 | Mus musculus | ribosomal protein S19(Rps19) |
| P25444 | 16898 | Mus musculus | ribosomal protein S2(Rps2) |
| P60867 | 67427 | Mus musculus | ribosomal protein S20(Rps20) |
| Q9CQR2 | 66481 | Mus musculus | ribosomal protein S21(Rps21) |
| P62267 | 66475 | Mus musculus | ribosomal protein S23(Rps23) |
| P62849 | 20088 | Mus musculus | ribosomal protein S24(Rps24) |
| P62852 | 75617 | Mus musculus | ribosomal protein S25(Rps25) |
| P62983 | 78294 | Mus musculus | ribosomal protein S27A(Rps27a) |
| P62858 | 54127 | Mus musculus | ribosomal protein S28(Rps28) |
| P62274 | 20090 | Mus musculus | ribosomal protein S29(Rps29) |
| P62908 | 27050 | Mus musculus | ribosomal protein S3(Rps3) |
| P97351 | 20091 | Mus musculus | ribosomal protein S3A1(Rps3a1) |
| P62702 | 20102 | Mus musculus | ribosomal protein S4, X-linked(Rps4x) |
| P62754 | 20104 | Mus musculus | ribosomal protein S6(Rps6) |
| P62242 | 20116 | Mus musculus | ribosomal protein S8(Rps8) |

| | | | |
|---|---|---|---|
| Q6ZWN5 | 76846 | Mus musculus | ribosomal protein S9(Rps9) |
| P14206 | 16785 | Mus musculus | ribosomal protein SA(Rpsa) |
| Q99PL5 | 81910 | Mus musculus | ribosome binding protein 1(Rrbp1) |
| Q9CRC3 | 67290 | Mus musculus | RIKEN cDNA 3110040N11 gene(3110040N11Rik) |
| Q91V76 | 70984 | Mus musculus | RIKEN cDNA 4931406C07 gene(4931406C07Rik) |
| Q8R0A7 | 234797 | Mus musculus | RIKEN cDNA 6430548M08 gene(6430548M08Rik) |
| Q8BFU3 | 235315 | Mus musculus | ring finger protein 214(Rnf214) |
| P62878 | 56438 | Mus musculus | ring-box 1(Rbx1) |
| Q9D394 | 52822 | Mus musculus | RUN and FYVE domain containing 3(Rufy3) |
| Q9CQK7 | 66521 | Mus musculus | RWD domain containing 1(Rwdd1) |
| P56565 | 20193 | Mus musculus | S100 calcium binding protein A1(S100a1) |
| P97352 | 20196 | Mus musculus | S100 calcium binding protein A13(S100a13) |
| P50114 | 20203 | Mus musculus | S100 protein, beta polypeptide, neural(S100b) |
| P50247 | 269378 | Mus musculus | S-adenosylhomocysteine hydrolase(Ahcy) |
| Q80SW1 | 229709 | Mus musculus | S-adenosylhomocysteine hydrolase-like 1(Ahcyl1) |
| Q80U72 | 105782 | Mus musculus | scribbled planar cell polarity(Scrib) |
| Q80UK0 | 228071 | Mus musculus | SEC14 and spectrin domains 1(Sestd1) |
| Q99J08 | 67815 | Mus musculus | SEC14-like lipid binding 2(Sec14l2) |
| Q9CZC8 | 69938 | Mus musculus | secernin 1(Scrn1) |
| P36536 | 20224 | Mus musculus | secretion associated Ras related GTPase 1A(Sar1a) |
| Q03517 | 20254 | Mus musculus | secretogranin II(Scg2) |
| P47867 | 20255 | Mus musculus | secretogranin III(Scg3) |
| Q8K021 | 107767 | Mus musculus | secretory carrier membrane protein 1(Scamp1) |
| Q9JKD3 | 56807 | Mus musculus | secretory carrier membrane protein 5(Scamp5) |
| P17563 | 20341 | Mus musculus | selenium binding protein 1(Selenbp1) |
| Q63836 | 20342 | Mus musculus | selenium binding protein 2(Selenbp2) |
| Q64105 | 20751 | Mus musculus | sepiapterin reductase(Spr) |
| Q8C1B7 | 52398 | Mus musculus | septin 11(Septin11) |
| Q9Z1S5 | 24050 | Mus musculus | septin 3(Septin3) |
| Q9Z2Q6 | 18951 | Mus musculus | septin 5(Septin5) |
| Q9R1T4 | 56526 | Mus musculus | septin 6(Septin6) |
| O55131 | 235072 | Mus musculus | septin 7(Septin7) |
| Q8CHH9 | 20362 | Mus musculus | septin 8(Septin8) |
| P07758 | 20700 | Mus musculus | serine (or cysteine) peptidase inhibitor, clade A, member 1A(Serpina1a) |
| Q00897 | 20703 | Mus musculus | serine (or cysteine) peptidase inhibitor, clade A, member 1D(Serpina1d) |

| | | | |
|---|---|---|---|
| Q00898 | 20704 | Mus musculus | serine (or cysteine) peptidase inhibitor, clade A, member 1E(Serpina1e) |
| P07759 | 20714 | Mus musculus | serine (or cysteine) peptidase inhibitor, clade A, member 3K(Serpina3k) |
| P22599 | 20701 | Mus musculus | serine (or cysteine) preptidase inhibitor, clade A, member 1B(Serpina1b) |
| P50431 | 20425 | Mus musculus | serine hydroxymethyltransferase 1 (soluble)(Shmt1) |
| Q9QZX7 | 27364 | Mus musculus | serine racemase(Srr) |
| Q9EQU5 | 56086 | Mus musculus | SET nuclear oncogene(Set) |
| Q8VD37 | 73094 | Mus musculus | SH3-domain GRB2-like (endophilin) interacting protein 1(Sgip1) |
| Q62420 | 20404 | Mus musculus | SH3-domain GRB2-like 2(Sh3gl2) |
| Q8R3V5 | 227700 | Mus musculus | SH3-domain GRB2-like endophilin B2(Sh3glb2) |
| Q91V61 | 94280 | Mus musculus | sideroflexin 3(Sfxn3) |
| Q9CPW5 | 66256 | Mus musculus | signal sequence receptor, beta(Ssr2) |
| P70297 | 20844 | Mus musculus | signal transducing adaptor molecule (SH3 domain and ITAM motif) 1(Stam) |
| P97797 | 19261 | Mus musculus | signal-regulatory protein alpha(Sirpa) |
| Q8VDQ8 | 64383 | Mus musculus | sirtuin 2(Sirt2) |
| Q91VZ6 | 98366 | Mus musculus | small ArfGAP 1(Smap1) |
| Q8BJU0 | 52551 | Mus musculus | small glutamine-rich tetratricopeptide repeat (TPR)-containing, alpha(Sgta) |
| Q62376 | 20637 | Mus musculus | small nuclear ribonucleoprotein 70 (U1)(Snrnp70) |
| P62317 | 107686 | Mus musculus | small nuclear ribonucleoprotein D2(Snrpd2) |
| P62305 | 20643 | Mus musculus | small nuclear ribonucleoprotein E(Snrpe) |
| Q62189 | 53607 | Mus musculus | small nuclear ribonucleoprotein polypeptide A(Snrpa) |
| Q9Z172 | 20610 | Mus musculus | small ubiquitin-like modifier 3(Sumo3) |
| Q3UZP4 | 75744 | Mus musculus | small VCP/p97-interacting protein(Svip) |
| P97952 | 20266 | Mus musculus | sodium channel, voltage-gated, type I, beta(Scn1b) |
| Q56A07 | 72821 | Mus musculus | sodium channel, voltage-gated, type II, beta(Scn2b) |
| P43006 | 20511 | Mus musculus | solute carrier family 1 (glial high affinity glutamate transporter), member 2(Slc1a2) |
| P56564 | 20512 | Mus musculus | solute carrier family 1 (glial high affinity glutamate transporter), member 3(Slc1a3) |
| Q9JIS8 | 20498 | Mus musculus | solute carrier family 12, member 4(Slc12a4) |
| Q91V14 | 57138 | Mus musculus | solute carrier family 12, member 5(Slc12a5) |
| Q3TXX4 | 72961 | Mus musculus | solute carrier family 17 (sodium-dependent inorganic phosphate cotransporter), member 7(Slc17a7) |
| P32037 | 20527 | Mus musculus | solute carrier family 2 (facilitated glucose transporter), member 3(Slc2a3) |
| Q9CR62 | 67863 | Mus musculus | solute carrier family 25 (mitochondrial carrier oxoglutarate carrier), member 11(Slc25a11) |
| P48962 | 11739 | Mus musculus | solute carrier family 25 (mitochondrial carrier, adenine nucleotide translocator), member 4(Slc25a4) |
| P51881 | 11740 | Mus musculus | solute carrier family 25 (mitochondrial carrier, adenine nucleotide translocator), member 5(Slc25a5) |
| Q9D6M3 | 68267 | Mus musculus | solute carrier family 25 (mitochondrial carrier, glutamate), member 22(Slc25a22) |

| | | | |
|---|---|---|---|
| Q8VEM8 | 18674 | Mus musculus | solute carrier family 25 (mitochondrial carrier, phosphate carrier), member 3(Slc25a3) |
| Q3V132 | 73333 | Mus musculus | solute carrier family 25 (mitochondrial carrier; adenine nucleotide translocator), member 31(Slc25a31) |
| P10852 | 17254 | Mus musculus | solute carrier family 3 (activators of dibasic and neutral amino acid transport), member 2(Slc3a2) |
| O35633 | 22348 | Mus musculus | solute carrier family 32 (GABA vesicular transporter), member 1(Slc32a1) |
| O88343 | 54403 | Mus musculus | solute carrier family 4 (anion exchanger), member 4(Slc4a4) |
| Q8JZR6 | 59033 | Mus musculus | solute carrier family 4 (anion exchanger), member 8(Slc4a8) |
| Q5DTL9 | 94229 | Mus musculus | solute carrier family 4, sodium bicarbonate cotransporter-like, member 10(Slc4a10) |
| Q6X893 | 100434 | Mus musculus | solute carrier family 44, member 1(Slc44a1) |
| Q8BJI1 | 229706 | Mus musculus | solute carrier family 6 (neurotransmitter transporter), member 17(Slc6a17) |
| P31648 | 232333 | Mus musculus | solute carrier family 6 (neurotransmitter transporter, GABA), member 1(Slc6a1) |
| P31650 | 243616 | Mus musculus | solute carrier family 6 (neurotransmitter transporter, GABA), member 11(Slc6a11) |
| P70414 | 20541 | Mus musculus | solute carrier family 8 (sodium/calcium exchanger), member 1(Slc8a1) |
| Q8K596 | 110891 | Mus musculus | solute carrier family 8 (sodium/calcium exchanger), member 2(Slc8a2) |
| Q9QXZ6 | 28248 | Mus musculus | solute carrier organic anion transporter family, member 1a1(Slco1a1) |
| Q64442 | 20322 | Mus musculus | sorbitol dehydrogenase(Sord) |
| Q6P069 | 109552 | Mus musculus | sorcin(Sri) |
| O70493 | 55988 | Mus musculus | sorting nexin 12(Snx12) |
| Q9CWK8 | 67804 | Mus musculus | sorting nexin 2(Snx2) |
| Q6P8X1 | 72183 | Mus musculus | sorting nexin 6(Snx6) |
| P16546 | 20740 | Mus musculus | spectrin alpha, non-erythrocytic 1(Sptan1) |
| Q62261 | 20742 | Mus musculus | spectrin beta, non-erythrocytic 1(Sptbn1) |
| Q9WTX5 | 21402 | Mus musculus | S-phase kinase-associated protein 1(Skp1) |
| Q8VIJ6 | 71514 | Mus musculus | splicing factor proline/glutamine rich (polypyrimidine tract binding protein associated)(Sfpq) |
| Q9JMD3 | 56018 | Mus musculus | StAR related lipid transfer domain containing 10(Stard10) |
| P54227 | 16765 | Mus musculus | stathmin 1(Stmn1) |
| P32020 | 114745 | Mus musculus | stem cell proliferation 2(Scpro2) |
| P32020 | 20280 | Mus musculus | sterol carrier protein 2, liver(Scp2) |
| Q99JB2 | 66592 | Mus musculus | stomatin (Epb7.2)-like 2(Stoml2) |
| Q60864 | 20867 | Mus musculus | stress-induced phosphoprotein 1(Stip1) |
| Q8K2B3 | 66945 | Mus musculus | succinate dehydrogenase complex, subunit A, flavoprotein (Fp)(Sdha) |
| Q9CQA3 | 67680 | Mus musculus | succinate dehydrogenase complex, subunit B, iron sulfur (Ip)(Sdhb) |
| Q9WUM5 | 56451 | Mus musculus | succinate-CoA ligase, GDP-forming, alpha subunit(Suclg1) |
| Q9Z2I9 | 20916 | Mus musculus | succinate-Coenzyme A ligase, ADP-forming, beta subunit(Sucla2) |
| Q9Z2I8 | 20917 | Mus musculus | succinate-Coenzyme A ligase, GDP-forming, beta subunit(Suclg2) |

| | | | |
|---|---|---|---|
| Q9R1T2 | 56459 | Mus musculus | SUMO1 activating enzyme subunit 1(Sae1) |
| P08228 | 20655 | Mus musculus | superoxide dismutase 1, soluble(Sod1) |
| P09671 | 20656 | Mus musculus | superoxide dismutase 2, mitochondrial(Sod2) |
| Q99L47 | 70356 | Mus musculus | suppression of tumorigenicity 13(St13) |
| P97496 | 20588 | Mus musculus | SWI/SNF related BAF chromatin remodeling complex subunit C1(Smarcc1) |
| O88935 | 20964 | Mus musculus | synapsin I(Syn1) |
| Q64332 | 20965 | Mus musculus | synapsin II(Syn2) |
| Q9JIS5 | 64051 | Mus musculus | synaptic vesicle glycoprotein 2a(Sv2a) |
| Q8BG39 | 64176 | Mus musculus | synaptic vesicle glycoprotein 2b(Sv2b) |
| O55100 | 20972 | Mus musculus | synaptogyrin 1(Syngr1) |
| Q8CHC4 | 104015 | Mus musculus | synaptojanin 1(Synj1) |
| Q9D6K5 | 24071 | Mus musculus | synaptojanin 2 binding protein(Synj2bp) |
| Q62277 | 20977 | Mus musculus | synaptophysin(Syp) |
| Q8CC35 | 104027 | Mus musculus | synaptopodin(Synpo) |
| Q8BGN8 | 72003 | Mus musculus | synaptoporin(Synpr) |
| P60879 | 20614 | Mus musculus | synaptosomal-associated protein 25(Snap25) |
| Q61548 | 20616 | Mus musculus | synaptosomal-associated protein 91(Snap91) |
| Q8R570 | 67826 | Mus musculus | synaptosomal-associated protein, 47(Snap47) |
| P46096 | 20979 | Mus musculus | synaptotagmin I(Syt1) |
| P46097 | 20980 | Mus musculus | synaptotagmin II(Syt2) |
| O35526 | 20907 | Mus musculus | syntaxin 1A (brain)(Stx1a) |
| P61264 | 56216 | Mus musculus | syntaxin 1B(Stx1b) |
| O70439 | 53331 | Mus musculus | syntaxin 7(Stx7) |
| O08599 | 20910 | Mus musculus | syntaxin binding protein 1(Stxbp1) |
| Q64324 | 20911 | Mus musculus | syntaxin binding protein 2(Stxbp2) |
| O55042 | 20617 | Mus musculus | synuclein, alpha(Snca) |
| Q91ZZ3 | 104069 | Mus musculus | synuclein, beta(Sncb) |
| Q9Z0F7 | 20618 | Mus musculus | synuclein, gamma(Sncg) |
| P58771 | 110086 | Mus musculus | T cell phenotype modifier 1(Tcpm1) |
| Q3UES3 | 74493 | Mus musculus | tankyrase, TRF1-interacting ankyrin-related ADP-ribose polymerase 2(Tnks2) |
| Q5SRX1 | 216810 | Mus musculus | target of myb1-like 2 (chicken)(Tom1l2) |
| P11983 | 21454 | Mus musculus | t-complex protein 1(Tcp1) |
| Q8BYI9 | 21960 | Mus musculus | tenascin R(Tnr) |
| Q9JKL5 | 57816 | Mus musculus | tescalcin(Tesc) |
| Q3URK3 | 52463 | Mus musculus | tet methylcytosine dioxygenase 1(Tet1) |

| | | | |
|---|---|---|---|
| Q922J6 | 70747 | Mus musculus | tetraspanin 2(Tspan2) |
| Q62283 | 21912 | Mus musculus | tetraspanin 7(Tspan7) |
| P10639 | 22166 | Mus musculus | thioredoxin 1(Txn1) |
| P97493 | 56551 | Mus musculus | thioredoxin 2(Txn2) |
| Q9CQM5 | 52700 | Mus musculus | thioredoxin domain containing 17(Txndc17) |
| Q8CDN6 | 53382 | Mus musculus | thioredoxin-like 1(Txnl1) |
| P52196 | 22117 | Mus musculus | thiosulfate sulfurtransferase, mitochondrial(Tst) |
| P20065 | 19241 | Mus musculus | thymosin, beta 4, X chromosome(Tmsb4x) |
| P01831 | 21838 | Mus musculus | thymus cell antigen 1, theta(Thy1) |
| A2ASS6 | 22138 | Mus musculus | titin(Ttn) |
| Q8VC26 | 380712 | Mus musculus | TLC domain containing 2(Tlcd2) |
| Q9QUK6 | 21898 | Mus musculus | toll-like receptor 4(Tlr4) |
| O55013 | 27096 | Mus musculus | trafficking protein particle complex 3(Trappc3) |
| Q93092 | 21351 | Mus musculus | transaldolase 1(Taldo1) |
| Q8R0A5 | 594844 | Mus musculus | transcription elongation factor A (SII)-like 3(Tceal3) |
| Q8CCT4 | 331532 | Mus musculus | transcription elongation factor A (SII)-like 5(Tceal5) |
| Q921I1 | 22041 | Mus musculus | transferrin(Trf) |
| Q9R1Q8 | 56370 | Mus musculus | transgelin 3(Tagln3) |
| P40142 | 21881 | Mus musculus | transketolase(Tkt) |
| Q9D4D4 | 74419 | Mus musculus | transketolase-like 2(Tktl2) |
| P62073 | 30059 | Mus musculus | translocase of inner mitochondrial membrane 10(Timm10) |
| Q99KF1 | 67511 | Mus musculus | transmembrane p24 trafficking protein 9(Tmed9) |
| Q3UBX0 | 68539 | Mus musculus | transmembrane protein 109(Tmem109) |
| Q8VEK0 | 69981 | Mus musculus | transmembrane protein 30A(Tmem30a) |
| P07309 | 22139 | Mus musculus | transthyretin(Ttr) |
| Q8VC30 | 225913 | Mus musculus | triokinase, FMN cyclase(Tkfc) |
| P17751 | 21991 | Mus musculus | triosephosphate isomerase 1(Tpi1) |
| Q8BVP1 | 244448 | Mus musculus | tripartite motif family-like 1(Triml1) |
| Q62318 | 21849 | Mus musculus | tripartite motif-containing 28(Trim28) |
| Q80X56 | 70928 | Mus musculus | tripartite motif-containing 69(Trim69) |
| O89023 | 12751 | Mus musculus | tripeptidyl peptidase I(Tpp1) |
| Q9JKK7 | 50876 | Mus musculus | tropomodulin 2(Tmod2) |
| P58771 | 22003 | Mus musculus | tropomyosin 1, alpha(Tpm1) |
| P58774 | 22004 | Mus musculus | tropomyosin 2, beta(Tpm2) |
| P21107 | 59069 | Mus musculus | tropomyosin 3, gamma(Tpm3) |

| | | | |
|---|---|---|---|
| Q6IRU2 | 326618 | Mus musculus | tropomyosin 4(Tpm4) |
| P20801 | 21925 | Mus musculus | troponin C2, fast(Tnnc2) |
| Q8C8T8 | 69499 | Mus musculus | TSR2 20S rRNA accumulation(Tsr2) |
| Q8BFR5 | 233870 | Mus musculus | Tu translation elongation factor, mitochondrial(Tufm) |
| Q9D1E6 | 66411 | Mus musculus | tubulin folding cofactor B(Tbcb) |
| Q7TQD2 | 72948 | Mus musculus | tubulin polymerization promoting protein(Tppp) |
| Q9CRB6 | 67971 | Mus musculus | tubulin polymerization-promoting protein family member 3(Tppp3) |
| P68369 | 22142 | Mus musculus | tubulin, alpha 1A(Tuba1a) |
| P05214 | 22144 | Mus musculus | tubulin, alpha 3A(Tuba3a) |
| P05214 | 22147 | Mus musculus | tubulin, alpha 3B(Tuba3b) |
| P68368 | 22145 | Mus musculus | tubulin, alpha 4A(Tuba4a) |
| P68368 | 53857 | Mus musculus | tubulin, alpha 8(Tuba8) |
| Q7TMM9 | 22151 | Mus musculus | tubulin, beta 2A class IIA(Tubb2a) |
| Q9CWF2 | 73710 | Mus musculus | tubulin, beta 2B class IIB(Tubb2b) |
| Q9ERD7 | 22152 | Mus musculus | tubulin, beta 3 class III(Tubb3) |
| Q9D6F9 | 22153 | Mus musculus | tubulin, beta 4A class IVA(Tubb4a) |
| P68372 | 227613 | Mus musculus | tubulin, beta 4B class IVB(Tubb4b) |
| P99024 | 22154 | Mus musculus | tubulin, beta 5 class I(Tubb5) |
| Q922F4 | 67951 | Mus musculus | tubulin, beta 6 class V(Tubb6) |
| Q9CYZ2 | 66314 | Mus musculus | tumor protein D52-like 2(Tpd52l2) |
| P63028 | 22070 | Mus musculus | tumor protein, translationally-controlled 1(Tpt1) |
| P68254 | 22630 | Mus musculus | tyrosine 3-monooxygenase/tryptophan 5-monooxygenase activation protein theta(Ywhaq) |
| Q9CQV8 | 54401 | Mus musculus | tyrosine 3-monooxygenase/tryptophan 5-monooxygenase activation protein, beta polypeptide(Ywhab) |
| P62259 | 22627 | Mus musculus | tyrosine 3-monooxygenase/tryptophan 5-monooxygenase activation protein, epsilon polypeptide(Ywhae) |
| P68510 | 22629 | Mus musculus | tyrosine 3-monooxygenase/tryptophan 5-monooxygenase activation protein, eta polypeptide(Ywhah) |
| P61982 | 22628 | Mus musculus | tyrosine 3-monooxygenase/tryptophan 5-monooxygenase activation protein, gamma polypeptide(Ywhag) |
| P63101 | 22631 | Mus musculus | tyrosine 3-monooxygenase/tryptophan 5-monooxygenase activation protein, zeta polypeptide(Ywhaz) |
| P97343 | 16589 | Mus musculus | U2AF homology motif (UHM) kinase 1(Uhmk1) |
| Q8R317 | 56085 | Mus musculus | ubiquilin 1(Ubqln1) |
| Q9QZM0 | 54609 | Mus musculus | ubiquilin 2(Ubqln2) |
| Q9DB77 | 67003 | Mus musculus | ubiquinol cytochrome c reductase core protein 2(Uqcrc2) |
| Q9D855 | 67530 | Mus musculus | ubiquinol-cytochrome c reductase binding protein(Uqcrb) |

| | | | |
|---|---|---|---|
| Q9CZ13 | 22273 | Mus musculus | ubiquinol-cytochrome c reductase core protein 1(Uqcrc1) |
| P99028 | 66576 | Mus musculus | ubiquinol-cytochrome c reductase hinge protein(Uqcrh) |
| Q9CQ69 | 22272 | Mus musculus | ubiquinol-cytochrome c reductase, complex III subunit VII(Uqcrq) |
| Q8R1I1 | 66152 | Mus musculus | ubiquinol-cytochrome c reductase, complex III subunit X(Uqcr10) |
| Q9CR68 | 66694 | Mus musculus | ubiquinol-cytochrome c reductase, Rieske iron-sulfur polypeptide 1(Uqcrfs1) |
| Q9JKB1 | 50933 | Mus musculus | ubiquitin carboxyl-terminal esterase L3 (ubiquitin thiolesterase)(Uchl3) |
| Q9R0P9 | 22223 | Mus musculus | ubiquitin carboxy-terminal hydrolase L1(Uchl1) |
| Q9JMA1 | 59025 | Mus musculus | ubiquitin specific peptidase 14(Usp14) |
| P56399 | 22225 | Mus musculus | ubiquitin specific peptidase 5 (isopeptidase T)(Usp5) |
| Q9CZY3 | 66589 | Mus musculus | ubiquitin-conjugating enzyme E2 variant 1(Ube2v1) |
| Q9D2M8 | 70620 | Mus musculus | ubiquitin-conjugating enzyme E2 variant 2(Ube2v2) |
| P61087 | 53323 | Mus musculus | ubiquitin-conjugating enzyme E2K(Ube2k) |
| P68037 | 22195 | Mus musculus | ubiquitin-conjugating enzyme E2L 3(Ube2l3) |
| P61082 | 22192 | Mus musculus | ubiquitin-conjugating enzyme E2M(Ube2m) |
| P61089 | 93765 | Mus musculus | ubiquitin-conjugating enzyme E2N(Ube2n) |
| Q8BGR9 | 79560 | Mus musculus | ubiquitin-like domain containing CTD phosphatase 1(Ublcp1) |
| Q02053 | 22201 | Mus musculus | ubiquitin-like modifier activating enzyme 1(Uba1) |
| Q9Z1F9 | 50995 | Mus musculus | ubiquitin-like modifier activating enzyme 2(Uba2) |
| P70691 | 22236 | Mus musculus | UDP glucuronosyltransferase 1 family, polypeptide A2(Ugt1a2) |
| P17717 | 22238 | Mus musculus | UDP glucuronosyltransferase 2 family, polypeptide B5(Ugt2b5) |
| P25688 | 22262 | Mus musculus | urate oxidase(Uox) |
| Q9Z1Z0 | 56041 | Mus musculus | USO1 vesicle docking factor(Uso1) |
| Q9D1C8 | 66914 | Mus musculus | vacuolar protein sorting 28(Vps28) |
| Q01853 | 269523 | Mus musculus | valosin containing protein(Vcp) |
| Q62059 | 13003 | Mus musculus | versican(Vcan) |
| Q62465 | 26949 | Mus musculus | vesicle amine transport 1(Vat1) |
| Q80TB8 | 270097 | Mus musculus | vesicle amine transport protein 1 like(Vat1l) |
| Q62442 | 22317 | Mus musculus | vesicle-associated membrane protein 1(Vamp1) |
| P63024 | 22319 | Mus musculus | vesicle-associated membrane protein 3(Vamp3) |
| Q9WV55 | 30960 | Mus musculus | vesicle-associated membrane protein, associated protein A(Vapa) |
| Q9QY76 | 56491 | Mus musculus | vesicle-associated membrane protein, associated protein B and C(Vapb) |
| P20152 | 22352 | Mus musculus | vimentin(Vim) |
| P62761 | 26950 | Mus musculus | visinin-like 1(Vsnl1) |
| P21614 | 14473 | Mus musculus | vitamin D binding protein(Gc) |
| Q60932 | 22333 | Mus musculus | voltage-dependent anion channel 1(Vdac1) |

| | | | |
|---|---|---|---|
| Q60930 | 22334 | Mus musculus | voltage-dependent anion channel 2(Vdac2) |
| Q60931 | 22335 | Mus musculus | voltage-dependent anion channel 3(Vdac3) |
| Q8R2E6 | 22297 | Mus musculus | vomeronasal 1 receptor 45(Vmn1r45) |
| P40336 | 30930 | Mus musculus | VPS26 retromer complex component A(Vps26a) |
| Q8C0E2 | 69091 | Mus musculus | VPS26 retromer complex component B(Vps26b) |
| Q9QZ88 | 56433 | Mus musculus | VPS29 retromer complex component(Vps29) |
| P63321 | 56044 | Mus musculus | v-ral simian leukemia viral oncogene A (ras related)(Rala) |
| Q9JIW9 | 64143 | Mus musculus | v-ral simian leukemia viral oncogene B(Ralb) |
| Q8R5H6 | 83767 | Mus musculus | WASP family, member 1(Wasf1) |
| O88342 | 22388 | Mus musculus | WD repeat domain 1(Wdr1) |
| P97765 | 22378 | Mus musculus | WW domain binding protein 2(Wbp2) |
| Q5GH66 | 319581 | Mus musculus | X-linked Kx blood group related 5(Xkr5) |
| P62960 | 22608 | Mus musculus | Y box protein 1(Ybx1) |
| Q9JKB3 | 56449 | Mus musculus | Y box protein 3(Ybx3) |
| Q04736 | 22612 | Mus musculus | YES proto-oncogene 1, Src family tyrosine kinase(Yes1) |
| Q9CQW1 | 56418 | Mus musculus | YKT6 v-SNARE homolog (S. cerevisiae)(Ykt6) |
| Q811F1 | 226470 | Mus musculus | zinc finger and BTB domain containing 41(Zbtb41) |
| Q9CWH1 | 73680 | Mus musculus | zinc finger and BTB domain containing 8a(Zbtb8a) |
| Q8C8V1 | 80292 | Mus musculus | ZXD family zinc finger C(Zxdc) |

Functional Annotation Chart
Help and Manual
Current Gene List: N-Hcy-proteins in vitro
Current Background: Mus musculus
1232 DAVID IDs

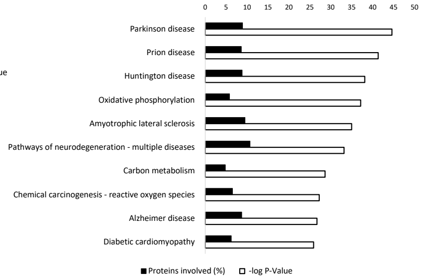

Options

130 chart r Right-click and select 'Save Target As' to download results Download File

| Sublist | Category | Term | RT | Genes | Count | % | P-Value | Fold Enrich | Bonferroni | Benjamini | FDR | Fisher Exact | -log P-Value | Term | Proteins in | -log P-Value |
|---|---|---|---|---|---|---|---|---|---|---|---|---|---|---|---|---|
| | KEGG_PAT | Parkinson | RT | | 110 | 8,9 | 2,60E-45 | 4,5 | 8,30E-43 | 8,30E-43 | 5,80E-43 | 3,80E-46 | 44,58503 | Parkinson | 8,9 | 44,58503 |
| | KEGG_PAT | Prion disea | RT | | 106 | 8,6 | 5,00E-42 | 4,4 | 1,60E-39 | 8,10E-40 | 5,70E-40 | 7,80E-43 | 41,30103 | Prion disea | 8,6 | 41,30103 |
| | KEGG_PAT | Huntingto | RT | | 109 | 8,8 | 7,90E-39 | 4 | 2,60E-36 | 8,50E-37 | 6,00E-37 | 1,40E-39 | 38,10237 | Huntingto | 8,8 | 38,10237 |
| | KEGG_PAT | Oxidative | RT | | 72 | 5,8 | 7,80E-38 | 5,7 | 2,50E-35 | 6,30E-36 | 4,40E-36 | 7,40E-39 | 37,10791 | Oxidative | 5,8 | 37,10791 |
| | KEGG_PAT | Amyotropl | RT | | 117 | 9,5 | 1,10E-35 | 3,5 | 3,60E-33 | 7,20E-34 | 5,00E-34 | 2,40E-36 | 34,95861 | Amyotropl | 9,5 | 34,95861 |
| | KEGG_PAT | Pathways | RT | | 132 | 10,7 | 6,90E-34 | 3,1 | 2,20E-31 | 3,70E-32 | 2,60E-32 | 1,80E-34 | 33,16115 | Pathways | 10,7 | 33,16115 |
| | KEGG_PAT | Carbon me | RT | | 59 | 4,8 | 2,40E-29 | 5,4 | 7,90E-27 | 1,10E-27 | 7,90E-28 | 2,60E-30 | 28,61979 | Carbon me | 4,8 | 28,61979 |
| | KEGG_PAT | Chemical c | RT | | 80 | 6,5 | 6,10E-28 | 3,9 | 2,00E-25 | 2,50E-26 | 1,70E-26 | 1,10E-28 | 27,21467 | Chemical c | 6,5 | 27,21467 |
| | KEGG_PAT | Alzheimer | RT | | 107 | 8,7 | 2,10E-27 | 3,1 | 6,70E-25 | 7,50E-26 | 5,30E-26 | 5,40E-28 | 26,67778 | Alzheimer | 8,7 | 26,67778 |
| | KEGG_PAT | Diabetic ca | RT | | 76 | 6,2 | 1,30E-26 | 3,9 | 4,10E-24 | 4,10E-25 | 2,90E-25 | 2,30E-27 | 25,88606 | Diabetic ca | 6,2 | 25,88606 |
| | KEGG_PAT | Metabolic | RT | | 261 | 21,2 | 5,50E-24 | 1,8 | 1,80E-21 | 1,60E-22 | 1,10E-22 | 2,90E-24 | 23,25964 | Metabolic | 21,2 | 23,25964 |
| | KEGG_PAT | Ribosome | RT | | 76 | 6,2 | 1,20E-20 | 3,2 | 3,90E-18 | 3,20E-19 | 2,30E-19 | 2,90E-21 | 19,92082 | Ribosome | 6,2 | 19,92082 |
| | KEGG_PAT | Biosynthes | RT | | 37 | 3 | 6,60E-18 | 5,3 | 2,10E-15 | 1,60E-16 | 1,10E-16 | 7,50E-19 | 17,18046 | Biosynthes | 3 | 17,18046 |
| | KEGG_PAT | Coronaviru | RT | | 82 | 6,7 | 9,60E-18 | 2,8 | 3,10E-15 | 2,20E-16 | 1,60E-16 | 2,90E-18 | 17,01773 | Coronaviru | 6,7 | 17,01773 |
| | KEGG_PAT | Thermoge | RT | | 66 | 5,4 | 2,80E-17 | 3,1 | 9,10E-15 | 6,00E-16 | 4,20E-16 | 7,20E-18 | 16,55284 | Thermoge | 5,4 | 16,55284 |
| | KEGG_PAT | Synaptic v | RT | | 35 | 2,8 | 1,80E-16 | 5,1 | 7,20E-14 | 3,70E-15 | 2,60E-15 | 2,20E-17 | 15,74473 | Synaptic v | 2,8 | 15,74473 |
| | KEGG_PAT | Salmonella | RT | | 65 | 5,3 | 3,90E-15 | 2,9 | 1,30E-12 | 7,30E-14 | 5,10E-14 | 1,10E-15 | 14,40894 | Salmonella | 5,3 | 14,40894 |
| | KEGG_PAT | Non-alcoh | RT | | 49 | 4 | 2,30E-14 | 3,4 | 7,30E-12 | 4,00E-13 | 2,80E-13 | 5,10E-15 | 13,63827 | Non-alcoh | 4 | 13,63827 |
| | KEGG_PAT | Cardiac mu | RT | | 35 | 2,8 | 3,90E-14 | 4,4 | 1,30E-11 | 6,60E-13 | 4,70E-13 | 6,00E-15 | 13,40894 | Cardiac mu | 2,8 | 13,40894 |
| | KEGG_PAT | Citrate cyc | RT | | 21 | 1,7 | 6,40E-14 | 7,4 | 2,10E-11 | 1,00E-12 | 7,30E-13 | 3,60E-15 | 13,19382 | Citrate cyc | 1,7 | 13,19382 |
| | KEGG_PAT | Retrograde | RT | | 46 | 3,7 | 2,00E-13 | 3,4 | 6,50E-11 | 3,10E-12 | 2,20E-12 | 4,60E-14 | 12,69897 | Retrograde | 3,7 | 12,69897 |
| | KEGG_PAT | Glycolysis , | RT | | 28 | 2,3 | 3,80E-12 | 4,7 | 1,20E-09 | 5,60E-11 | 3,90E-11 | 5,30E-13 | 11,42022 | Glycolysis , | 2,3 | 11,42022 |
| | KEGG_PAT | Endocrine | RT | | 26 | 2,1 | 1,50E-11 | 4,8 | 5,00E-09 | 2,20E-10 | 1,50E-10 | 2,10E-12 | 10,82391 | Endocrine | 2,1 | 10,82391 |
| | KEGG_PAT | Valine, leu | RT | | 24 | 1,9 | 1,50E-10 | 4,7 | 4,80E-08 | 2,00E-09 | 1,40E-09 | 2,00E-11 | 9,823909 | Valine, leu | 1,9 | 9,823909 |
| | KEGG_PAT | Pyruvate n | RT | | 21 | 1,7 | 1,80E-10 | 5,4 | 5,70E-08 | 2,30E-09 | 1,60E-09 | 2,00E-11 | 9,744727 | Pyruvate n | 1,7 | 9,744727 |
| | KEGG_PAT | Phagosom | RT | | 44 | 3,6 | 2,00E-09 | 2,7 | 6,40E-07 | 2,50E-08 | 1,70E-08 | 6,10E-10 | 8,69897 | Phagosom | 3,6 | 8,69897 |
| | KEGG_PAT | Motor pro | RT | | 46 | 3,7 | 2,60E-09 | 2,6 | 8,40E-07 | 3,10E-08 | 2,20E-08 | 8,30E-10 | 8,585027 | Motor pro | 3,7 | 8,585027 |
| | KEGG_PAT | Endocytosi | RT | | 56 | 4,5 | 3,20E-09 | 2,3 | 1,00E-06 | 3,70E-08 | 2,60E-08 | 1,20E-09 | 8,49485 | Endocytosi | 4,5 | 8,49485 |
| | KEGG_PAT | 2-Oxocarb | RT | | 16 | 1,3 | 5,80E-08 | 5,3 | 1,90E-05 | 6,40E-07 | 4,50E-07 | 6,60E-09 | 7,236572 | 2-Oxocarb | 1,3 | 7,236572 |
| | KEGG_PAT | Gap juncti | RT | | 26 | 2,1 | 6,20E-08 | 3,4 | 2,00E-05 | 6,60E-07 | 4,70E-07 | 1,40E-08 | 7,207608 | Gap juncti | 2,1 | 7,207608 |
| | KEGG_PAT | Bacterial i | RT | | 24 | 1,9 | 1,20E-07 | 3,5 | 3,90E-05 | 1,20E-06 | 8,80E-07 | 2,60E-08 | 6,920819 | Bacterial i | 1,9 | 6,920819 |
| | KEGG_PAT | Glutamate | RT | | 30 | 2,4 | 2,10E-07 | 2,9 | 6,80E-05 | 2,10E-06 | 1,50E-06 | 5,90E-08 | 6,677781 | Glutamate | 2,4 | 6,677781 |
| | KEGG_PAT | Adrenergic | RT | | 36 | 2,9 | 2,20E-07 | 2,6 | 7,00E-05 | 2,10E-06 | 1,50E-06 | 7,00E-08 | 6,657577 | Adrenergic | 2,9 | 6,657577 |
| | KEGG_PAT | Protein pr | RT | | 38 | 3,1 | 4,80E-07 | 2,4 | 1,50E-04 | 4,50E-06 | 3,20E-06 | 1,70E-07 | 6,318759 | Protein pr | 3,1 | 6,318759 |
| | KEGG_PAT | Tight junct | RT | | 37 | 3 | 6,50E-07 | 2,4 | 2,10E-04 | 5,90E-06 | 4,20E-06 | 2,30E-07 | 6,187087 | Tight junct | 3 | 6,187087 |
| | KEGG_PAT | GABAergic | RT | | 25 | 2 | 6,60E-07 | 3,1 | 2,10E-04 | 5,90E-06 | 4,20E-06 | 1,70E-07 | 6,180456 | GABAergic | 2 | 6,180456 |
| | KEGG_PAT | Alcoholism | RT | | 42 | 3,4 | 7,30E-07 | 2,3 | 2,40E-04 | 6,40E-06 | 4,50E-06 | 2,80E-07 | 6,136677 | Alcoholism | 3,4 | 6,136677 |
| | KEGG_PAT | HIF-1 signa | RT | | 29 | 2,4 | 8,90E-07 | 2,8 | 2,90E-04 | 7,60E-06 | 5,30E-06 | 2,60E-07 | 6,05061 | HIF-1 signa | 2,4 | 6,05061 |
| | KEGG_PAT | Fc gamma | RT | | 24 | 1,9 | 5,60E-06 | 2,9 | 1,80E-03 | 4,50E-05 | 3,20E-05 | 1,60E-06 | 5,251812 | Fc gamma | 1,9 | 5,251812 |
| | KEGG_PAT | Gastric aci | RT | | 21 | 1,7 | 5,60E-06 | 3,1 | 1,80E-03 | 4,50E-05 | 3,20E-05 | 1,40E-06 | 5,251812 | Gastric aci | 1,7 | 5,251812 |
| | KEGG_PAT | Propanoat | RT | | 13 | 1,1 | 6,60E-06 | 4,7 | 2,10E-03 | 5,10E-05 | 3,60E-05 | 9,30E-07 | 5,180456 | Propanoat | 1,1 | 5,180456 |
| | KEGG_PAT | cGMP-PKG | RT | | 35 | 2,8 | 6,70E-06 | 2,3 | 2,20E-03 | 5,10E-05 | 3,60E-05 | 2,50E-06 | 5,173925 | cGMP-PKG | 2,8 | 5,173925 |
| | KEGG_PAT | Drug meta | RT | | 24 | 1,9 | 6,80E-06 | 2,8 | 2,20E-03 | 5,10E-05 | 3,60E-05 | 2,00E-06 | 5,167491 | Drug meta | 1,9 | 5,167491 |
| | KEGG_PAT | Dopamine | RT | | 30 | 2,4 | 6,90E-06 | 2,5 | 2,20E-03 | 5,10E-05 | 3,60E-05 | 2,40E-06 | 5,161151 | Dopamine | 2,4 | 5,161151 |
| | KEGG_PAT | Proximal t | RT | | 11 | 0,9 | 7,10E-06 | 5,6 | 2,30E-03 | 5,10E-05 | 3,60E-05 | 7,50E-07 | 5,148742 | Proximal t | 0,9 | 5,148742 |
| | KEGG_PAT | Butanoate | RT | | 12 | 1 | 9,00E-06 | 5 | 2,90E-03 | 6,30E-05 | 4,40E-05 | 1,20E-06 | 5,045757 | Butanoate | 1 | 5,045757 |
| | KEGG_PAT | Glyoxylate | RT | | 13 | 1,1 | 9,70E-06 | 4,6 | 3,10E-03 | 6,70E-05 | 4,70E-05 | 1,40E-06 | 5,013228 | Glyoxylate | 1,1 | 5,013228 |
| | KEGG_PAT | Cytoskelet | RT | | 42 | 3,4 | 1,40E-05 | 2 | 4,60E-03 | 9,70E-05 | 6,80E-05 | 6,30E-06 | 4,853872 | Cytoskelet | 3,4 | 4,853872 |
| | KEGG_PAT | Neutrophil | RT | | 39 | 3,2 | 1,50E-05 | 2,1 | 4,90E-03 | 1,00E-04 | 7,00E-05 | 6,40E-06 | 4,823909 | Neutrophi | 3,2 | 4,823909 |
| | KEGG_PAT | Vasopressi | RT | | 15 | 1,2 | 1,60E-05 | 3,8 | 5,20E-03 | 1,00E-04 | 7,30E-05 | 3,10E-06 | 4,79588 | Vasopressi | 1,2 | 4,79588 |
| | KEGG_PAT | Arginine b | RT | | 10 | 0,8 | 3,70E-05 | 5,3 | 1,20E-02 | 2,40E-04 | 1,70E-04 | 4,30E-06 | 4,431798 | Arginine b | 0,8 | 4,431798 |
| | KEGG_PAT | Fructose a | RT | | 13 | 1,1 | 3,90E-05 | 4 | 1,20E-02 | 2,40E-04 | 1,70E-04 | 6,80E-06 | 4,408935 | Fructose a | 1,1 | 4,408935 |
| | KEGG_PAT | Long-term | RT | | 17 | 1,4 | 4,80E-05 | 3,2 | 1,50E-02 | 2,90E-04 | 2,10E-04 | 1,20E-05 | 4,318759 | Long-term | 1,4 | 4,318759 |
| | KEGG_PAT | Glutathion | RT | | 19 | 1,5 | 5,20E-05 | 2,9 | 1,70E-02 | 3,10E-04 | 2,20E-04 | 1,40E-05 | 4,283997 | Glutathion | 1,5 | 4,283997 |
| | KEGG_PAT | Thyroid ho | RT | | 19 | 1,5 | 6,30E-05 | 2,9 | 2,00E-02 | 3,70E-04 | 2,60E-04 | 1,80E-05 | 4,200659 | Thyroid ho | 1,5 | 4,200659 |
| | KEGG_PAT | Oxytocin s | RT | | 30 | 2,4 | 8,00E-05 | 2,2 | 2,50E-02 | 4,60E-04 | 3,20E-04 | 3,20E-05 | 4,09691 | Oxytocin s | 2,4 | 4,09691 |
| | KEGG_PAT | Regulation | RT | | 40 | 3,2 | 8,90E-05 | 1,9 | 2,80E-02 | 5,00E-04 | 3,50E-04 | 4,20E-05 | 4,05061 | Regulation | 3,2 | 4,05061 |
| | KEGG_PAT | Alanine, as | RT | | 13 | 1,1 | 9,40E-05 | 3,7 | 3,00E-02 | 5,20E-04 | 3,70E-04 | 1,90E-05 | 4,026872 | Alanine, as | 1,1 | 4,026872 |
| | KEGG_PAT | Sulfur met | RT | | 7 | 0,6 | 1,50E-04 | 7,1 | 4,80E-02 | 8,40E-04 | 5,90E-04 | 1,00E-05 | 3,823909 | Sulfur met | 0,6 | 3,823909 |
| | KEGG_PAT | Long-term | RT | | 17 | 1,4 | 2,00E-04 | 2,8 | 6,20E-02 | 1,10E-03 | 7,50E-04 | 5,70E-05 | 3,69897 | Long-term | 1,4 | 3,69897 |
| | KEGG_PAT | Cysteine a | RT | | 15 | 1,2 | 2,40E-04 | 3,1 | 7,50E-02 | 1,30E-03 | 9,00E-04 | 6,30E-05 | 3,619789 | Cysteine a | 1,2 | 3,619789 |
| | KEGG_PAT | Collecting | RT | | 10 | 0,8 | 3,60E-04 | 4,2 | 1,10E-01 | 1,90E-03 | 1,30E-03 | 6,20E-05 | 3,443697 | Collecting | 0,8 | 3,443697 |
| | KEGG_PAT | Circadian e | RT | | 21 | 1,7 | 3,80E-04 | 2,4 | 1,20E-01 | 2,00E-03 | 1,40E-03 | 1,40E-04 | 3,420216 | Circadian e | 1,7 | 3,420216 |
| | KEGG_PAT | Viral carcir | RT | | 37 | 3 | 4,80E-04 | 1,8 | 1,40E-01 | 2,40E-03 | 1,70E-03 | 2,40E-04 | 3,318759 | Viral carcir | 3 | 3,318759 |
| | KEGG_PAT | Fatty acid | RT | | 14 | 1,1 | 4,80E-04 | 3 | 1,40E-01 | 2,40E-03 | 1,70E-03 | 1,30E-04 | 3,318759 | Fatty acid | 1,1 | 3,318759 |
| | KEGG_PAT | Pentose pl | RT | | 11 | 0,9 | 5,40E-04 | 3,6 | 1,60E-01 | 2,60E-03 | 1,80E-03 | 1,10E-04 | 3,267606 | Pentose pl | 0,9 | 3,267606 |
| | KEGG_PAT | Tyrosine m | RT | | 12 | 1 | 5,40E-04 | 3,4 | 1,60E-01 | 2,60E-03 | 1,80E-03 | 1,20E-04 | 3,267606 | Tyrosine m | 1 | 3,267606 |
| | KEGG_PAT | Proteasom | RT | | 13 | 1,1 | 6,40E-04 | 3,1 | 1,90E-01 | 3,10E-03 | 2,20E-03 | 1,60E-04 | 3,19382 | Proteasom | 1,1 | 3,19382 |
| | KEGG_PAT | Mineral ab | RT | | 14 | 1,1 | 8,60E-04 | 2,9 | 2,40E-01 | 4,00E-03 | 2,80E-03 | 2,40E-04 | 3,065502 | Mineral ab | 1,1 | 3,065502 |
| | KEGG_PAT | Metabolisı | RT | | 17 | 1,4 | 9,00E-04 | 2,5 | 2,50E-01 | 4,10E-03 | 2,90E-03 | 3,00E-04 | 3,045757 | Metabolisı | 1,4 | 3,045757 |
| | KEGG_PAT | Spinocereb | RT | | 25 | 2 | 1,40E-03 | 2 | 3,70E-01 | 6,50E-03 | 4,50E-03 | 6,40E-04 | 2,853872 | Spinocereb | 2 | 2,853872 |
| | KEGG_PAT | Drug meta | RT | | 16 | 1,3 | 1,40E-03 | 2,5 | 3,70E-01 | 6,50E-03 | 4,50E-03 | 4,80E-04 | 2,853872 | Drug meta | 1,3 | 2,853872 |
| | KEGG_PAT | Glucagon s | RT | | 20 | 1,6 | 1,90E-03 | 2,2 | 4,60E-01 | 8,30E-03 | 5,90E-03 | 7,60E-04 | 2,721246 | Glucagon s | 1,6 | 2,721246 |
| | KEGG_PAT | Amphetam | RT | | 15 | 1,2 | 2,70E-03 | 2,4 | 5,80E-01 | 1,10E-02 | 8,00E-03 | 9,20E-04 | 2,568636 | Amphetam | 1,2 | 2,568636 |
| | KEGG_PAT | Central ca | RT | | 15 | 1,2 | 2,70E-03 | 2,4 | 5,80E-01 | 1,10E-02 | 8,00E-03 | 9,20E-04 | 2,568636 | Central ca | 1,2 | 2,568636 |
| | KEGG_PAT | Oocyte me | RT | | 22 | 1,8 | 2,70E-03 | 2 | 5,80E-01 | 1,10E-02 | 8,00E-03 | 1,20E-03 | 2,568636 | Oocyte me | 1,8 | 2,568636 |
| | KEGG_PAT | Fluid shea | RT | | 25 | 2 | 2,80E-03 | 1,9 | 6,00E-01 | 1,20E-02 | 8,30E-03 | 1,30E-03 | 2,552842 | Fluid shea | 2 | 2,552842 |
| | KEGG_PAT | Insulin sec | RT | | 17 | 1,4 | 3,50E-03 | 2,2 | 6,70E-01 | 1,40E-02 | 1,00E-02 | 1,30E-03 | 2,455932 | Insulin sec | 1,4 | 2,455932 |
| | KEGG_PAT | Estrogen s | RT | | 23 | 1,9 | 3,50E-03 | 1,9 | 6,80E-01 | 1,40E-02 | 1,00E-02 | 1,60E-03 | 2,455932 | Estrogen s | 1,9 | 2,455932 |
| | KEGG_PAT | Human im | RT | | 35 | 2,8 | 4,70E-03 | 1,6 | 7,80E-01 | 1,90E-02 | 1,30E-02 | 2,60E-03 | 2,327902 | Human im | 2,8 | 2,327902 |
| | KEGG_PAT | Lipoic acid | RT | | 7 | 0,6 | 4,80E-03 | 4,1 | 7,90E-01 | 1,90E-02 | 1,40E-02 | 8,50E-04 | 2,318759 | Lipoic acid | 0,6 | 2,318759 |
| | KEGG_PAT | Cholinergi | RT | | 20 | 1,6 | 4,90E-03 | 2 | 8,00E-01 | 1,90E-02 | 1,40E-02 | 2,20E-03 | 2,309804 | Cholinergi | 1,6 | 2,309804 |
| | KEGG_PAT | Tryptopha | RT | | 12 | 1 | 5,30E-03 | 2,6 | 8,20E-01 | 2,10E-02 | 1,40E-02 | 1,70E-03 | 2,275724 | Tryptopha | 1 | 2,275724 |
| | KEGG_PAT | Legionello | RT | | 13 | 1,1 | 5,90E-03 | 2,4 | 8,50E-01 | 2,30E-02 | 1,60E-02 | 2,00E-03 | 2,229148 | Legionello | 1,1 | 2,229148 |
| | KEGG_PAT | Morphine | RT | | 17 | 1,4 | 6,80E-03 | 2,1 | 8,90E-01 | 2,60E-02 | 1,80E-02 | 2,90E-03 | 2,167491 | Morphine | 1,4 | 2,167491 |
| | KEGG_PAT | Adherens j | RT | | 17 | 1,4 | 6,80E-03 | 2,1 | 8,90E-01 | 2,60E-02 | 1,80E-02 | 2,90E-03 | 2,167491 | Adherens j | 1,4 | 2,167491 |
| | KEGG_PAT | Arginine a | RT | | 12 | 1 | 7,10E-03 | 2,5 | 9,00E-01 | 2,60E-02 | 1,90E-02 | 2,40E-03 | 2,148742 | Arginine a | 1 | 2,148742 |
| | KEGG_PAT | Fatty acid | RT | | 13 | 1,1 | 7,70E-03 | 2,4 | 9,20E-01 | 2,80E-02 | 2,00E-02 | 2,80E-03 | 2,113509 | Fatty acid | 1,1 | 2,113509 |
| | KEGG_PAT | Ferroptosi | RT | | 10 | 0,8 | 8,80E-03 | 2,7 | 9,40E-01 | 3,20E-02 | 2,20E-02 | 2,60E-03 | 2,055517 | Ferroptosi | 0,8 | 2,055517 |
| | KEGG_PAT | Aldosteror | RT | | 18 | 1,5 | 1,00E-02 | 1,9 | 9,60E-01 | 3,60E-02 | 2,50E-02 | 4,50E-03 | 2 | Aldosteror | 1,5 | 2 |
| | KEGG_PAT | Human cyt | RT | | 35 | 2,8 | 1,00E-02 | 1,5 | 9,70E-01 | 3,70E-02 | 2,60E-02 | 6,10E-03 | 2 | Human cyt | 2,8 | 2 |
| | KEGG_PAT | Apelin sigr | RT | | 22 | 1,8 | 1,10E-02 | 1,8 | 9,70E-01 | 3,90E-02 | 2,80E-02 | 5,60E-03 | 1,958607 | Apelin sigr | 1,8 | 1,958607 |
| | KEGG_PAT | One carbo | RT | | 9 | 0,7 | 1,40E-02 | 2,7 | 9,90E-01 | 5,00E-02 | 3,50E-02 | 4,30E-03 | 1,853872 | One carbo | 0,7 | 1,853872 |
| | KEGG_PAT | Aldosteror | RT | | 9 | 0,7 | 1,70E-02 | 2,7 | 1,00E+00 | 5,80E-02 | 4,10E-02 | 5,20E-03 | 1,769551 | Aldosteror | 0,7 | 1,769551 |
| | KEGG_PAT | Serotonerg | RT | | 21 | 1,7 | 1,70E-02 | 1,7 | 1,00E+00 | 5,90E-02 | 4,10E-02 | 8,90E-03 | 1,769551 | Serotonerg | 1,7 | 1,769551 |
| | KEGG_PAT | Leukocyte | RT | | 19 | 1,5 | 1,80E-02 | 1,8 | 1,00E+00 | 6,00E-02 | 4,20E-02 | 8,80E-03 | 1,744727 | Leukocyte | 1,5 | 1,744727 |
| | KEGG_PAT | Dilated ca | RT | | 17 | 1,4 | 1,90E-02 | 1,9 | 1,00E+00 | 6,50E-02 | 4,60E-02 | 9,30E-03 | 1,721246 | Dilated ca | 1,4 | 1,721246 |
| | KEGG_PAT | beta-Alani | RT | | 8 | 0,6 | 2,00E-02 | 2,8 | 1,00E+00 | 6,70E-02 | 4,70E-02 | 5,90E-03 | 1,69897 | beta-Alani | 0,6 | 1,69897 |
| | KEGG_PAT | Antigen pr | RT | | 15 | 1,2 | 2,10E-02 | 1,9 | 1,00E+00 | 6,80E-02 | 4,80E-02 | 9,50E-03 | 1,677781 | Antigen pr | 1,2 | 1,677781 |
| | KEGG_PAT | Salivary se | RT | | 15 | 1,2 | 2,10E-02 | 1,9 | 1,00E+00 | 6,80E-02 | 4,80E-02 | 9,50E-03 | 1,677781 | Salivary se | 1,2 | 1,677781 |
| | KEGG_PAT | Glycine, se | RT | | 9 | 0,7 | 2,30E-02 | 2,5 | 1,00E+00 | 7,30E-02 | 5,10E-02 | 7,50E-03 | 1,638272 | Glycine, se | 0,7 | 1,638272 |
| | KEGG_PAT | Pancreatic | RT | | 18 | 1,5 | 2,60E-02 | 1,8 | 1,00E+00 | 8,00E-02 | 5,70E-02 | 1,30E-02 | 1,585027 | Pancreatic | 1,5 | 1,585027 |
| | KEGG_PAT | cAMP sign | RT | | 30 | 2,4 | 2,60E-02 | 1,5 | 1,00E+00 | 8,00E-02 | 5,70E-02 | 1,60E-02 | 1,585027 | cAMP sign | 2,4 | 1,585027 |
| | KEGG_PAT | Cholestero | RT | | 10 | 0,8 | 3,10E-02 | 2,2 | 1,00E+00 | 9,40E-02 | 6,60E-02 | 1,20E-02 | 1,508638 | Cholestero | 0,8 | 1,508638 |
| | KEGG_PAT | VEGF signa | RT | | 11 | 0,9 | 3,10E-02 | 2,1 | 1,00E+00 | 9,40E-02 | 6,60E-02 | 1,20E-02 | 1,508638 | VEGF signa | 0,9 | 1,508638 |
| | KEGG_PAT | Pentose ar | RT | | 8 | 0,6 | 3,70E-02 | 2,5 | 1,00E+00 | 1,10E-01 | 7,90E-02 | 1,20E-02 | 1,431798 | Pentose ar | 0,6 | 1,431798 |
| | KEGG_PAT | Relaxin sig | RT | | 19 | 1,5 | 3,90E-02 | 1,6 | 1,00E+00 | 1,20E-01 | 8,30E-02 | 2,10E-02 | 1,408935 | Relaxin sig | 1,5 | 1,408935 |
| | KEGG_PAT | Peroxisom | RT | | 14 | 1,1 | 4,40E-02 | 1,8 | 1,00E+00 | 1,30E-01 | 9,10E-02 | 2,10E-02 | 1,356547 | Peroxisom | 1,1 | 1,356547 |
| | KEGG_PAT | Systemic lu | RT | | 21 | 1,7 | 4,40E-02 | 1,6 | 1,00E+00 | 1,30E-01 | 9,10E-02 | 2,50E-02 | 1,356547 | Systemic lu | 1,7 | 1,356547 |
| | KEGG_PAT | Tuberculos | RT | | 24 | 1,9 | 4,90E-02 | 1,5 | 1,00E+00 | 1,40E-01 | 1,00E-01 | 3,00E-02 | 1,309804 | Tuberculos | 1,9 | 1,309804 |
| | KEGG_PAT | PPAR signa | RT | | 14 | 1,1 | 5,10E-02 | 1,8 | 1,00E+00 | 1,50E-01 | 1,00E-01 | 2,50E-02 | 1,29243 | PPAR signa | 1,1 | 1,29243 |
| | KEGG_PAT | Platinum c | RT | | 13 | 1,1 | 5,40E-02 | 1,8 | 1,00E+00 | 1,50E-01 | 1,10E-01 | 2,60E-02 | 1,267606 | Platinum c | 1,1 | 1,267606 |
| | KEGG_PAT | African try | RT | | 8 | 0,6 | 5,40E-02 | 2,3 | 1,00E+00 | 1,50E-01 | 1,10E-01 | 2,00E-02 | 1,267606 | African try | 0,6 | 1,267606 |
| | KEGG_PAT | Sphingolip | RT | | 18 | 1,5 | 5,40E-02 | 1,6 | 1,00E+00 | 1,50E-01 | 1,10E-01 | 3,00E-02 | 1,267606 | Sphingolip | 1,5 | 1,267606 |
| | KEGG_PAT | Chemical c | RT | | 14 | 1,1 | 5,50E-02 | 1,7 | 1,00E+00 | 1,50E-01 | 1,10E-01 | 2,80E-02 | 1,259637 | Chemical c | 1,1 | 1,259637 |
| | KEGG_PAT | Amoebiasi | RT | | 16 | 1,3 | 5,50E-02 | 1,7 | 1,00E+00 | 1,50E-01 | 1,10E-01 | 2,90E-02 | 1,259637 | Amoebiasi | 1,3 | 1,259637 |
| | KEGG_PAT | Yersinia in | RT | | 19 | 1,5 | 5,70E-02 | 1,6 | 1,00E+00 | 1,60E-01 | 1,10E-01 | 3,30E-02 | 1,244125 | Yersinia in | 1,5 | 1,244125 |
| | KEGG_PAT | Nitrogen n | RT | | 5 | 0,4 | 5,80E-02 | 3,3 | 1,00E+00 | 1,60E-01 | 1,10E-01 | 1,40E-02 | 1,236572 | Nitrogen n | 0,4 | 1,236572 |
| | KEGG_PAT | Melanoge | RT | | 15 | 1,2 | 5,90E-02 | 1,7 | 1,00E+00 | 1,60E-01 | 1,10E-01 | 3,10E-02 | 1,229148 | Melanoge | 1,2 | 1,229148 |
| | KEGG_PAT | Hypertrop | RT | | 15 | 1,2 | 5,90E-02 | 1,7 | 1,00E+00 | 1,60E-01 | 1,10E-01 | 3,10E-02 | 1,229148 | Hypertrop | 1,2 | 1,229148 |
| | KEGG_PAT | Carbohydr | RT | | 9 | 0,7 | 6,60E-02 | 2,1 | 1,00E+00 | 1,80E-01 | 1,20E-01 | 2,70E-02 | 1,180456 | Carbohydr | 0,7 | 1,180456 |
| | KEGG_PAT | Thyroid ho | RT | | 17 | 1,4 | 6,60E-02 | 1,6 | 1,00E+00 | 1,80E-01 | 1,20E-01 | 3,70E-02 | 1,180456 | Thyroid ho | 1,4 | 1,180456 |
| | KEGG_PAT | Necroptos | RT | | 23 | 1,9 | 6,80E-02 | 1,5 | 1,00E+00 | 1,80E-01 | 1,30E-01 | 4,20E-02 | 1,167491 | Necroptos | 1,9 | 1,167491 |
| | KEGG_PAT | Lipid and a | RT | | 27 | 2,2 | 7,30E-02 | 1,4 | 1,00E+00 | 1,90E-01 | 1,30E-01 | 4,70E-02 | 1,136677 | Lipid and a | 2,2 | 1,136677 |
| | KEGG_PAT | Folate bios | RT | | 6 | 0,5 | 7,60E-02 | 2,6 | 1,00E+00 | 2,00E-01 | 1,40E-01 | 2,40E-02 | 1,119186 | Folate bios | 0,5 | 1,119186 |
| | KEGG_PAT | Biosynthes | RT | | 20 | 1,6 | 8,60E-02 | 1,5 | 1,00E+00 | 2,20E-01 | 1,60E-01 | 5,30E-02 | 1,065502 | Biosynthes | 1,6 | 1,065502 |
| | KEGG_PAT | Virion - He | RT | | 9 | 0,7 | 8,70E-02 | 1,9 | 1,00E+00 | 2,20E-01 | 1,60E-01 | 3,90E-02 | 1,060481 | Virion - He | 0,7 | 1,060481 |
| | KEGG_PAT | Focal adhe | RT | | 25 | 2 | 8,80E-02 | 1,4 | 1,00E+00 | 2,20E-01 | 1,60E-01 | 5,80E-02 | 1,055517 | Focal adhe | 2 | 1,055517 |
| | KEGG_PAT | Parathyroi | RT | | 16 | 1,3 | 9,10E-02 | 1,5 | 1,00E+00 | 2,30E-01 | 1,60E-01 | 5,20E-02 | 1,040959 | Parathyroi | 1,3 | 1,040959 |
| | KEGG_PAT | Proteoglyc | RT | | 25 | 2 | 9,60E-02 | 1,4 | 1,00E+00 | 2,40E-01 | 1,70E-01 | 6,30E-02 | 1,017729 | Proteoglyc | 2 | 1,017729 |
